# The power to resolve relationships: identifying incongruence and precision of reduced representation and genome-wide data in phylogenomics and population genomics

**DOI:** 10.1101/2025.07.09.663802

**Authors:** Joshua M. Felton, Chloe Jelley, Julianna J. Harden, Justin Scholten, Leland Graber, Michelle Heeney, Yanã C. Rizzieri, Chelsea D. Specht, Jacob B. Landis

## Abstract

Target capture of ultraconserved elements (UCEs) and taxon-specific probes are widely used reduced-representation methods in phylogenomics and, increasingly, in population genomics for their ability to retrieve hundreds to thousands of homologous loci across divergent taxa. Meanwhile, declining costs and improved computational methods have made genome resequencing more accessible for non-model species, enabling the generation of datasets that can address evolutionary and ecological questions from micro- to macroevolutionary scales. Whether target capture approaches to likewise generate datasets that can address questions across broad hierarchical scales remains unclear. Here, we assess the efficacy of data collection (i.e., single nucleotide polymorphism (SNP) retention), predicted genetic variation across samples (i.e., heterozygosity), and phylogenetic congruence between data generated using reduced-representation methods and genome resequencing, leveraging publicly available datasets from plants and animals.

We found that SNP retention varied by locus type, with genome-wide datasets retaining the highest proportion of SNPs and UCEs the lowest proportion. Heterozygosity also differed, with Benchmarking Universal Single-Copy Orthologs (BUSCOs) producing the lowest estimates, followed by UCEs; the inclusion of supercontig flanking regions raised heterozygosity values moderately.

Across all phylogenetic trees, UCE datasets had the lowest bootstrap support, followed by BUSCOs and single copy orthologous genes. Population structure analyses frequently underestimated the number of ancestral populations in reduced-representation datasets, often identifying fewer populations than genome-wide datasets and assigning samples to different clusters. These discrepancies underscore the challenges of relying solely on reduced-representation methods for robust inferences of genetic diversity, phylogenetic relationships, and population structure.

## Introduction

Advances in high throughput sequencing have made it feasible to collect thousands of markers, yet the optimal choice of which loci to sequence and how many are required for phylogenomic and population genomic analyses is an ongoing debate in evolutionary biology, largely due to the best set of markers being lineage and/or context dependent (Hughes et al., 2006; Ai & Kang, 2015; Borowiec et al., 2024). Slower-evolving markers are better suited for resolving higher-level relationships (*e.g.*, across angiosperms, Zuntini et al., 2024; all vertebrates, Fong & Fujita, 2011), as strong purifying selection at these loci greatly reduces the probability that mutations become fixed, slowing the accumulation of substitutions and minimizing saturation over long evolutionary timescales. In contrast, faster-evolving loci provide greater signal to distinguish among closely related species and have been useful for resolving relationships in rapidly radiating plant lineages such as *Costus (*Costaceae) (Valderrama et al., 2020) and *Oenothera* section *Calylophus* (Onagraceae) (Cooper et al., 2023). Furthermore, when investigating microevolutionary relationships (*i.e.*, population genomics), the accuracy of population assignments improves as the number of variable loci that distinguish groups increases. For example, Haasl & Payseur (2011) showed that datasets containing 1,000 SNPs, selected with a criterion of minor allele frequency >0.1, predict ancestral population structure (K) with high accuracy; while smaller datasets of only 35 SNPs or 1,000 SNPs without a minor allele frequency >0.1 yielded lower K-values and consistently underestimating the number of populations. Upwards of 1,000 SNPs were required to obtain accurate and robust estimates of heterozygosity in *Amphirrhox longifolia* (Nazareno et al., 2017).

Whole genome sequencing provides the most comprehensive view of genetic variation within and between species by capturing changes across the entire genome. Having whole genome sequences to compare is particularly valuable for studies requiring high-resolution genotypic characterization of particular regions, especially for regions with high repeat densities (e.g. the S-locus; Vekemans et al., 2021). Although whole genome sequencing remains relatively costly, especially to generate datasets with many samples or for species with large genomes, the declining price of sequencing and improvements in computational tools have led to resequencing being a cost-effective option for species with genomes around 1Gb or smaller (Song, Buckler, et al., 2023). Genome resequencing has enabled significant advances in addressing evolutionary questions within and between species such as characterizing the divergence of traits in *Vitis vinifera* from wild relatives during crop domestication (Zhou et al., 2017), phylogeographic inquiries regarding how ecological forces sculpt genetic divergence (Hou et al., 2020), testing if ancient admixture or recent gene flow are driving phylogenetic discordance (Stubbs et al., 2023), and the ability to identify sex-associated SNPs across multiple species within *Salix* (Wang et al., 2022). We define genome resequencing as sequencing with a coverage depth of 10–20x, which provides sufficient resolution for robust genotype calls across the genome (Kardos & Waples 2024) and “genome skimming” being the term used for data with 0.01-4x coverage (Straub et al., 2012). While the lower coverage of genome skimming introduces uncertainty in genotype calls, it is generally adequate for recovering organellar and high-copy genes and thus can produce a viable dataset depending on the sampling and scale of question.

In contrast to sequencing entire genomes, a common method to obtain molecular markers for evolutionary studies, particularly in phylogenomics, is target enrichment or target capture (Yardeni et al., 2022). This approach uses ultra-conserved elements (UCEs) (Faircloth et al., 2012) or taxon-specific regions to identify putative single-copy loci from transcriptomes (Weitemier et al., 2014; Sass et al., 2016) or transcriptome and genomic regions of closely related species (Gates et al., 2021). Generally, this approach is used for phylogenomic analyses above the species level (Ball et al., 2023; Pyrcz et al., 2023) however, recent studies have used this approach for population genomic analyses (Boluda et al., 2021; White et al., 2021). Those interested in using UCEs should note that these loci and other single-copy orthologous loci can be highly conserved due to their critical role in fitness and can therefore be subject to strong purifying selection (Faircloth et al., 2015). This high conservation makes them particularly easy to identify as homologous and to align across deep time and therefore effective for resolving deeper evolutionary relationships among broad taxonomic groups. However, in rapidly radiating lineages, these markers might be too conserved to provide sufficient evolutionary signal to resolve relationships (Giarla & Esselstyn, 2015). Additionally, purifying selection over time can increase the potential for convergence (homoplasy), causing different species to independently evolve similar amino acid sequences (Edwards, 2009). To address these limitations, incorporating supercontigs (the combination of overlapping contigs that include both exons and any captured intronic regions) can increase sequence variation for known homologous loci and thus provide more phylogenetic signal than using the conserved marker alone (Bagley et al., 2020).

The choice of loci, whether under strong selection (adaptive or purifying) or in neutrally evolving regions, combined with the taxonomic scale of the sampling directly influences the quantity of loci necessary for reliable genetic analysis and to resolve evolutionary relationships. Advances in sequencing technologies and the increased availability of high-throughput platforms now enable researchers to generate datasets capable of simultaneously addressing both macro- and microevolutionary questions (Villaverde et al., 2018). However, performance assessments of different marker types have been focused on feasibility of either micro- or macroevolutionary questions (Slimp et al., 2021; Szarmach et al., 2021; Mo et al., 2025). At the micro-evolutionary scale, studies demonstrate that genome resequencing provides a more accurate assessment of population structure due to its continuous genomic coverage allowing for the detection of fine-scale introgression events (Yoshida et al., 2016; Sandercock et al., 2022). In contrast, reduced-representation methods can miss these events because of their limited and uneven sampling of loci (Szarmach et al., 2021). While previous studies have demonstrated the advantages of genome resequencing among species undergoing rapid radiations (Valderrama et al., 2022; de Jong et al., 2023), we further explore the precision of different marker types to SNPs across the genome via genome resequencing for population genomic and phylogenomic metrics.

Using published UCE probe sets for plant and animal clades and single-copy orthologous genes as a proxy for taxon-specific probes, we tested if reduced representation loci are equally informative to, and concordant with, genome-wide markers pulled from genome resequencing data in terms of commonly inferred metrics and support for both micro- and macroevolutionary studies. We first test the impact of SNP calling programs and filtering paradigms on SNP retention in order to investigate the best approaches to build a SNP dataset from either whole genome or reduced representation sequences. Next, we assess the ability of various SNP datasets to recover well-supported and topologically congruent phylogenies in an effort to detect their ability to generate signal for the study of macroevolutionary patterns. We then investigate the ability of SNP datasets to reveal microevolutionary patterns by comparing estimates of heterozygosity, population-level clustering along principal components, and detection of ancestral contributions to current population structure. Evaluating the reliability of data generated through reduced representation approaches compared to datasets generated by comprehensive genome resequencing will guide researchers in efficiently allocating resources to produce high-quality data for their biological studies.

## Results

To generate the data for this study, we use *in silico* methods to call SNPs for seven different marker types (UCE, UCE plus supercontigs, BUSCO, BUSCO plus supercontigs, single copy, single copy plus supercontigs, and genome-wide) from publicly available genome resequencing data sets (Fig. 1). These marker types were selected because they represent commonly used approaches in phylogenomics that are increasingly being used in population genomics, varying in terms of genomic distribution, constraint, and informativeness (Faircloth et al., 2012; Simão et al., 2015; Zhang et al., 2020). We then applied a consistent filtering pipeline across marker types, including removal of low-quality variants and filtering for linkage disequilibrium (LD), to retain largely unlinked, high-confidence SNPs.

**Figure 1.**
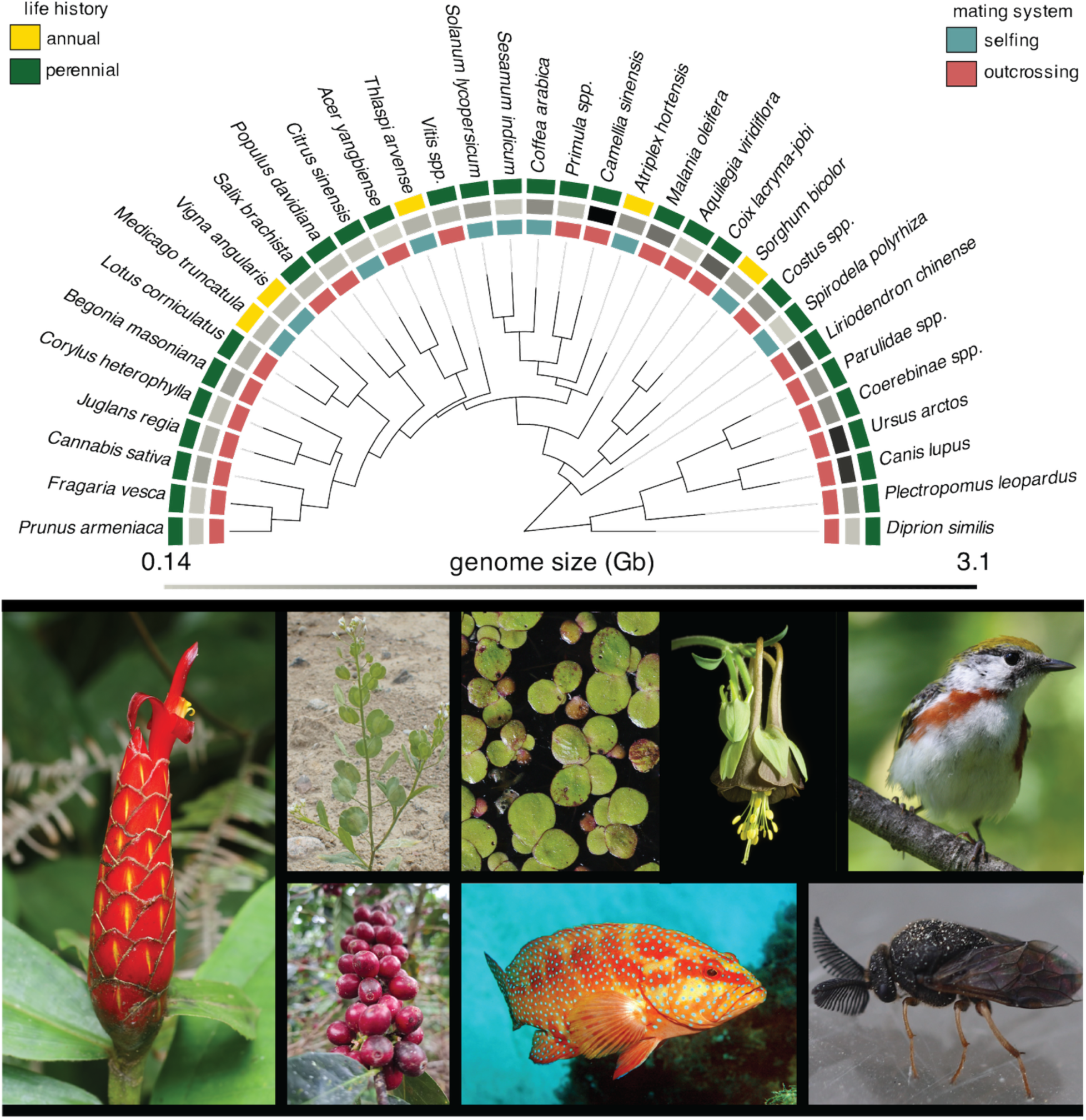
Phylogenetic distribution of the taxa analyzed in this study, highlighting variation in mating system, life history, and genome size. Genome size spanned from 0.14 – 3.1 Gb, (ii) mating system, with teal indicating selfing and red indicating outcrossing, and (iii) life history, with yellow marking annuals and dark green marking perennials. Photographs below the tree highlight the morphological breadth of the sampled taxa. This phylogenetic framework anchors all subsequent comparisons of SNP retention and phylogenetic resolution across loci types Photo credits (clockwise from top left): J. Smith, *Costus spiralis* inflorescence (Wikimedia Commons, CC BY 2.0); L. Chen, *Thalpsi arvense* rosette (Wikimedia Commons, CC BY 2.0).

We hypothesized that after filtering for linkage disequilibrium (LD), the number of SNPs derived from UCEs and BUSCOs would be reduced compared to whole-genome SNPs due to overall size of the original dataset. We also hypothesized that the recovered SNPs from reduced representation loci would provide less phylogenetic resolution for closely related species or rapidly evolving lineages due to the inherently conserved nature of targeting only annotated genes which are typically highly conserved. For phylogenetic inference, we anticipated that while broad evolutionary relationships will be consistent across different marker types, the level of support for deep nodes and nodes with short branches will likely vary depending on the number of loci and the number of informative characters within each locus. Datasets from whole-genome resequencing include rapidly evolving regions and thus may offer more robust resolution of recent nodes (Xu et al., 2022), while reduced representation datasets having more conserved loci may provide strong resolution at most evolutionary levels but struggle with relationships below the species level due to their slower evolutionary rates and lack of informative characters. At the population level, we hypothesize that SNPs recovered from resequencing data will be able to capture fine-scale population differentiation due to their broader representation of the genome, while SNPs recovered from the capture of conserved loci might reveal a narrow view of population structure. We also predict that certain species-specific traits, such as reproductive strategies (i.e. selfing v. outcrossing) and genome size, will influence SNP retention after filtering for linkage due to differential recombination potential inherent to these traits.

### Impact of SNP calling and filtering on retention

SNP retention refers to the number of high-quality, informative single nucleotide polymorphisms that remain after variant calling and subsequent filtering steps. In non-model species, where reference genomes may be incomplete or divergent, many SNPs are often discarded due to low coverage across samples, low certainty of the variant call, as well as proximity to other variants. This can affect downstream analyses of genetic diversity, population structure, and evolutionary relationships. Given recent discussions in the literature about the best approach for SNP calling in non-model species, we first compared SNP retention with BCFtools (Danecek et al., 2021) and GATK (Van der Auwera & O’Connor, 2020). Using identical filtering criteria, we found that BCFtools retained more high-quality SNPs than GATK (Fig. S1). Of the total SNPs detected, 101,859 (39.7%) were identified by both methods. BCFtools uniquely retained 97,425 SNPs (37.9%), while GATK uniquely retained 57,547 SNPs (22.4%). For the remainer of this study we largely proceeded using datasets generated with GATK, partially due to computational constraints where GATK’s joint genotyping was beneficial. Due to the reasonable similarity between the two programs, for several of the genome-wide datasets involving species with large genomes, BCFtools was used to ensure full completion of the SNP calling pipeline (Table S1).

Across all datasets analyzed, marker type had a significant impact on SNP retention (χ² = 156.22, p < 0.001). Post hoc Dunn’s tests revealed that all reduced-representation datasets retained significantly fewer SNPs compared to the genome-wide dataset (p < 0.001). Among the reduced datasets, BUSCO, and single-copy loci (with and without supercontigs) retained a similar number of SNPs across all four categories, while UCE loci exhibited the lowest SNP retention, compared to all other datasets (p < 0.001). The inclusion of supercontigs increased UCE SNP retention (p < 0.001), however UCEs with and without the supercontigs still had the fewest SNPs compared to other reduced representation marker types (Fig. 2) (Table S2).

**Figure 2.**
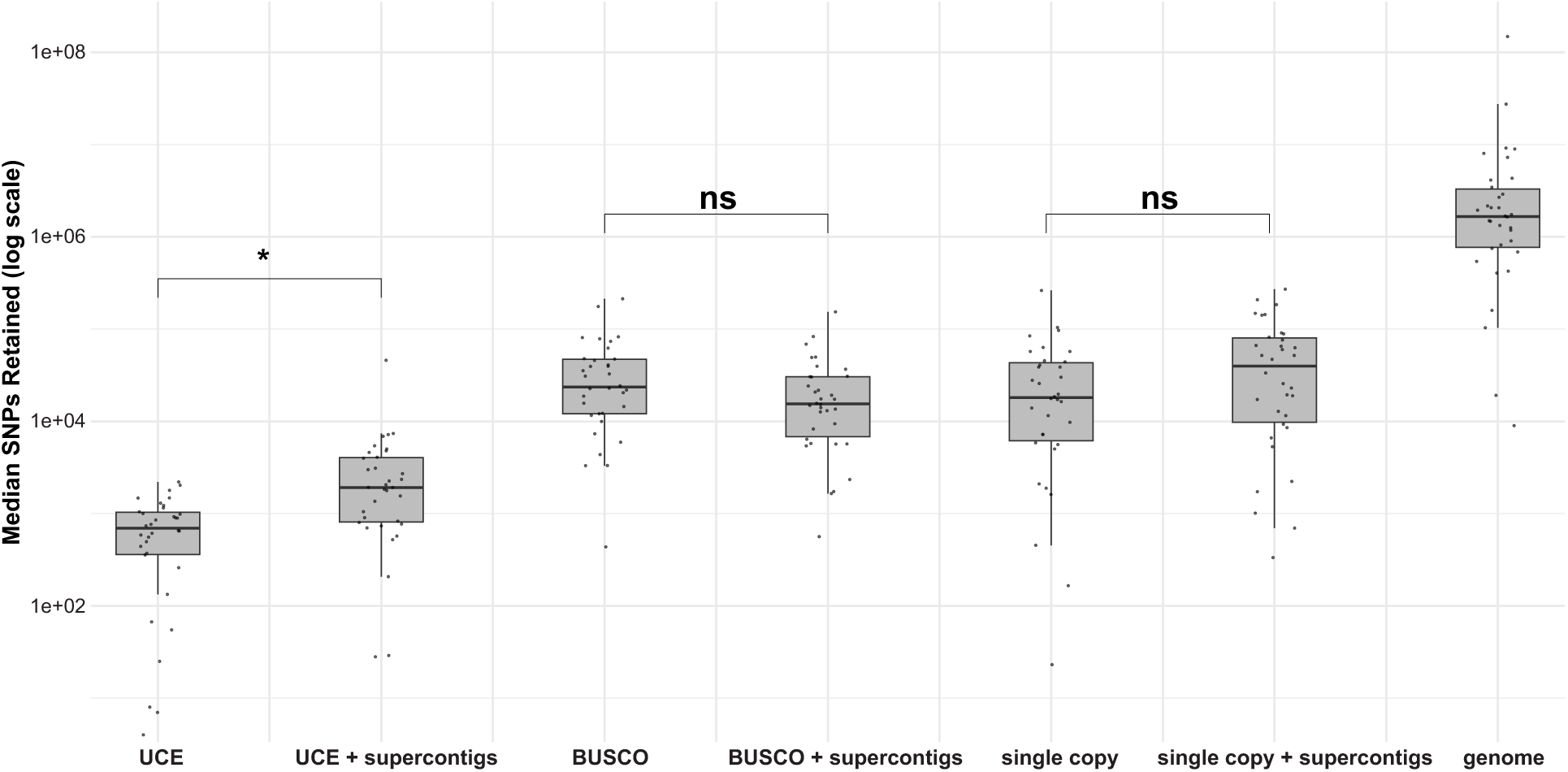
SNP retention across different loci types after filtering for linkage disequilibrium. Genome-wide datasets retained the highest proportion of SNPs compared to all other loci types. BUSCO and single-copy loci, with or without supercontigs, retained a similar proportion of SNPs but fewer compared to genome-wide datasets. UCE loci exhibited the lowest SNP retention, with UCEs with supercontigs retaining slightly more SNPs than UCEs alone. Significance levels are represented as follows: * (p < 0.05), and "ns" (p ≥ 0.05).

### Phylogenetic inference and Topological Congruence testing

To assess whether phylogenetic resolution depends on the number of SNPs rather than the informativeness of the SNP, we pruned the genome-wide dataset to match the number of SNPs in the UCE plus supercontigs dataset. Across the pruned genome replicates there were no differences in nodes shared with the UCE + supercontigs dataset, however all pruned datasets had fewer supported nodes compared to trees inferred from the entire genome SNP dataset (V = 0, p < 0.001) (Fig. S2).

Phylogenetic inference using the SNPs filtered for LD showed that the number of supported nodes (BS ≥ 70 %) varied widely across datasets (F(6, 238) = 3.376, p < 0.00326). Trees inferred from the whole genome dataset had a significantly greater proportion of supported nodes (mean = .741) versus all other datasets (p < 0.0001 for all pairwise comparisons). On average, UCEs had the lowest proportion of supported nodes among the reduced representation datasets (mean = 0.233) albeit the inclusion of supercontigs in the UCE datasets did increase their support (mean = 0.391). In contrast, the BUSCOs yielded the highest proportion of supported nodes (mean = 0.587) across reduced representation approaches. No differences were detected between the BUSCO and BUSCOs plus supercontigs datasets (p = 1.000) Trees generated from the single copy loci (mean = 0.532) and single copy loci plus supercontigs had moderate node support (mean = 0.542), with no significant difference detected with the addition of supercontigs (p = 1.000). (Fig. 3a; S3).

**Figure 3.**
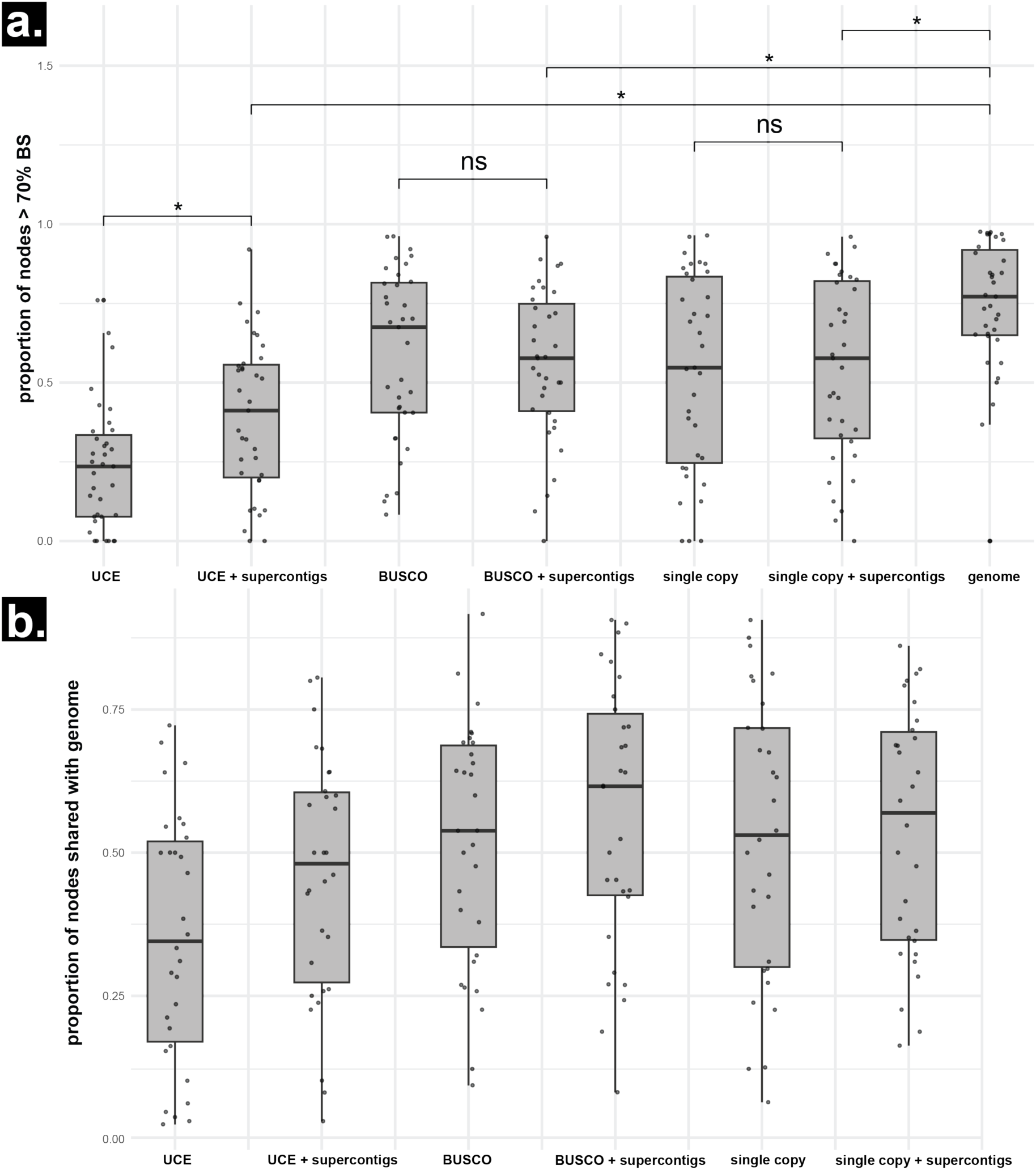
(a) Proportion of well-supported nodes (BS ≥ 70%) across datasets. Phylogenies inferred from the genome dataset had the highest proportion of supported nodes. Among reduced representation datasets, BUSCO genes yielded the highest node support, while UCEs had the lowest. The inclusion of supercontigs increased support for UCEs but had no significant effect on BUSCOs or single-copy loci. Significance levels are represented as follows: * (p < 0.05), and "ns" (p ≥ 0.05). **(b) Proportion of shared nodes (compared to genome-wide SNPs) across datasets.** UCE datasets showed the lowest overlap (mean = 0.352, p < 0.01). Supercontigs improved UCE performance (mean = 0.457) but remained lower than BUSCO and single-copy datasets. BUSCO (mean = 0.516) and single-copy (mean = 0.524) datasets had higher concordance, with modest gains from supercontigs (means = 0.565 and 0.536, respectively). No significant differences were found between BUSCO and single-copy datasets or with inclusion of supercontigs (p > 0.4).

To assess consistency in phylogenetic inference across loci types, we analyzed the proportion of shared nodes between the phylogenies generated with reduced representation datasets and those generated by whole genome data. Trees inferred from UCE data had the lowest mean proportion of nodes in common (mean = 0.352), significantly lower than all other datasets (p < 0.01 for all pairwise comparisons). Inclusion of supercontigs moderately improved UCE performance (mean = 0.457), though the number of shared nodes were still significantly lower than those generated from BUSCO and single-copy datasets (Fig. 4). BUSCO datasets yielded a phylogeny with significantly higher proportion of shared nodes with the whole genome phylogeny as compared to the UCE phylogeny (mean = 0.516), with further improvement when supercontigs were included (mean = 0.565). Similarly, trees from single-copy loci showed relatively high topological congruence with the whole genome dataset (mean = 0.524), and the inclusion of supercontigs with the single-copy loci produced a modest increase (mean = 0.536), though the difference was not significant (p = 1.000). No significant differences in topological congruence were observed between BUSCO and single-copy datasets or between their corresponding supercontig-enhanced counterparts (all p > 0.4 after Bonferroni correction) (Fig. 3b).

**Figure 4.**
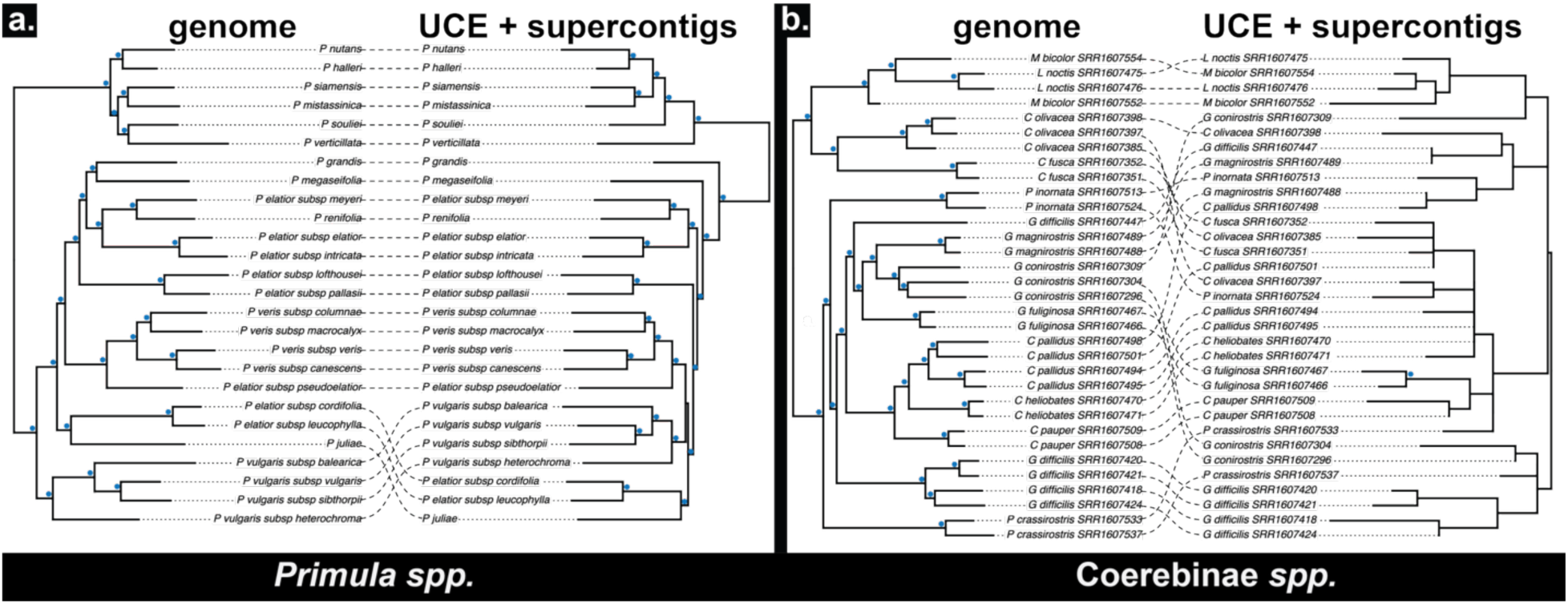
Topological incongruence to genome wide datasets was the most with UCE + supercontigs datasets. Maximum-likelihood trees inferred from genome-wide SNPs (left) are paired with trees built from UCEs plus their flanking supercontigs (right) for (a) *Primula* spp., which showed the smallest departure from the genome-wide topology, and (b) Coerebinae spp., which exhibited the largest. Nodes ≥ 70 BS are labeled with a blue dot. Dashed lines connect identical taxa; the crossings of dashed lines note where there are topological differences.

### Estimation of site-specific heterozygosity

Observed heterozygosity, measured as the proportion of heterozygous sites among all high-quality mapped positions, varied significantly by marker type (Fig. 5). BUSCO sequences exhibited the lowest proportion of heterozygous sites (mean = 0.00706), with whole genome heterozygosity similarly low (mean = 0.00860). In contrast, single-copy sequences yielded the highest proportion of heterozygous sites (mean = 0.0146), with single-copy plus supercontigs producing comparable values (mean = 0.0148) to single-copy alone. UCEs, whether exon-only (mean = 0.00899) or including supercontigs (mean = 0.0108), exhibited intermediate levels of heterozygosity amongst the seven datasets.

**Figure 5.**
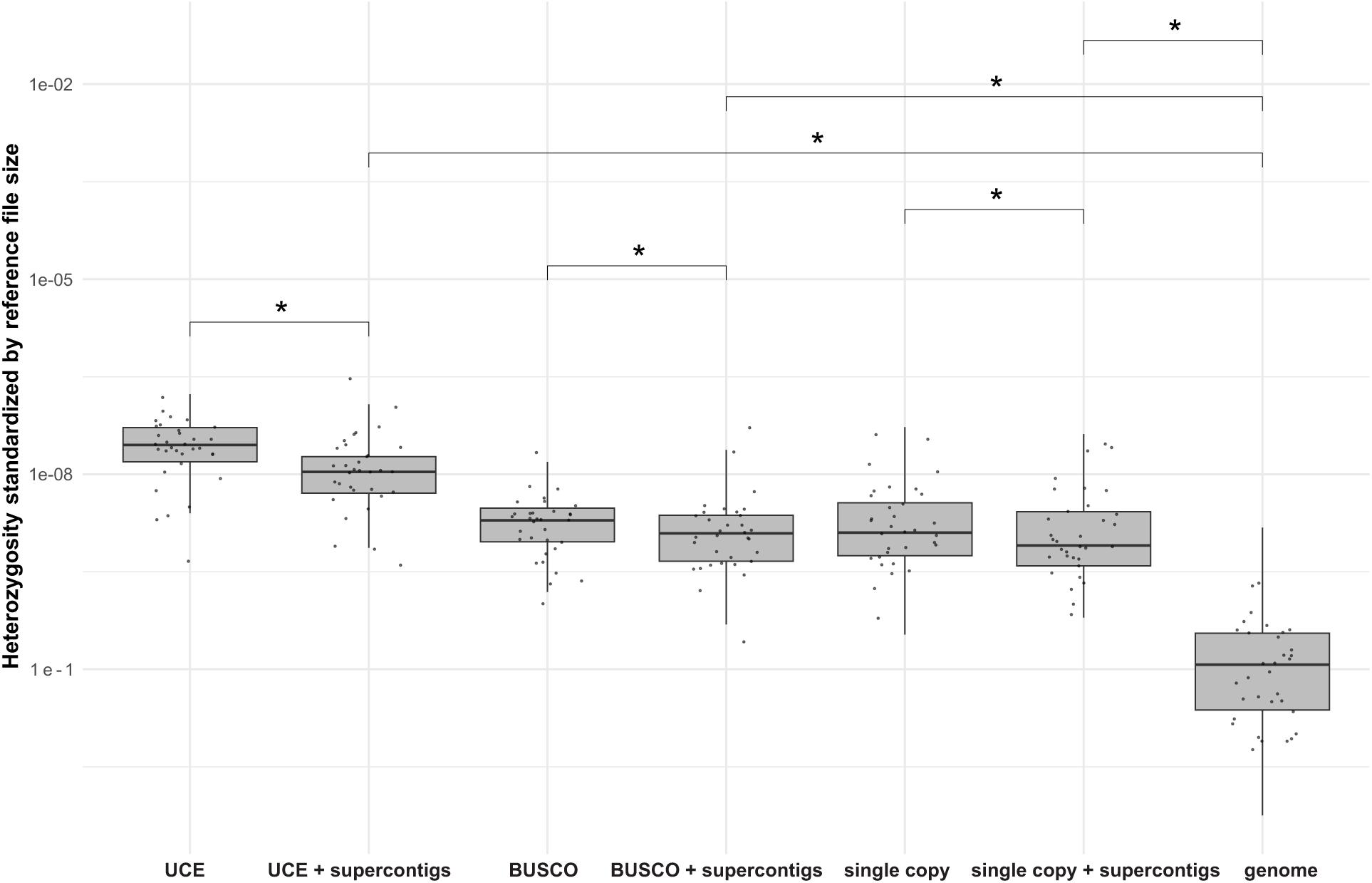
Heterozygosity estimates varied significantly marker types. Genome-wide sequences exhibited the lowest proportion of heterozygous sites. Heterozygosity estimates are normalized by number of sites in the reference file for each marker type. Significance levels are represented as follows: * (p < 0.05), and "ns" (p ≥ 0.05).

Pairwise Wilcoxon tests further supported the observed differences in heterozygosity among dataset types, revealing significant differences in most comparisons (Table S3). BUSCO sequences had significantly lower heterozygosity than all other datasets (p < 0.0001 for UCE, single-copy, and genome comparisons). UCE sequences, with or without supercontigs, showed significantly lower heterozygosity than single-copy datasets (p < 0.0001). The inclusion of supercontigs generally increased heterozygosity across marker types, particularly in single-copy loci. Notably, comparisons between BUSCO plus supercontigs and UCE plus supercontigs, as well as between single-copy and single-copy plus supercontigs, were not statistically significant (p = 1.000).

### Inferring population clusters

#### a. Clustering based on principal components and explained variance

Principal Component Analysis (PCA) are often used to reveal clustering of populations or other evolutionary units based on observed variation in the dataset. Such clustering can be used to assign relevant spatial or temporal characteristics. For all tested datasets, the variance among units explained by PCs 1–4 did not differ significantly between dataset types (Fig. S4). While the variance explained by PCs was generally uniform across datasets, the amount of variance explained by the first PC (PC1) varied across species, reflecting differences in genetic structure. PC1 accounted for less than 10% of the total variance in *Sesamum indicum* and *Corylus heterophylla* (9.52% and 9.82%, respectively), suggesting weak or diffuse genetic structure. In contrast, species like *Atriplex*, *Fragaria*, and *Spirodela* showed much higher PC1 variance (53.65%, 43.7%, and 25.57%, respectively), indicating more pronounced genetic differentiation captured along a single axis (Fig. S5). Clustering with 95% confidence ellipses across datasets based on the first two PCs found that UCEs showed significant overlap across clusters, highlighting the limited variation in those loci and indicating a potential lack of ability to resolve variance among distinct populations (Fig. 6).

**Figure 6.**
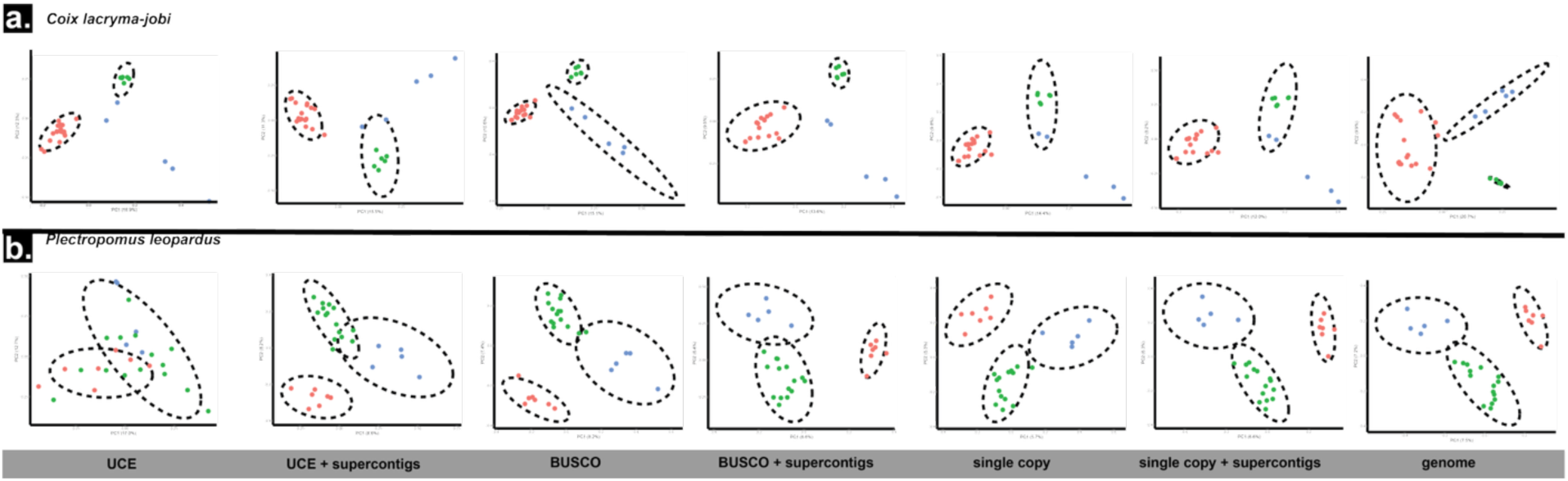
Genetic clustering based on the first two principal components. PCA plots are shown for (a) *Coix lacryma-jobi* and (b) *Plectropomus leopardus.* Individuals are colored according to kmeans clusters defined from genome wide SNPs, and dashed ellipses represent 95 % confidence limits calculated from SNPs in each dataset. UCE SNPs provide the least clustering resolution.

#### b. Detecting ancestral (K) and current population structure

To assess how different marker sets influence the detection of population structure, we compared both the number of inferred ancestral populations (K) and the estimated ancestry proportions across datasets. Here, K refers to the number of genetic clusters inferred the model-based population analysis STRUCTURE, representing historical gene pools that have contributed to the genetic makeup of the individuals in the dataset.

Across taxa, reduced-representation panels under- and overestimated K by compared to genome-wide SNPs (Fig. S6). When we fixed K at the genome-wide optimum, ancestry coefficients from BUSCO and single-copy loci were largely congruent with those from genome-wide data; UCE-only analyses, however, failed to detect minor admixture proportions in samples where one cluster represented over 90% of ancestry (Fig. 7; Fig. S7. In these cases, UCE-derived profiles collapsed individuals into a single homogeneous cluster, whereas genome-wide SNPs revealed residual structure.

**Figure 7.**
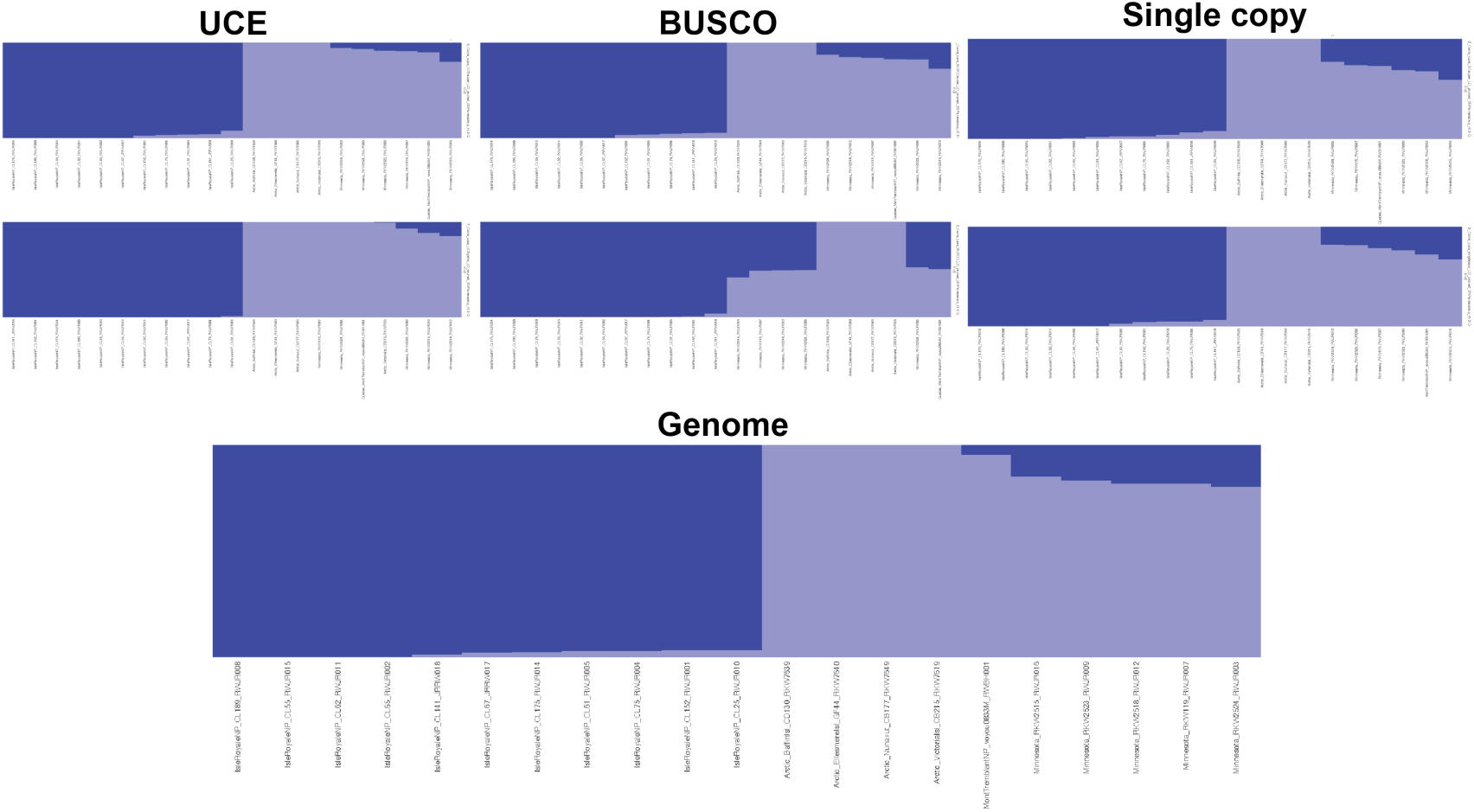
Reduced representation yields largely congruent model based genetic structure clustering. Structure plots for *Canis lupus* at K = 2. Supercontigs were included in the top plot for each reduced representation approach.

## Discussion

In this study, we evaluate the congruence between SNPs obtained from reduced representation markers and those obtained from whole genome sequencing data for population genomic and phylogenomic analyses. Our analyses reveal that marker choice exerts a pervasive influence on both phylogenetic and population-genomic inferences. Across reduced representation datasets, UCEs showed the lowest congruence with genome-wide SNPs with results that varied significantly with respect to heterozygosity estimates and population clustering. We thus recommend against using UCEs for population level inferences. However, in light of UCE topological incongruence and lack of support for phylogenetic results, we observed that incorporating supercontigs generally improved phylogenetic resolution.

Genome resequencing offers the most comprehensive dataset for studying evolutionary processes, capturing variation in both coding regions and noncoding sequences. Resequencing also allows for the recovery of the organellar genomes, either through SNP calling or assembly with programs such as getOrganelle (Jin et al., 2020); whereas with hyb-seq, uncaptured barcoded libraries must be pooled and sequenced with the captured libraries with an approximate 70/30 (Vera-Paz et al., 2022) or 3:1 (Hatami et al., 2022) ratio to ensure enough reads mapping to the organellar genomes to allow assembly or SNP calling due to higher efficiencies of the recent probe kits (Devault & Enk, 2023).

Linkage disequilibrium (LD) describes the nonrandom association of alleles at different loci due to proximity on the chromosome, genetic drift, selection, or past demographic events (Slatkin, 2008). Because LD creates statistical dependence among SNPs, many SNP-based phylogenetic and population-genetic methods, such as SVDquartets (Chifman & Kubatko, 2014), SNAPP and SNAPPER (Bouckaert et al., 2019), D-statistics (Green et al., 2010, Durand et al., 2011, Hahn & Hibbins, 2019) and HyDe (Blischak et al., 2018), require LD pruning to satisfy the assumptions of independence. While LD filtering is essential in population genomics, we argue that LD pruning is equally necessary for SNP-based phylogenomic approaches. The theoretical foundation of bootstrapping for maximum likelihood phylogenetic trees assumes that sites in a sequence alignment evolve independently (Felsenstein, 1985). In real datasets, physical linkage causes alignment sites to evolve together rather than independently, resulting in redundant signal that can artificially inflate bootstrap support values and exaggerate support on short internal branches in genome-scale alignments with millions of sites (Galtier, 2004; Roycroft et al., 2020). For SNP-based datasets, we suggest filtering for linkage to mitigate the effects of non-independence on support values. Filtering for LD also decreases computational time by excluding unnecessary genetic data from large datasets, reducing data sets by millions of sites in some cases (Calus & Vandenplas, 2018). Moreover, even after LD pruning, invariant sites and clusters of linked sites can still distort branch-length estimation and downstream analyses; to address this, one can incorporate a sampling fraction during tree inference (Minh et al., 2020) or turn to divergence-time estimation frameworks that model rate variation (Suissa et al., 2023).

### Macroevolutionary analyses

Of all reduced-representation markers, single-copy and BUSCO loci most closely recapitulated the genome-wide topologies for phylogenetic and clustering analyses, whereas UCEs showed greater incongruence (Fig. 3; 4; S5). Our results align with Cooper et al. (2023) and Valderrama et al. (2020), who found that taxon specific probes were effective to resolve recently diverged clades in *Oenothera* and *Costus*. BUSCO markers are routinely used to assess genome assembly quality and can be extracted through a simple pipeline, making them valuable for deep-time evolutionary studies (Alam et al., 2025). In contrast, UCEs, despite their appeal as universal markers, often lack the resolution required for more recent divergences. While they have proven highly effective at deeper taxonomic levels (e.g., family and order), their utility at the species level is mixed and often limited below the species level. This limitation is also apparent in species radiations, such as within Darwin’s finches in the Coerebinae (Lamichhaney et al., 2015) (Fig. 4b).

The Angiosperm353 was designed to be a universal probe set for targeted sequencing in any angiosperm group, from species-level studies to the entire Angiospermae (Johnson et al., 2019). Previous studies have shown that the Angiosperm353 are able to confidently resolve higher-level phylogenetic relationships within the Gentianales, Sapindales, Dipsacales and within the rosids (Antonelli et al., 2021; Lee et al., 2021; Joyce et al., 2023; Zuntini et al., 2024). However, within genera, Angiosperm353 probes are not a one-size-fits-all solution. In the rapidly radiating clade *Veronica* sect. *Hebe,* the median gene concordance factor (gCF) ranged from 0 to 2.13% and the median site concordance factors (sCF) were similarly low across their data sets with medians between 36.2 and 37.8% (Thomas et al., 2021). Similar low concordance factors were found in Ochnaceae, where bootstrap support was 100% or close across many genera but gCFs were below 50% (Shah et al., 2021). In our *in-silco* capture of UCE probes, some lineages, such as *Corylus* fared better with UCEs (Fig. S5), but overall UCEs often fail to capture the full phylogenetic signal in complex groups and below the species level. Including supercontigs did help increase signal across all datasets, boosting the average contig length in our datasets by as much as 242%. Nonetheless, UCEs with supercontigs only represented 0.08% (*Geospiza*) to 47% (*Fragaria*) of each total genome size, highlighting the lack of genomic spread of UCEs.

To determine whether phylogenetic resolution is primarily influenced by the informativeness of specific loci or simply by the total number of sites analyzed, we downsampled genome-wide SNP datasets to match the number of sites found in our UCE + supercontig datasets. This approach allowed us to disentangle the effect of locus type from the number of sites. Universal probe sets span only a few hundred to a few thousand loci (e.g., 353 in angiosperms, 5,060 in Tetrapoda, 1,172 in Coleoptera; Johnson et al., 2019; Faircloth et al., 2012). Our downsampling results indicate that phylogenetic resolution scales primarily with the total number of independent sites rather than with specific characteristics of targeted loci. While it is true that some targeted loci may contain highly informative sites that disproportionately impact tree topology (Shen et al., 2017), the breadth of genome-wide SNP data more reliably captures enough unlinked variation to resolve recent divergences. These findings suggest that future phylogenetic studies using reduced-representation methods should consider employing larger or taxon-specific probe sets to maximize the number of informative sites and improve resolution.

### Microevolutionary analyses

BUSCOs inferred heterozygosity estimates were similar to the entire genome, both of which found lower heterozygosity than other reduced representation markers (Fig. 5). ANGSD derives heterozygosity from the site frequency spectrum across all high-quality mapped sites in the BAM files, not solely from variant positions (Korneliussen et al., 2014).

The *in-silico* UCE and single-copy datasets, by design, sample shorter, gene-centric regions and thus exclude large tracts of invariant sequences. UCEs (especially when including supercontigs) and single-copy orthologs incorporate more neutrally evolving or moderately constrained flanking sequences. Those regions can harbor both rare and intermediate-frequency alleles, boosting genetic diversity relative to BUSCOs and the genome-wide dataset. Although whole-genome resequencing encompasses both conserved and variable regions, the total number of sites passing the threshold swells with many filtered invariants due to missing data, minor allele frequency or those in linkage disequilibrium, attenuating the heterozygosity signal.

Our heterozygosity incongruence highlights the critical impact of marker choice in heterozygosity assessments. UCEs and single copy orthologous genes inherently concentrate on variable windows and under-represent invariant sites, artificially inflating heterozygosity. This bias has important downstream consequences: overestimated heterozygosity can obscure signals of recent bottlenecks or inbreeding, and meta-analyses that aggregate heterozygosity across studies may misjudge relative levels of genetic diversity (Alves et al., 2023).

Complementary metrics such as allelic richness, F-statistics, and admixture statistics (Leroy et al., 2018) should accompany heterozygosity estimates to provide a comprehensive view of genetic diversity.

The variance explained by principal components was largely congruent across all datasets (Fig. S3), highlighting the ability of reduced representation markers in capturing population structure from model-free methods. With dimensionality reduction methods such as PCA, proximity reflects genetic similarity, and UCEs did not provide enough discriminatory power to resolve fine-scale differences that genome-wide data revealed; information crucial for subsequent FST analyses (Fig. 6b). Thus, while UCEs generally recover major clusters (Duckett et al., 2023), their resolution weakens at finer scales, particularly below the species level. Reduced-representation datasets consistently underestimated the true number of ancestral populations (*K*) compared to genome-wide SNPs. Because methods like STRUCTURE rely on allele frequency differences, their power to detect subtle population structure declines as the number of loci decreases (Graham et al., 2020). In our best-K comparisons (Fig. S6), UCE and BUSCO loci frequently collapsed distinct subpopulations into a single cluster. The underestimation of *K* has significant implications, as it can obscure the true complexity of population structure leading downstream conservation impacts such as over-harvesting, loss of unique genetic diversity and misallocation of conservation resources (Cullingham et al., 2020). The differences in ancestry proportions across reduced representation approaches may potentially be due to the effects of balancing selection, as certain loci maintained by selection may obscure demographic patterns that are more apparent in neutrally evolving regions. Future studies that subset the genome into different loci should further investigate this trend.

### Suggestions

Our findings imply that researchers seeking to resolve recent divergences, detect fine-scale structure, or estimate genetic diversity should prioritize genome-wide resequencing whenever feasible. Genome resequencing is already relatively inexpensive for organisms with genomes smaller than one gigabase (Song, Ning, et al., 2023). However, large genomes pose significant computational and financial challenges, making strategies such as genotyping-by-sequencing (GBS) or Hyb-Seq more practical alternatives (Dodsworth et al., 2019). Large genome sizes can also complicate approaches like genome skimming due to increased data volume and deeper sequencing requirements, whereas capture-based methods like Hyb-Seq are generally unaffected by genome size (Pezzini et al., 2023). Researchers should be mindful that low-coverage or RAD-seq reduced-representation datasets, though efficient for large genomes, can introduce batch effects that reduce consistency and reproducibility compared to full resequencing (Leigh et al., 2018; Lou & Therkildsen, 2022). Nonetheless, in studies spanning deep time, UCEs remain highly valuable. Looking forward, we anticipate a future in which inference of demography, phylogeny, and adaptive evolution from a single dataset becomes standard; resequencing approaches will be crucial to ensure that newly generated accessions or species can be seamlessly integrated into these pipelines, enabling continual expansion and updating of analyses. For now, our work provides a practical framework for choosing markers. If one’s question focuses on microevolutionary processes such as local adaptation, introgression, or cryptic structure, genome-wide SNPs are indispensable. When budgets limit sequencing depth, carefully designed taxon-specific probes with supercontigs are a better choice compared to UCEs in resolving both shallow and deep nodes, albeit with reduced power for population-level inferences.

## Methods

### Data sampling and selection

We obtained publicly available genome sequencing datasets from the Sequence Read Archive (SRA) and other nucleotide archives (e.g., ENA), with our main inclusion criterion being datasets of 25-50 accessions; some studies were focused on relationships between species while others were below the species level. To ensure a comprehensive dataset, information on reproductive strategies (outcrossing vs. selfing), life history traits (annual vs. perennial), genome size (small: > 0.5 Gb, medium 0.5 to 1.5 Gb, large: < 1.5 Gb), and divergence times from TimeTree 5.0 (Kumar et al., 2022) were collected for each taxon. A summary of this data can be found in Table S1. Raw reads were cleaned using fastp version 0.23.4 (Chen et al., 2018) with automatic detection of adapters, a minimum quality threshold of 20, and a minimum length of 75 bp. Reference genomes were either retrieved using information from the original study or from the NCBI genome database (Table S1).

### Dataset generation

We generated each dataset *in silico* following the specific steps outlined below.

#### BUSCO loci

To create Benchmarking Universal Single-Copy Orthologs (BUSCO) reference sequences, 10 lineage specific libraries were selected based on their taxonomic relevance to the study organisms (e.g. embryophyta, insecta, tetrapoda) with BUSCO version 5.5.0 (Manni et al., 2021). Genome mode was used with the Metaeuk pipeline to generate sequences, references, spanning complete, duplicated, and fragmented sequences, which were concatenated into a single file. To create a BUSCO reference that included flanking regions, we used the retrieve sequences function with the supercontigs argument in Hybpiper version 2.1.6 (Johnson et al., 2016) on a subset of samples. When selecting samples to be used for each of our taxonomic datasets, we did not include a priori information on population structure or phylogenetic topology, however we used samples that had the highest amount of data and that represented a spread across species within the dataset to maximize loci recovery to retrieve supercontigs using hybpiper. We did not include a priori information on population structure or phylogenetic topology; however, we used samples that had the highest amount of data and that represented a spread across species within the dataset to maximize loci recovery. For each locus, the longest supercontig was extracted using a custom script using bioawk (see data availability statement). These supercontigs were then concatenated into a final reference file, which was used for subsequent analyses for our BUSCO plus supercontigs dataset.

#### Ultra Conserved Elements (UCE) loci

For angiosperm datasets, we used the Mega 353 target file (McLay et al., 2021) with all included targets. For non-plant datasets, lineage appropriate UCE target files were used (Table S1). Hybpiper was run on a subset of samples to generate a reference file for the UCE loci and for the UCE supercontigs using the same steps as for the BUSCO reference files.

#### Single copy orthologous genes

Protein-coding reference genome sequences (CDS files) were downloaded from Genbank and RNA-Seq raw reads for focal taxa and closely related species were downloaded from SRA (Table S1). *De novo* transcriptome assemblies were generated, keeping contigs that were a minimum of 300 bp using Trinity version 2.14.0 (Grabherr et al., 2011). Following assembly, we generated a supertranscripts file using the gene splice modeler. To identify single copy orthologous sequences, we used OrthoFinder version 2.5.4 with default settings except DIAMOND for sequence similarity (Emms & Kelly, 2019). The longest single copy orthologous contig was extracted and concatenated into a reference file. To remove any overlap with our BUSCO reference, a reciprocal BLAST search (Altschul et al., 1990) was performed between the BUSCO reference and the single copy reference, retaining only non-reciprocal hits for our single copy reference as reciprocal hits were assumed to have been included in our BUSCO reference. We then used the reference as a target file to extract supercontigs which were then concatenated into a single copy reference with supercontigs (Fig. 8).

**Figure 8.**
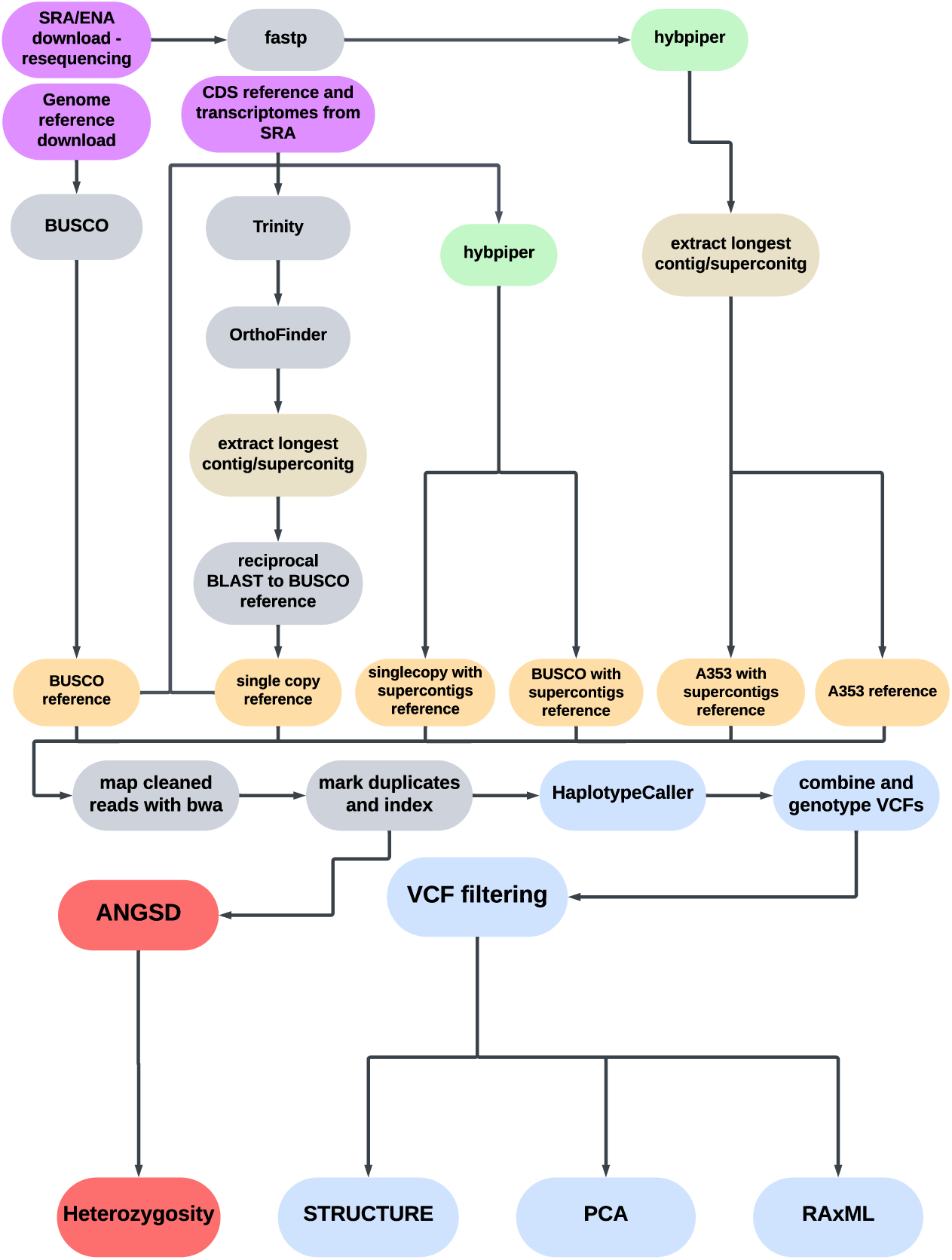
Data processing workflow for downstream SNP analysis. Each marker type was extracted in silico from publicly available resequencing data to generate reference files. Supercontig reference files were constructed by selecting the longest supercontig from hybpiper for each sequence.

### Read mapping and SNP calling

Resequencing reads were aligned to the appropriate reference using bwa-mem2 version 2.2.1 (Vasimuddin et al., 2019), followed by converting outputs to BAM and sorting using samtools version 1.19.2 (Danecek et al., 2021). Duplicate reads were marked using piccard (http://broadinstitute.github.io/picard/) bundled with GATK version 4.2.3.0 (Van der Auwera & O’Connor, 2020) and then indexed with samtools. For a subset of taxonomic datasets (*Spirodela* and *Costus*), a comparative analysis of SNP calls was performed using HaplotypeCaller from GATK version 4.2.3.0 and mpileup from BCFtools version 1.19.2 (Danecek et al., 2021) to assess the number of SNPs retained after filtering. This approach was motivated by two factors: first, recent studies have questioned the suitability of GATK in non-model systems, suggesting it may produce a higher proportion of false positives(Lefouili & Nam, 2022); while GATK does produce more false positives, with hard filtering, GATK produced fewer false positives. Due to computational constraints (i.e., runtimes and storage requirements), BCFtools mpileup was used for large-genome SNP calling to facilitate timely analysis. We have noted in Table S1 the datasets where SNPs were generated with BCFtools. Individual gVCF files were combined using GATK’s GenomicsDBImport (version 4.2.3.0). Joint genotyping was then performed using GATK’s GenotypeGVCFs, generating a final VCF file that contained SNP calls for all samples.

### SNP filtering

To retain only high quality, unlinked biallelic SNPs the following filtering parameters were applied: Maximum missing data threshold of 0.7 was selected to retain loci with sufficient representation across individuals, a minor allele frequency threshold of 0.05 was used to remove highly rare alleles, with a minimum of 2 alleles and a maximum of 2 alleles per site, and read depth filters (minimum 5, maximum 200) were chosen to balance the inclusion of informative loci and the exclusion of potential sequencing artifacts or coverage biases. To evaluate differences in SNP retention across dataset types, we conducted a Kruskal-Wallis rank sum test in R version 4.4 (R Core Team, 2024) using the kruskal.test function. Pairwise comparisons were performed using Dunn’s test with Bonferroni correction in the FSA package, version 0.9.4 (Ogle et al., 2025) and visualized with ggplot2 version 3.5.1 (Wickham, 2016).

### Linkage disequilibrium (LD) inference and plotting

Linkage disequilibrium (LD) was inferred by calculating r² values for SNP pairs using POPLDDECAY version 3.40 (Zhang et al., 2019) For each dataset, we calculated r² values and plotted the LD decay curve using the Plot_OnePop.pl script provided with PopLDdecay, SNPs were then filtered based on the threshold where the LD plot curve approached its asymptote (Table S1) in Plink version 1.9 (Chang et al., 2015) using a 20 kb window with a step size of 10 kb. For downstream analyses, we converted the VCF file to FASTA format using vcf2phylip version 2.8 (Ortiz, 2019) to generate a sequence alignment. To prepare the dataset for population structure analysis, we generated a STRUCTURE file by running the filtered VCF with the populations program in Stacks version 2.66 (Catchen et al., 2013). For larger genome wide datasets, VCF files were thinned to 100,000 SNPs using the --thin-count option in PLINK (Chang et al., 2015) before being converted back into VCF format for downstream analyses to reduce computational time without introducing biases due to too strict of filtering (Suissa et al., 2023).

### Phylogenetic inference

Maximum likelihood trees were inferred with RAxML-NG version 8.2.11 (Kozlov et al., 2019) under a genotypic model of evolution where all substitution rates were treated as distinct (GTGTR) with 100 bootstrap replicates. While this parameter-rich model increases computational demands, it was chosen to maximize model realism given the SNP data used in this analysis. To evaluate topological congruence between trees derived from different datasets, we used the comparePhylo function in the R package ape version 5.8 (Paradis & Schliep, 2018). To compare the proportion of shared nodes between trees inferred from reduced representation datasets to the entire genome dataset, we conducted post-hoc pairwise comparisons of supported nodes using the *emmeans* version 1.10.4 in R (Lenth, 2024) with a Tukey adjustment for multiple comparisons.

To determine whether the robustness of our genomic phylogenetic analyses depends on the number of characters or the specific sites used, we downsampled our genome VCF file for the *Populus davidiana* and *Costus* datasets to the number of SNPs retained after filtering for each of their respective Angiosperm353 with supercontigs VCF using JVarkit (https://github.com/lindenb/jvarkit). We ran 1000 iterations for each species, which randomly selected sites from the file to create the subset files. The VCF subset files were converted to a FASTA file using vcf2phylip version 2.8. Phylogenetic trees were inferred using RAxML-NG version 8.2.11 with GTGTR and 100 bootstrap iterations. Unique topologies were scored across the downsampled trees using phytools version 2.3-0 (Revell, 2024) and compared with a Wilcoxon rank sum test in R.

We evaluated the effect of marker type on the number of supported nodes (>70% BS) using a linear mixed-effects model from lmerTest version 3.1-3 in R Statistical Software (v4.4.0; R Core Team, 2024). The model included marker type as a fixed effect and species as a random intercept to account for species-specific variation. To assess the significance of the marker type, we conducted an ANOVA on the mixed model using the anova() function from the car package version 3.1-3 (Fox & Weisberg, 2019).

Post-hoc pairwise comparisons were conducted to evaluate differences in supported nodes across marker types. We used the emmeans package version 1.10.6 to obtain estimated marginal means and performed pairwise comparisons with Bonferroni correction for multiple testing.

### Heterozygosity

Global heterozygosity for each dataset was calculated using ANGSD version 0.935 based on high-quality sites with minimum mapping and base quality scores of 30 and 20, respectively (Korneliussen et al., 2014). We estimated the maximum likelihood site frequency spectrum using the realSFS function and then calculated heterozygosity as the proportion of heterozygous sites out of the total sites for each individual. A bash script was used to output these values in table format for easy comparison across individuals.

### Clustering for population-level analyses: Principal Component Analysis & STRUCTURE

To infer population clusters we performed Principal Component Analyses (PCA) on our SNP data using the R package SNPRelate version 1.22.0 (Zheng et al., 2012). We calculated the proportion of variance explained by each principal component and used an elbow plot of within-cluster sum of squares. We tested values from 1 to 10 for the number of clusters based on conservative assumptions, aiming to ensure that the clustering solution would not overfit the data while still capturing meaningful patterns. Based on this analysis, we applied *k*-means clustering on the first two principal components to assign individuals to the optimal number of clusters for subsequent pairwise comparisons across datasets. We visualized the first two principal components with 95% confidence ellipses generated for each dataset using ggplot2 version 3.5.1 (Wickham, 2016). with each sample colored based on the clustering assignments from the genome-wide SNPs. To compare the variance percentages explained by different datasets across species, we fitted a linear mixed-effects model with dataset and species as fixed effects, and species as a random intercept using lmerTest version 3.1-3 (Kuznetsova et al., 2017). Pairwise comparisons were then conducted using estimated marginal means from the model. The analysis aimed to assess whether there were significant differences in variance explained by each dataset across species. In tandem with PCA, population structure was analyzed with the R package LEA version 3.10.2 (Frichot & François, 2015). We used the cross-entropy criterion from the *snmf* function to infer the best fit of ancestral populations. We tested a range of *K* values from 1 to 25 and chose the number of populations that minimized cross-entropy.

## Data availability

Additional figures and tables supporting the findings of this study are available in the supplemental materials. Supplemental Table S1 provides a comprehensive list of all study systems, including the SRA accession numbers, reference genomes, original publications, and additional relevant information. Scripts used in this study can be accessed on GitHub at XXXX

## Acknowledgements

This project developed during a semester-long discussion group on phylogenomics at Cornell University in spring 2023 led by J.B.L. We would like to thank additional participants in the discussion group, including Sylvana Ross, Annette Kang, and Lidane Audrey Noronha for their invaluable comments throughout our dry lab and discussion sessions. Special thanks to Cornell University College of Agriculture and Life Sciences (CALS) IT department and the Cornell BioHPC for providing server space during the semester. *Solanum* samples are based upon work supported by the National Science Foundation Graduate Research Fellowship under Grant No. 2139899 to M.H. J.F. was supported by the National Science Foundation Graduate Research Fellowship under Grant No. 2139899

## References

Ai, B., & Kang, M. (2015). How Many Genes are Needed to Resolve Phylogenetic Incongruence? Evolutionary Bioinformatics Online, 11, 185–188. 10.4137/EBO.S26047

Alam, M. N. U., Román-Palacios, C., Copetti, D., & Wing, R. A. (2025). Universal orthologs infer deep phylogenies and improve genome quality assessments (p. 2025.02.17.638702). bioRxiv. 10.1101/2025.02.17.638702

Altschul, S. F., Gish, W., Miller, W., Myers, E. W., & Lipman, D. J. (1990). Basic local alignment search tool. Journal of Molecular Biology, 215(3), 403–410. 10.1016/S0022-2836(05)80360-2

Alves, F., Banks, S. C., Edworthy, M., Stojanovic, D., Langmore, N. E., & Heinsohn, R. (2023). Using conservation genetics to prioritise management options for an endangered songbird. Heredity, 130(5), 289–301. 10.1038/s41437-023-00609-6

Antonelli, A., Clarkson, J. J., Kainulainen, K., Maurin, O., Brewer, G. E., Davis, A. P., Epitawalage, N., Goyder, D. J., Livshultz, T., Persson, C., Pokorny, L., Straub, S. C. K., Struwe, L., Zuntini, A. R., Forest, F., & Baker, W. J. (2021). Settling a family feud: A high-level phylogenomic framework for the Gentianales based on 353 nuclear genes and partial plastomes. American Journal of Botany, 108(7), 1143–1165. 10.1002/ajb2.1697

Bagley, J. C., Uribe-Convers, S., Carlsen, M. M., & Muchhala, N. (2020). Utility of targeted sequence capture for phylogenomics in rapid, recent angiosperm radiations: Neotropical *Burmeistera* bellflowers as a case study. Molecular Phylogenetics and Evolution, 152, 106769. 10.1016/j.ympev.2020.106769

Ball, L. D., Bedoya, A. M., Taylor, C. M., & Lagomarsino, L. P. (2023). A target enrichment probe set for resolving phylogenetic relationships in the coffee family, Rubiaceae. Applications in Plant Sciences, 11(6), e11554. 10.1002/aps3.11554

Blischak, P. D., Chifman, J., Wolfe, A. D., & Kubatko, L. S. (2018). HyDe: A Python Package for Genome-Scale Hybridization Detection. Systematic Biology, 67(5), 821–829. 10.1093/sysbio/syy023

Boluda, C. G., Christe, C., Randriarisoa, A., Gautier, L., & Naciri, Y. (2021). Species Delimitation and Conservation in Taxonomically Challenging Lineages: The Case of Two Clades of Capurodendron (Sapotaceae) in Madagascar. Plants, 10(8), Article 8. 10.3390/plants10081702

Borowiec, M. L., Miles Zhang, Y., Neves, K., Ramalho, M. O., Fisher, B. L., Lucky, A., & Moreau, C. S. (2024). Evaluating UCE data adequacy and integrating uncertainty in a comprehensive phylogeny of ants. 10.1101/2024.07.03.601921

Bouckaert, R., Vaughan, T. G., Barido-Sottani, J., Duchêne, S., Fourment, M., Gavryushkina, A., Heled, J., Jones, G., Kühnert, D., Maio, N. D., Matschiner, M., Mendes, F. K., Müller, N. F., Ogilvie, H. A., Plessis, L. du, Popinga, A., Rambaut, A., Rasmussen, D., Siveroni, I., … Drummond, A. J. (2019). BEAST 2.5: An advanced software platform for Bayesian evolutionary analysis. PLOS Computational Biology, 15(4), e1006650. 10.1371/journal.pcbi.1006650

Calus, M. P. L., & Vandenplas, J. (2018). SNPrune: An efficient algorithm to prune large SNP array and sequence datasets based on high linkage disequilibrium. Genetics Selection Evolution, 50(1), 34. 10.1186/s12711-018-0404-z

Catchen, J., Hohenlohe, P. A., Bassham, S., Amores, A., & Cresko, W. A. (2013). Stacks: An analysis tool set for population genomics. Molecular Ecology, 22(11), 3124–3140. 10.1111/mec.12354

Chang, C. C., Chow, C. C., Tellier, L. C., Vattikuti, S., Purcell, S. M., & Lee, J. J. (2015). Second-generation PLINK: rising to the challenge of larger and richer datasets. GigaScience, 4(1), s13742–015-0047–0048. 10.1186/s13742-015-0047-8

Chen, S., Zhou, Y., Chen, Y., & Gu, J. (2018). fastp: An ultra-fast all-in-one FASTQ preprocessor. Bioinformatics, 34(17), i884–i890. 10.1093/bioinformatics/bty560

Chifman, J., & Kubatko, L. (2014). Quartet Inference from SNP Data Under the Coalescent Model. Bioinformatics, 30(23), 3317–3324. 10.1093/bioinformatics/btu530

Cooper, B. J., Moore, M. J., Douglas, N. A., Wagner, W. L., Johnson, M. G., Overson, R. P., Kinosian, S. P., McDonnell, A. J., Levin, R. A., Raguso, R. A., Flores Olvera, H., Ochoterena, H., Fant, J. B., Skogen, K. A., & Wickett, N. J. (2023). Target Enrichment and Extensive Population Sampling Help Untangle the Recent, Rapid Radiation of Oenothera Sect. Calylophus. Systematic Biology, 72(2), 249–263. 10.1093/sysbio/syac032

Cullingham, C. I., Miller, J. M., Peery, R. M., Dupuis, J. R., Malenfant, R. M., Gorrell, J. C., & Janes, J. K. (2020). Confidently identifying the correct K value using the ΔK method: When does K = 2? Molecular Ecology, 29(5), 862–869. 10.1111/mec.15374

Danecek, P., Bonfield, J. K., Liddle, J., Marshall, J., Ohan, V., Pollard, M. O., Whitwham, A., Keane, T., McCarthy, S. A., Davies, R. M., & Li, H. (2021). Twelve years of SAMtools and BCFtools. GigaScience, 10(2), giab008. 10.1093/gigascience/giab008

de Jong, M. J., Niamir, A., Wolf, M., Kitchener, A. C., Lecomte, N., Seryodkin, I. V., Fain, S. R., Hagen, S. B., Saarma, U., & Janke, A. (2023). Range-wide whole-genome resequencing of the brown bear reveals drivers of intraspecies divergence. Communications Biology, 6(1), 1–16. 10.1038/s42003-023-04514-w

Devault, A., & Enk, J. (2023). A universal targeted sequencing system for any high-throughput sequencing platform.

Dodsworth, S., Pokorny, L., Johnson, M. G., Kim, J. T., Maurin, O., Wickett, N. J., Forest, F., & Baker, W. J. (2019). Hyb-Seq for Flowering Plant Systematics. Trends in Plant Science, 24(10), 887–891. 10.1016/j.tplants.2019.07.011

Duckett, D. J., Calder, K., Sullivan, J., Tank, D. C., & Carstens, B. C. (2023). Reduced representation approaches produce similar results to whole genome sequencing for some common phylogeographic analyses. PLOS ONE, 18(11), e0291941. 10.1371/journal.pone.0291941

Durand, E. Y., Patterson, N., Reich, D., & Slatkin, M. (2011). Testing for Ancient Admixture between Closely Related Populations. Molecular Biology and Evolution, 28(8), 2239– 2252. 10.1093/molbev/msr048

Edwards, S. V. (2009). Natural selection and phylogenetic analysis. Proceedings of the National Academy of Sciences, 106(22), 8799–8800. 10.1073/pnas.0904103106

Emms, D. M., & Kelly, S. (2019). OrthoFinder: Phylogenetic orthology inference for comparative genomics. Genome Biology, 20(1), 238. 10.1186/s13059-019-1832-y

Faircloth, B. C., Branstetter, M. G., White, N. D., & Brady, S. G. (2015). Target enrichment of ultraconserved elements from arthropods provides a genomic perspective on relationships among Hymenoptera. Molecular Ecology Resources, 15(3), 489–501. 10.1111/1755-0998.12328

Faircloth, B. C., McCormack, J. E., Crawford, N. G., Harvey, M. G., Brumfield, R. T., & Glenn, T. C. (2012). Ultraconserved Elements Anchor Thousands of Genetic Markers Spanning Multiple Evolutionary Timescales. Systematic Biology, 61(5), 717–726. 10.1093/sysbio/sys004

Felsenstein, J. (1985). Confidence limits on phylogenies: An approach using the bootstrap. Evolution, 39(4), 783–791.

Fong, J. J., & Fujita, M. K. (2011). Evaluating phylogenetic informativeness and data-type usage for new protein-coding genes across Vertebrata. Molecular Phylogenetics and Evolution, 61(2), 300–307. 10.1016/j.ympev.2011.06.016

Fox, J., & Weisberg, S. (2019). An R Companion to Applied Regression (Third). Sage. https://www.john-fox.ca/Companion/

Frichot, E., & François, O. (2015). LEA: An R package for landscape and ecological association studies. Methods in Ecology and Evolution, 6(8), 925–929. 10.1111/2041-210X.12382

Galtier, N. (2004). Sampling Properties of the Bootstrap Support in Molecular Phylogeny: Influence of Nonindependence Among Sites. Systematic Biology, 53(1), 38–46. 10.1080/10635150490264680

Gates, D. J., Pilson, D., & Smith, S. D. (2021). Inferring the history of hybridization: A case study in Iochrominae (Solanaceae). 10.32942/osf.io/jhev4

Giarla, T. C., & Esselstyn, J. A. (2015). The Challenges of Resolving a Rapid, Recent Radiation: Empirical and Simulated Phylogenomics of Philippine Shrews. Systematic Biology, 64(5), 727–740. 10.1093/sysbio/syv029

Grabherr, M. G., Haas, B. J., Yassour, M., Levin, J. Z., Thompson, D. A., Amit, I., Adiconis, X., Fan, L., Raychowdhury, R., Zeng, Q., Chen, Z., Mauceli, E., Hacohen, N., Gnirke, A., Rhind, N., di Palma, F., Birren, B. W., Nusbaum, C., Lindblad-Toh, K., … Regev, A. (2011). Full-length transcriptome assembly from RNA-Seq data without a reference genome. Nature Biotechnology, 29(7), 644–652. 10.1038/nbt.1883

Graham, C. F., Boreham, D. R., Manzon, R. G., Stott, W., Wilson, J. Y., & Somers, C. M. (2020). How “simple” methodological decisions affect interpretation of population structure based on reduced representation library DNA sequencing: A case study using the lake whitefish. PLOS ONE, 15(1), e0226608. 10.1371/journal.pone.0226608

Green, R. E., Krause, J., Briggs, A. W., Maricic, T., Stenzel, U., Kircher, M., Patterson, N., Li, H., Zhai, W., Fritz, M. H.-Y., Hansen, N. F., Durand, E. Y., Malaspinas, A.-S., Jensen, J. D., Marques-Bonet, T., Alkan, C., Prüfer, K., Meyer, M., Burbano, H. A., … Pääbo, S. (2010). A Draft Sequence of the Neandertal Genome. Science, 328(5979), 710–722. 10.1126/science.1188021

Haasl, R. J., & Payseur, B. A. (2011). Multi-locus inference of population structure: A comparison between single nucleotide polymorphisms and microsatellites. Heredity, 106(1), 158–171. 10.1038/hdy.2010.21

Hahn, M. W., & Hibbins, M. S. (2019). A Three-Sample Test for Introgression. Molecular Biology and Evolution, 36(12), 2878–2882. 10.1093/molbev/msz178

Hatami, E., Jones, K. E., & Kilian, N. (2022). New Insights Into the Relationships Within Subtribe Scorzonerinae (Cichorieae, Asteraceae) Using Hybrid Capture Phylogenomics (Hyb-Seq). Frontiers in Plant Science, 13. 10.3389/fpls.2022.851716

Hou, Z., Li, A., & Zhang, J. (2020). Genetic architecture, demographic history, and genomic differentiation of Populus davidiana revealed by whole-genome resequencing. Evolutionary Applications, 13(10), 2582–2596. 10.1111/eva.13046

Hughes, C. E., Eastwood, R. J., & Donovan Bailey, C. (2006). From famine to feast? Selecting nuclear DNA sequence loci for plant species-level phylogeny reconstruction. Philosophical Transactions of the Royal Society B: Biological Sciences, 361(1465), 211–225. 10.1098/rstb.2005.1735

Jin, J.-J., Yu, W.-B., Yang, J.-B., Song, Y., dePamphilis, C. W., Yi, T.-S., & Li, D.-Z. (2020). GetOrganelle: A fast and versatile toolkit for accurate de novo assembly of organelle genomes. Genome Biology, 21(1), 241. 10.1186/s13059-020-02154-5

Johnson, M. G., Gardner, E. M., Liu, Y., Medina, R., Goffinet, B., Shaw, A. J., Zerega, N. J. C., & Wickett, N. J. (2016). HybPiper: Extracting coding sequence and introns for phylogenetics from high-throughput sequencing reads using target enrichment. Applications in Plant Sciences, 4(7), 1600016. 10.3732/apps.1600016

Johnson, M. G., Pokorny, L., Dodsworth, S., Botigué, L. R., Cowan, R. S., Devault, A., Eiserhardt, W. L., Epitawalage, N., Forest, F., Kim, J. T., Leebens-Mack, J. H., Leitch, I. J., Maurin, O., Soltis, D. E., Soltis, P. S., Wong, G. K., Baker, W. J., & Wickett, N. J. (2019). A Universal Probe Set for Targeted Sequencing of 353 Nuclear Genes from Any Flowering Plant Designed Using k-Medoids Clustering. Systematic Biology, 68(4), 594– 606. 10.1093/sysbio/syy086

Kardos, M., & Waples, R. S. (2024). Low-coverage sequencing and Wahlund effect severely bias estimates of inbreeding, heterozygosity and effective population size in North American wolves. Molecular Ecology, n/a(n/a), e17415. 10.1111/mec.17415

Korneliussen, T. S., Albrechtsen, A., & Nielsen, R. (2014). ANGSD: Analysis of Next Generation Sequencing Data. BMC Bioinformatics, 15(1), 356. 10.1186/s12859-014-0356-4

Kozlov, A. M., Darriba, D., Flouri, T., Morel, B., & Stamatakis, A. (2019). RAxML-NG: A fast, scalable and user-friendly tool for maximum likelihood phylogenetic inference. Bioinformatics, 35(21), 4453–4455. 10.1093/bioinformatics/btz305

Kuznetsova, A., Brockhoff, P. B., & Christensen, R. H. B. (2017). lmerTest Package: Tests in Linear Mixed Effects Models. Journal of Statistical Software, 82(13), 1–26. 10.18637/jss.v082.i13

Lamichhaney, S., Berglund, J., Almén, M. S., Maqbool, K., Grabherr, M., Martinez-Barrio, A., Promerová, M., Rubin, C.-J., Wang, C., Zamani, N., Grant, B. R., Grant, P. R., Webster, M. T., & Andersson, L. (2015). Evolution of Darwin’s finches and their beaks revealed by genome sequencing. Nature, 518(7539), 371–375. 10.1038/nature14181

Lee, A. K., Gilman, I. S., Srivastav, M., Lerner, A. D., Donoghue, M. J., & Clement, W. L. (2021). Reconstructing Dipsacales phylogeny using Angiosperms353: Issues and insights. American Journal of Botany, 108(7), 1122–1142. 10.1002/ajb2.1695

Lefouili, M., & Nam, K. (2022). The evaluation of Bcftools mpileup and GATK HaplotypeCaller for variant calling in non-human species. Scientific Reports, 12(1), 11331. 10.1038/s41598-022-15563-2

Leigh, D. M., Lischer, H. E. L., Grossen, C., & Keller, L. F. (2018). Batch effects in a multiyear sequencing study: False biological trends due to changes in read lengths. Molecular Ecology Resources, 18(4), 778–788. 10.1111/1755-0998.12779

Lenth, R. V. (2024). emmeans: Estimated Marginal Means, aka Least-Squares Means. https://rvlenth.github.io/emmeans/

Leroy, G., Carroll, E. L., Bruford, M. W., DeWoody, J. A., Strand, A., Waits, L., & Wang, J. (2018). Next-generation metrics for monitoring genetic erosion within populations of conservation concern. Evolutionary Applications, 11(7), 1066–1083. 10.1111/eva.12564

Lou, R. N., & Therkildsen, N. O. (2022). Batch effects in population genomic studies with low-coverage whole genome sequencing data: Causes, detection and mitigation. Molecular Ecology Resources, 22(5), 1678–1692. 10.1111/1755-0998.13559

Manni, M., Berkeley, M. R., Seppey, M., Simão, F. A., & Zdobnov, E. M. (2021). BUSCO Update: Novel and Streamlined Workflows along with Broader and Deeper Phylogenetic Coverage for Scoring of Eukaryotic, Prokaryotic, and Viral Genomes. Molecular Biology and Evolution, 38(10), 4647–4654. 10.1093/molbev/msab199

McLay, T. G. B., Birch, J. L., Gunn, B. F., Ning, W., Tate, J. A., Nauheimer, L., Joyce, E. M., Simpson, L., Schmidt-Lebuhn, A. N., Baker, W. J., Forest, F., & Jackson, C. J. (2021). New targets acquired: Improving locus recovery from the Angiosperms353 probe set. Applications in Plant Sciences, 9(7). 10.1002/aps3.11420

Minh, B. Q., Schmidt, H. A., Chernomor, O., Schrempf, D., Woodhams, M. D., von Haeseler, A., & Lanfear, R. (2020). IQ-TREE 2: New Models and Efficient Methods for Phylogenetic Inference in the Genomic Era. Molecular Biology and Evolution, 37(5), 1530–1534. 10.1093/molbev/msaa015

Mo, Z.-Q., Fu, C.-N., Twyford, A. D., Hollingsworth, P. M., Zhang, T., Yang, J.-B., Li, D.-Z., & Gao, L.-M. (2025). Evaluating the utility of deep genome skimming for phylogenomic analyses: A case study in the species-rich genus Rhododendron. Plant Diversity, S2468265925000836. 10.1016/j.pld.2025.04.006

Nazareno, A. G., Bemmels, J. B., Dick, C. W., & Lohmann, L. G. (2017). Minimum sample sizes for population genomics: An empirical study from an Amazonian plant species. Molecular Ecology Resources, 17(6), 1136–1147. 10.1111/1755-0998.12654

Ogle, D. H., Doll, J. C., Wheeler, A. P., & Dinno, A. (2025). FSA: Simple Fisheries Stock Assessment Methods. 10.32614/CRAN.package.FSA

Ortiz, E. M. (2019). vcf2phylip v2.0: Convert a VCF matrix into several matrix formats for phylogenetic analysis. (Version v2.0) [Computer software]. Zenodo. 10.5281/zenodo.2540861

Paradis, E., & Schliep, K. (2018). ape 5.0: An environment for modern phylogenetics and evolutionary analyses in R. Bioinformatics, 35(3), 526–528. 10.1093/bioinformatics/bty633

Pezzini, F. F., Ferrari, G., Forrest, L. L., Hart, M. L., Nishii, K., & Kidner, C. A. (2023). Target capture and genome skimming for plant diversity studies. Applications in Plant Sciences, 11(4), e11537. 10.1002/aps3.11537

Pyrcz, T. W., Lachowska-Cierlik, D., Willmott, K. R., Mrozek, A., Mahecha-Jiménez, O., Fåhraeus, C., Boyer, P., Martín, S., & Espeland, M. (2023). A new genus in the diverse Andean Pedaliodes complex uncovered using target enrichment (Lepidoptera, Nymphalidae). Systematic Entomology, 48(1), 163–177. 10.1111/syen.12568

R Core Team. (2024). R: A Language and Environment for Statistical Computing. R Foundation for Statistical Computing. https://www.R-project.org/

Revell, L. J. (2024). phytools 2.0: An updated R ecosystem for phylogenetic comparative methods (and other things). PeerJ, 12, e16505. 10.7717/peerj.16505

Roycroft, E. J., Moussalli, A., & Rowe, K. C. (2020). Phylogenomics Uncovers Confidence and Conflict in the Rapid Radiation of Australo-Papuan Rodents. Systematic Biology, 69(3), 431–444. 10.1093/sysbio/syz044

Sandercock, A. M., Westbrook, J. W., Zhang, Q., Johnson, H. A., Saielli, T. M., Scrivani, J. A., Fitzsimmons, S. F., Collins, K., Perkins, M. T., Craddock, J. H., Schmutz, J., Grimwood, J., & Holliday, J. A. (2022). Frozen in time: Rangewide genomic diversity, structure, and demographic history of relict American chestnut populations. Molecular Ecology, 31(18), 4640–4655. 10.1111/mec.16629

Sass, C., Iles, W. J. D., Barrett, C. F., Smith, S. Y., & Specht, C. D. (2016). Revisiting the Zingiberales: Using multiplexed exon capture to resolve ancient and recent phylogenetic splits in a charismatic plant lineage. PeerJ, 4, e1584. 10.7717/peerj.1584

Shah, T., Schneider, J. V., Zizka, G., Maurin, O., Baker, W., Forest, F., Brewer, G. E., Savolainen, V., Darbyshire, I., & Larridon, I. (2021). Joining forces in Ochnaceae phylogenomics: A tale of two targeted sequencing probe kits. American Journal of Botany, 108(7), 1201–1216. 10.1002/ajb2.1682

Shen, X.-X., Hittinger, C. T., & Rokas, A. (2017). Contentious relationships in phylogenomic studies can be driven by a handful of genes. Nature Ecology & Evolution, 1(5), 1–10. 10.1038/s41559-017-0126

Simão, F. A., Waterhouse, R. M., Ioannidis, P., Kriventseva, E. V., & Zdobnov, E. M. (2015). BUSCO: Assessing genome assembly and annotation completeness with single-copy orthologs. Bioinformatics, 31(19), 3210–3212. 10.1093/bioinformatics/btv351

Slatkin, M. (2008). Linkage disequilibrium—Understanding the evolutionary past and mapping the medical future. Nature Reviews Genetics, 9(6), 477–485. 10.1038/nrg2361

Slimp, M., Williams, L. D., Hale, H., & Johnson, M. G. (2021). On the potential of Angiosperms353 for population genomic studies. Applications in Plant Sciences, 9(7), 10.1002/aps3.11419. 10.1002/aps3.11419

Song, B., Buckler, E. S., & Stitzer, M. C. (2023). New whole-genome alignment tools are needed for tapping into plant diversity. *Trends in Plant Science*, S1360138523002753. 10.1016/j.tplants.2023.08.013

Song, B., Ning, W., Wei, D., Jiang, M., Zhu, K., Wang, X., Edwards, D., Odeny, D. A., & Cheng, S. (2023). Plant genome resequencing and population genomics: Current status and future prospects. Molecular Plant, 16(8), 1252–1268. 10.1016/j.molp.2023.07.009

Straub, S. C. K., Parks, M., Weitemier, K., Fishbein, M., Cronn, R. C., & Liston, A. (2012). Navigating the tip of the genomic iceberg: Next-generation sequencing for plant systematics. American Journal of Botany, 99(2), 349–364. 10.3732/ajb.1100335

Stubbs, R. L., Theodoridis, S., Mora-Carrera, E., Keller, B., Yousefi, N., Potente, G., Léveillé-Bourret, É., Celep, F., Kochjarová, J., Tedoradze, G., Eaton, D. A. R., & Conti, E. (2023). Whole-genome analyses disentangle reticulate evolution of primroses in a biodiversity hotspot. New Phytologist, 237(2), 656–671. 10.1111/nph.18525

Suissa, J. S., De La Cerda, G. Y., Graber, L. C., Jelley, C., Wickell, D., Phillips, H. R., Grinage, A. D., Moreau, C. S., Specht, C. D., Doyle, J. J., & Landis, J. B. (2023). Comparative phylogenomic analyses of SNP versus full locus datasets: Insights and recommendations for researchers. 10.1101/2023.09.02.556036

Szarmach, S. J., Brelsford, A., Witt, C. C., & Toews, D. P. L. (2021). Comparing divergence landscapes from reduced-representation and whole genome resequencing in the yellow-rumped warbler (Setophaga coronata) species complex. Molecular Ecology, 30(23), 5994–6005. 10.1111/mec.15940

Thomas, A. E., Igea, J., Meudt, H. M., Albach, D. C., Lee, W. G., & Tanentzap, A. J. (2021). Using target sequence capture to improve the phylogenetic resolution of a rapid radiation in New Zealand Veronica. American Journal of Botany, 108(7), 1289–1306. 10.1002/ajb2.1678

Valderrama, E., Landis, J. B., Skinner, D., Maas, P. J. M., Maas-van de Kramer, H., André, T., Grunder, N., Sass, C., Pinilla-Vargas, M., Guan, C. J., Phillips, H. R., Almeida, A. M. R. de, & Specht, C. D. (2022). The genetic mechanisms underlying the convergent evolution of pollination syndromes in the Neotropical radiation of Costus L. Frontiers in Plant Science, 13. https://www.frontiersin.org/articles/10.3389/fpls.2022.874322

Valderrama, E., Sass, C., Pinilla-Vargas, M., Skinner, D., Maas, P. J. M., Maas-van de Kamer, H., Landis, J. B., Guan, C. J., & Specht, C. D. (2020). Unraveling the Spiraling Radiation: A Phylogenomic Analysis of Neotropical Costus L. Frontiers in Plant Science, 11. 10.3389/fpls.2020.01195

Van der Auwera, G. A., & O’Connor, B. D. (2020). Genomics in the cloud: Using Docker, GATK, and WDL in Terra. O’Reilly Media.

Vasimuddin, Md., Misra, S., Li, H., & Aluru, S. (2019). Efficient Architecture-Aware Acceleration of BWA-MEM for Multicore Systems. 2019 IEEE International Parallel and Distributed Processing Symposium (IPDPS), 314–324. 10.1109/IPDPS.2019.00041

Vekemans, X., Castric, V., Hipperson, H., Müller, N. A., Westerdahl, H., & Cronk, Q. (2021). Whole-genome sequencing and genome regions of special interest: Lessons from major histocompatibility complex, sex determination, and plant self-incompatibility. Molecular Ecology, 30(23), 6072–6086. 10.1111/mec.16020

Vera-Paz, S. I., Díaz Contreras Díaz, D. D., Jost, M., Wanke, S., Rossado, A. J., Hernández-Gutiérrez, R., Salazar, G. A., Magallón, S., Gouda, E. J., Ramírez-Morillo, I. M., Donadío, S., & Granados Mendoza, C. (2022). New plastome structural rearrangements discovered in core Tillandsioideae (Bromeliaceae) support recently adopted taxonomy. Frontiers in Plant Science, 13. 10.3389/fpls.2022.924922

Villaverde, T., Pokorny, L., Olsson, S., Rincón-Barrado, M., Johnson, M. G., Gardner, E. M., Wickett, N. J., Molero, J., Riina, R., & Sanmartín, I. (2018). Bridging the micro- and macroevolutionary levels in phylogenomics: Hyb-Seq solves relationships from populations to species and above. New Phytologist, 220(2), 636–650. 10.1111/nph.15312

Wang, D., Li, Y., Li, M., Yang, W., Ma, X., Zhang, L., Wang, Y., Feng, Y., Zhang, Y., Zhou, R., Sanderson, B. J., Keefover-Ring, K., Yin, T., Smart, L. B., DiFazio, S. P., Liu, J., Olson, M., & Ma, T. (2022). Repeated turnovers keep sex chromosomes young in willows. Genome Biology, 23(1), 200. 10.1186/s13059-022-02769-w

Weitemier, K., Straub, S. C. K., Cronn, R. C., Fishbein, M., Schmickl, R., McDonnell, A., & Liston, A. (2014). Hyb-Seq: Combining target enrichment and genome skimming for plant phylogenomics. Applications in Plant Sciences, 2(9), 1400042. 10.3732/apps.1400042

White, D. M., Huang, J.-P., Jara-Muñoz, O. A., MadriñáN, S., Ree, R. H., & Mason-Gamer, R. J. (2021). The Origins of Coca: Museum Genomics Reveals Multiple Independent Domestications from Progenitor Erythroxylum gracilipes. Systematic Biology, 70(1), 1–13. 10.1093/sysbio/syaa074

Wickham, H. (with Sievert, C.). (2016). ggplot2: Elegant graphics for data analysis (Second edition). Springer international publishing.

Xu, B., Liao, M., Deng, H., Yan, C., Lv, Y., Gao, Y., Ju, W., Zhang, J., Jiang, L., Li, X., & Gao, X. (2022). Chromosome-level de novo genome assembly and whole-genome resequencing of the threatened species Acanthochlamys bracteata (Velloziaceae) provide insights into alpine plant divergence in a biodiversity hotspot. Molecular Ecology Resources, 22(4), 1582–1595. 10.1111/1755-0998.13562

Yardeni, G., Viruel, J., Paris, M., Hess, J., Groot Crego, C., de La Harpe, M., Rivera, N., Barfuss, M. H. J., Till, W., Guzmán-Jacob, V., Krömer, T., Lexer, C., Paun, O., & Leroy, T. (2022). Taxon-specific or universal? Using target capture to study the evolutionary history of rapid radiations. Molecular Ecology Resources, 22(3), 927–945. 10.1111/1755-0998.13523

Yoshida, K., Miyagi, R., Mori, S., Takahashi, A., Makino, T., Toyoda, A., Fujiyama, A., & Kitano, J. (2016). Whole-genome sequencing reveals small genomic regions of introgression in an introduced crater lake population of threespine stickleback. Ecology and Evolution, 6(7), 2190–2204. 10.1002/ece3.2047

Zhang, C., Dong, S.-S., Xu, J.-Y., He, W.-M., & Yang, T.-L. (2019). PopLDdecay: A fast and effective tool for linkage disequilibrium decay analysis based on variant call format files. Bioinformatics, 35(10), 1786–1788. 10.1093/bioinformatics/bty875

Zhang, C., Scornavacca, C., Molloy, E. K., & Mirarab, S. (2020). ASTRAL-Pro: Quartet-Based Species-Tree Inference despite Paralogy. Molecular Biology and Evolution, 37(11), 3292–3307. 10.1093/molbev/msaa139

Zheng, X., Levine, D., Shen, J., Gogarten, S. M., Laurie, C., & Weir, B. S. (2012). A high-performance computing toolset for relatedness and principal component analysis of SNP data. Bioinformatics, 28(24), 3326–3328. 10.1093/bioinformatics/bts606

Zhou, Y., Massonnet, M., Sanjak, J. S., Cantu, D., & Gaut, B. S. (2017). Evolutionary genomics of grape (Vitis vinifera ssp. Vinifera) domestication. Proceedings of the National Academy of Sciences, 114(44), 11715–11720. 10.1073/pnas.1709257114

Zuntini, A. R., Carruthers, T., Maurin, O., Bailey, P. C., Leempoel, K., Brewer, G. E., Epitawalage, N., Françoso, E., Gallego-Paramo, B., McGinnie, C., Negrão, R., Roy, S. R., Simpson, L., Toledo Romero, E., Barber, V. M. A., Botigué, L., Clarkson, J. J., Cowan, R. S., Dodsworth, S., … Baker, W. J. (2024). Phylogenomics and the rise of the angiosperms. Nature, 629(8013), 843–850. 10.1038/s41586-024-07324-0

