## Supplemental Material captions for "The power to resolve relationships: identifying incongruence and precision of reduced representation and genome-wide data in phylogenomics and population genomics"

### Tables

**Table S1. Summary of taxa included in the comparative genomic analysis.** For each taxon, we provide the number of samples, genome size, reference genome used, UCE set, BUSCO library, LD pruning threshold ($r^2$), divergence time (in millions of years ago, MYA), species used to identify single-copy orthologs, reproductive mode (outcrossing vs. selfing), life history (annual vs. perennial), and the original publication from which the data were sourced.

**Table S2: Results of Dunn's post hoc tests comparing SNP retention across marker types.** Comparisons are based on pairwise Z-scores, unadjusted p-values (P.unadj), and Bonferri corrected p-values (P.adj).

**Table S3: Pairwise Wilcoxon rank-sum tests comparing heterozygosity across dataset types.** Comparisons of dataset types, test statistics, raw p-values, and bonferri corrected p-values.

### Figures

**S1: Comparison of SNP retention between GATK HaplotypeCaller and BCFtools mpileup following strict filtering.** While GATK initially identified more SNPs, BCFtools retained more SNPs after applying equivalent filtering thresholds. Filtering included retaining only biallelic SNPs with a minor allele count (MAC) $\geq 3$, minor allele frequency (MAF) $\geq 0.05$, depth between 5 and 200, and no more than 30% missing data per site. Individuals with >50% missing data were excluded. The number of SNPs retained, including shared and unique variants for each caller, is shown.

**S2: Co-phylo plots from 5 most common topologies from downsampled replicates in *Populus and Costus*.** Pruning the genome-wide SNP dataset to match the number of SNPs in the UCE + supercontigs dataset led to topological incongruence in both taxonomic datasets.

**S3: Maximum likelihood phylogenies for each taxonomic and marker dataset.** Each panel (e.g., S5a, S5b) shows the best ML tree inferred from one of the seven datasets (UCE, UCE + supercontigs, BUSCO, BUSCO + supercontigs, single-copy, single-copy + supercontigs, and genome-wide SNPs). Trees were inferred using RAxML-NG with 100 bootstrap replicates. Tip and branch colors reflect individual cluster assignments from principal component analysis.

**S4: Estimated marginal means of the percentage of variance explained by principal components 1–4 across dataset types.** Error bars represent 95% confidence intervals.

**S5: Principal component analysis (PCA) plots across all taxon–dataset combinations.** Plots display PC1 and PC2 from PCA of SNP datasets representing different marker types for each taxonomic group. Individual samples are colored by their genome-wide cluster assignment, derived from k-means clustering based on the genome-wide dataset. Local clustering was applied to each dataset independently to generate 95% confidence ellipses, shown as dashed lines.

**S6: Best K value determination for each dataset based on sparse non-negative matrix factorization (sNMF).** Each panel (e.g., S4a, S4b) corresponds to a different taxon. Cross-entropy values were calculated across a range of clusters (K = 1 to 25). Lower cross-entropy values indicate better model fit. The best K (red dot) was selected based on the point at which cross-entropy stabilized or reached a minimum.

**S7: STRUCTURE plots for all taxon–dataset combinations.** Barplots represent individual ancestry coefficients (Q-values) inferred from STRUCTURE-like analyses across marker types for each taxonomic group. The number of ancestral clusters (K) contributing to the current genetic material was fixed at the genome-wide optimum for each taxonomic group.
