## Supplementary figures and images for "The power to resolve relationships: identifying incongruence and precision of reduced representation and genome-wide data in phylogenomics and population genomics"

### Supplemental Figure 1

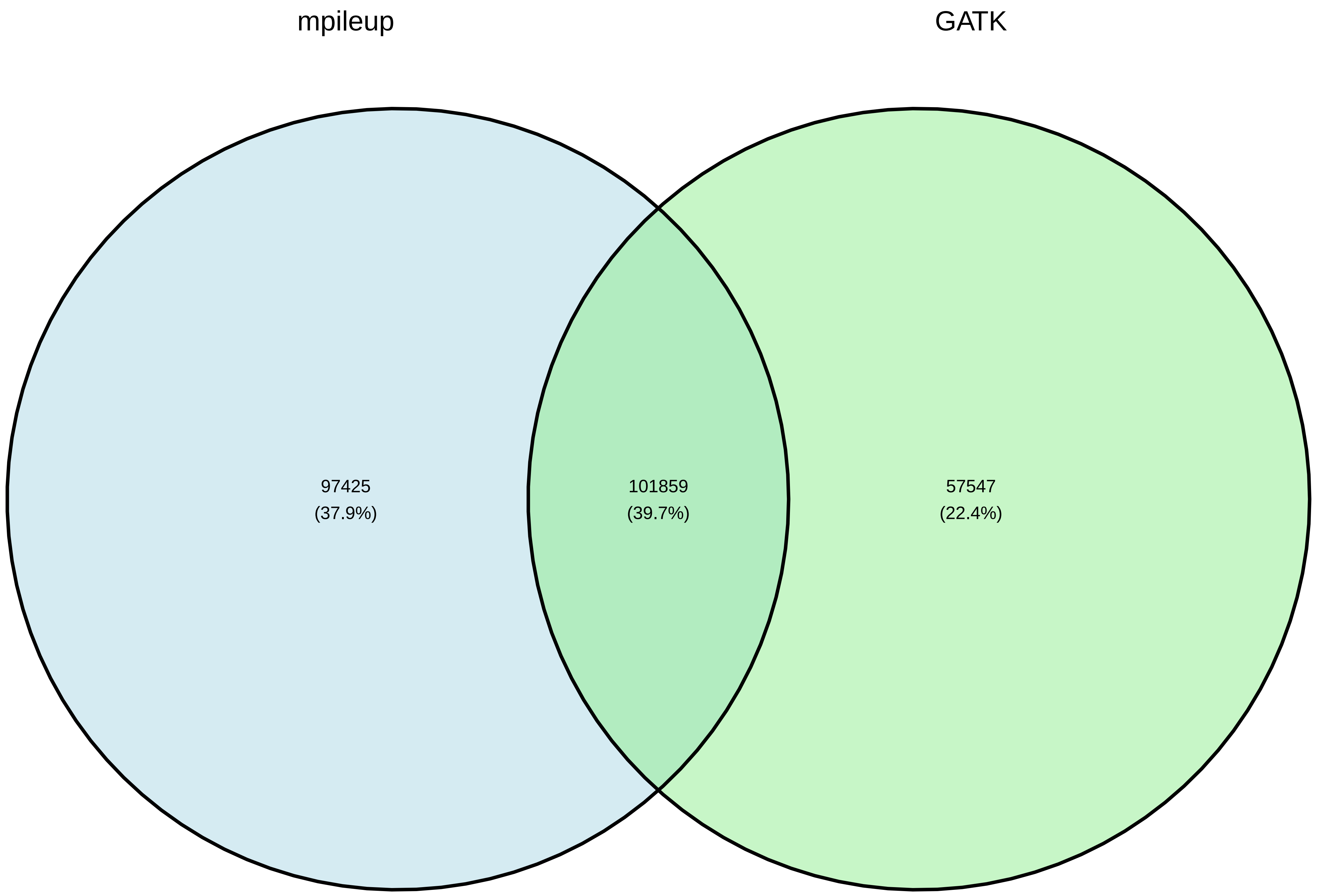

### Supplemental Figure 2

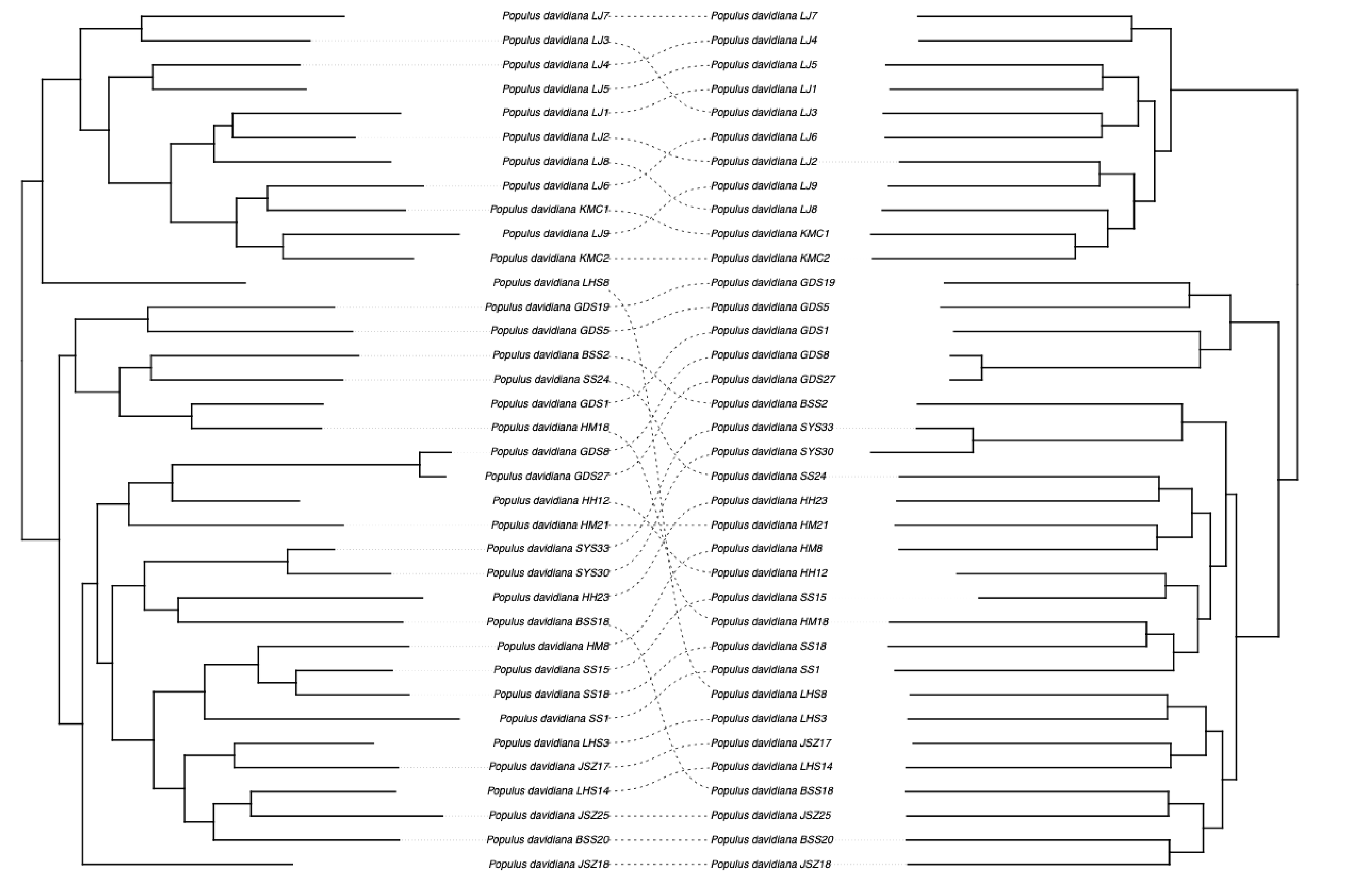

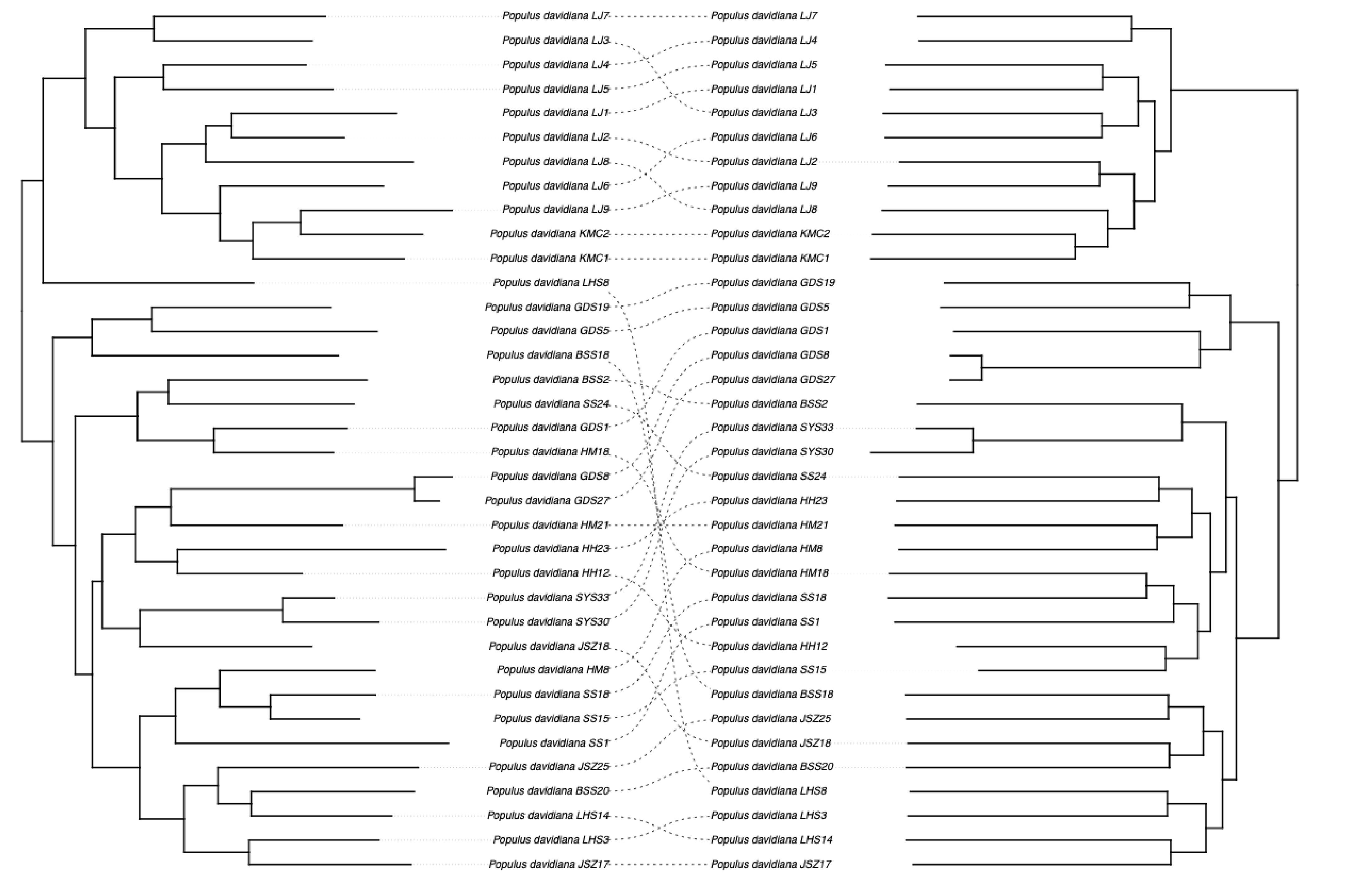

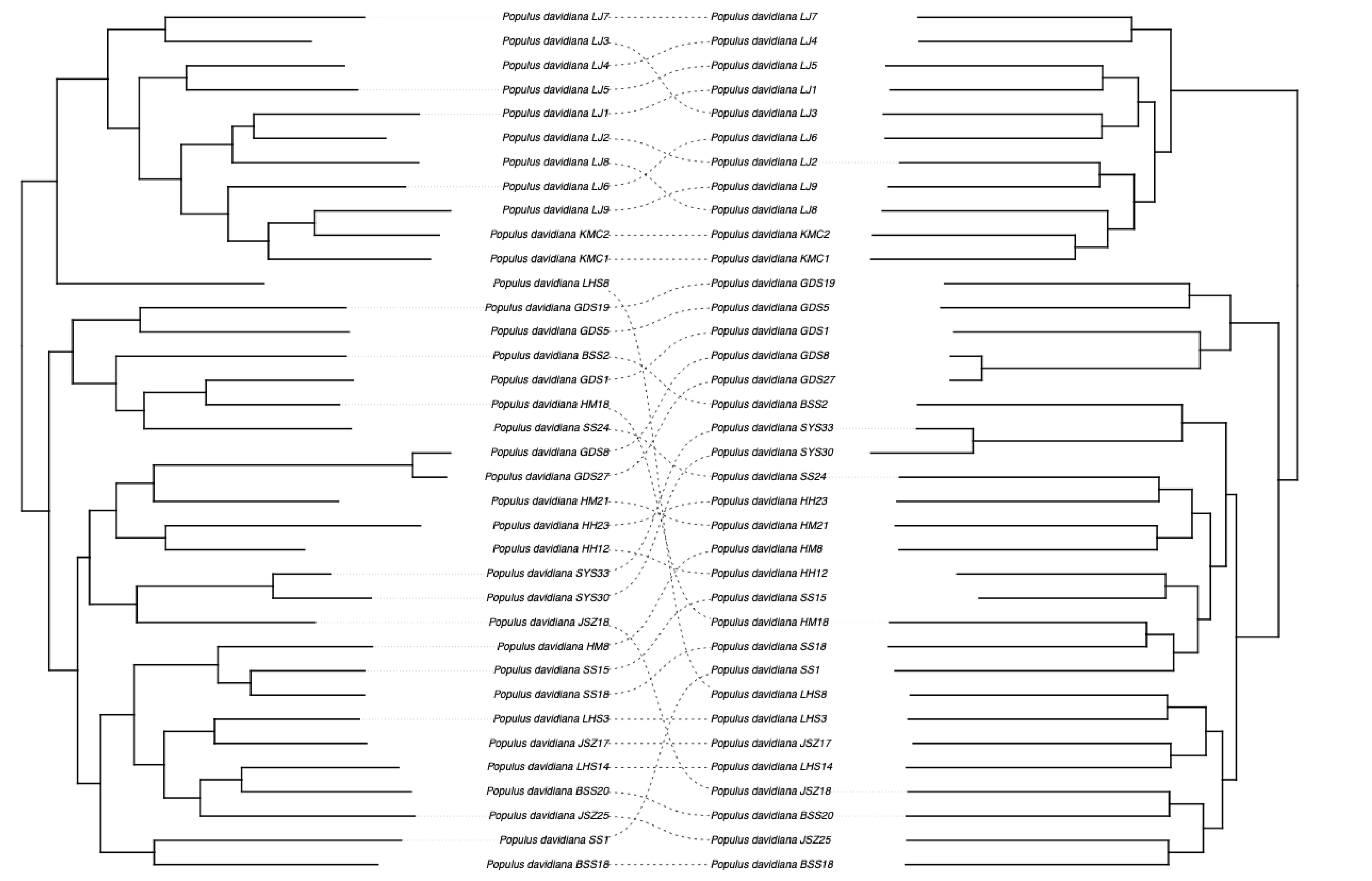

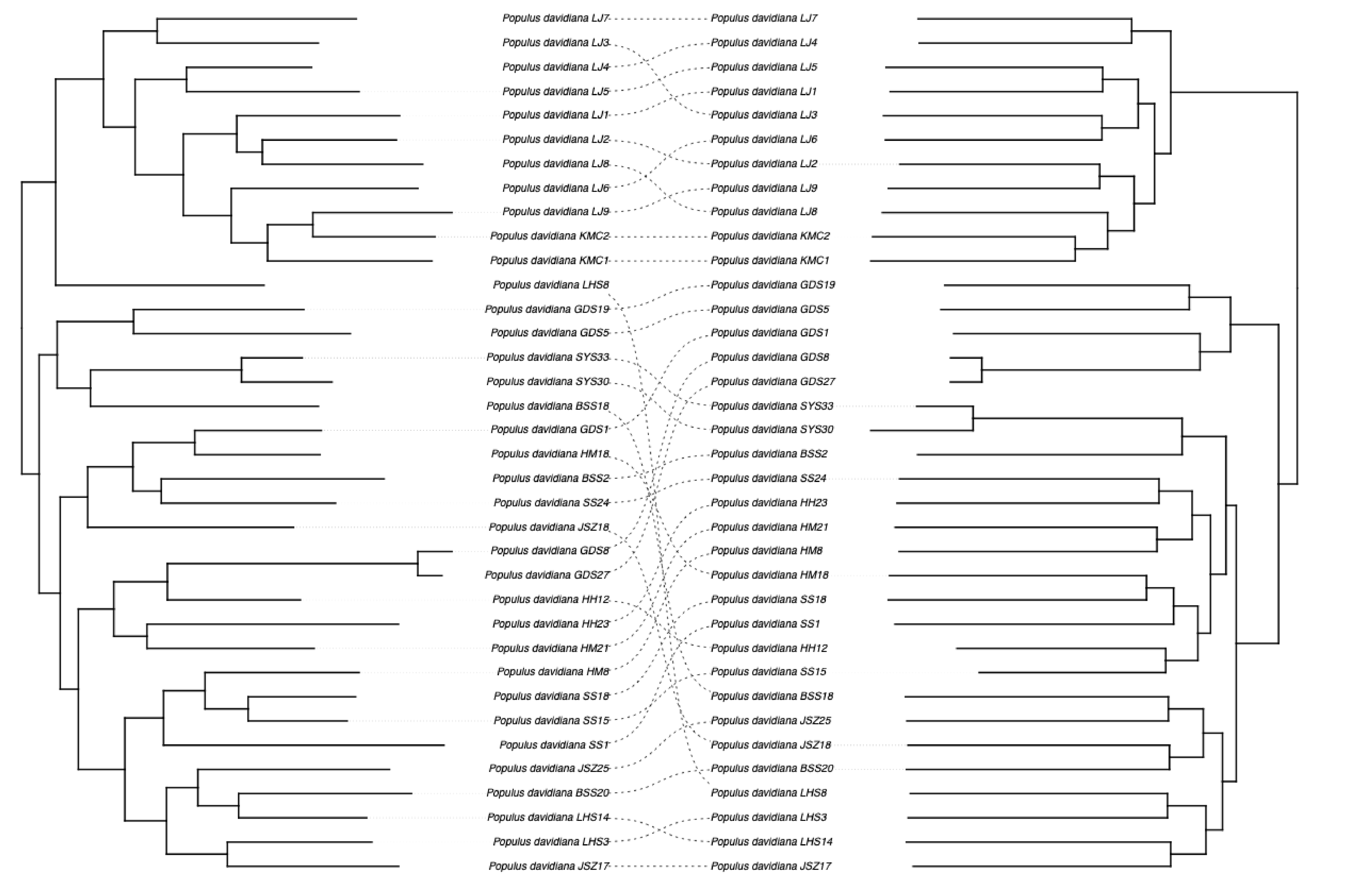

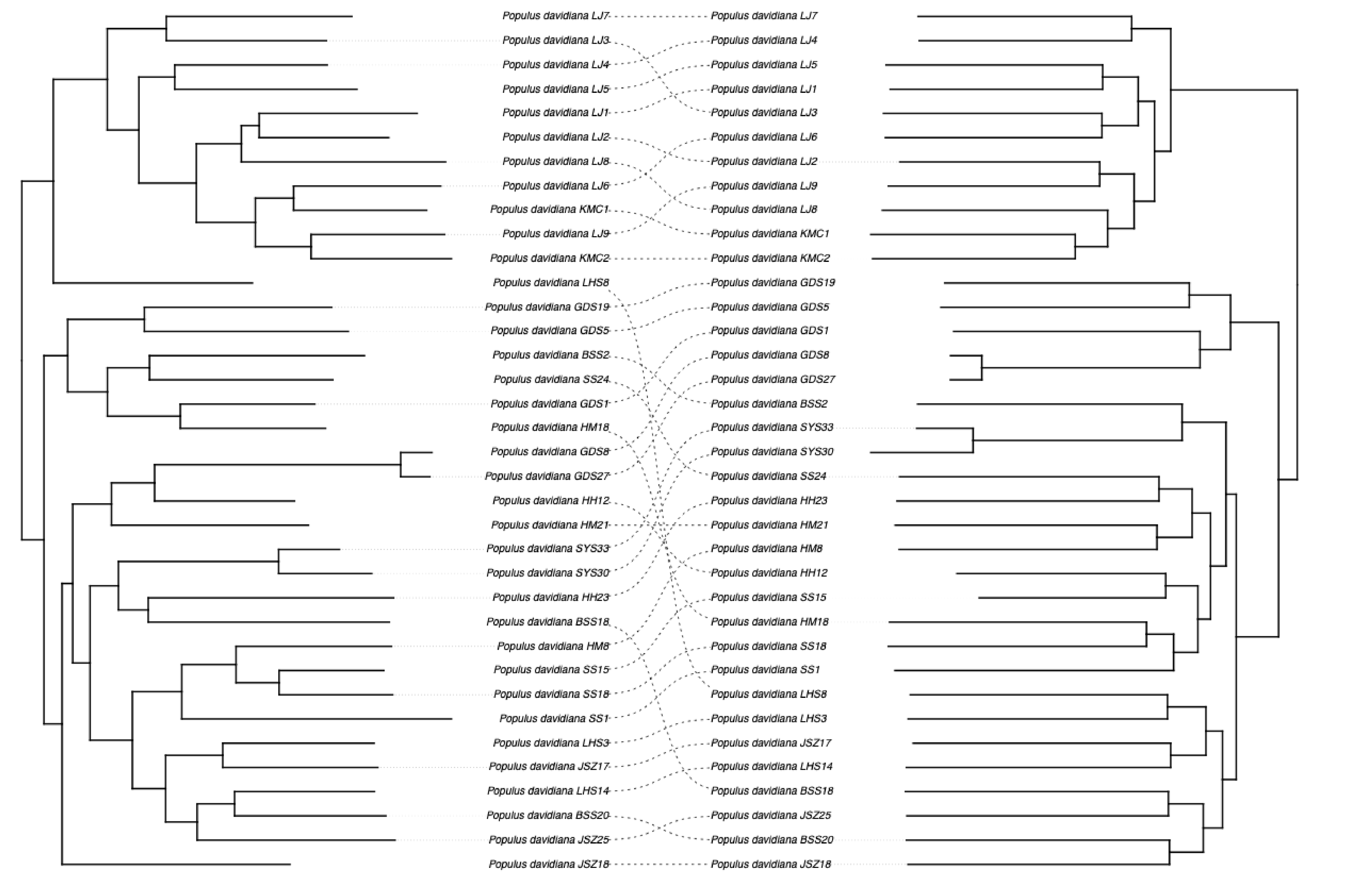

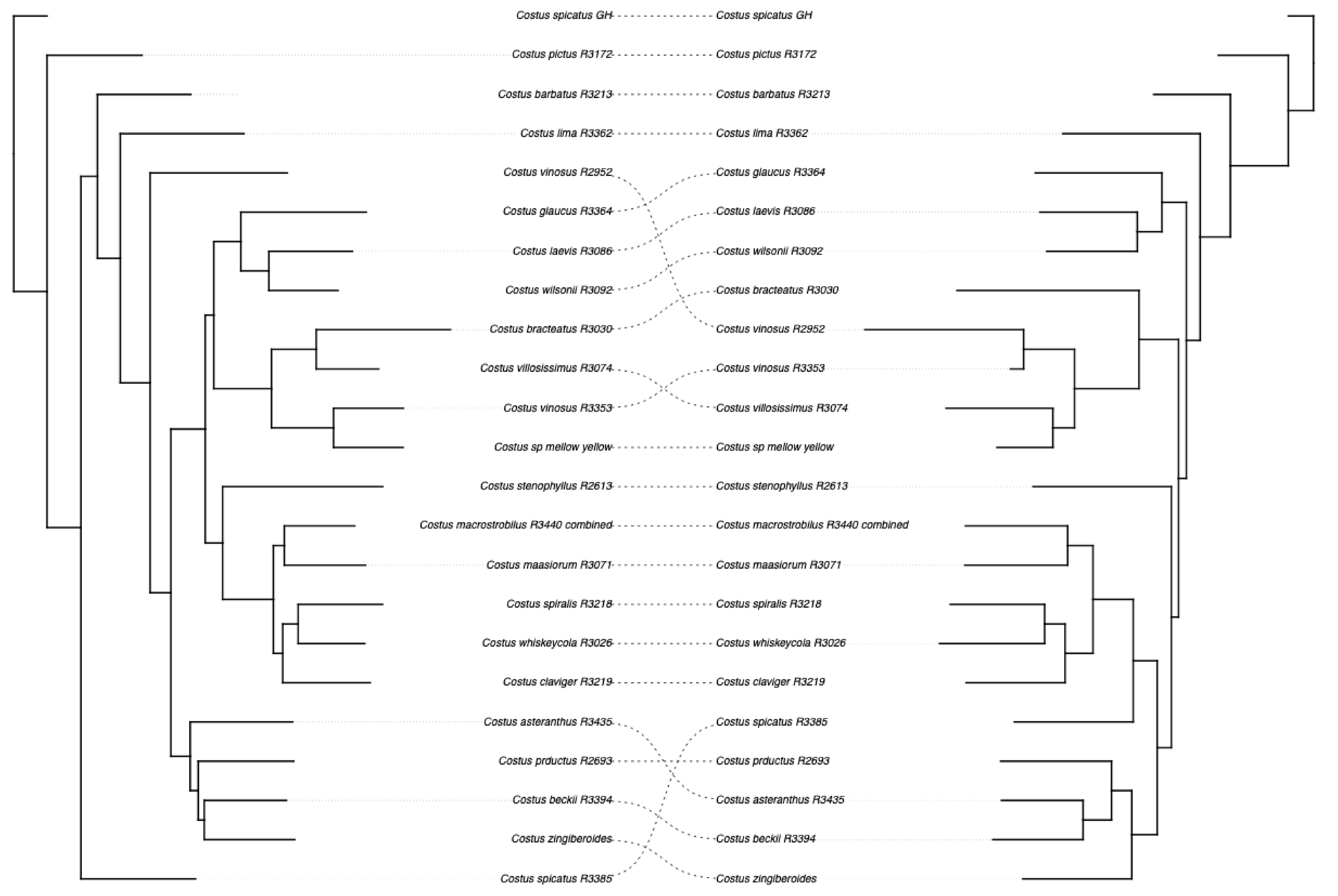

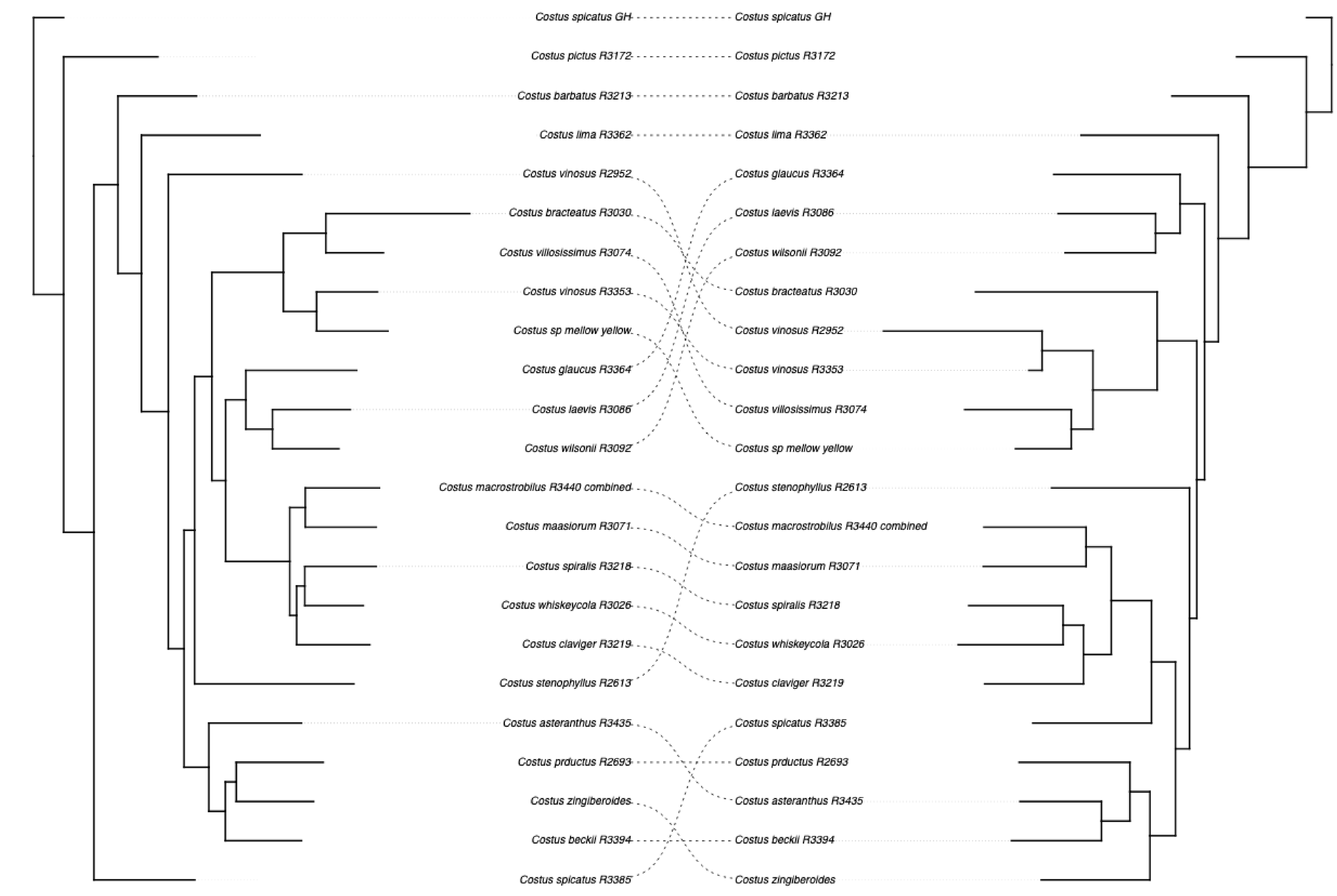

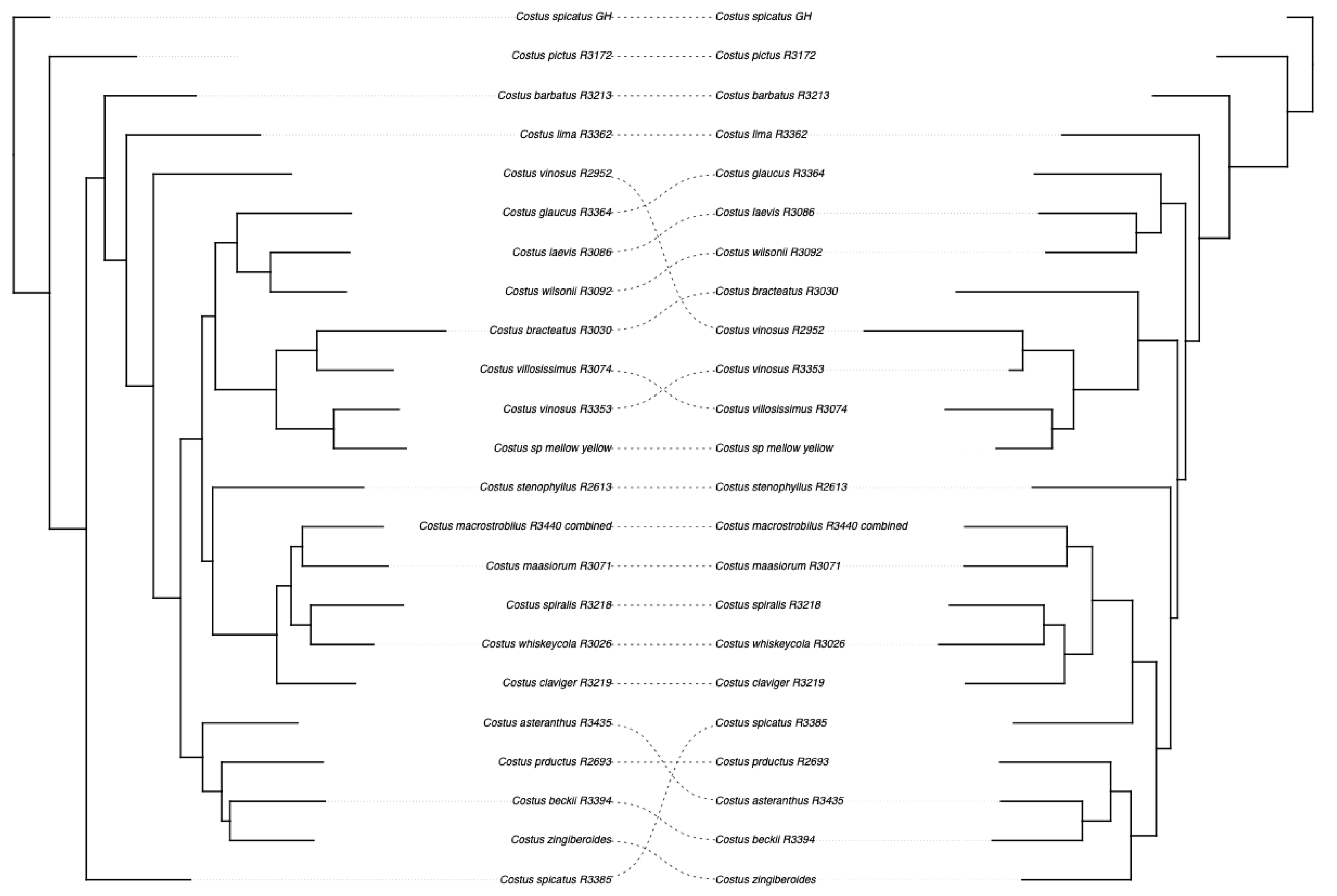

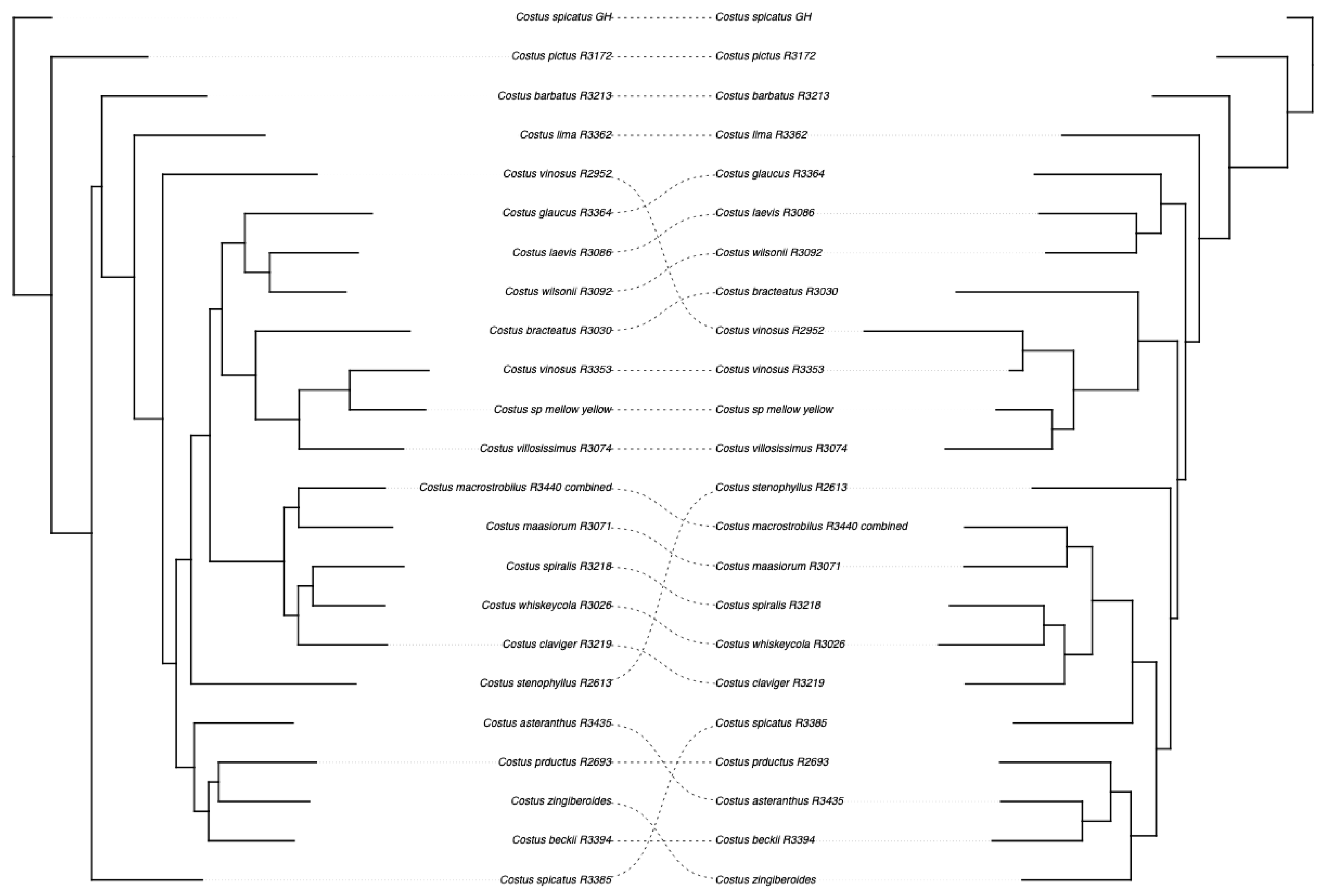

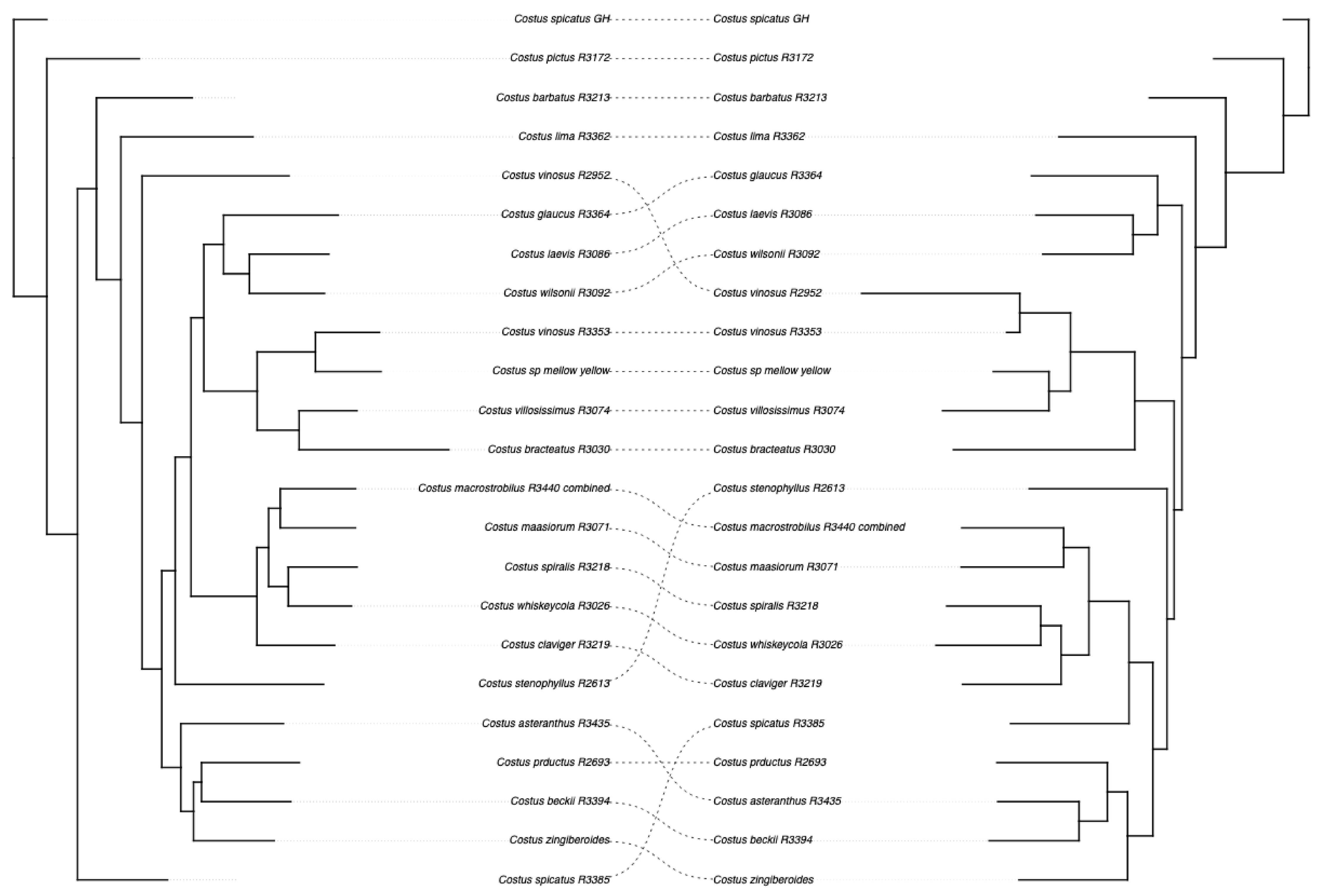

### Supplemental Figure 4

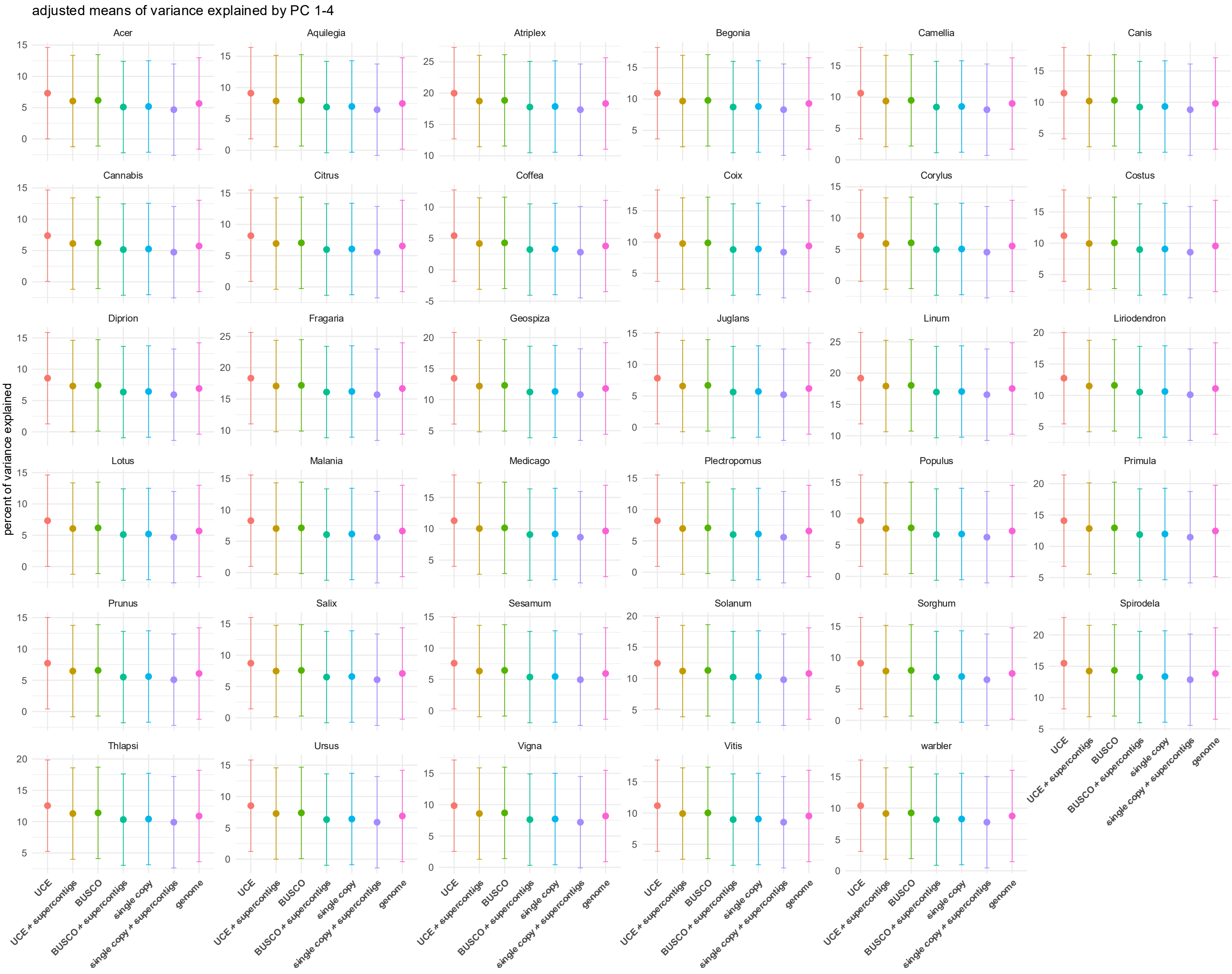

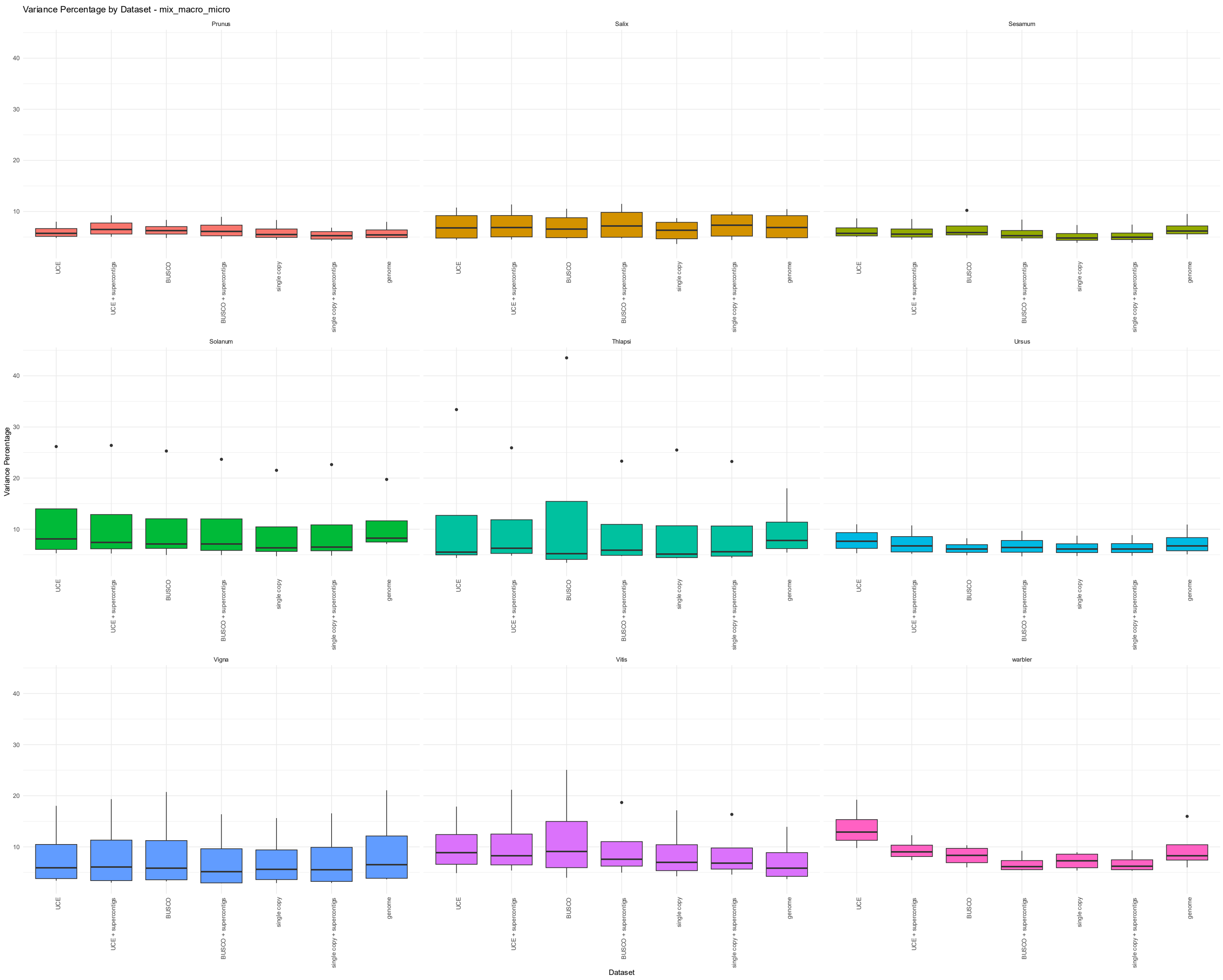

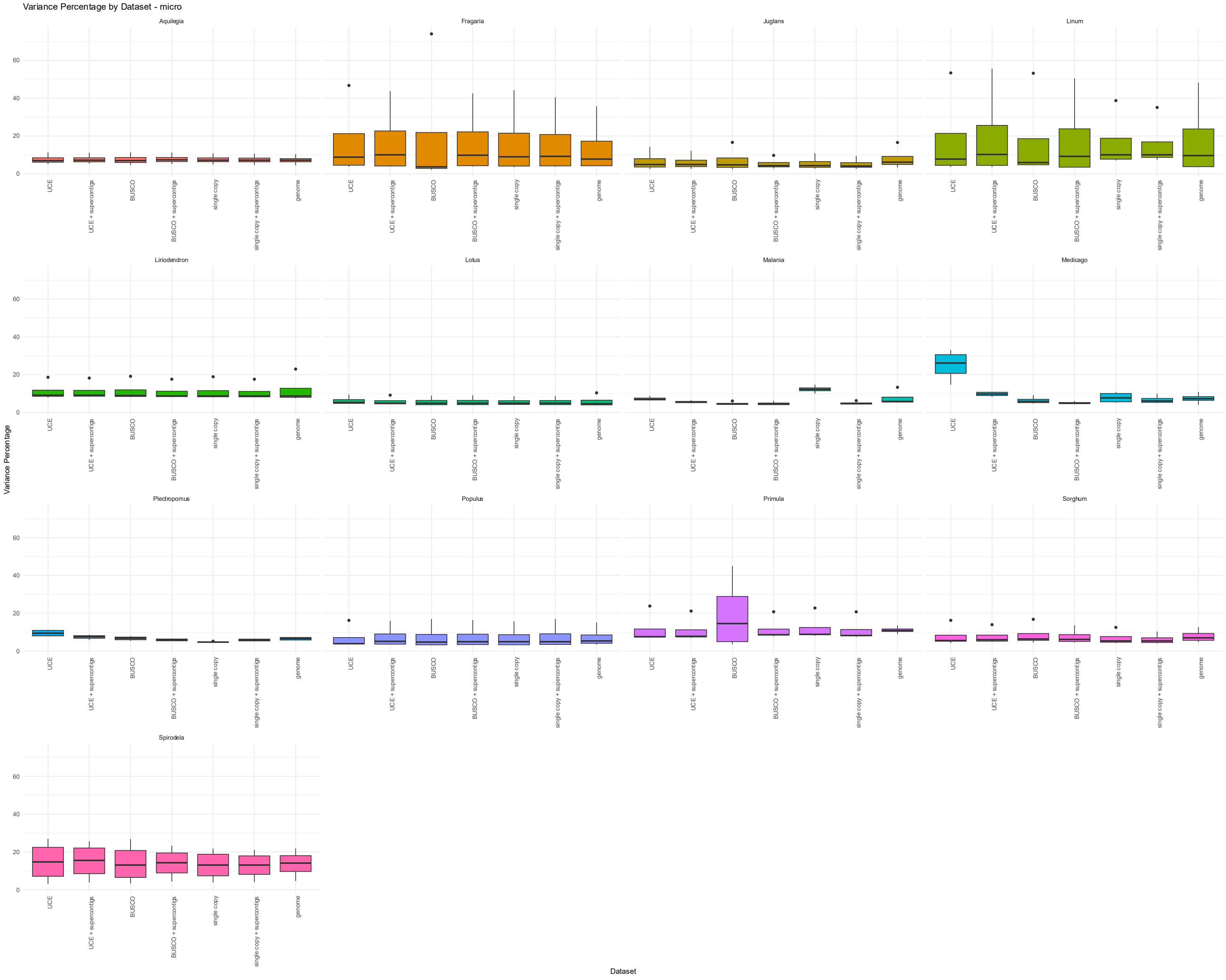

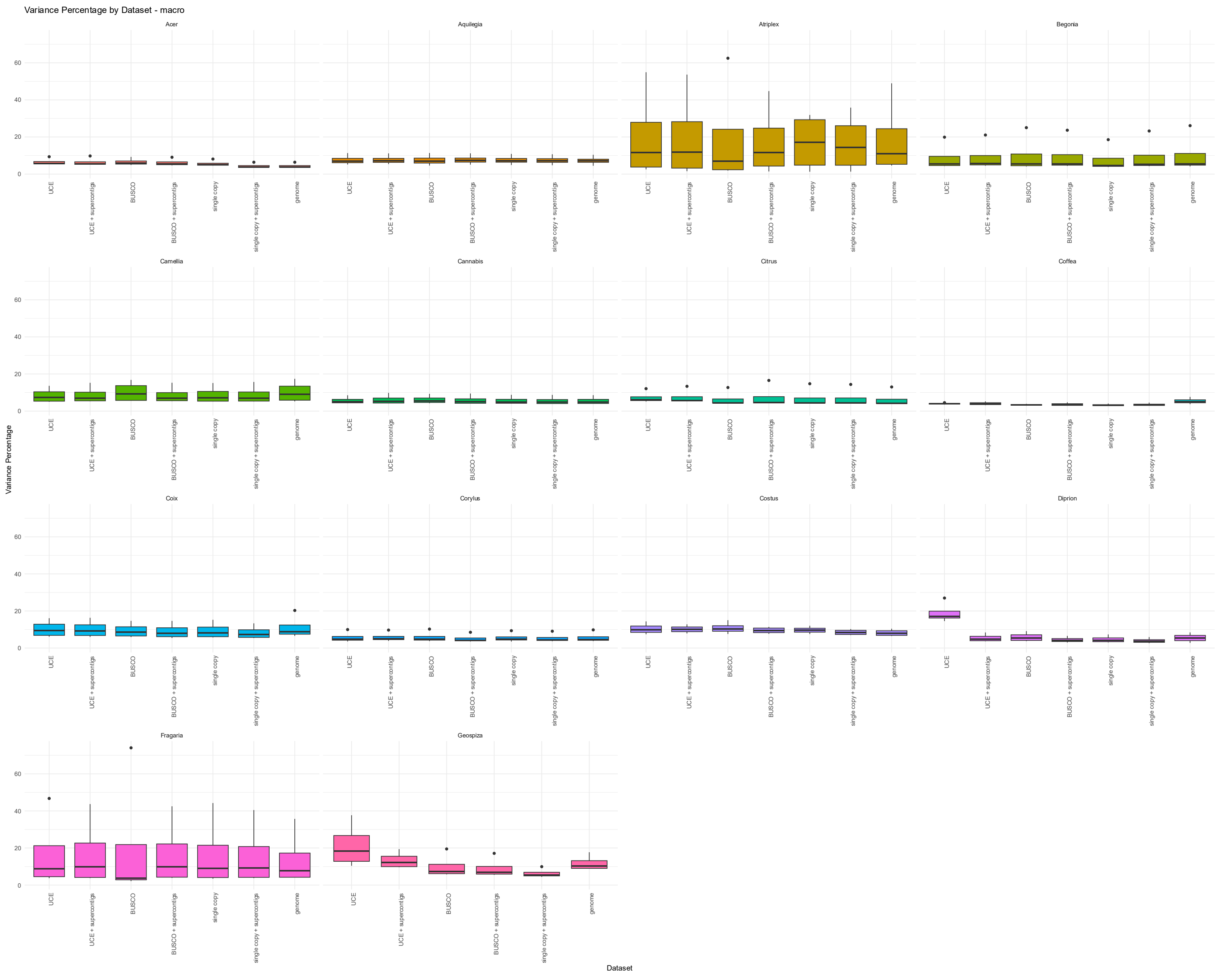
