## Supplemental Figure 3 for "The power to resolve relationships: identifying incongruence and precision of reduced representation and genome-wide data in phylogenomics and population genomics"

Acer_yangbiense – Acer_UCE

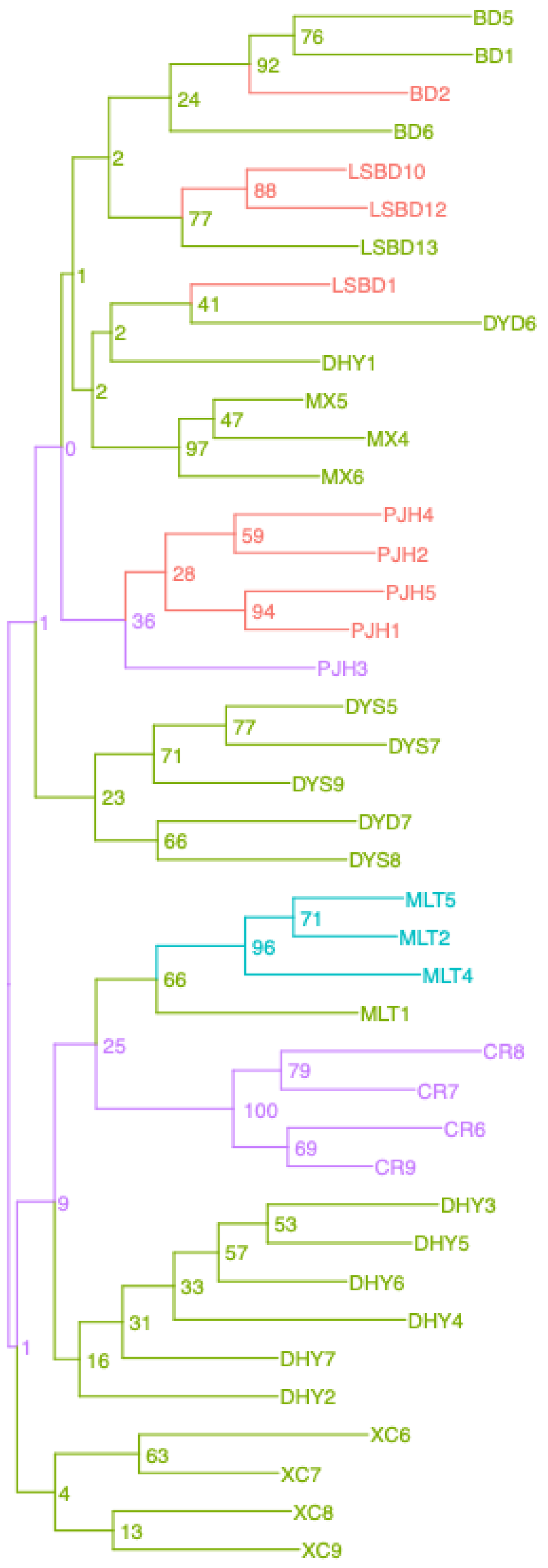


Acer_yangbiense – Acer_BUSCO

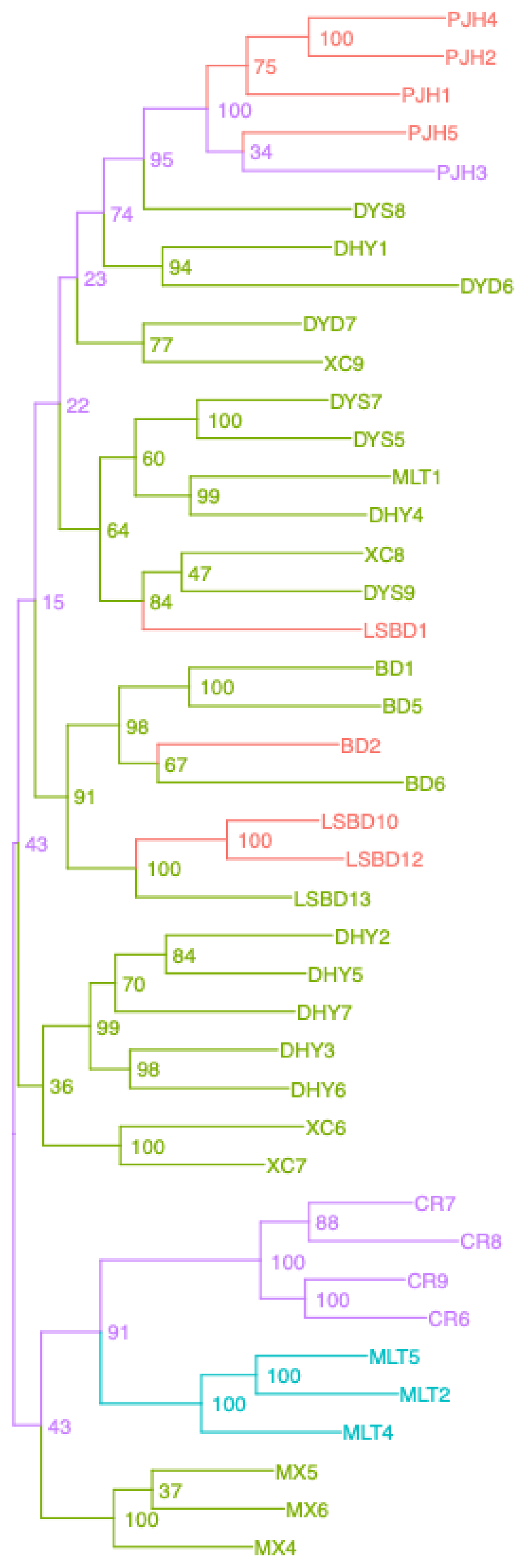


Acer_yangbiense – Acer_singlecopy

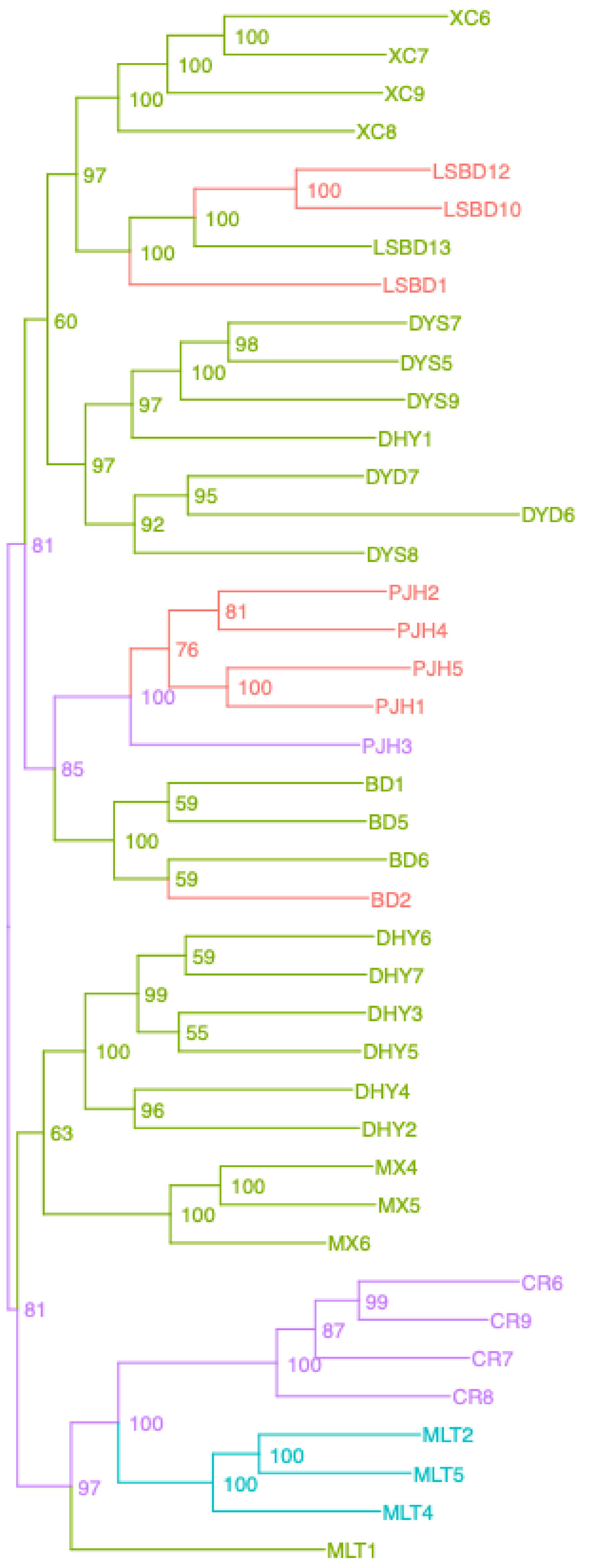


Acer_yangbiense – Acer_UCEsuper

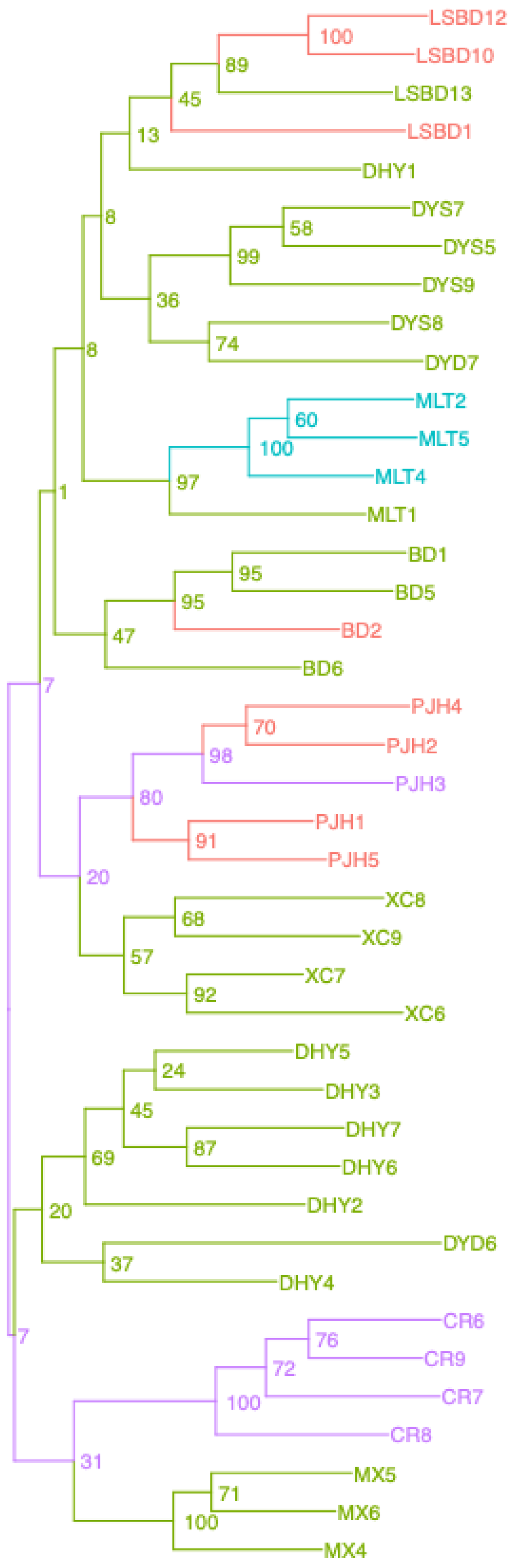


Acer_yangbiense – Acer_BUSCOsuper

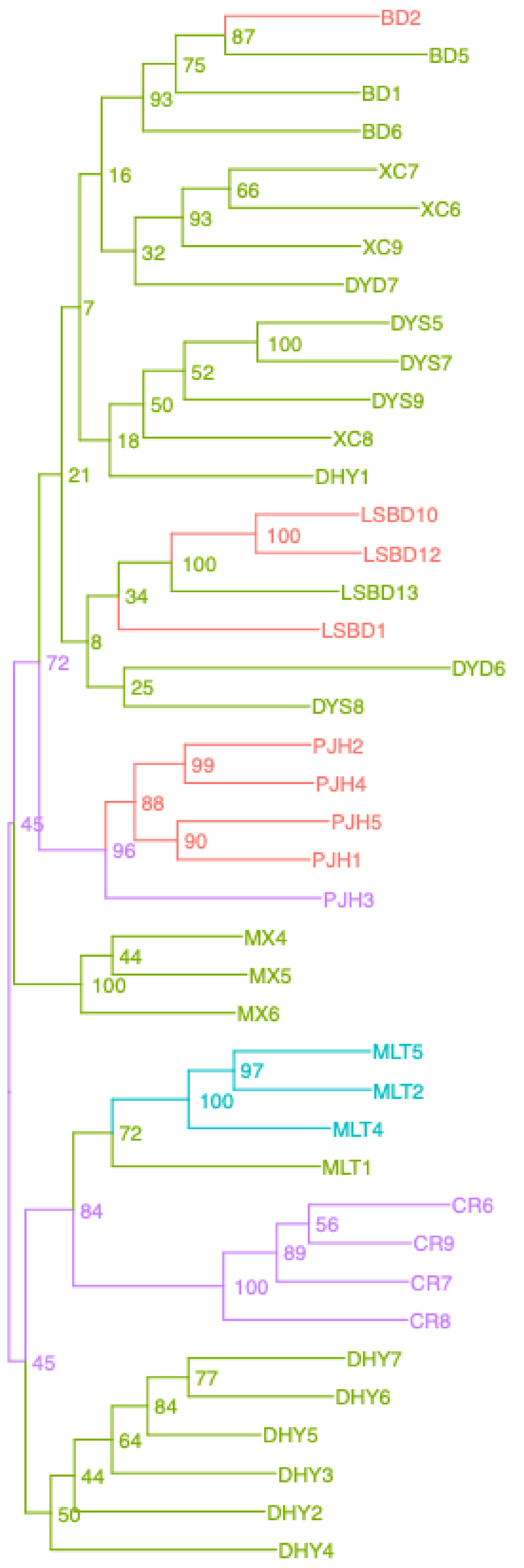


Acer_yangbiense – Acer_SCsuper

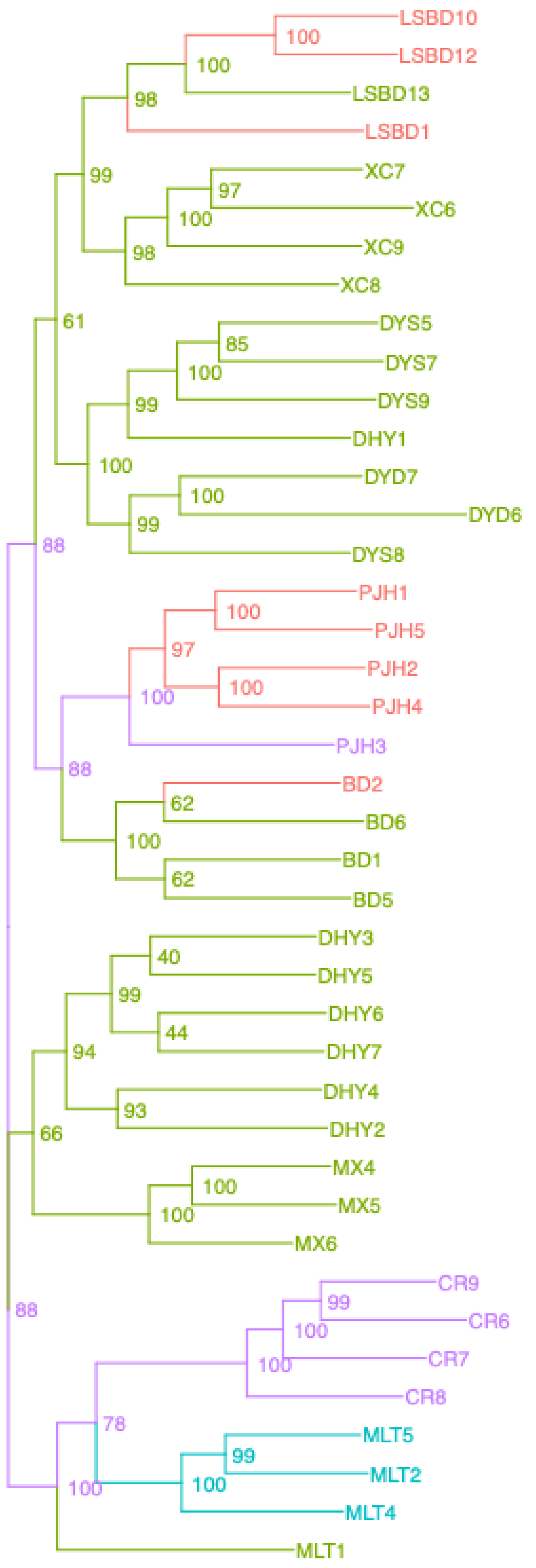


Acer_yangbiense – Acer_genome

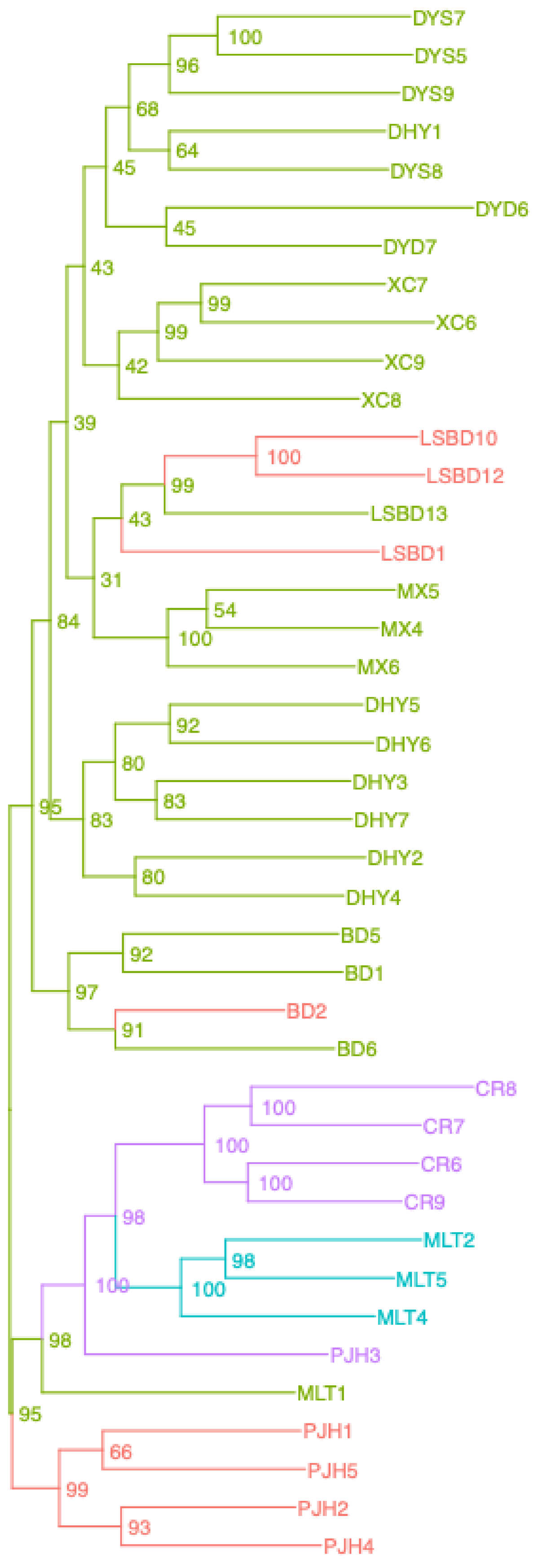

rooted_trees – Aquilegia_A353gene

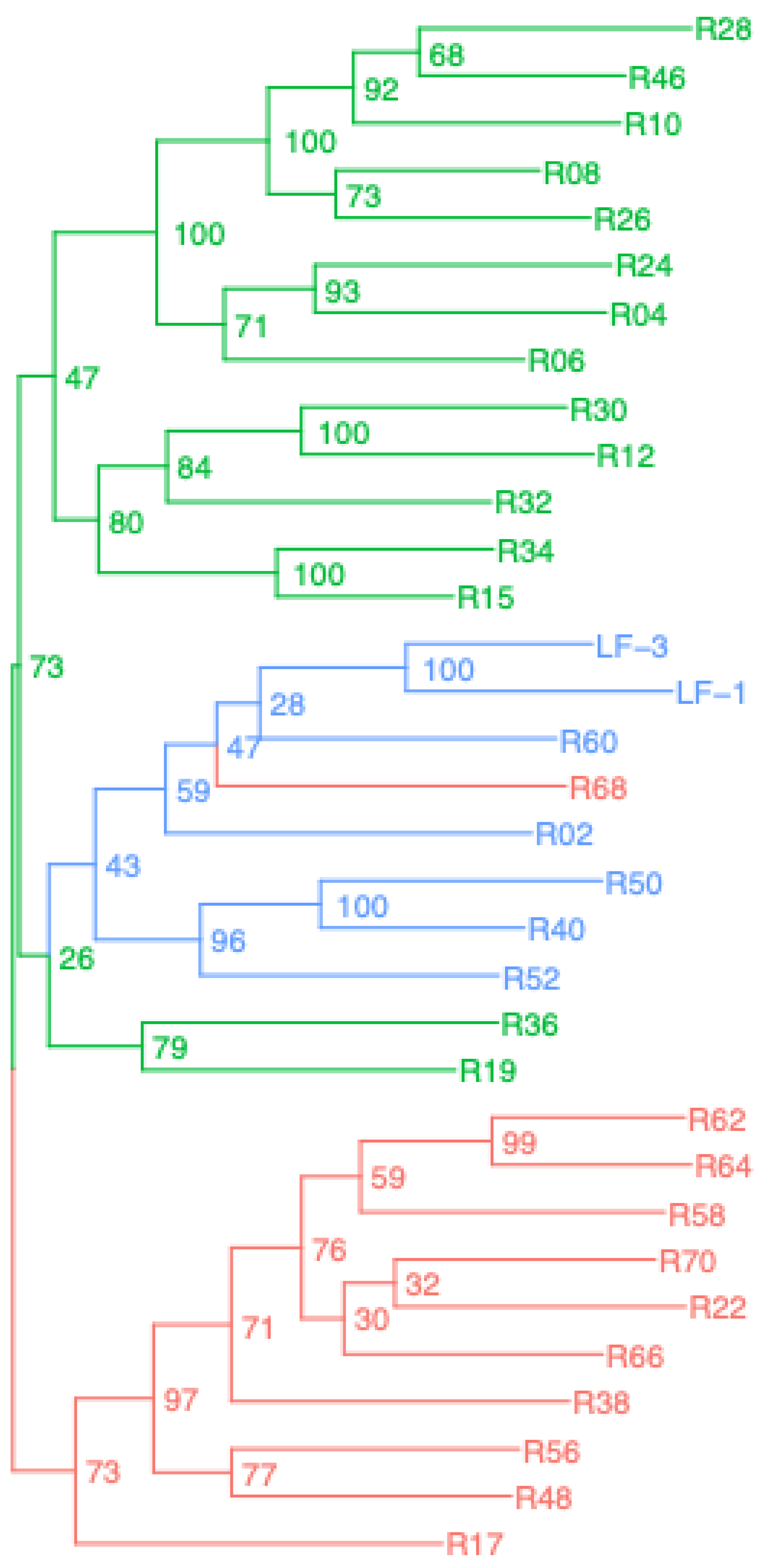


rooted_trees – Aquilegia_BUSCO

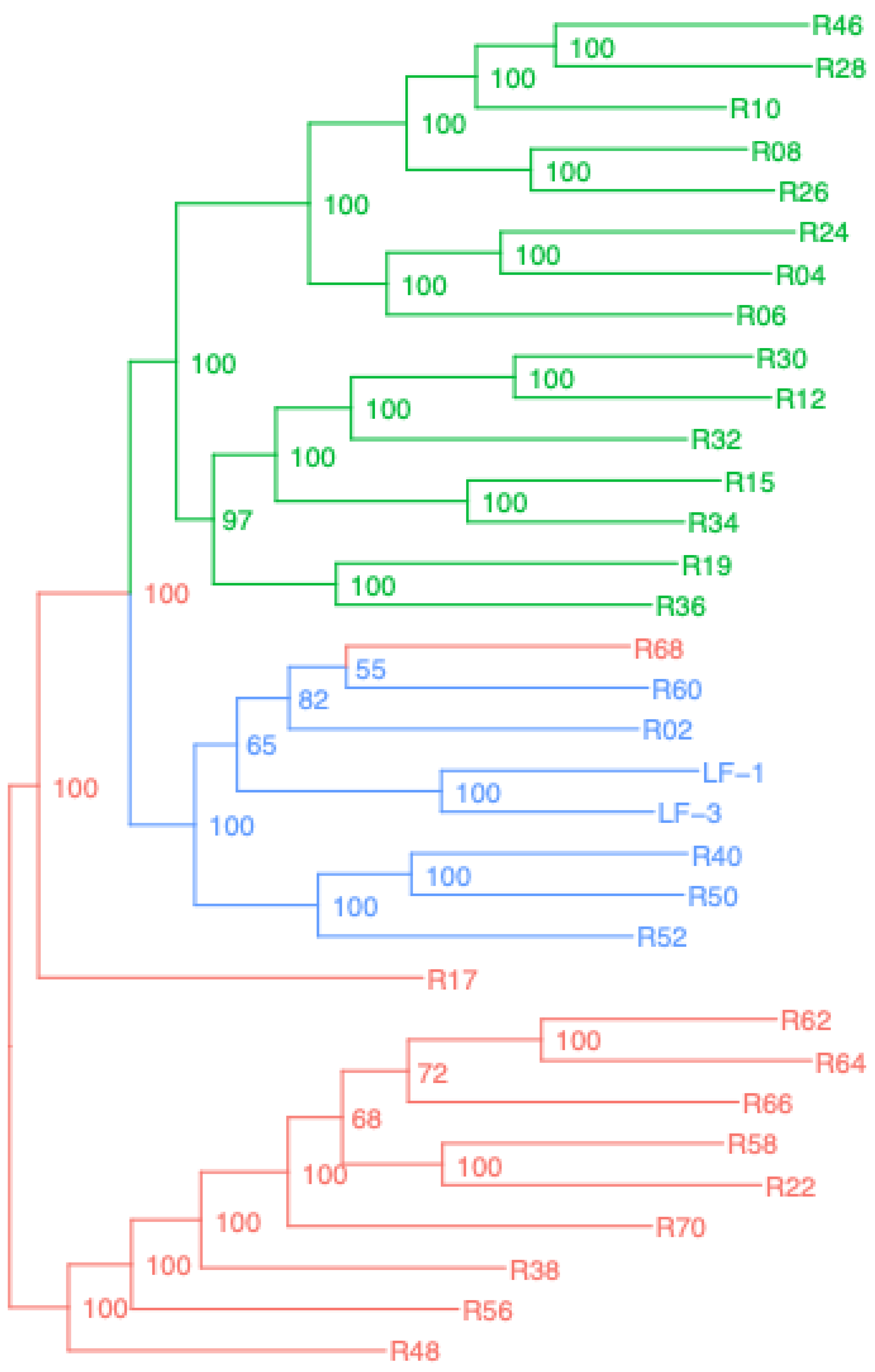


rooted_trees – Aquilegia_singlecopy

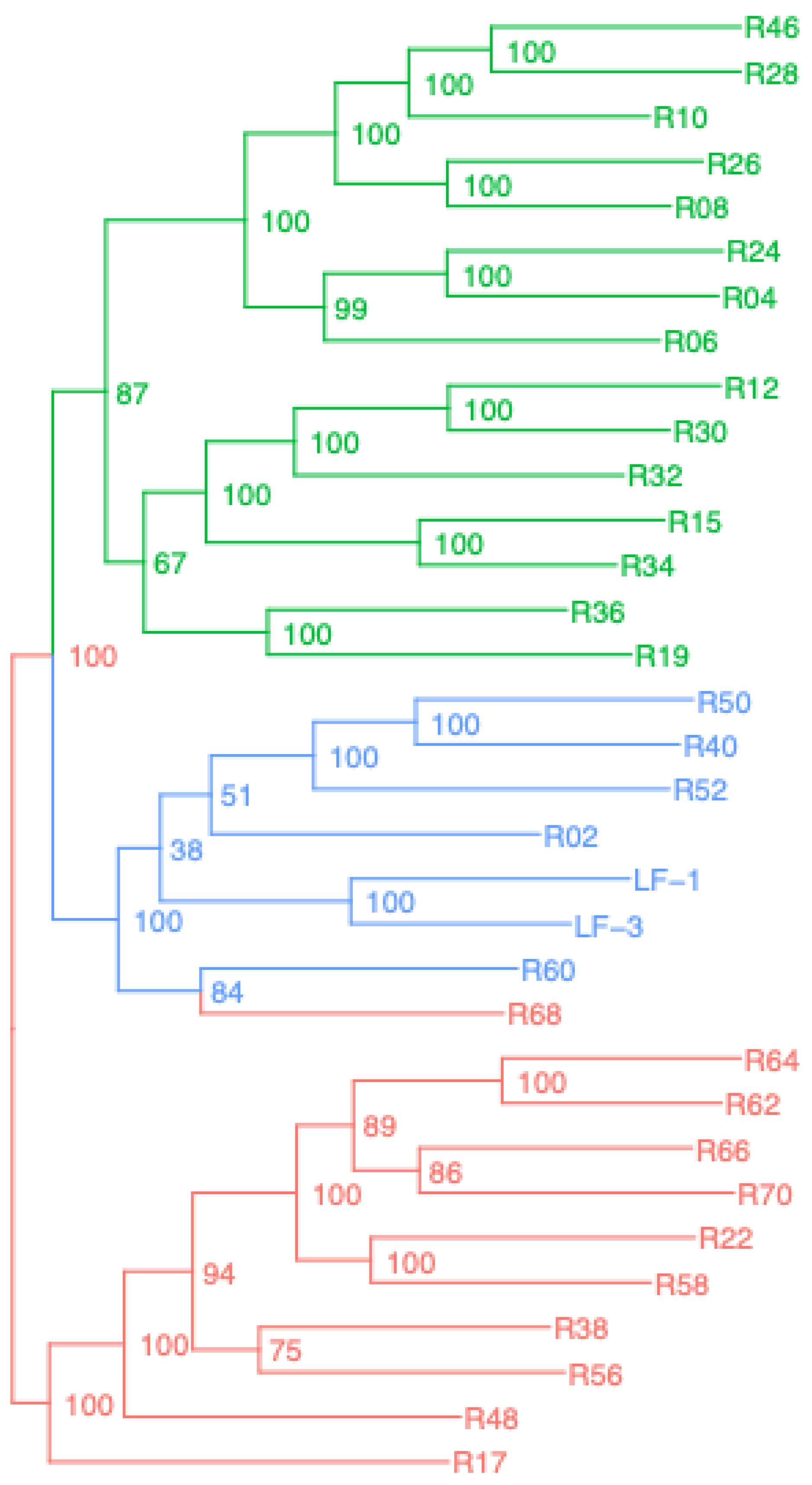


rooted_trees – Aquilegia_A353super

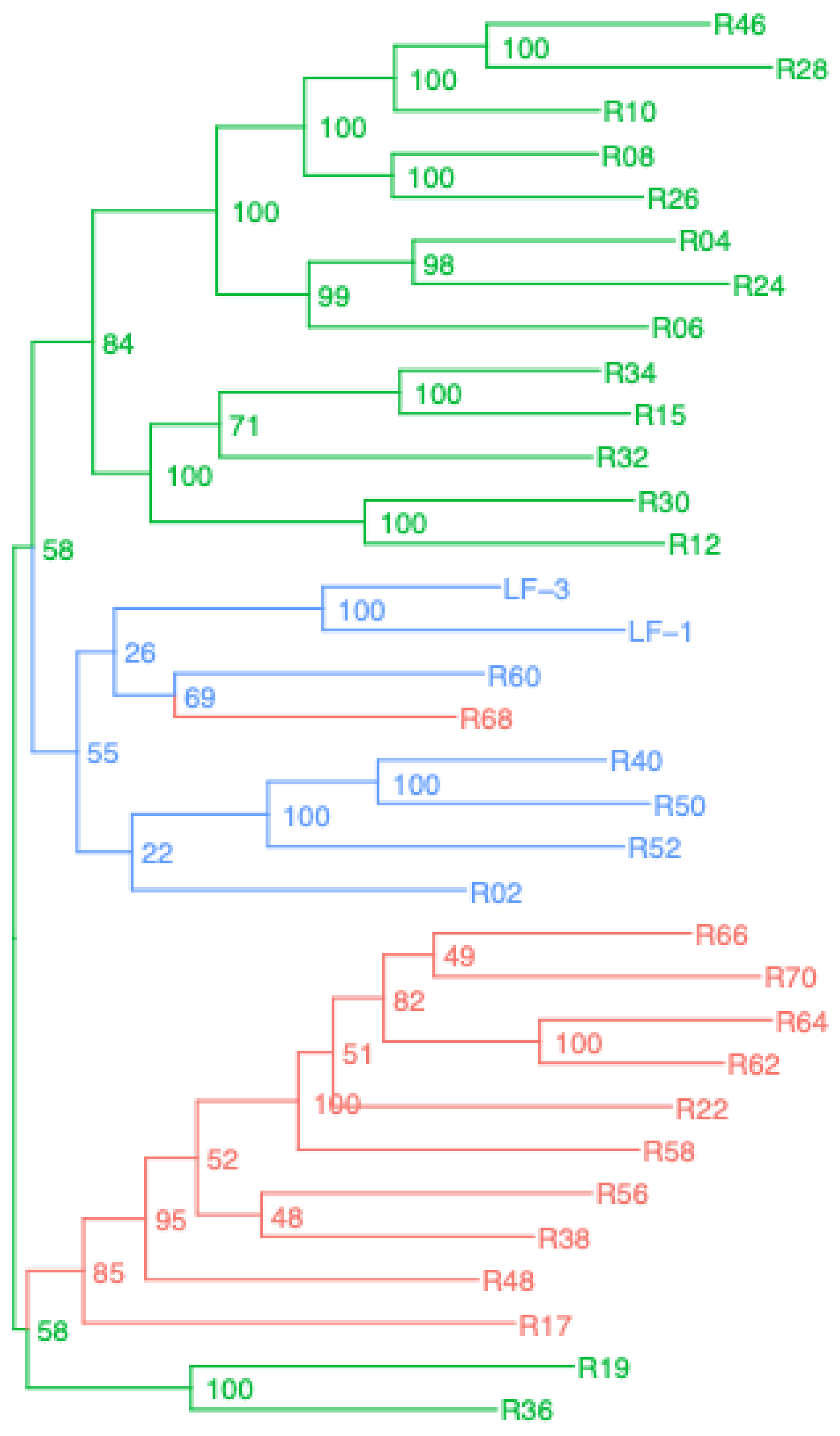


rooted_trees – Aquilegia_BUSCOsuper

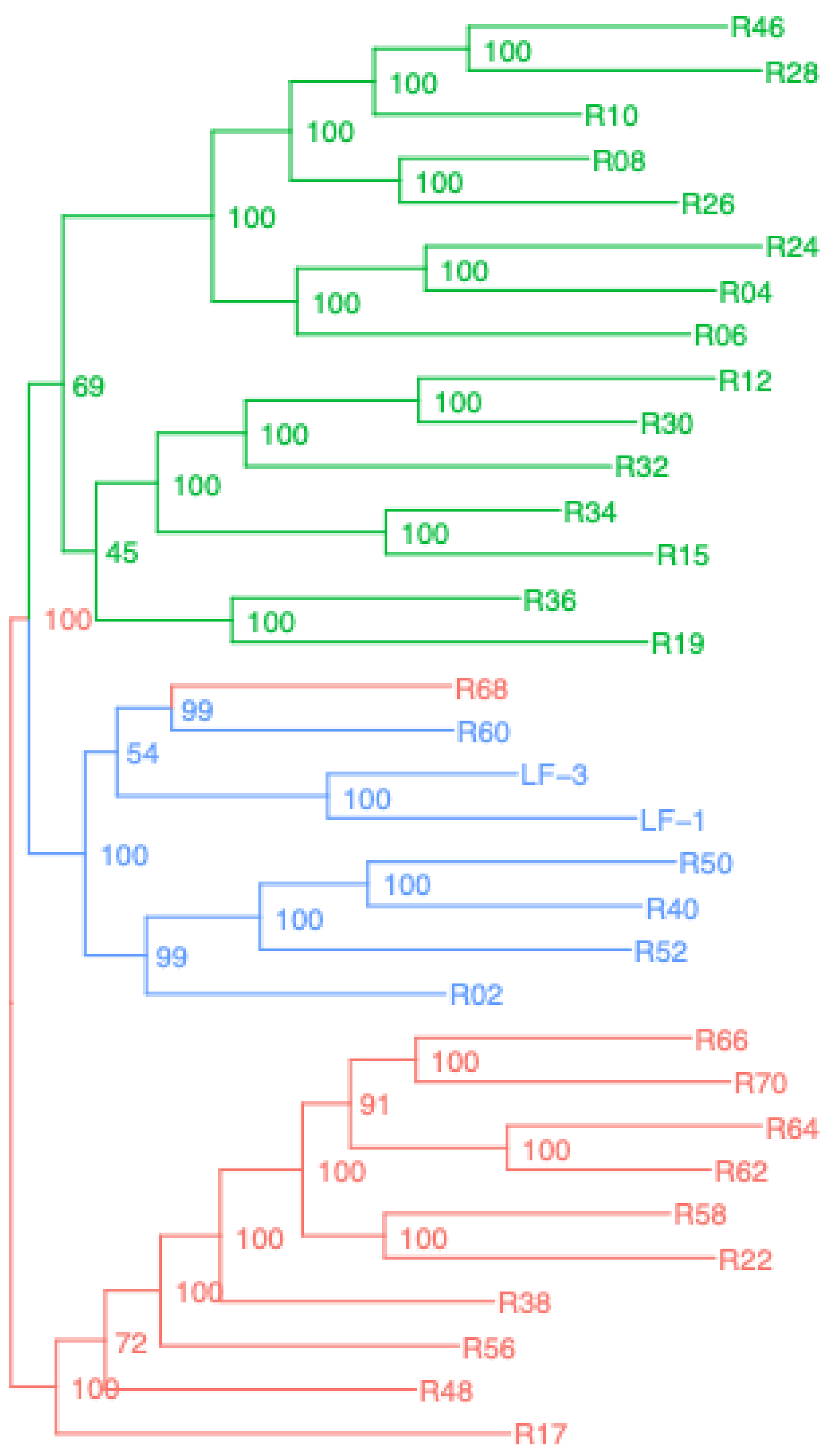


rooted_trees – Aquilegia_SCsuper

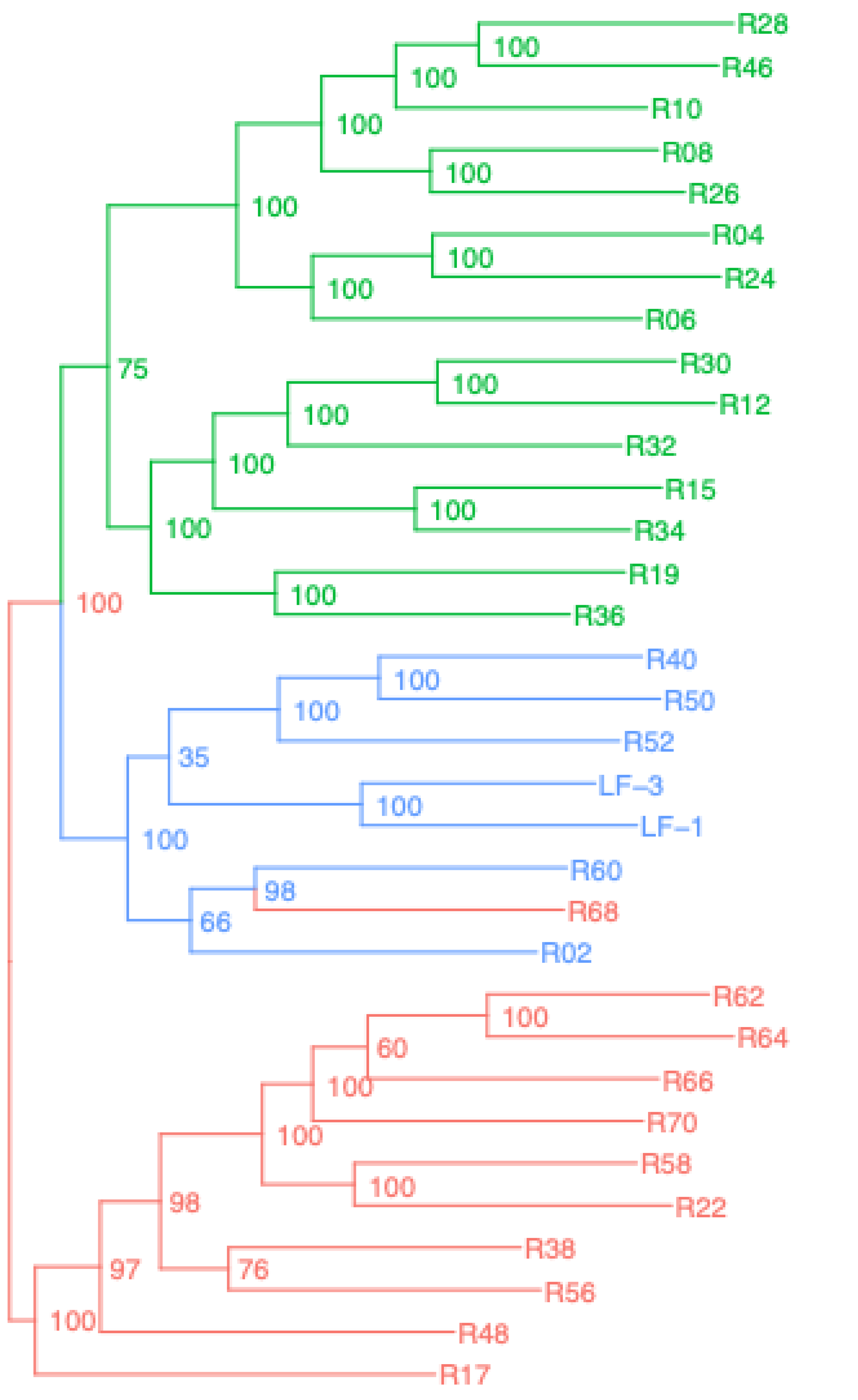


rooted_trees – Aquilegia_genome

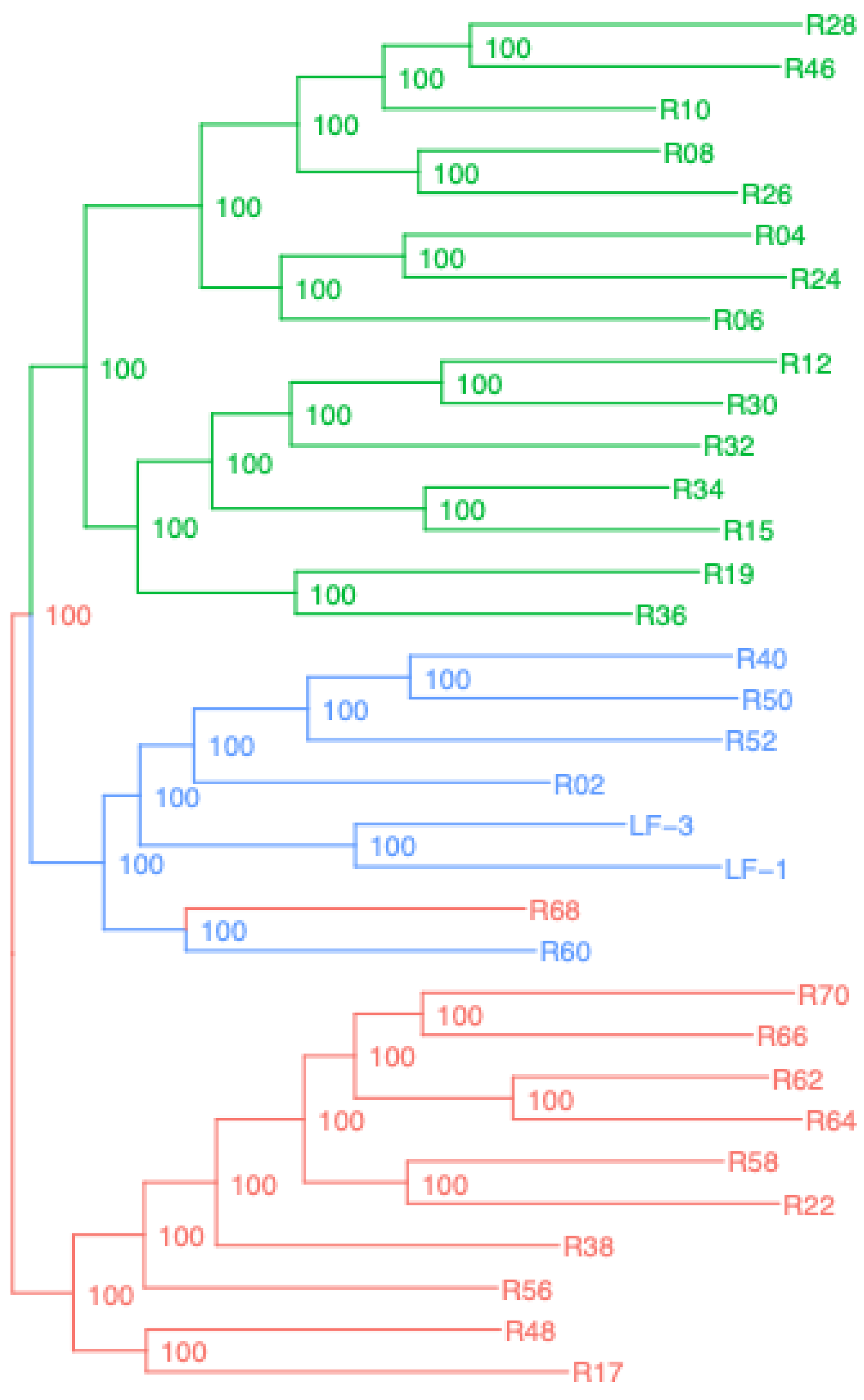

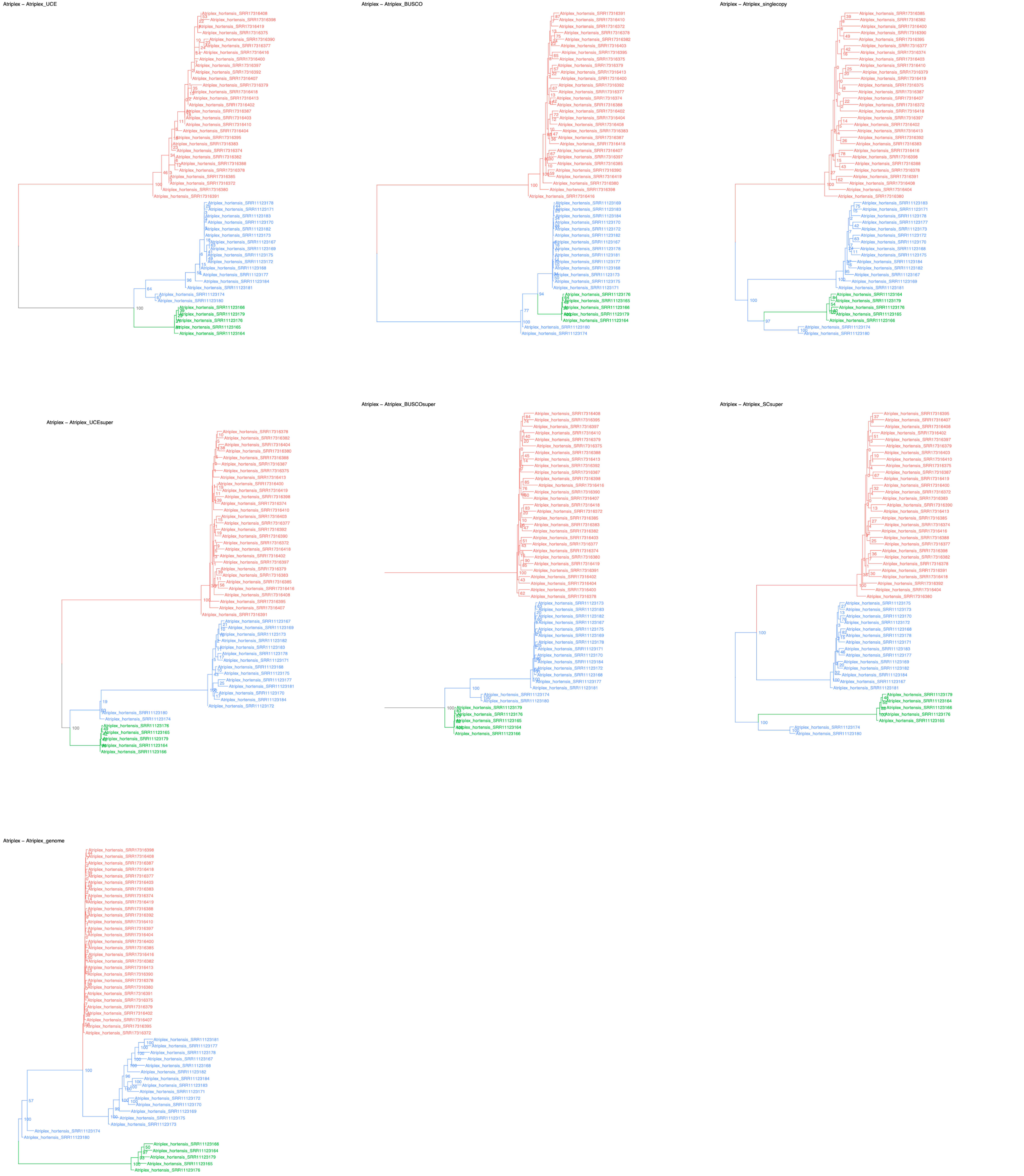
Atriplex – Atriplex_UCE
Atriplex – Atriplex_BUSCO
Atriplex – Atriplex_singlecopy
Atriplex – Atriplex_UCEsuper
Atriplex – Atriplex_BUSCOsuper
Atriplex – Atriplex_SCsuper
Atriplex – Atriplex_genome

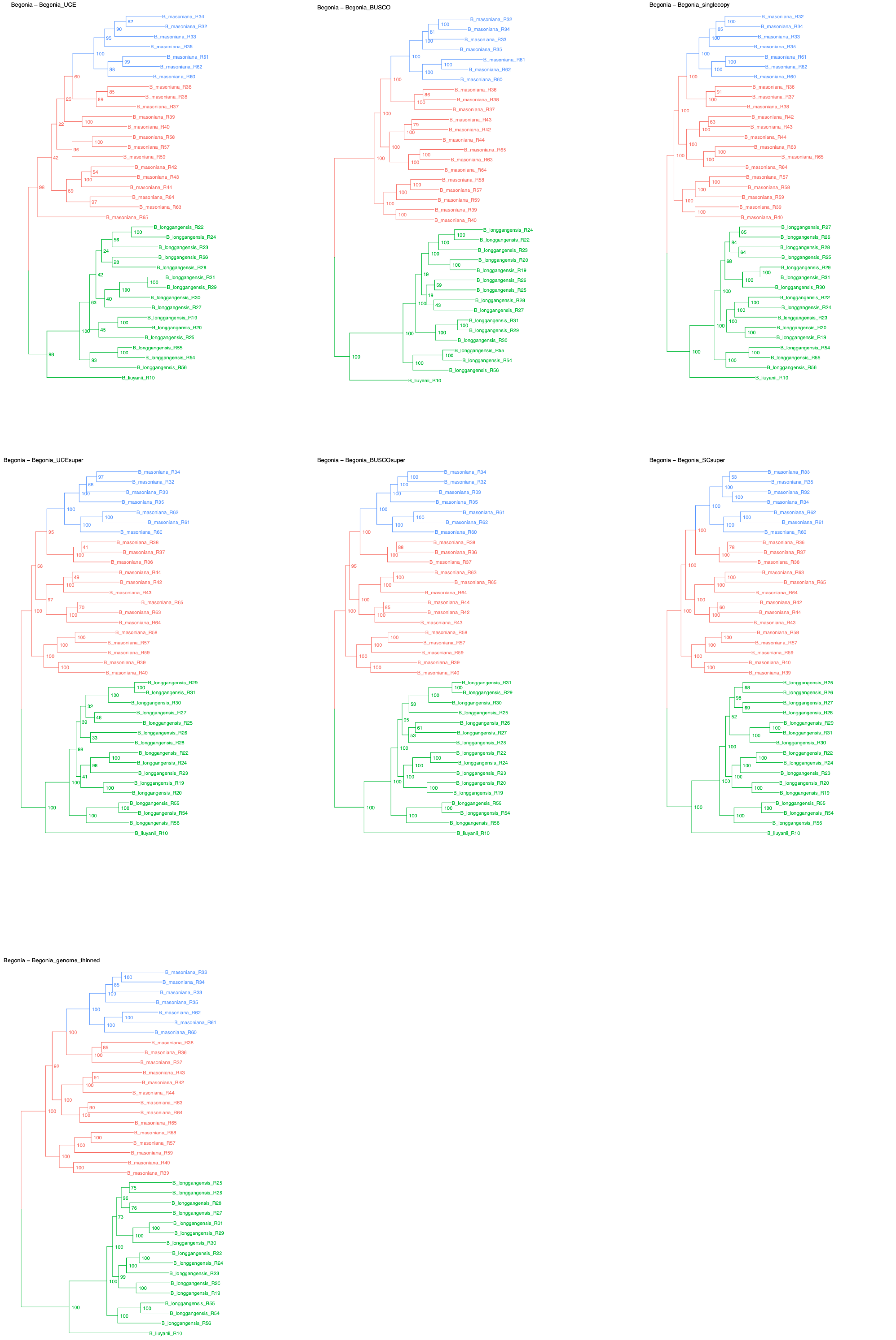
Begonia – Begonia_UCE
B_masoniana_R34
B_masoniana_R32
B_masoniana_R33
B_masoniana_R35
B_masoniana_R61
B_masoniana_R62
B_masoniana_R60
B_masoniana_R36
B_masoniana_R38
B_masoniana_R37
B_masoniana_R39
B_masoniana_R40
B_masoniana_R58
B_masoniana_R57
B_masoniana_R59
B_masoniana_R42
B_masoniana_R43
B_masoniana_R44
B_masoniana_R64
B_masoniana_R63
B_masoniana_R65
B_longgangensis_R22
B_longgangensis_R24
B_longgangensis_R23
B_longgangensis_R26
B_longgangensis_R28
B_longgangensis_R31
B_longgangensis_R29
B_longgangensis_R30
B_longgangensis_R27
B_longgangensis_R19
B_longgangensis_R20
B_longgangensis_R25
B_longgangensis_R55
B_longgangensis_R54
B_longgangensis_R56
B_liuyanii_R10
Begonia – Begonia_BUSCO
B_masoniana_R32
B_masoniana_R34
B_masoniana_R33
B_masoniana_R35
B_masoniana_R61
B_masoniana_R62
B_masoniana_R60
B_masoniana_R36
B_masoniana_R38
B_masoniana_R37
B_masoniana_R43
B_masoniana_R42
B_masoniana_R44
B_masoniana_R65
B_masoniana_R63
B_masoniana_R64
B_masoniana_R58
B_masoniana_R57
B_masoniana_R59
B_masoniana_R39
B_masoniana_R40
B_longgangensis_R24
B_longgangensis_R22
B_longgangensis_R23
B_longgangensis_R20
B_longgangensis_R19
B_longgangensis_R26
B_longgangensis_R25
B_longgangensis_R28
B_longgangensis_R27
B_longgangensis_R31
B_longgangensis_R29
B_longgangensis_R30
B_longgangensis_R55
B_longgangensis_R54
B_longgangensis_R56
B_liuyanii_R10
Begonia – Begonia_singlecopy
B_masoniana_R32
B_masoniana_R34
B_masoniana_R33
B_masoniana_R35
B_masoniana_R61
B_masoniana_R62
B_masoniana_R60
B_masoniana_R36
B_masoniana_R37
B_masoniana_R38
B_masoniana_R42
B_masoniana_R43
B_masoniana_R44
B_masoniana_R63
B_masoniana_R65
B_masoniana_R64
B_masoniana_R57
B_masoniana_R58
B_masoniana_R59
B_masoniana_R39
B_masoniana_R40
B_longgangensis_R27
B_longgangensis_R26
B_longgangensis_R28
B_longgangensis_R25
B_longgangensis_R29
B_longgangensis_R31
B_longgangensis_R30
B_longgangensis_R22
B_longgangensis_R24
B_longgangensis_R23
B_longgangensis_R20
B_longgangensis_R19
B_longgangensis_R54
B_longgangensis_R55
B_longgangensis_R56
B_liuyanii_R10
Begonia – Begonia_UCEsuper
B_masoniana_R34
B_masoniana_R32
B_masoniana_R33
B_masoniana_R35
B_masoniana_R62
B_masoniana_R61
B_masoniana_R60
B_masoniana_R38
B_masoniana_R37
B_masoniana_R36
B_masoniana_R44
B_masoniana_R42
B_masoniana_R43
B_masoniana_R65
B_masoniana_R63
B_masoniana_R64
B_masoniana_R58
B_masoniana_R57
B_masoniana_R59
B_masoniana_R39
B_masoniana_R40
B_longgangensis_R29
B_longgangensis_R31
B_longgangensis_R30
B_longgangensis_R27
B_longgangensis_R25
B_longgangensis_R26
B_longgangensis_R28
B_longgangensis_R22
B_longgangensis_R24
B_longgangensis_R23
B_longgangensis_R19
B_longgangensis_R20
B_longgangensis_R55
B_longgangensis_R54
B_longgangensis_R56
B_liuyanii_R10
Begonia – Begonia_BUSCOsuper
B_masoniana_R34
B_masoniana_R32
B_masoniana_R33
B_masoniana_R35
B_masoniana_R61
B_masoniana_R62
B_masoniana_R60
B_masoniana_R38
B_masoniana_R36
B_masoniana_R37
B_masoniana_R63
B_masoniana_R65
B_masoniana_R64
B_masoniana_R44
B_masoniana_R42
B_masoniana_R43
B_masoniana_R58
B_masoniana_R57
B_masoniana_R59
B_masoniana_R39
B_masoniana_R40
B_longgangensis_R31
B_longgangensis_R29
B_longgangensis_R30
B_longgangensis_R25
B_longgangensis_R26
B_longgangensis_R27
B_longgangensis_R28
B_longgangensis_R22
B_longgangensis_R24
B_longgangensis_R23
B_longgangensis_R20
B_longgangensis_R19
B_longgangensis_R55
B_longgangensis_R54
B_longgangensis_R56
B_liuyanii_R10
Begonia – Begonia_SCsuper
B_masoniana_R33
B_masoniana_R35
B_masoniana_R32
B_masoniana_R34
B_masoniana_R62
B_masoniana_R61
B_masoniana_R60
B_masoniana_R36
B_masoniana_R37
B_masoniana_R38
B_masoniana_R63
B_masoniana_R65
B_masoniana_R64
B_masoniana_R42
B_masoniana_R44
B_masoniana_R43
B_masoniana_R58
B_masoniana_R57
B_masoniana_R59
B_masoniana_R40
B_masoniana_R39
B_longgangensis_R25
B_longgangensis_R26
B_longgangensis_R27
B_longgangensis_R28
B_longgangensis_R29
B_longgangensis_R31
B_longgangensis_R30
B_longgangensis_R22
B_longgangensis_R24
B_longgangensis_R23
B_longgangensis_R20
B_longgangensis_R19
B_longgangensis_R55
B_longgangensis_R54
B_longgangensis_R56
B_liuyanii_R10
Begonia – Begonia_genome_thinned
B_masoniana_R32
B_masoniana_R34
B_masoniana_R33
B_masoniana_R35
B_masoniana_R62
B_masoniana_R61
B_masoniana_R60
B_masoniana_R38
B_masoniana_R36
B_masoniana_R37
B_masoniana_R43
B_masoniana_R42
B_masoniana_R44
B_masoniana_R63
B_masoniana_R64
B_masoniana_R65
B_masoniana_R58
B_masoniana_R57
B_masoniana_R59
B_masoniana_R40
B_masoniana_R39
B_longgangensis_R25
B_longgangensis_R26
B_longgangensis_R28
B_longgangensis_R27
B_longgangensis_R31
B_longgangensis_R29
B_longgangensis_R30
B_longgangensis_R22
B_longgangensis_R24
B_longgangensis_R23
B_longgangensis_R20
B_longgangensis_R19
B_longgangensis_R55
B_longgangensis_R54
B_longgangensis_R56
B_liuyanii_R10

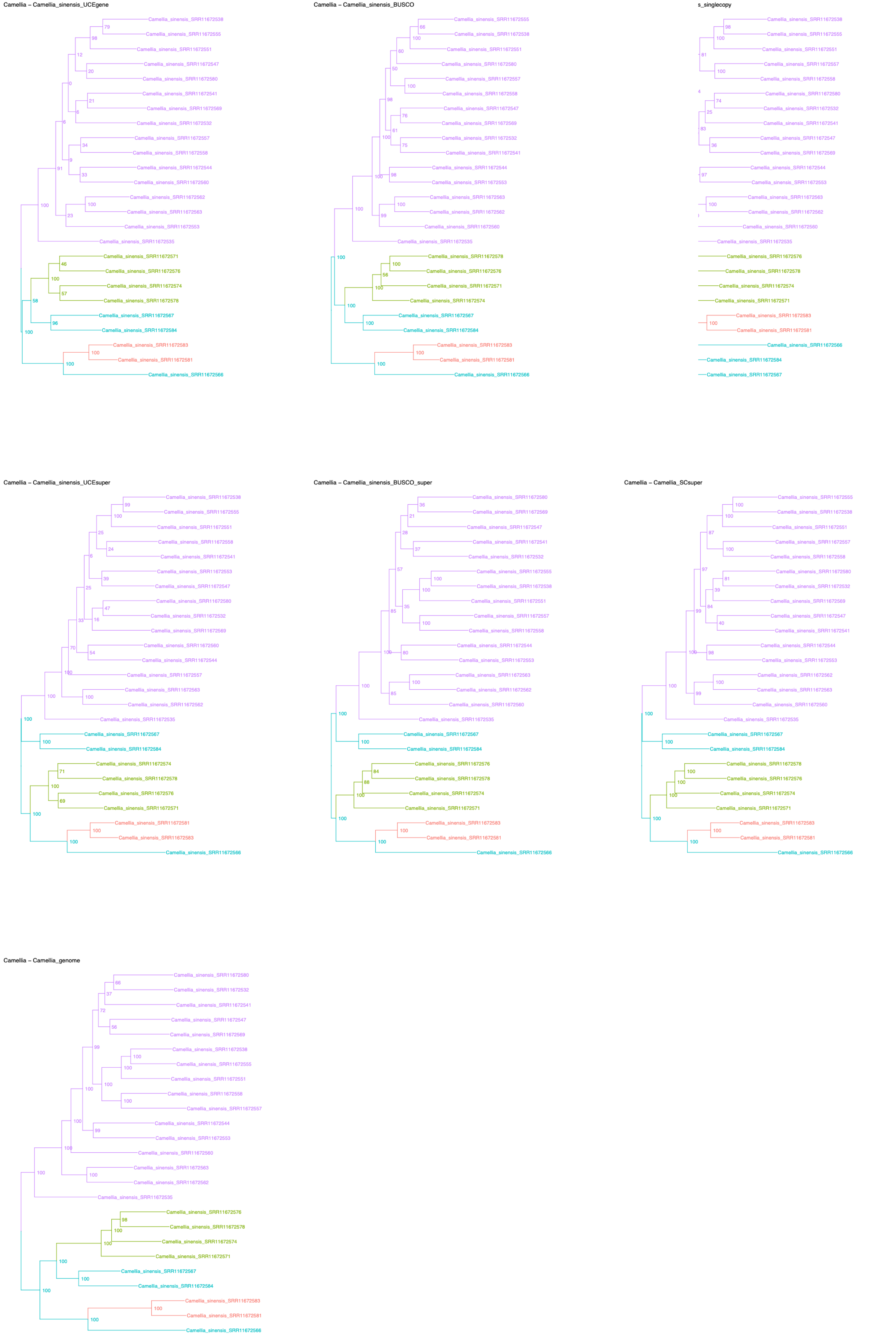
Camellia – Camellia_sinensis_UCEgene
Camellia_sinensis_SRR11672538
Camellia_sinensis_SRR11672555
Camellia_sinensis_SRR11672551
Camellia_sinensis_SRR11672547
Camellia_sinensis_SRR11672580
Camellia_sinensis_SRR11672541
Camellia_sinensis_SRR11672569
Camellia_sinensis_SRR11672532
Camellia_sinensis_SRR11672557
Camellia_sinensis_SRR11672558
Camellia_sinensis_SRR11672544
Camellia_sinensis_SRR11672560
Camellia_sinensis_SRR11672562
Camellia_sinensis_SRR11672563
Camellia_sinensis_SRR11672553
Camellia_sinensis_SRR11672535
Camellia_sinensis_SRR11672571
Camellia_sinensis_SRR11672576
Camellia_sinensis_SRR11672574
Camellia_sinensis_SRR11672578
Camellia_sinensis_SRR11672567
Camellia_sinensis_SRR11672584
Camellia_sinensis_SRR11672583
Camellia_sinensis_SRR11672581
Camellia_sinensis_SRR11672566
Camellia – Camellia_sinensis_BUSCO
Camellia_sinensis_SRR11672555
Camellia_sinensis_SRR11672538
Camellia_sinensis_SRR11672551
Camellia_sinensis_SRR11672580
Camellia_sinensis_SRR11672557
Camellia_sinensis_SRR11672558
Camellia_sinensis_SRR11672547
Camellia_sinensis_SRR11672569
Camellia_sinensis_SRR11672532
Camellia_sinensis_SRR11672541
Camellia_sinensis_SRR11672544
Camellia_sinensis_SRR11672553
Camellia_sinensis_SRR11672563
Camellia_sinensis_SRR11672562
Camellia_sinensis_SRR11672560
Camellia_sinensis_SRR11672535
Camellia_sinensis_SRR11672578
Camellia_sinensis_SRR11672576
Camellia_sinensis_SRR11672571
Camellia_sinensis_SRR11672574
Camellia_sinensis_SRR11672567
Camellia_sinensis_SRR11672584
Camellia_sinensis_SRR11672583
Camellia_sinensis_SRR11672581
Camellia_sinensis_SRR11672566
s_singlecopy
Camellia_sinensis_SRR11672538
Camellia_sinensis_SRR11672555
Camellia_sinensis_SRR11672551
Camellia_sinensis_SRR11672557
Camellia_sinensis_SRR11672558
Camellia_sinensis_SRR11672580
Camellia_sinensis_SRR11672532
Camellia_sinensis_SRR11672541
Camellia_sinensis_SRR11672547
Camellia_sinensis_SRR11672569
Camellia_sinensis_SRR11672544
Camellia_sinensis_SRR11672553
Camellia_sinensis_SRR11672563
Camellia_sinensis_SRR11672562
Camellia_sinensis_SRR11672560
Camellia_sinensis_SRR11672535
Camellia_sinensis_SRR11672576
Camellia_sinensis_SRR11672578
Camellia_sinensis_SRR11672574
Camellia_sinensis_SRR11672571
Camellia_sinensis_SRR11672583
Camellia_sinensis_SRR11672581
Camellia_sinensis_SRR11672566
Camellia_sinensis_SRR11672584
Camellia_sinensis_SRR11672567
Camellia – Camellia_sinensis_UCEsuper
Camellia_sinensis_SRR11672538
Camellia_sinensis_SRR11672555
Camellia_sinensis_SRR11672551
Camellia_sinensis_SRR11672558
Camellia_sinensis_SRR11672541
Camellia_sinensis_SRR11672553
Camellia_sinensis_SRR11672547
Camellia_sinensis_SRR11672580
Camellia_sinensis_SRR11672532
Camellia_sinensis_SRR11672569
Camellia_sinensis_SRR11672560
Camellia_sinensis_SRR11672544
Camellia_sinensis_SRR11672557
Camellia_sinensis_SRR11672563
Camellia_sinensis_SRR11672562
Camellia_sinensis_SRR11672535
Camellia_sinensis_SRR11672567
Camellia_sinensis_SRR11672584
Camellia_sinensis_SRR11672574
Camellia_sinensis_SRR11672578
Camellia_sinensis_SRR11672576
Camellia_sinensis_SRR11672571
Camellia_sinensis_SRR11672581
Camellia_sinensis_SRR11672583
Camellia_sinensis_SRR11672566
Camellia – Camellia_sinensis_BUSCO_super
Camellia_sinensis_SRR11672580
Camellia_sinensis_SRR11672569
Camellia_sinensis_SRR11672547
Camellia_sinensis_SRR11672541
Camellia_sinensis_SRR11672532
Camellia_sinensis_SRR11672555
Camellia_sinensis_SRR11672538
Camellia_sinensis_SRR11672551
Camellia_sinensis_SRR11672557
Camellia_sinensis_SRR11672558
Camellia_sinensis_SRR11672544
Camellia_sinensis_SRR11672553
Camellia_sinensis_SRR11672563
Camellia_sinensis_SRR11672562
Camellia_sinensis_SRR11672560
Camellia_sinensis_SRR11672535
Camellia_sinensis_SRR11672567
Camellia_sinensis_SRR11672584
Camellia_sinensis_SRR11672576
Camellia_sinensis_SRR11672578
Camellia_sinensis_SRR11672574
Camellia_sinensis_SRR11672571
Camellia_sinensis_SRR11672583
Camellia_sinensis_SRR11672581
Camellia_sinensis_SRR11672566
Camellia – Camellia_SCsuper
Camellia_sinensis_SRR11672555
Camellia_sinensis_SRR11672538
Camellia_sinensis_SRR11672551
Camellia_sinensis_SRR11672557
Camellia_sinensis_SRR11672558
Camellia_sinensis_SRR11672580
Camellia_sinensis_SRR11672532
Camellia_sinensis_SRR11672569
Camellia_sinensis_SRR11672547
Camellia_sinensis_SRR11672541
Camellia_sinensis_SRR11672544
Camellia_sinensis_SRR11672553
Camellia_sinensis_SRR11672562
Camellia_sinensis_SRR11672563
Camellia_sinensis_SRR11672560
Camellia_sinensis_SRR11672535
Camellia_sinensis_SRR11672567
Camellia_sinensis_SRR11672584
Camellia_sinensis_SRR11672578
Camellia_sinensis_SRR11672576
Camellia_sinensis_SRR11672574
Camellia_sinensis_SRR11672571
Camellia_sinensis_SRR11672583
Camellia_sinensis_SRR11672581
Camellia_sinensis_SRR11672566
Camellia – Camellia_genome
Camellia_sinensis_SRR11672580
Camellia_sinensis_SRR11672532
Camellia_sinensis_SRR11672541
Camellia_sinensis_SRR11672547
Camellia_sinensis_SRR11672569
Camellia_sinensis_SRR11672538
Camellia_sinensis_SRR11672555
Camellia_sinensis_SRR11672551
Camellia_sinensis_SRR11672558
Camellia_sinensis_SRR11672557
Camellia_sinensis_SRR11672544
Camellia_sinensis_SRR11672553
Camellia_sinensis_SRR11672560
Camellia_sinensis_SRR11672563
Camellia_sinensis_SRR11672562
Camellia_sinensis_SRR11672535
Camellia_sinensis_SRR11672576
Camellia_sinensis_SRR11672578
Camellia_sinensis_SRR11672574
Camellia_sinensis_SRR11672571
Camellia_sinensis_SRR11672567
Camellia_sinensis_SRR11672584
Camellia_sinensis_SRR11672583
Camellia_sinensis_SRR11672581
Camellia_sinensis_SRR11672566

Cannabis – Cannabis_UCE

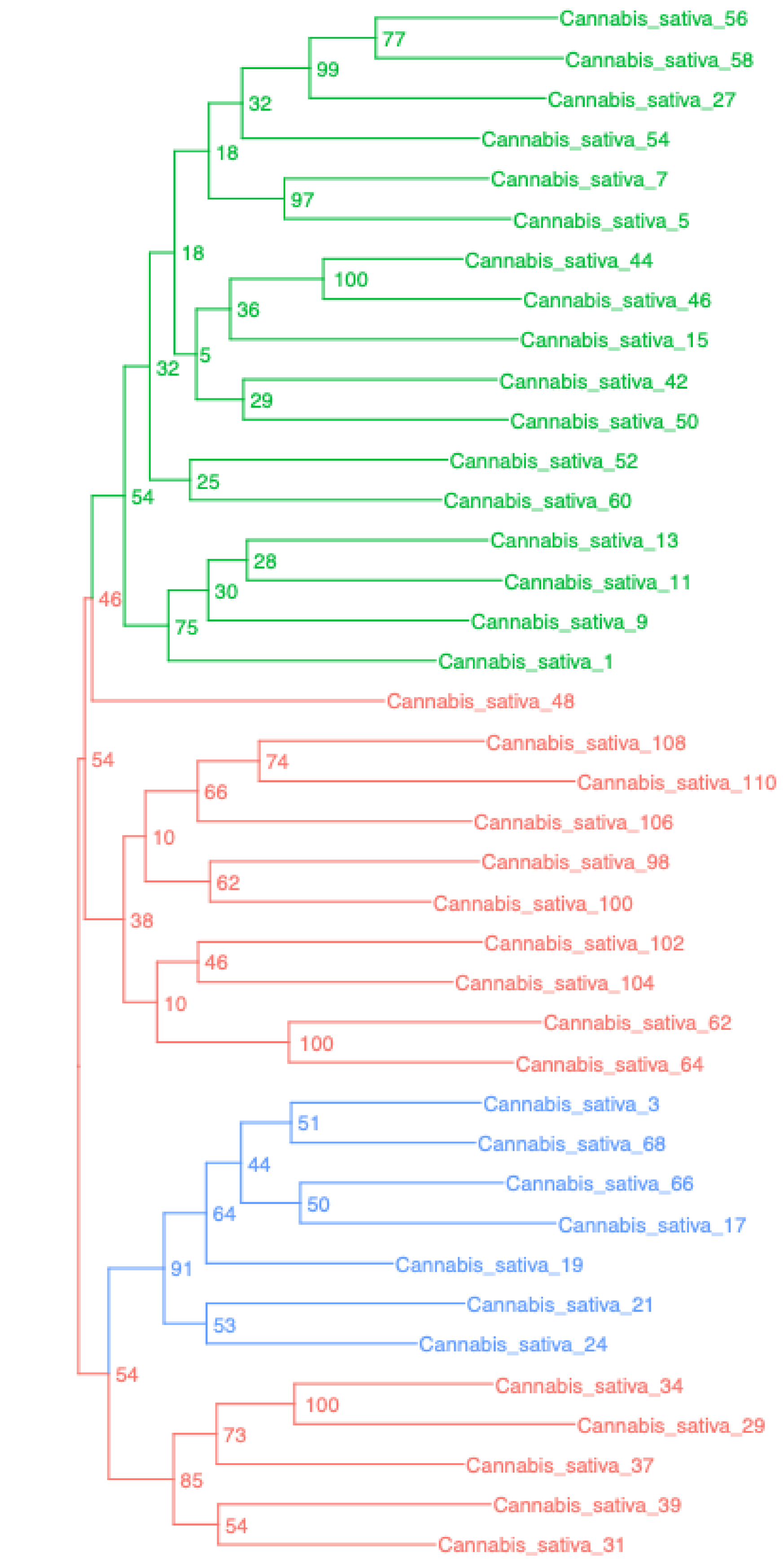


Cannabis – Cannabis_BUSCO

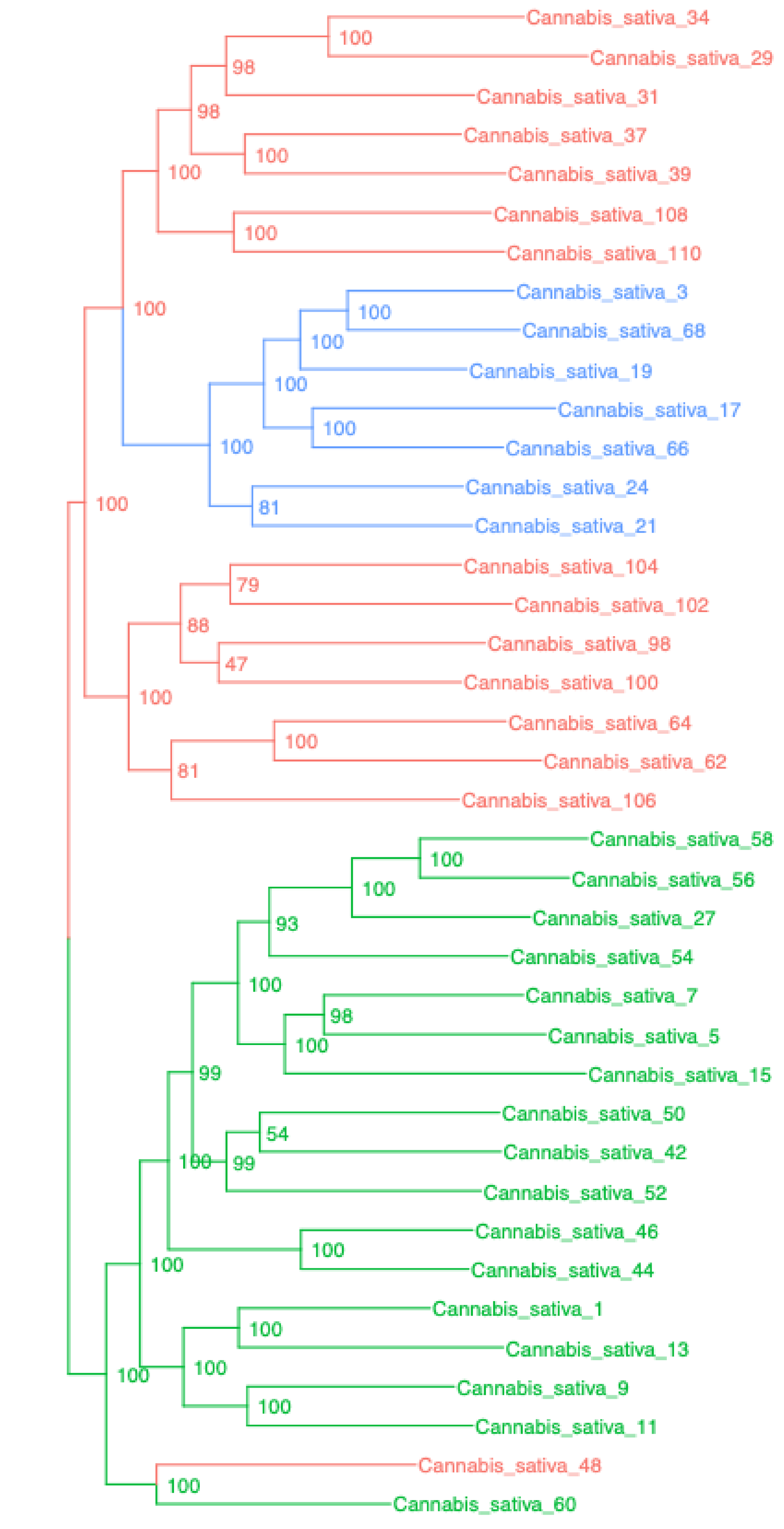


Cannabis – Cannabis_singlecopy

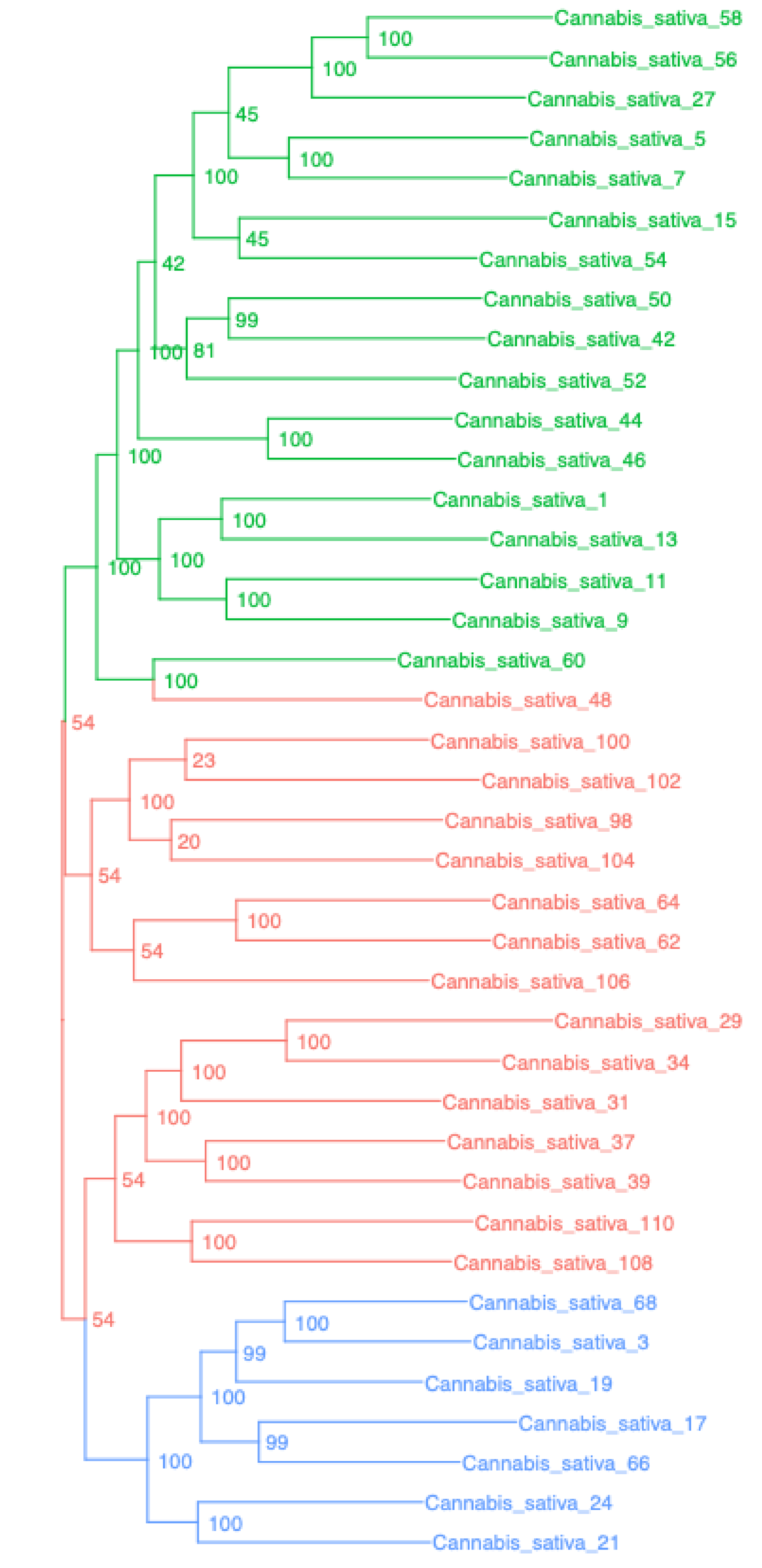


Cannabis – Cannabis_UCEsuper

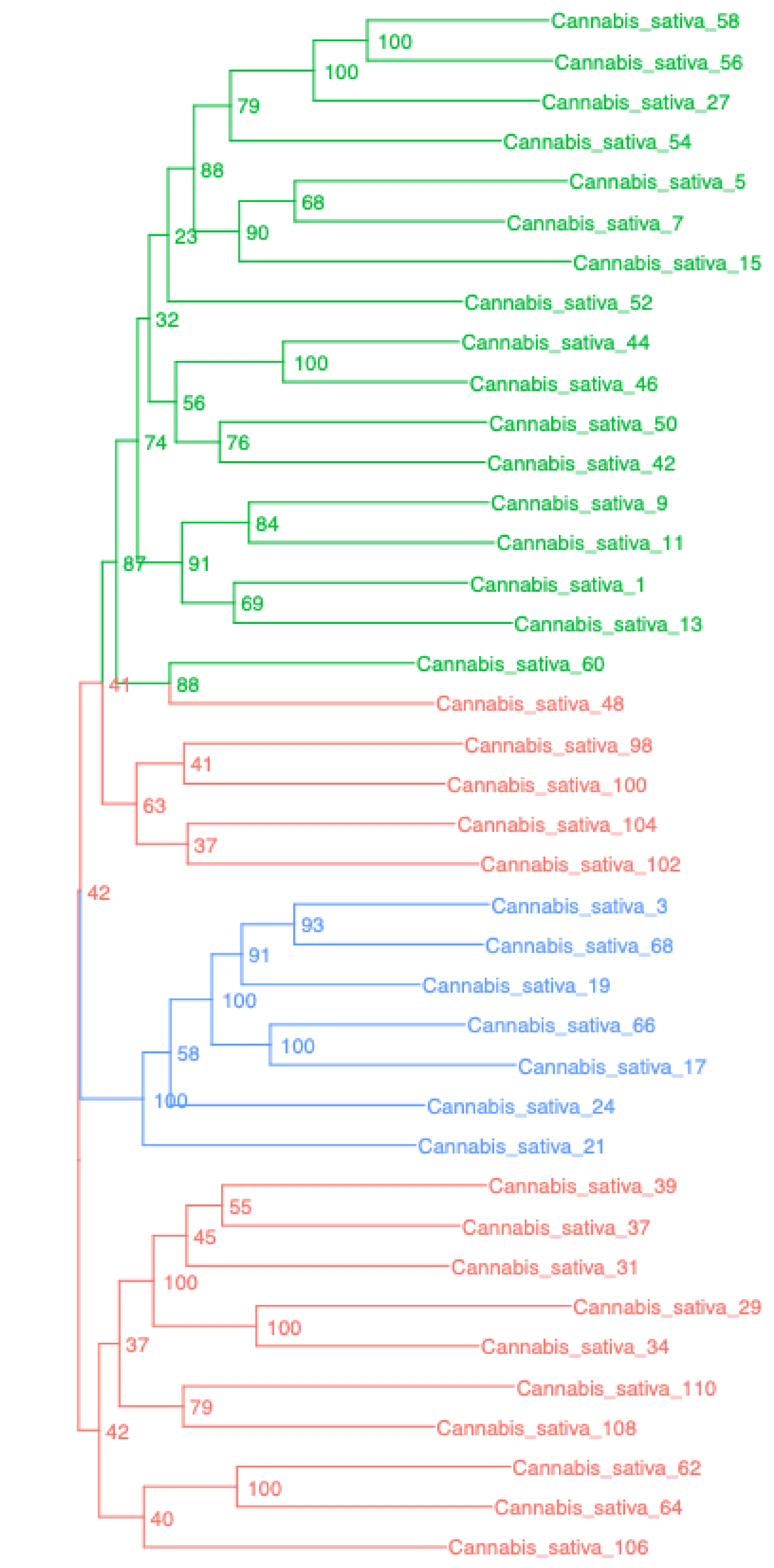


Cannabis – Cannabis_BUSCOsuper

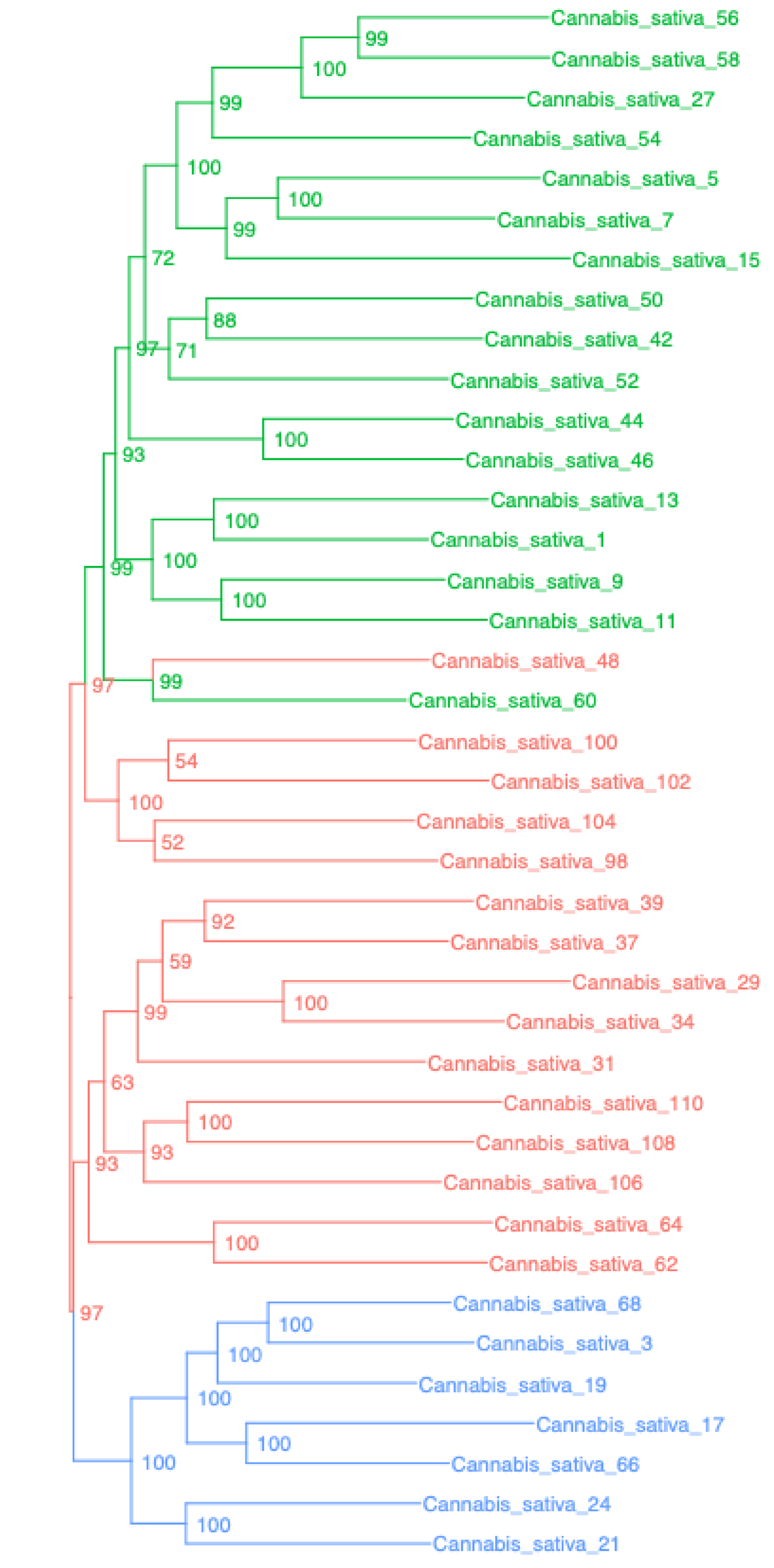


Cannabis – Cannabis_SCsuper

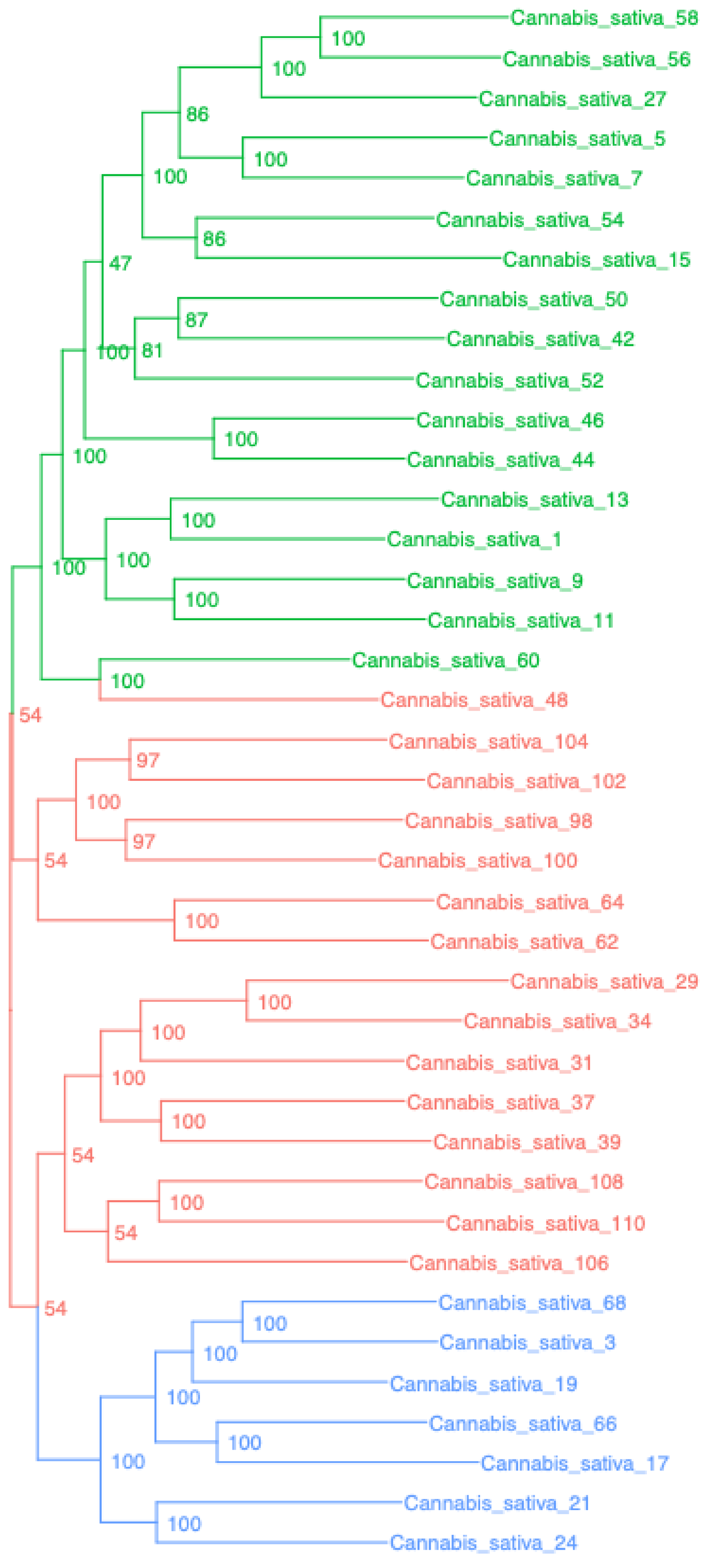


Cannabis – Cannabis_genome_thinned

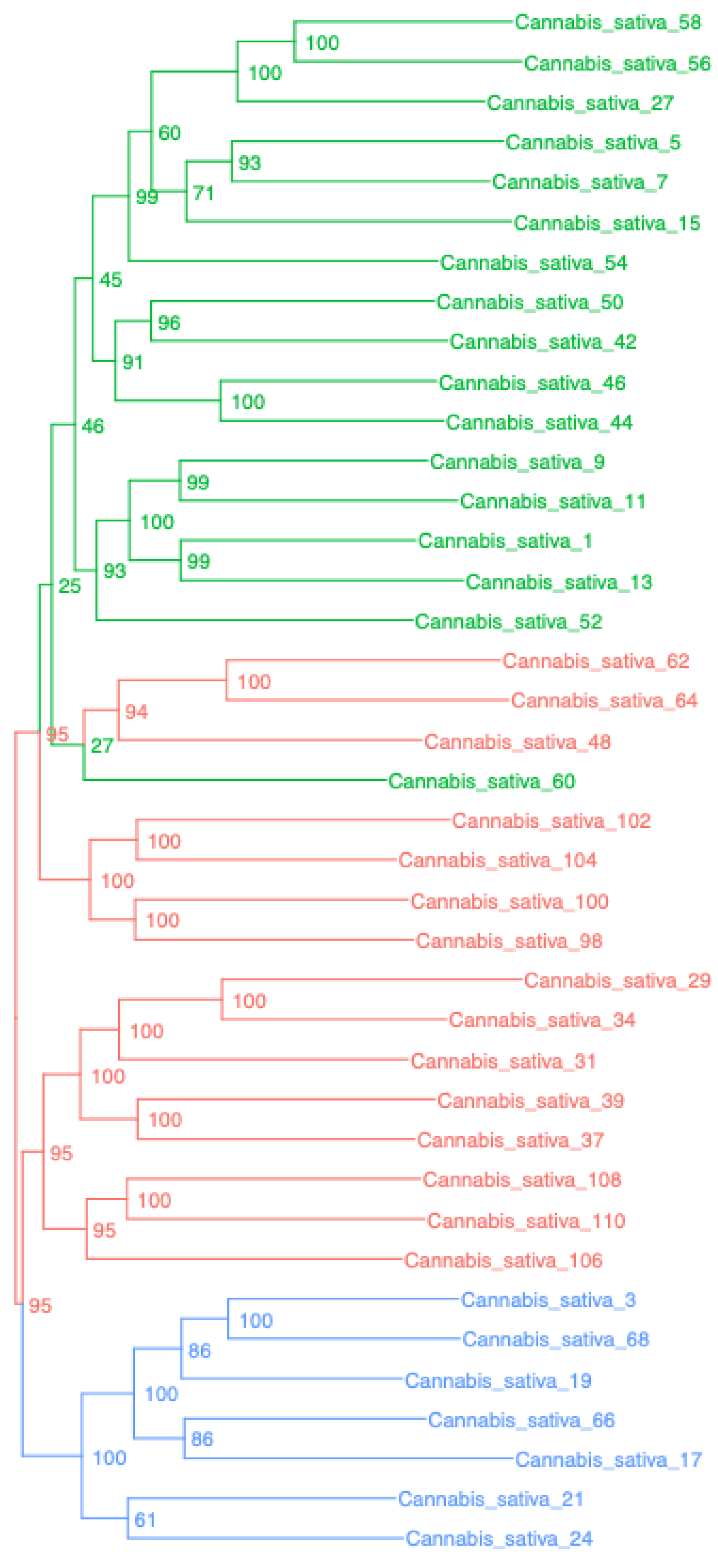

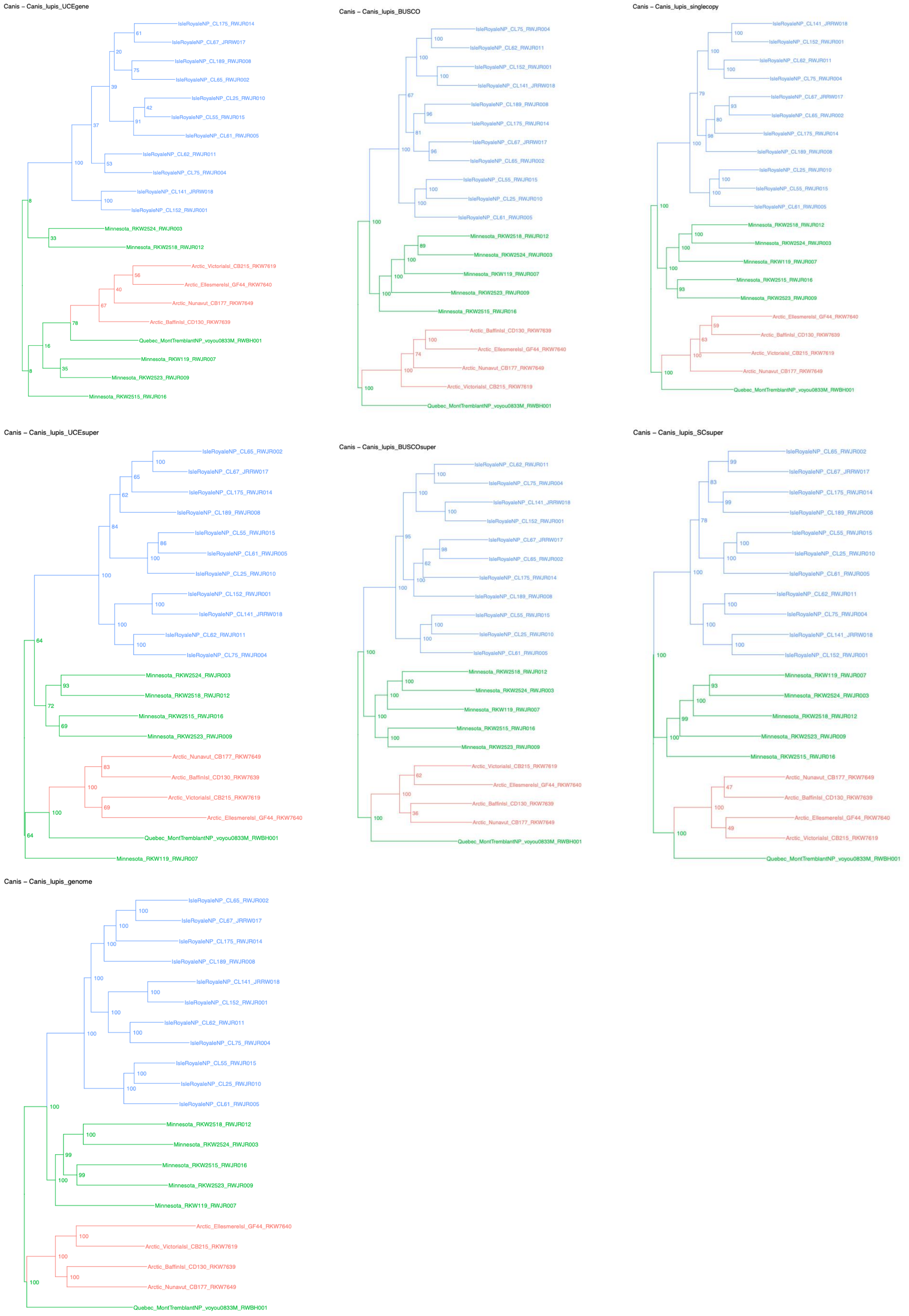
Canis – Canis_lupis_UCEgene
IsleRoyaleNP_CL175_RWJR014
IsleRoyaleNP_CL67_JRRW017
IsleRoyaleNP_CL189_RWJR008
IsleRoyaleNP_CL65_RWJR002
IsleRoyaleNP_CL25_RWJR010
IsleRoyaleNP_CL55_RWJR015
IsleRoyaleNP_CL61_RWJR005
IsleRoyaleNP_CL62_RWJR011
IsleRoyaleNP_CL75_RWJR004
IsleRoyaleNP_CL141_JRRW018
IsleRoyaleNP_CL152_RWJR001
Minnesota_RKW2524_RWJR003
Minnesota_RKW2518_RWJR012
Arctic_VictoriaIsl_CB215_RKW7619
Arctic_EllesmereIsl_GF44_RKW7640
Arctic_Nunavut_CB177_RKW7649
Arctic_BaffinIsl_CD130_RKW7639
Quebec_MontTremblantNP_voyou0833M_RWBH001
Minnesota_RKW119_RWJR007
Minnesota_RKW2523_RWJR009
Minnesota_RKW2515_RWJR016
Canis – Canis_lupis_BUSCO
IsleRoyaleNP_CL75_RWJR004
IsleRoyaleNP_CL62_RWJR011
IsleRoyaleNP_CL152_RWJR001
IsleRoyaleNP_CL141_JRRW018
IsleRoyaleNP_CL189_RWJR008
IsleRoyaleNP_CL175_RWJR014
IsleRoyaleNP_CL67_JRRW017
IsleRoyaleNP_CL65_RWJR002
IsleRoyaleNP_CL55_RWJR015
IsleRoyaleNP_CL25_RWJR010
IsleRoyaleNP_CL61_RWJR005
Minnesota_RKW2518_RWJR012
Minnesota_RKW2524_RWJR003
Minnesota_RKW119_RWJR007
Minnesota_RKW2523_RWJR009
Minnesota_RKW2515_RWJR016
Arctic_BaffinIsl_CD130_RKW7639
Arctic_EllesmereIsl_GF44_RKW7640
Arctic_Nunavut_CB177_RKW7649
Arctic_VictoriaIsl_CB215_RKW7619
Quebec_MontTremblantNP_voyou0833M_RWBH001
Canis – Canis_lupis_singlecopy
IsleRoyaleNP_CL141_JRRW018
IsleRoyaleNP_CL152_RWJR001
IsleRoyaleNP_CL62_RWJR011
IsleRoyaleNP_CL75_RWJR004
IsleRoyaleNP_CL67_JRRW017
IsleRoyaleNP_CL65_RWJR002
IsleRoyaleNP_CL175_RWJR014
IsleRoyaleNP_CL189_RWJR008
IsleRoyaleNP_CL25_RWJR010
IsleRoyaleNP_CL55_RWJR015
IsleRoyaleNP_CL61_RWJR005
Minnesota_RKW2518_RWJR012
Minnesota_RKW2524_RWJR003
Minnesota_RKW119_RWJR007
Minnesota_RKW2515_RWJR016
Minnesota_RKW2523_RWJR009
Arctic_EllesmereIsl_GF44_RKW7640
Arctic_BaffinIsl_CD130_RKW7639
Arctic_VictoriaIsl_CB215_RKW7619
Arctic_Nunavut_CB177_RKW7649
Quebec_MontTremblantNP_voyou0833M_RWBH001
Canis – Canis_lupis_UCEsuper
IsleRoyaleNP_CL65_RWJR002
IsleRoyaleNP_CL67_JRRW017
IsleRoyaleNP_CL175_RWJR014
IsleRoyaleNP_CL189_RWJR008
IsleRoyaleNP_CL55_RWJR015
IsleRoyaleNP_CL61_RWJR005
IsleRoyaleNP_CL25_RWJR010
IsleRoyaleNP_CL152_RWJR001
IsleRoyaleNP_CL141_JRRW018
IsleRoyaleNP_CL62_RWJR011
IsleRoyaleNP_CL75_RWJR004
Minnesota_RKW2524_RWJR003
Minnesota_RKW2518_RWJR012
Minnesota_RKW2515_RWJR016
Minnesota_RKW2523_RWJR009
Arctic_Nunavut_CB177_RKW7649
Arctic_BaffinIsl_CD130_RKW7639
Arctic_VictoriaIsl_CB215_RKW7619
Arctic_EllesmereIsl_GF44_RKW7640
Quebec_MontTremblantNP_voyou0833M_RWBH001
Minnesota_RKW119_RWJR007
Canis – Canis_lupis_BUSCOsuper
IsleRoyaleNP_CL62_RWJR011
IsleRoyaleNP_CL75_RWJR004
IsleRoyaleNP_CL141_JRRW018
IsleRoyaleNP_CL152_RWJR001
IsleRoyaleNP_CL67_JRRW017
IsleRoyaleNP_CL65_RWJR002
IsleRoyaleNP_CL175_RWJR014
IsleRoyaleNP_CL189_RWJR008
IsleRoyaleNP_CL55_RWJR015
IsleRoyaleNP_CL25_RWJR010
IsleRoyaleNP_CL61_RWJR005
Minnesota_RKW2518_RWJR012
Minnesota_RKW2524_RWJR003
Minnesota_RKW119_RWJR007
Minnesota_RKW2515_RWJR016
Minnesota_RKW2523_RWJR009
Arctic_VictoriaIsl_CB215_RKW7619
Arctic_EllesmereIsl_GF44_RKW7640
Arctic_BaffinIsl_CD130_RKW7639
Arctic_Nunavut_CB177_RKW7649
Quebec_MontTremblantNP_voyou0833M_RWBH001
Canis – Canis_lupis_SCsuper
IsleRoyaleNP_CL65_RWJR002
IsleRoyaleNP_CL67_JRRW017
IsleRoyaleNP_CL175_RWJR014
IsleRoyaleNP_CL189_RWJR008
IsleRoyaleNP_CL55_RWJR015
IsleRoyaleNP_CL25_RWJR010
IsleRoyaleNP_CL61_RWJR005
IsleRoyaleNP_CL62_RWJR011
IsleRoyaleNP_CL75_RWJR004
IsleRoyaleNP_CL141_JRRW018
IsleRoyaleNP_CL152_RWJR001
Minnesota_RKW119_RWJR007
Minnesota_RKW2524_RWJR003
Minnesota_RKW2518_RWJR012
Minnesota_RKW2523_RWJR009
Minnesota_RKW2515_RWJR016
Arctic_Nunavut_CB177_RKW7649
Arctic_BaffinIsl_CD130_RKW7639
Arctic_EllesmereIsl_GF44_RKW7640
Arctic_VictoriaIsl_CB215_RKW7619
Quebec_MontTremblantNP_voyou0833M_RWBH001
Canis – Canis_lupis_genome
IsleRoyaleNP_CL65_RWJR002
IsleRoyaleNP_CL67_JRRW017
IsleRoyaleNP_CL175_RWJR014
IsleRoyaleNP_CL189_RWJR008
IsleRoyaleNP_CL141_JRRW018
IsleRoyaleNP_CL152_RWJR001
IsleRoyaleNP_CL62_RWJR011
IsleRoyaleNP_CL75_RWJR004
IsleRoyaleNP_CL55_RWJR015
IsleRoyaleNP_CL25_RWJR010
IsleRoyaleNP_CL61_RWJR005
Minnesota_RKW2518_RWJR012
Minnesota_RKW2524_RWJR003
Minnesota_RKW2515_RWJR016
Minnesota_RKW2523_RWJR009
Minnesota_RKW119_RWJR007
Arctic_EllesmereIsl_GF44_RKW7640
Arctic_VictoriaIsl_CB215_RKW7619
Arctic_BaffinIsl_CD130_RKW7639
Arctic_Nunavut_CB177_RKW7649
Quebec_MontTremblantNP_voyou0833M_RWBH001

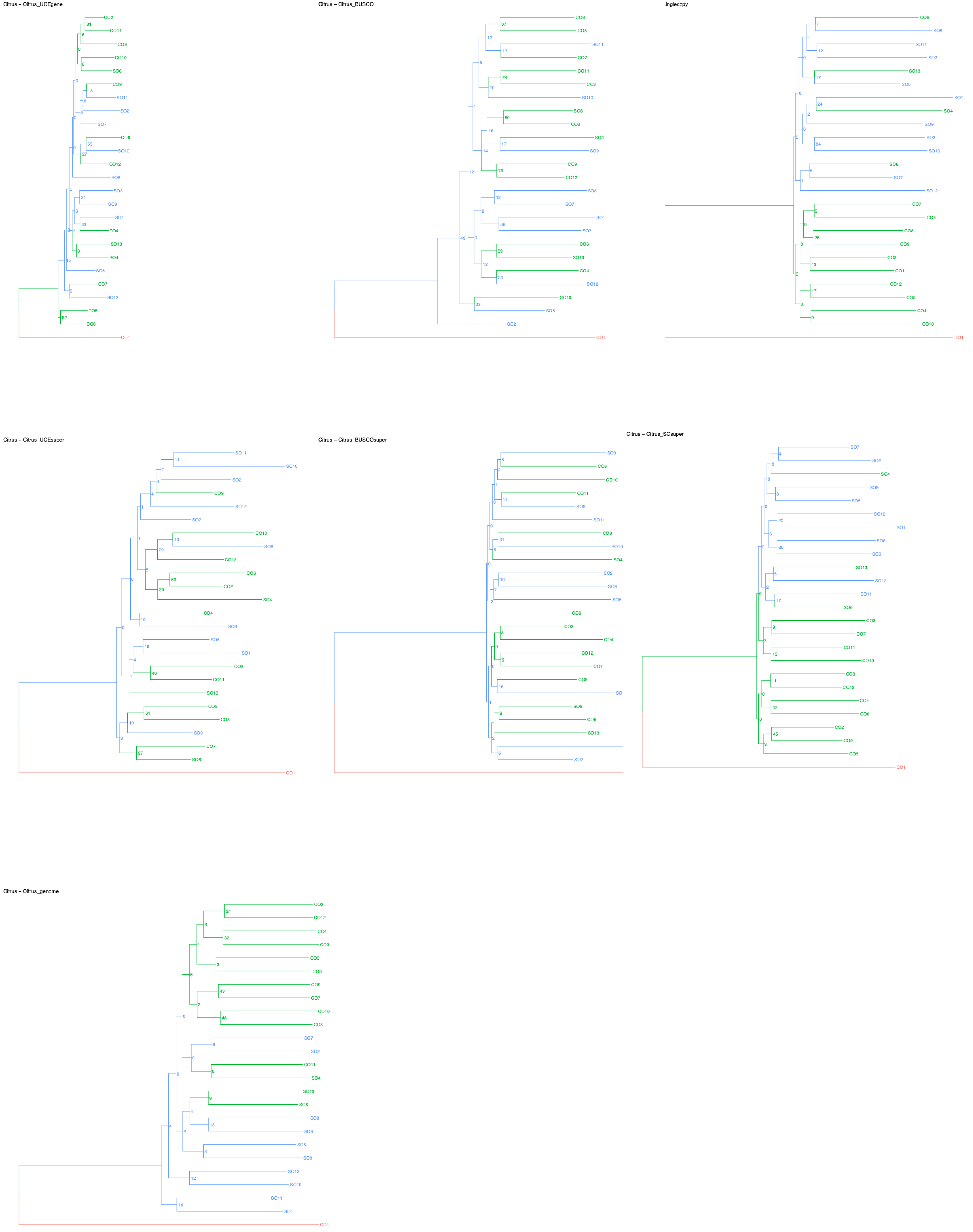
Citrus – Citrus_UCEgene
CO2
CO11
CO3
CO10
SO6
CO9
SO11
SO2
SO7
CO6
SO10
CO12
SO8
SO3
SO9
SO1
CO4
SO13
SO4
SO5
CO7
SO12
CO5
CO8
CO1
Citrus – Citrus_BUSCO
CO8
CO5
SO11
CO7
CO11
CO3
SO10
SO6
CO2
SO4
SO9
CO9
CO12
SO8
SO7
SO1
SO3
CO6
SO13
CO4
SO12
CO10
SO5
SO2
CO1
singlecopy
CO6
SO8
SO11
SO2
SO13
SO5
SO4
SO9
SO3
SO10
SO6
SO7
SO12
CO7
CO3
CO8
CO9
CO2
CO11
CO12
CO5
CO4
CO10
Citrus – Citrus_UCEsuper
SO11
SO10
SO2
CO9
SO12
SO7
CO10
SO8
CO12
CO6
CO2
SO4
CO4
SO3
SO5
SO1
CO3
CO11
SO13
CO5
CO8
SO9
CO7
SO6
CO1
Citrus – Citrus_BUSCOsuper
SO3
CO6
CO10
CO11
SO5
SO11
CO3
SO10
SO4
SO2
SO9
SO8
CO9
CO2
CO4
CO12
CO7
CO8
SO6
CO5
SO13
SO7
Citrus – Citrus_SCsuper
SO7
SO2
SO4
SO9
SO5
SO10
SO1
SO8
SO3
SO13
SO12
SO11
SO6
CO3
CO7
CO11
CO10
CO8
CO12
CO4
CO6
CO2
CO9
CO5
CO1
Citrus – Citrus_genome
CO2
CO12
CO4
CO3
CO5
CO6
CO9
CO7
CO10
CO8
SO7
SO2
CO11
SO4
SO13
SO6
SO8
SO3
SO5
SO9
SO12
SO10
SO11
SO1
CO1

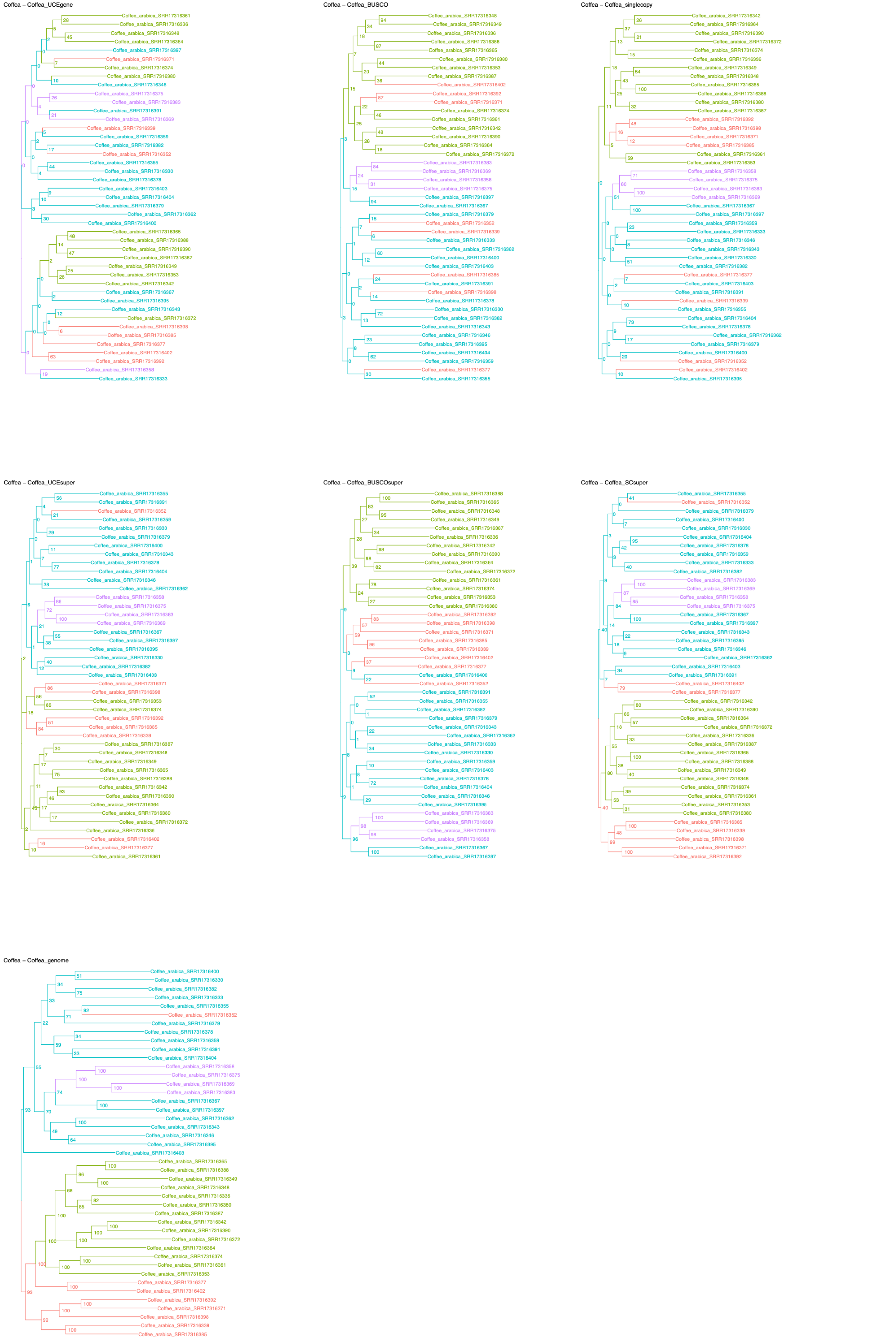

Coffea – Coffea_UCEgene
Coffea – Coffea_BUSCO
Coffea – Coffea_singlecopy
Coffea – Coffea_UCEsuper
Coffea – Coffea_BUSCOsuper
Coffea – Coffea_SCsuper
Coffea – Coffea_genome

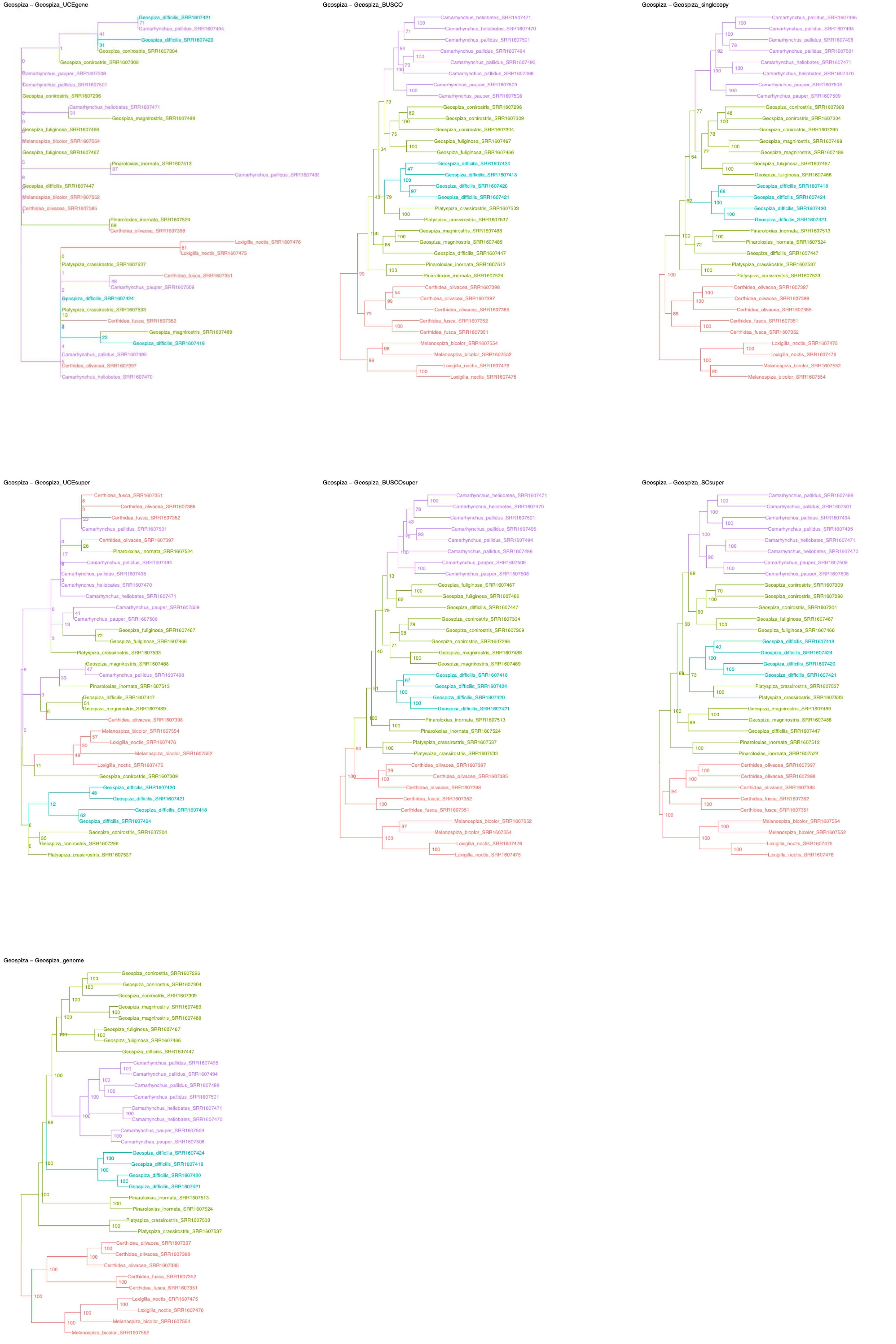
Geospiza – Geospiza_UCEgene
Geospiza – Geospiza_BUSCO
Geospiza – Geospiza_singlecopy
Geospiza – Geospiza_UCEsuper
Geospiza – Geospiza_BUSCOsuper
Geospiza – Geospiza_SCsuper
Geospiza – Geospiza_genome

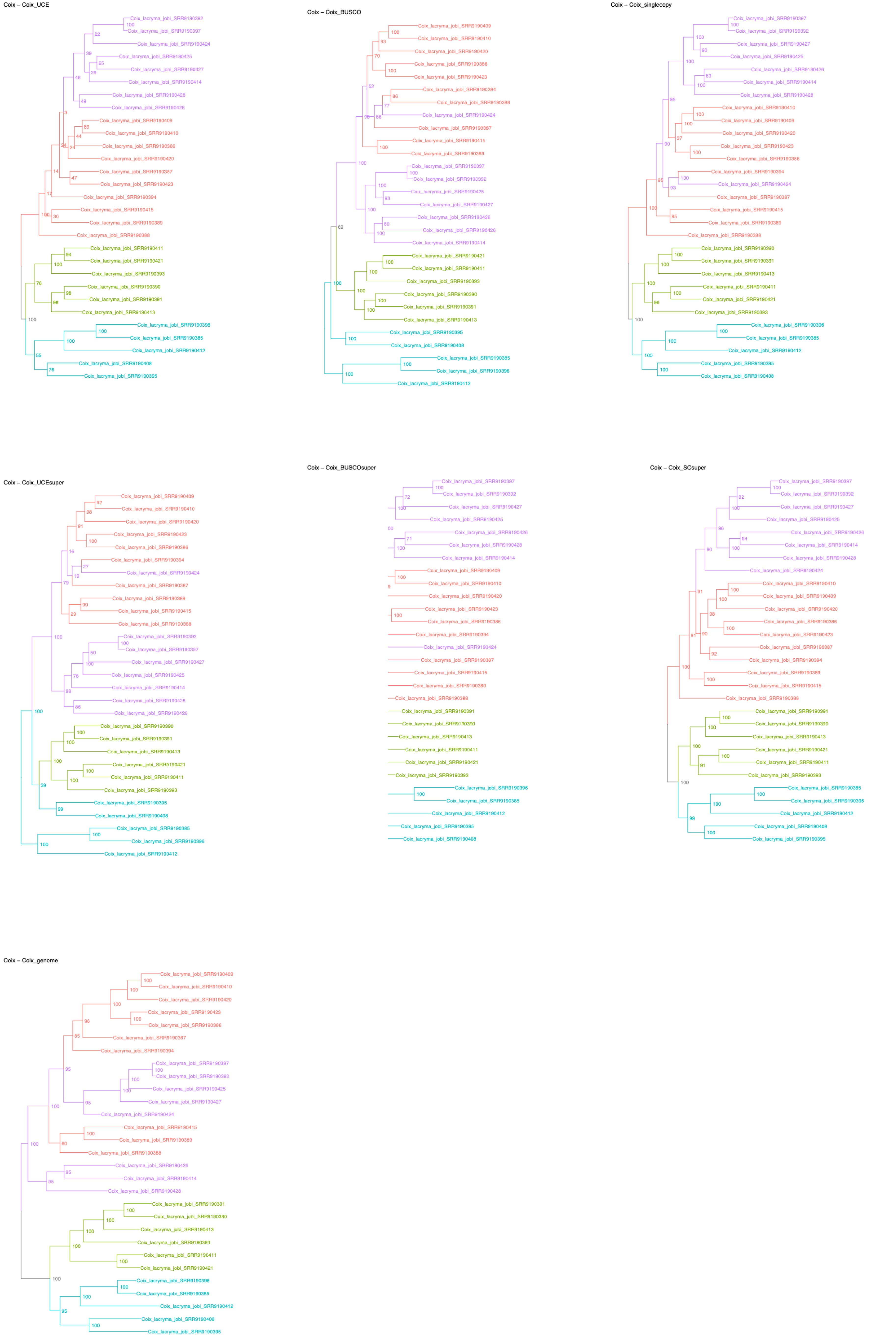
Coix – Coix_UCE
Coix_lacryma_jobi_SRR9190392
Coix_lacryma_jobi_SRR9190397
Coix_lacryma_jobi_SRR9190424
Coix_lacryma_jobi_SRR9190425
Coix_lacryma_jobi_SRR9190427
Coix_lacryma_jobi_SRR9190414
Coix_lacryma_jobi_SRR9190428
Coix_lacryma_jobi_SRR9190426
Coix_lacryma_jobi_SRR9190409
Coix_lacryma_jobi_SRR9190410
Coix_lacryma_jobi_SRR9190386
Coix_lacryma_jobi_SRR9190420
Coix_lacryma_jobi_SRR9190387
Coix_lacryma_jobi_SRR9190423
Coix_lacryma_jobi_SRR9190394
Coix_lacryma_jobi_SRR9190415
Coix_lacryma_jobi_SRR9190389
Coix_lacryma_jobi_SRR9190388
Coix_lacryma_jobi_SRR9190411
Coix_lacryma_jobi_SRR9190421
Coix_lacryma_jobi_SRR9190393
Coix_lacryma_jobi_SRR9190390
Coix_lacryma_jobi_SRR9190391
Coix_lacryma_jobi_SRR9190413
Coix_lacryma_jobi_SRR9190396
Coix_lacryma_jobi_SRR9190385
Coix_lacryma_jobi_SRR9190412
Coix_lacryma_jobi_SRR9190408
Coix_lacryma_jobi_SRR9190395
Coix – Coix_BUSCO
Coix_lacryma_jobi_SRR9190409
Coix_lacryma_jobi_SRR9190410
Coix_lacryma_jobi_SRR9190420
Coix_lacryma_jobi_SRR9190386
Coix_lacryma_jobi_SRR9190423
Coix_lacryma_jobi_SRR9190394
Coix_lacryma_jobi_SRR9190388
Coix_lacryma_jobi_SRR9190424
Coix_lacryma_jobi_SRR9190387
Coix_lacryma_jobi_SRR9190415
Coix_lacryma_jobi_SRR9190389
Coix_lacryma_jobi_SRR9190397
Coix_lacryma_jobi_SRR9190392
Coix_lacryma_jobi_SRR9190425
Coix_lacryma_jobi_SRR9190427
Coix_lacryma_jobi_SRR9190428
Coix_lacryma_jobi_SRR9190426
Coix_lacryma_jobi_SRR9190414
Coix_lacryma_jobi_SRR9190421
Coix_lacryma_jobi_SRR9190411
Coix_lacryma_jobi_SRR9190393
Coix_lacryma_jobi_SRR9190390
Coix_lacryma_jobi_SRR9190391
Coix_lacryma_jobi_SRR9190413
Coix_lacryma_jobi_SRR9190395
Coix_lacryma_jobi_SRR9190408
Coix_lacryma_jobi_SRR9190385
Coix_lacryma_jobi_SRR9190396
Coix_lacryma_jobi_SRR9190412
Coix – Coix_singlecopy
Coix_lacryma_jobi_SRR9190397
Coix_lacryma_jobi_SRR9190392
Coix_lacryma_jobi_SRR9190427
Coix_lacryma_jobi_SRR9190425
Coix_lacryma_jobi_SRR9190426
Coix_lacryma_jobi_SRR9190414
Coix_lacryma_jobi_SRR9190428
Coix_lacryma_jobi_SRR9190410
Coix_lacryma_jobi_SRR9190409
Coix_lacryma_jobi_SRR9190420
Coix_lacryma_jobi_SRR9190423
Coix_lacryma_jobi_SRR9190386
Coix_lacryma_jobi_SRR9190394
Coix_lacryma_jobi_SRR9190424
Coix_lacryma_jobi_SRR9190387
Coix_lacryma_jobi_SRR9190415
Coix_lacryma_jobi_SRR9190389
Coix_lacryma_jobi_SRR9190388
Coix_lacryma_jobi_SRR9190390
Coix_lacryma_jobi_SRR9190391
Coix_lacryma_jobi_SRR9190413
Coix_lacryma_jobi_SRR9190411
Coix_lacryma_jobi_SRR9190421
Coix_lacryma_jobi_SRR9190393
Coix_lacryma_jobi_SRR9190396
Coix_lacryma_jobi_SRR9190385
Coix_lacryma_jobi_SRR9190412
Coix_lacryma_jobi_SRR9190395
Coix_lacryma_jobi_SRR9190408
Coix – Coix_UCEsuper
Coix_lacryma_jobi_SRR9190409
Coix_lacryma_jobi_SRR9190410
Coix_lacryma_jobi_SRR9190420
Coix_lacryma_jobi_SRR9190423
Coix_lacryma_jobi_SRR9190386
Coix_lacryma_jobi_SRR9190394
Coix_lacryma_jobi_SRR9190424
Coix_lacryma_jobi_SRR9190387
Coix_lacryma_jobi_SRR9190389
Coix_lacryma_jobi_SRR9190415
Coix_lacryma_jobi_SRR9190388
Coix_lacryma_jobi_SRR9190392
Coix_lacryma_jobi_SRR9190397
Coix_lacryma_jobi_SRR9190427
Coix_lacryma_jobi_SRR9190425
Coix_lacryma_jobi_SRR9190414
Coix_lacryma_jobi_SRR9190428
Coix_lacryma_jobi_SRR9190426
Coix_lacryma_jobi_SRR9190390
Coix_lacryma_jobi_SRR9190391
Coix_lacryma_jobi_SRR9190413
Coix_lacryma_jobi_SRR9190421
Coix_lacryma_jobi_SRR9190411
Coix_lacryma_jobi_SRR9190393
Coix_lacryma_jobi_SRR9190395
Coix_lacryma_jobi_SRR9190408
Coix_lacryma_jobi_SRR9190385
Coix_lacryma_jobi_SRR9190396
Coix_lacryma_jobi_SRR9190412
Coix – Coix_BUSCOsuper
Coix_lacryma_jobi_SRR9190397
Coix_lacryma_jobi_SRR9190392
Coix_lacryma_jobi_SRR9190427
Coix_lacryma_jobi_SRR9190425
Coix_lacryma_jobi_SRR9190426
Coix_lacryma_jobi_SRR9190428
Coix_lacryma_jobi_SRR9190414
Coix_lacryma_jobi_SRR9190409
Coix_lacryma_jobi_SRR9190410
Coix_lacryma_jobi_SRR9190420
Coix_lacryma_jobi_SRR9190423
Coix_lacryma_jobi_SRR9190386
Coix_lacryma_jobi_SRR9190394
Coix_lacryma_jobi_SRR9190424
Coix_lacryma_jobi_SRR9190387
Coix_lacryma_jobi_SRR9190415
Coix_lacryma_jobi_SRR9190389
Coix_lacryma_jobi_SRR9190388
Coix_lacryma_jobi_SRR9190391
Coix_lacryma_jobi_SRR9190390
Coix_lacryma_jobi_SRR9190413
Coix_lacryma_jobi_SRR9190411
Coix_lacryma_jobi_SRR9190421
Coix_lacryma_jobi_SRR9190393
Coix_lacryma_jobi_SRR9190396
Coix_lacryma_jobi_SRR9190385
Coix_lacryma_jobi_SRR9190412
Coix_lacryma_jobi_SRR9190395
Coix_lacryma_jobi_SRR9190408
Coix – Coix_SCsuper
Coix_lacryma_jobi_SRR9190397
Coix_lacryma_jobi_SRR9190392
Coix_lacryma_jobi_SRR9190427
Coix_lacryma_jobi_SRR9190425
Coix_lacryma_jobi_SRR9190426
Coix_lacryma_jobi_SRR9190414
Coix_lacryma_jobi_SRR9190428
Coix_lacryma_jobi_SRR9190424
Coix_lacryma_jobi_SRR9190410
Coix_lacryma_jobi_SRR9190409
Coix_lacryma_jobi_SRR9190420
Coix_lacryma_jobi_SRR9190386
Coix_lacryma_jobi_SRR9190423
Coix_lacryma_jobi_SRR9190387
Coix_lacryma_jobi_SRR9190394
Coix_lacryma_jobi_SRR9190389
Coix_lacryma_jobi_SRR9190415
Coix_lacryma_jobi_SRR9190388
Coix_lacryma_jobi_SRR9190391
Coix_lacryma_jobi_SRR9190390
Coix_lacryma_jobi_SRR9190413
Coix_lacryma_jobi_SRR9190421
Coix_lacryma_jobi_SRR9190411
Coix_lacryma_jobi_SRR9190393
Coix_lacryma_jobi_SRR9190385
Coix_lacryma_jobi_SRR9190396
Coix_lacryma_jobi_SRR9190412
Coix_lacryma_jobi_SRR9190408
Coix_lacryma_jobi_SRR9190395
Coix – Coix_genome
Coix_lacryma_jobi_SRR9190409
Coix_lacryma_jobi_SRR9190410
Coix_lacryma_jobi_SRR9190420
Coix_lacryma_jobi_SRR9190423
Coix_lacryma_jobi_SRR9190386
Coix_lacryma_jobi_SRR9190387
Coix_lacryma_jobi_SRR9190394
Coix_lacryma_jobi_SRR9190397
Coix_lacryma_jobi_SRR9190392
Coix_lacryma_jobi_SRR9190425
Coix_lacryma_jobi_SRR9190427
Coix_lacryma_jobi_SRR9190424
Coix_lacryma_jobi_SRR9190415
Coix_lacryma_jobi_SRR9190389
Coix_lacryma_jobi_SRR9190388
Coix_lacryma_jobi_SRR9190426
Coix_lacryma_jobi_SRR9190414
Coix_lacryma_jobi_SRR9190428
Coix_lacryma_jobi_SRR9190391
Coix_lacryma_jobi_SRR9190390
Coix_lacryma_jobi_SRR9190413
Coix_lacryma_jobi_SRR9190393
Coix_lacryma_jobi_SRR9190411
Coix_lacryma_jobi_SRR9190421
Coix_lacryma_jobi_SRR9190396
Coix_lacryma_jobi_SRR9190385
Coix_lacryma_jobi_SRR9190412
Coix_lacryma_jobi_SRR9190408
Coix_lacryma_jobi_SRR9190395

Costus – Costus_UCEgene

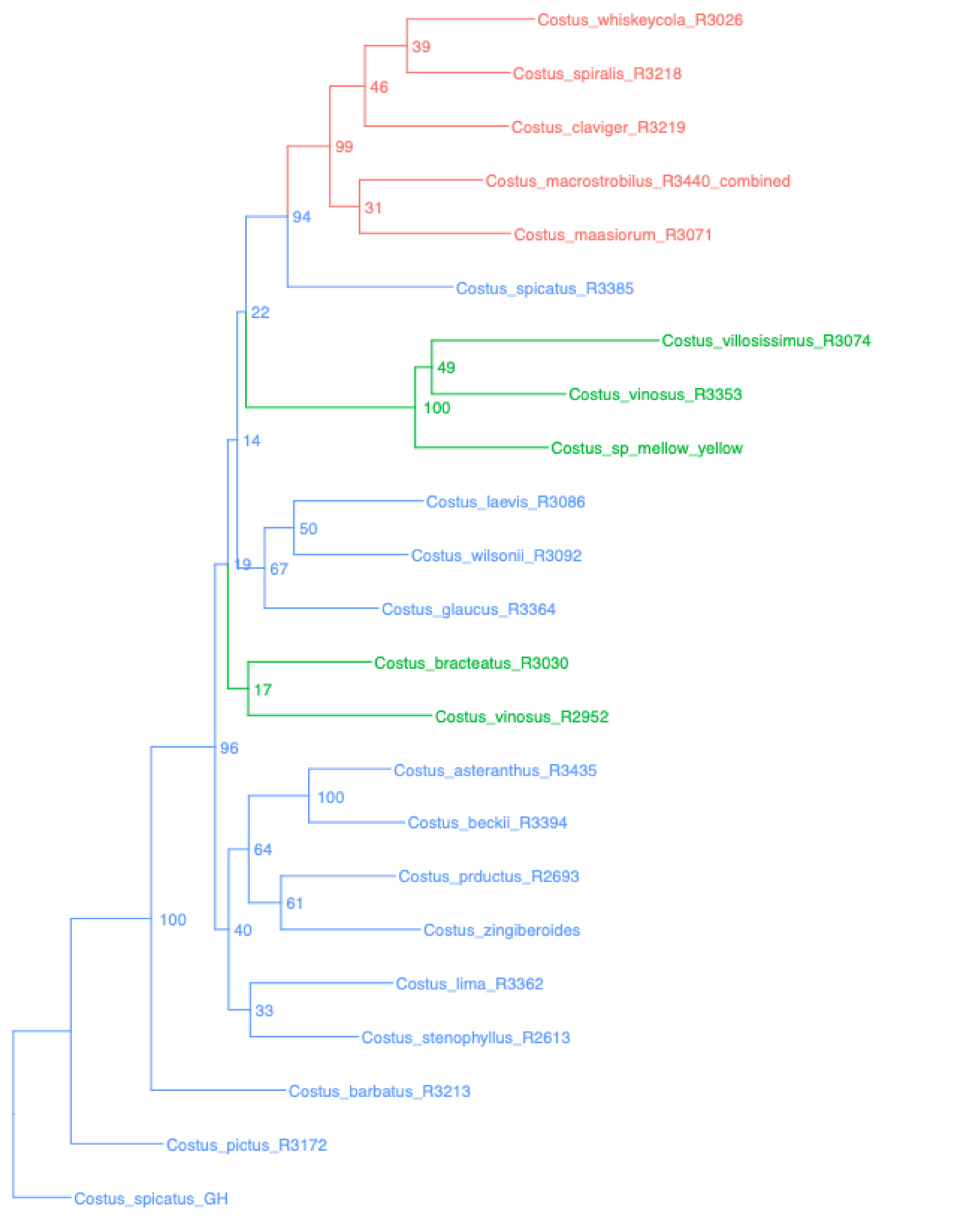


Costus – Costus_BUSCO

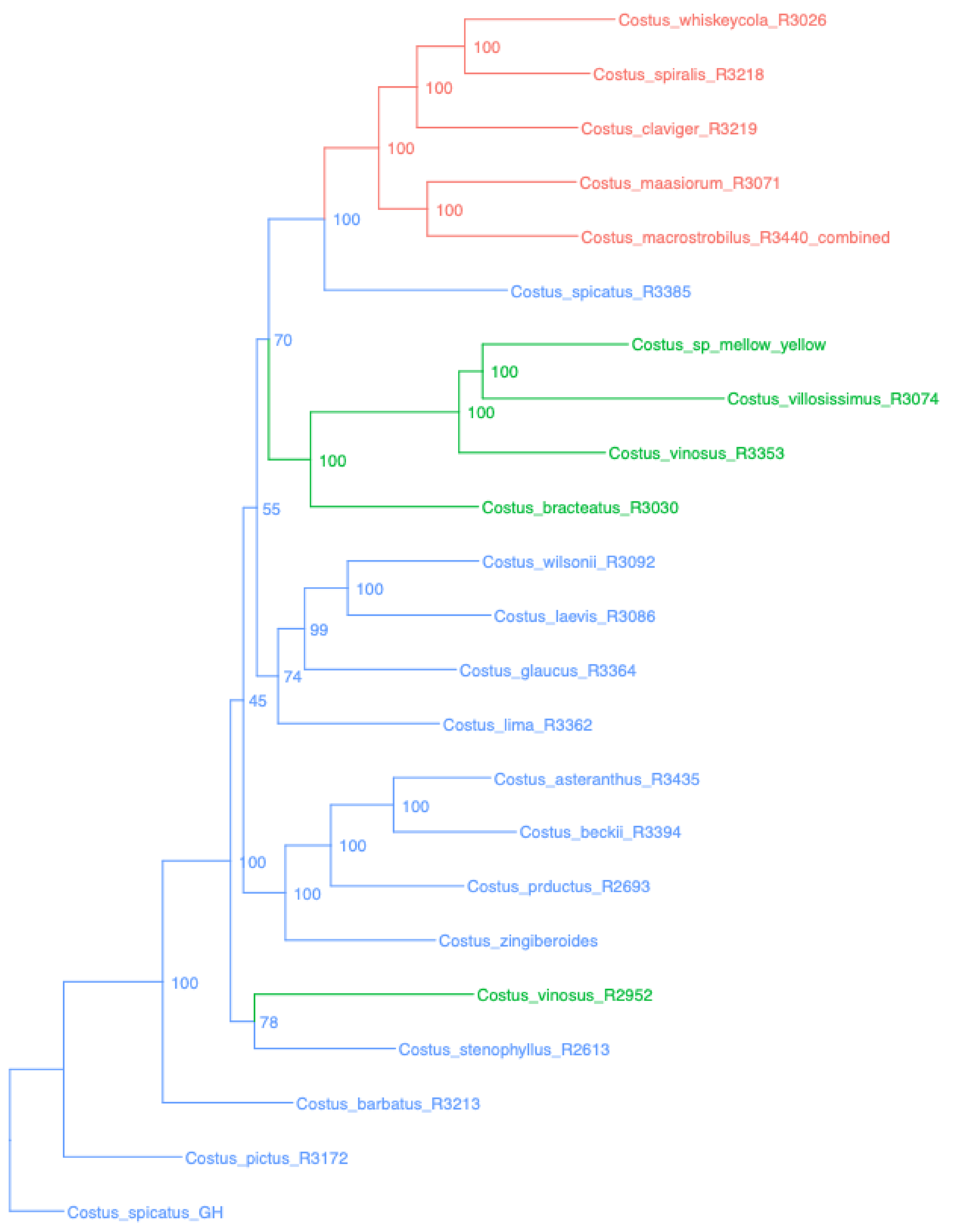


Costus – Costus_singlecopy

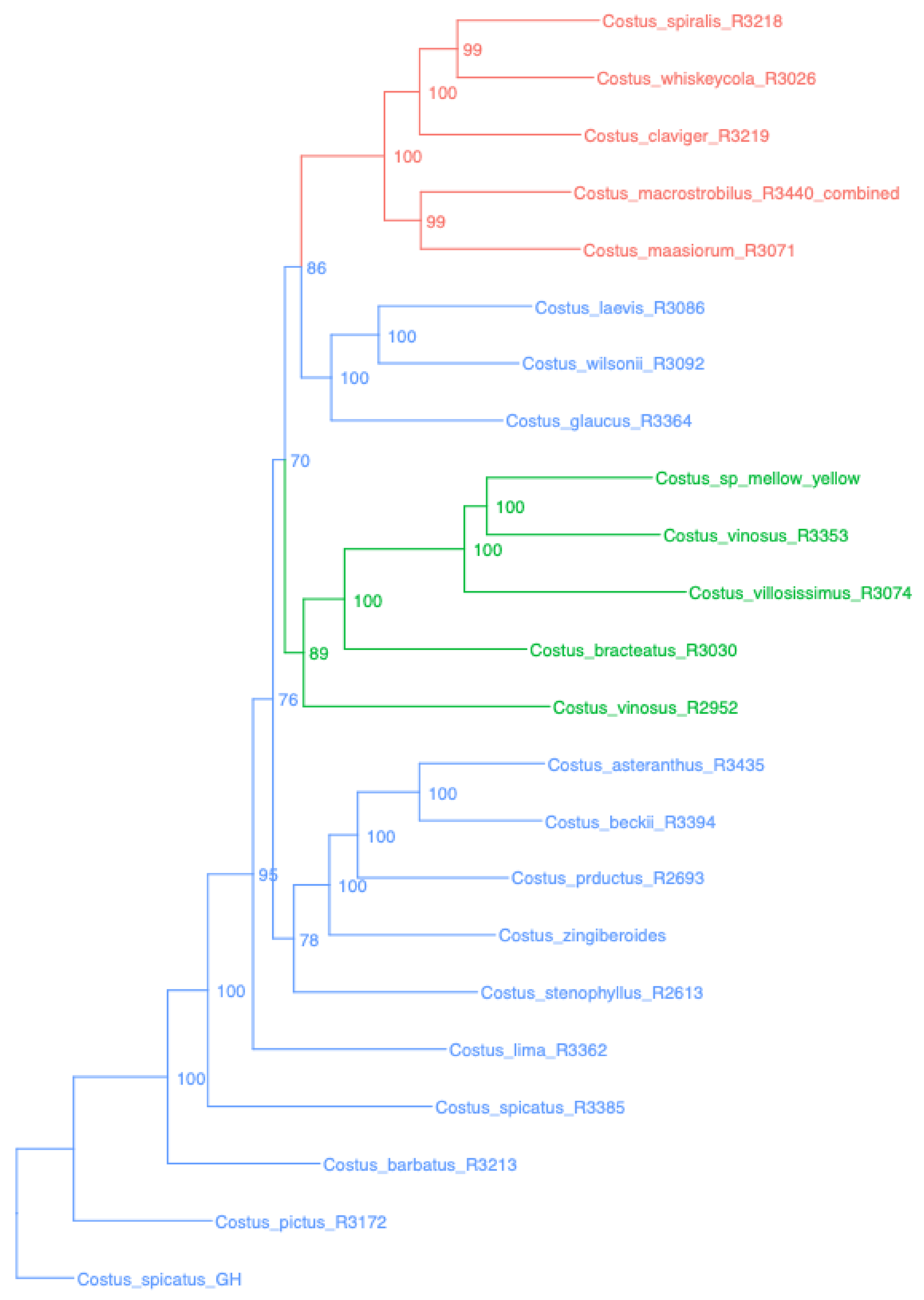


Costus – Costus_UCEsuper

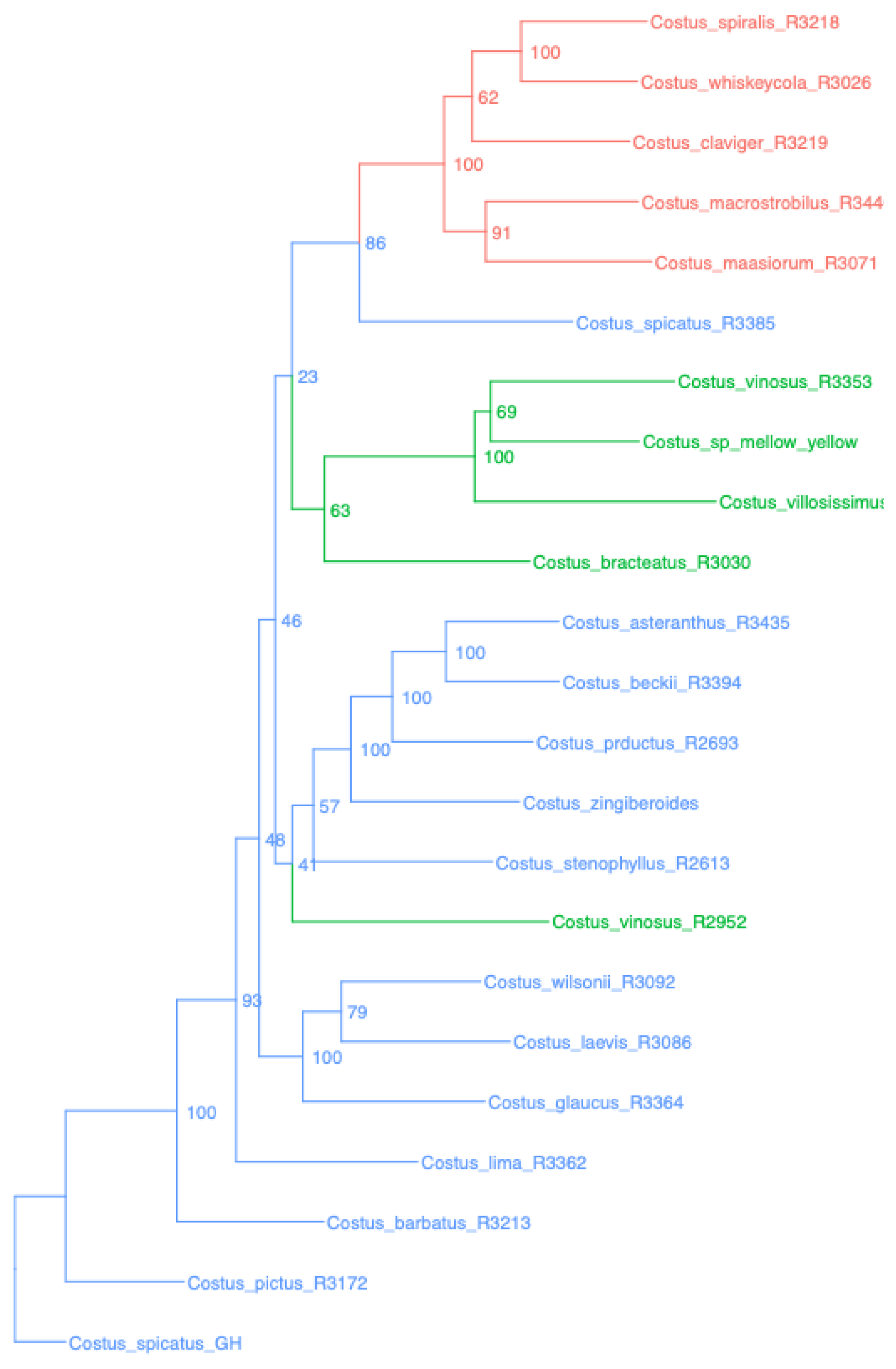


Costus – Costus_BUSCOsuper

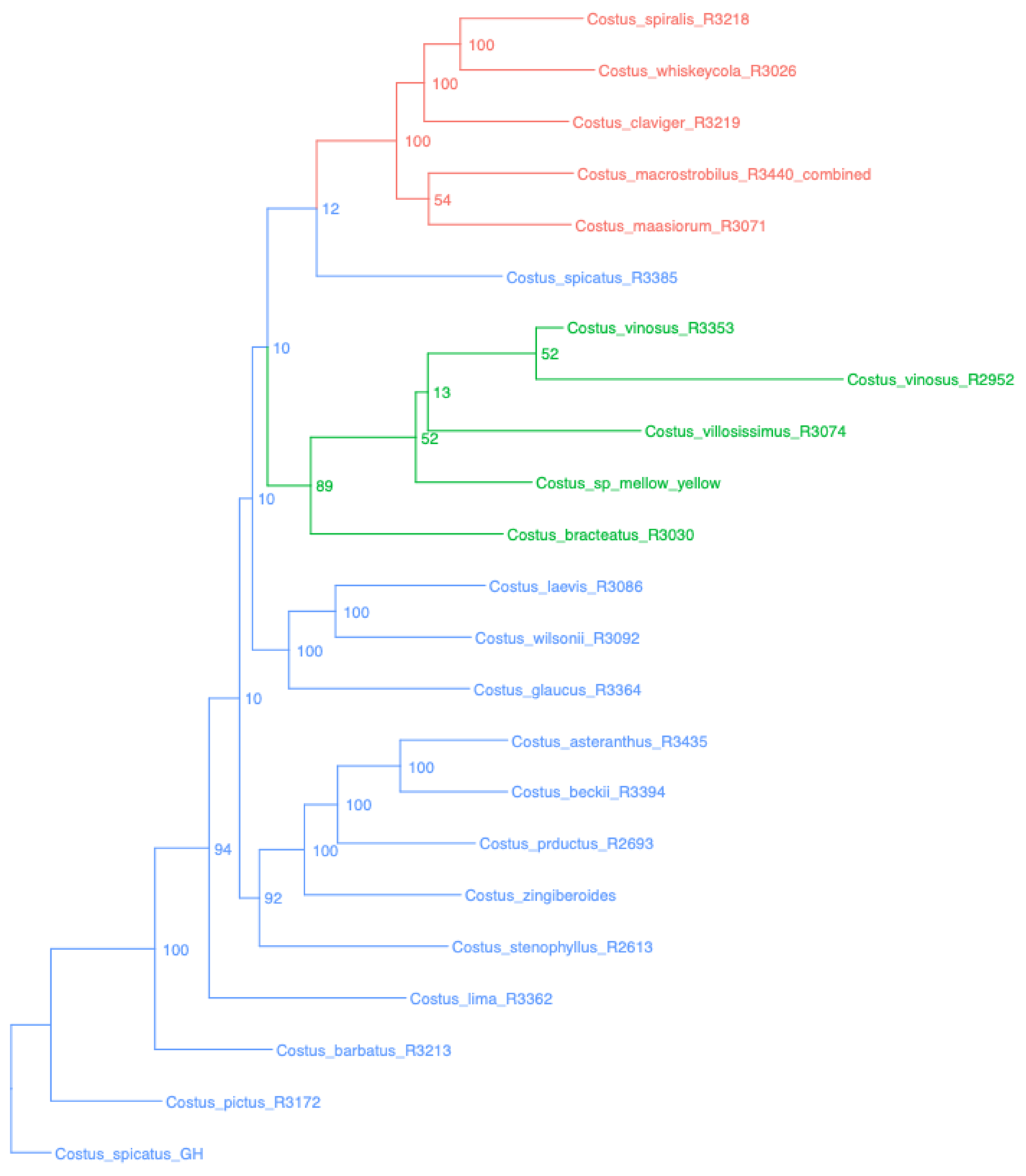


Costus – Costus_SCsuper

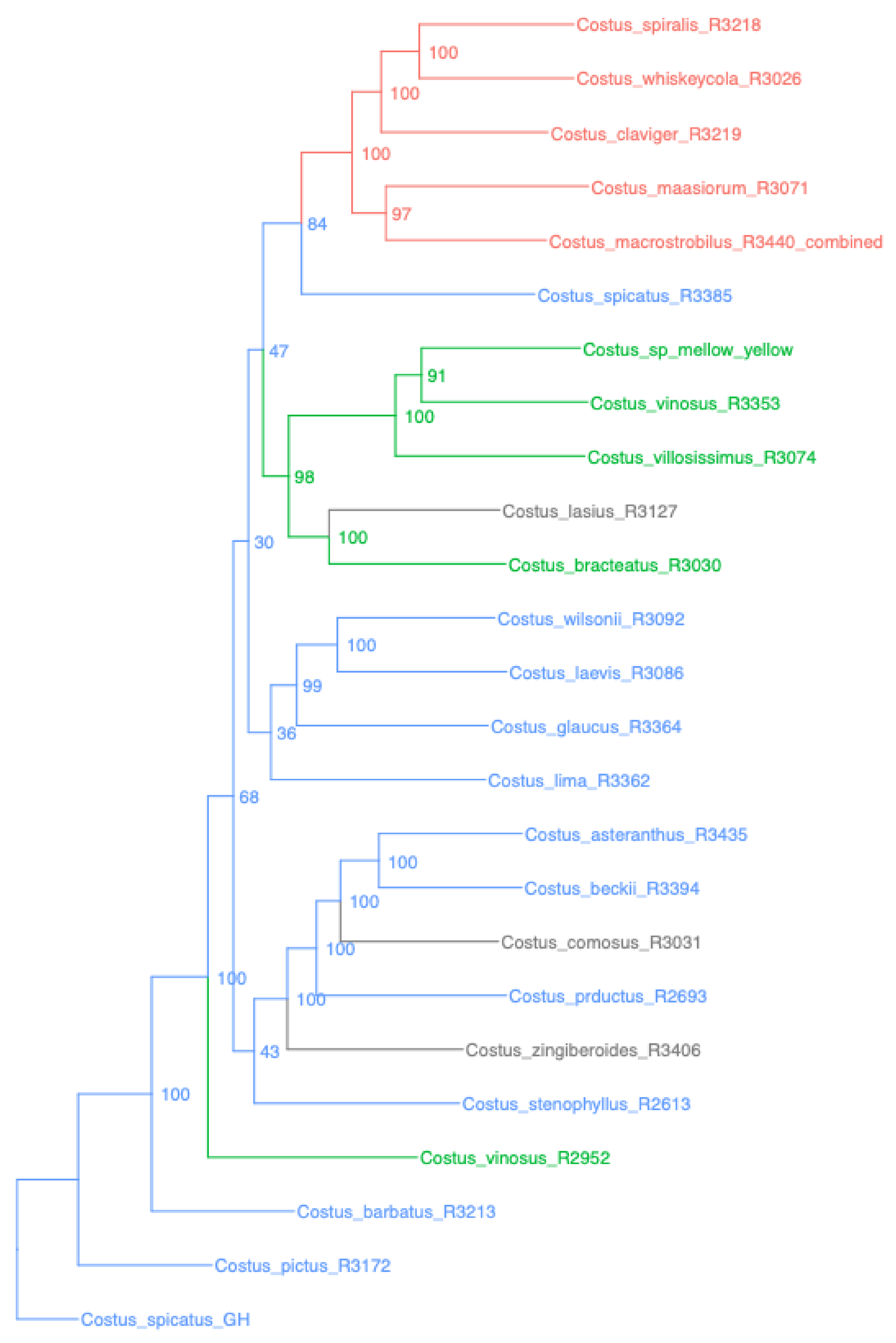


Costus – Costus_genome

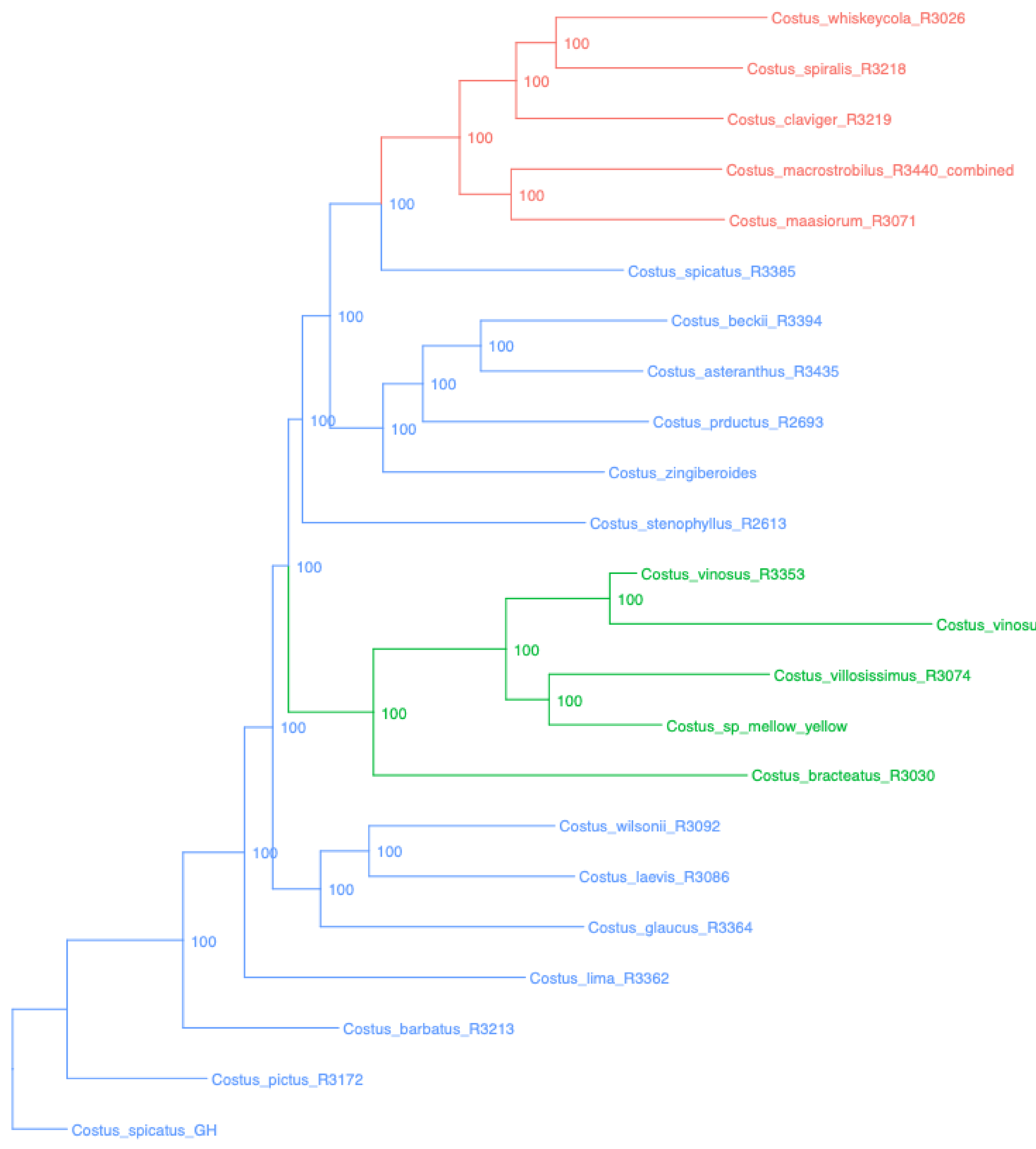

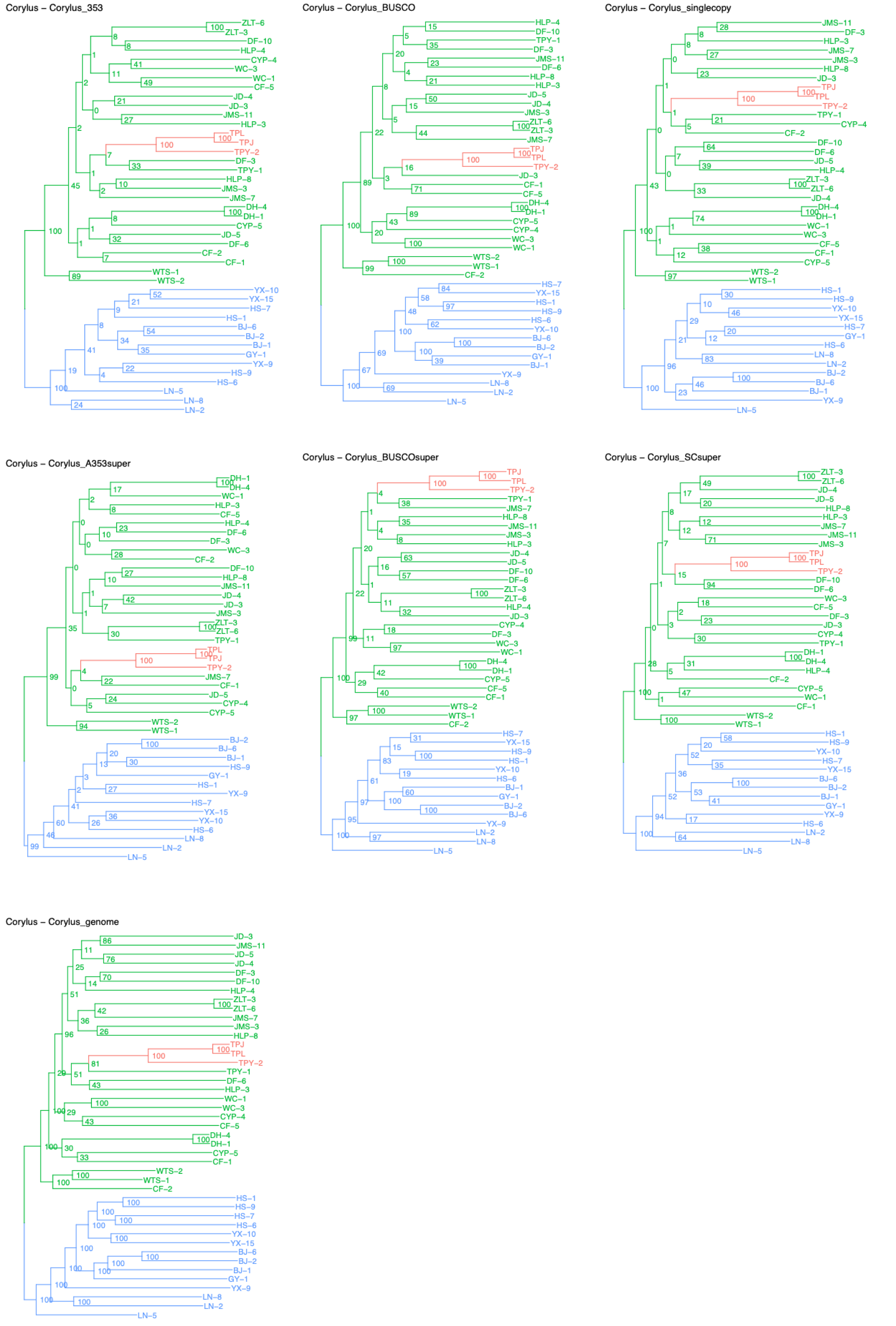
Corylus – Corylus_353
Corylus – Corylus_BUSCO
Corylus – Corylus_singlecopy
Corylus – Corylus_A353super
Corylus – Corylus_BUSCOsuper
Corylus – Corylus_SCsuper
Corylus – Corylus_genome

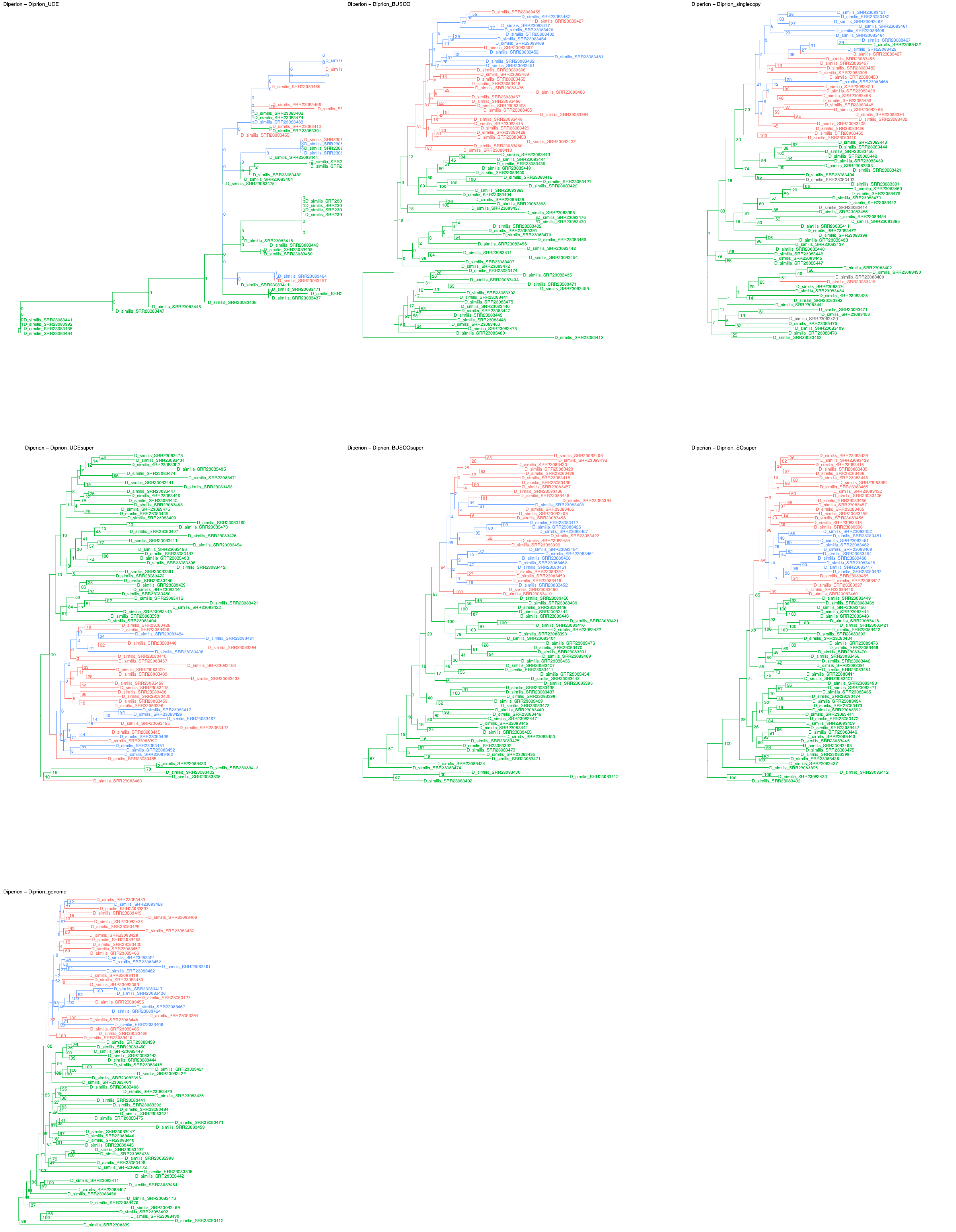
Diperion – Diprion_UCE
Diperion – Diprion_BUSCO
Diperion – Diprion_singlecopy
Diperion – Diprion_UCEsuper
Diperion – Diprion_BUSCOsuper
Diperion – Diprion_SCsuper
Diperion – Diprion_genome

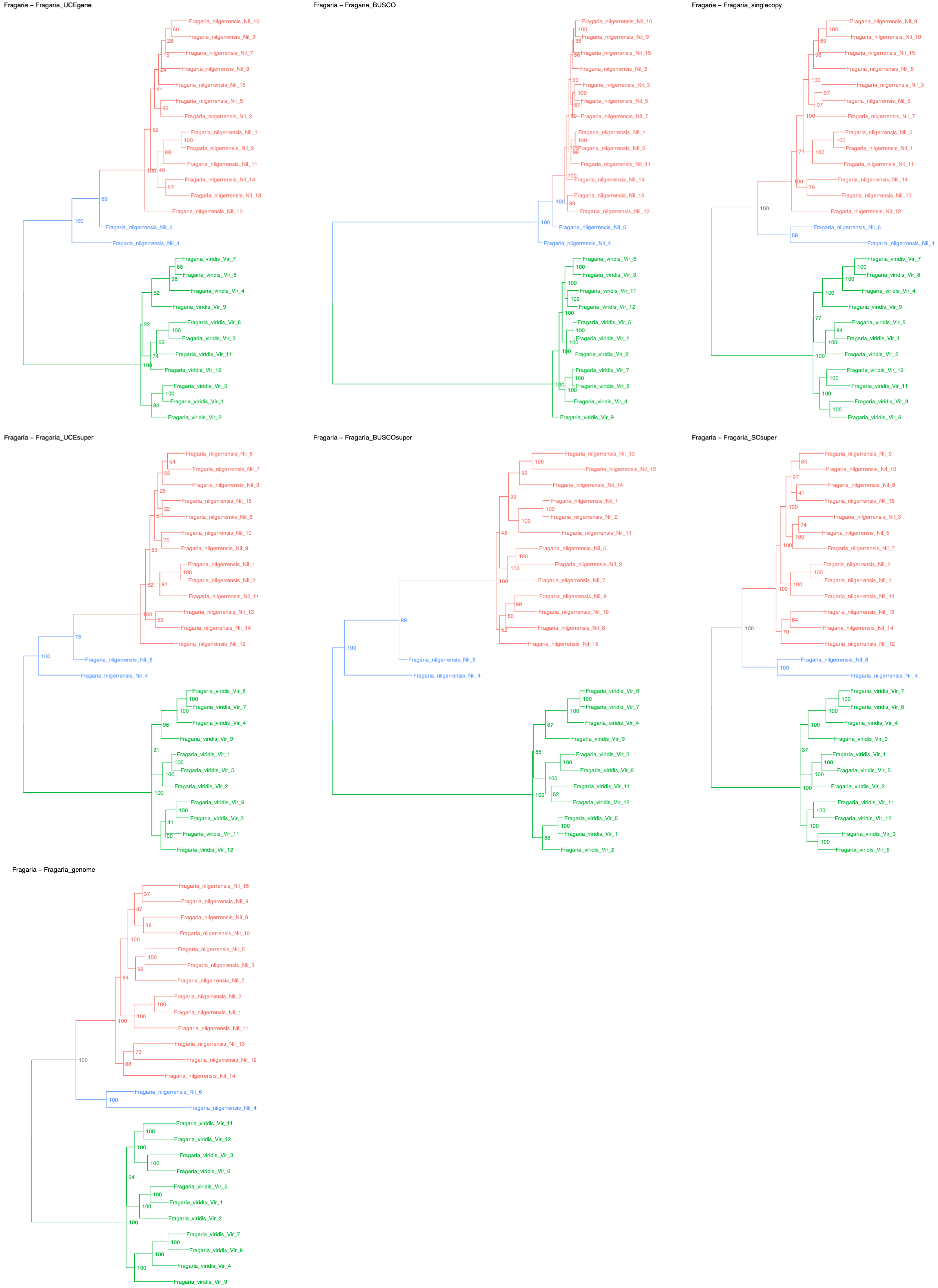
Fragaria – Fragaria_UCEgene
Fragaria_nilgerrensis_Nil_10
Fragaria_nilgerrensis_Nil_9
Fragaria_nilgerrensis_Nil_7
Fragaria_nilgerrensis_Nil_8
Fragaria_nilgerrensis_Nil_15
Fragaria_nilgerrensis_Nil_5
Fragaria_nilgerrensis_Nil_3
Fragaria_nilgerrensis_Nil_1
Fragaria_nilgerrensis_Nil_2
Fragaria_nilgerrensis_Nil_11
Fragaria_nilgerrensis_Nil_14
Fragaria_nilgerrensis_Nil_13
Fragaria_nilgerrensis_Nil_12
Fragaria_nilgerrensis_Nil_6
Fragaria_nilgerrensis_Nil_4
Fragaria_viridis_Vir_7
Fragaria_viridis_Vir_8
Fragaria_viridis_Vir_4
Fragaria_viridis_Vir_9
Fragaria_viridis_Vir_6
Fragaria_viridis_Vir_3
Fragaria_viridis_Vir_11
Fragaria_viridis_Vir_12
Fragaria_viridis_Vir_5
Fragaria_viridis_Vir_1
Fragaria_viridis_Vir_2
Fragaria – Fragaria_BUSCO
Fragaria_nilgerrensis_Nil_10
Fragaria_nilgerrensis_Nil_9
Fragaria_nilgerrensis_Nil_15
Fragaria_nilgerrensis_Nil_8
Fragaria_nilgerrensis_Nil_3
Fragaria_nilgerrensis_Nil_5
Fragaria_nilgerrensis_Nil_7
Fragaria_nilgerrensis_Nil_1
Fragaria_nilgerrensis_Nil_2
Fragaria_nilgerrensis_Nil_11
Fragaria_nilgerrensis_Nil_14
Fragaria_nilgerrensis_Nil_13
Fragaria_nilgerrensis_Nil_12
Fragaria_nilgerrensis_Nil_6
Fragaria_nilgerrensis_Nil_4
Fragaria_viridis_Vir_6
Fragaria_viridis_Vir_3
Fragaria_viridis_Vir_11
Fragaria_viridis_Vir_12
Fragaria_viridis_Vir_5
Fragaria_viridis_Vir_1
Fragaria_viridis_Vir_2
Fragaria_viridis_Vir_7
Fragaria_viridis_Vir_8
Fragaria_viridis_Vir_4
Fragaria_viridis_Vir_9
Fragaria – Fragaria_singlecopy
Fragaria_nilgerrensis_Nil_9
Fragaria_nilgerrensis_Nil_10
Fragaria_nilgerrensis_Nil_15
Fragaria_nilgerrensis_Nil_8
Fragaria_nilgerrensis_Nil_3
Fragaria_nilgerrensis_Nil_5
Fragaria_nilgerrensis_Nil_7
Fragaria_nilgerrensis_Nil_2
Fragaria_nilgerrensis_Nil_1
Fragaria_nilgerrensis_Nil_11
Fragaria_nilgerrensis_Nil_14
Fragaria_nilgerrensis_Nil_13
Fragaria_nilgerrensis_Nil_12
Fragaria_nilgerrensis_Nil_6
Fragaria_nilgerrensis_Nil_4
Fragaria_viridis_Vir_7
Fragaria_viridis_Vir_8
Fragaria_viridis_Vir_4
Fragaria_viridis_Vir_9
Fragaria_viridis_Vir_5
Fragaria_viridis_Vir_1
Fragaria_viridis_Vir_2
Fragaria_viridis_Vir_12
Fragaria_viridis_Vir_11
Fragaria_viridis_Vir_3
Fragaria_viridis_Vir_6
Fragaria – Fragaria_UCEsuper
Fragaria_nilgerrensis_Nil_5
Fragaria_nilgerrensis_Nil_7
Fragaria_nilgerrensis_Nil_3
Fragaria_nilgerrensis_Nil_15
Fragaria_nilgerrensis_Nil_8
Fragaria_nilgerrensis_Nil_10
Fragaria_nilgerrensis_Nil_9
Fragaria_nilgerrensis_Nil_1
Fragaria_nilgerrensis_Nil_2
Fragaria_nilgerrensis_Nil_11
Fragaria_nilgerrensis_Nil_13
Fragaria_nilgerrensis_Nil_14
Fragaria_nilgerrensis_Nil_12
Fragaria_nilgerrensis_Nil_6
Fragaria_nilgerrensis_Nil_4
Fragaria_viridis_Vir_8
Fragaria_viridis_Vir_7
Fragaria_viridis_Vir_4
Fragaria_viridis_Vir_9
Fragaria_viridis_Vir_1
Fragaria_viridis_Vir_5
Fragaria_viridis_Vir_2
Fragaria_viridis_Vir_6
Fragaria_viridis_Vir_3
Fragaria_viridis_Vir_11
Fragaria_viridis_Vir_12
Fragaria – Fragaria_BUSCOsuper
Fragaria_nilgerrensis_Nil_13
Fragaria_nilgerrensis_Nil_12
Fragaria_nilgerrensis_Nil_14
Fragaria_nilgerrensis_Nil_1
Fragaria_nilgerrensis_Nil_2
Fragaria_nilgerrensis_Nil_11
Fragaria_nilgerrensis_Nil_5
Fragaria_nilgerrensis_Nil_3
Fragaria_nilgerrensis_Nil_7
Fragaria_nilgerrensis_Nil_9
Fragaria_nilgerrensis_Nil_10
Fragaria_nilgerrensis_Nil_8
Fragaria_nilgerrensis_Nil_15
Fragaria_nilgerrensis_Nil_6
Fragaria_nilgerrensis_Nil_4
Fragaria_viridis_Vir_8
Fragaria_viridis_Vir_7
Fragaria_viridis_Vir_4
Fragaria_viridis_Vir_9
Fragaria_viridis_Vir_3
Fragaria_viridis_Vir_6
Fragaria_viridis_Vir_11
Fragaria_viridis_Vir_12
Fragaria_viridis_Vir_5
Fragaria_viridis_Vir_1
Fragaria_viridis_Vir_2
Fragaria – Fragaria_SCsuper
Fragaria_nilgerrensis_Nil_9
Fragaria_nilgerrensis_Nil_10
Fragaria_nilgerrensis_Nil_8
Fragaria_nilgerrensis_Nil_15
Fragaria_nilgerrensis_Nil_3
Fragaria_nilgerrensis_Nil_5
Fragaria_nilgerrensis_Nil_7
Fragaria_nilgerrensis_Nil_2
Fragaria_nilgerrensis_Nil_1
Fragaria_nilgerrensis_Nil_11
Fragaria_nilgerrensis_Nil_13
Fragaria_nilgerrensis_Nil_14
Fragaria_nilgerrensis_Nil_12
Fragaria_nilgerrensis_Nil_6
Fragaria_nilgerrensis_Nil_4
Fragaria_viridis_Vir_7
Fragaria_viridis_Vir_8
Fragaria_viridis_Vir_4
Fragaria_viridis_Vir_9
Fragaria_viridis_Vir_1
Fragaria_viridis_Vir_5
Fragaria_viridis_Vir_2
Fragaria_viridis_Vir_11
Fragaria_viridis_Vir_12
Fragaria_viridis_Vir_3
Fragaria_viridis_Vir_6
Fragaria – Fragaria_genome
Fragaria_nilgerrensis_Nil_15
Fragaria_nilgerrensis_Nil_9
Fragaria_nilgerrensis_Nil_8
Fragaria_nilgerrensis_Nil_10
Fragaria_nilgerrensis_Nil_5
Fragaria_nilgerrensis_Nil_3
Fragaria_nilgerrensis_Nil_7
Fragaria_nilgerrensis_Nil_2
Fragaria_nilgerrensis_Nil_1
Fragaria_nilgerrensis_Nil_11
Fragaria_nilgerrensis_Nil_13
Fragaria_nilgerrensis_Nil_12
Fragaria_nilgerrensis_Nil_14
Fragaria_nilgerrensis_Nil_6
Fragaria_nilgerrensis_Nil_4
Fragaria_viridis_Vir_11
Fragaria_viridis_Vir_12
Fragaria_viridis_Vir_3
Fragaria_viridis_Vir_6
Fragaria_viridis_Vir_5
Fragaria_viridis_Vir_1
Fragaria_viridis_Vir_2
Fragaria_viridis_Vir_7
Fragaria_viridis_Vir_8
Fragaria_viridis_Vir_4
Fragaria_viridis_Vir_9

Juglans – Juglans_UCEgene

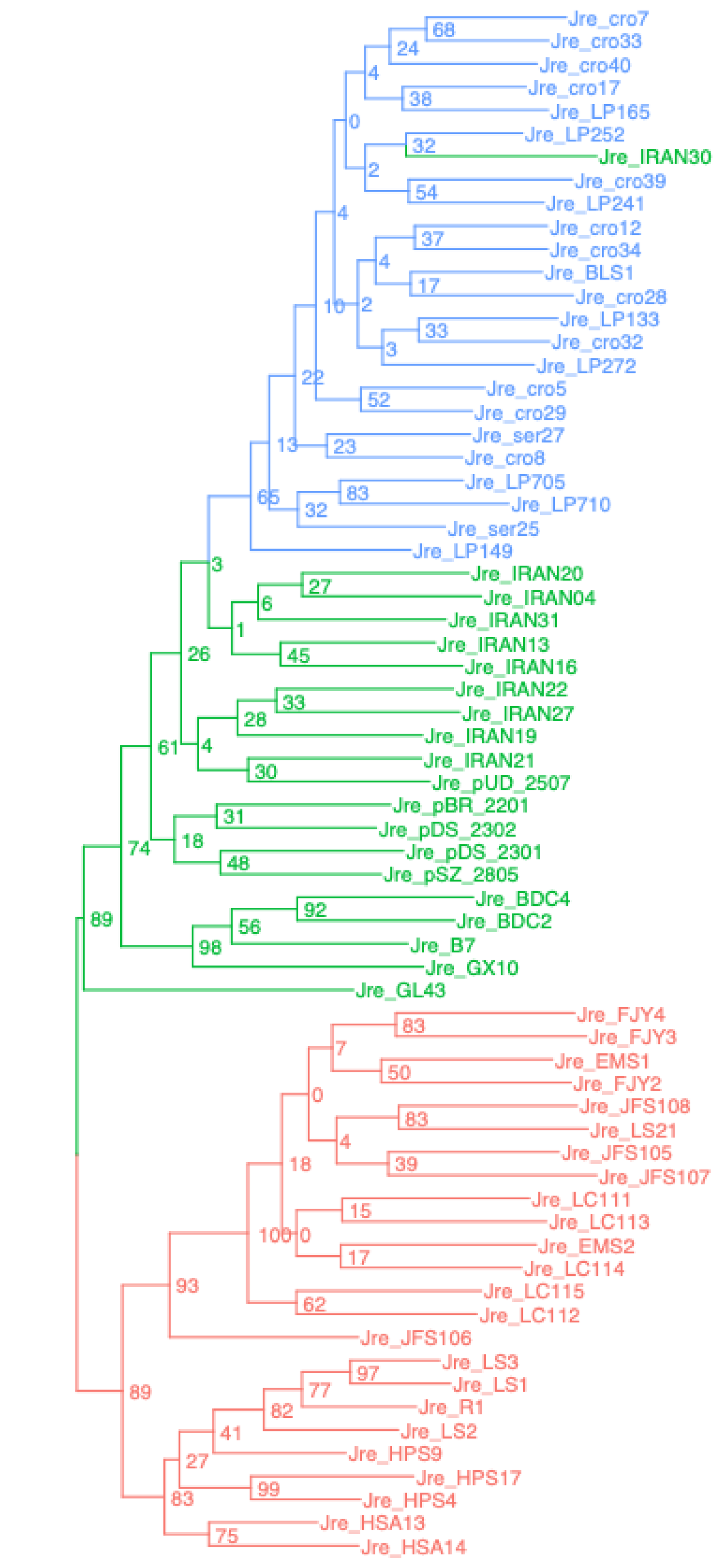


Juglans – Juglans_BUSCO

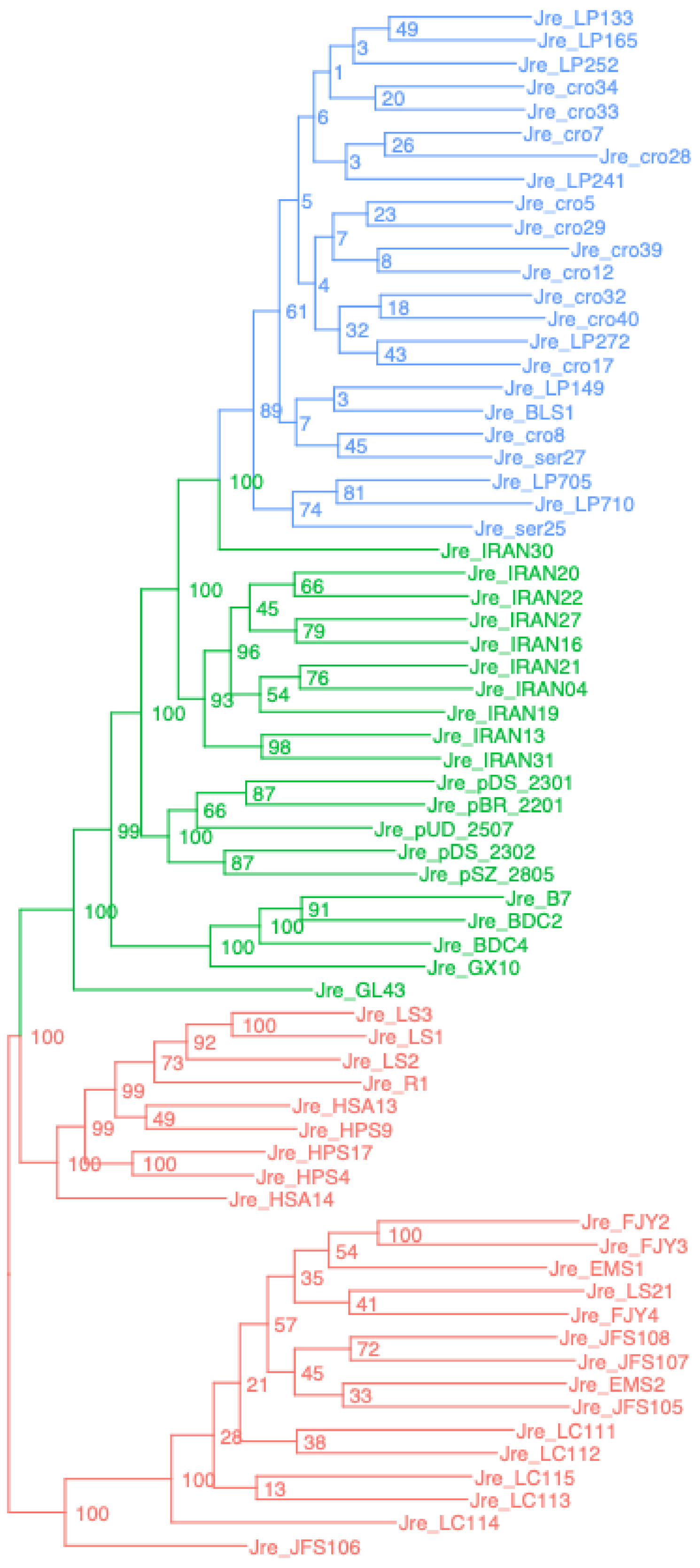


Juglans – Juglans_singlecopy

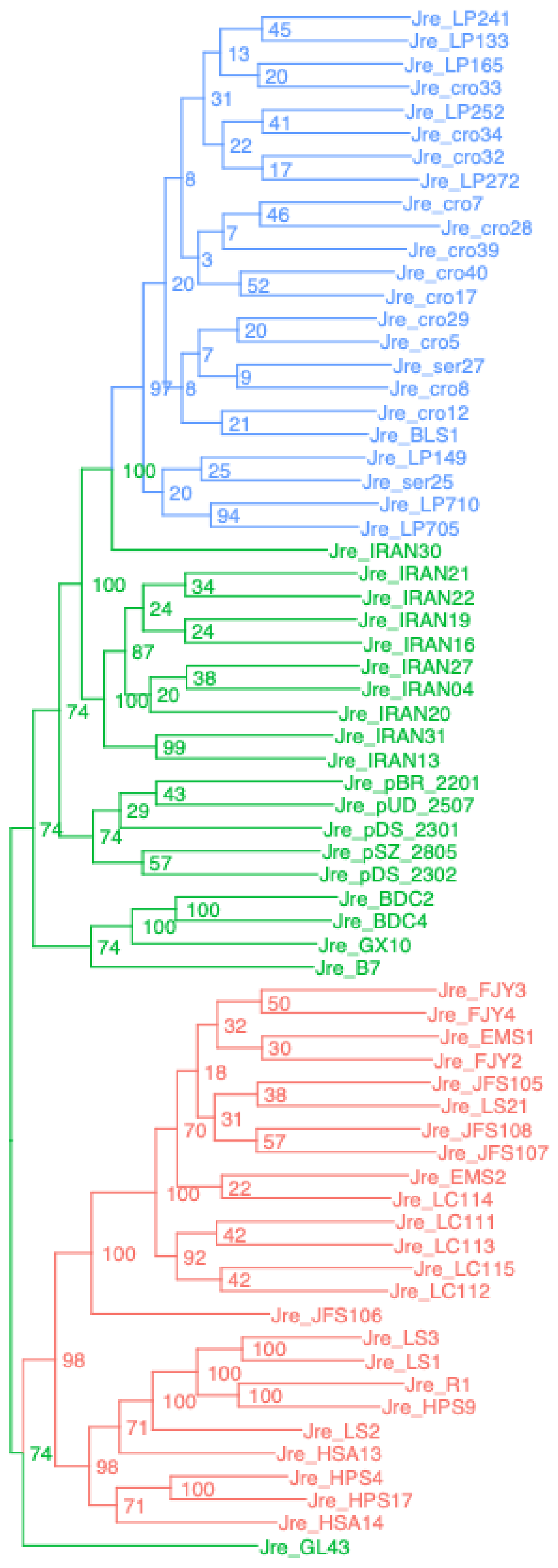


Juglans – Juglans_UCEsuper

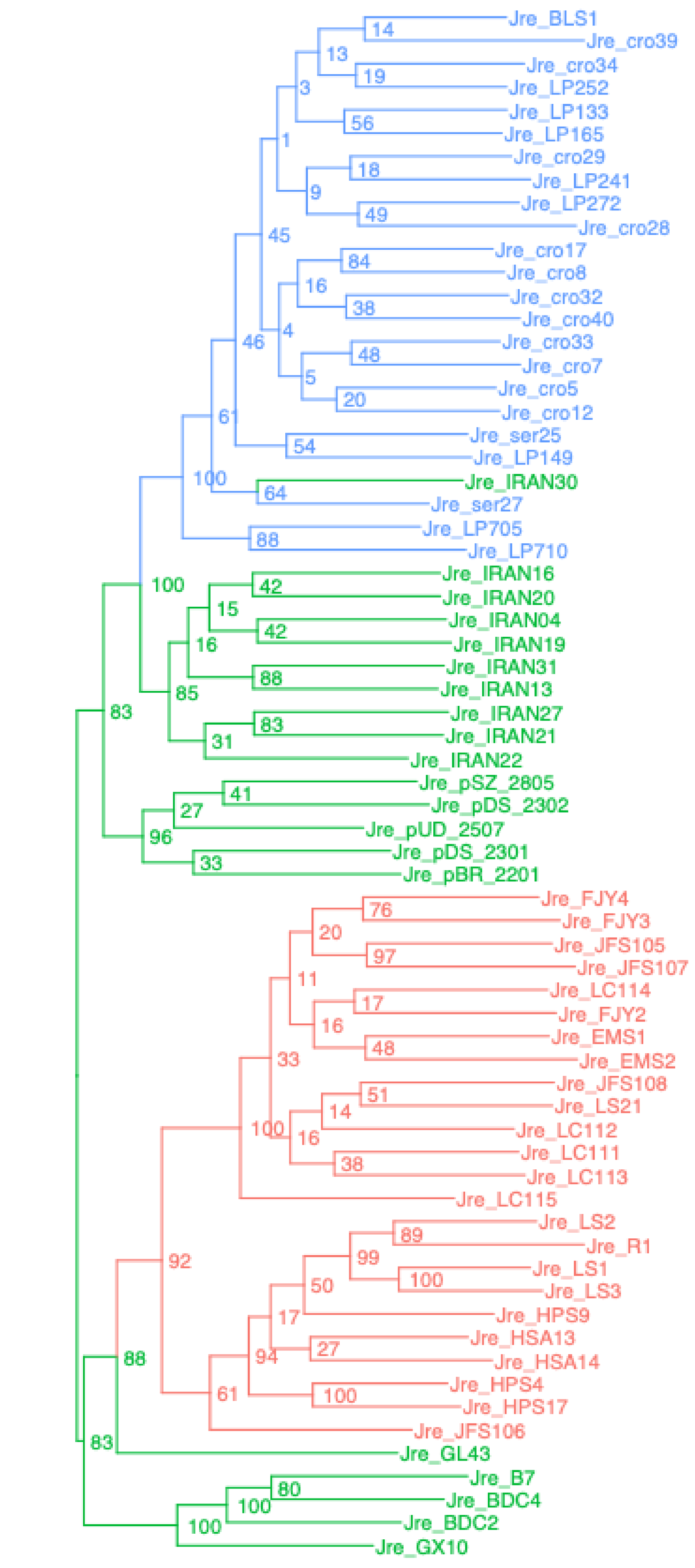


Juglans – Juglans_BUSCOsuper

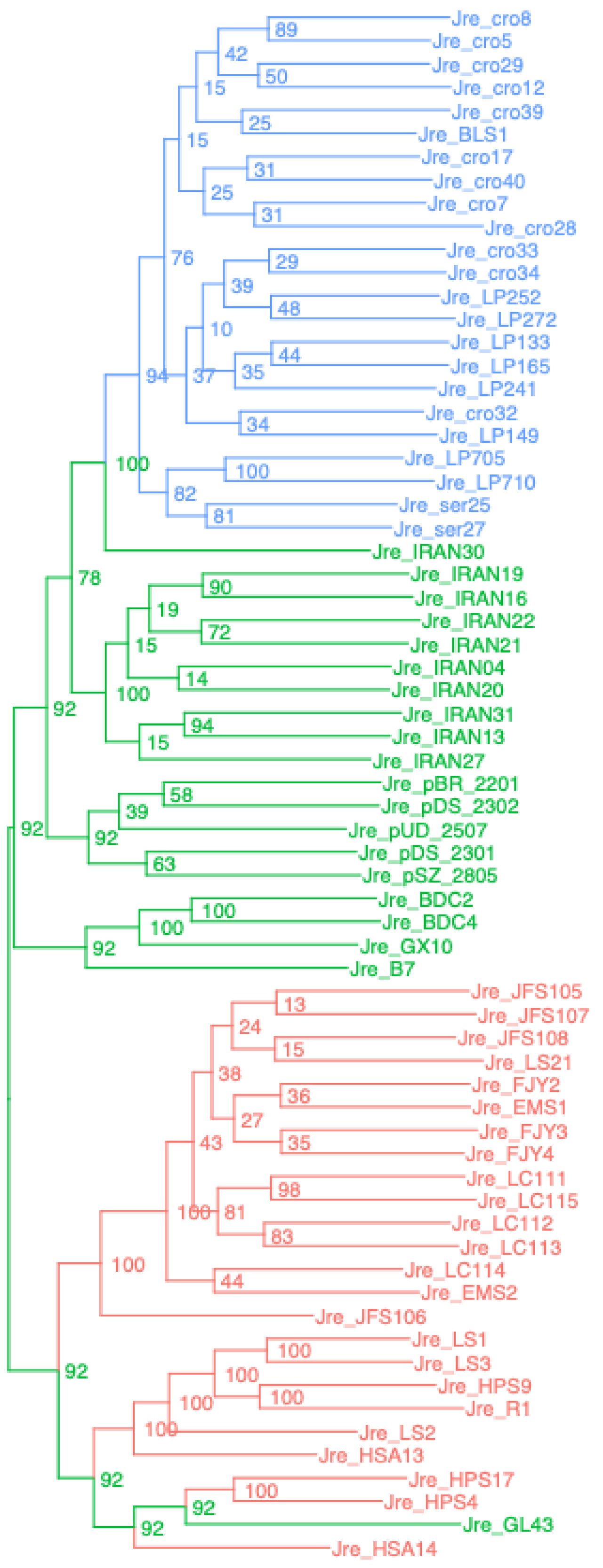


Juglans – Juglans_SCsuper

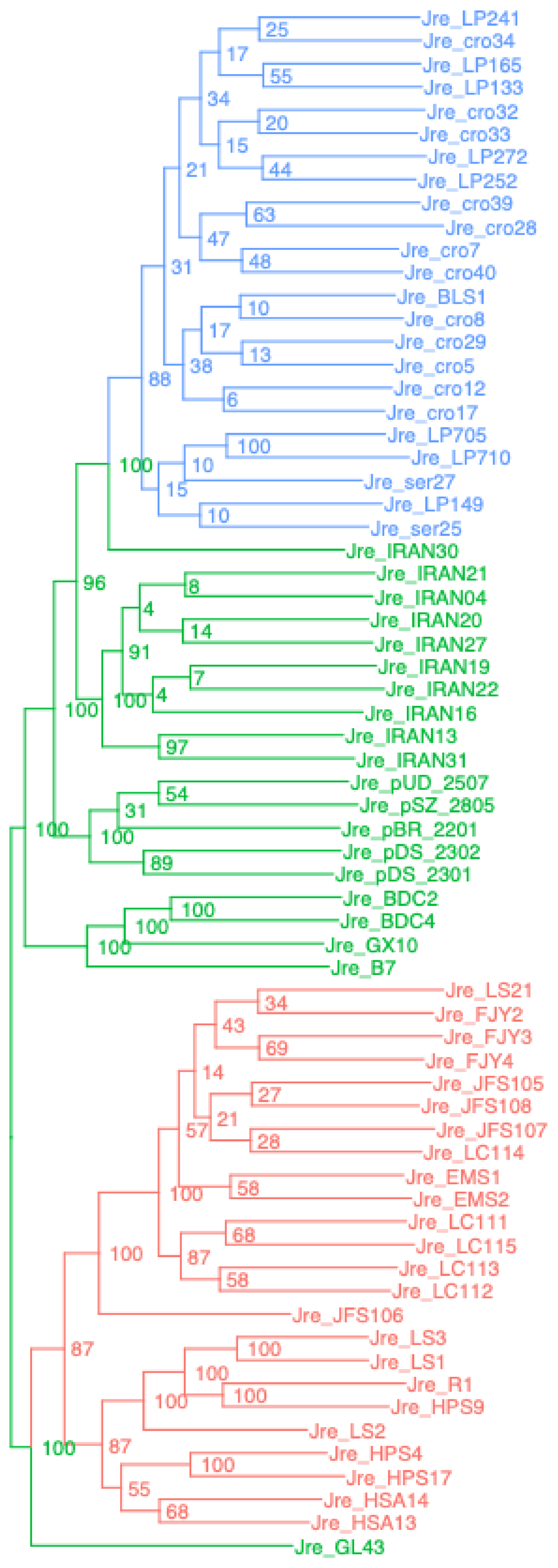


Juglans – Juglans_genome

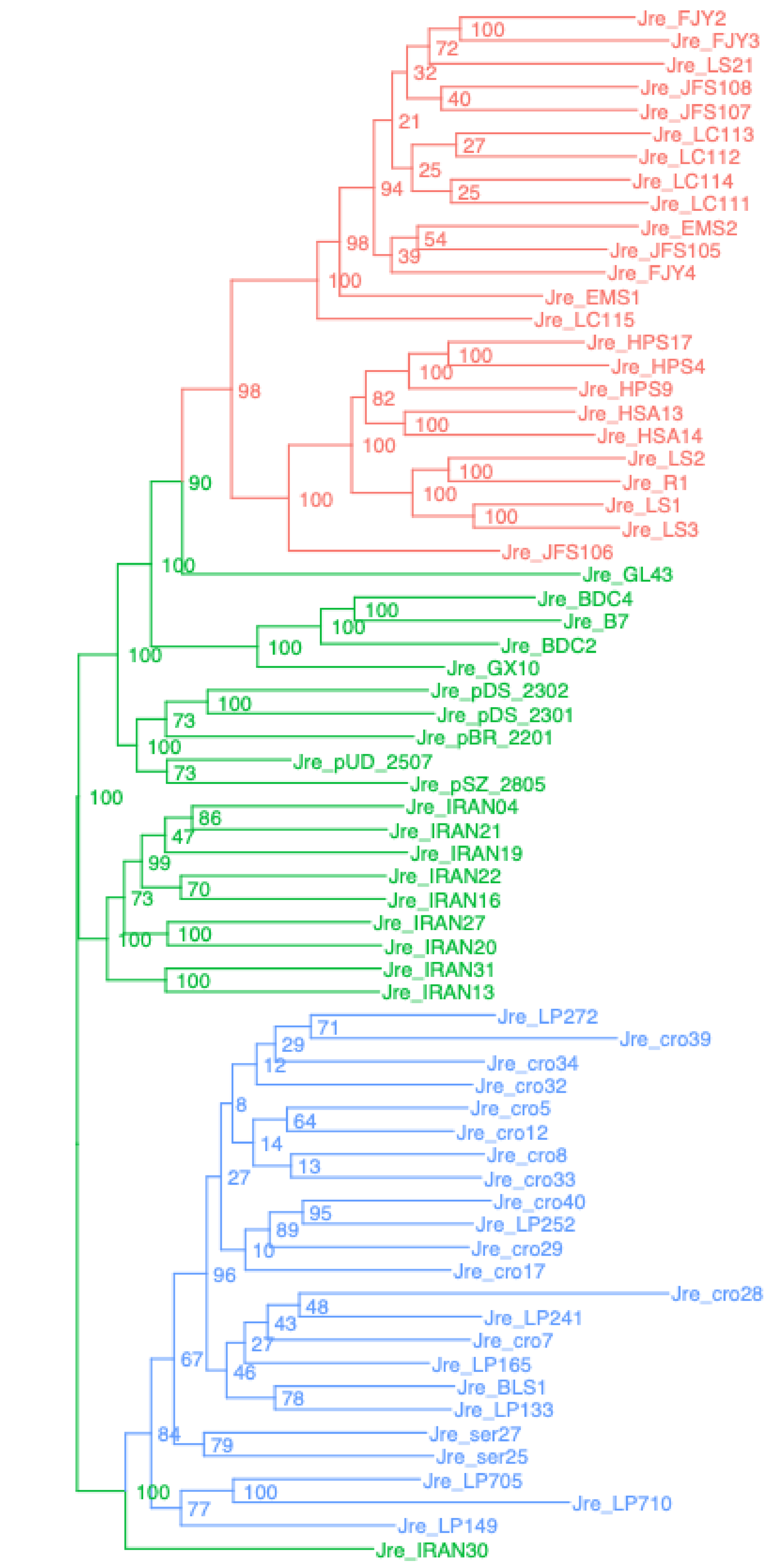

Liriodendron – Liriodendron_UCEgene

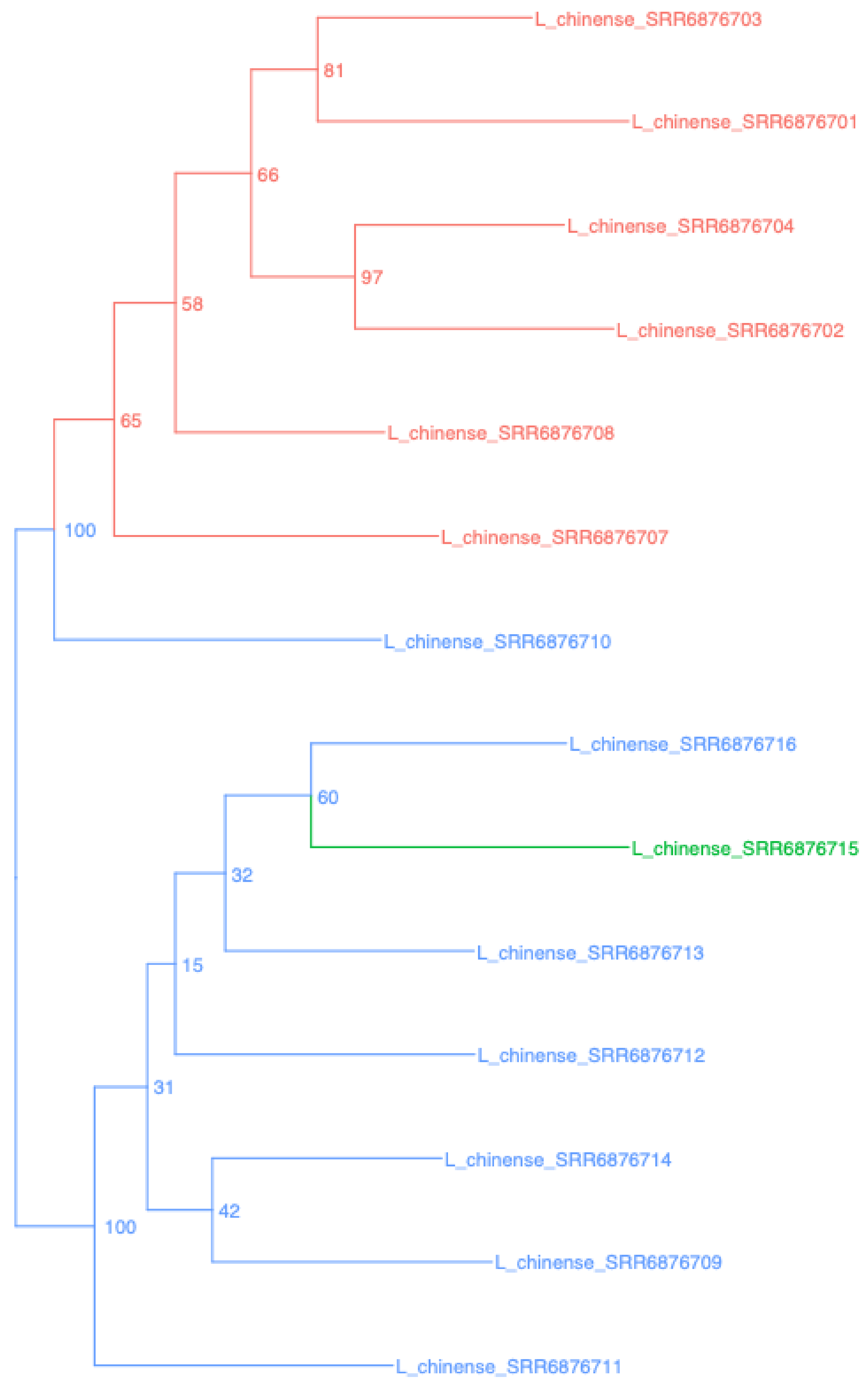


Liriodendron – Liriodendron_BUSCO

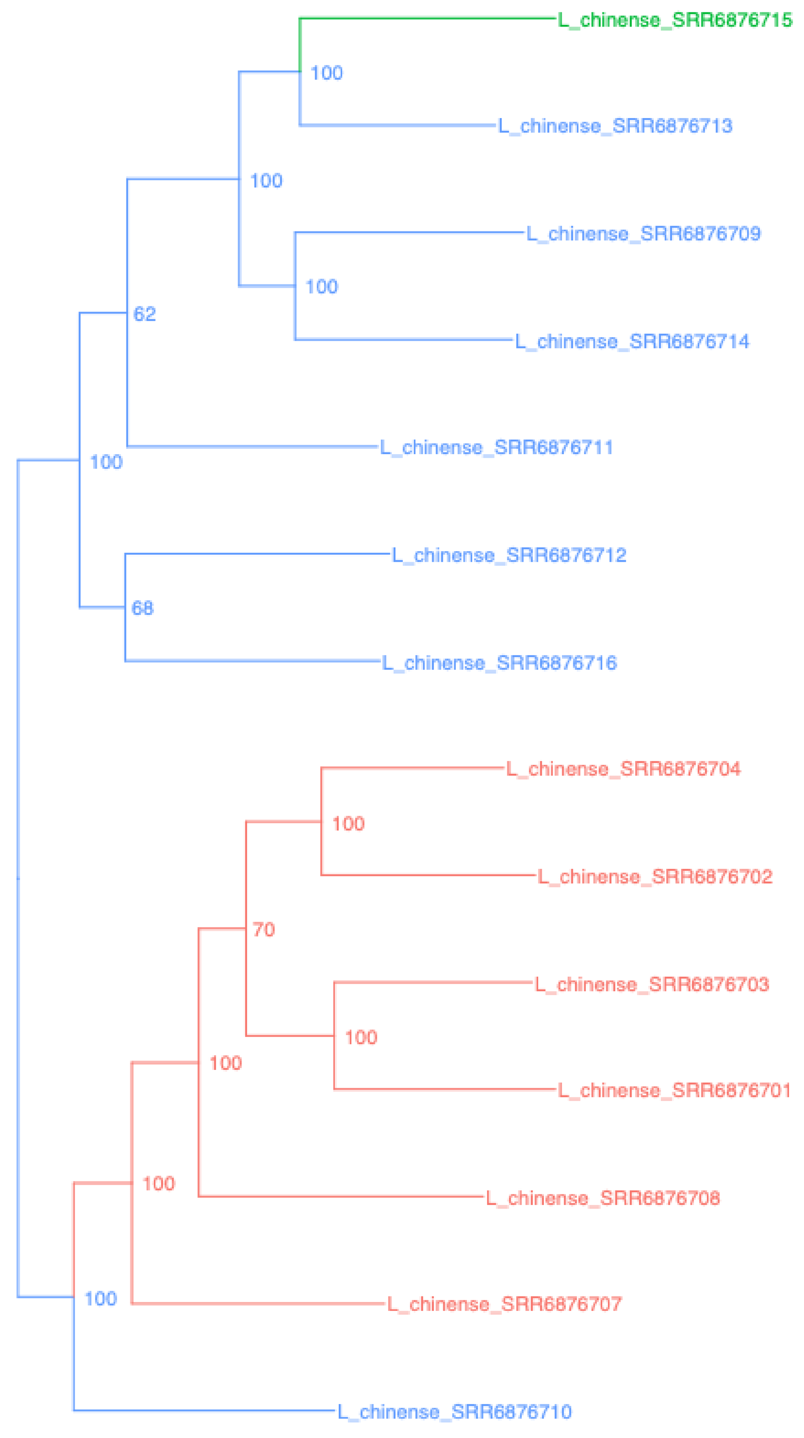


Liriodendron – Liriodendron_chinense_singlecopy

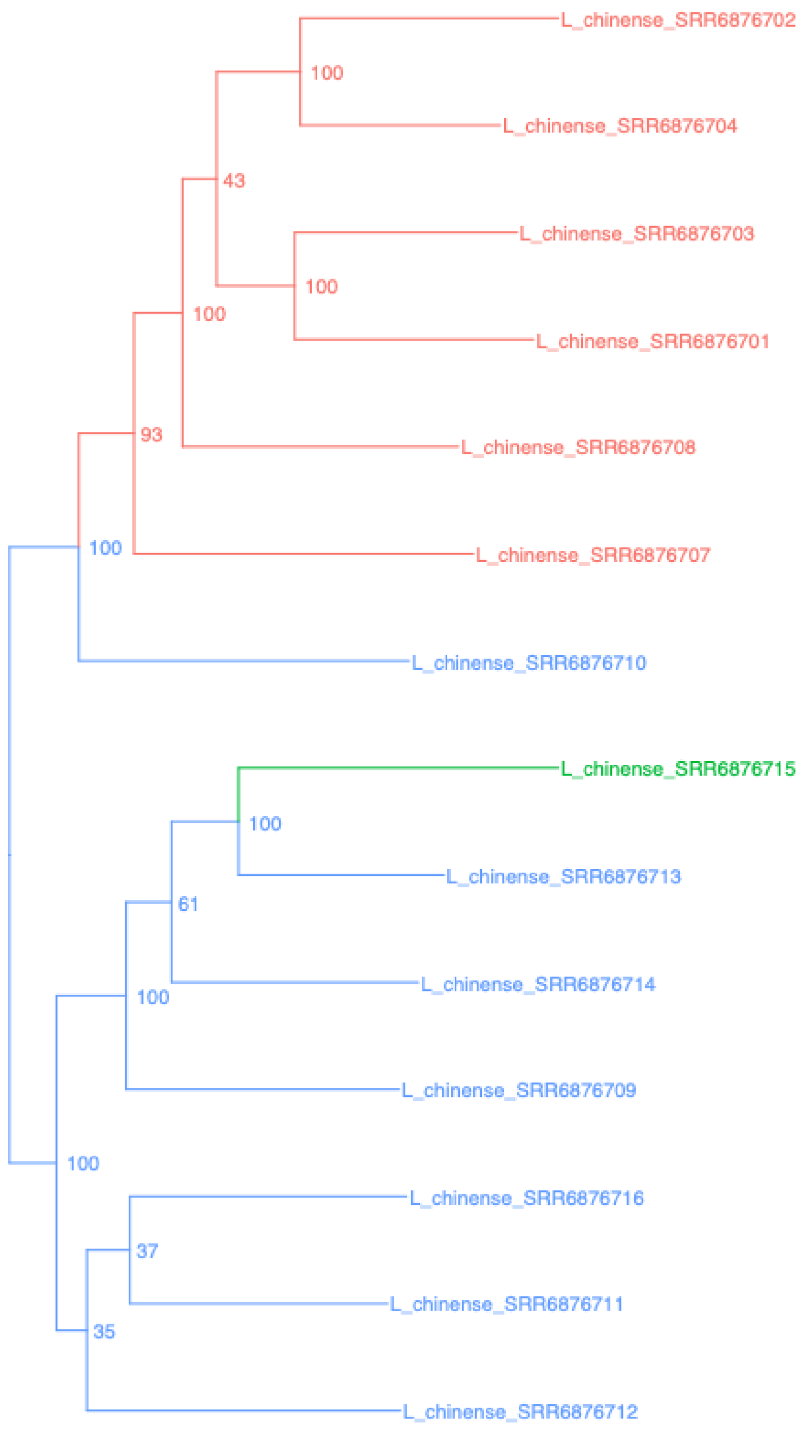


Liriodendron – Liriodendron_UCEsuper

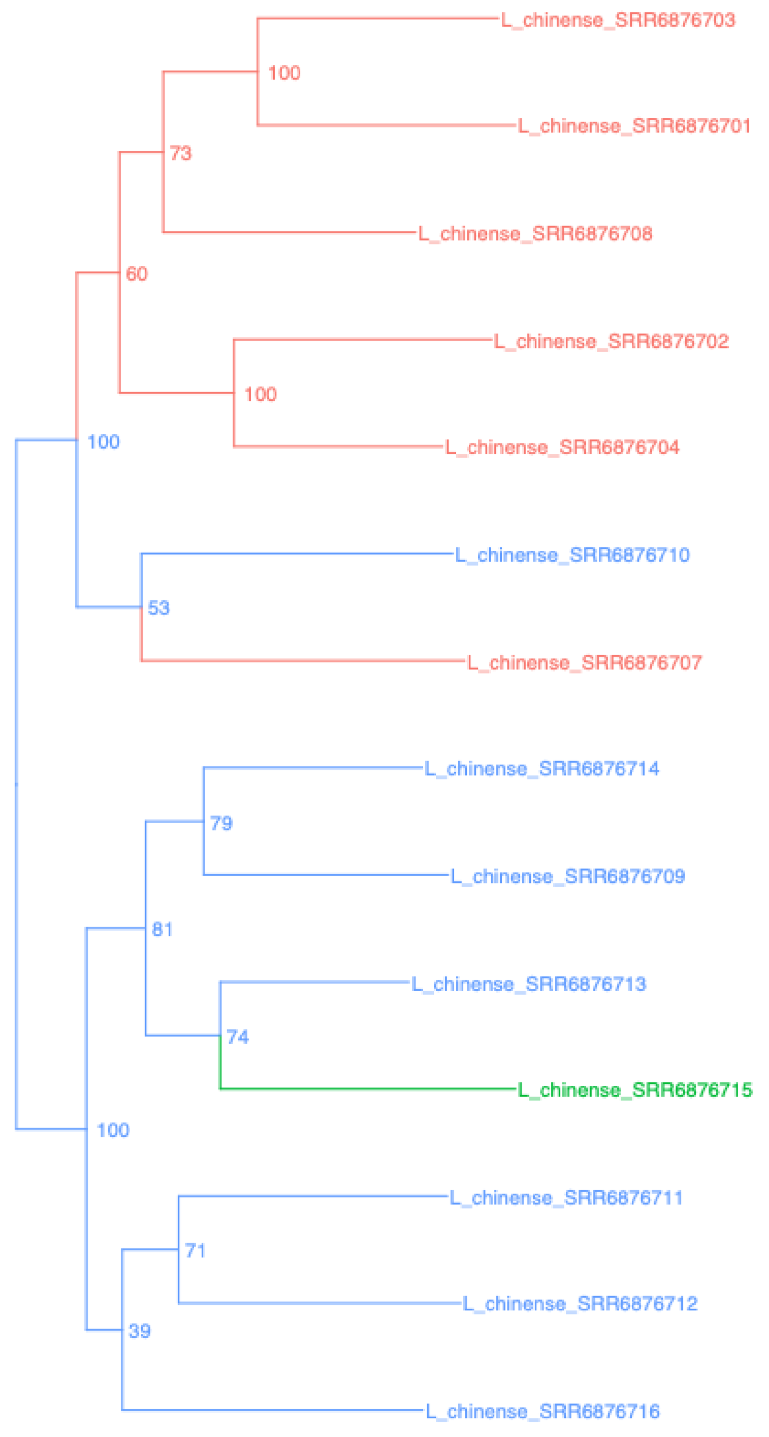


Liriodendron – Liriodendron_BUSCOsuper

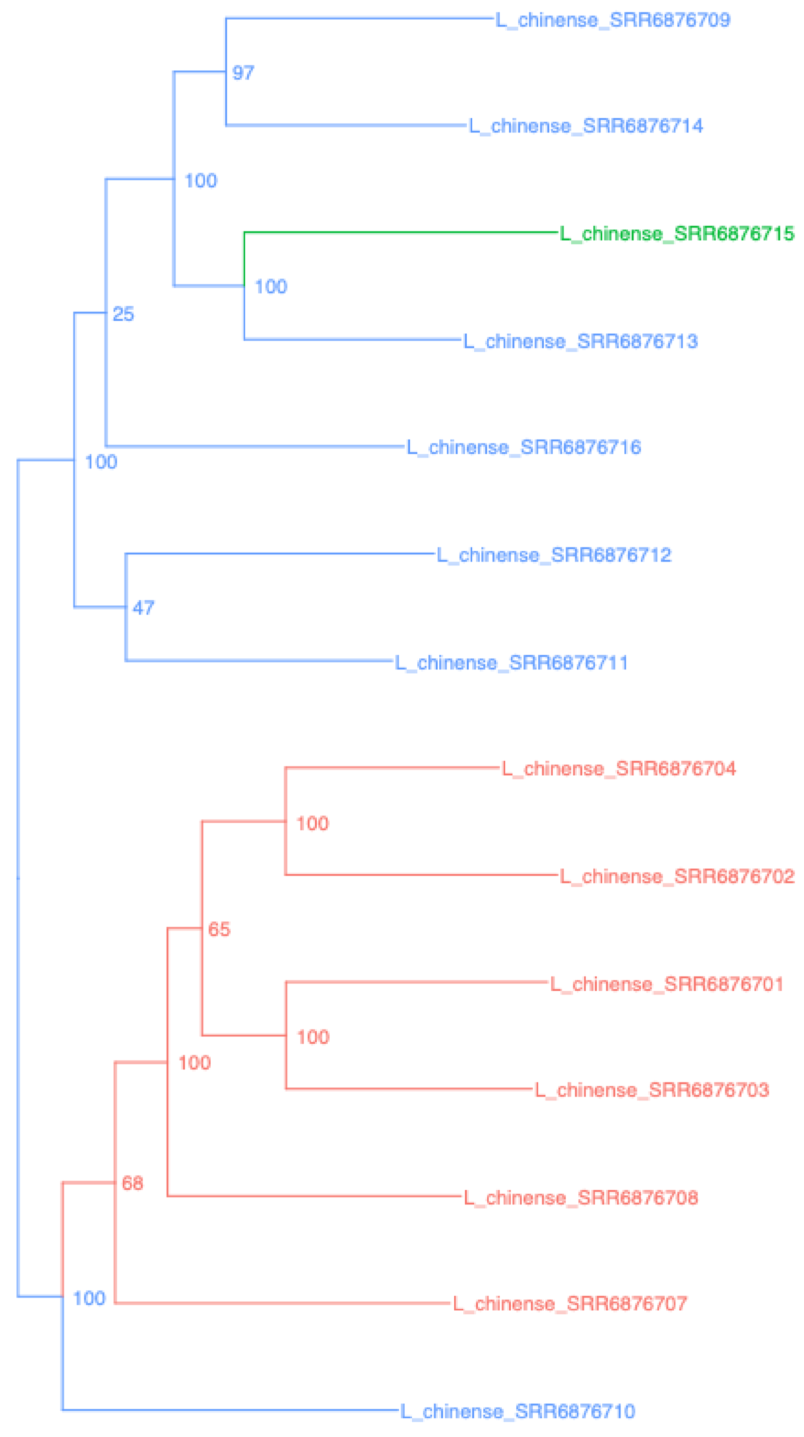


Liriodendron – Liriodendron_SCsuper

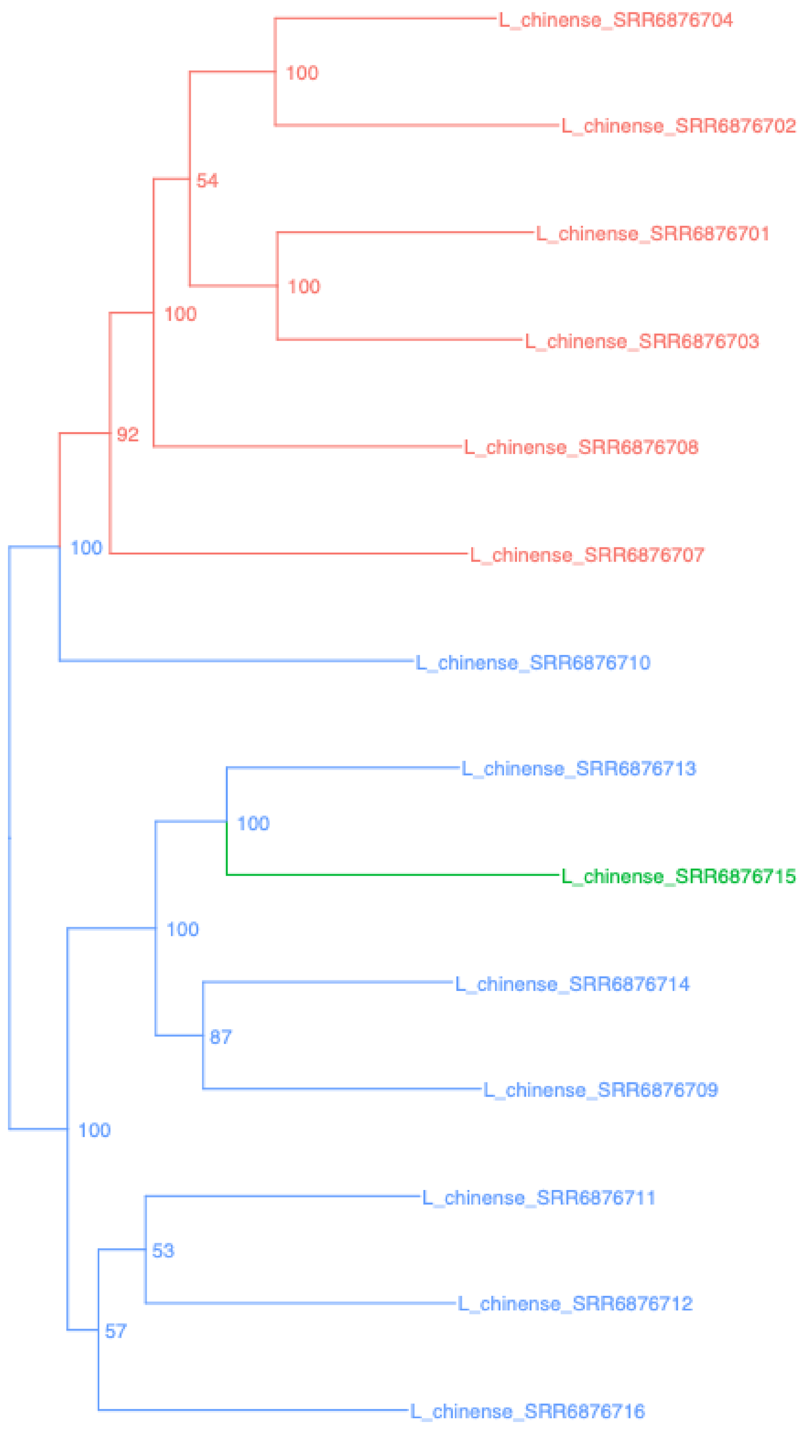


Liriodendron – Liriodendron_genome

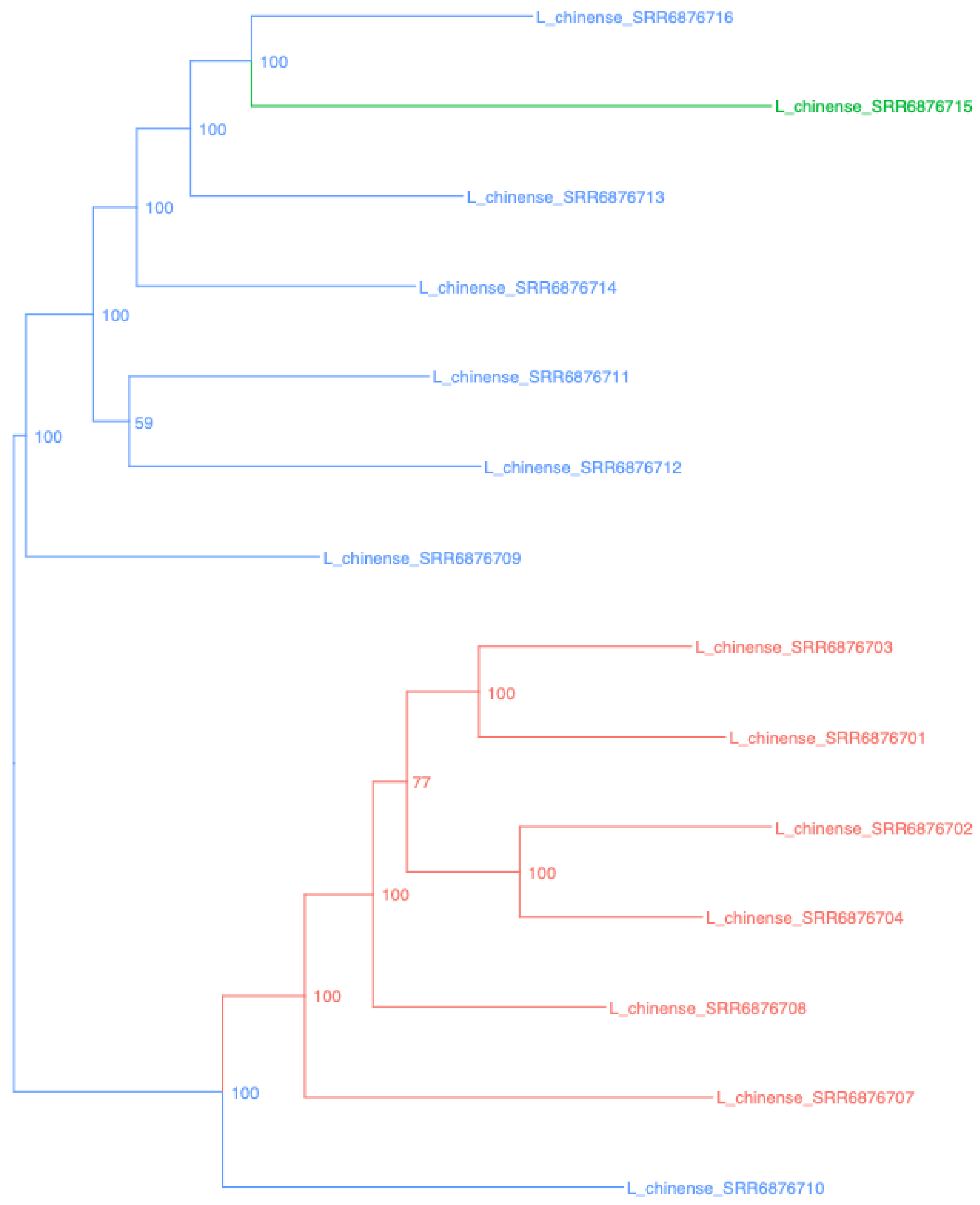

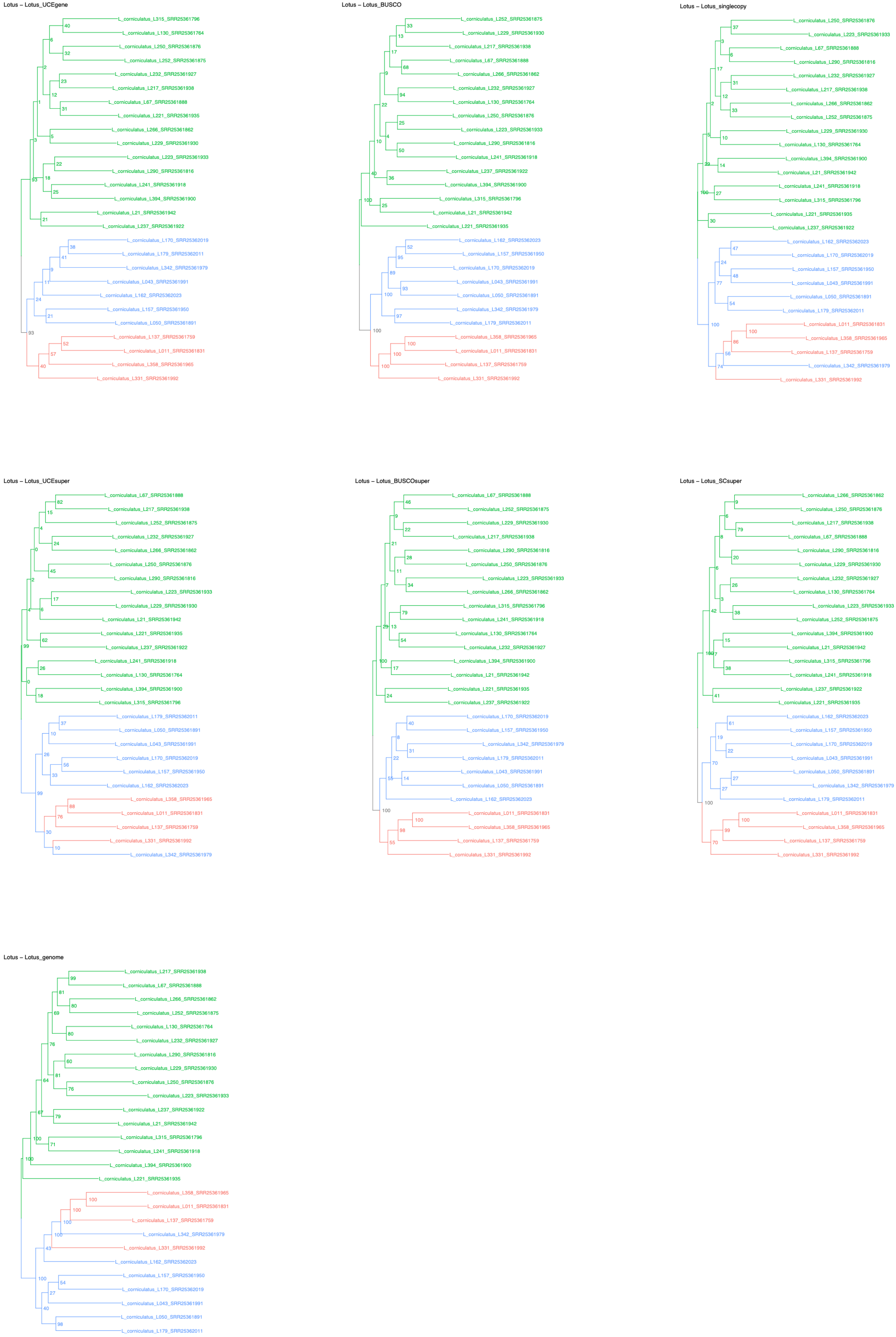

Lotus – Lotus_UCEgene
L_corniculatus_L315_SRR25361796
L_corniculatus_L130_SRR25361764
L_corniculatus_L250_SRR25361876
L_corniculatus_L252_SRR25361875
L_corniculatus_L232_SRR25361927
L_corniculatus_L217_SRR25361938
L_corniculatus_L67_SRR25361888
L_corniculatus_L221_SRR25361935
L_corniculatus_L266_SRR25361862
L_corniculatus_L229_SRR25361930
L_corniculatus_L223_SRR25361933
L_corniculatus_L290_SRR25361816
L_corniculatus_L241_SRR25361918
L_corniculatus_L394_SRR25361900
L_corniculatus_L21_SRR25361942
L_corniculatus_L237_SRR25361922
L_corniculatus_L170_SRR25362019
L_corniculatus_L179_SRR25362011
L_corniculatus_L342_SRR25361979
L_corniculatus_L043_SRR25361991
L_corniculatus_L162_SRR25362023
L_corniculatus_L157_SRR25361950
L_corniculatus_L050_SRR25361891
L_corniculatus_L137_SRR25361759
L_corniculatus_L011_SRR25361831
L_corniculatus_L358_SRR25361965
L_corniculatus_L331_SRR25361992
Lotus – Lotus_BUSCO
L_corniculatus_L252_SRR25361875
L_corniculatus_L229_SRR25361930
L_corniculatus_L217_SRR25361938
L_corniculatus_L67_SRR25361888
L_corniculatus_L266_SRR25361862
L_corniculatus_L232_SRR25361927
L_corniculatus_L130_SRR25361764
L_corniculatus_L250_SRR25361876
L_corniculatus_L223_SRR25361933
L_corniculatus_L290_SRR25361816
L_corniculatus_L241_SRR25361918
L_corniculatus_L237_SRR25361922
L_corniculatus_L394_SRR25361900
L_corniculatus_L315_SRR25361796
L_corniculatus_L21_SRR25361942
L_corniculatus_L221_SRR25361935
L_corniculatus_L162_SRR25362023
L_corniculatus_L157_SRR25361950
L_corniculatus_L170_SRR25362019
L_corniculatus_L043_SRR25361991
L_corniculatus_L050_SRR25361891
L_corniculatus_L342_SRR25361979
L_corniculatus_L179_SRR25362011
L_corniculatus_L358_SRR25361965
L_corniculatus_L011_SRR25361831
L_corniculatus_L137_SRR25361759
L_corniculatus_L331_SRR25361992
Lotus – Lotus_singlecopy
L_corniculatus_L250_SRR25361876
L_corniculatus_L223_SRR25361933
L_corniculatus_L67_SRR25361888
L_corniculatus_L290_SRR25361816
L_corniculatus_L232_SRR25361927
L_corniculatus_L217_SRR25361938
L_corniculatus_L266_SRR25361862
L_corniculatus_L252_SRR25361875
L_corniculatus_L229_SRR25361930
L_corniculatus_L130_SRR25361764
L_corniculatus_L394_SRR25361900
L_corniculatus_L21_SRR25361942
L_corniculatus_L241_SRR25361918
L_corniculatus_L315_SRR25361796
L_corniculatus_L221_SRR25361935
L_corniculatus_L237_SRR25361922
L_corniculatus_L162_SRR25362023
L_corniculatus_L170_SRR25362019
L_corniculatus_L157_SRR25361950
L_corniculatus_L043_SRR25361991
L_corniculatus_L050_SRR25361891
L_corniculatus_L179_SRR25362011
L_corniculatus_L011_SRR25361831
L_corniculatus_L358_SRR25361965
L_corniculatus_L137_SRR25361759
L_corniculatus_L342_SRR25361979
L_corniculatus_L331_SRR25361992
Lotus – Lotus_UCEsuper
L_corniculatus_L67_SRR25361888
L_corniculatus_L217_SRR25361938
L_corniculatus_L252_SRR25361875
L_corniculatus_L232_SRR25361927
L_corniculatus_L266_SRR25361862
L_corniculatus_L250_SRR25361876
L_corniculatus_L290_SRR25361816
L_corniculatus_L223_SRR25361933
L_corniculatus_L229_SRR25361930
L_corniculatus_L21_SRR25361942
L_corniculatus_L221_SRR25361935
L_corniculatus_L237_SRR25361922
L_corniculatus_L241_SRR25361918
L_corniculatus_L130_SRR25361764
L_corniculatus_L394_SRR25361900
L_corniculatus_L315_SRR25361796
L_corniculatus_L179_SRR25362011
L_corniculatus_L050_SRR25361891
L_corniculatus_L043_SRR25361991
L_corniculatus_L170_SRR25362019
L_corniculatus_L157_SRR25361950
L_corniculatus_L162_SRR25362023
L_corniculatus_L358_SRR25361965
L_corniculatus_L011_SRR25361831
L_corniculatus_L137_SRR25361759
L_corniculatus_L331_SRR25361992
L_corniculatus_L342_SRR25361979
Lotus – Lotus_BUSCOsuper
L_corniculatus_L67_SRR25361888
L_corniculatus_L252_SRR25361875
L_corniculatus_L229_SRR25361930
L_corniculatus_L217_SRR25361938
L_corniculatus_L290_SRR25361816
L_corniculatus_L250_SRR25361876
L_corniculatus_L223_SRR25361933
L_corniculatus_L266_SRR25361862
L_corniculatus_L315_SRR25361796
L_corniculatus_L241_SRR25361918
L_corniculatus_L130_SRR25361764
L_corniculatus_L232_SRR25361927
L_corniculatus_L394_SRR25361900
L_corniculatus_L21_SRR25361942
L_corniculatus_L221_SRR25361935
L_corniculatus_L237_SRR25361922
L_corniculatus_L170_SRR25362019
L_corniculatus_L157_SRR25361950
L_corniculatus_L342_SRR25361979
L_corniculatus_L179_SRR25362011
L_corniculatus_L043_SRR25361991
L_corniculatus_L050_SRR25361891
L_corniculatus_L162_SRR25362023
L_corniculatus_L011_SRR25361831
L_corniculatus_L358_SRR25361965
L_corniculatus_L137_SRR25361759
L_corniculatus_L331_SRR25361992
Lotus – Lotus_SCsuper
L_corniculatus_L266_SRR25361862
L_corniculatus_L250_SRR25361876
L_corniculatus_L217_SRR25361938
L_corniculatus_L67_SRR25361888
L_corniculatus_L290_SRR25361816
L_corniculatus_L229_SRR25361930
L_corniculatus_L232_SRR25361927
L_corniculatus_L130_SRR25361764
L_corniculatus_L223_SRR25361933
L_corniculatus_L252_SRR25361875
L_corniculatus_L394_SRR25361900
L_corniculatus_L21_SRR25361942
L_corniculatus_L315_SRR25361796
L_corniculatus_L241_SRR25361918
L_corniculatus_L237_SRR25361922
L_corniculatus_L221_SRR25361935
L_corniculatus_L162_SRR25362023
L_corniculatus_L157_SRR25361950
L_corniculatus_L170_SRR25362019
L_corniculatus_L043_SRR25361991
L_corniculatus_L050_SRR25361891
L_corniculatus_L342_SRR25361979
L_corniculatus_L179_SRR25362011
L_corniculatus_L011_SRR25361831
L_corniculatus_L358_SRR25361965
L_corniculatus_L137_SRR25361759
L_corniculatus_L331_SRR25361992
Lotus – Lotus_genome
L_corniculatus_L217_SRR25361938
L_corniculatus_L67_SRR25361888
L_corniculatus_L266_SRR25361862
L_corniculatus_L252_SRR25361875
L_corniculatus_L130_SRR25361764
L_corniculatus_L232_SRR25361927
L_corniculatus_L290_SRR25361816
L_corniculatus_L229_SRR25361930
L_corniculatus_L250_SRR25361876
L_corniculatus_L223_SRR25361933
L_corniculatus_L237_SRR25361922
L_corniculatus_L21_SRR25361942
L_corniculatus_L315_SRR25361796
L_corniculatus_L241_SRR25361918
L_corniculatus_L394_SRR25361900
L_corniculatus_L221_SRR25361935
L_corniculatus_L358_SRR25361965
L_corniculatus_L011_SRR25361831
L_corniculatus_L137_SRR25361759
L_corniculatus_L342_SRR25361979
L_corniculatus_L331_SRR25361992
L_corniculatus_L162_SRR25362023
L_corniculatus_L157_SRR25361950
L_corniculatus_L170_SRR25362019
L_corniculatus_L043_SRR25361991
L_corniculatus_L050_SRR25361891
L_corniculatus_L179_SRR25362011

Malania – Malania_UCEgene

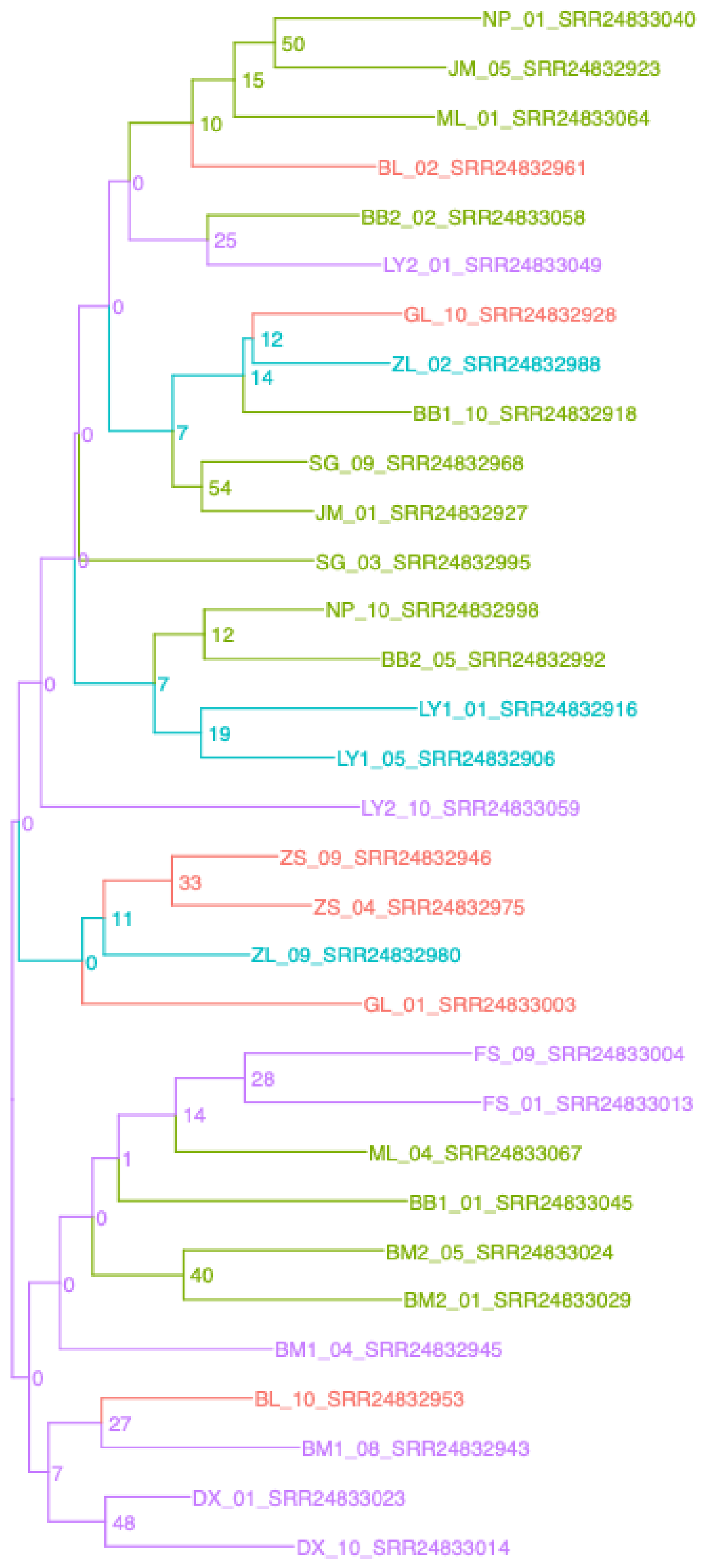


Malania – Malania_BUSCO

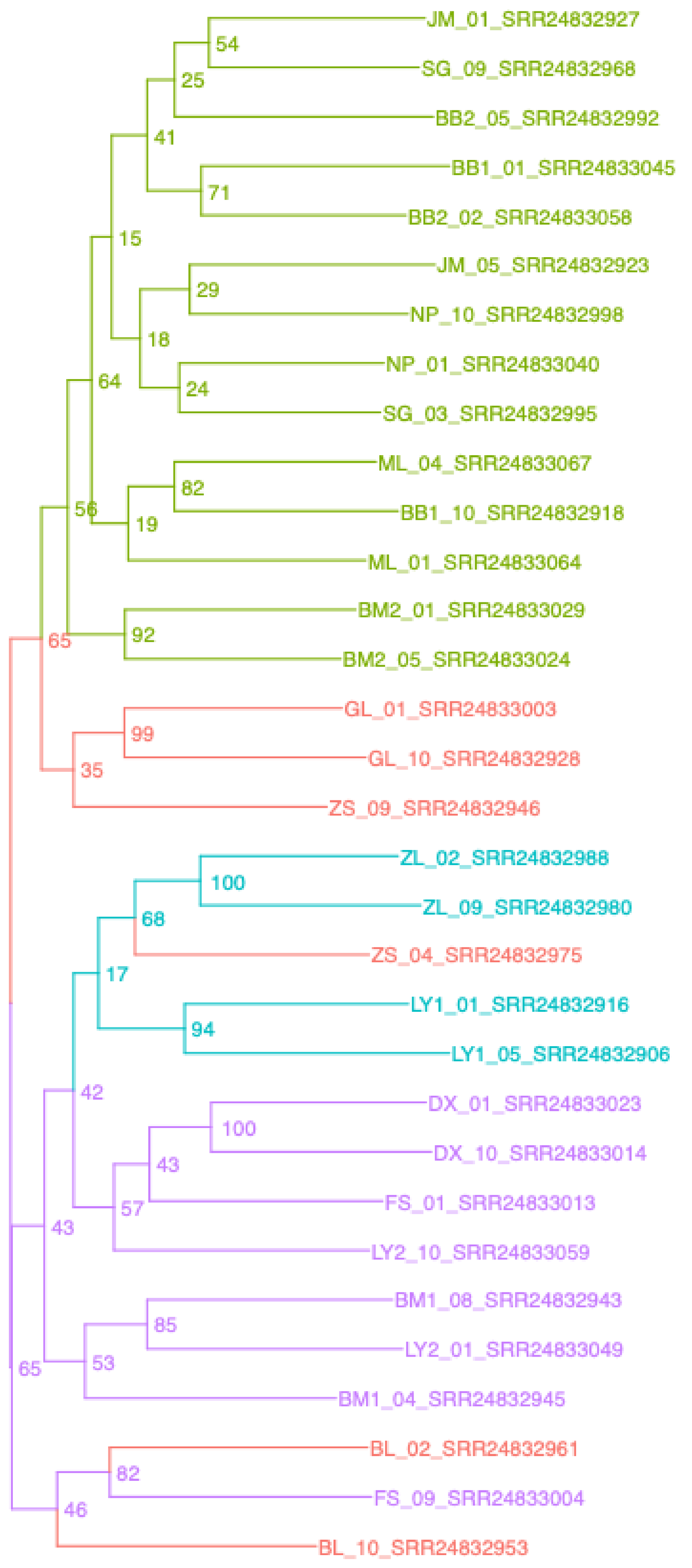


Malania – Malania_singlecopy

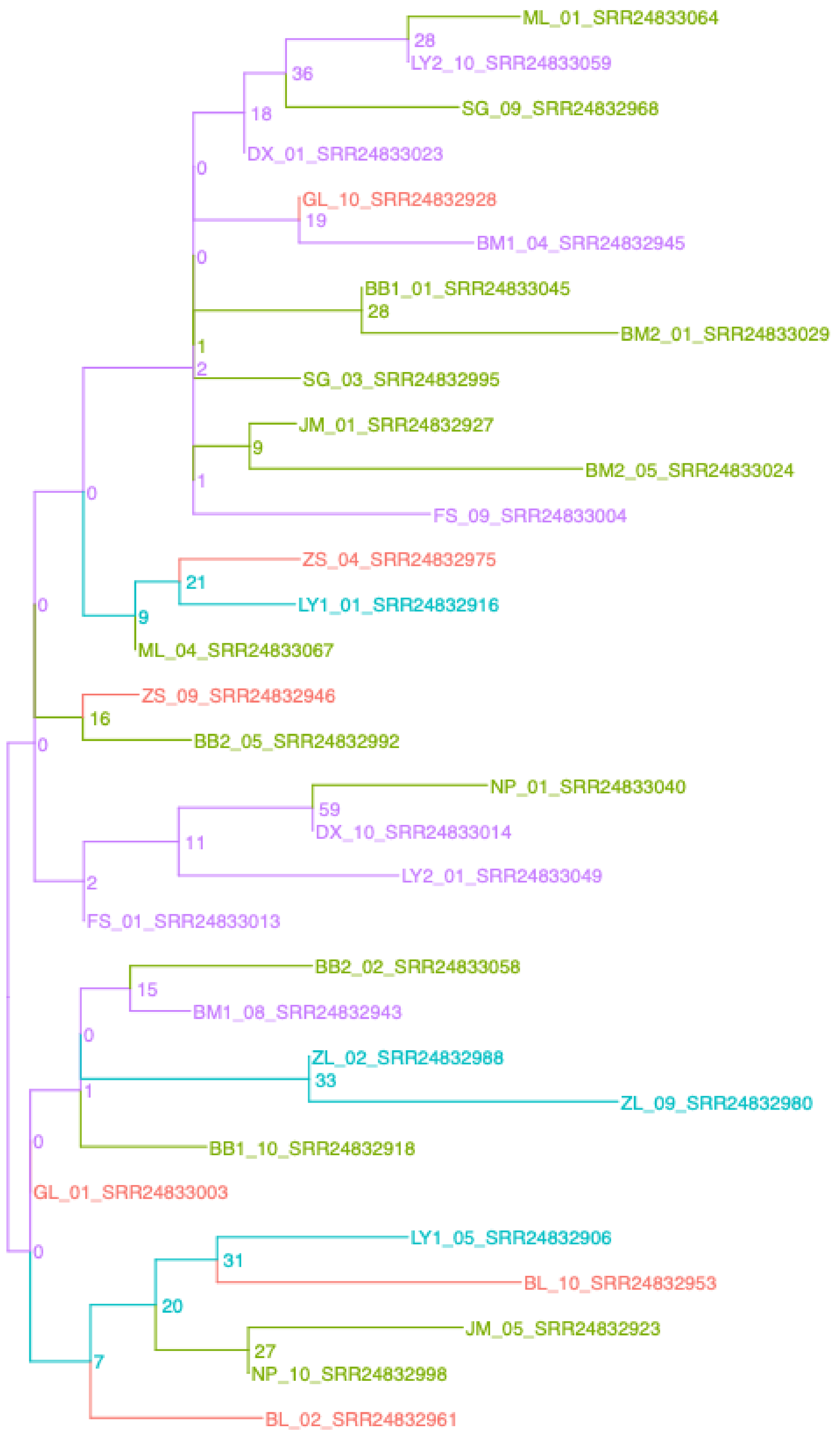


Malania – Malania_UCEsuper

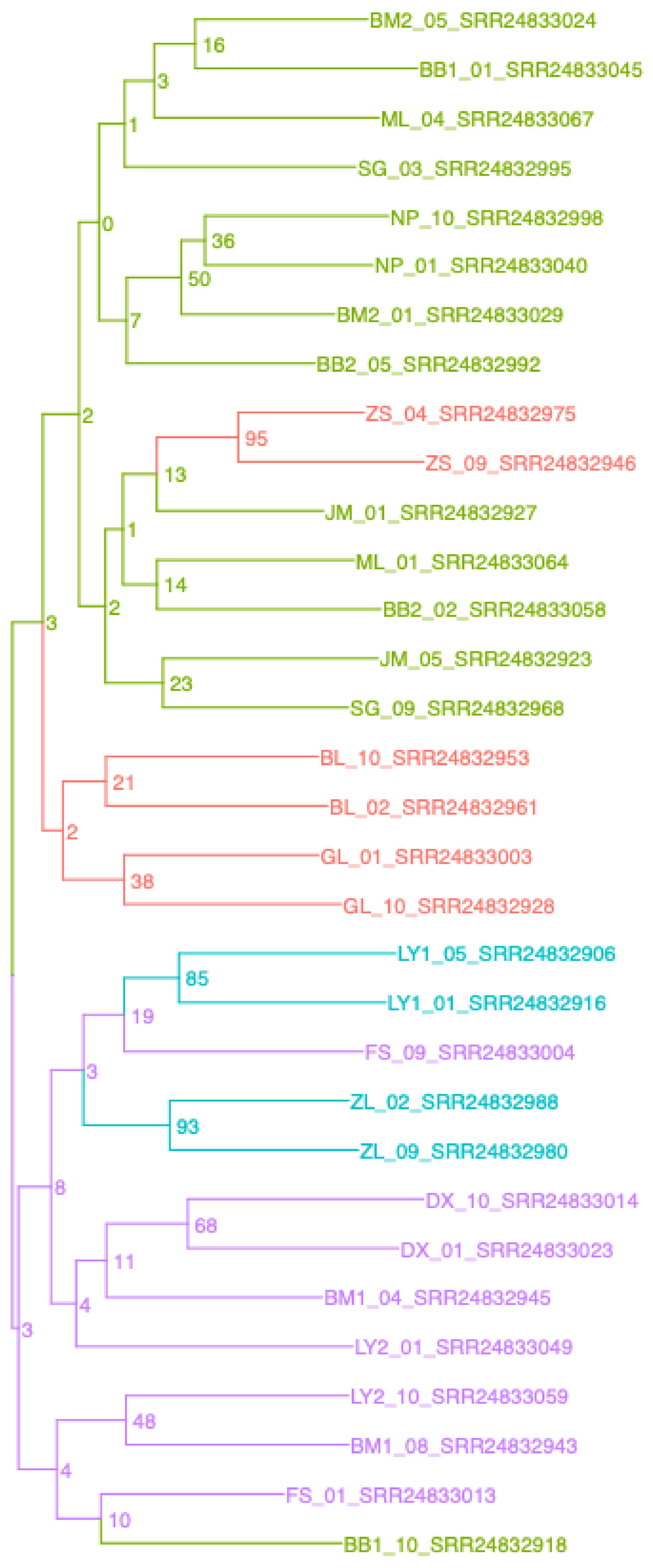


Malania – Malania_BUSCOsuper

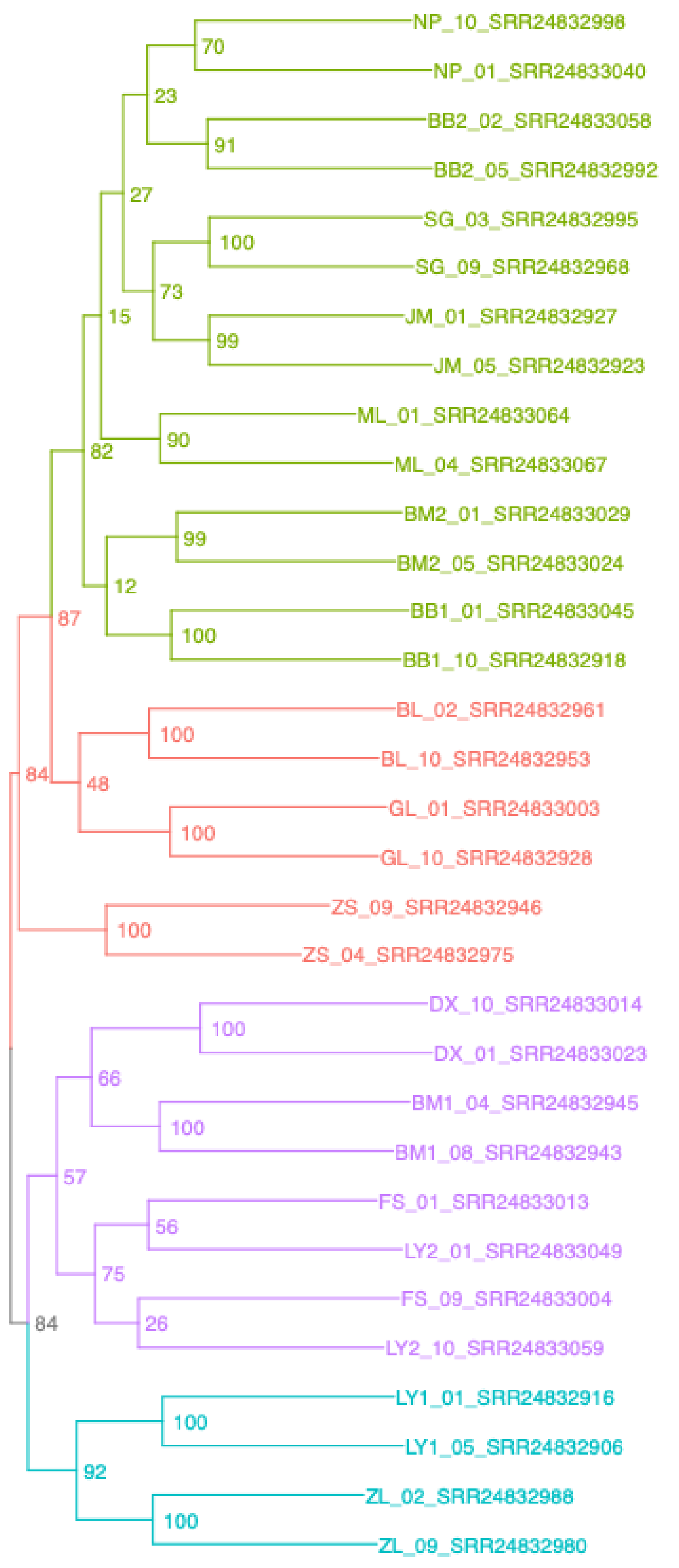


Malania – Malania_SCsuper

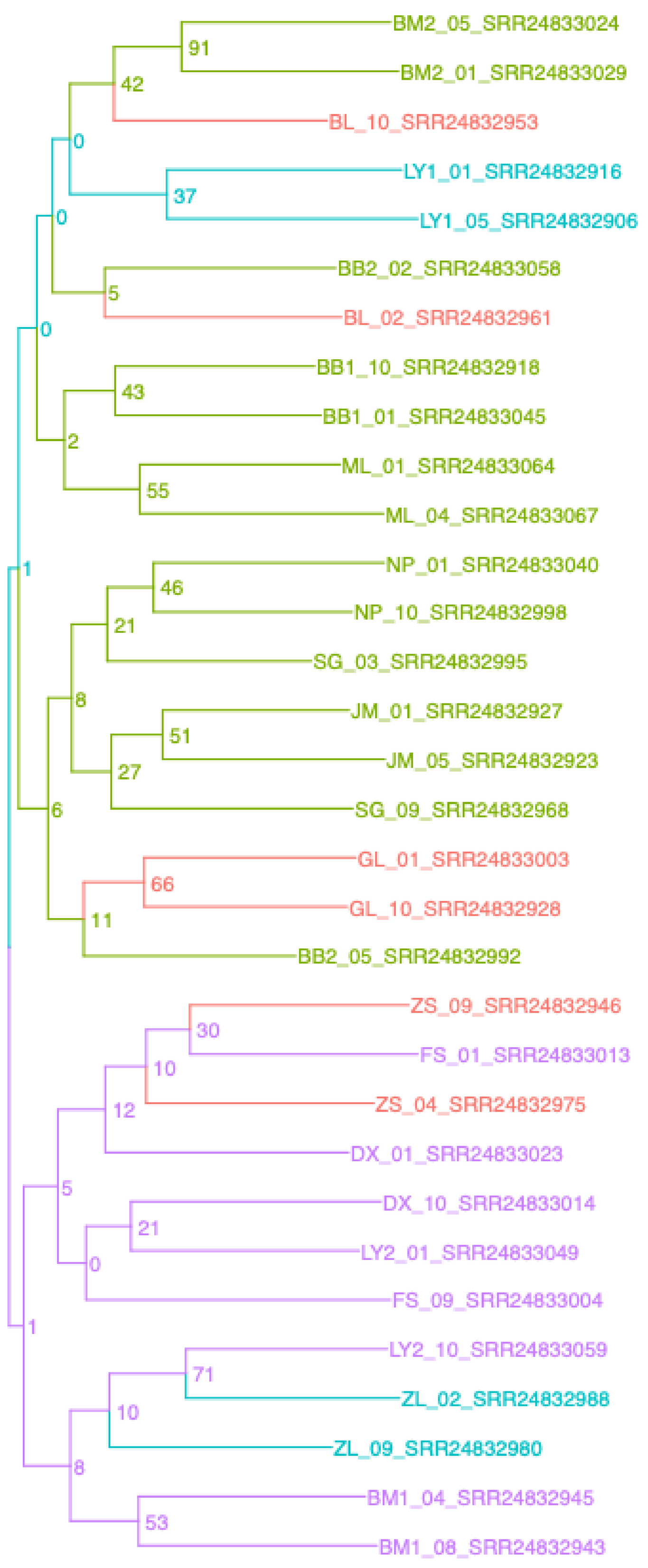


Malania – Malania_genome

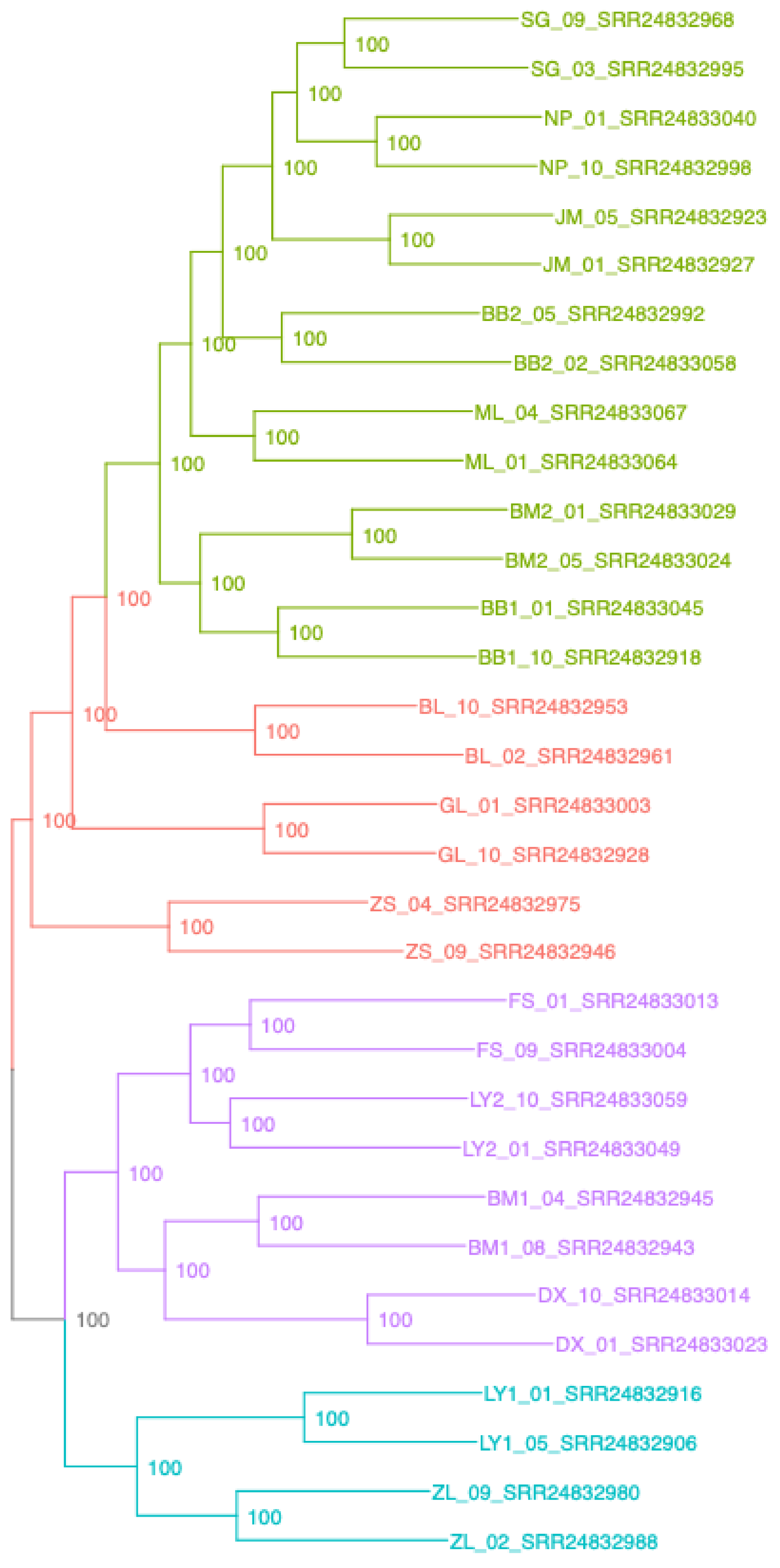

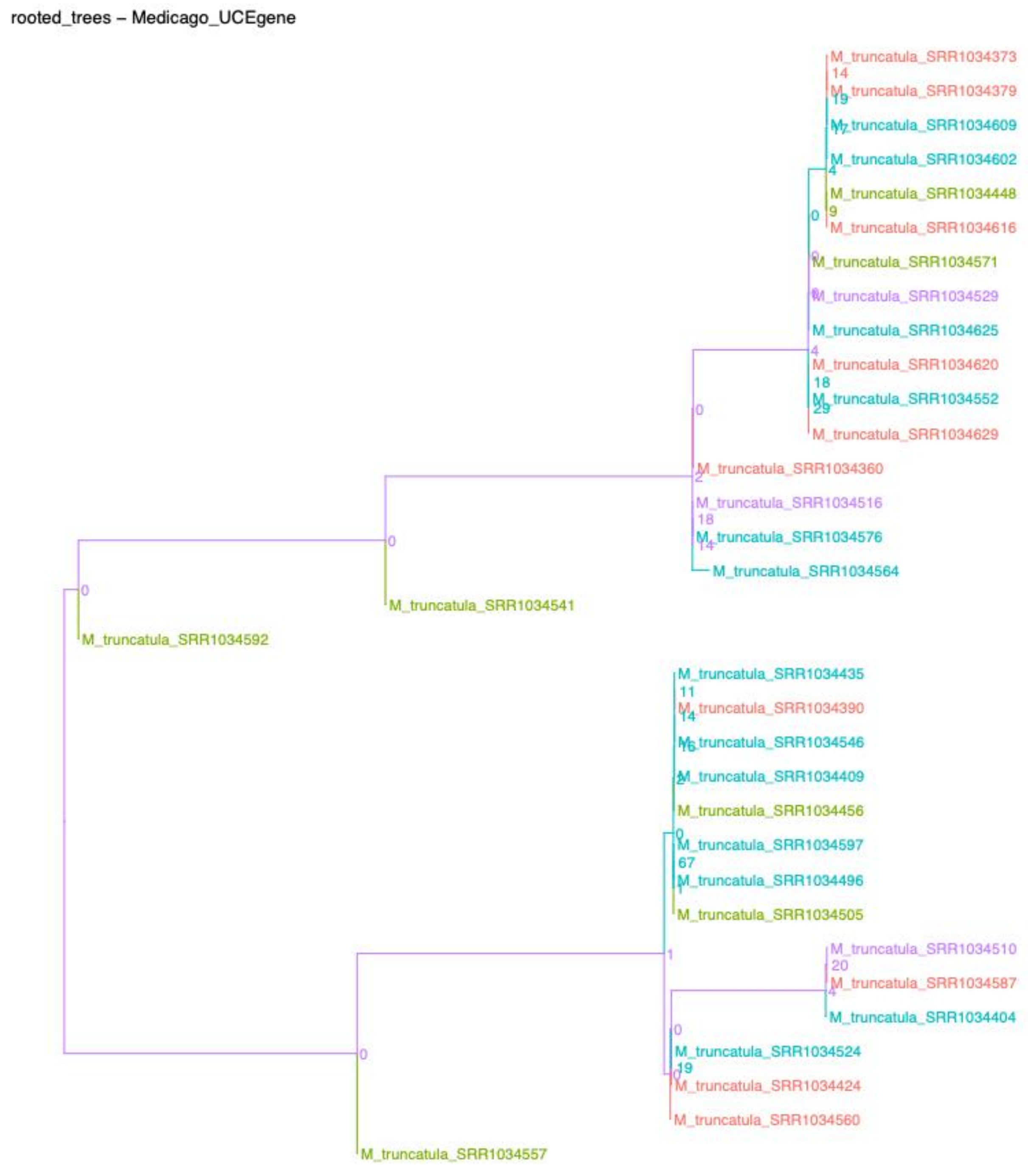
rooted_trees – Medicago_UCEgene

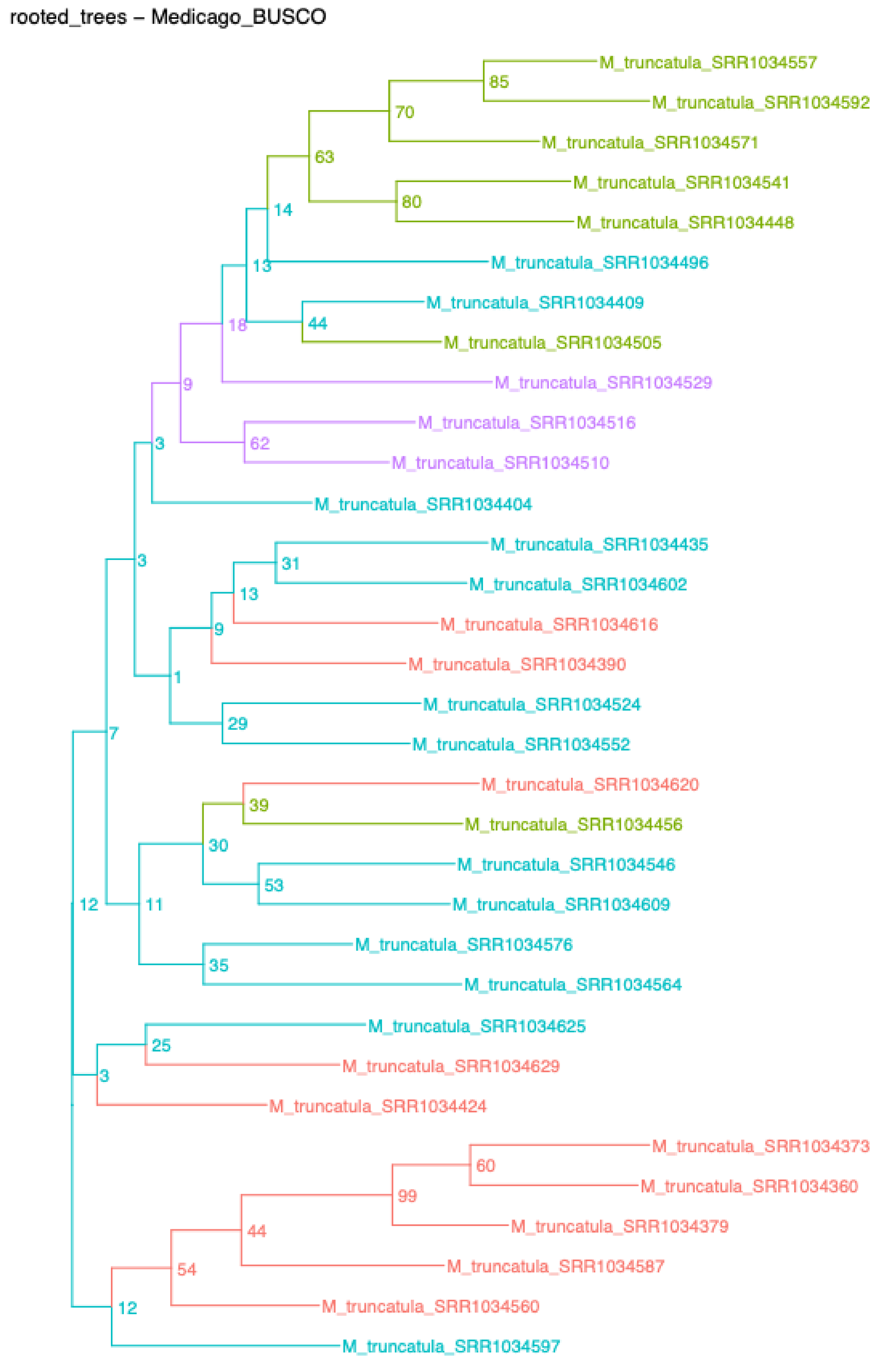
rooted_trees – Medicago_BUSCO

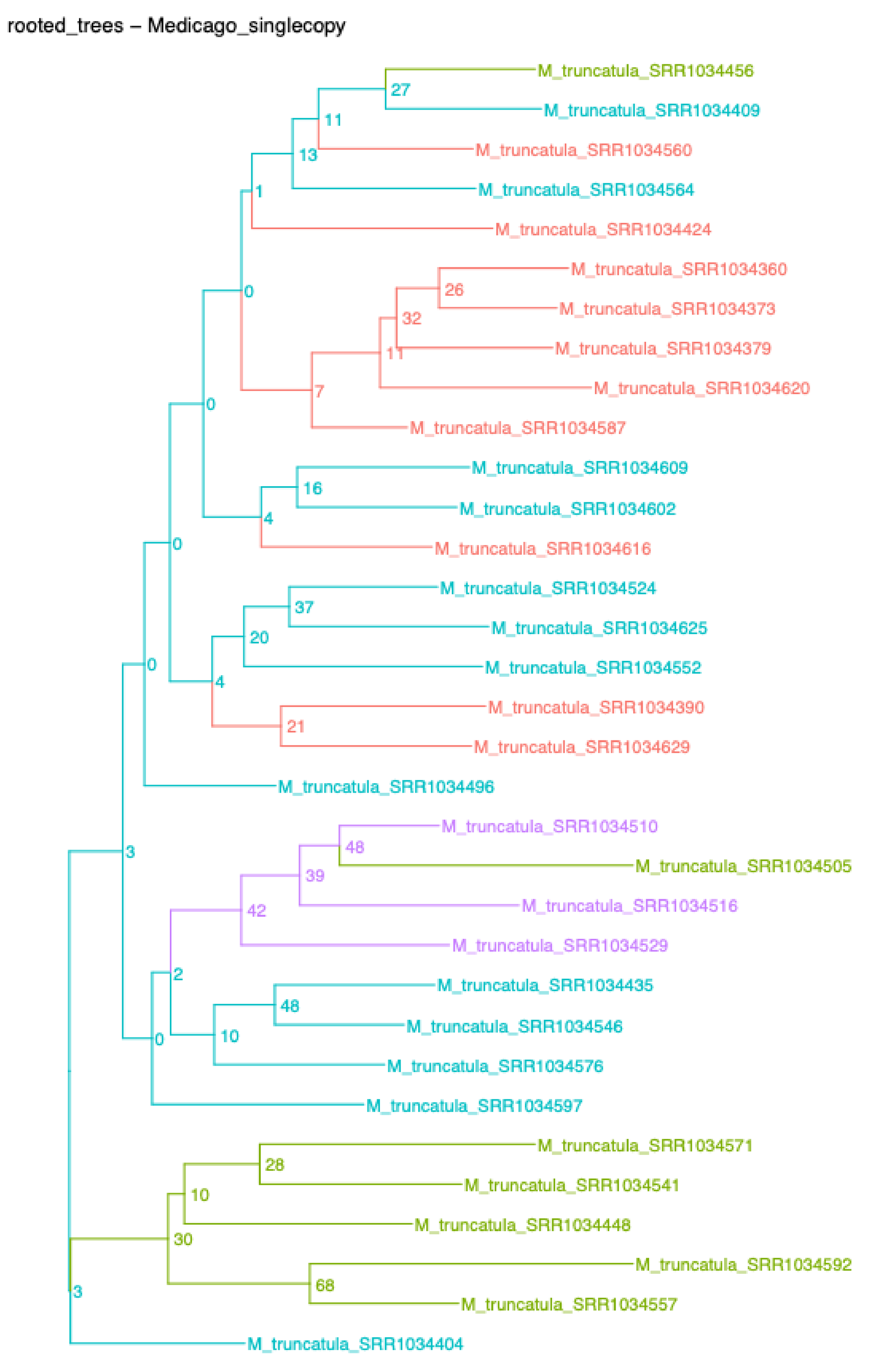
rooted_trees – Medicago_singlecopy

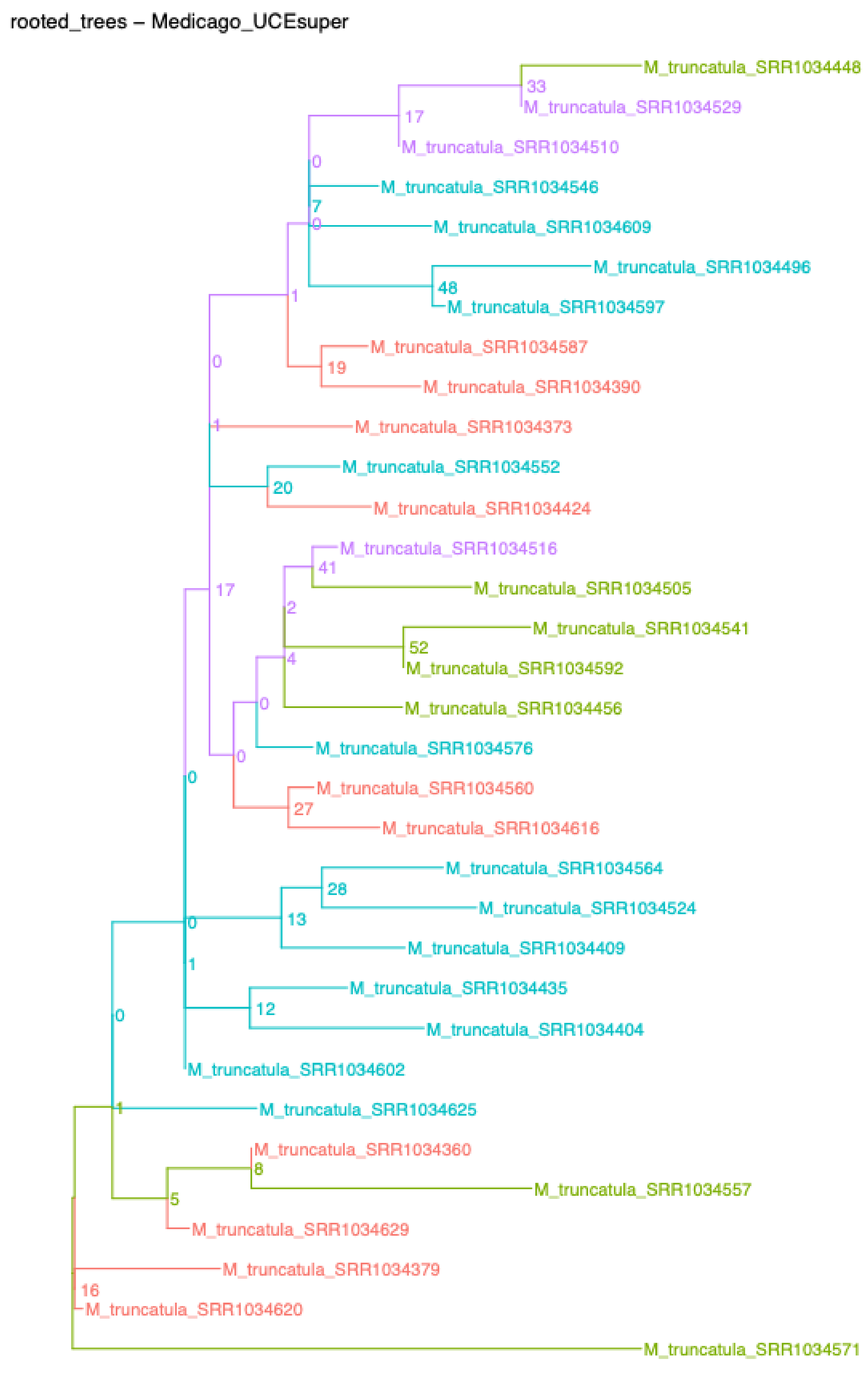
rooted_trees – Medicago_UCEsuper

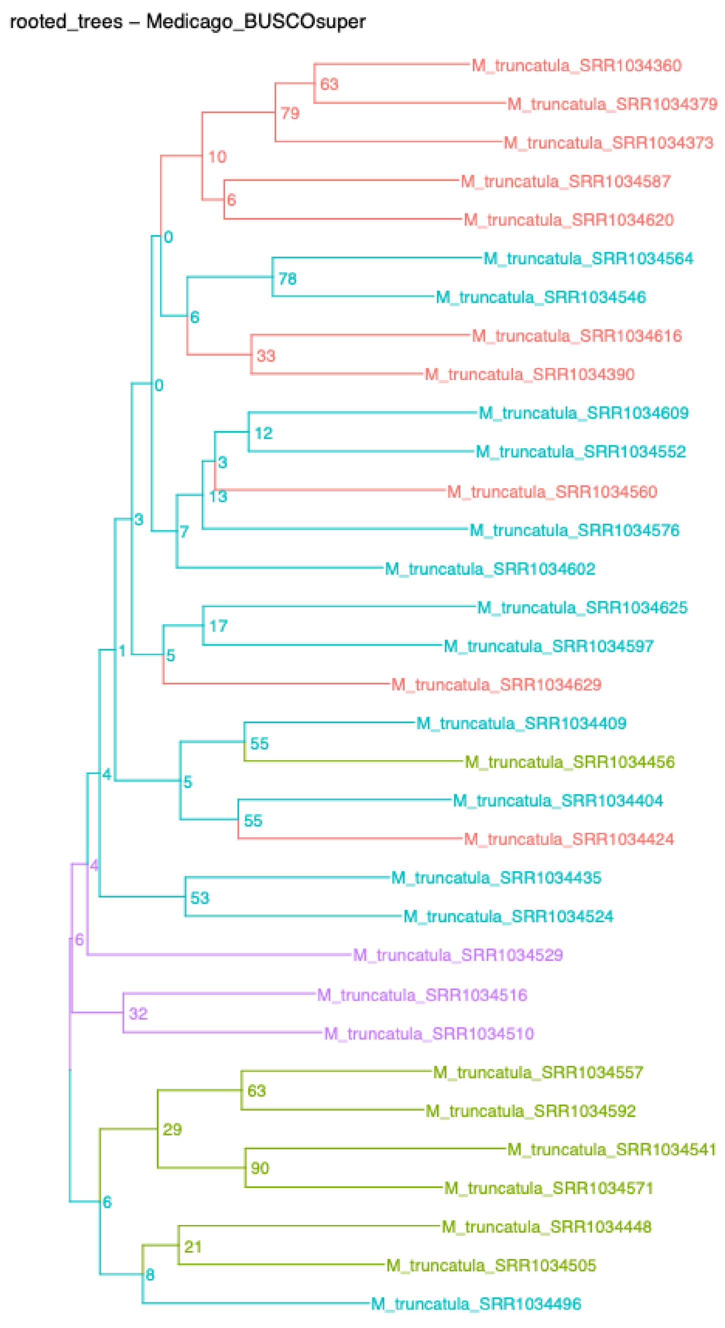
rooted_trees – Medicago_BUSCOsuper

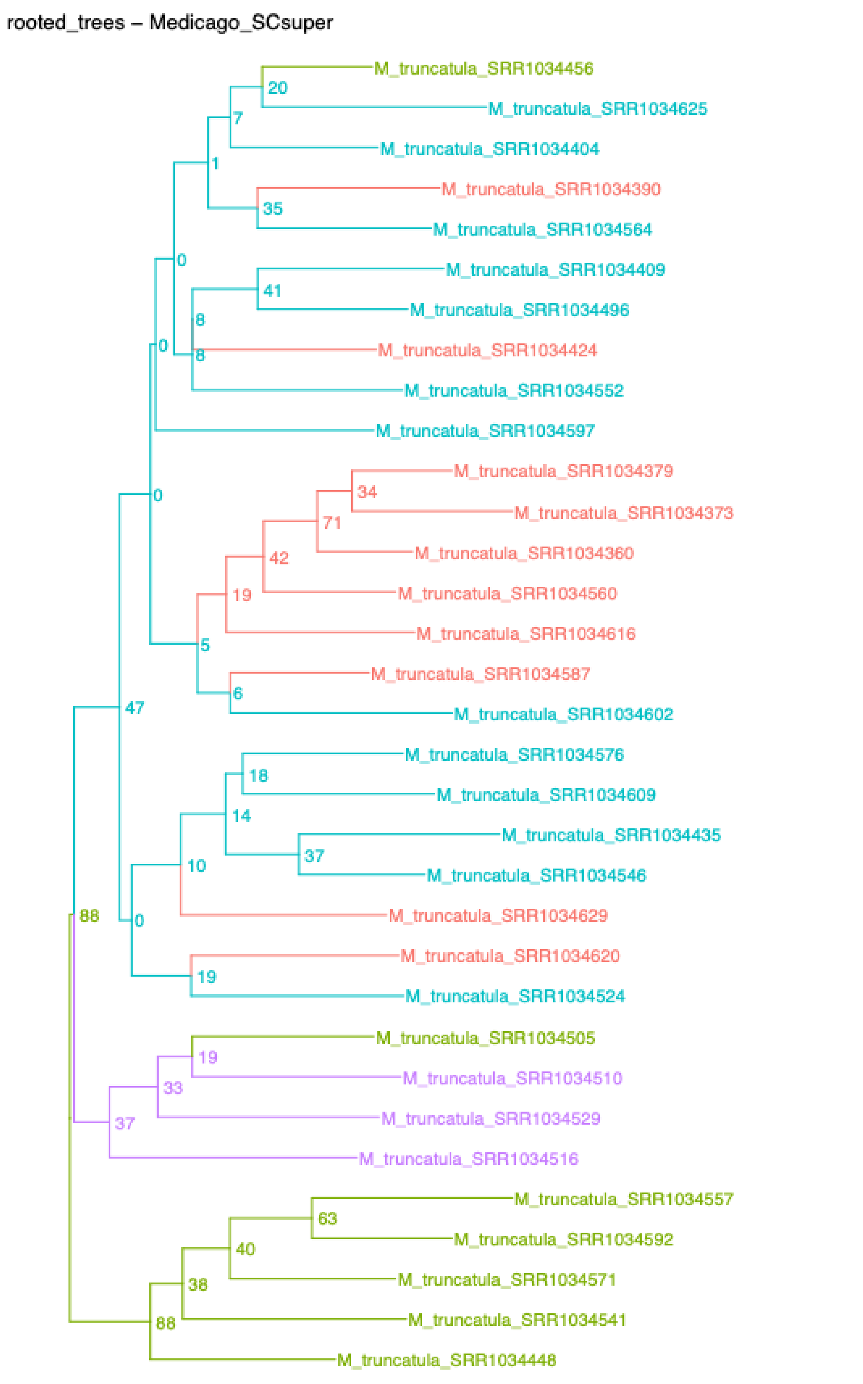
rooted_trees – Medicago_SCsuper

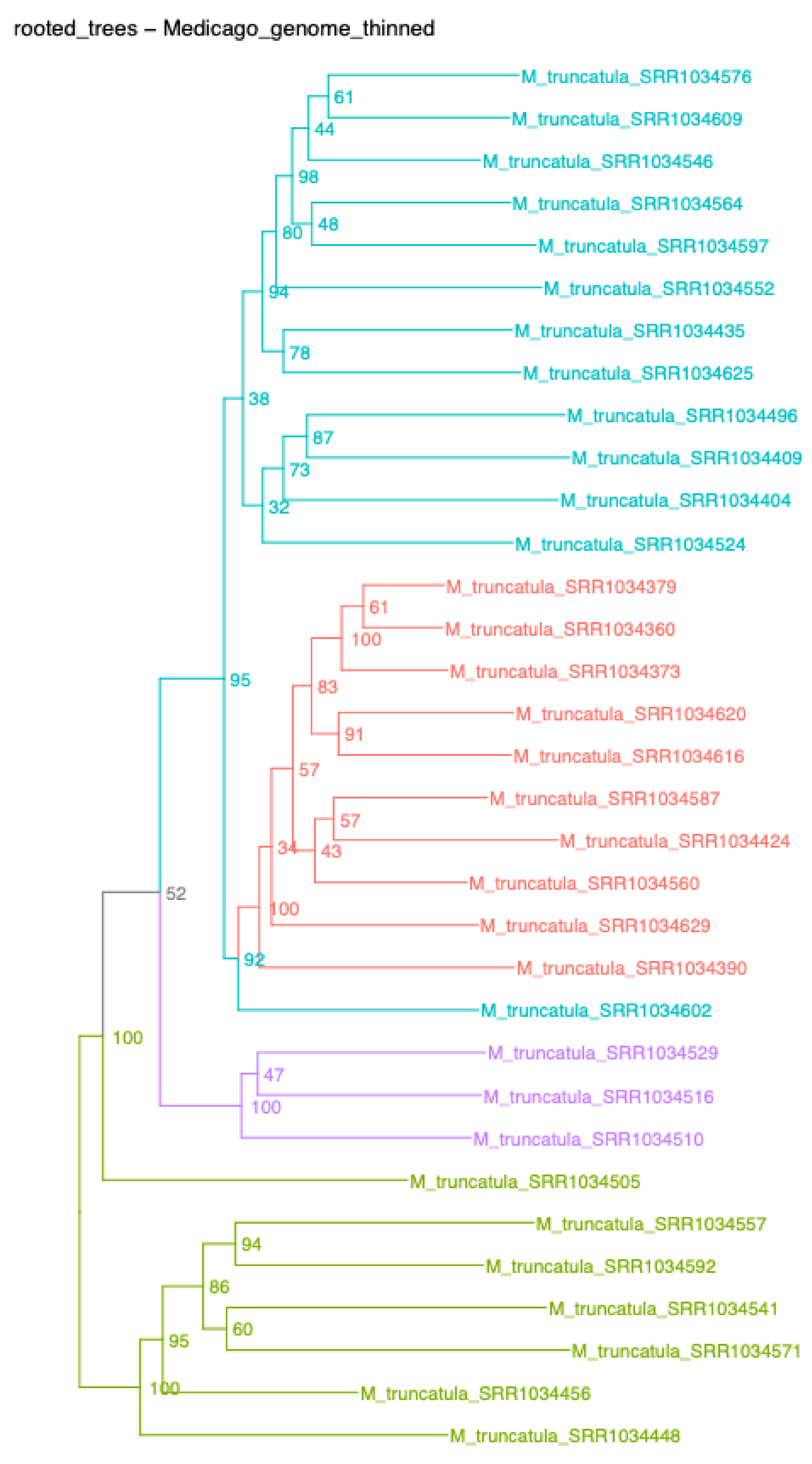
rooted_trees – Medicago_genome_thinned

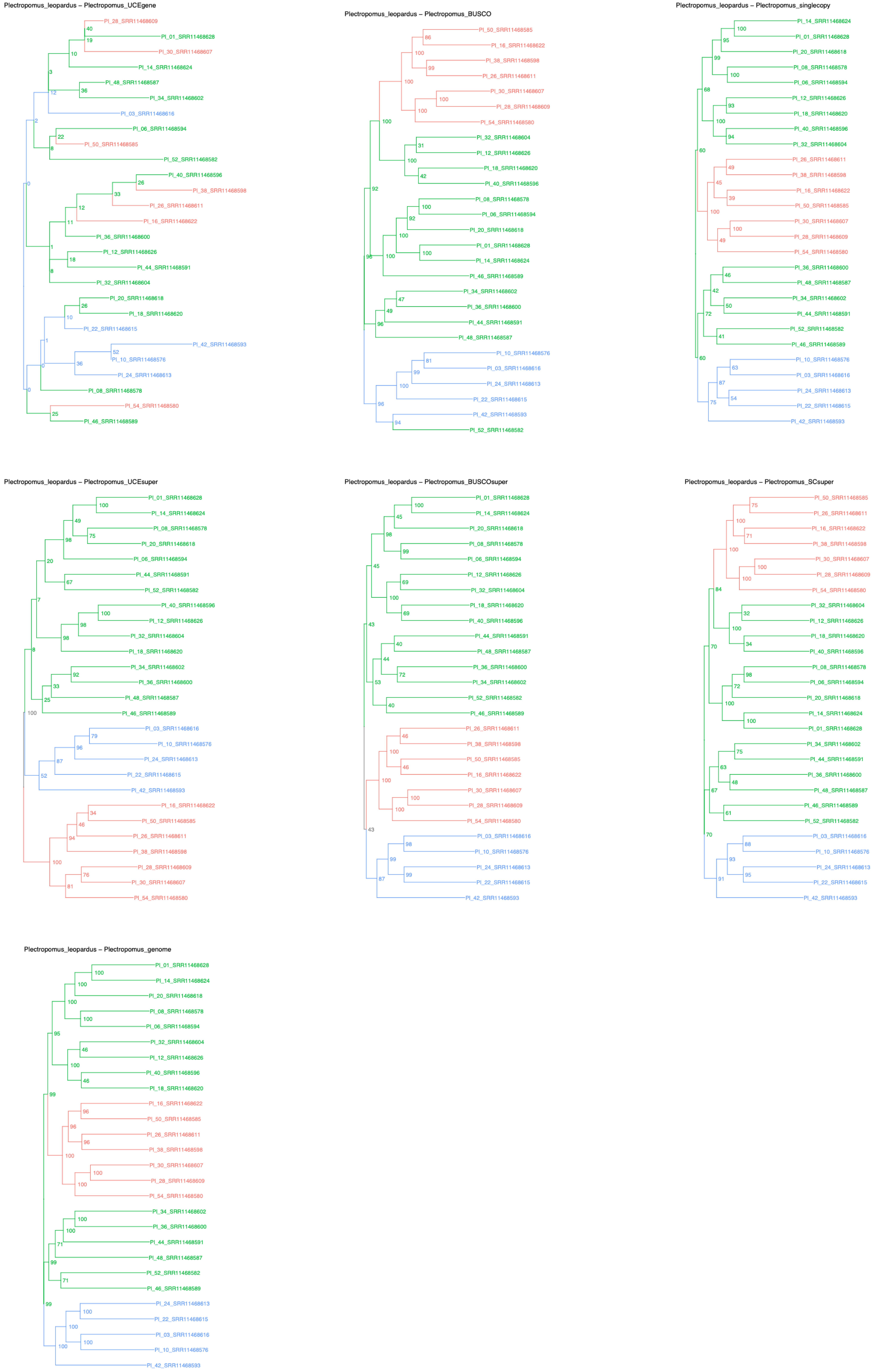
Plectropomus_leopardus – Plectropomus_UCEgene
Pl_28_SRR11468609
Pl_01_SRR11468628
Pl_30_SRR11468607
Pl_14_SRR11468624
Pl_48_SRR11468587
Pl_34_SRR11468602
Pl_03_SRR11468616
Pl_06_SRR11468594
Pl_50_SRR11468585
Pl_52_SRR11468582
Pl_40_SRR11468596
Pl_38_SRR11468598
Pl_26_SRR11468611
Pl_16_SRR11468622
Pl_36_SRR11468600
Pl_12_SRR11468626
Pl_44_SRR11468591
Pl_32_SRR11468604
Pl_20_SRR11468618
Pl_18_SRR11468620
Pl_22_SRR11468615
Pl_42_SRR11468593
Pl_10_SRR11468576
Pl_24_SRR11468613
Pl_08_SRR11468578
Pl_54_SRR11468580
Pl_46_SRR11468589
Plectropomus_leopardus – Plectropomus_BUSCO
Pl_50_SRR11468585
Pl_16_SRR11468622
Pl_38_SRR11468598
Pl_26_SRR11468611
Pl_30_SRR11468607
Pl_28_SRR11468609
Pl_54_SRR11468580
Pl_32_SRR11468604
Pl_12_SRR11468626
Pl_18_SRR11468620
Pl_40_SRR11468596
Pl_08_SRR11468578
Pl_06_SRR11468594
Pl_20_SRR11468618
Pl_01_SRR11468628
Pl_14_SRR11468624
Pl_46_SRR11468589
Pl_34_SRR11468602
Pl_36_SRR11468600
Pl_44_SRR11468591
Pl_48_SRR11468587
Pl_10_SRR11468576
Pl_03_SRR11468616
Pl_24_SRR11468613
Pl_22_SRR11468615
Pl_42_SRR11468593
Pl_52_SRR11468582
Plectropomus_leopardus – Plectropomus_singlecopy
Pl_14_SRR11468624
Pl_01_SRR11468628
Pl_20_SRR11468618
Pl_08_SRR11468578
Pl_06_SRR11468594
Pl_12_SRR11468626
Pl_18_SRR11468620
Pl_40_SRR11468596
Pl_32_SRR11468604
Pl_26_SRR11468611
Pl_38_SRR11468598
Pl_16_SRR11468622
Pl_50_SRR11468585
Pl_30_SRR11468607
Pl_28_SRR11468609
Pl_54_SRR11468580
Pl_36_SRR11468600
Pl_48_SRR11468587
Pl_34_SRR11468602
Pl_44_SRR11468591
Pl_52_SRR11468582
Pl_46_SRR11468589
Pl_10_SRR11468576
Pl_03_SRR11468616
Pl_24_SRR11468613
Pl_22_SRR11468615
Pl_42_SRR11468593
Plectropomus_leopardus – Plectropomus_UCEsuper
Pl_01_SRR11468628
Pl_14_SRR11468624
Pl_08_SRR11468578
Pl_20_SRR11468618
Pl_06_SRR11468594
Pl_44_SRR11468591
Pl_52_SRR11468582
Pl_40_SRR11468596
Pl_12_SRR11468626
Pl_32_SRR11468604
Pl_18_SRR11468620
Pl_34_SRR11468602
Pl_36_SRR11468600
Pl_48_SRR11468587
Pl_46_SRR11468589
Pl_03_SRR11468616
Pl_10_SRR11468576
Pl_24_SRR11468613
Pl_22_SRR11468615
Pl_42_SRR11468593
Pl_16_SRR11468622
Pl_50_SRR11468585
Pl_26_SRR11468611
Pl_38_SRR11468598
Pl_28_SRR11468609
Pl_30_SRR11468607
Pl_54_SRR11468580
Plectropomus_leopardus – Plectropomus_BUSCOsuper
Pl_01_SRR11468628
Pl_14_SRR11468624
Pl_20_SRR11468618
Pl_08_SRR11468578
Pl_06_SRR11468594
Pl_12_SRR11468626
Pl_32_SRR11468604
Pl_18_SRR11468620
Pl_40_SRR11468596
Pl_44_SRR11468591
Pl_48_SRR11468587
Pl_36_SRR11468600
Pl_34_SRR11468602
Pl_52_SRR11468582
Pl_46_SRR11468589
Pl_26_SRR11468611
Pl_38_SRR11468598
Pl_50_SRR11468585
Pl_16_SRR11468622
Pl_30_SRR11468607
Pl_28_SRR11468609
Pl_54_SRR11468580
Pl_03_SRR11468616
Pl_10_SRR11468576
Pl_24_SRR11468613
Pl_22_SRR11468615
Pl_42_SRR11468593
Plectropomus_leopardus – Plectropomus_SCsuper
Pl_50_SRR11468585
Pl_26_SRR11468611
Pl_16_SRR11468622
Pl_38_SRR11468598
Pl_30_SRR11468607
Pl_28_SRR11468609
Pl_54_SRR11468580
Pl_32_SRR11468604
Pl_12_SRR11468626
Pl_18_SRR11468620
Pl_40_SRR11468596
Pl_08_SRR11468578
Pl_06_SRR11468594
Pl_20_SRR11468618
Pl_14_SRR11468624
Pl_01_SRR11468628
Pl_34_SRR11468602
Pl_44_SRR11468591
Pl_36_SRR11468600
Pl_48_SRR11468587
Pl_46_SRR11468589
Pl_52_SRR11468582
Pl_03_SRR11468616
Pl_10_SRR11468576
Pl_24_SRR11468613
Pl_22_SRR11468615
Pl_42_SRR11468593
Plectropomus_leopardus – Plectropomus_genome
Pl_01_SRR11468628
Pl_14_SRR11468624
Pl_20_SRR11468618
Pl_08_SRR11468578
Pl_06_SRR11468594
Pl_32_SRR11468604
Pl_12_SRR11468626
Pl_40_SRR11468596
Pl_18_SRR11468620
Pl_16_SRR11468622
Pl_50_SRR11468585
Pl_26_SRR11468611
Pl_38_SRR11468598
Pl_30_SRR11468607
Pl_28_SRR11468609
Pl_54_SRR11468580
Pl_34_SRR11468602
Pl_36_SRR11468600
Pl_44_SRR11468591
Pl_48_SRR11468587
Pl_52_SRR11468582
Pl_46_SRR11468589
Pl_24_SRR11468613
Pl_22_SRR11468615
Pl_03_SRR11468616
Pl_10_SRR11468576
Pl_42_SRR11468593

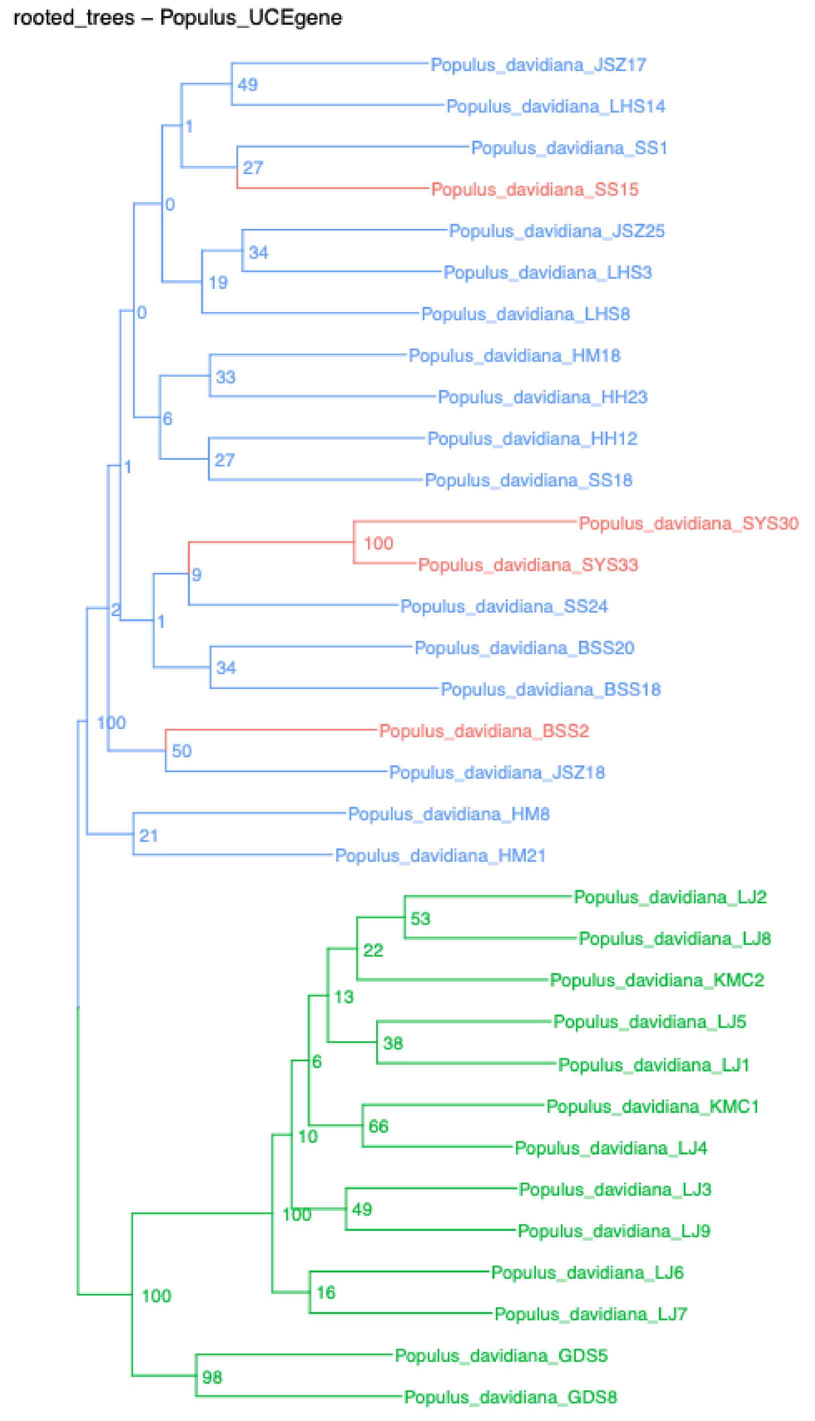
rooted_trees – Populus_UCEgene
Populus_davidiana_JSZ17
Populus_davidiana_LHS14
Populus_davidiana_SS1
Populus_davidiana_SS15
Populus_davidiana_JSZ25
Populus_davidiana_LHS3
Populus_davidiana_LHS8
Populus_davidiana_HM18
Populus_davidiana_HH23
Populus_davidiana_HH12
Populus_davidiana_SS18
Populus_davidiana_SYS30
Populus_davidiana_SYS33
Populus_davidiana_SS24
Populus_davidiana_BSS20
Populus_davidiana_BSS18
Populus_davidiana_BSS2
Populus_davidiana_JSZ18
Populus_davidiana_HM8
Populus_davidiana_HM21
Populus_davidiana_LJ2
Populus_davidiana_LJ8
Populus_davidiana_KMC2
Populus_davidiana_LJ5
Populus_davidiana_LJ1
Populus_davidiana_KMC1
Populus_davidiana_LJ4
Populus_davidiana_LJ3
Populus_davidiana_LJ9
Populus_davidiana_LJ6
Populus_davidiana_LJ7
Populus_davidiana_GDS5
Populus_davidiana_GDS8

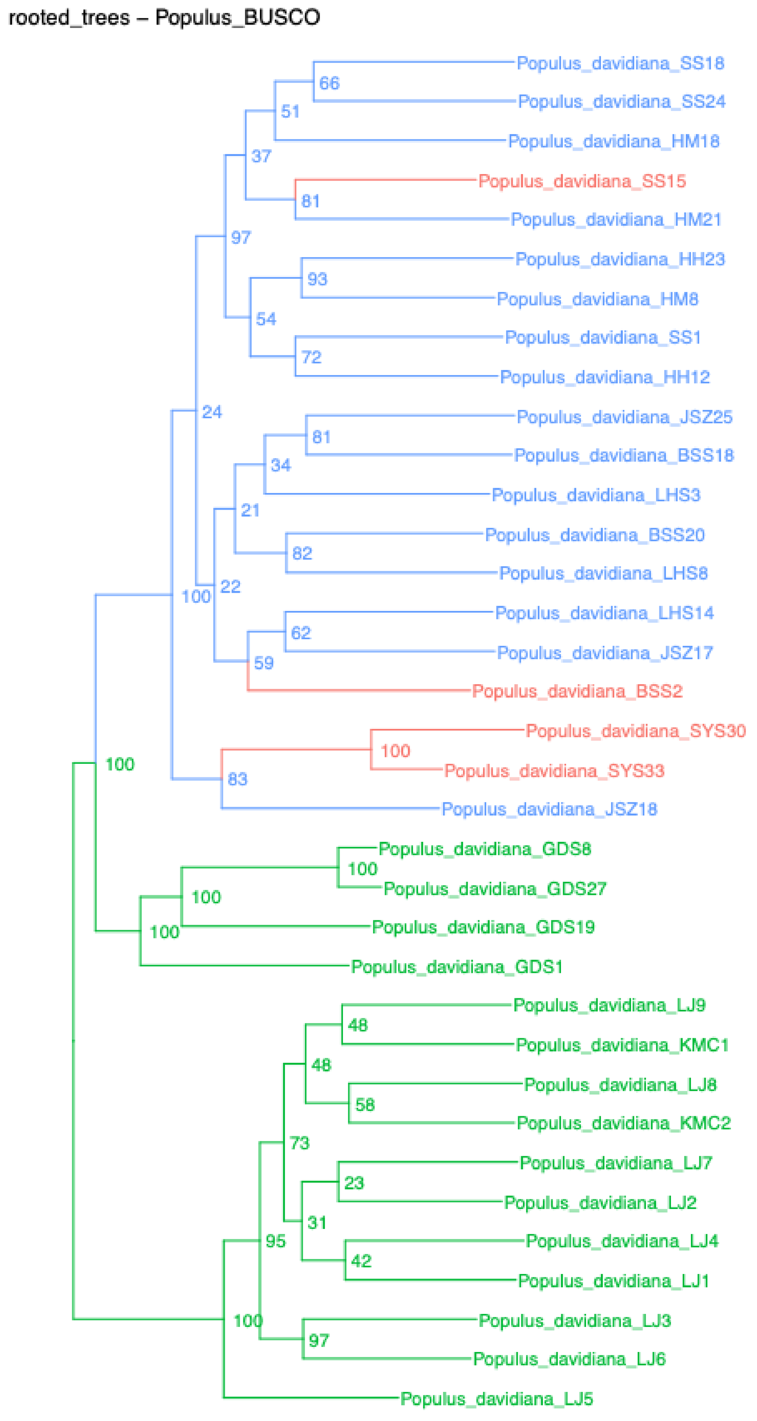
rooted_trees – Populus_BUSCO
Populus_davidiana_SS18
Populus_davidiana_SS24
Populus_davidiana_HM18
Populus_davidiana_SS15
Populus_davidiana_HM21
Populus_davidiana_HH23
Populus_davidiana_HM8
Populus_davidiana_SS1
Populus_davidiana_HH12
Populus_davidiana_JSZ25
Populus_davidiana_BSS18
Populus_davidiana_LHS3
Populus_davidiana_BSS20
Populus_davidiana_LHS8
Populus_davidiana_LHS14
Populus_davidiana_JSZ17
Populus_davidiana_BSS2
Populus_davidiana_SYS30
Populus_davidiana_SYS33
Populus_davidiana_JSZ18
Populus_davidiana_GDS8
Populus_davidiana_GDS27
Populus_davidiana_GDS19
Populus_davidiana_GDS1
Populus_davidiana_LJ9
Populus_davidiana_KMC1
Populus_davidiana_LJ8
Populus_davidiana_KMC2
Populus_davidiana_LJ7
Populus_davidiana_LJ2
Populus_davidiana_LJ4
Populus_davidiana_LJ1
Populus_davidiana_LJ3
Populus_davidiana_LJ6
Populus_davidiana_LJ5

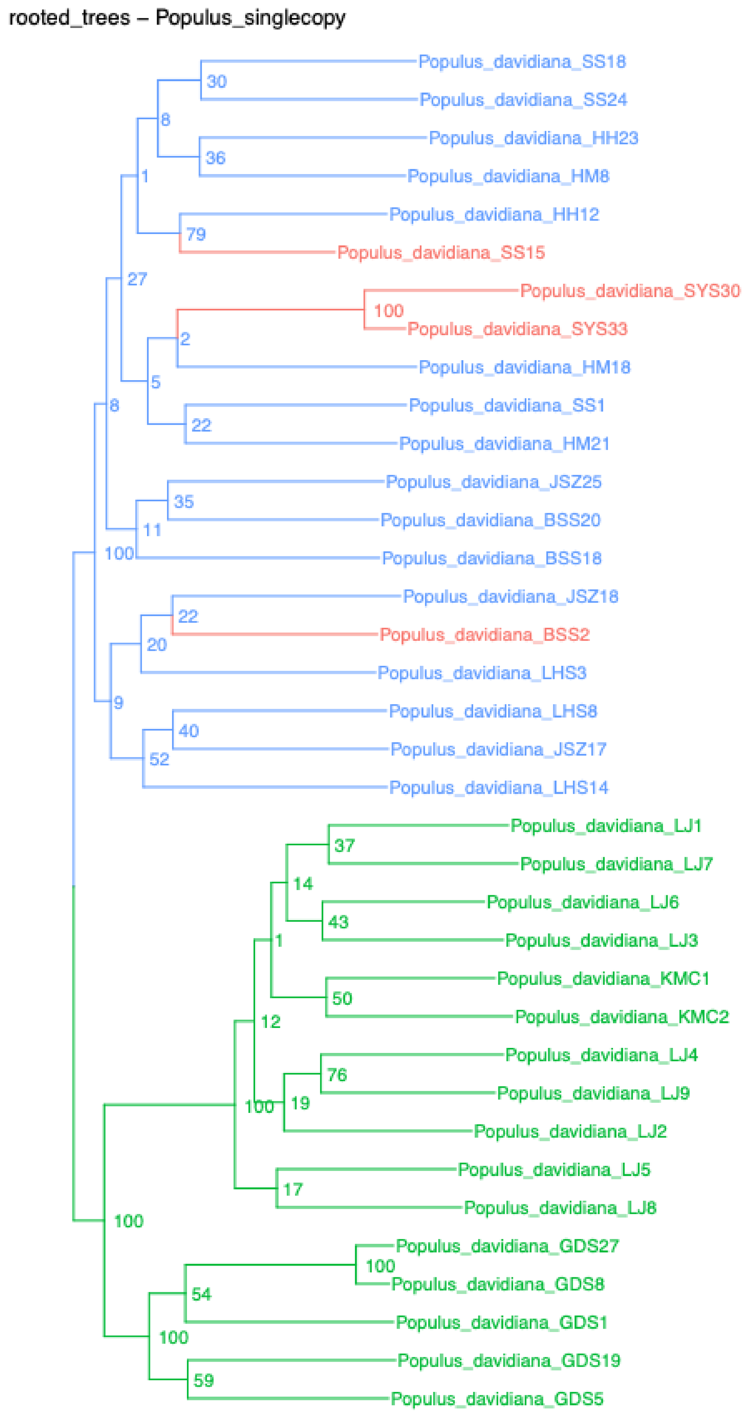
rooted_trees – Populus_singlecopy
Populus_davidiana_SS18
Populus_davidiana_SS24
Populus_davidiana_HH23
Populus_davidiana_HM8
Populus_davidiana_HH12
Populus_davidiana_SS15
Populus_davidiana_SYS30
Populus_davidiana_SYS33
Populus_davidiana_HM18
Populus_davidiana_SS1
Populus_davidiana_HM21
Populus_davidiana_JSZ25
Populus_davidiana_BSS20
Populus_davidiana_BSS18
Populus_davidiana_JSZ18
Populus_davidiana_BSS2
Populus_davidiana_LHS3
Populus_davidiana_LHS8
Populus_davidiana_JSZ17
Populus_davidiana_LHS14
Populus_davidiana_LJ1
Populus_davidiana_LJ7
Populus_davidiana_LJ6
Populus_davidiana_LJ3
Populus_davidiana_KMC1
Populus_davidiana_KMC2
Populus_davidiana_LJ4
Populus_davidiana_LJ9
Populus_davidiana_LJ2
Populus_davidiana_LJ5
Populus_davidiana_LJ8
Populus_davidiana_GDS27
Populus_davidiana_GDS8
Populus_davidiana_GDS1
Populus_davidiana_GDS19
Populus_davidiana_GDS5

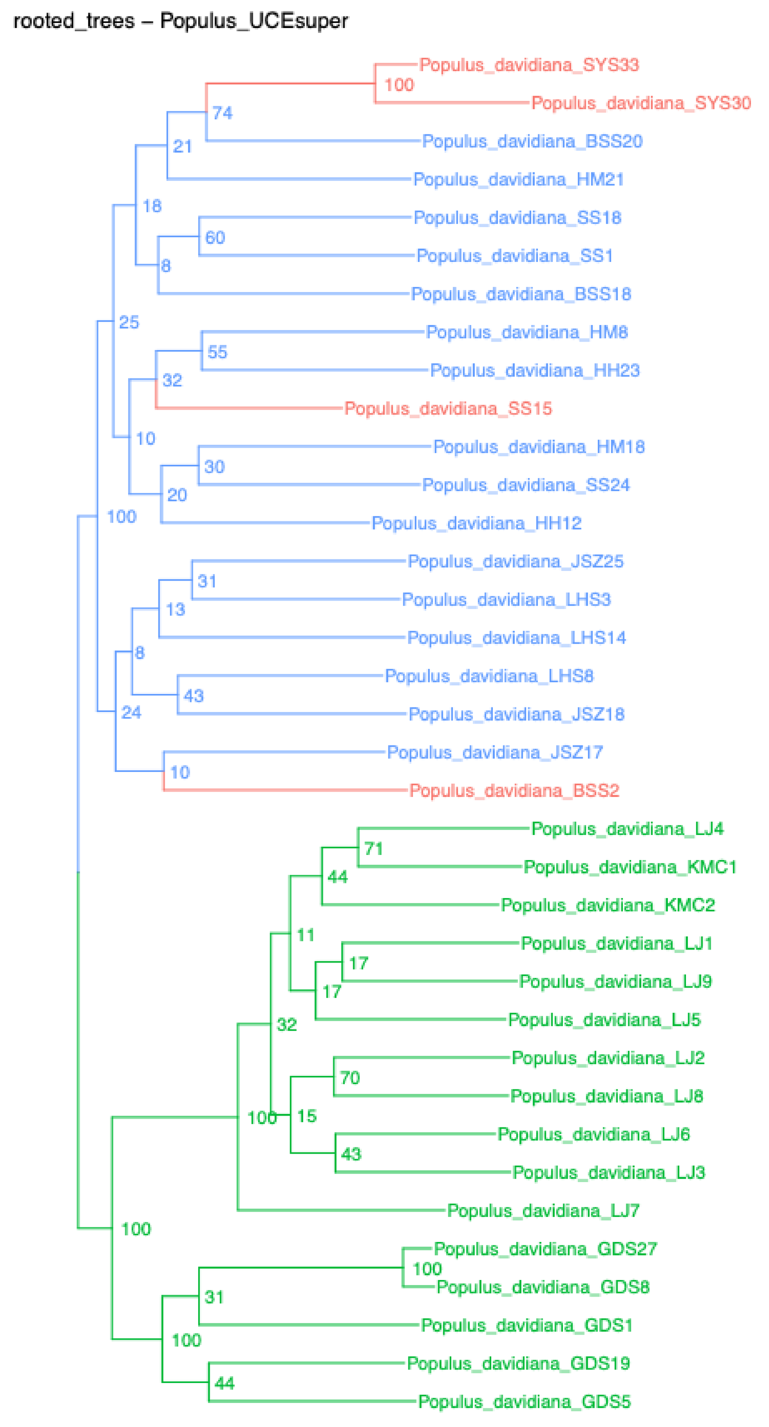
rooted_trees – Populus_UCEsuper
Populus_davidiana_SYS33
Populus_davidiana_SYS30
Populus_davidiana_BSS20
Populus_davidiana_HM21
Populus_davidiana_SS18
Populus_davidiana_SS1
Populus_davidiana_BSS18
Populus_davidiana_HM8
Populus_davidiana_HH23
Populus_davidiana_SS15
Populus_davidiana_HM18
Populus_davidiana_SS24
Populus_davidiana_HH12
Populus_davidiana_JSZ25
Populus_davidiana_LHS3
Populus_davidiana_LHS14
Populus_davidiana_LHS8
Populus_davidiana_JSZ18
Populus_davidiana_JSZ17
Populus_davidiana_BSS2
Populus_davidiana_LJ4
Populus_davidiana_KMC1
Populus_davidiana_KMC2
Populus_davidiana_LJ1
Populus_davidiana_LJ9
Populus_davidiana_LJ5
Populus_davidiana_LJ2
Populus_davidiana_LJ8
Populus_davidiana_LJ6
Populus_davidiana_LJ3
Populus_davidiana_LJ7
Populus_davidiana_GDS27
Populus_davidiana_GDS8
Populus_davidiana_GDS1
Populus_davidiana_GDS19
Populus_davidiana_GDS5

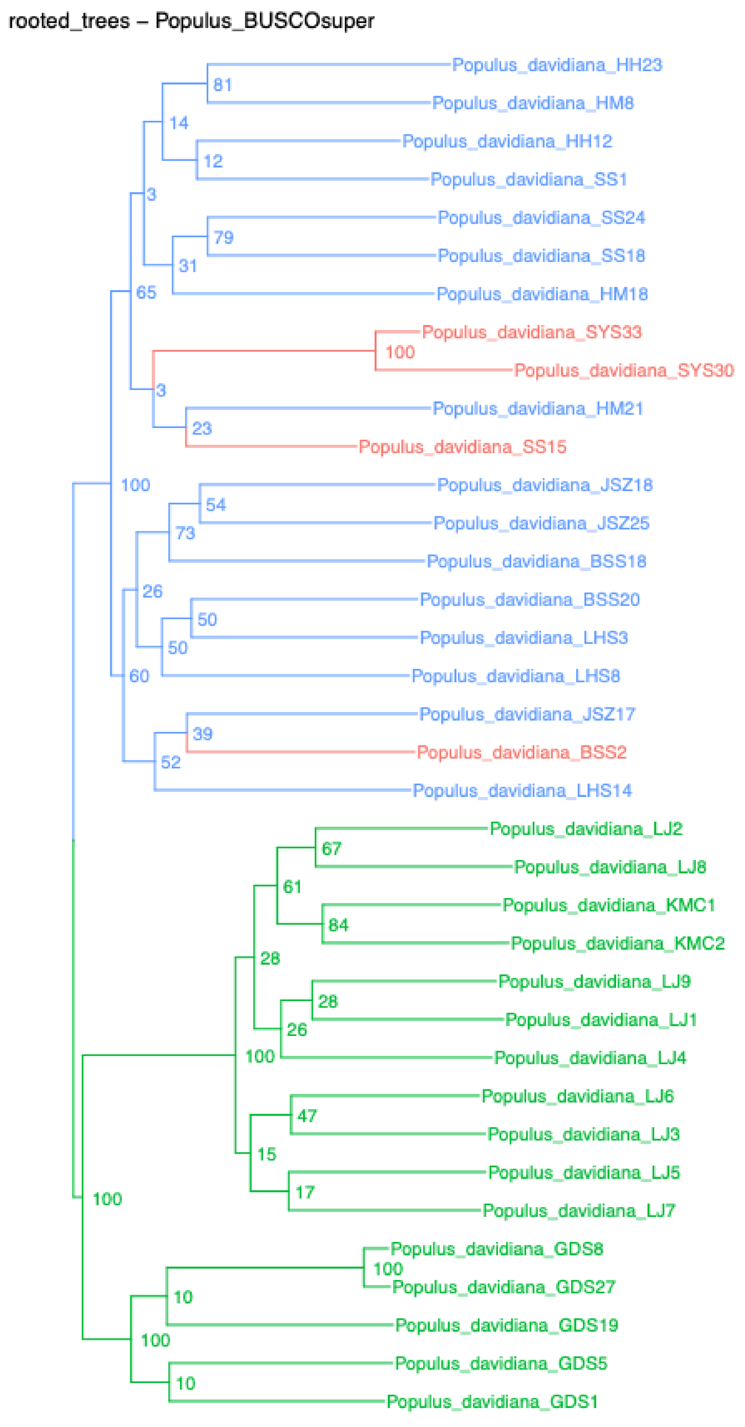
rooted_trees – Populus_BUSCOsuper
Populus_davidiana_HH23
Populus_davidiana_HM8
Populus_davidiana_HH12
Populus_davidiana_SS1
Populus_davidiana_SS24
Populus_davidiana_SS18
Populus_davidiana_HM18
Populus_davidiana_SYS33
Populus_davidiana_SYS30
Populus_davidiana_HM21
Populus_davidiana_SS15
Populus_davidiana_JSZ18
Populus_davidiana_JSZ25
Populus_davidiana_BSS18
Populus_davidiana_BSS20
Populus_davidiana_LHS3
Populus_davidiana_LHS8
Populus_davidiana_JSZ17
Populus_davidiana_BSS2
Populus_davidiana_LHS14
Populus_davidiana_LJ2
Populus_davidiana_LJ8
Populus_davidiana_KMC1
Populus_davidiana_KMC2
Populus_davidiana_LJ9
Populus_davidiana_LJ1
Populus_davidiana_LJ4
Populus_davidiana_LJ6
Populus_davidiana_LJ3
Populus_davidiana_LJ5
Populus_davidiana_LJ7
Populus_davidiana_GDS8
Populus_davidiana_GDS27
Populus_davidiana_GDS19
Populus_davidiana_GDS5
Populus_davidiana_GDS1

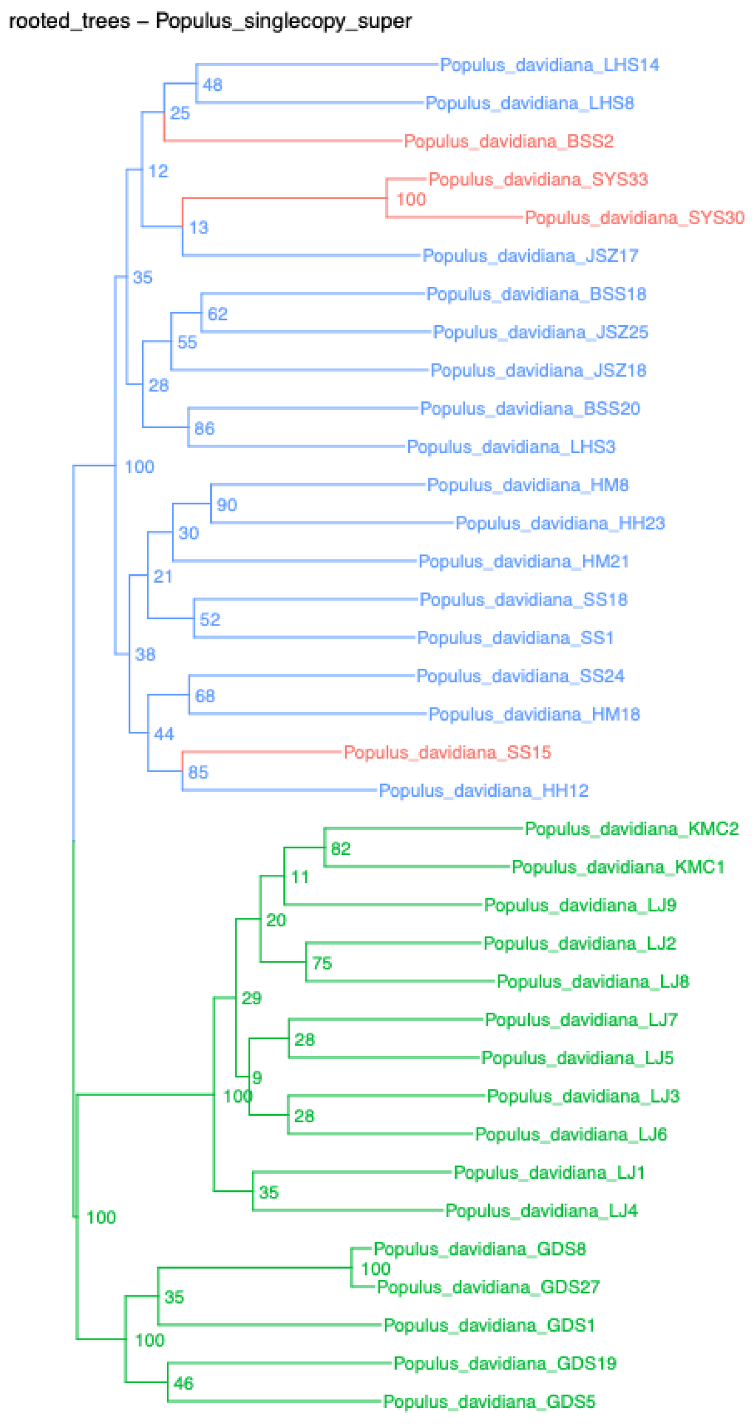
rooted_trees – Populus_singlecopy_super
Populus_davidiana_LHS14
Populus_davidiana_LHS8
Populus_davidiana_BSS2
Populus_davidiana_SYS33
Populus_davidiana_SYS30
Populus_davidiana_JSZ17
Populus_davidiana_BSS18
Populus_davidiana_JSZ25
Populus_davidiana_JSZ18
Populus_davidiana_BSS20
Populus_davidiana_LHS3
Populus_davidiana_HM8
Populus_davidiana_HH23
Populus_davidiana_HM21
Populus_davidiana_SS18
Populus_davidiana_SS1
Populus_davidiana_SS24
Populus_davidiana_HM18
Populus_davidiana_SS15
Populus_davidiana_HH12
Populus_davidiana_KMC2
Populus_davidiana_KMC1
Populus_davidiana_LJ9
Populus_davidiana_LJ2
Populus_davidiana_LJ8
Populus_davidiana_LJ7
Populus_davidiana_LJ5
Populus_davidiana_LJ3
Populus_davidiana_LJ6
Populus_davidiana_LJ1
Populus_davidiana_LJ4
Populus_davidiana_GDS8
Populus_davidiana_GDS27
Populus_davidiana_GDS1
Populus_davidiana_GDS19
Populus_davidiana_GDS5

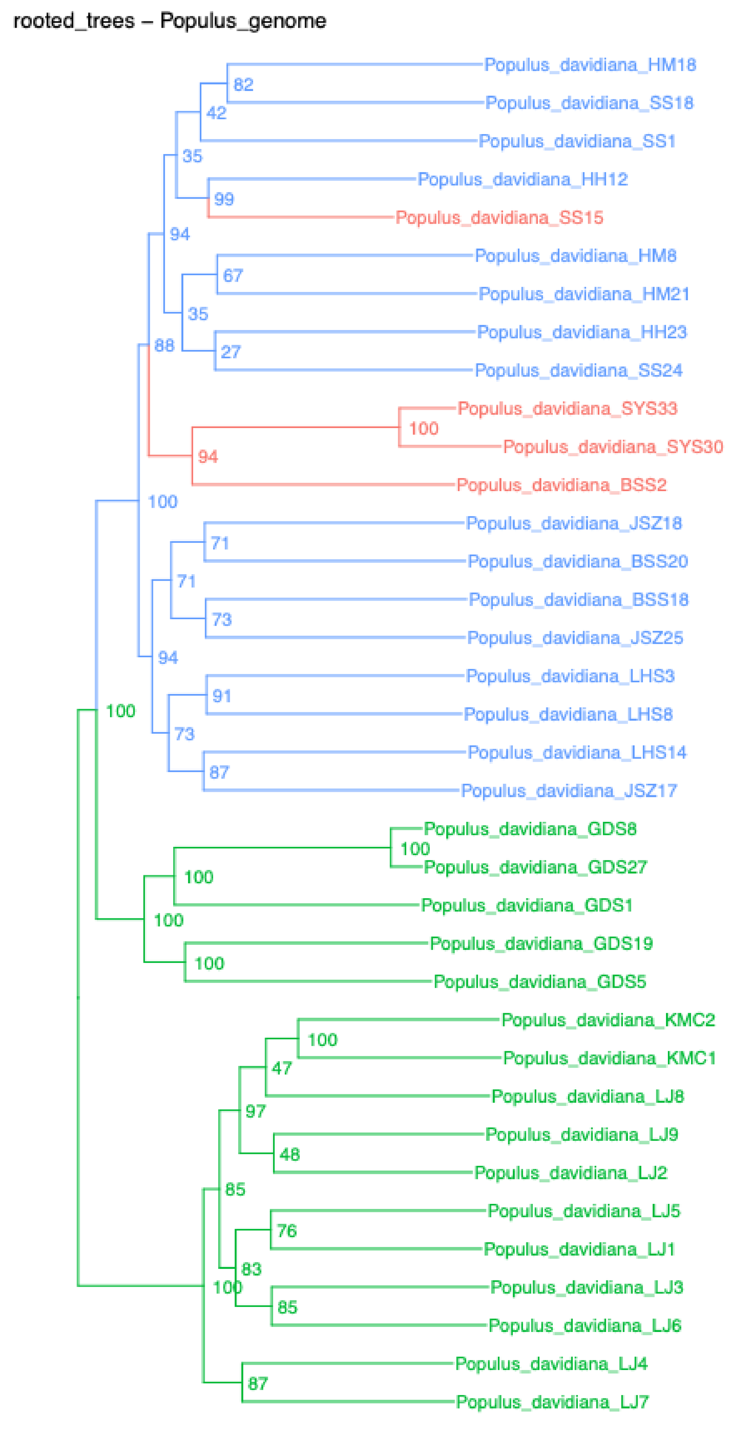
rooted_trees – Populus_genome
Populus_davidiana_HM18
Populus_davidiana_SS18
Populus_davidiana_SS1
Populus_davidiana_HH12
Populus_davidiana_SS15
Populus_davidiana_HM8
Populus_davidiana_HM21
Populus_davidiana_HH23
Populus_davidiana_SS24
Populus_davidiana_SYS33
Populus_davidiana_SYS30
Populus_davidiana_BSS2
Populus_davidiana_JSZ18
Populus_davidiana_BSS20
Populus_davidiana_BSS18
Populus_davidiana_JSZ25
Populus_davidiana_LHS3
Populus_davidiana_LHS8
Populus_davidiana_LHS14
Populus_davidiana_JSZ17
Populus_davidiana_GDS8
Populus_davidiana_GDS27
Populus_davidiana_GDS1
Populus_davidiana_GDS19
Populus_davidiana_GDS5
Populus_davidiana_KMC2
Populus_davidiana_KMC1
Populus_davidiana_LJ8
Populus_davidiana_LJ9
Populus_davidiana_LJ2
Populus_davidiana_LJ5
Populus_davidiana_LJ1
Populus_davidiana_LJ3
Populus_davidiana_LJ6
Populus_davidiana_LJ4
Populus_davidiana_LJ7

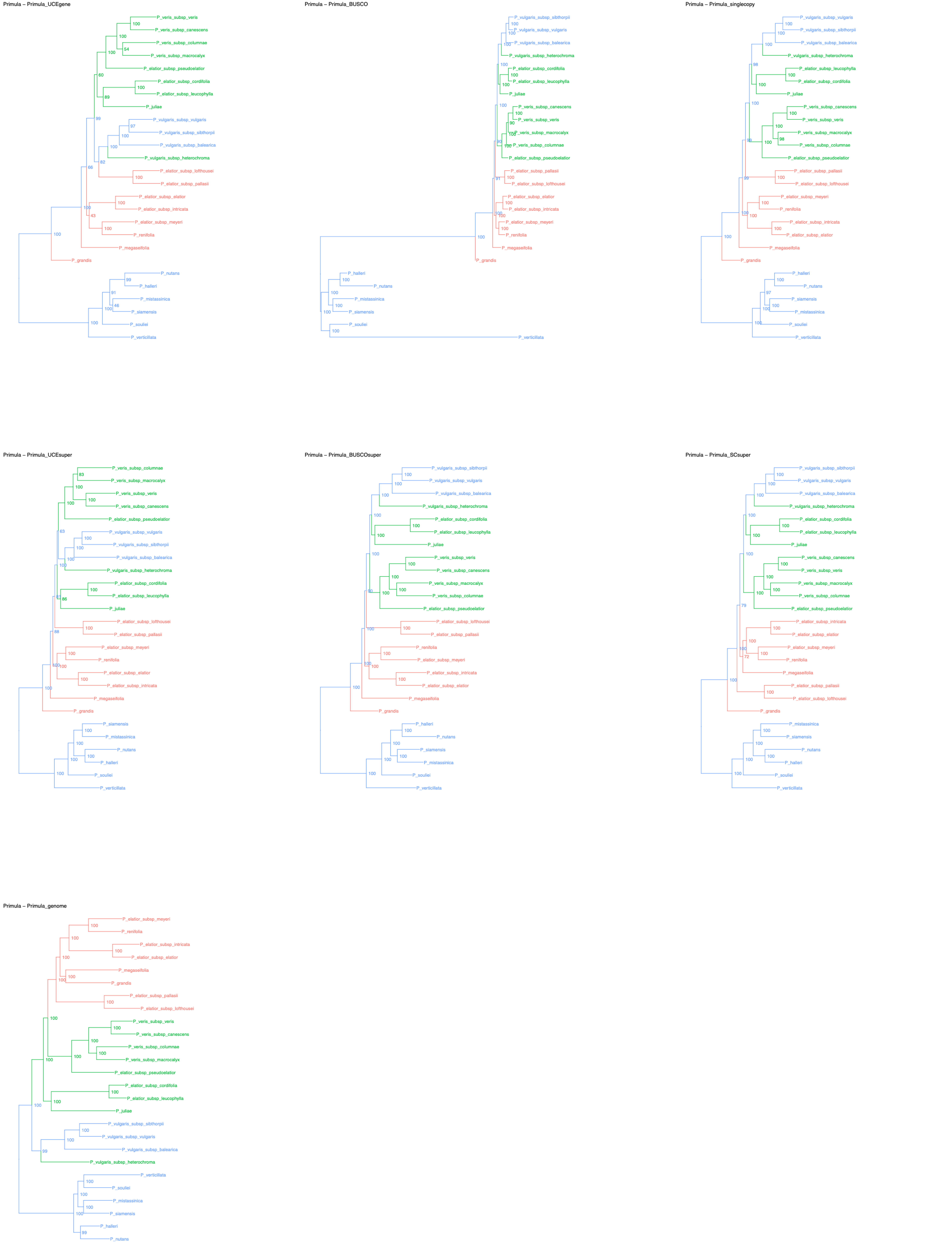
Primula – Primula_UCEgene
P_veris_subsp_veris
P_veris_subsp_canescens
P_veris_subsp_columnae
P_veris_subsp_macrocalyx
P_elatior_subsp_pseudoelatior
P_elatior_subsp_cordifolia
P_elatior_subsp_leucophylla
P_juliae
P_vulgaris_subsp_vulgaris
P_vulgaris_subsp_sibthorpii
P_vulgaris_subsp_balearica
P_vulgaris_subsp_heterochroma
P_elatior_subsp_lofthousei
P_elatior_subsp_pallasii
P_elatior_subsp_elatior
P_elatior_subsp_intricata
P_elatior_subsp_meyeri
P_renifolia
P_megaseifolia
P_grandis
P_nutans
P_halleri
P_mistassinica
P_siamensis
P_souliei
P_verticillata
Primula – Primula_BUSCO
P_vulgaris_subsp_sibthorpii
P_vulgaris_subsp_vulgaris
P_vulgaris_subsp_balearica
P_vulgaris_subsp_heterochroma
P_elatior_subsp_cordifolia
P_elatior_subsp_leucophylla
P_juliae
P_veris_subsp_canescens
P_veris_subsp_veris
P_veris_subsp_macrocalyx
P_veris_subsp_columnae
P_elatior_subsp_pseudoelatior
P_elatior_subsp_pallasii
P_elatior_subsp_lofthousei
P_elatior_subsp_elatior
P_elatior_subsp_intricata
P_elatior_subsp_meyeri
P_renifolia
P_megaseifolia
P_grandis
P_halleri
P_nutans
P_mistassinica
P_siamensis
P_souliei
P_verticillata
Primula – Primula_singlecopy
P_vulgaris_subsp_vulgaris
P_vulgaris_subsp_sibthorpii
P_vulgaris_subsp_balearica
P_vulgaris_subsp_heterochroma
P_elatior_subsp_leucophylla
P_elatior_subsp_cordifolia
P_juliae
P_veris_subsp_canescens
P_veris_subsp_veris
P_veris_subsp_macrocalyx
P_veris_subsp_columnae
P_elatior_subsp_pseudoelatior
P_elatior_subsp_pallasii
P_elatior_subsp_lofthousei
P_elatior_subsp_meyeri
P_renifolia
P_elatior_subsp_intricata
P_elatior_subsp_elatior
P_megaseifolia
P_grandis
P_halleri
P_nutans
P_siamensis
P_mistassinica
P_souliei
P_verticillata
Primula – Primula_UCEsuper
P_veris_subsp_columnae
P_veris_subsp_macrocalyx
P_veris_subsp_veris
P_veris_subsp_canescens
P_elatior_subsp_pseudoelatior
P_vulgaris_subsp_vulgaris
P_vulgaris_subsp_sibthorpii
P_vulgaris_subsp_balearica
P_vulgaris_subsp_heterochroma
P_elatior_subsp_cordifolia
P_elatior_subsp_leucophylla
P_juliae
P_elatior_subsp_lofthousei
P_elatior_subsp_pallasii
P_elatior_subsp_meyeri
P_renifolia
P_elatior_subsp_elatior
P_elatior_subsp_intricata
P_megaseifolia
P_grandis
P_siamensis
P_mistassinica
P_nutans
P_halleri
P_souliei
P_verticillata
Primula – Primula_BUSCOsuper
P_vulgaris_subsp_sibthorpii
P_vulgaris_subsp_vulgaris
P_vulgaris_subsp_balearica
P_vulgaris_subsp_heterochroma
P_elatior_subsp_cordifolia
P_elatior_subsp_leucophylla
P_juliae
P_veris_subsp_veris
P_veris_subsp_canescens
P_veris_subsp_macrocalyx
P_veris_subsp_columnae
P_elatior_subsp_pseudoelatior
P_elatior_subsp_lofthousei
P_elatior_subsp_pallasii
P_renifolia
P_elatior_subsp_meyeri
P_elatior_subsp_intricata
P_elatior_subsp_elatior
P_megaseifolia
P_grandis
P_halleri
P_nutans
P_siamensis
P_mistassinica
P_souliei
P_verticillata
Primula – Primula_SCsuper
P_vulgaris_subsp_sibthorpii
P_vulgaris_subsp_vulgaris
P_vulgaris_subsp_balearica
P_vulgaris_subsp_heterochroma
P_elatior_subsp_cordifolia
P_elatior_subsp_leucophylla
P_juliae
P_veris_subsp_canescens
P_veris_subsp_veris
P_veris_subsp_macrocalyx
P_veris_subsp_columnae
P_elatior_subsp_pseudoelatior
P_elatior_subsp_intricata
P_elatior_subsp_elatior
P_elatior_subsp_meyeri
P_renifolia
P_megaseifolia
P_elatior_subsp_pallasii
P_elatior_subsp_lofthousei
P_grandis
P_mistassinica
P_siamensis
P_nutans
P_halleri
P_souliei
P_verticillata
Primula – Primula_genome
P_elatior_subsp_meyeri
P_renifolia
P_elatior_subsp_intricata
P_elatior_subsp_elatior
P_megaseifolia
P_grandis
P_elatior_subsp_pallasii
P_elatior_subsp_lofthousei
P_veris_subsp_veris
P_veris_subsp_canescens
P_veris_subsp_columnae
P_veris_subsp_macrocalyx
P_elatior_subsp_pseudoelatior
P_elatior_subsp_cordifolia
P_elatior_subsp_leucophylla
P_juliae
P_vulgaris_subsp_sibthorpii
P_vulgaris_subsp_vulgaris
P_vulgaris_subsp_balearica
P_vulgaris_subsp_heterochroma
P_verticillata
P_souliei
P_mistassinica
P_siamensis
P_halleri
P_nutans

v2 – Prunus_UCEgene

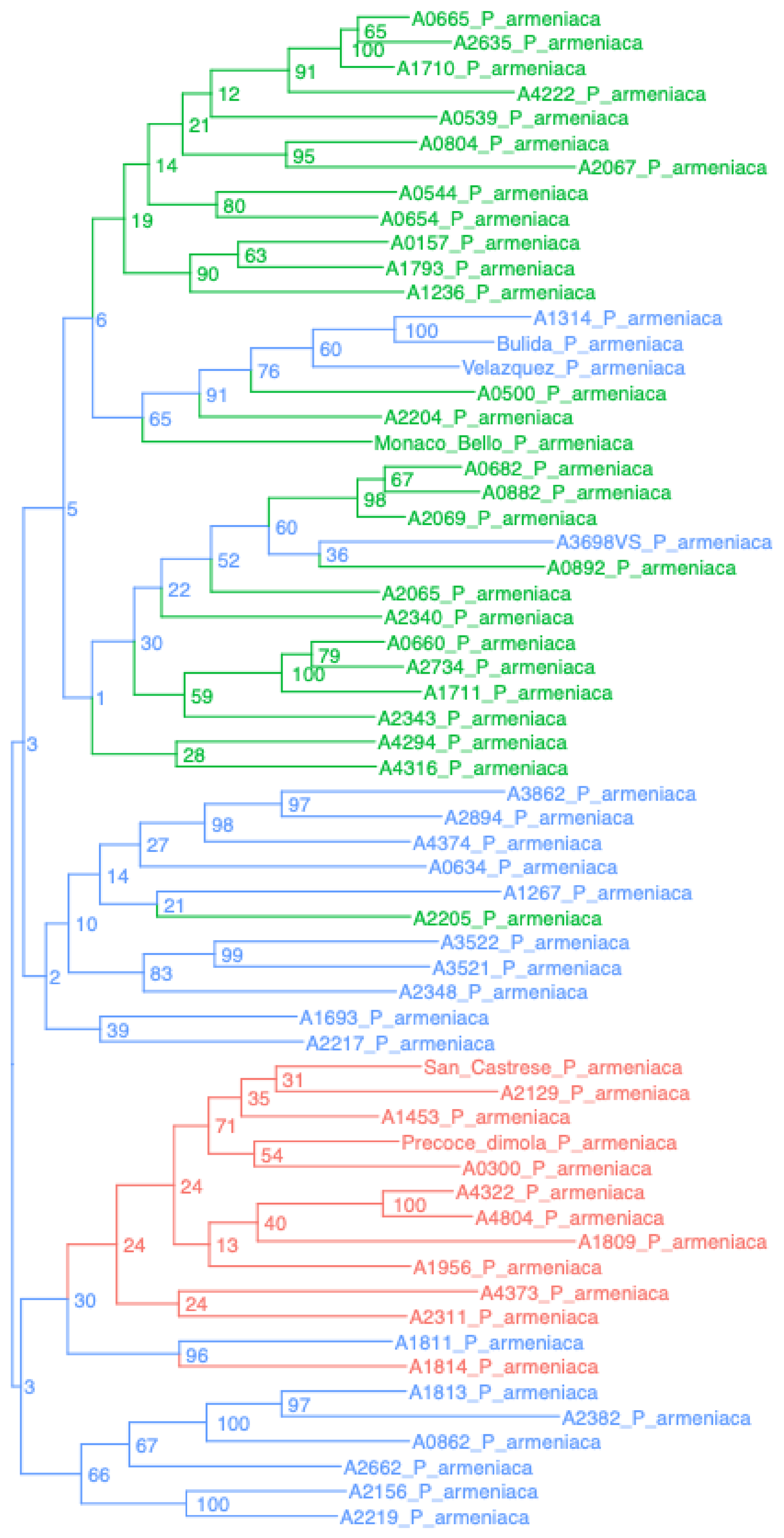


v2 – Prunus_BUSCO

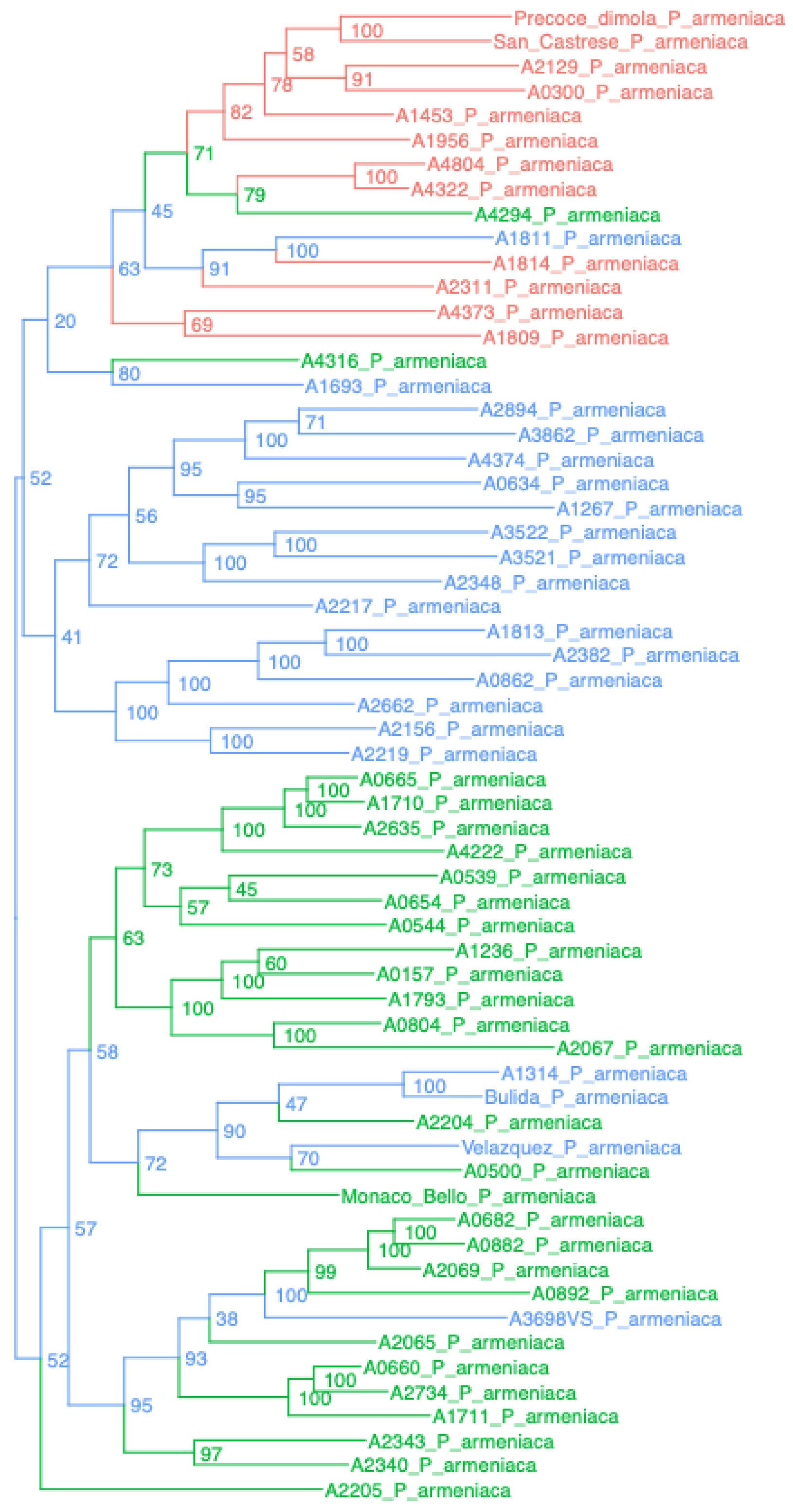


v2 – Prunus_singlecopy

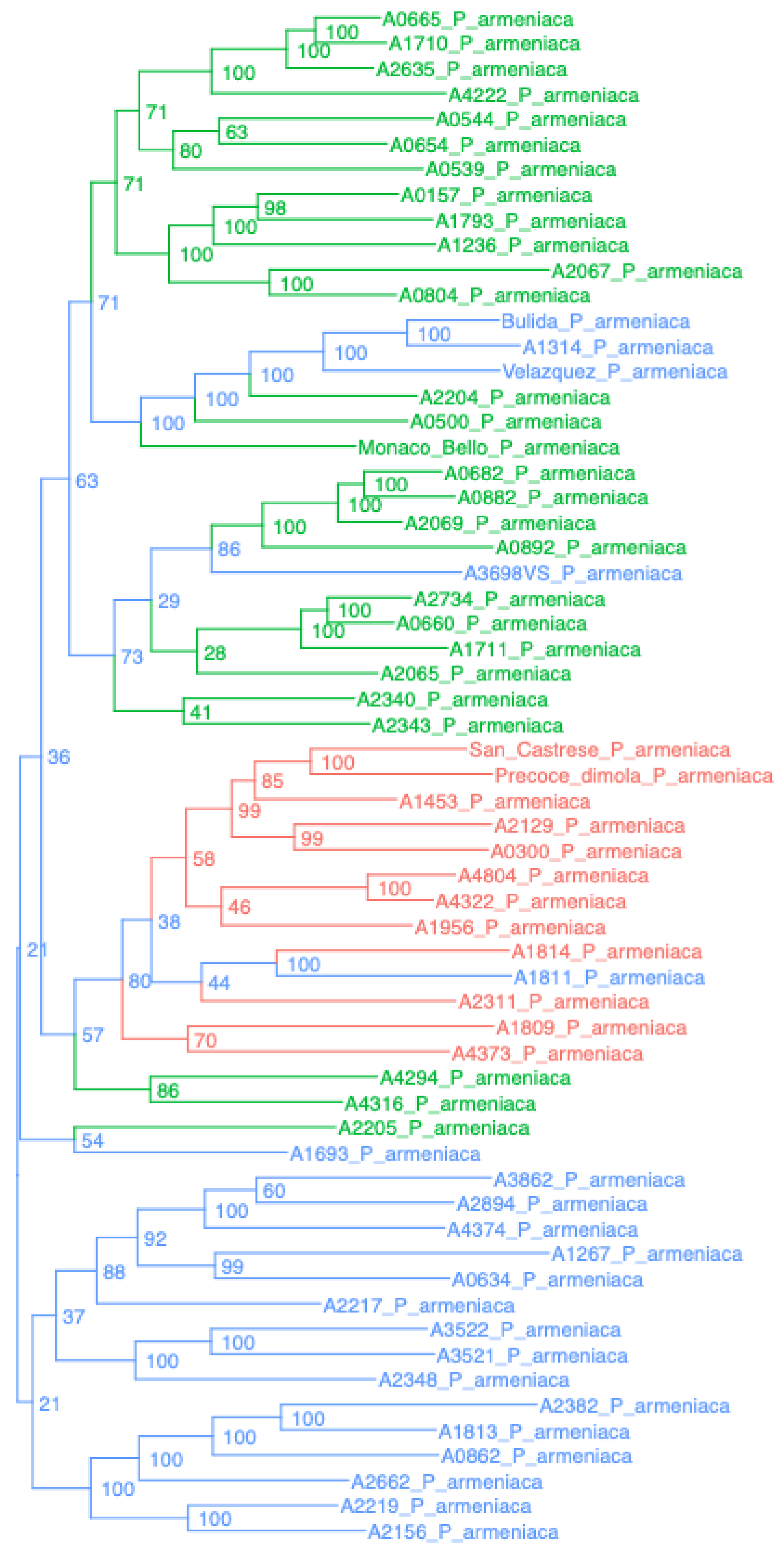


v2 – Prunus_UCEsuper

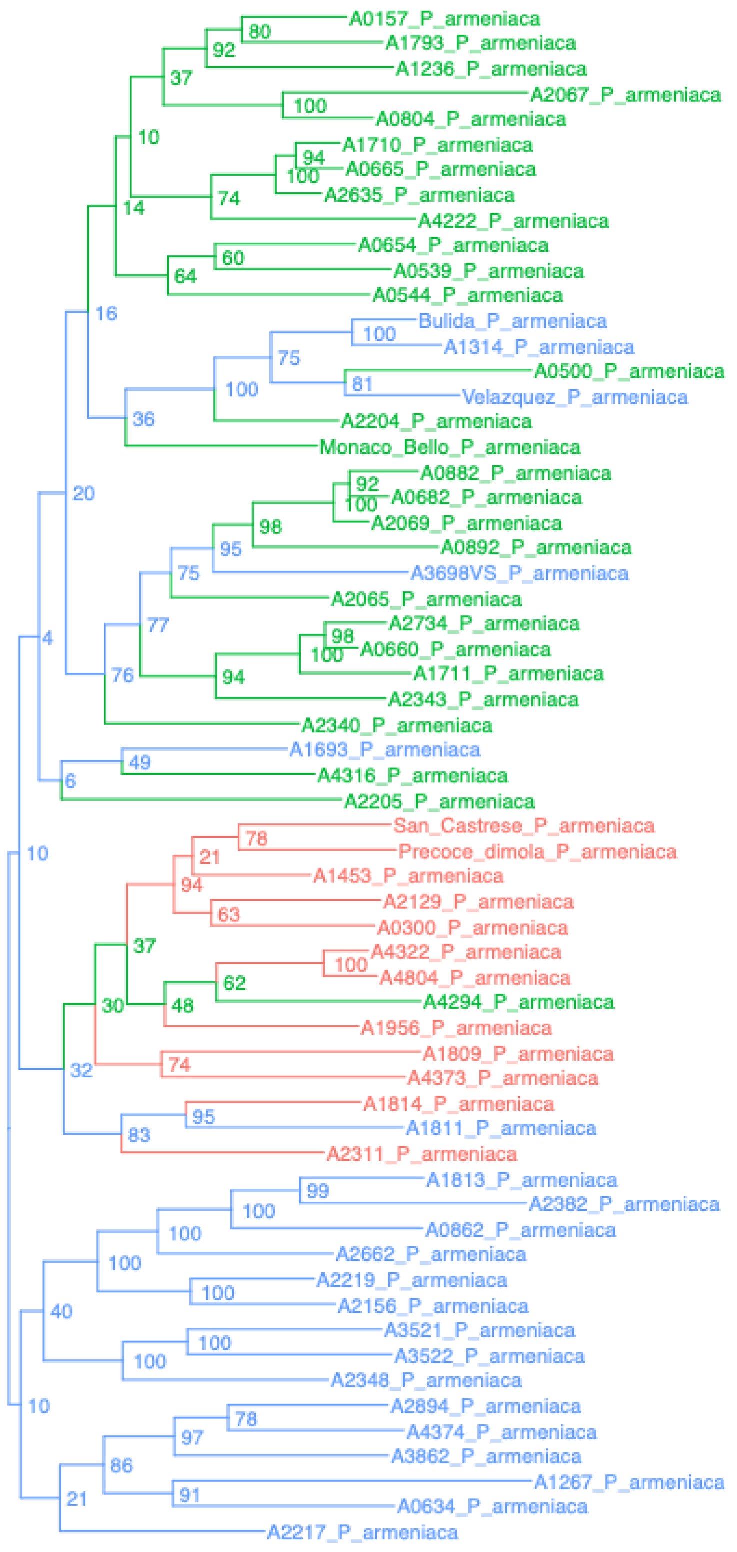


v2 – Prunus_BUSCOsuper

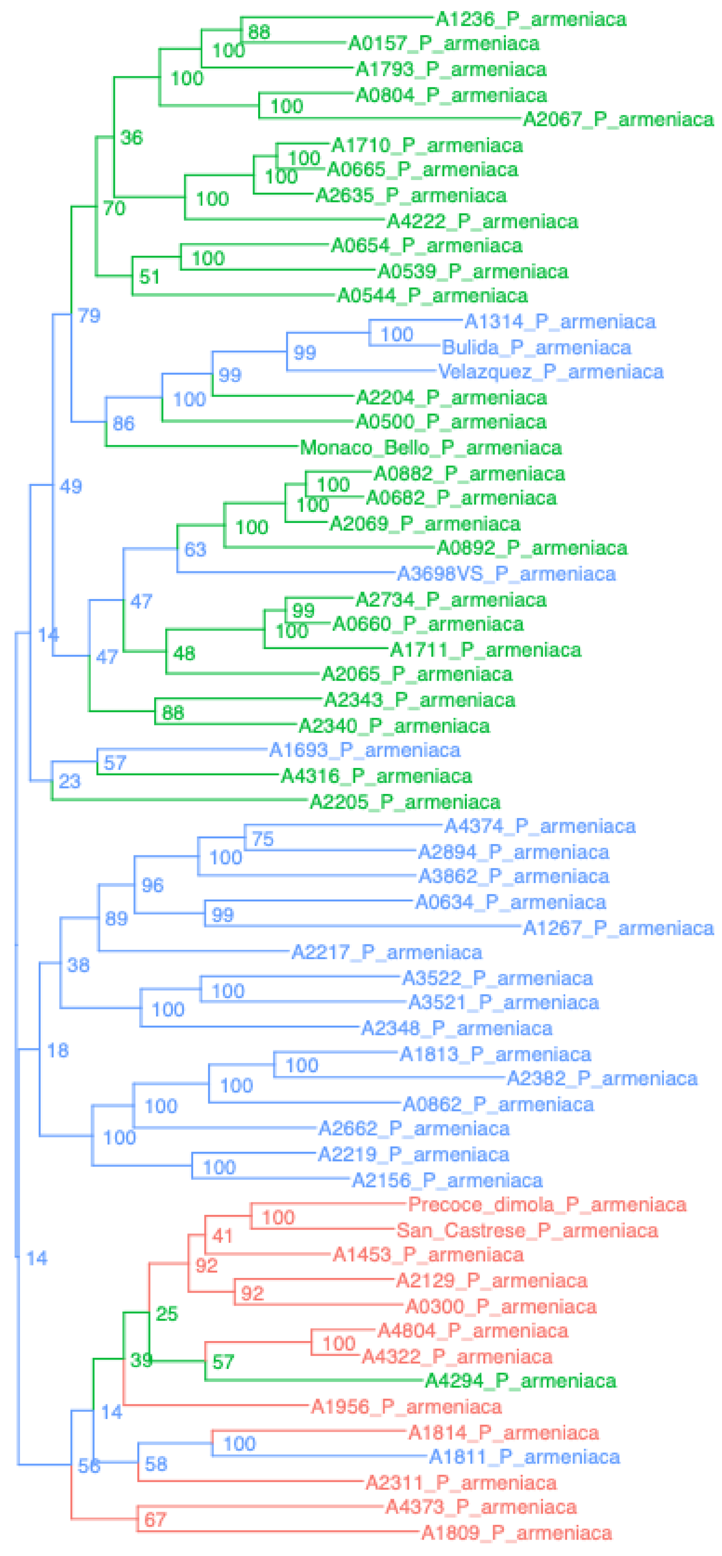


v2 – Prunus_SCsuper

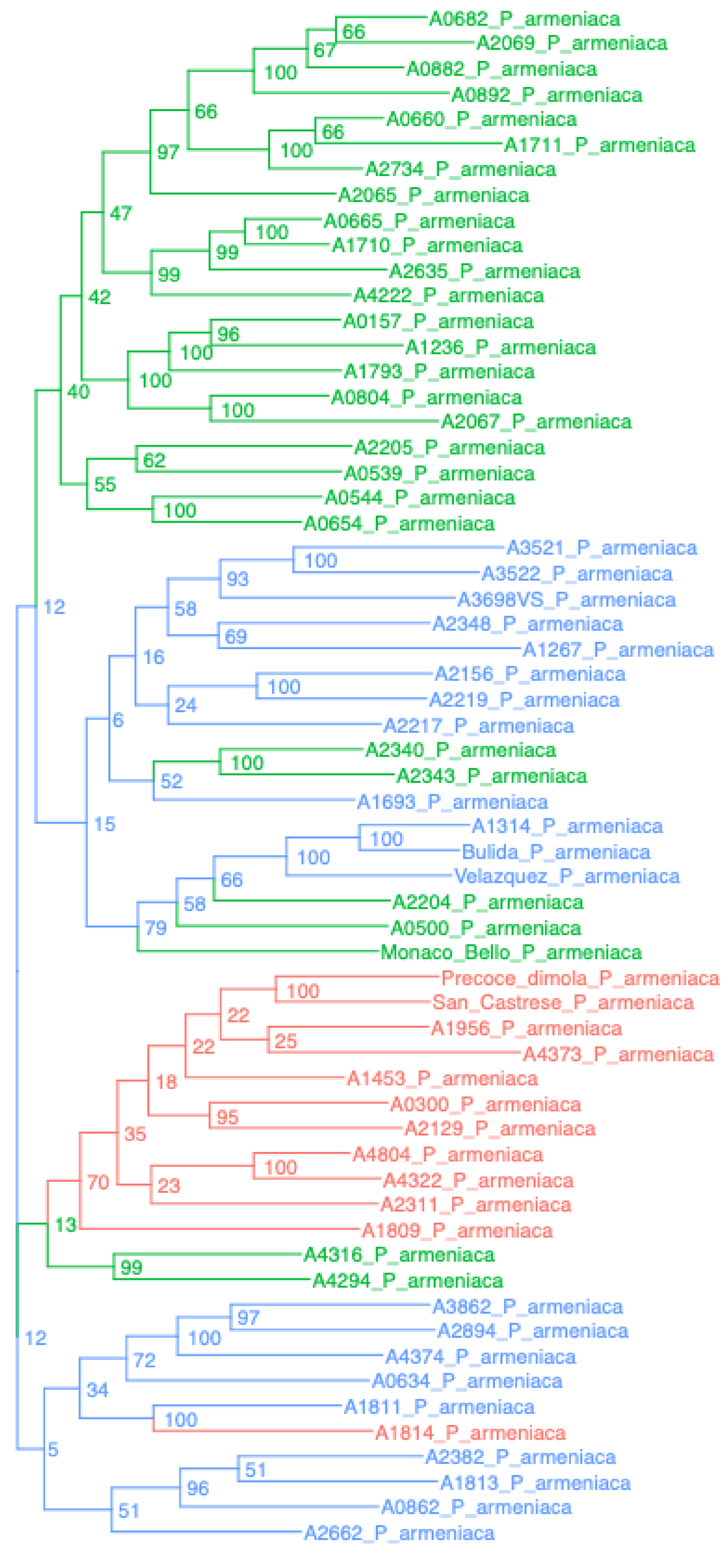


v2 – Prunus_armeniaca_genome

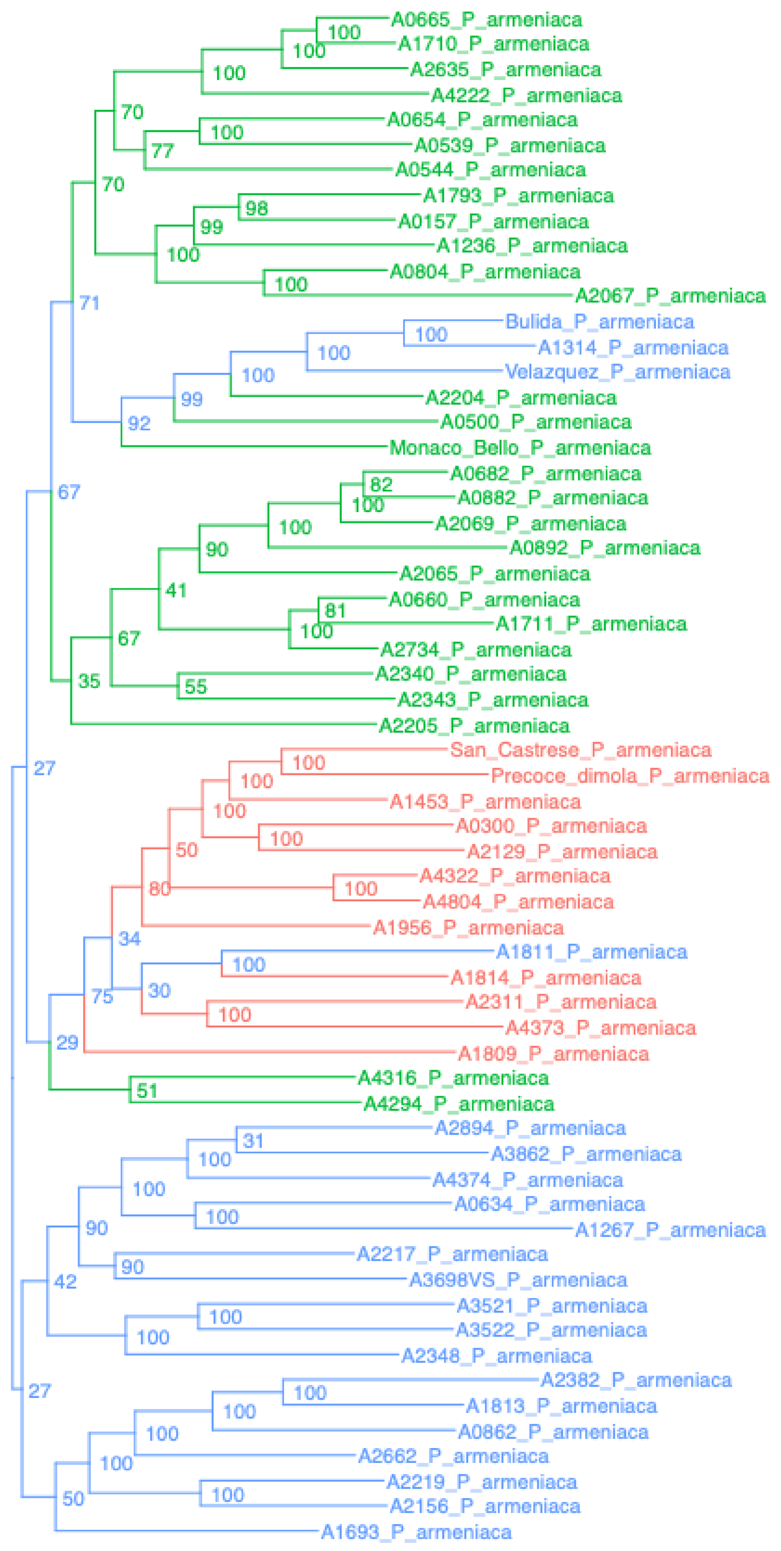

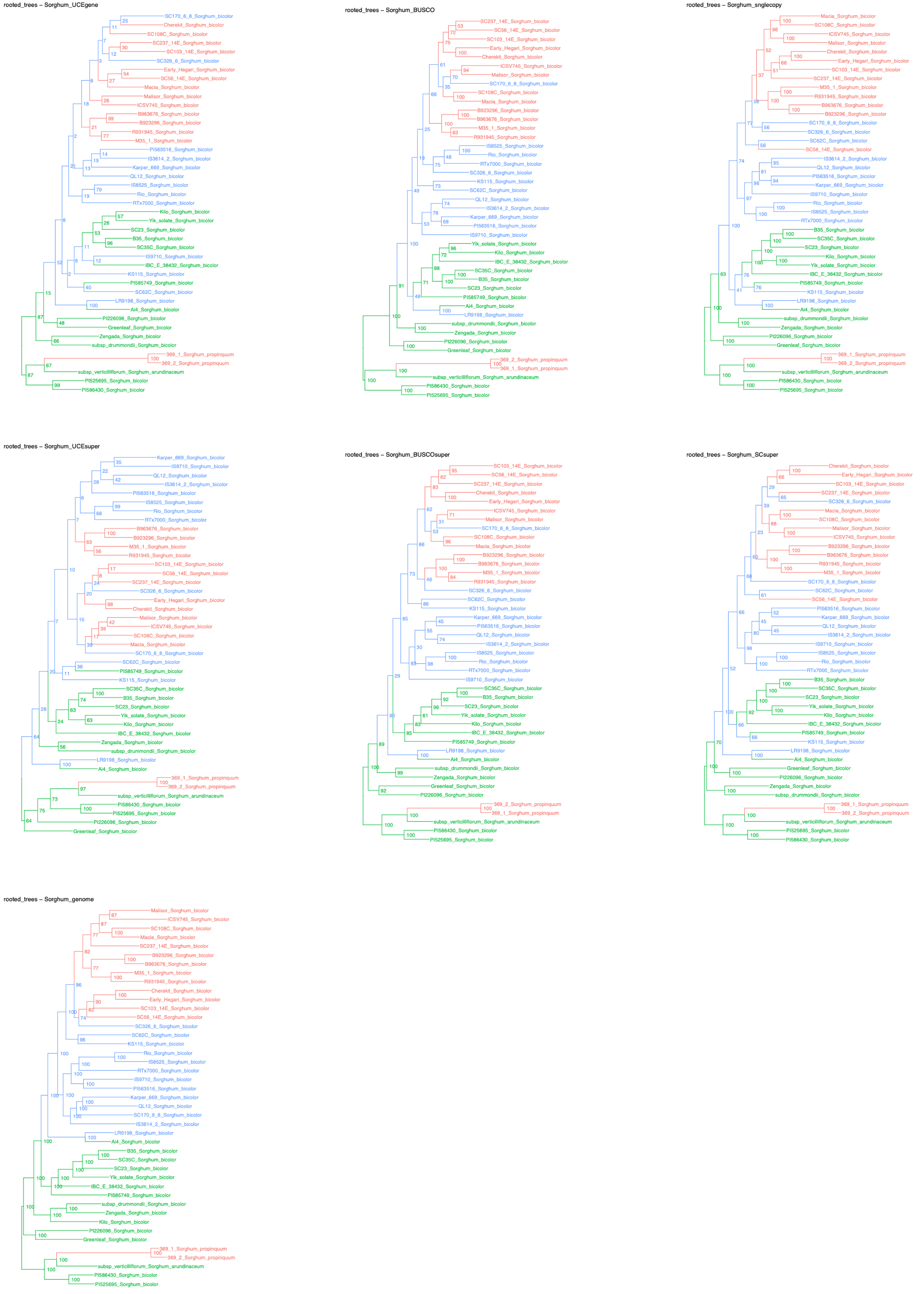
rooted_trees – Sorghum_UCEgene
rooted_trees – Sorghum_BUSCO
rooted_trees – Sorghum_snglecopy
rooted_trees – Sorghum_UCEsuper
rooted_trees – Sorghum_BUSCOsuper
rooted_trees – Sorghum_SCsuper
rooted_trees – Sorghum_genome

v2 – Vitis_UCEgene

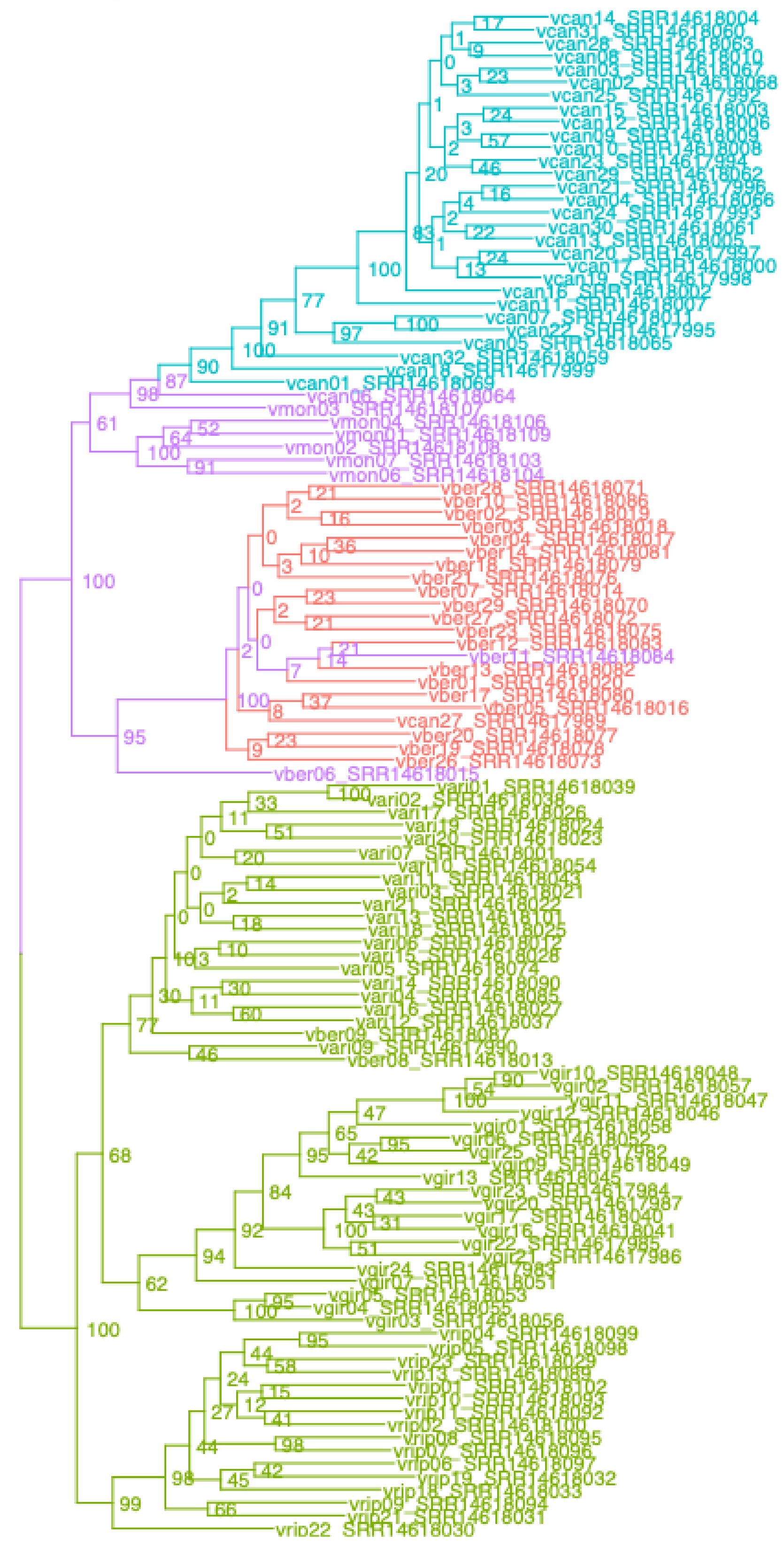

v2 – Vitis_BUSCO

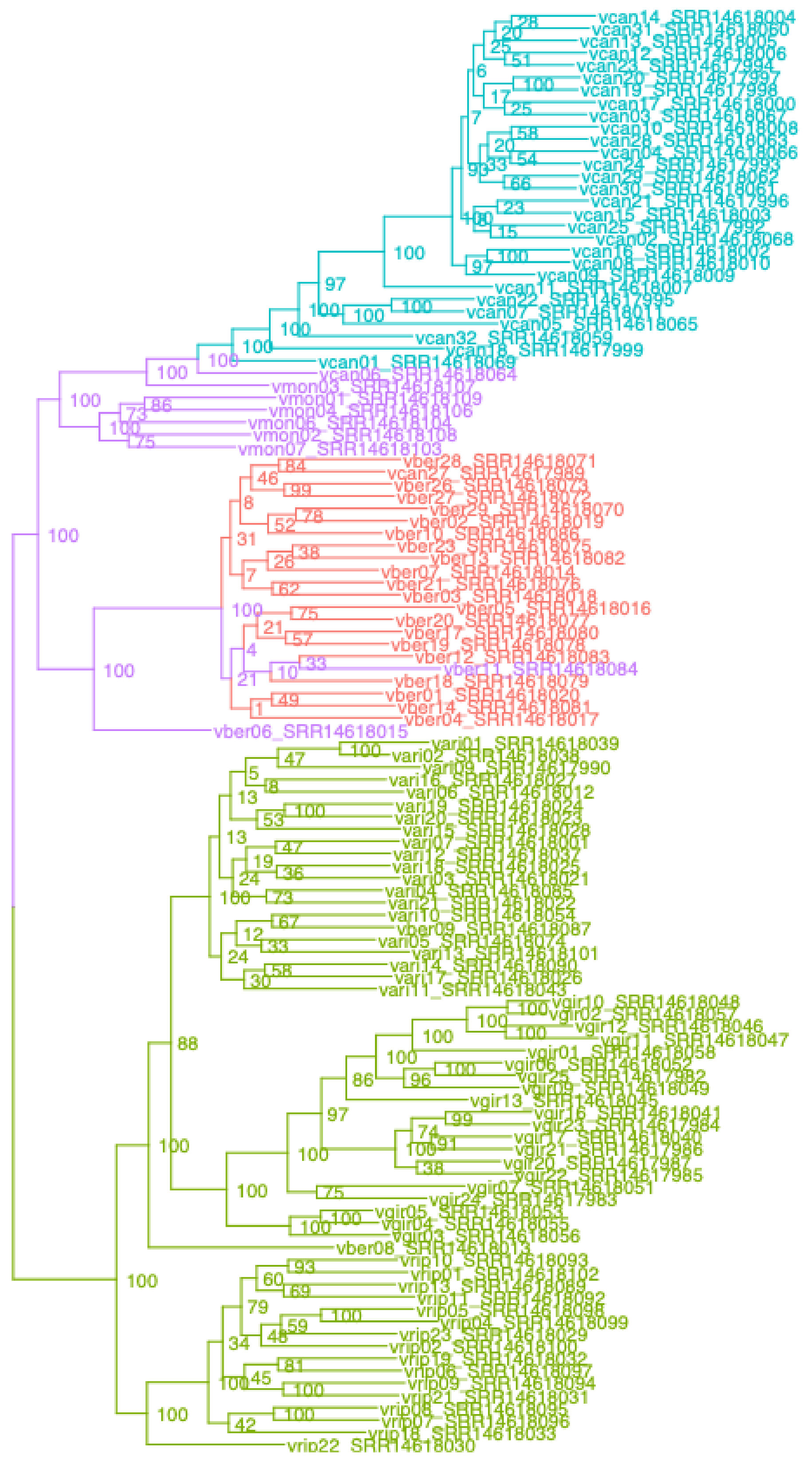

v2 – Vitis_singlecopy

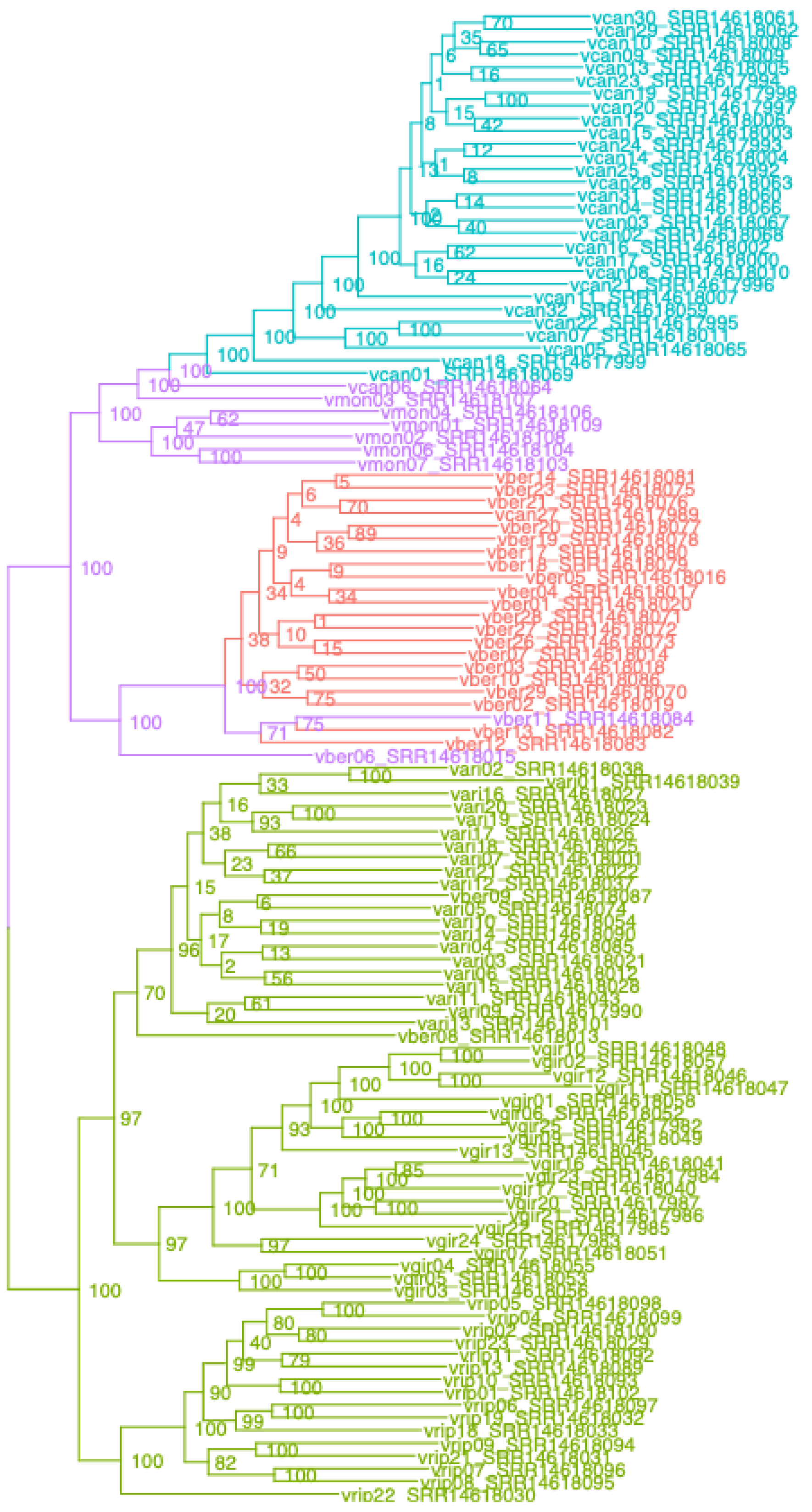

v2 – Vitis_UCEsuper

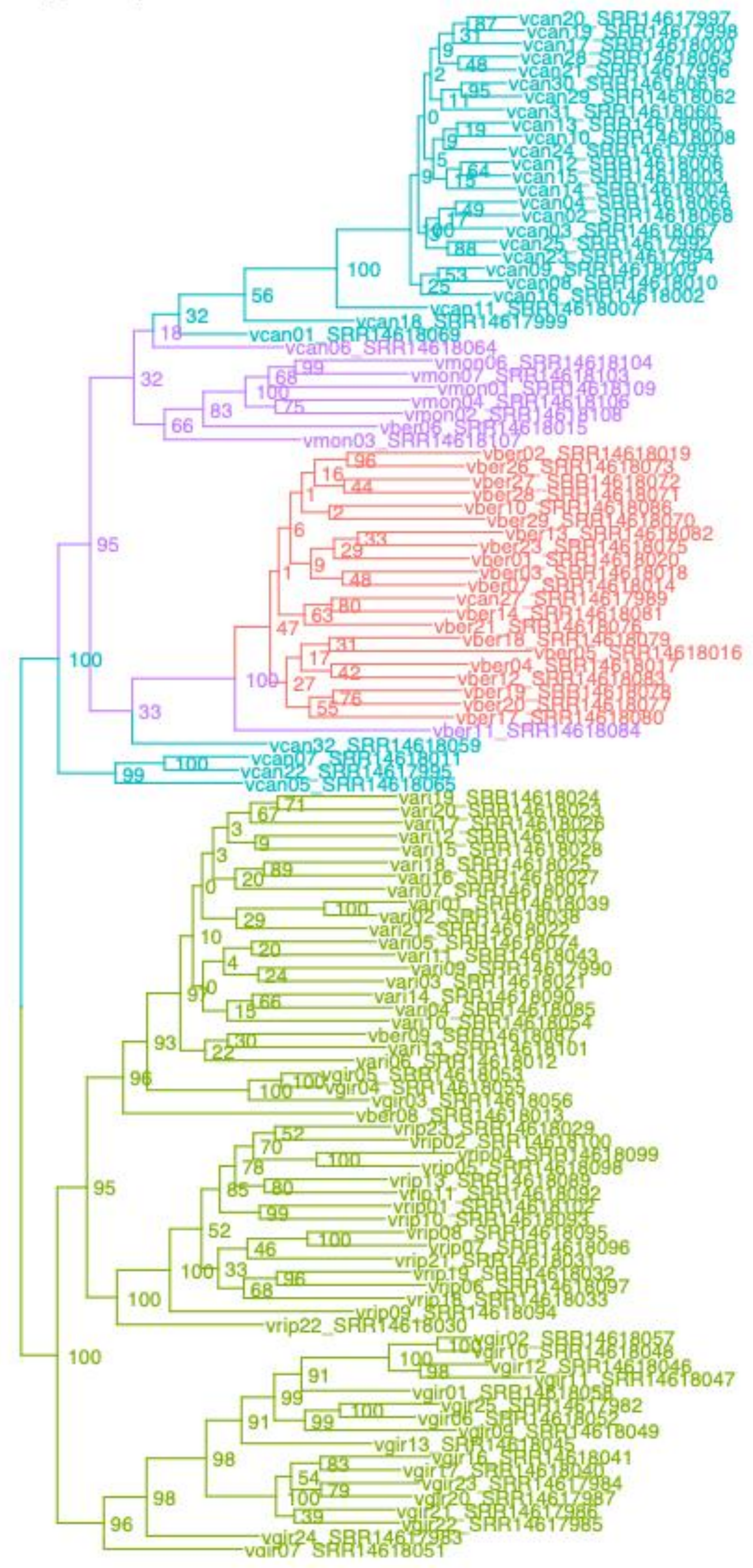

v2 – Vitis_BUSCOsuper

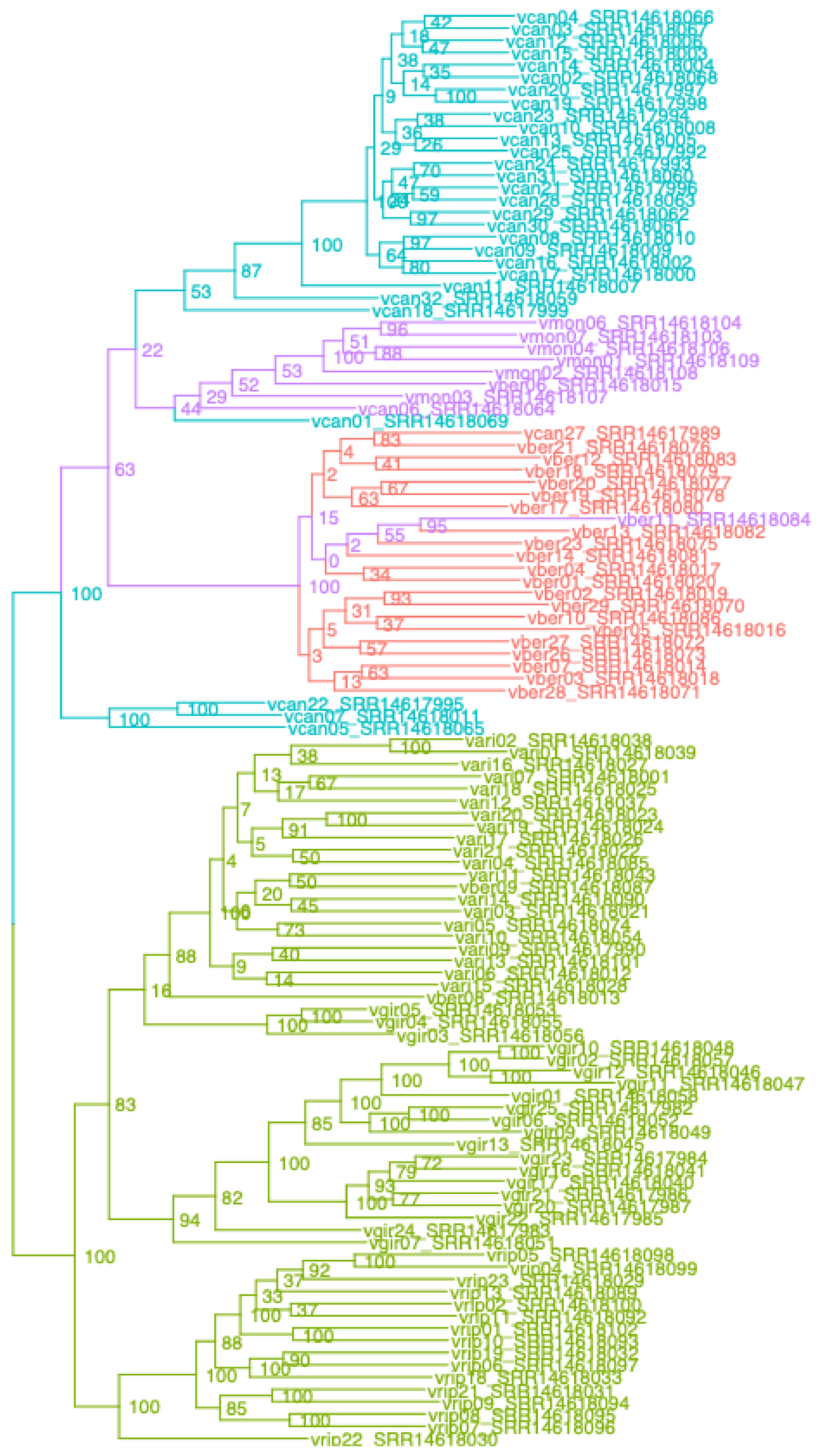

v2 – Vitis_SCsuper

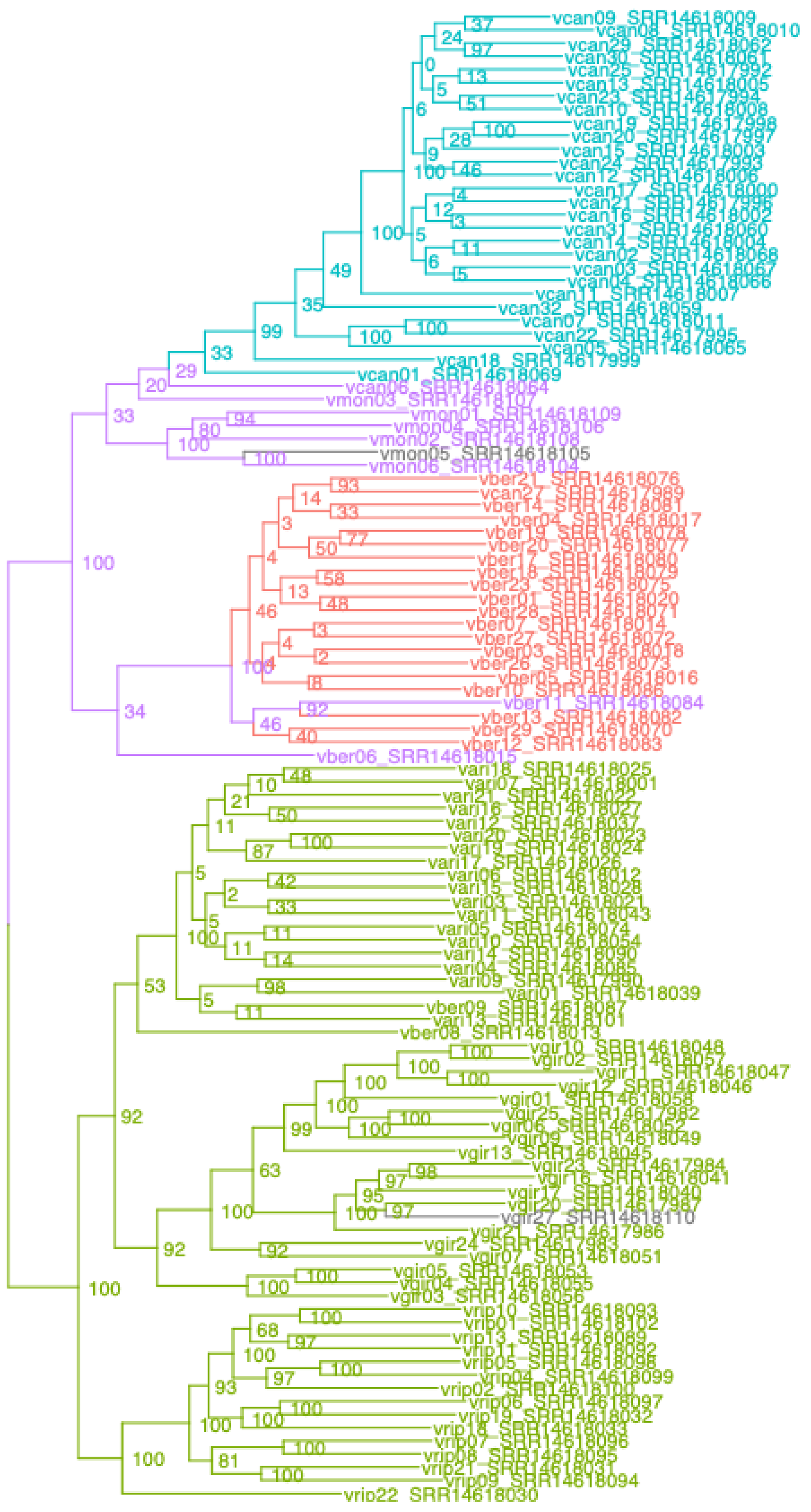

v2 – Vitis_genome

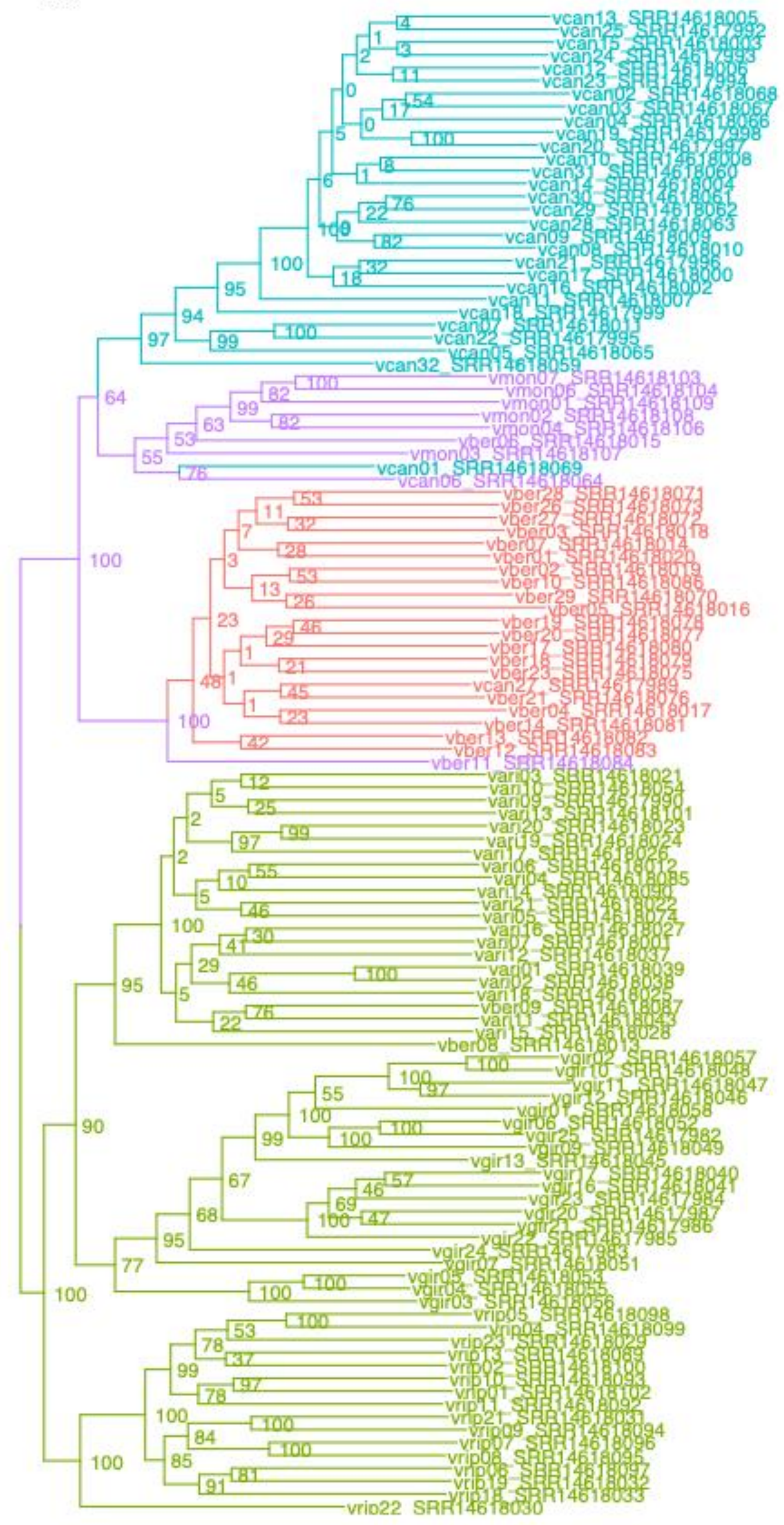

salix – Salix_UCE

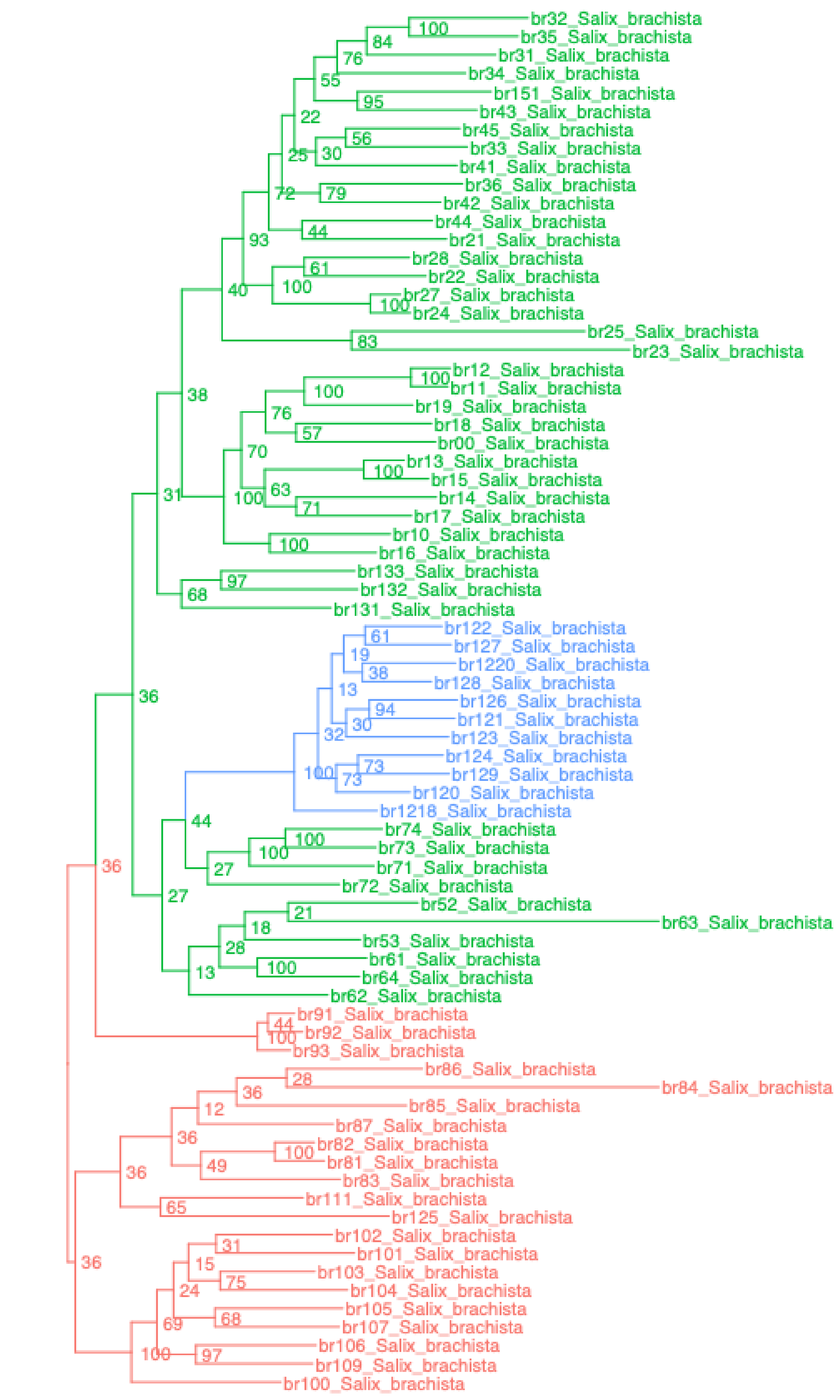

salix – Salix_BUSCO

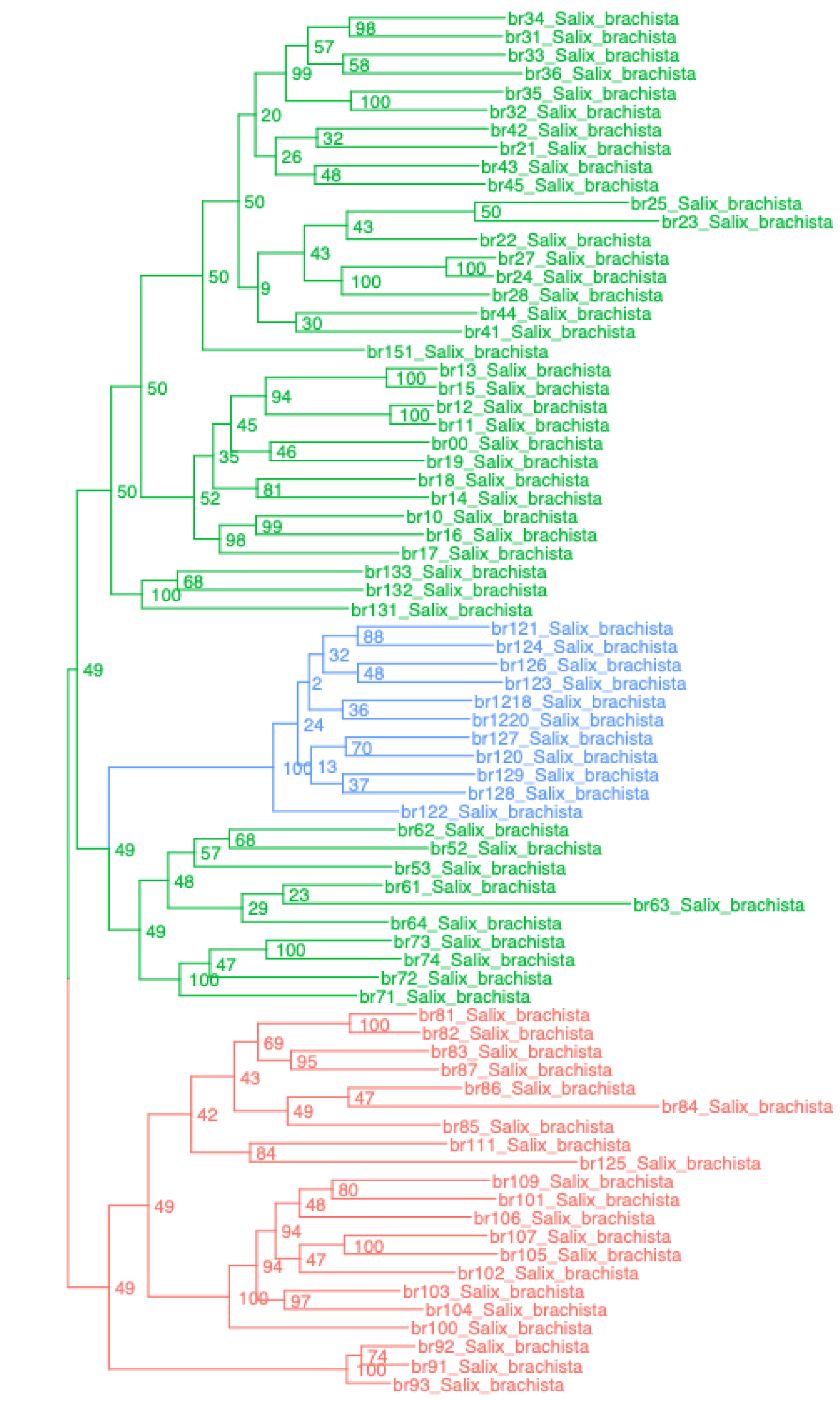

salix – Salix_singlecopy

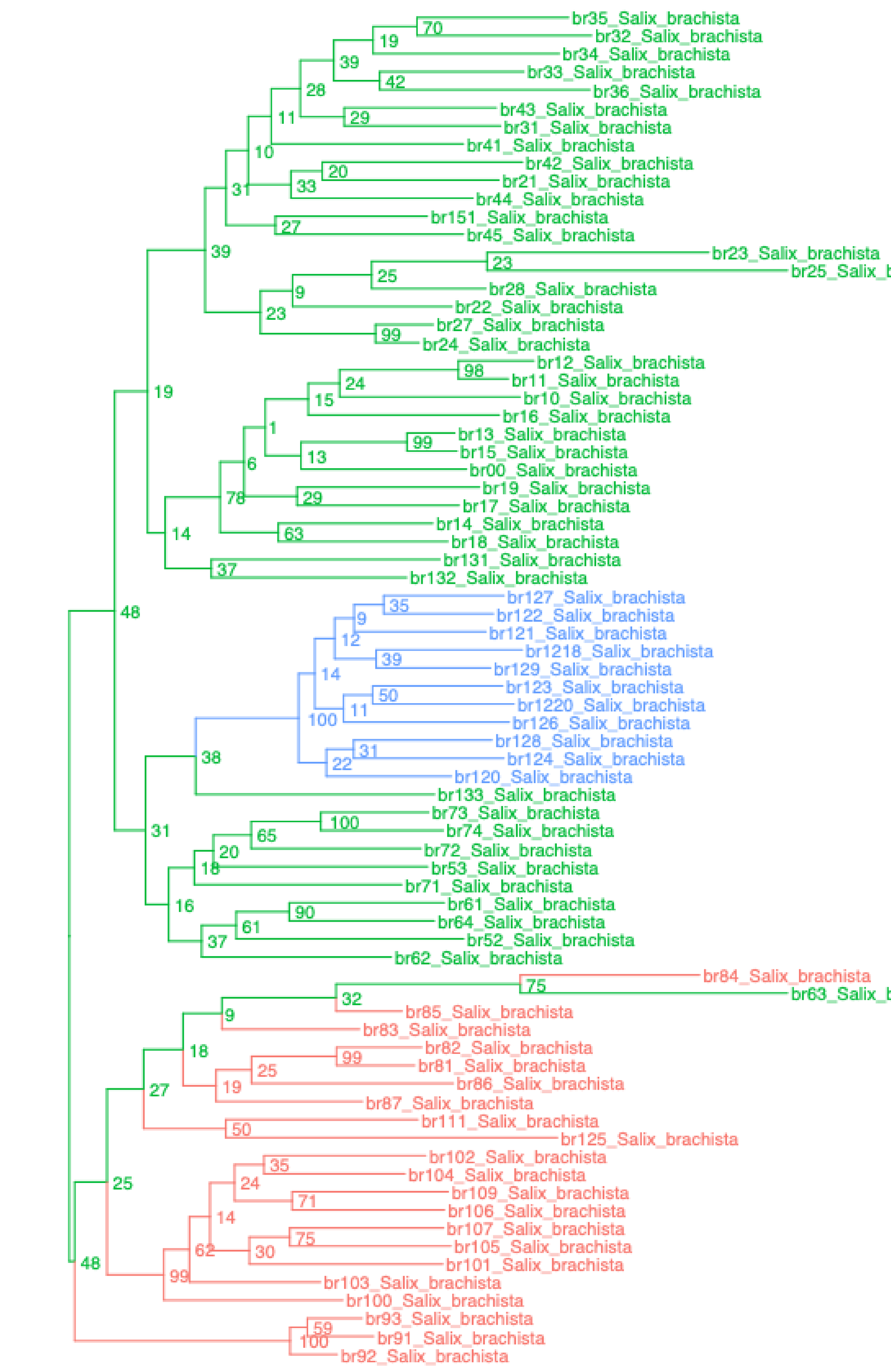

salix – Salix_UCEsuper

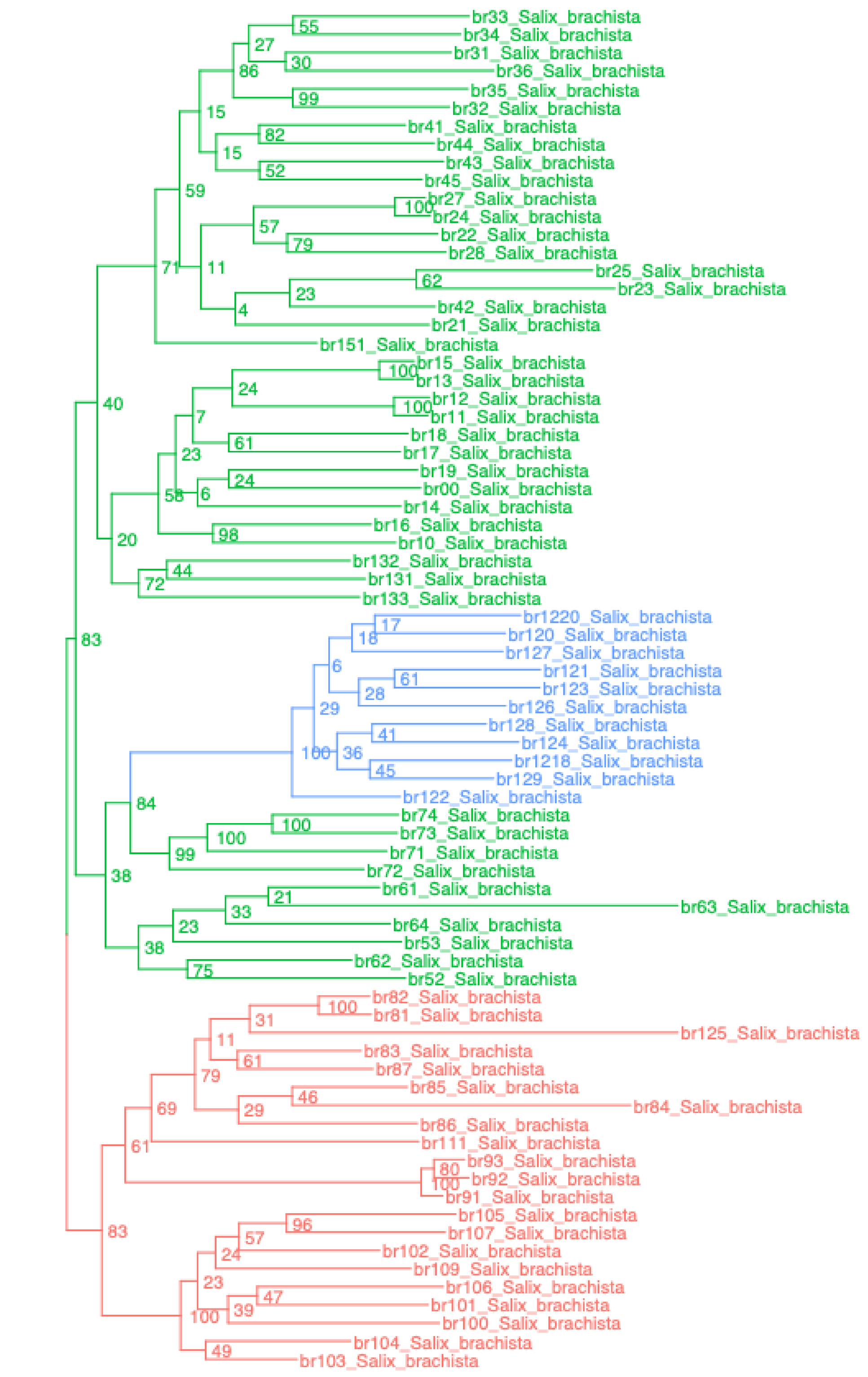

salix – Salix_BUSCOsuper

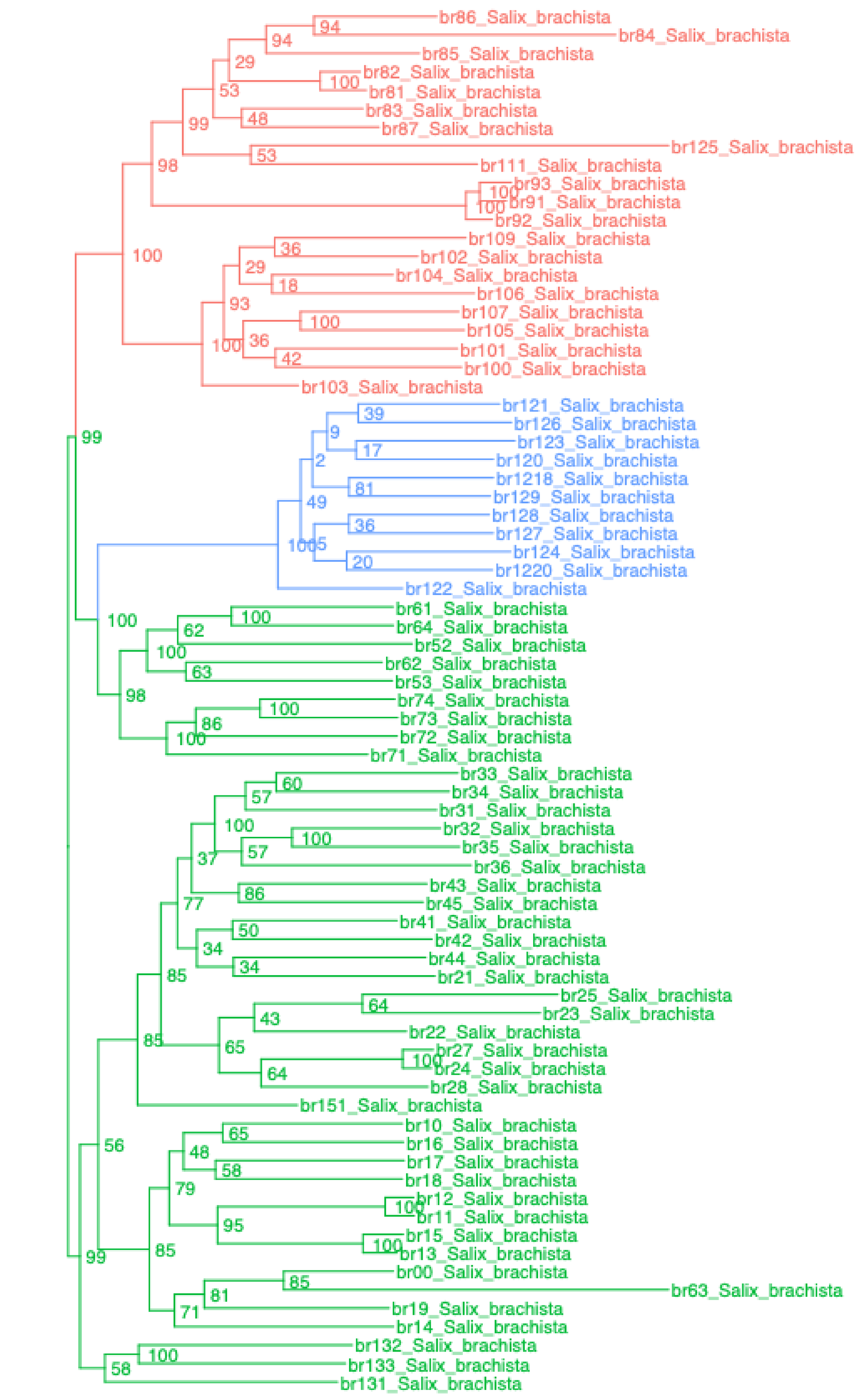

salix – Salix_SCsuper

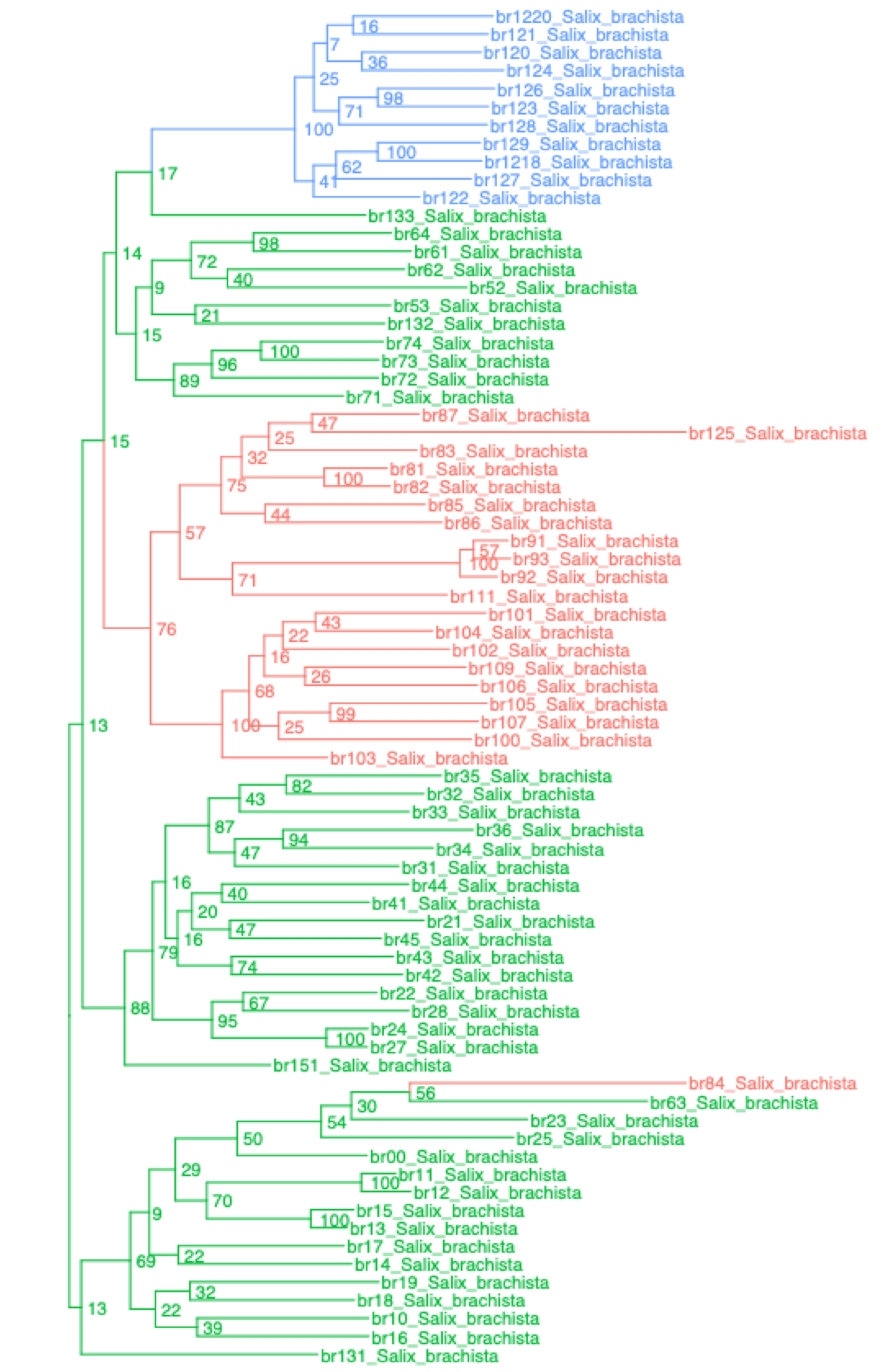

salix – Salix_genome

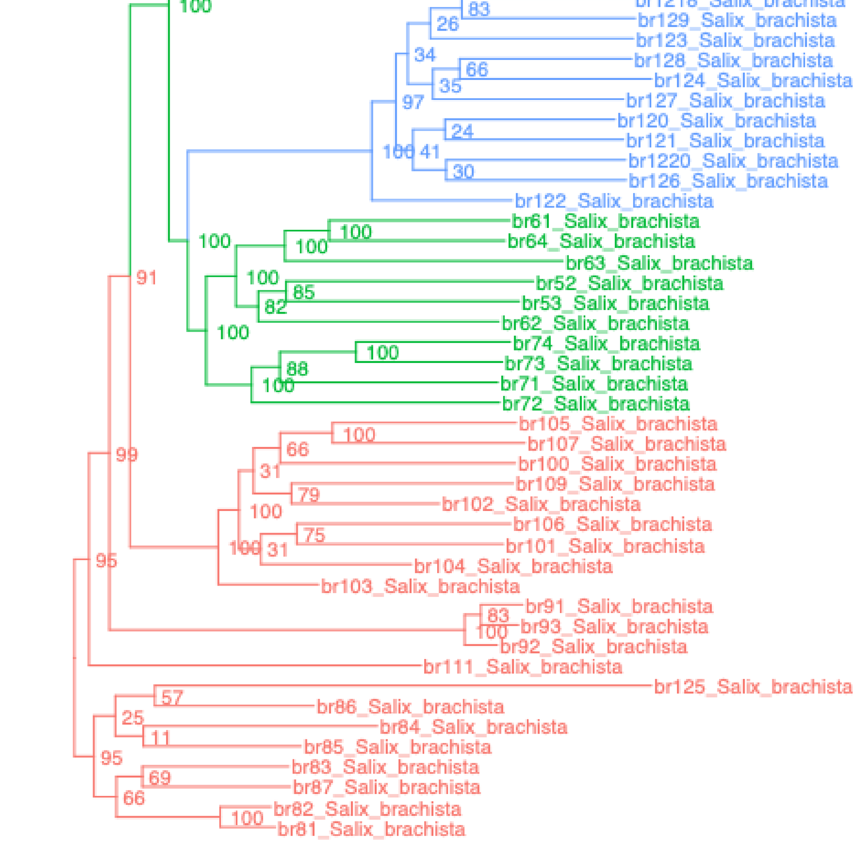

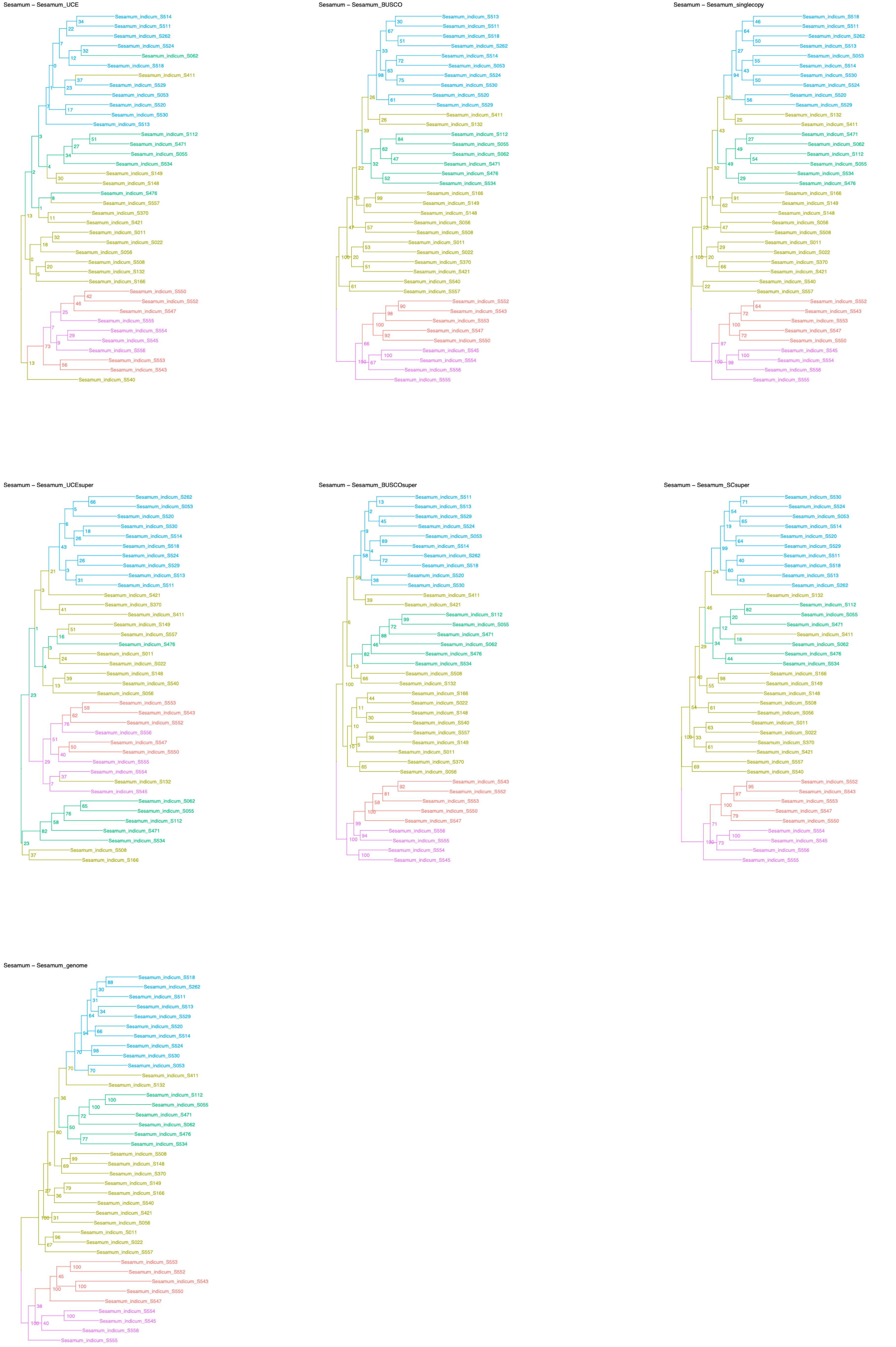

Sesamum – Sesamum_UCE
Sesamum – Sesamum_BUSCO
Sesamum – Sesamum_singlecopy
Sesamum – Sesamum_UCEsuper
Sesamum – Sesamum_BUSCOsuper
Sesamum – Sesamum_SCsuper
Sesamum – Sesamum_genome

rooted_trees – Thlapsi_UCEgene

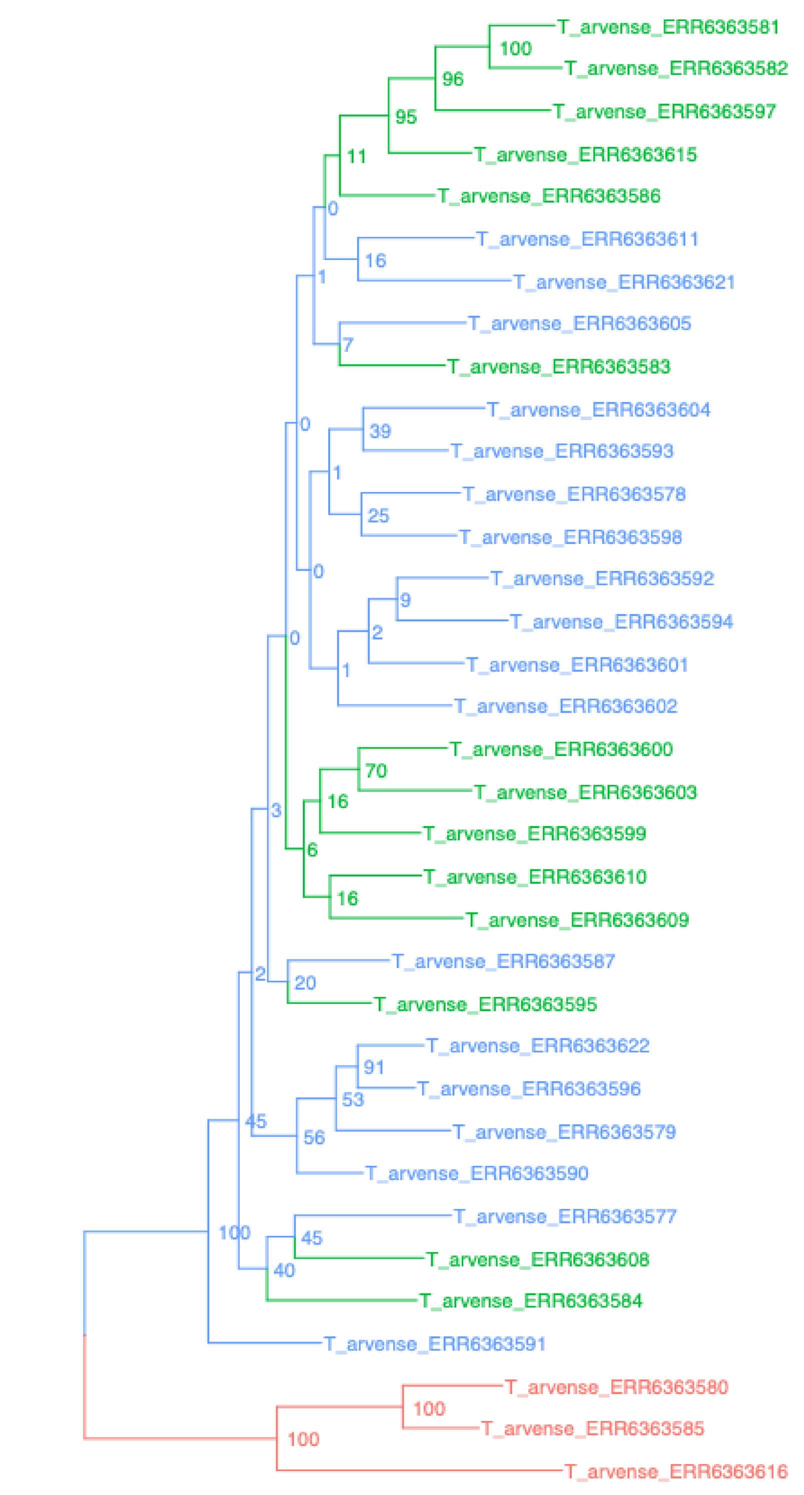


rooted_trees – Thlapsi_BUSCO

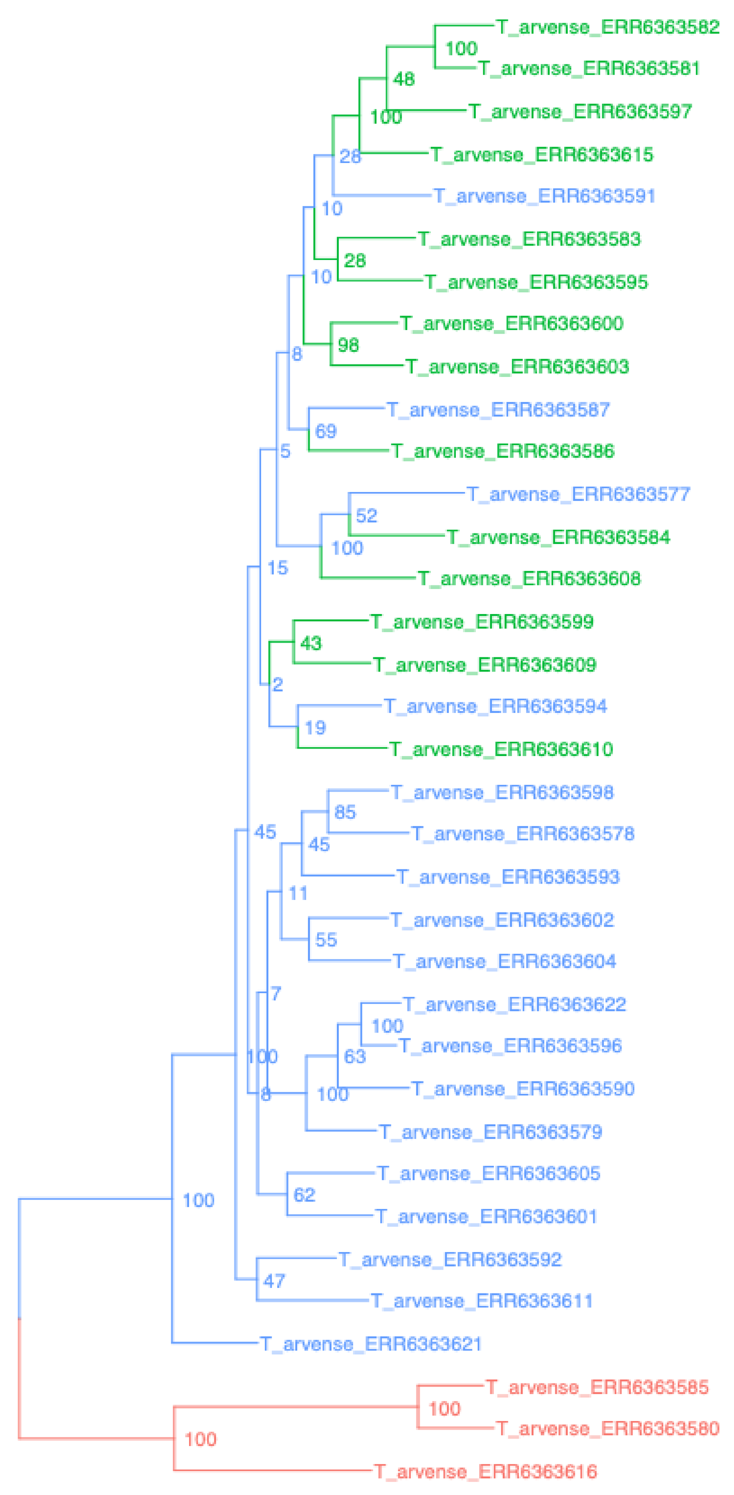


rooted_trees – Thlapsi_singlecopy

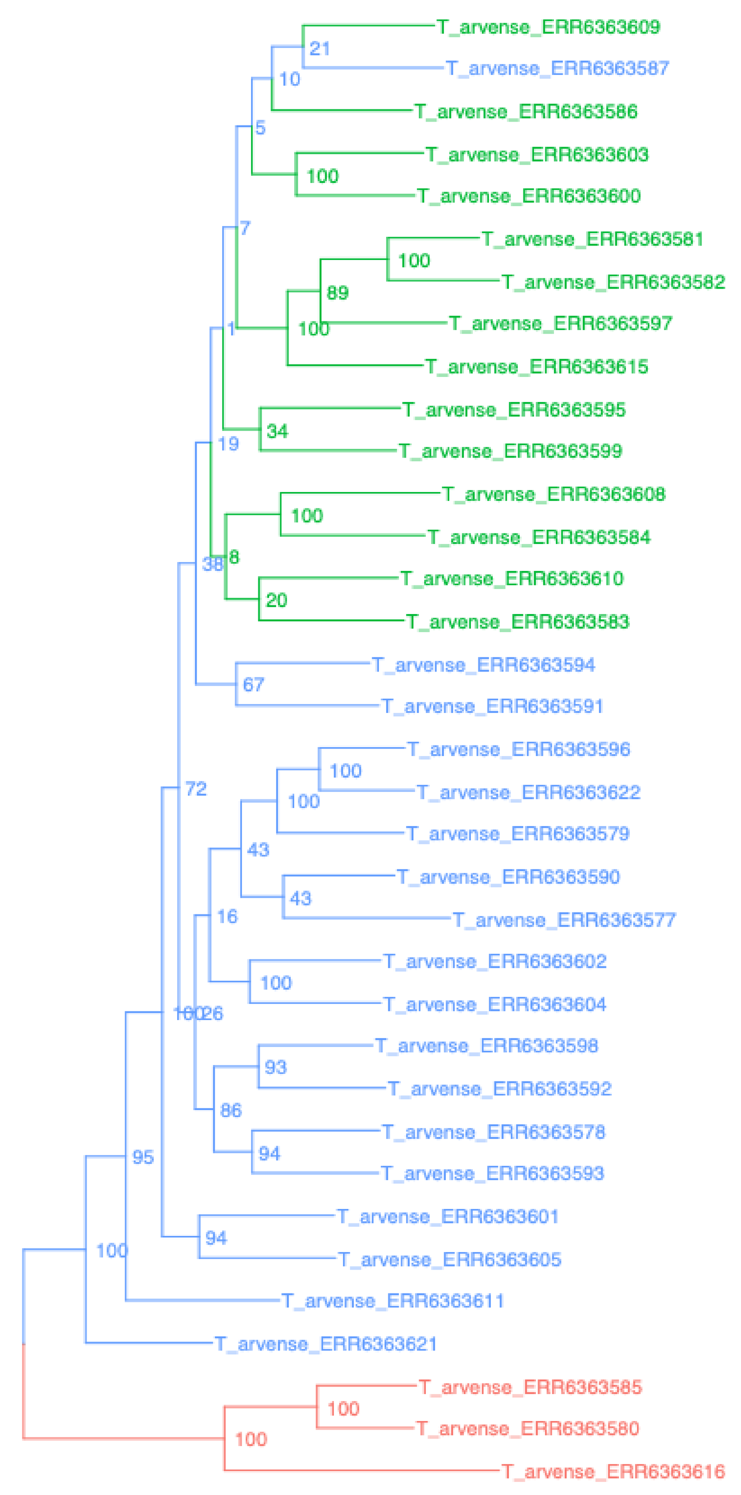


rooted_trees – Thlapsi_UCEsuper

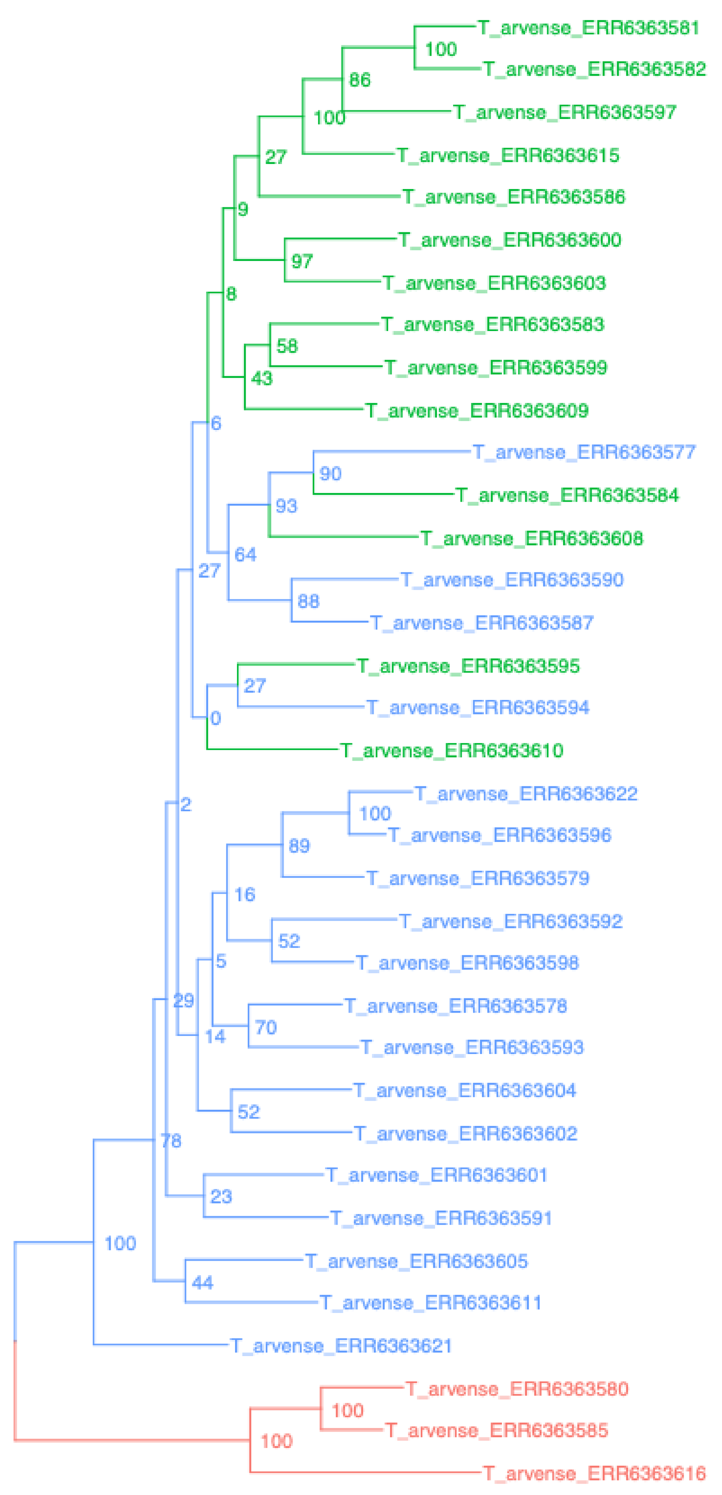


rooted_trees – Thlapsi_BUSCOsuper

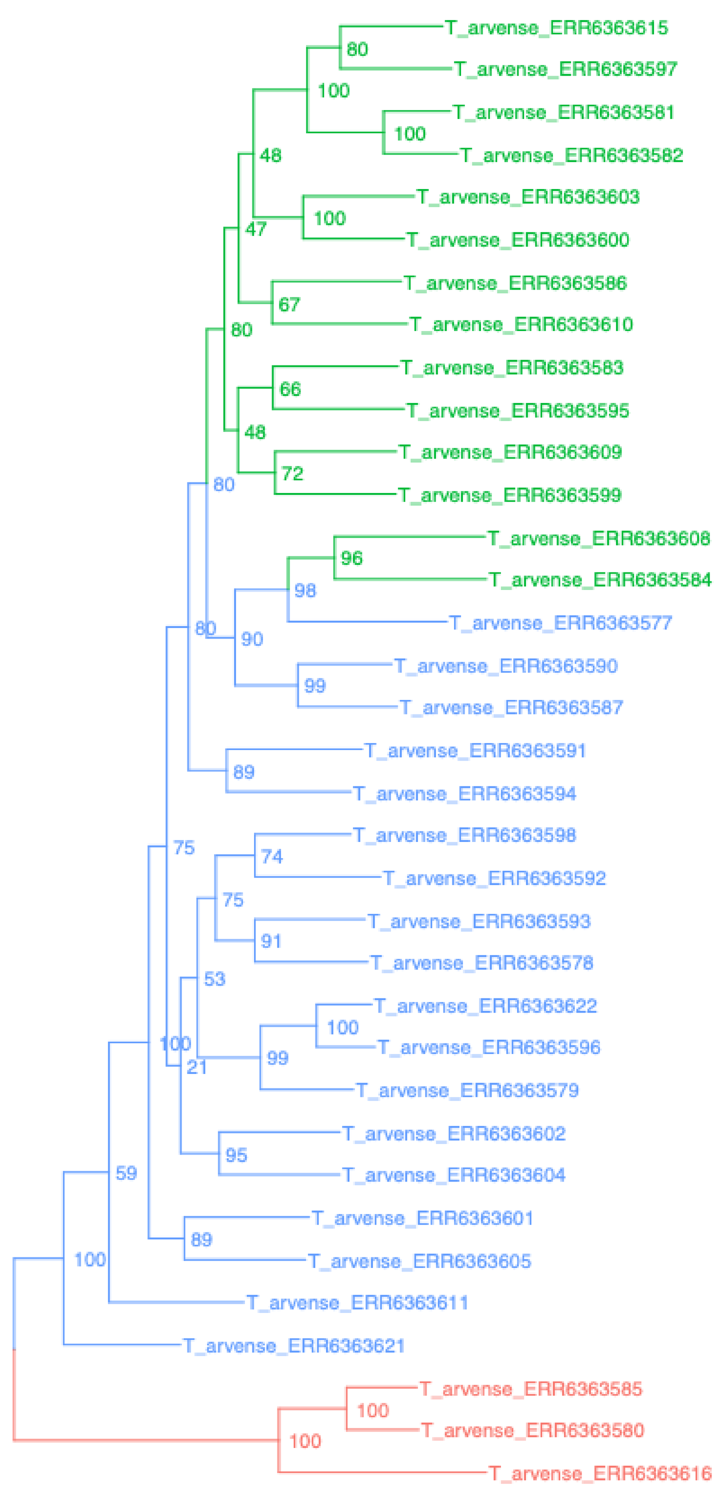


rooted_trees – Thlapsi_SCsuper

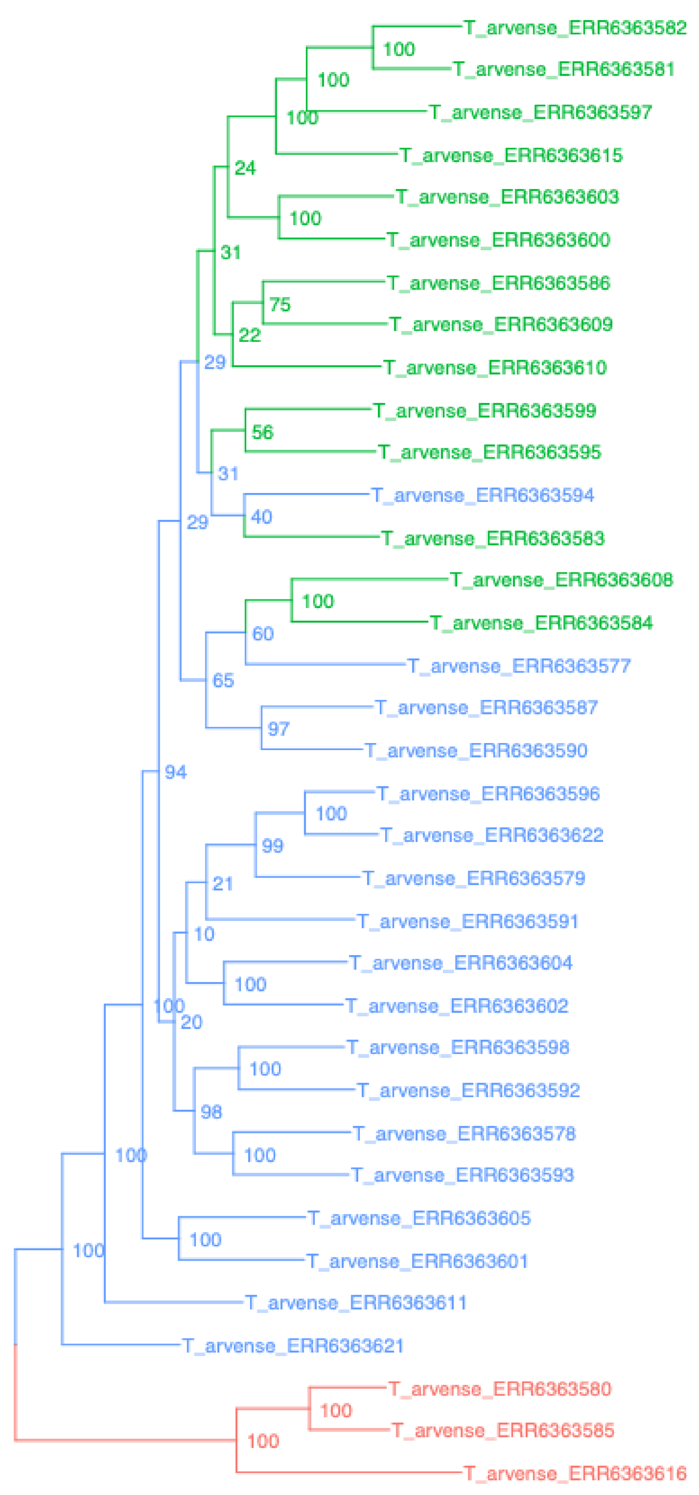


rooted_trees – Thlapsi_genome

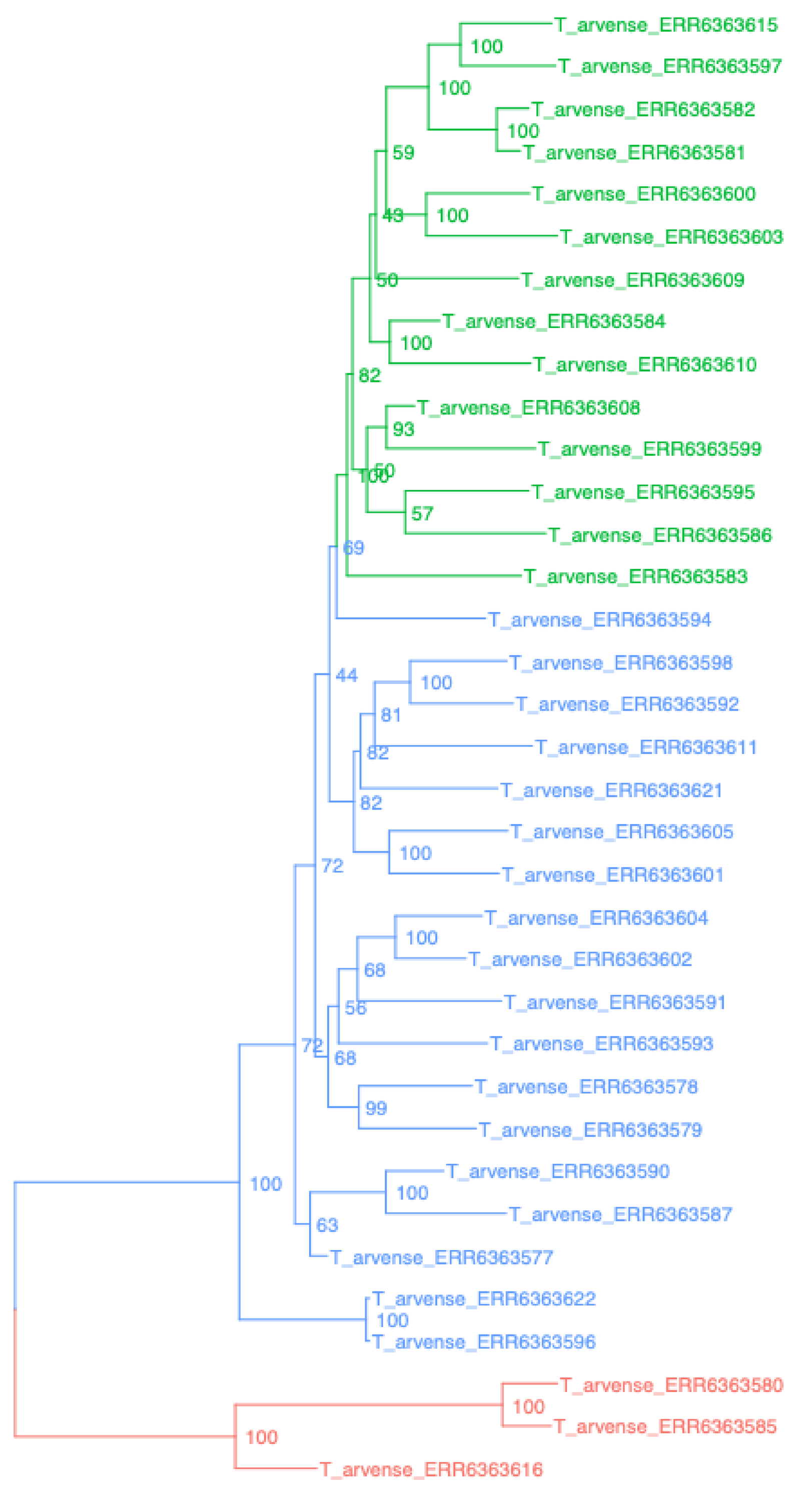

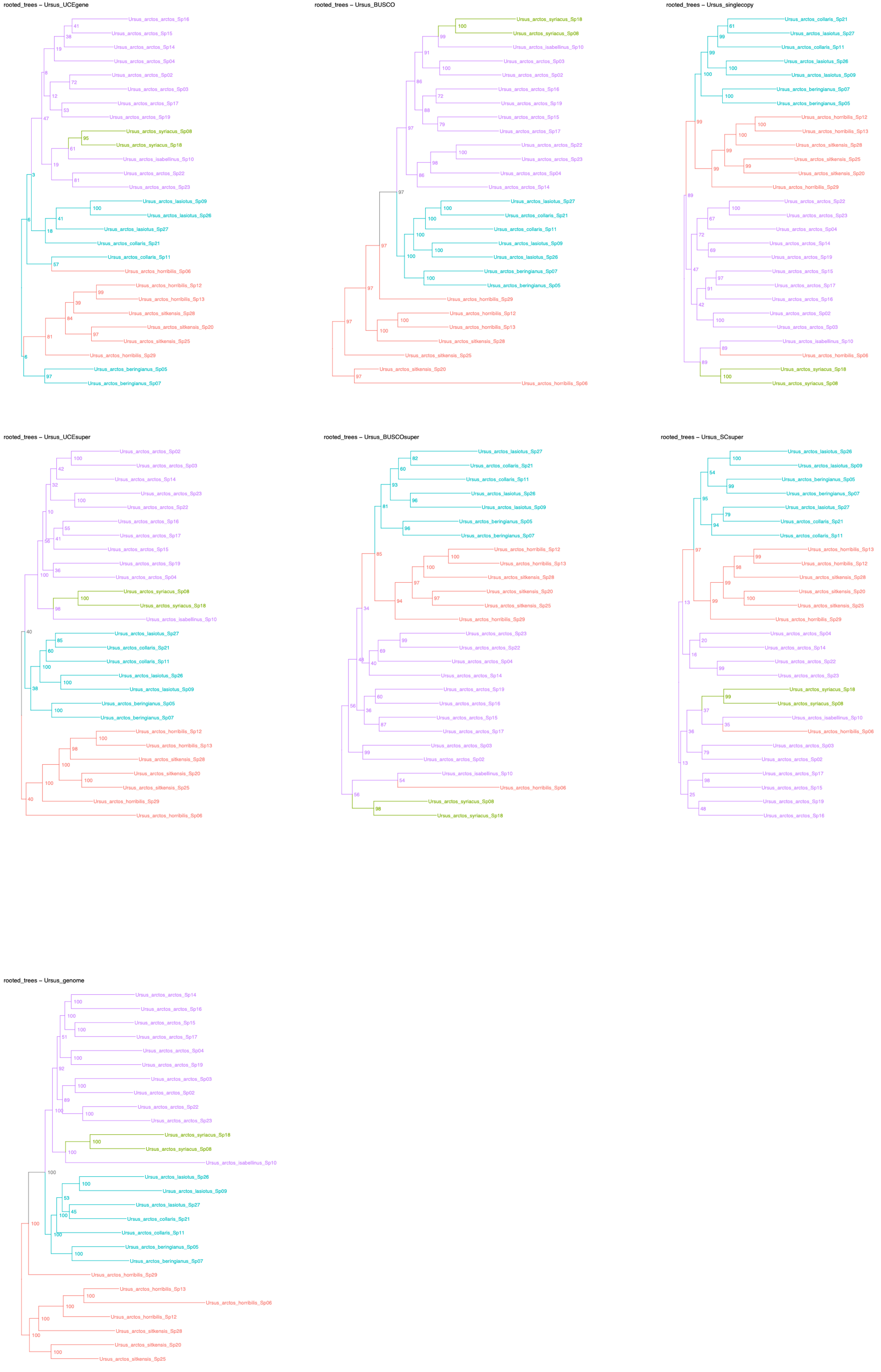
rooted_trees – Ursus_UCEgene
Ursus_arctos_arctos_Sp16
Ursus_arctos_arctos_Sp15
Ursus_arctos_arctos_Sp14
Ursus_arctos_arctos_Sp04
Ursus_arctos_arctos_Sp02
Ursus_arctos_arctos_Sp03
Ursus_arctos_arctos_Sp17
Ursus_arctos_arctos_Sp19
Ursus_arctos_syriacus_Sp08
Ursus_arctos_syriacus_Sp18
Ursus_arctos_isabellinus_Sp10
Ursus_arctos_arctos_Sp22
Ursus_arctos_arctos_Sp23
Ursus_arctos_lasiotus_Sp09
Ursus_arctos_lasiotus_Sp26
Ursus_arctos_lasiotus_Sp27
Ursus_arctos_collaris_Sp21
Ursus_arctos_collaris_Sp11
Ursus_arctos_horribilis_Sp06
Ursus_arctos_horribilis_Sp12
Ursus_arctos_horribilis_Sp13
Ursus_arctos_sitkensis_Sp28
Ursus_arctos_sitkensis_Sp20
Ursus_arctos_sitkensis_Sp25
Ursus_arctos_horribilis_Sp29
Ursus_arctos_beringianus_Sp05
Ursus_arctos_beringianus_Sp07
rooted_trees – Ursus_BUSCO
Ursus_arctos_syriacus_Sp18
Ursus_arctos_syriacus_Sp08
Ursus_arctos_isabellinus_Sp10
Ursus_arctos_arctos_Sp03
Ursus_arctos_arctos_Sp02
Ursus_arctos_arctos_Sp16
Ursus_arctos_arctos_Sp19
Ursus_arctos_arctos_Sp15
Ursus_arctos_arctos_Sp17
Ursus_arctos_arctos_Sp22
Ursus_arctos_arctos_Sp23
Ursus_arctos_arctos_Sp04
Ursus_arctos_arctos_Sp14
Ursus_arctos_lasiotus_Sp27
Ursus_arctos_collaris_Sp21
Ursus_arctos_collaris_Sp11
Ursus_arctos_lasiotus_Sp09
Ursus_arctos_lasiotus_Sp26
Ursus_arctos_beringianus_Sp07
Ursus_arctos_beringianus_Sp05
Ursus_arctos_horribilis_Sp29
Ursus_arctos_horribilis_Sp12
Ursus_arctos_horribilis_Sp13
Ursus_arctos_sitkensis_Sp28
Ursus_arctos_sitkensis_Sp25
Ursus_arctos_sitkensis_Sp20
Ursus_arctos_horribilis_Sp06
rooted_trees – Ursus_singlecopy
Ursus_arctos_collaris_Sp21
Ursus_arctos_lasiotus_Sp27
Ursus_arctos_collaris_Sp11
Ursus_arctos_lasiotus_Sp26
Ursus_arctos_lasiotus_Sp09
Ursus_arctos_beringianus_Sp07
Ursus_arctos_beringianus_Sp05
Ursus_arctos_horribilis_Sp12
Ursus_arctos_horribilis_Sp13
Ursus_arctos_sitkensis_Sp28
Ursus_arctos_sitkensis_Sp25
Ursus_arctos_sitkensis_Sp20
Ursus_arctos_horribilis_Sp29
Ursus_arctos_arctos_Sp22
Ursus_arctos_arctos_Sp23
Ursus_arctos_arctos_Sp04
Ursus_arctos_arctos_Sp14
Ursus_arctos_arctos_Sp19
Ursus_arctos_arctos_Sp15
Ursus_arctos_arctos_Sp17
Ursus_arctos_arctos_Sp16
Ursus_arctos_arctos_Sp02
Ursus_arctos_arctos_Sp03
Ursus_arctos_isabellinus_Sp10
Ursus_arctos_horribilis_Sp06
Ursus_arctos_syriacus_Sp18
Ursus_arctos_syriacus_Sp08
rooted_trees – Ursus_UCEsuper
Ursus_arctos_arctos_Sp02
Ursus_arctos_arctos_Sp03
Ursus_arctos_arctos_Sp14
Ursus_arctos_arctos_Sp23
Ursus_arctos_arctos_Sp22
Ursus_arctos_arctos_Sp16
Ursus_arctos_arctos_Sp17
Ursus_arctos_arctos_Sp15
Ursus_arctos_arctos_Sp19
Ursus_arctos_arctos_Sp04
Ursus_arctos_syriacus_Sp08
Ursus_arctos_syriacus_Sp18
Ursus_arctos_isabellinus_Sp10
Ursus_arctos_lasiotus_Sp27
Ursus_arctos_collaris_Sp21
Ursus_arctos_collaris_Sp11
Ursus_arctos_lasiotus_Sp26
Ursus_arctos_lasiotus_Sp09
Ursus_arctos_beringianus_Sp05
Ursus_arctos_beringianus_Sp07
Ursus_arctos_horribilis_Sp12
Ursus_arctos_horribilis_Sp13
Ursus_arctos_sitkensis_Sp28
Ursus_arctos_sitkensis_Sp20
Ursus_arctos_sitkensis_Sp25
Ursus_arctos_horribilis_Sp29
Ursus_arctos_horribilis_Sp06
rooted_trees – Ursus_BUSCOsuper
Ursus_arctos_lasiotus_Sp27
Ursus_arctos_collaris_Sp21
Ursus_arctos_collaris_Sp11
Ursus_arctos_lasiotus_Sp26
Ursus_arctos_lasiotus_Sp09
Ursus_arctos_beringianus_Sp05
Ursus_arctos_beringianus_Sp07
Ursus_arctos_horribilis_Sp12
Ursus_arctos_horribilis_Sp13
Ursus_arctos_sitkensis_Sp28
Ursus_arctos_sitkensis_Sp20
Ursus_arctos_sitkensis_Sp25
Ursus_arctos_horribilis_Sp29
Ursus_arctos_arctos_Sp23
Ursus_arctos_arctos_Sp22
Ursus_arctos_arctos_Sp04
Ursus_arctos_arctos_Sp14
Ursus_arctos_arctos_Sp19
Ursus_arctos_arctos_Sp16
Ursus_arctos_arctos_Sp15
Ursus_arctos_arctos_Sp17
Ursus_arctos_arctos_Sp03
Ursus_arctos_arctos_Sp02
Ursus_arctos_isabellinus_Sp10
Ursus_arctos_horribilis_Sp06
Ursus_arctos_syriacus_Sp08
Ursus_arctos_syriacus_Sp18
rooted_trees – Ursus_SCsuper
Ursus_arctos_lasiotus_Sp26
Ursus_arctos_lasiotus_Sp09
Ursus_arctos_beringianus_Sp05
Ursus_arctos_beringianus_Sp07
Ursus_arctos_lasiotus_Sp27
Ursus_arctos_collaris_Sp21
Ursus_arctos_collaris_Sp11
Ursus_arctos_horribilis_Sp13
Ursus_arctos_horribilis_Sp12
Ursus_arctos_sitkensis_Sp28
Ursus_arctos_sitkensis_Sp20
Ursus_arctos_sitkensis_Sp25
Ursus_arctos_horribilis_Sp29
Ursus_arctos_arctos_Sp04
Ursus_arctos_arctos_Sp14
Ursus_arctos_arctos_Sp22
Ursus_arctos_arctos_Sp23
Ursus_arctos_syriacus_Sp18
Ursus_arctos_syriacus_Sp08
Ursus_arctos_isabellinus_Sp10
Ursus_arctos_horribilis_Sp06
Ursus_arctos_arctos_Sp03
Ursus_arctos_arctos_Sp02
Ursus_arctos_arctos_Sp17
Ursus_arctos_arctos_Sp15
Ursus_arctos_arctos_Sp19
Ursus_arctos_arctos_Sp16
rooted_trees – Ursus_genome
Ursus_arctos_arctos_Sp14
Ursus_arctos_arctos_Sp16
Ursus_arctos_arctos_Sp15
Ursus_arctos_arctos_Sp17
Ursus_arctos_arctos_Sp04
Ursus_arctos_arctos_Sp19
Ursus_arctos_arctos_Sp03
Ursus_arctos_arctos_Sp02
Ursus_arctos_arctos_Sp22
Ursus_arctos_arctos_Sp23
Ursus_arctos_syriacus_Sp18
Ursus_arctos_syriacus_Sp08
Ursus_arctos_isabellinus_Sp10
Ursus_arctos_lasiotus_Sp26
Ursus_arctos_lasiotus_Sp09
Ursus_arctos_lasiotus_Sp27
Ursus_arctos_collaris_Sp21
Ursus_arctos_collaris_Sp11
Ursus_arctos_beringianus_Sp05
Ursus_arctos_beringianus_Sp07
Ursus_arctos_horribilis_Sp29
Ursus_arctos_horribilis_Sp13
Ursus_arctos_horribilis_Sp06
Ursus_arctos_horribilis_Sp12
Ursus_arctos_sitkensis_Sp28
Ursus_arctos_sitkensis_Sp20
Ursus_arctos_sitkensis_Sp25

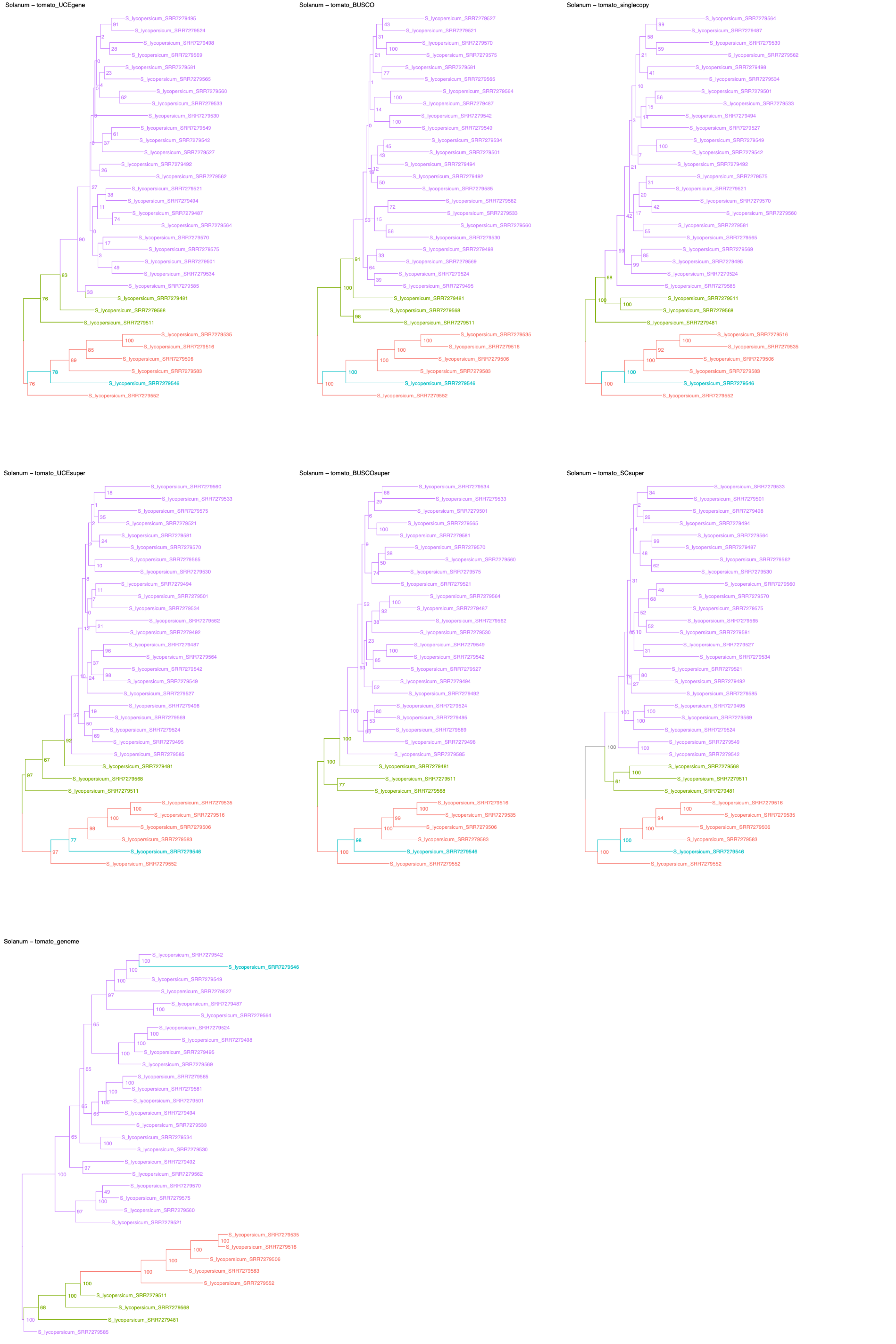
Solanum – tomato_UCEgene
S_lycopersicum_SRR7279495
S_lycopersicum_SRR7279524
S_lycopersicum_SRR7279498
S_lycopersicum_SRR7279569
S_lycopersicum_SRR7279581
S_lycopersicum_SRR7279565
S_lycopersicum_SRR7279560
S_lycopersicum_SRR7279533
S_lycopersicum_SRR7279530
S_lycopersicum_SRR7279549
S_lycopersicum_SRR7279542
S_lycopersicum_SRR7279527
S_lycopersicum_SRR7279492
S_lycopersicum_SRR7279562
S_lycopersicum_SRR7279521
S_lycopersicum_SRR7279494
S_lycopersicum_SRR7279487
S_lycopersicum_SRR7279564
S_lycopersicum_SRR7279570
S_lycopersicum_SRR7279575
S_lycopersicum_SRR7279501
S_lycopersicum_SRR7279534
S_lycopersicum_SRR7279585
S_lycopersicum_SRR7279481
S_lycopersicum_SRR7279568
S_lycopersicum_SRR7279511
S_lycopersicum_SRR7279535
S_lycopersicum_SRR7279516
S_lycopersicum_SRR7279506
S_lycopersicum_SRR7279583
S_lycopersicum_SRR7279546
S_lycopersicum_SRR7279552
Solanum – tomato_BUSCO
S_lycopersicum_SRR7279527
S_lycopersicum_SRR7279521
S_lycopersicum_SRR7279570
S_lycopersicum_SRR7279575
S_lycopersicum_SRR7279581
S_lycopersicum_SRR7279565
S_lycopersicum_SRR7279564
S_lycopersicum_SRR7279487
S_lycopersicum_SRR7279542
S_lycopersicum_SRR7279549
S_lycopersicum_SRR7279534
S_lycopersicum_SRR7279501
S_lycopersicum_SRR7279494
S_lycopersicum_SRR7279492
S_lycopersicum_SRR7279585
S_lycopersicum_SRR7279562
S_lycopersicum_SRR7279533
S_lycopersicum_SRR7279560
S_lycopersicum_SRR7279530
S_lycopersicum_SRR7279498
S_lycopersicum_SRR7279569
S_lycopersicum_SRR7279524
S_lycopersicum_SRR7279495
S_lycopersicum_SRR7279481
S_lycopersicum_SRR7279568
S_lycopersicum_SRR7279511
S_lycopersicum_SRR7279535
S_lycopersicum_SRR7279516
S_lycopersicum_SRR7279506
S_lycopersicum_SRR7279583
S_lycopersicum_SRR7279546
S_lycopersicum_SRR7279552
Solanum – tomato_singlecopy
S_lycopersicum_SRR7279564
S_lycopersicum_SRR7279487
S_lycopersicum_SRR7279530
S_lycopersicum_SRR7279562
S_lycopersicum_SRR7279498
S_lycopersicum_SRR7279534
S_lycopersicum_SRR7279501
S_lycopersicum_SRR7279533
S_lycopersicum_SRR7279494
S_lycopersicum_SRR7279527
S_lycopersicum_SRR7279549
S_lycopersicum_SRR7279542
S_lycopersicum_SRR7279492
S_lycopersicum_SRR7279575
S_lycopersicum_SRR7279521
S_lycopersicum_SRR7279570
S_lycopersicum_SRR7279560
S_lycopersicum_SRR7279581
S_lycopersicum_SRR7279565
S_lycopersicum_SRR7279569
S_lycopersicum_SRR7279495
S_lycopersicum_SRR7279524
S_lycopersicum_SRR7279585
S_lycopersicum_SRR7279511
S_lycopersicum_SRR7279568
S_lycopersicum_SRR7279481
S_lycopersicum_SRR7279516
S_lycopersicum_SRR7279535
S_lycopersicum_SRR7279506
S_lycopersicum_SRR7279583
S_lycopersicum_SRR7279546
S_lycopersicum_SRR7279552
Solanum – tomato_UCEsuper
S_lycopersicum_SRR7279560
S_lycopersicum_SRR7279533
S_lycopersicum_SRR7279575
S_lycopersicum_SRR7279521
S_lycopersicum_SRR7279581
S_lycopersicum_SRR7279570
S_lycopersicum_SRR7279565
S_lycopersicum_SRR7279530
S_lycopersicum_SRR7279494
S_lycopersicum_SRR7279501
S_lycopersicum_SRR7279534
S_lycopersicum_SRR7279562
S_lycopersicum_SRR7279492
S_lycopersicum_SRR7279487
S_lycopersicum_SRR7279564
S_lycopersicum_SRR7279542
S_lycopersicum_SRR7279549
S_lycopersicum_SRR7279527
S_lycopersicum_SRR7279498
S_lycopersicum_SRR7279569
S_lycopersicum_SRR7279524
S_lycopersicum_SRR7279495
S_lycopersicum_SRR7279585
S_lycopersicum_SRR7279481
S_lycopersicum_SRR7279568
S_lycopersicum_SRR7279511
S_lycopersicum_SRR7279535
S_lycopersicum_SRR7279516
S_lycopersicum_SRR7279506
S_lycopersicum_SRR7279583
S_lycopersicum_SRR7279546
S_lycopersicum_SRR7279552
Solanum – tomato_BUSCOsuper
S_lycopersicum_SRR7279534
S_lycopersicum_SRR7279533
S_lycopersicum_SRR7279501
S_lycopersicum_SRR7279565
S_lycopersicum_SRR7279581
S_lycopersicum_SRR7279570
S_lycopersicum_SRR7279560
S_lycopersicum_SRR7279575
S_lycopersicum_SRR7279521
S_lycopersicum_SRR7279564
S_lycopersicum_SRR7279487
S_lycopersicum_SRR7279562
S_lycopersicum_SRR7279530
S_lycopersicum_SRR7279549
S_lycopersicum_SRR7279542
S_lycopersicum_SRR7279527
S_lycopersicum_SRR7279494
S_lycopersicum_SRR7279492
S_lycopersicum_SRR7279524
S_lycopersicum_SRR7279495
S_lycopersicum_SRR7279569
S_lycopersicum_SRR7279498
S_lycopersicum_SRR7279585
S_lycopersicum_SRR7279481
S_lycopersicum_SRR7279511
S_lycopersicum_SRR7279568
S_lycopersicum_SRR7279516
S_lycopersicum_SRR7279535
S_lycopersicum_SRR7279506
S_lycopersicum_SRR7279583
S_lycopersicum_SRR7279546
S_lycopersicum_SRR7279552
Solanum – tomato_SCsuper
S_lycopersicum_SRR7279533
S_lycopersicum_SRR7279501
S_lycopersicum_SRR7279498
S_lycopersicum_SRR7279494
S_lycopersicum_SRR7279564
S_lycopersicum_SRR7279487
S_lycopersicum_SRR7279562
S_lycopersicum_SRR7279530
S_lycopersicum_SRR7279560
S_lycopersicum_SRR7279570
S_lycopersicum_SRR7279575
S_lycopersicum_SRR7279565
S_lycopersicum_SRR7279581
S_lycopersicum_SRR7279527
S_lycopersicum_SRR7279534
S_lycopersicum_SRR7279521
S_lycopersicum_SRR7279492
S_lycopersicum_SRR7279585
S_lycopersicum_SRR7279495
S_lycopersicum_SRR7279569
S_lycopersicum_SRR7279524
S_lycopersicum_SRR7279549
S_lycopersicum_SRR7279542
S_lycopersicum_SRR7279568
S_lycopersicum_SRR7279511
S_lycopersicum_SRR7279481
S_lycopersicum_SRR7279516
S_lycopersicum_SRR7279535
S_lycopersicum_SRR7279506
S_lycopersicum_SRR7279583
S_lycopersicum_SRR7279546
S_lycopersicum_SRR7279552
Solanum – tomato_genome
S_lycopersicum_SRR7279542
S_lycopersicum_SRR7279546
S_lycopersicum_SRR7279549
S_lycopersicum_SRR7279527
S_lycopersicum_SRR7279487
S_lycopersicum_SRR7279564
S_lycopersicum_SRR7279524
S_lycopersicum_SRR7279498
S_lycopersicum_SRR7279495
S_lycopersicum_SRR7279569
S_lycopersicum_SRR7279565
S_lycopersicum_SRR7279581
S_lycopersicum_SRR7279501
S_lycopersicum_SRR7279494
S_lycopersicum_SRR7279533
S_lycopersicum_SRR7279534
S_lycopersicum_SRR7279530
S_lycopersicum_SRR7279492
S_lycopersicum_SRR7279562
S_lycopersicum_SRR7279570
S_lycopersicum_SRR7279575
S_lycopersicum_SRR7279560
S_lycopersicum_SRR7279521
S_lycopersicum_SRR7279535
S_lycopersicum_SRR7279516
S_lycopersicum_SRR7279506
S_lycopersicum_SRR7279583
S_lycopersicum_SRR7279552
S_lycopersicum_SRR7279511
S_lycopersicum_SRR7279568
S_lycopersicum_SRR7279481
S_lycopersicum_SRR7279585

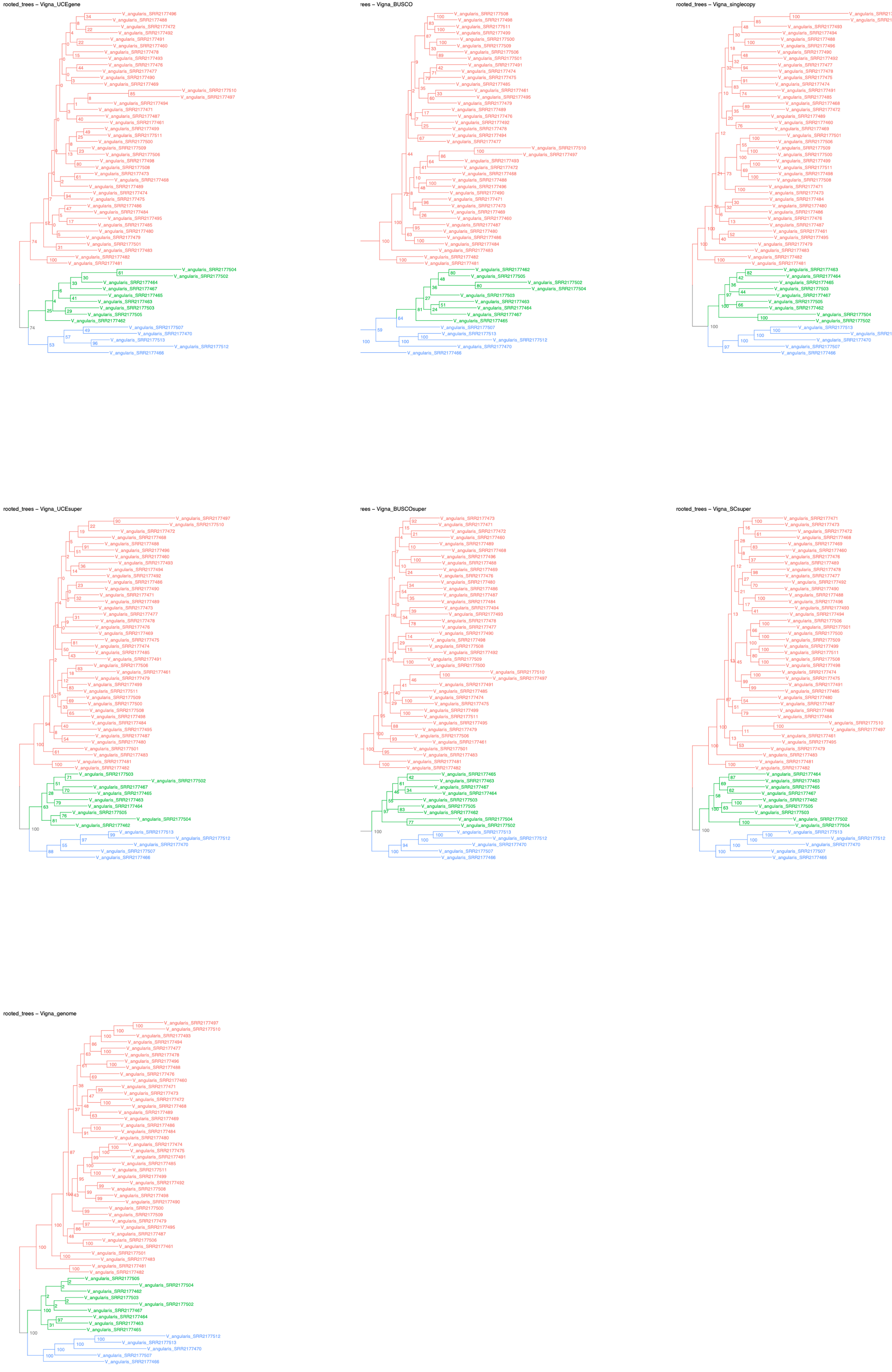
rooted_trees – Vigna_UCEgene
rees – Vigna_BUSCO
rooted_trees – Vigna_singlecopy
rooted_trees – Vigna_UCEsuper
rees – Vigna_BUSCOsuper
rooted_trees – Vigna_SCsuper
rooted_trees – Vigna_genome

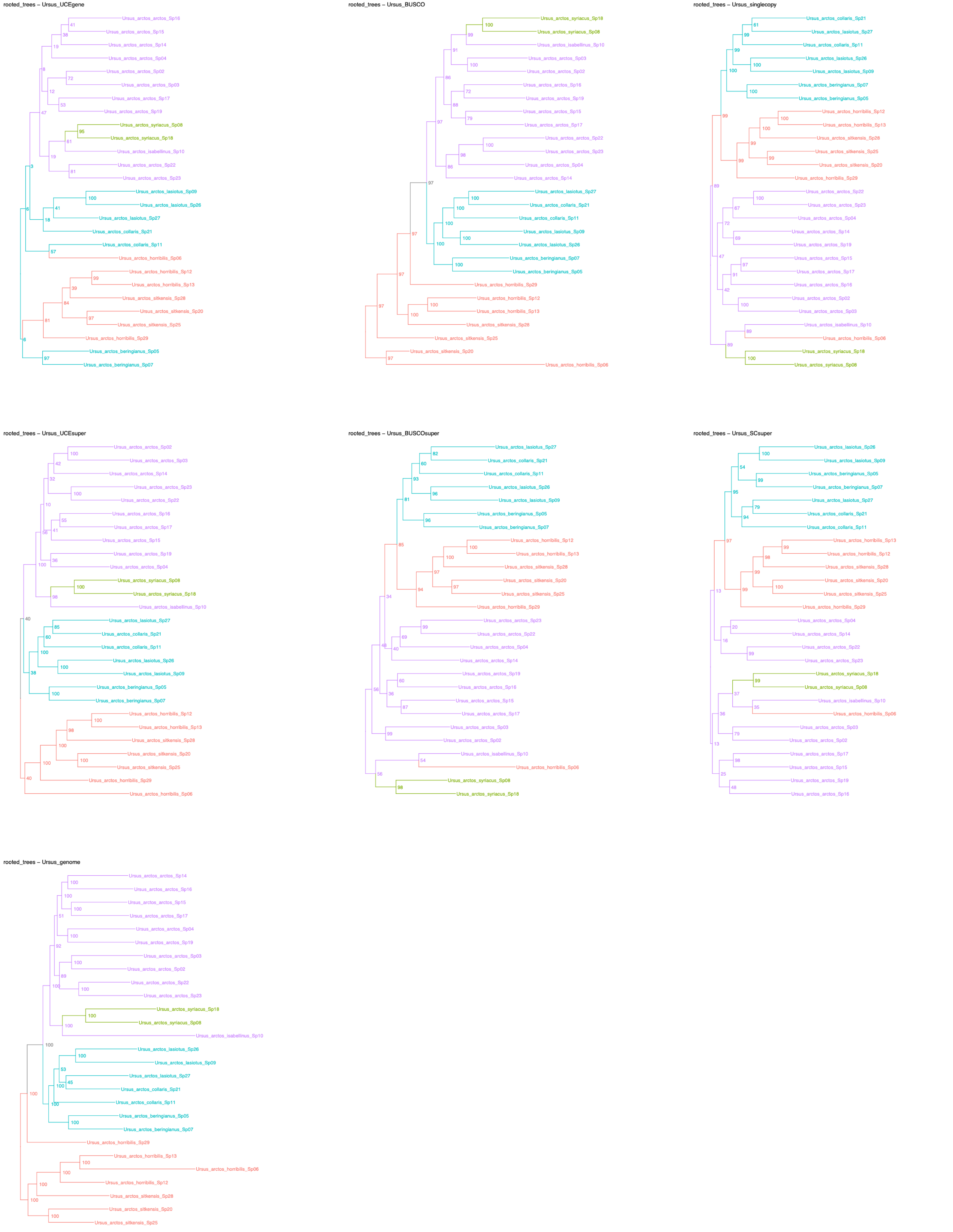

rooted_trees – Ursus_UCEgene
Ursus_arctos_arctos_Sp16
Ursus_arctos_arctos_Sp15
Ursus_arctos_arctos_Sp14
Ursus_arctos_arctos_Sp04
Ursus_arctos_arctos_Sp02
Ursus_arctos_arctos_Sp03
Ursus_arctos_arctos_Sp17
Ursus_arctos_arctos_Sp19
Ursus_arctos_syriacus_Sp08
Ursus_arctos_syriacus_Sp18
Ursus_arctos_isabellinus_Sp10
Ursus_arctos_arctos_Sp22
Ursus_arctos_arctos_Sp23
Ursus_arctos_lasiotus_Sp09
Ursus_arctos_lasiotus_Sp26
Ursus_arctos_lasiotus_Sp27
Ursus_arctos_collaris_Sp21
Ursus_arctos_collaris_Sp11
Ursus_arctos_horribilis_Sp06
Ursus_arctos_horribilis_Sp12
Ursus_arctos_horribilis_Sp13
Ursus_arctos_sitkensis_Sp28
Ursus_arctos_sitkensis_Sp20
Ursus_arctos_sitkensis_Sp25
Ursus_arctos_horribilis_Sp29
Ursus_arctos_beringianus_Sp05
Ursus_arctos_beringianus_Sp07
rooted_trees – Ursus_BUSCO
Ursus_arctos_syriacus_Sp18
Ursus_arctos_syriacus_Sp08
Ursus_arctos_isabellinus_Sp10
Ursus_arctos_arctos_Sp03
Ursus_arctos_arctos_Sp02
Ursus_arctos_arctos_Sp16
Ursus_arctos_arctos_Sp19
Ursus_arctos_arctos_Sp15
Ursus_arctos_arctos_Sp17
Ursus_arctos_arctos_Sp22
Ursus_arctos_arctos_Sp23
Ursus_arctos_arctos_Sp04
Ursus_arctos_arctos_Sp14
Ursus_arctos_lasiotus_Sp27
Ursus_arctos_collaris_Sp21
Ursus_arctos_collaris_Sp11
Ursus_arctos_lasiotus_Sp09
Ursus_arctos_lasiotus_Sp26
Ursus_arctos_beringianus_Sp07
Ursus_arctos_beringianus_Sp05
Ursus_arctos_horribilis_Sp29
Ursus_arctos_horribilis_Sp12
Ursus_arctos_horribilis_Sp13
Ursus_arctos_sitkensis_Sp28
Ursus_arctos_sitkensis_Sp25
Ursus_arctos_sitkensis_Sp20
Ursus_arctos_horribilis_Sp06
rooted_trees – Ursus_singlecopy
Ursus_arctos_collaris_Sp21
Ursus_arctos_lasiotus_Sp27
Ursus_arctos_collaris_Sp11
Ursus_arctos_lasiotus_Sp26
Ursus_arctos_lasiotus_Sp09
Ursus_arctos_beringianus_Sp07
Ursus_arctos_beringianus_Sp05
Ursus_arctos_horribilis_Sp12
Ursus_arctos_horribilis_Sp13
Ursus_arctos_sitkensis_Sp28
Ursus_arctos_sitkensis_Sp25
Ursus_arctos_sitkensis_Sp20
Ursus_arctos_horribilis_Sp29
Ursus_arctos_arctos_Sp22
Ursus_arctos_arctos_Sp23
Ursus_arctos_arctos_Sp04
Ursus_arctos_arctos_Sp14
Ursus_arctos_arctos_Sp19
Ursus_arctos_arctos_Sp15
Ursus_arctos_arctos_Sp17
Ursus_arctos_arctos_Sp16
Ursus_arctos_arctos_Sp02
Ursus_arctos_arctos_Sp03
Ursus_arctos_isabellinus_Sp10
Ursus_arctos_horribilis_Sp06
Ursus_arctos_syriacus_Sp18
Ursus_arctos_syriacus_Sp08
rooted_trees – Ursus_UCEsuper
Ursus_arctos_arctos_Sp02
Ursus_arctos_arctos_Sp03
Ursus_arctos_arctos_Sp14
Ursus_arctos_arctos_Sp23
Ursus_arctos_arctos_Sp22
Ursus_arctos_arctos_Sp16
Ursus_arctos_arctos_Sp17
Ursus_arctos_arctos_Sp15
Ursus_arctos_arctos_Sp19
Ursus_arctos_arctos_Sp04
Ursus_arctos_syriacus_Sp08
Ursus_arctos_syriacus_Sp18
Ursus_arctos_isabellinus_Sp10
Ursus_arctos_lasiotus_Sp27
Ursus_arctos_collaris_Sp21
Ursus_arctos_collaris_Sp11
Ursus_arctos_lasiotus_Sp26
Ursus_arctos_lasiotus_Sp09
Ursus_arctos_beringianus_Sp05
Ursus_arctos_beringianus_Sp07
Ursus_arctos_horribilis_Sp12
Ursus_arctos_horribilis_Sp13
Ursus_arctos_sitkensis_Sp28
Ursus_arctos_sitkensis_Sp20
Ursus_arctos_sitkensis_Sp25
Ursus_arctos_horribilis_Sp29
Ursus_arctos_horribilis_Sp06
rooted_trees – Ursus_BUSCOsuper
Ursus_arctos_lasiotus_Sp27
Ursus_arctos_collaris_Sp21
Ursus_arctos_collaris_Sp11
Ursus_arctos_lasiotus_Sp26
Ursus_arctos_lasiotus_Sp09
Ursus_arctos_beringianus_Sp05
Ursus_arctos_beringianus_Sp07
Ursus_arctos_horribilis_Sp12
Ursus_arctos_horribilis_Sp13
Ursus_arctos_sitkensis_Sp28
Ursus_arctos_sitkensis_Sp20
Ursus_arctos_sitkensis_Sp25
Ursus_arctos_horribilis_Sp29
Ursus_arctos_arctos_Sp23
Ursus_arctos_arctos_Sp22
Ursus_arctos_arctos_Sp04
Ursus_arctos_arctos_Sp14
Ursus_arctos_arctos_Sp19
Ursus_arctos_arctos_Sp16
Ursus_arctos_arctos_Sp15
Ursus_arctos_arctos_Sp17
Ursus_arctos_arctos_Sp03
Ursus_arctos_arctos_Sp02
Ursus_arctos_isabellinus_Sp10
Ursus_arctos_horribilis_Sp06
Ursus_arctos_syriacus_Sp08
Ursus_arctos_syriacus_Sp18
rooted_trees – Ursus_SCsuper
Ursus_arctos_lasiotus_Sp26
Ursus_arctos_lasiotus_Sp09
Ursus_arctos_beringianus_Sp05
Ursus_arctos_beringianus_Sp07
Ursus_arctos_lasiotus_Sp27
Ursus_arctos_collaris_Sp21
Ursus_arctos_collaris_Sp11
Ursus_arctos_horribilis_Sp13
Ursus_arctos_horribilis_Sp12
Ursus_arctos_sitkensis_Sp28
Ursus_arctos_sitkensis_Sp20
Ursus_arctos_sitkensis_Sp25
Ursus_arctos_horribilis_Sp29
Ursus_arctos_arctos_Sp04
Ursus_arctos_arctos_Sp14
Ursus_arctos_arctos_Sp22
Ursus_arctos_arctos_Sp23
Ursus_arctos_syriacus_Sp18
Ursus_arctos_syriacus_Sp08
Ursus_arctos_isabellinus_Sp10
Ursus_arctos_horribilis_Sp06
Ursus_arctos_arctos_Sp03
Ursus_arctos_arctos_Sp02
Ursus_arctos_arctos_Sp17
Ursus_arctos_arctos_Sp15
Ursus_arctos_arctos_Sp19
Ursus_arctos_arctos_Sp16
rooted_trees – Ursus_genome
Ursus_arctos_arctos_Sp14
Ursus_arctos_arctos_Sp16
Ursus_arctos_arctos_Sp15
Ursus_arctos_arctos_Sp17
Ursus_arctos_arctos_Sp04
Ursus_arctos_arctos_Sp19
Ursus_arctos_arctos_Sp03
Ursus_arctos_arctos_Sp02
Ursus_arctos_arctos_Sp22
Ursus_arctos_arctos_Sp23
Ursus_arctos_syriacus_Sp18
Ursus_arctos_syriacus_Sp08
Ursus_arctos_isabellinus_Sp10
Ursus_arctos_lasiotus_Sp26
Ursus_arctos_lasiotus_Sp09
Ursus_arctos_lasiotus_Sp27
Ursus_arctos_collaris_Sp21
Ursus_arctos_collaris_Sp11
Ursus_arctos_beringianus_Sp05
Ursus_arctos_beringianus_Sp07
Ursus_arctos_horribilis_Sp29
Ursus_arctos_horribilis_Sp13
Ursus_arctos_horribilis_Sp06
Ursus_arctos_horribilis_Sp12
Ursus_arctos_sitkensis_Sp28
Ursus_arctos_sitkensis_Sp20
Ursus_arctos_sitkensis_Sp25

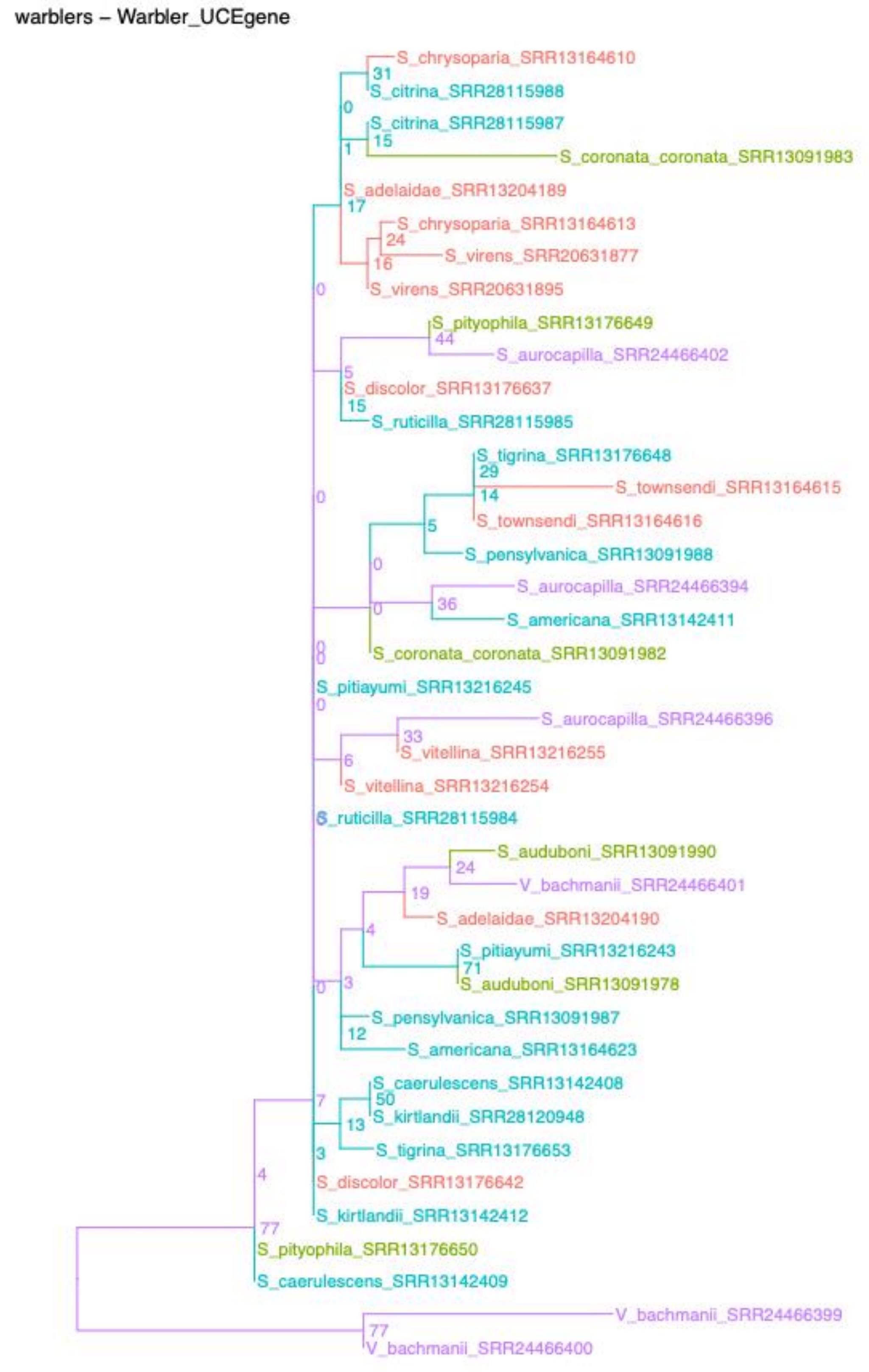
warblers – Warbler_UCEgene

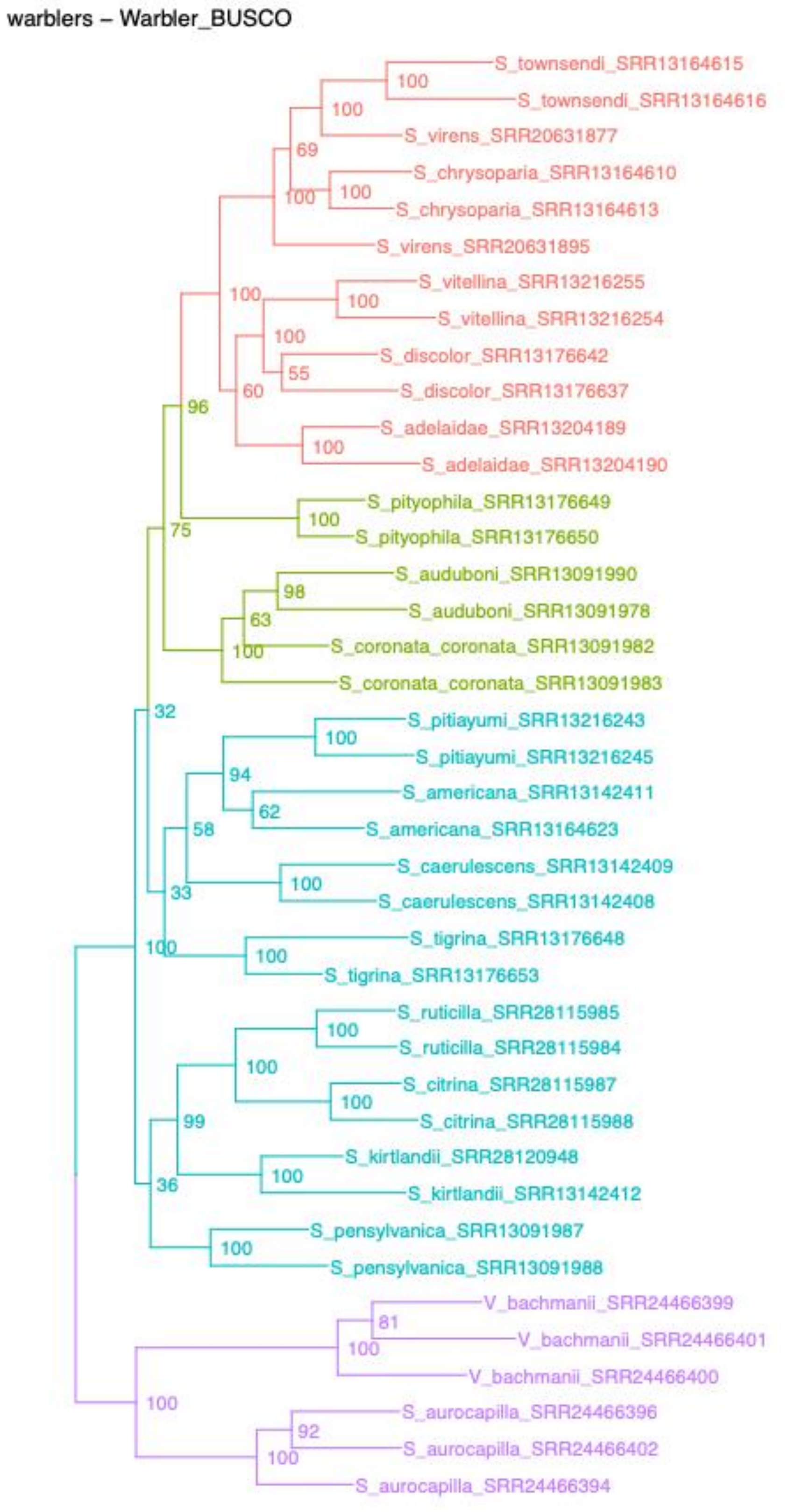
warblers – Warbler_BUSCO

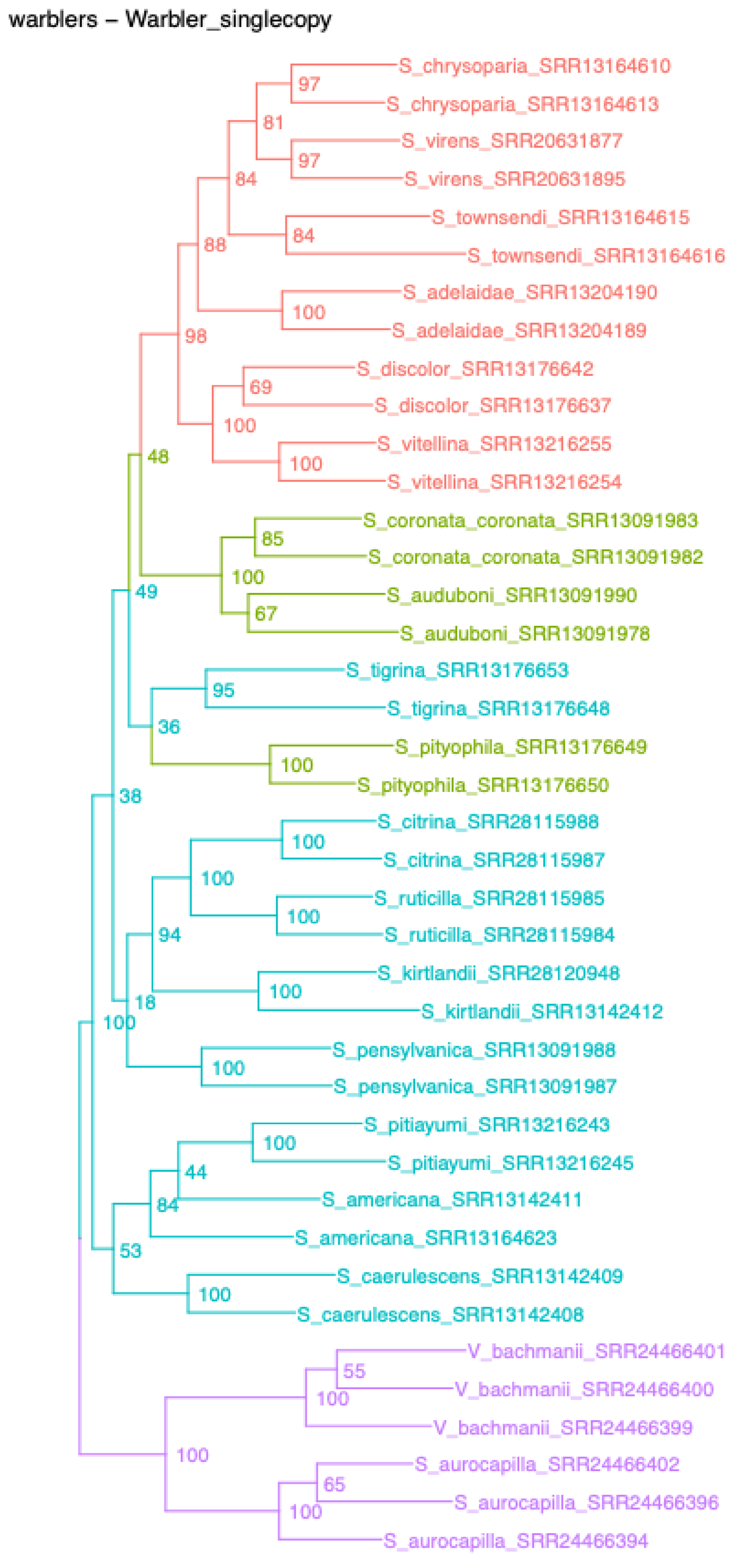
warblers – Warbler_singlecopy

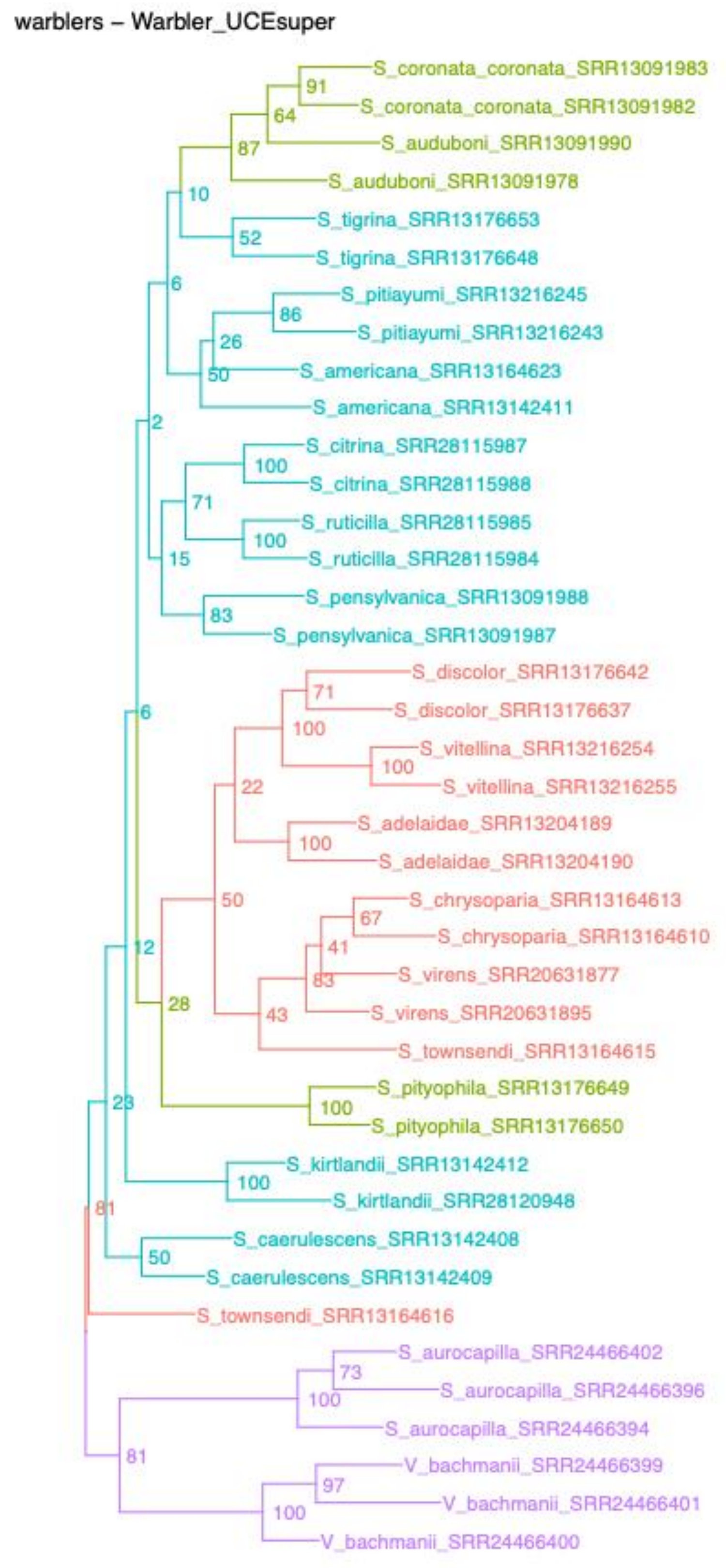
warblers – Warbler_UCEsuper

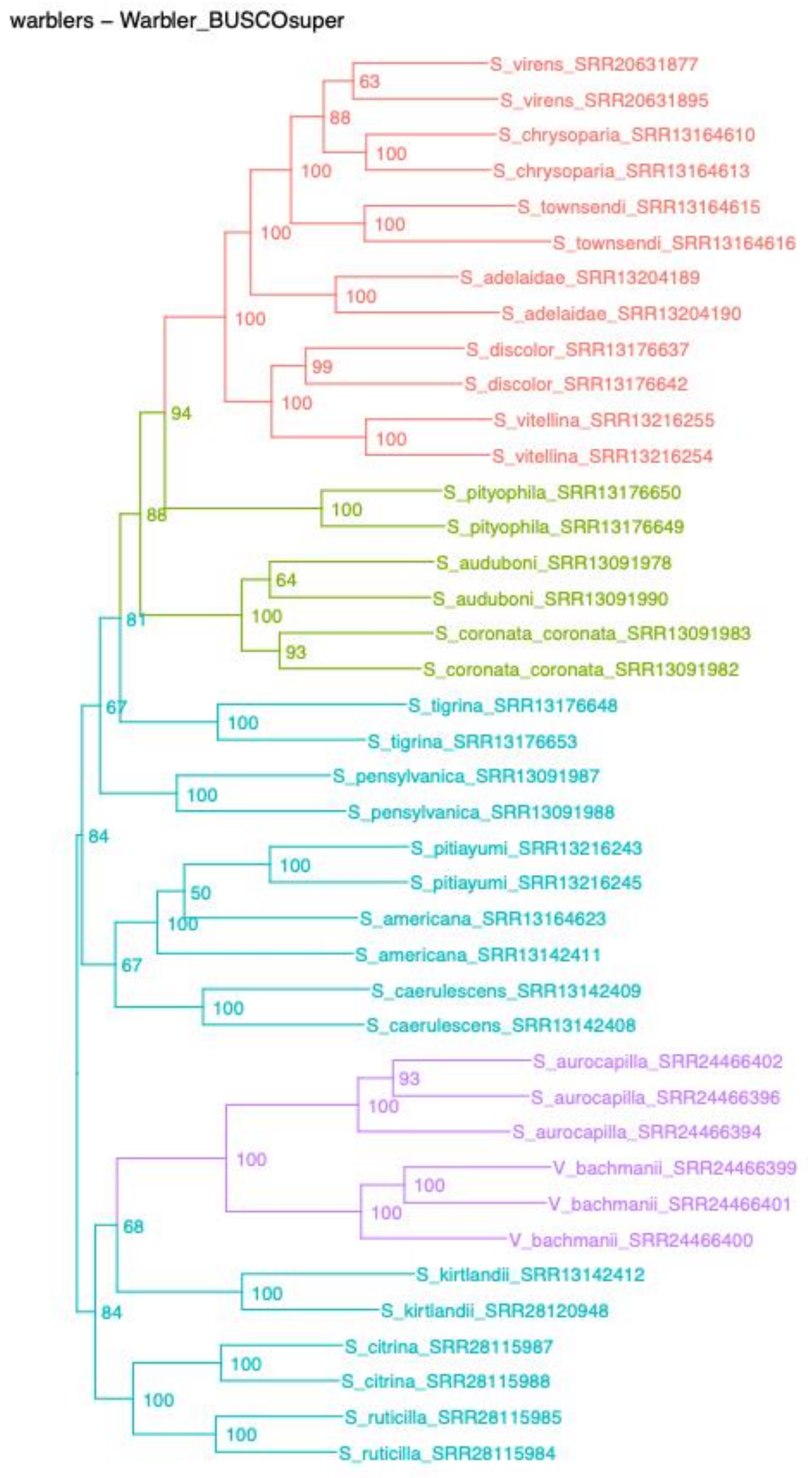
warblers – Warbler_BUSCOsuper

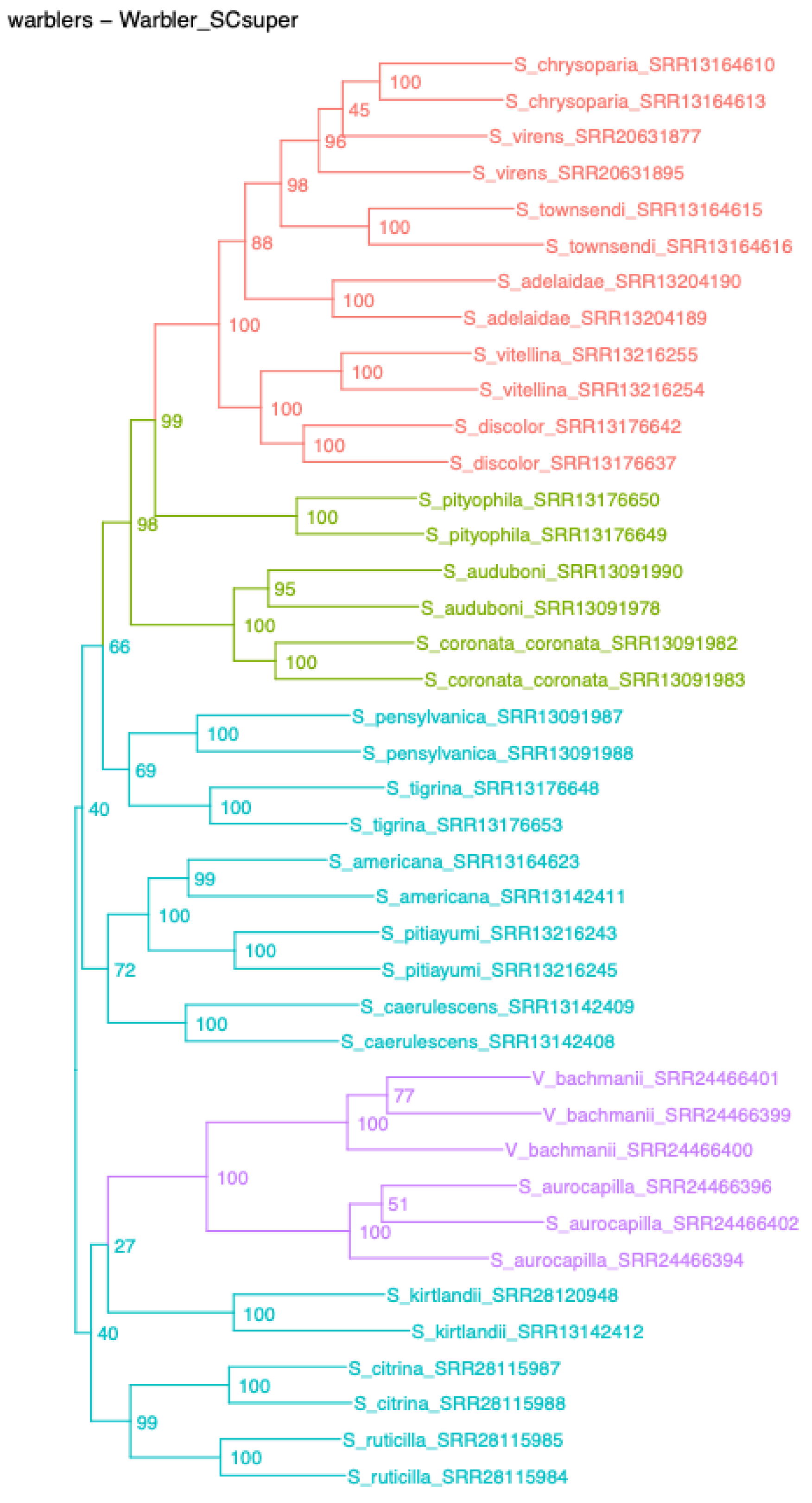
warblers – Warbler_SCsuper

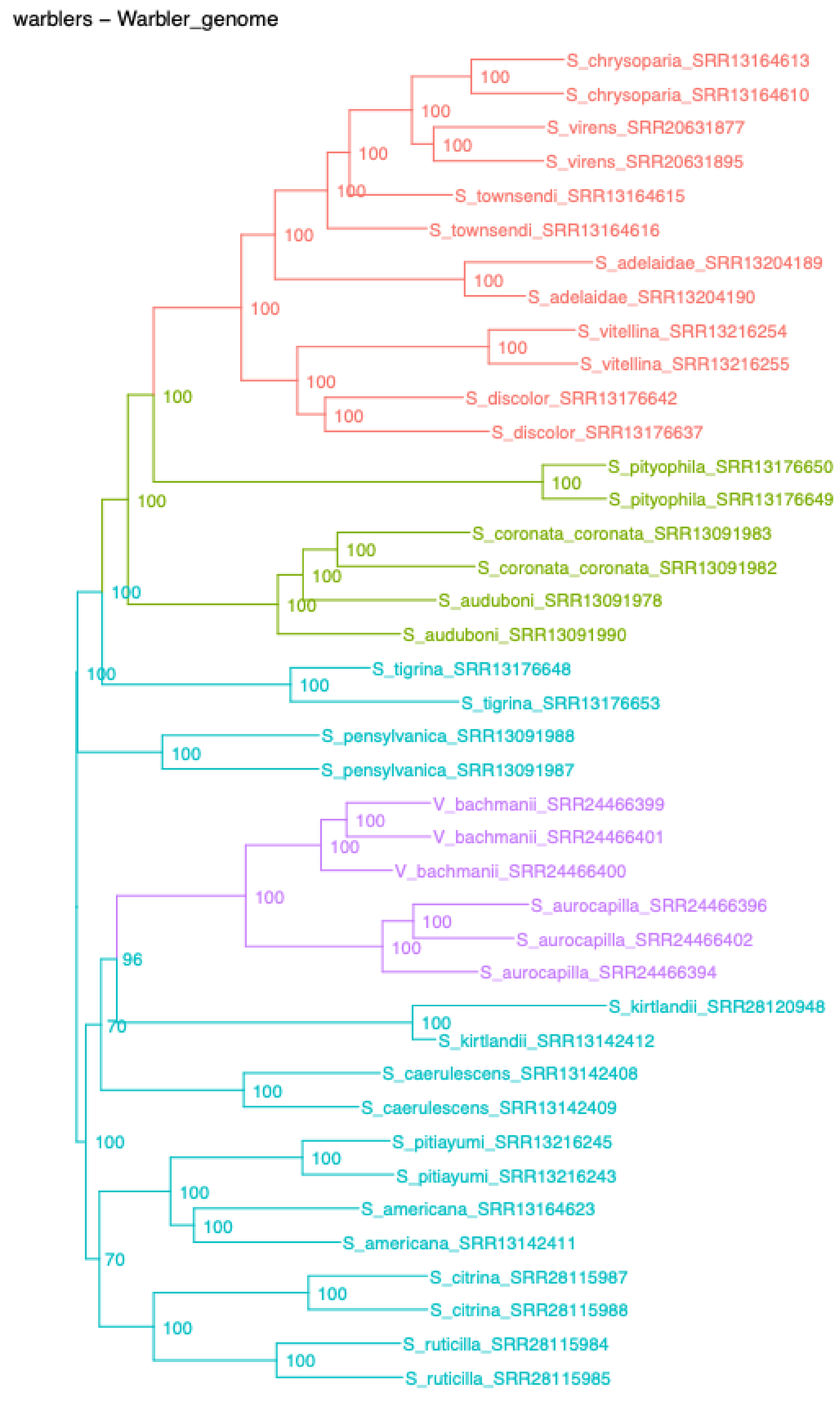
warblers – Warbler_genome
