## Supplemental Figure 5 for "The power to resolve relationships: identifying incongruence and precision of reduced representation and genome-wide data in phylogenomics and population genomics"

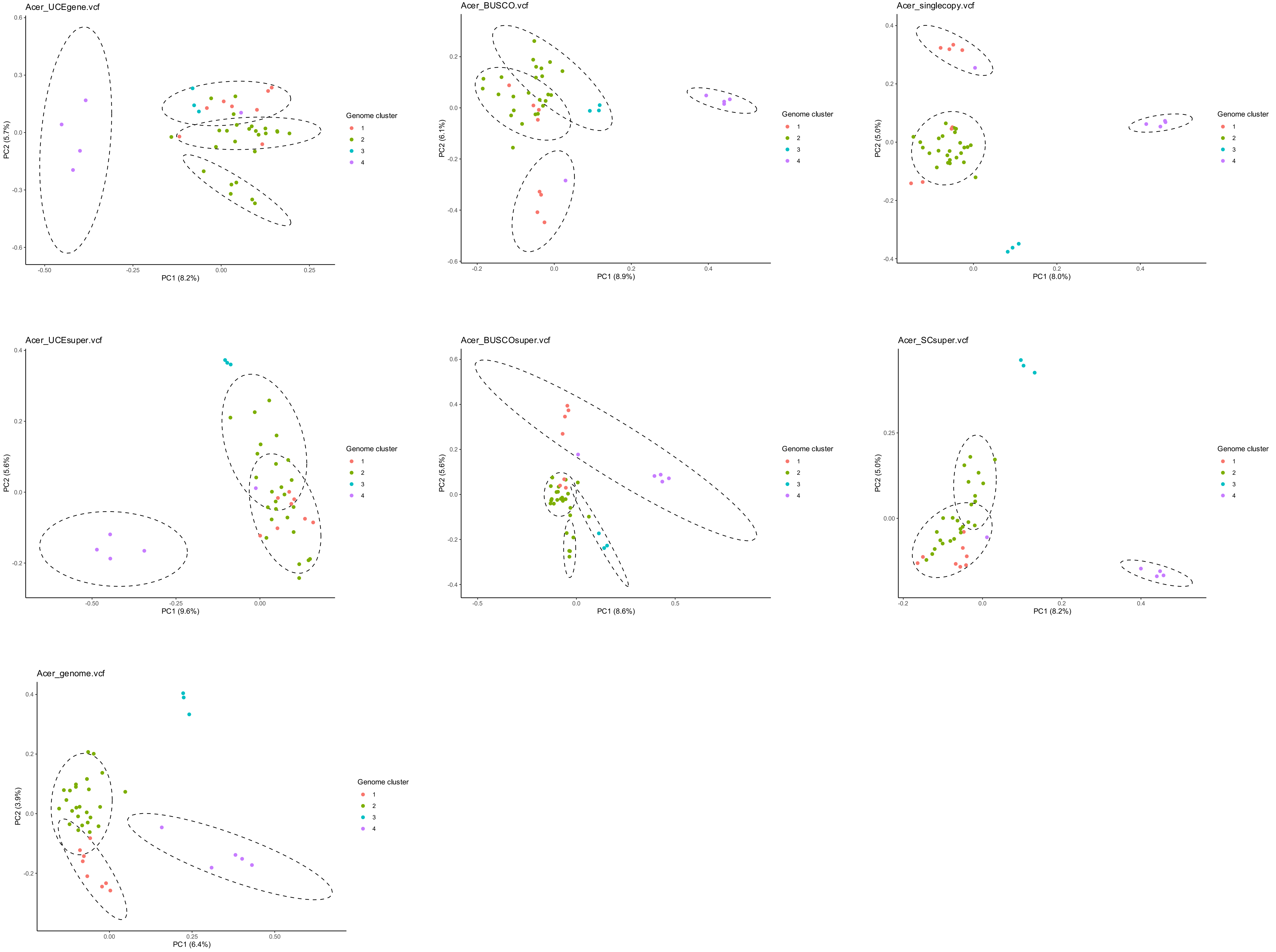
Acer_UCEgene.vcf
PC1 (8.2%)
PC2 (5.7%)
Genome cluster
1
2
3
4
Acer_BUSCO.vcf
PC1 (8.9%)
PC2 (6.1%)
Genome cluster
1
2
3
4
Acer_singlecopy.vcf
PC1 (8.0%)
PC2 (5.0%)
Genome cluster
1
2
3
4
Acer_UCEsuper.vcf
PC1 (9.6%)
PC2 (5.6%)
Genome cluster
1
2
3
4
Acer_BUSCOsuper.vcf
PC1 (8.6%)
PC2 (5.6%)
Genome cluster
1
2
3
4
Acer_SCsuper.vcf
PC1 (8.2%)
PC2 (5.0%)
Genome cluster
1
2
3
4
Acer_genome.vcf
PC1 (6.4%)
PC2 (3.9%)
Genome cluster
1
2
3
4

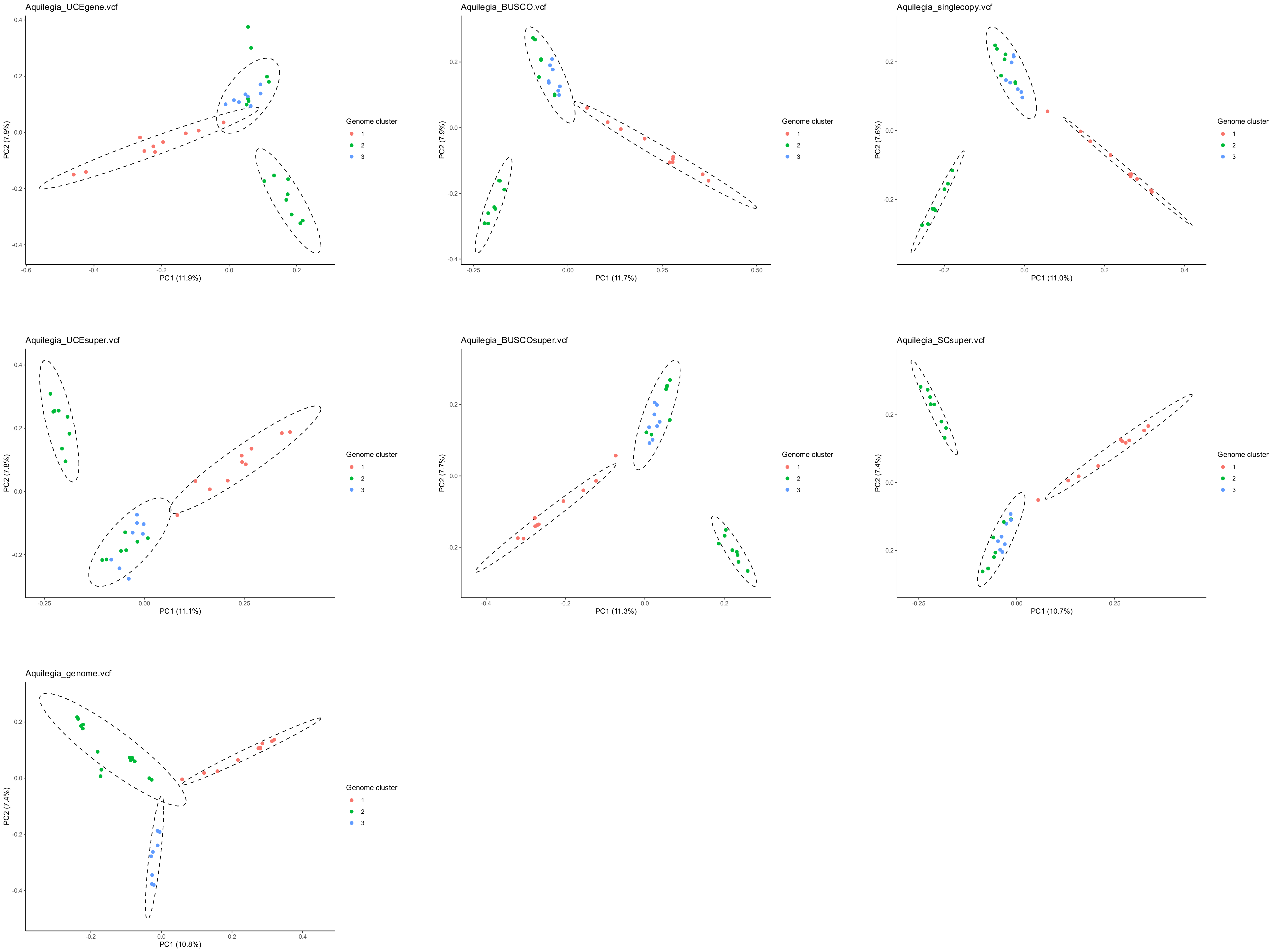

Aquilegia_UCEgene.vcf
PC1 (11.9%)
PC2 (7.9%)
Aquilegia_BUSCO.vcf
PC1 (11.7%)
PC2 (7.9%)
Aquilegia_singlecopy.vcf
PC1 (11.0%)
PC2 (7.6%)
Aquilegia_UCEsuper.vcf
PC1 (11.1%)
PC2 (7.8%)
Aquilegia_BUSCOsuper.vcf
PC1 (11.3%)
PC2 (7.7%)
Aquilegia_SCsuper.vcf
PC1 (10.7%)
PC2 (7.4%)
Aquilegia_genome.vcf
PC1 (10.8%)
PC2 (7.4%)
Genome cluster
1
2
3

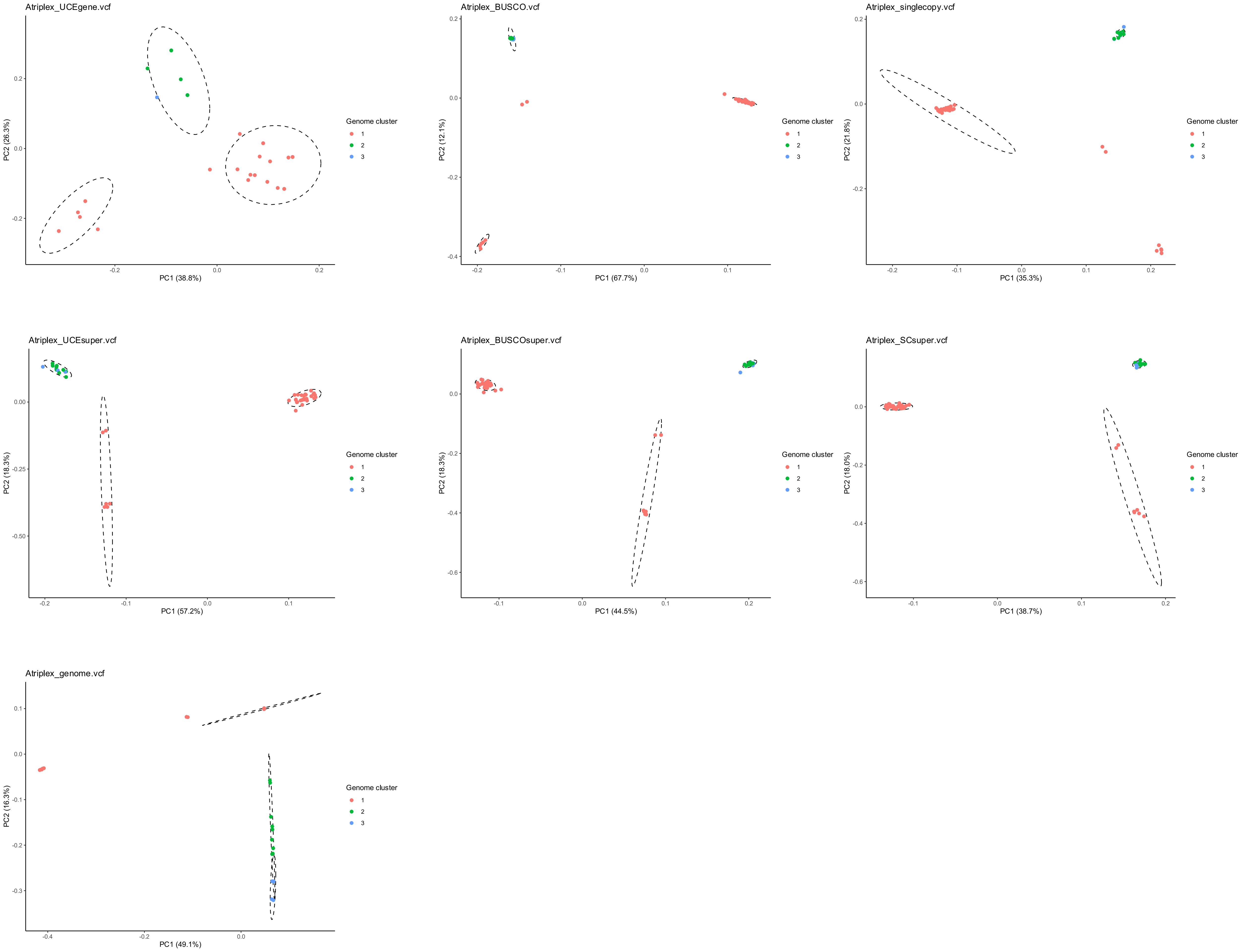

Atriplex_UCEgene.vcf
PC1 (38.8%)
PC2 (26.3%)
Atriplex_BUSCO.vcf
PC1 (67.7%)
PC2 (12.1%)
Atriplex_singlecopy.vcf
PC1 (35.3%)
PC2 (21.8%)
Atriplex_UCEsuper.vcf
PC1 (57.2%)
PC2 (18.3%)
Atriplex_BUSCOsuper.vcf
PC1 (44.5%)
PC2 (18.3%)
Atriplex_SCsuper.vcf
PC1 (38.7%)
PC2 (18.0%)
Atriplex_genome.vcf
PC1 (49.1%)
PC2 (16.3%)
Genome cluster
1
2
3

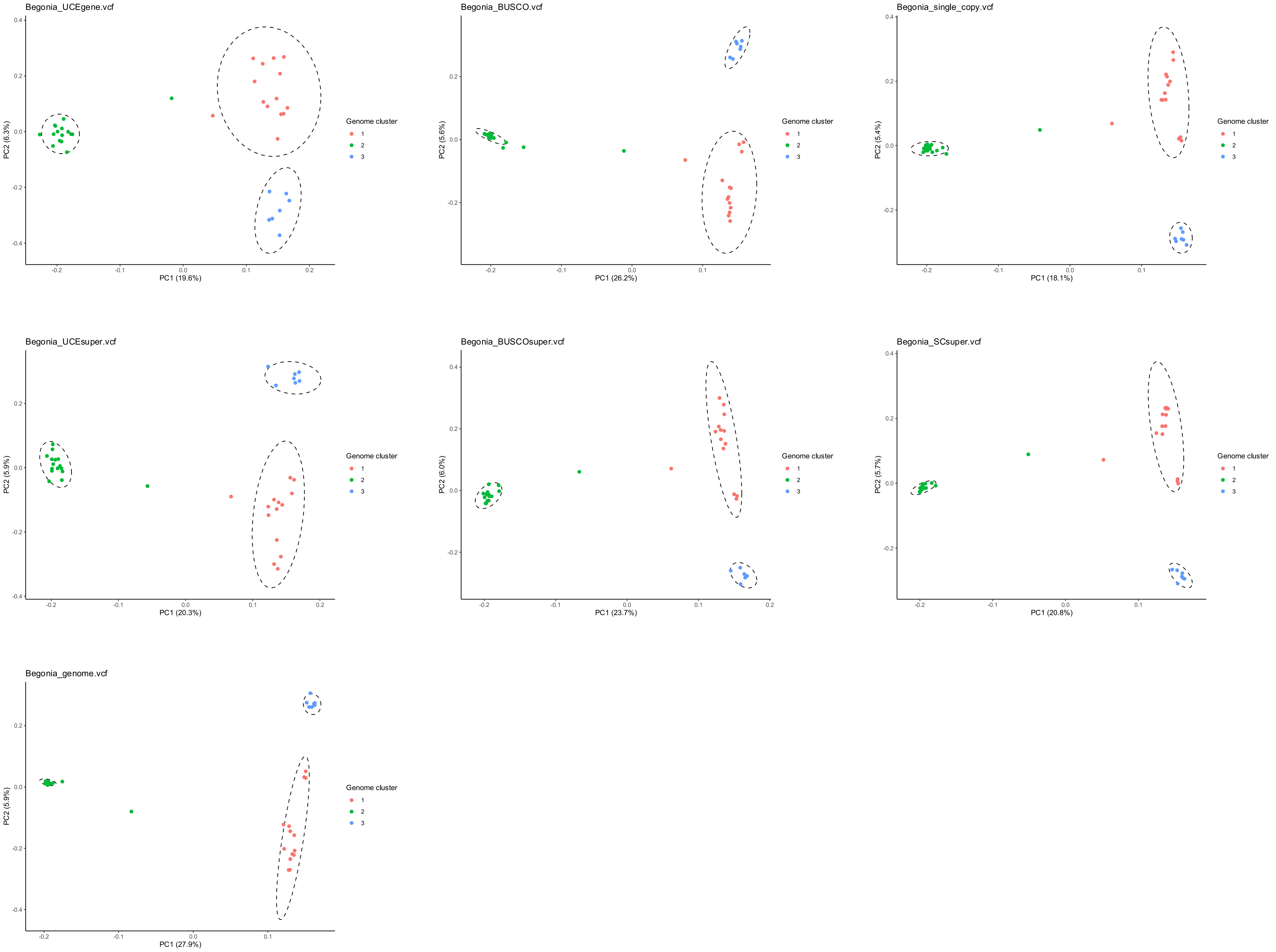

Begonia_UCEgene.vcf
PC1 (19.6%)
PC2 (6.3%)
Begonia_BUSCO.vcf
PC1 (26.2%)
PC2 (5.6%)
Begonia_single_copy.vcf
PC1 (18.1%)
PC2 (5.4%)
Begonia_UCEsuper.vcf
PC1 (20.3%)
PC2 (5.9%)
Begonia_BUSCOsuper.vcf
PC1 (23.7%)
PC2 (6.0%)
Begonia_SCsuper.vcf
PC1 (20.8%)
PC2 (5.7%)
Begonia_genome.vcf
PC1 (27.9%)
PC2 (5.9%)
Genome cluster
1
2
3

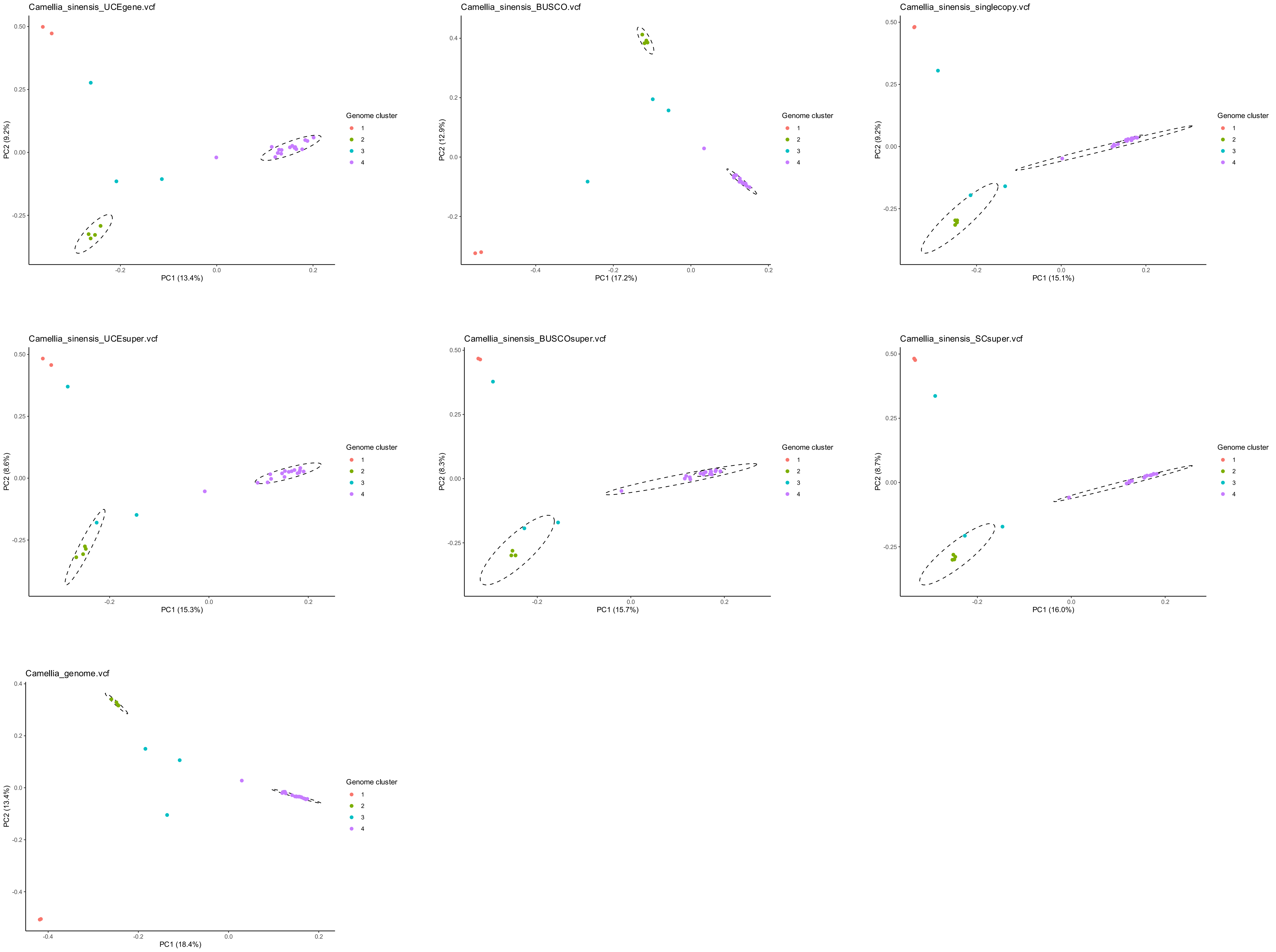

Camellia_sinensis_UCEgene.vcf
PC1 (13.4%)
PC2 (9.2%)
Camellia_sinensis_BUSCO.vcf
PC1 (17.2%)
PC2 (12.9%)
Camellia_sinensis_singlecopy.vcf
PC1 (15.1%)
PC2 (9.2%)
Camellia_sinensis_UCEsuper.vcf
PC1 (15.3%)
PC2 (8.6%)
Camellia_sinensis_BUSCOsuper.vcf
PC1 (15.7%)
PC2 (8.3%)
Camellia_sinensis_SCsuper.vcf
PC1 (16.0%)
PC2 (8.7%)
Camellia_genome.vcf
PC1 (18.4%)
PC2 (13.4%)
Genome cluster
1
2
3
4

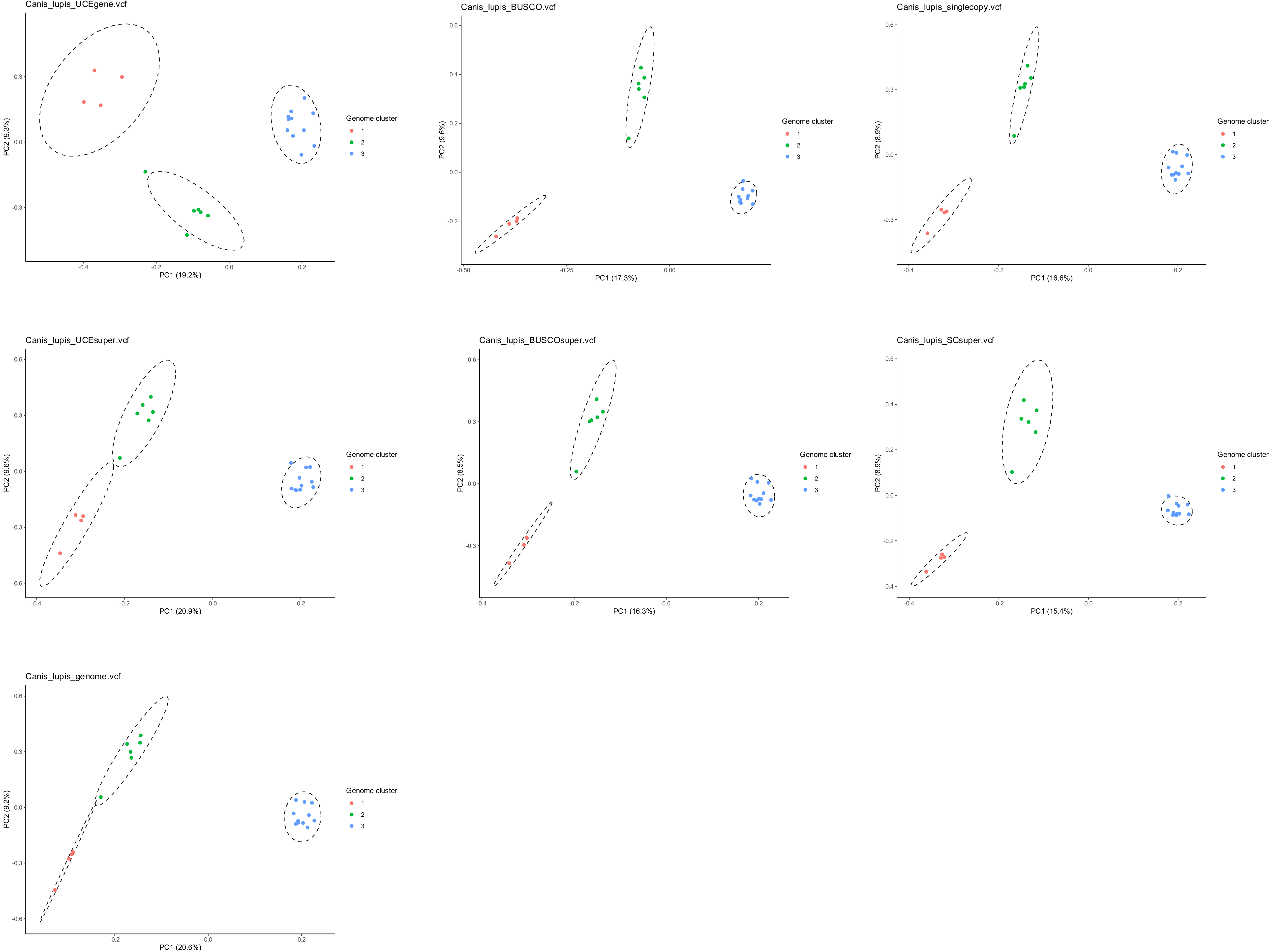
Canis_lupis_UCEgene.vcf
PC1 (19.2%)
PC2 (9.3%)
Genome cluster
1
2
3
Canis_lupis_BUSCO.vcf
PC1 (17.3%)
PC2 (9.6%)
Genome cluster
1
2
3
Canis_lupis_singlecopy.vcf
PC1 (16.6%)
PC2 (8.9%)
Genome cluster
1
2
3
Canis_lupis_UCEsuper.vcf
PC1 (20.9%)
PC2 (9.6%)
Genome cluster
1
2
3
Canis_lupis_BUSCOsuper.vcf
PC1 (16.3%)
PC2 (8.5%)
Genome cluster
1
2
3
Canis_lupis_SCsuper.vcf
PC1 (15.4%)
PC2 (8.9%)
Genome cluster
1
2
3
Canis_lupis_genome.vcf
PC1 (20.6%)
PC2 (9.2%)
Genome cluster
1
2
3

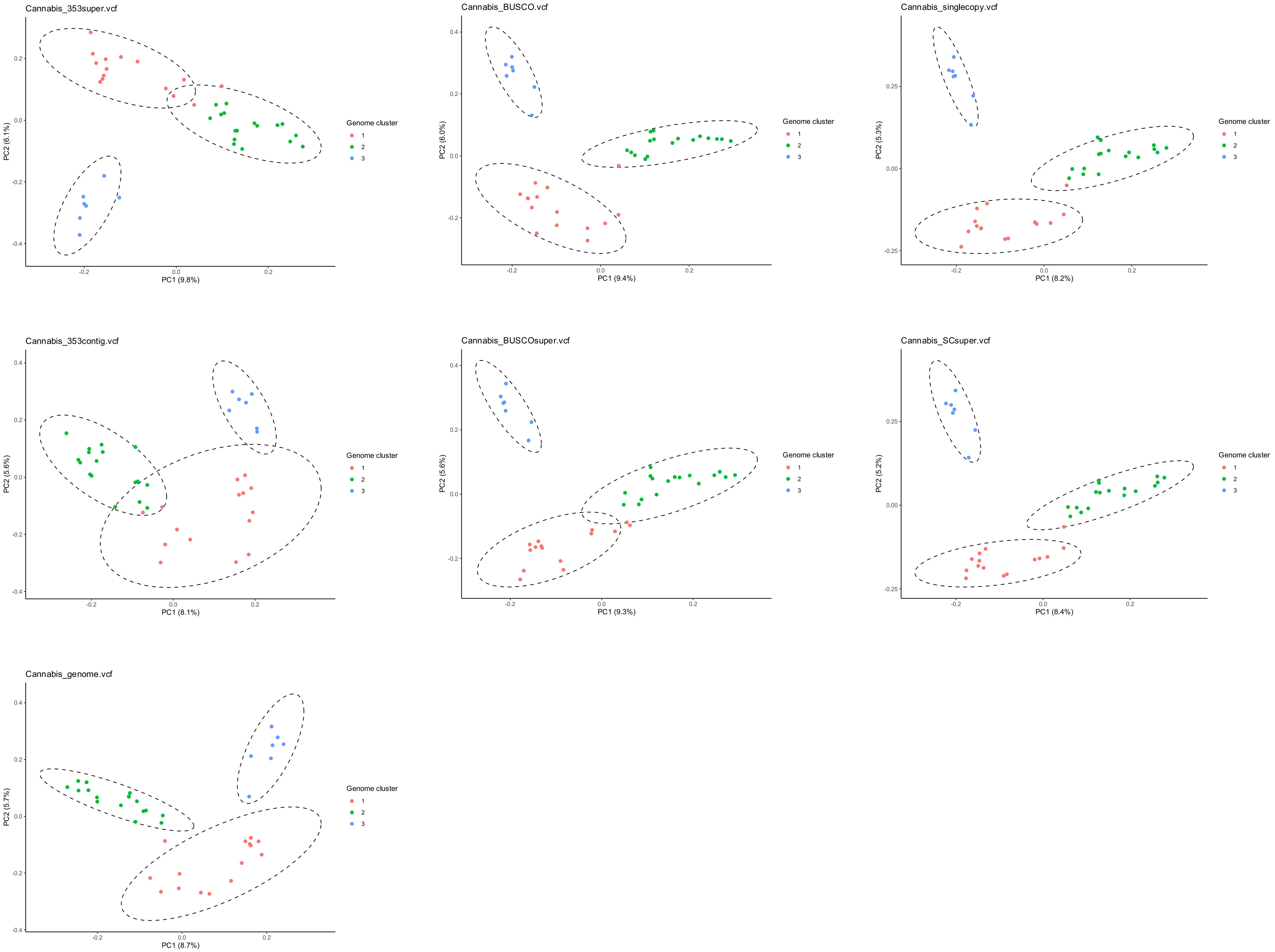
Cannabis_353super.vcf
PC2 (6.1%)
PC1 (9.8%)
Genome cluster
1
2
3
Cannabis_BUSCO.vcf
PC2 (6.0%)
PC1 (9.4%)
Genome cluster
1
2
3
Cannabis_singlecopy.vcf
PC2 (5.3%)
PC1 (8.2%)
Genome cluster
1
2
3
Cannabis_353contig.vcf
PC2 (5.6%)
PC1 (8.1%)
Genome cluster
1
2
3
Cannabis_BUSCOsuper.vcf
PC2 (5.6%)
PC1 (9.3%)
Genome cluster
1
2
3
Cannabis_SCsuper.vcf
PC2 (5.2%)
PC1 (8.4%)
Genome cluster
1
2
3
Cannabis_genome.vcf
PC2 (5.7%)
PC1 (8.7%)
Genome cluster
1
2
3

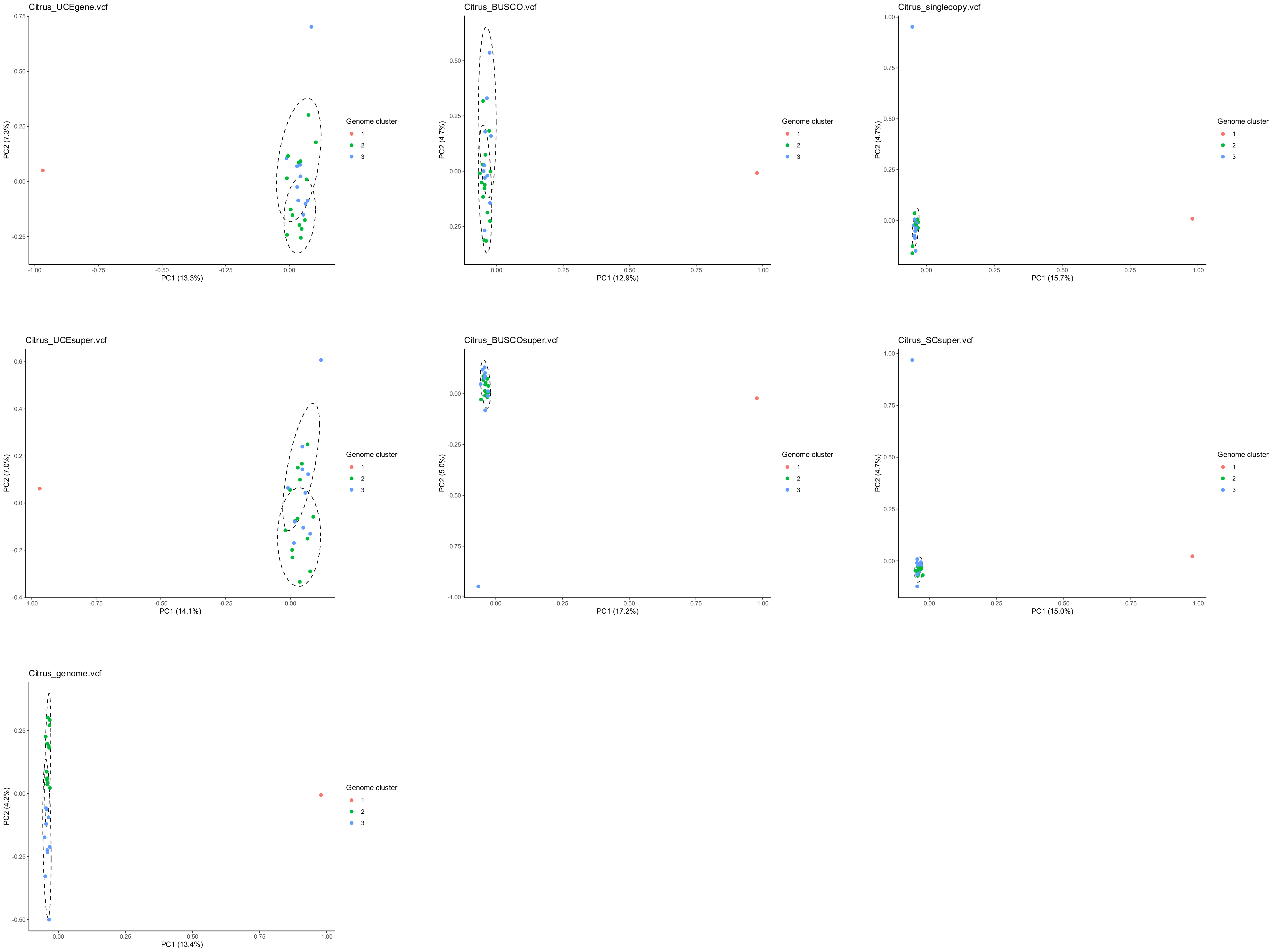
Citrus_UCEgene.vcf
PC1 (13.3%)
PC2 (7.3%)
Genome cluster
1
2
3
Citrus_BUSCO.vcf
PC1 (12.9%)
PC2 (4.7%)
Genome cluster
1
2
3
Citrus_singlecopy.vcf
PC1 (15.7%)
PC2 (4.7%)
Genome cluster
1
2
3
Citrus_UCEsuper.vcf
PC1 (14.1%)
PC2 (7.0%)
Genome cluster
1
2
3
Citrus_BUSCOsuper.vcf
PC1 (17.2%)
PC2 (5.0%)
Genome cluster
1
2
3
Citrus_SCsuper.vcf
PC1 (15.0%)
PC2 (4.7%)
Genome cluster
1
2
3
Citrus_genome.vcf
PC1 (13.4%)
PC2 (4.2%)
Genome cluster
1
2
3

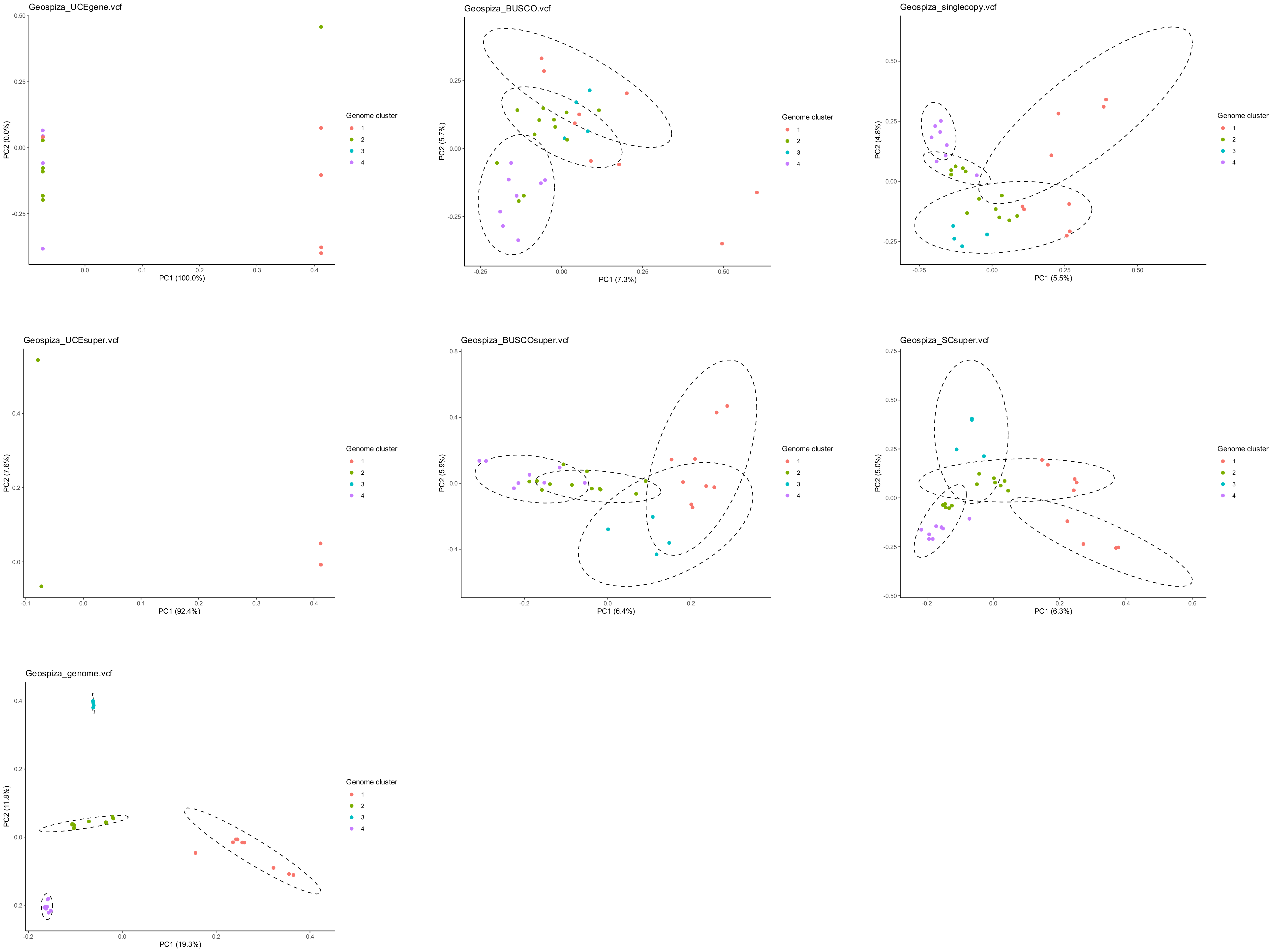
Geospiza_UCEgene.vcf
PC1 (100.0%)
PC2 (0.0%)
Geospiza_BUSCO.vcf
PC1 (7.3%)
PC2 (5.7%)
Geospiza_singlecopy.vcf
PC1 (5.5%)
PC2 (4.8%)
Geospiza_UCEsuper.vcf
PC1 (92.4%)
PC2 (7.6%)
Geospiza_BUSCOsuper.vcf
PC1 (6.4%)
PC2 (5.9%)
Geospiza_SCsuper.vcf
PC1 (6.3%)
PC2 (5.0%)
Geospiza_genome.vcf
PC1 (19.3%)
PC2 (11.8%)
Genome cluster
1
2
3
4

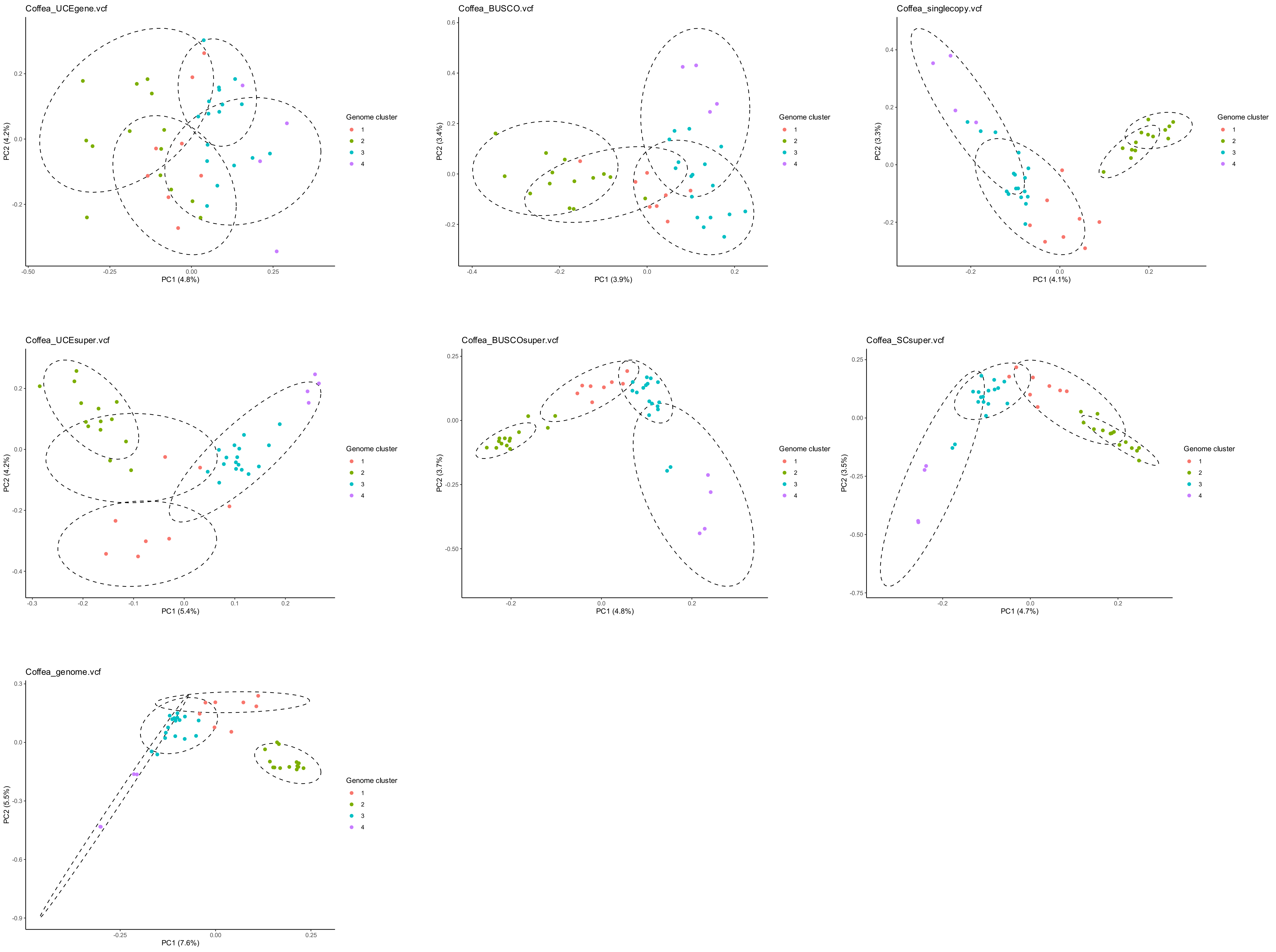
Coffea_UCEgene.vcf
PC1 (4.8%)
PC2 (4.2%)
Coffea_BUSCO.vcf
PC1 (3.9%)
PC2 (3.4%)
Coffea_singlecopy.vcf
PC1 (4.1%)
PC2 (3.3%)
Coffea_UCEsuper.vcf
PC1 (5.4%)
PC2 (4.2%)
Coffea_BUSCOsuper.vcf
PC1 (4.8%)
PC2 (3.7%)
Coffea_SCsuper.vcf
PC1 (4.7%)
PC2 (3.5%)
Coffea_genome.vcf
PC1 (7.6%)
PC2 (5.5%)
Genome cluster
1
2
3
4

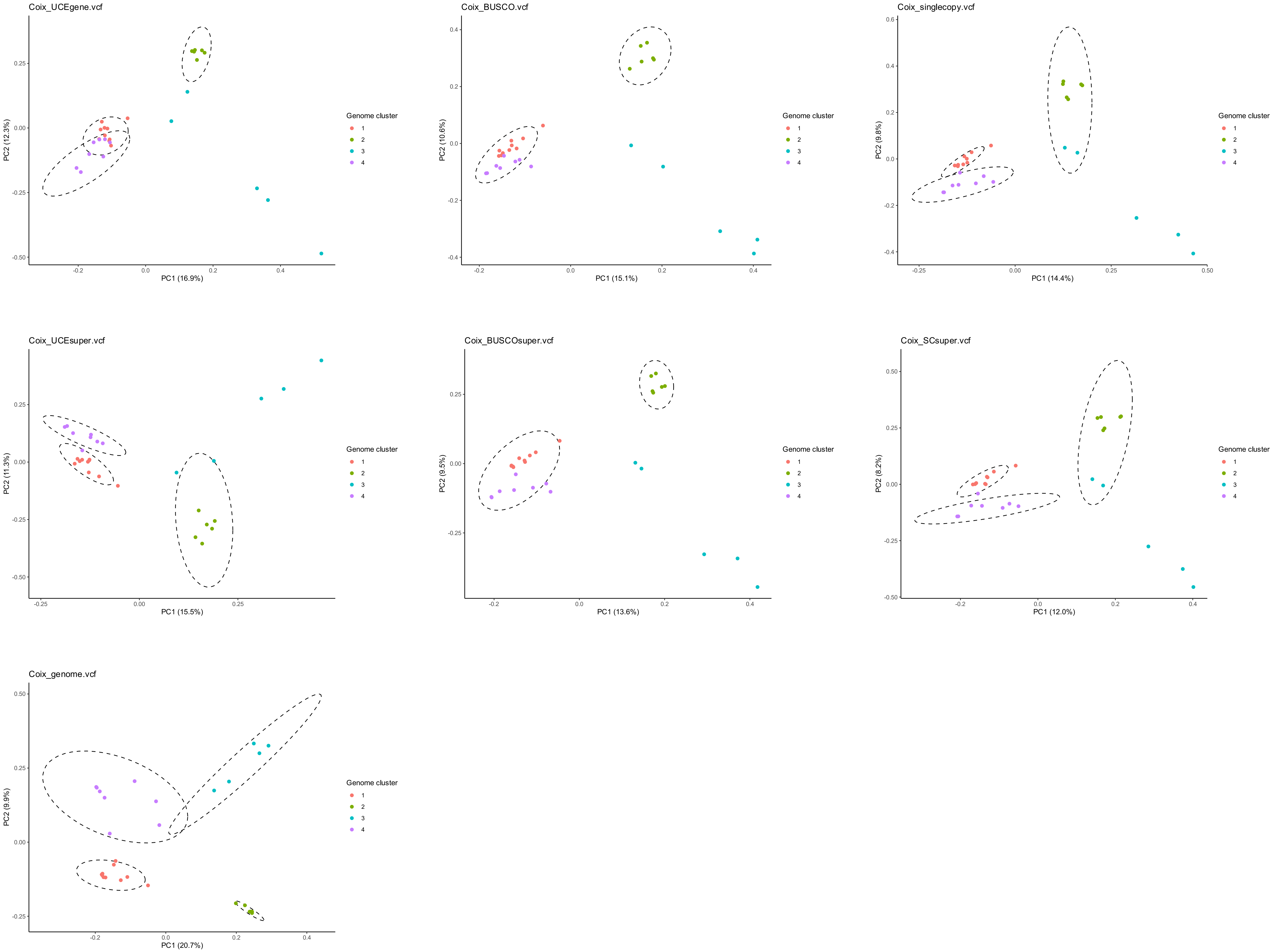

Coix_UCEgene.vcf
PC1 (16.9%)
PC2 (12.3%)
Genome cluster
1
2
3
4
Coix_BUSCO.vcf
PC1 (15.1%)
PC2 (10.6%)
Genome cluster
1
2
3
4
Coix_singlecopy.vcf
PC1 (14.4%)
PC2 (9.8%)
Genome cluster
1
2
3
4
Coix_UCEsuper.vcf
PC1 (15.5%)
PC2 (11.3%)
Genome cluster
1
2
3
4
Coix_BUSCOsuper.vcf
PC1 (13.6%)
PC2 (9.5%)
Genome cluster
1
2
3
4
Coix_SCsuper.vcf
PC1 (12.0%)
PC2 (8.2%)
Genome cluster
1
2
3
4
Coix_genome.vcf
PC1 (20.7%)
PC2 (9.9%)
Genome cluster
1
2
3
4

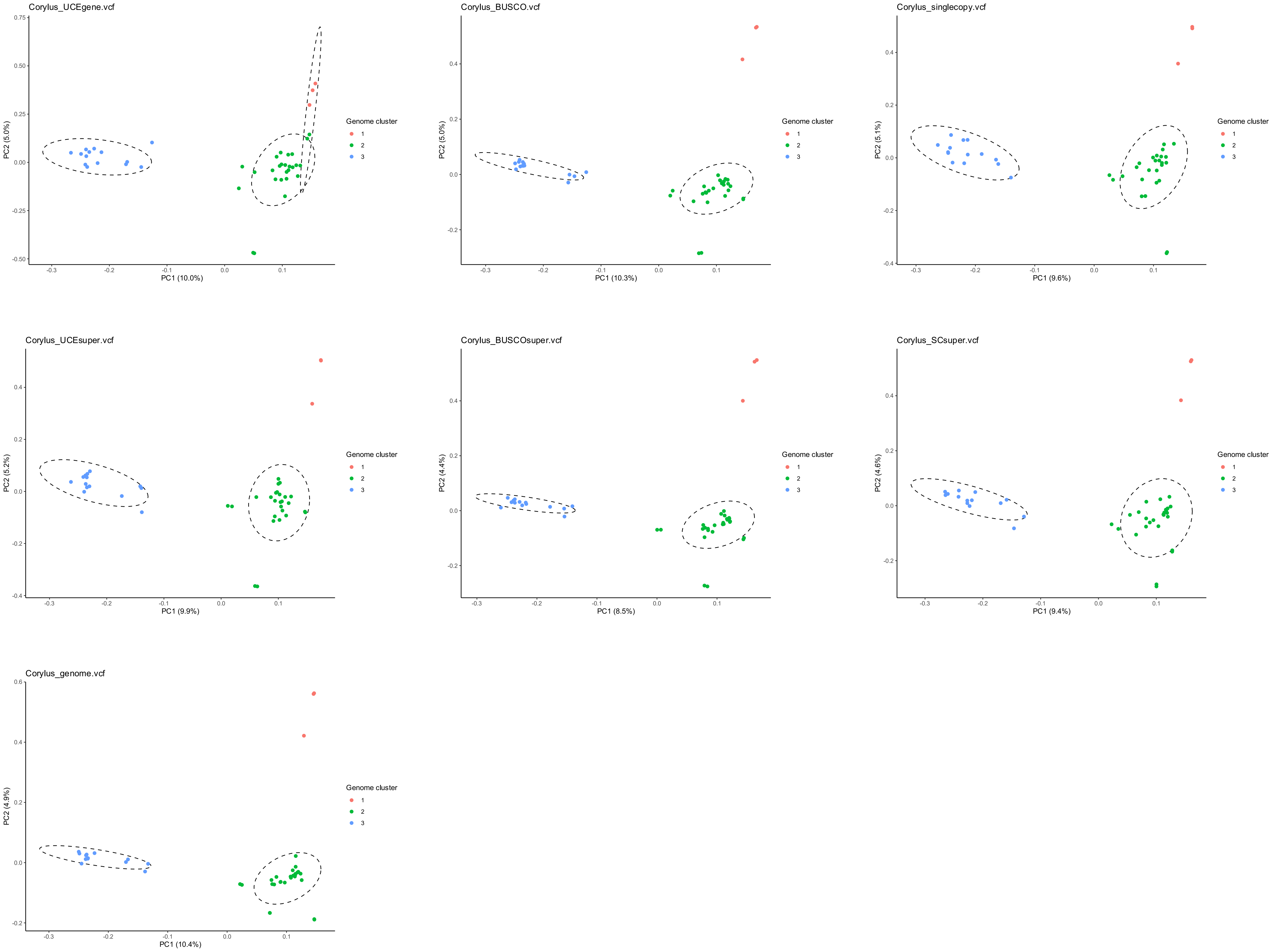
Corylus_UCEgene.vcf
PC1 (10.0%)
PC2 (5.0%)
Genome cluster
1
2
3
Corylus_BUSCO.vcf
PC1 (10.3%)
PC2 (5.0%)
Genome cluster
1
2
3
Corylus_singlecopy.vcf
PC1 (9.6%)
PC2 (5.1%)
Genome cluster
1
2
3
Corylus_UCEsuper.vcf
PC1 (9.9%)
PC2 (5.2%)
Genome cluster
1
2
3
Corylus_BUSCOsuper.vcf
PC1 (8.5%)
PC2 (4.4%)
Genome cluster
1
2
3
Corylus_SCsuper.vcf
PC1 (9.4%)
PC2 (4.6%)
Genome cluster
1
2
3
Corylus_genome.vcf
PC1 (10.4%)
PC2 (4.9%)
Genome cluster
1
2
3

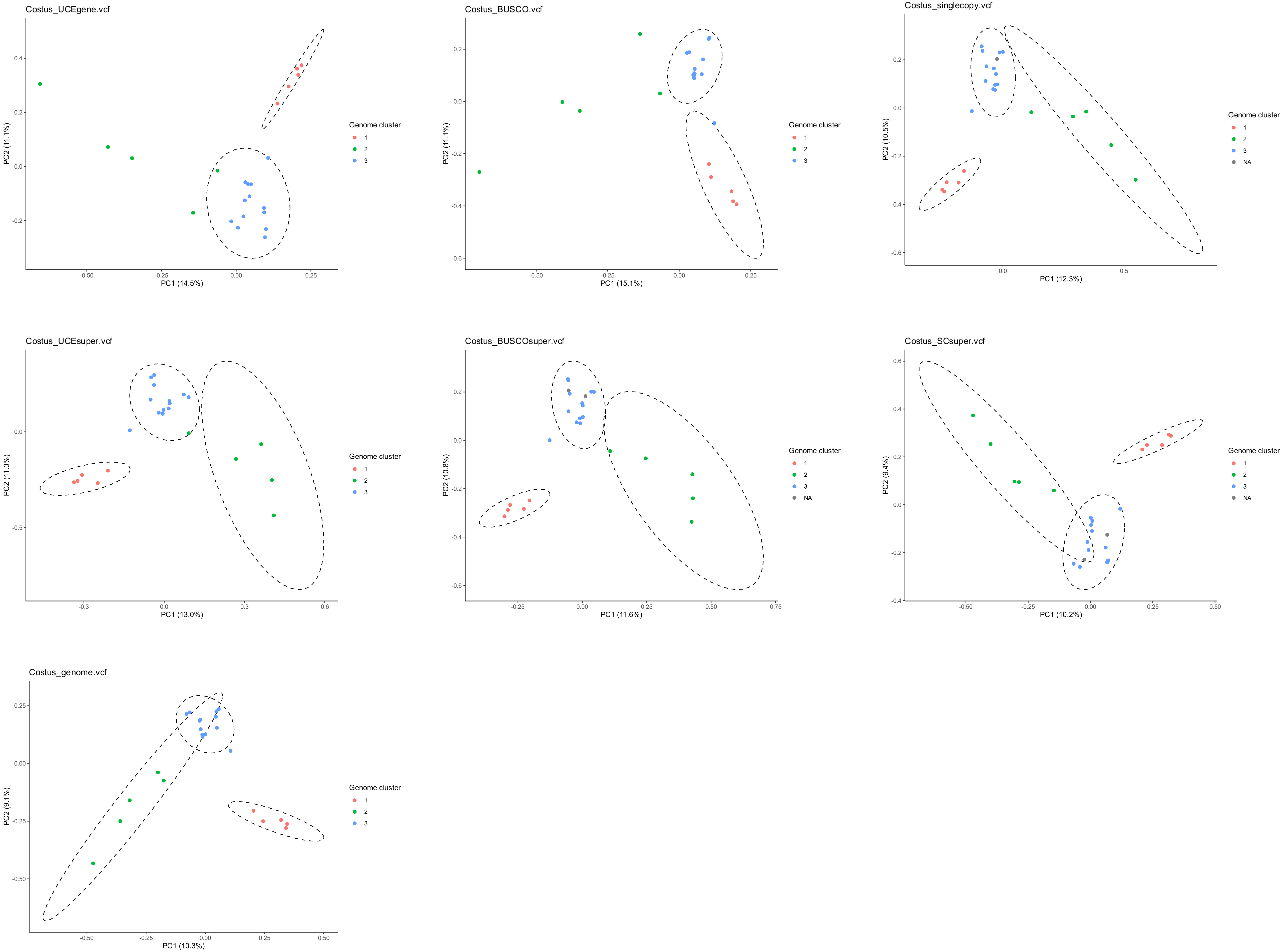

Costus_UCEgene.vcf
PC1 (14.5%)
PC2 (11.1%)
Genome cluster
1
2
3
Costus_BUSCO.vcf
PC1 (15.1%)
PC2 (11.1%)
Genome cluster
1
2
3
Costus_singlecopy.vcf
PC1 (12.3%)
PC2 (10.5%)
Genome cluster
1
2
3
NA
Costus_UCEsuper.vcf
PC1 (13.0%)
PC2 (11.0%)
Genome cluster
1
2
3
Costus_BUSCOsuper.vcf
PC1 (11.6%)
PC2 (10.8%)
Genome cluster
1
2
3
NA
Costus_SCsuper.vcf
PC1 (10.2%)
PC2 (9.4%)
Genome cluster
1
2
3
NA
Costus_genome.vcf
PC1 (10.3%)
PC2 (9.1%)
Genome cluster
1
2
3

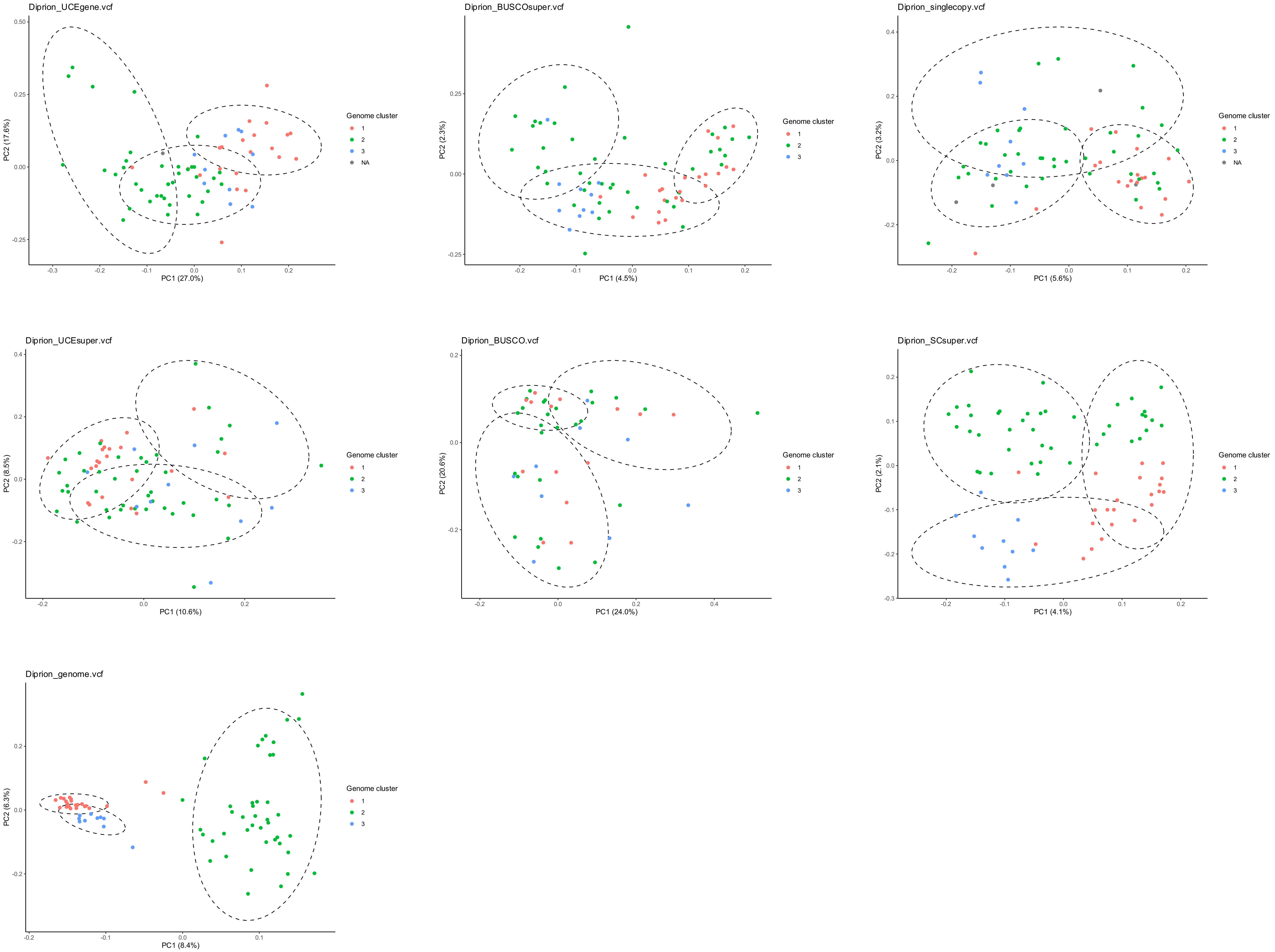

Diprion_UCEgene.vcf
PC1 (27.0%)
PC2 (17.6%)
Genome cluster
1
2
3
NA
Diprion_BUSCOsuper.vcf
PC1 (4.5%)
PC2 (2.3%)
Genome cluster
1
2
3
Diprion_singlecopy.vcf
PC1 (5.6%)
PC2 (3.2%)
Genome cluster
1
2
3
NA
Diprion_UCEsuper.vcf
PC1 (10.6%)
PC2 (8.5%)
Genome cluster
1
2
3
Diprion_BUSCO.vcf
PC1 (24.0%)
PC2 (20.6%)
Genome cluster
1
2
3
Diprion_SCsuper.vcf
PC1 (4.1%)
PC2 (2.1%)
Genome cluster
1
2
3
Diprion_genome.vcf
PC1 (8.4%)
PC2 (6.3%)
Genome cluster
1
2
3

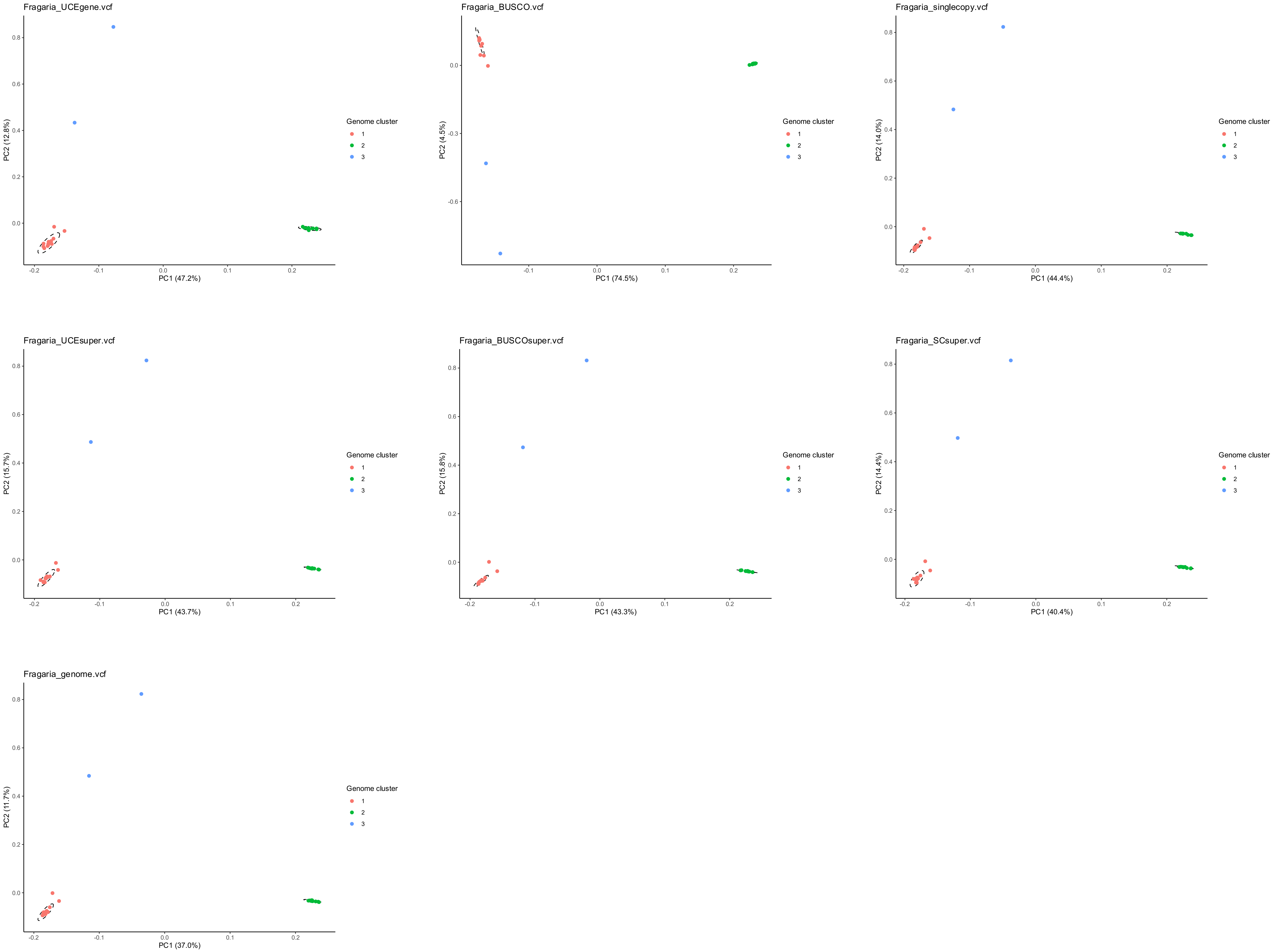
Fragaria_UCEgene.vcf
PC2 (12.8%)
PC1 (47.2%)
Genome cluster
1
2
3
Fragaria_BUSCO.vcf
PC2 (4.5%)
PC1 (74.5%)
Genome cluster
1
2
3
Fragaria_singlecopy.vcf
PC2 (14.0%)
PC1 (44.4%)
Genome cluster
1
2
3
Fragaria_UCEsuper.vcf
PC2 (15.7%)
PC1 (43.7%)
Genome cluster
1
2
3
Fragaria_BUSCOsuper.vcf
PC2 (15.8%)
PC1 (43.3%)
Genome cluster
1
2
3
Fragaria_SCsuper.vcf
PC2 (14.4%)
PC1 (40.4%)
Genome cluster
1
2
3
Fragaria_genome.vcf
PC2 (11.7%)
PC1 (37.0%)
Genome cluster
1
2
3

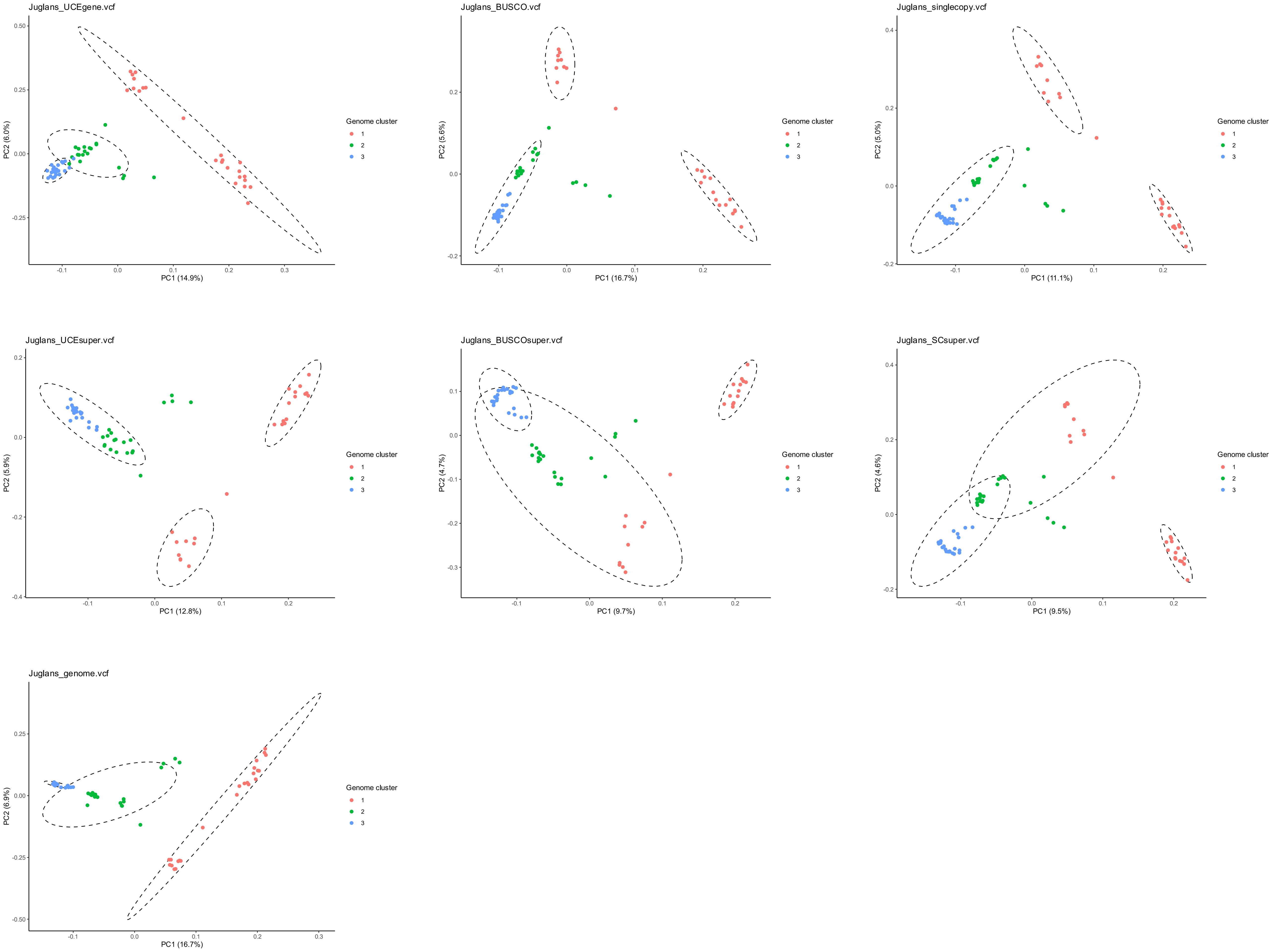
Juglans_UCEgene.vcf
PC1 (14.9%)
PC2 (6.0%)
Genome cluster
1
2
3
Juglans_BUSCO.vcf
PC1 (16.7%)
PC2 (5.6%)
Genome cluster
1
2
3
Juglans_singlecopy.vcf
PC1 (11.1%)
PC2 (5.0%)
Genome cluster
1
2
3
Juglans_UCEsuper.vcf
PC1 (12.8%)
PC2 (5.9%)
Genome cluster
1
2
3
Juglans_BUSCOsuper.vcf
PC1 (9.7%)
PC2 (4.7%)
Genome cluster
1
2
3
Juglans_SCsuper.vcf
PC1 (9.5%)
PC2 (4.6%)
Genome cluster
1
2
3
Juglans_genome.vcf
PC1 (16.7%)
PC2 (6.9%)
Genome cluster
1
2
3

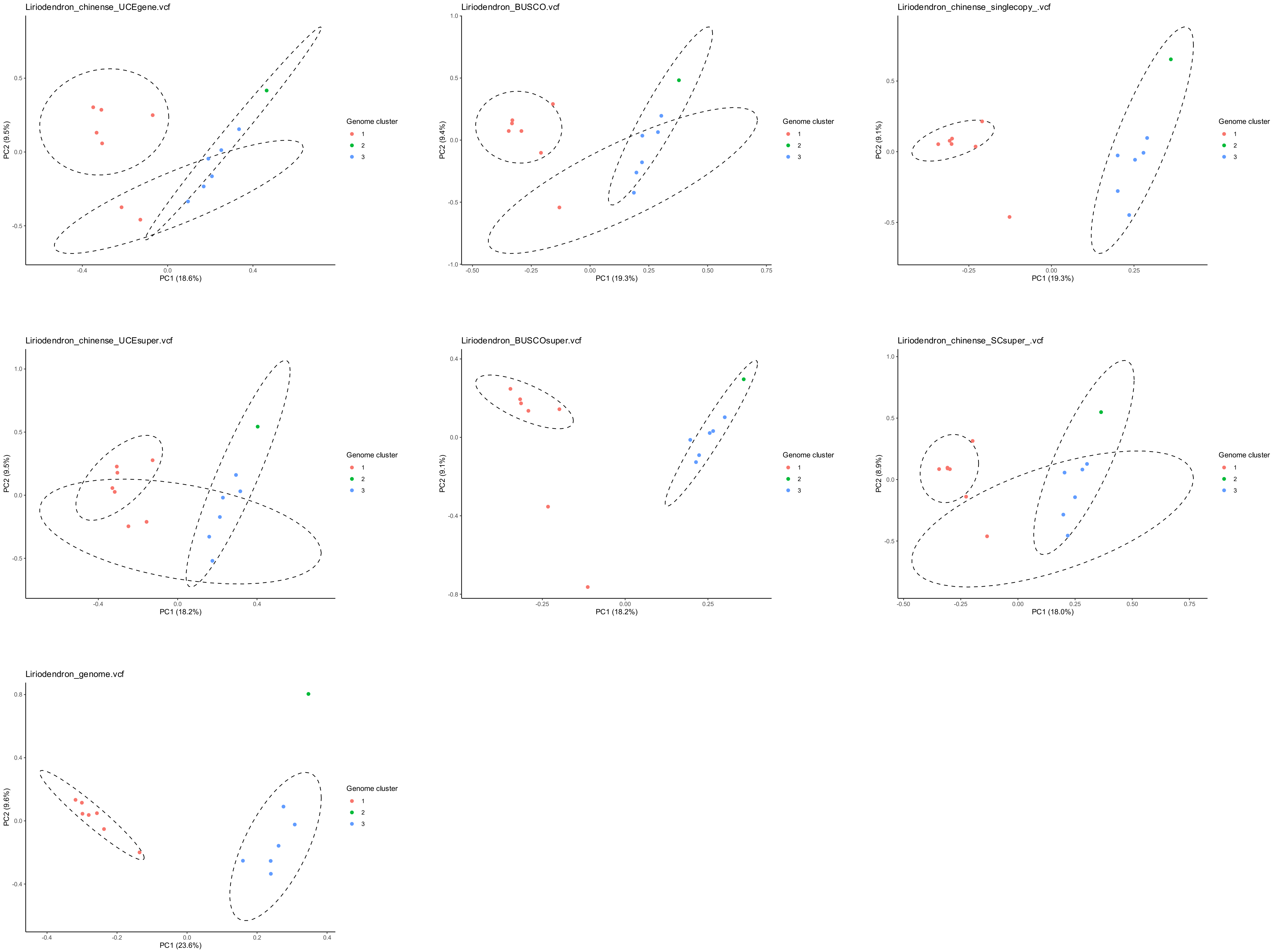

Liriodendron_chinense_UCEgene.vcf
PC1 (18.6%)
PC2 (9.5%)
Genome cluster
1
2
3
Liriodendron_BUSCO.vcf
PC1 (19.3%)
PC2 (9.4%)
Genome cluster
1
2
3
Liriodendron_chinense_singlecopy_.vcf
PC1 (19.3%)
PC2 (9.1%)
Genome cluster
1
2
3
Liriodendron_chinense_UCEsuper.vcf
PC1 (18.2%)
PC2 (9.5%)
Genome cluster
1
2
3
Liriodendron_BUSCOsuper.vcf
PC1 (18.2%)
PC2 (9.1%)
Genome cluster
1
2
3
Liriodendron_chinense_SCsuper_.vcf
PC1 (18.0%)
PC2 (8.9%)
Genome cluster
1
2
3
Liriodendron_genome.vcf
PC1 (23.6%)
PC2 (9.6%)
Genome cluster
1
2
3

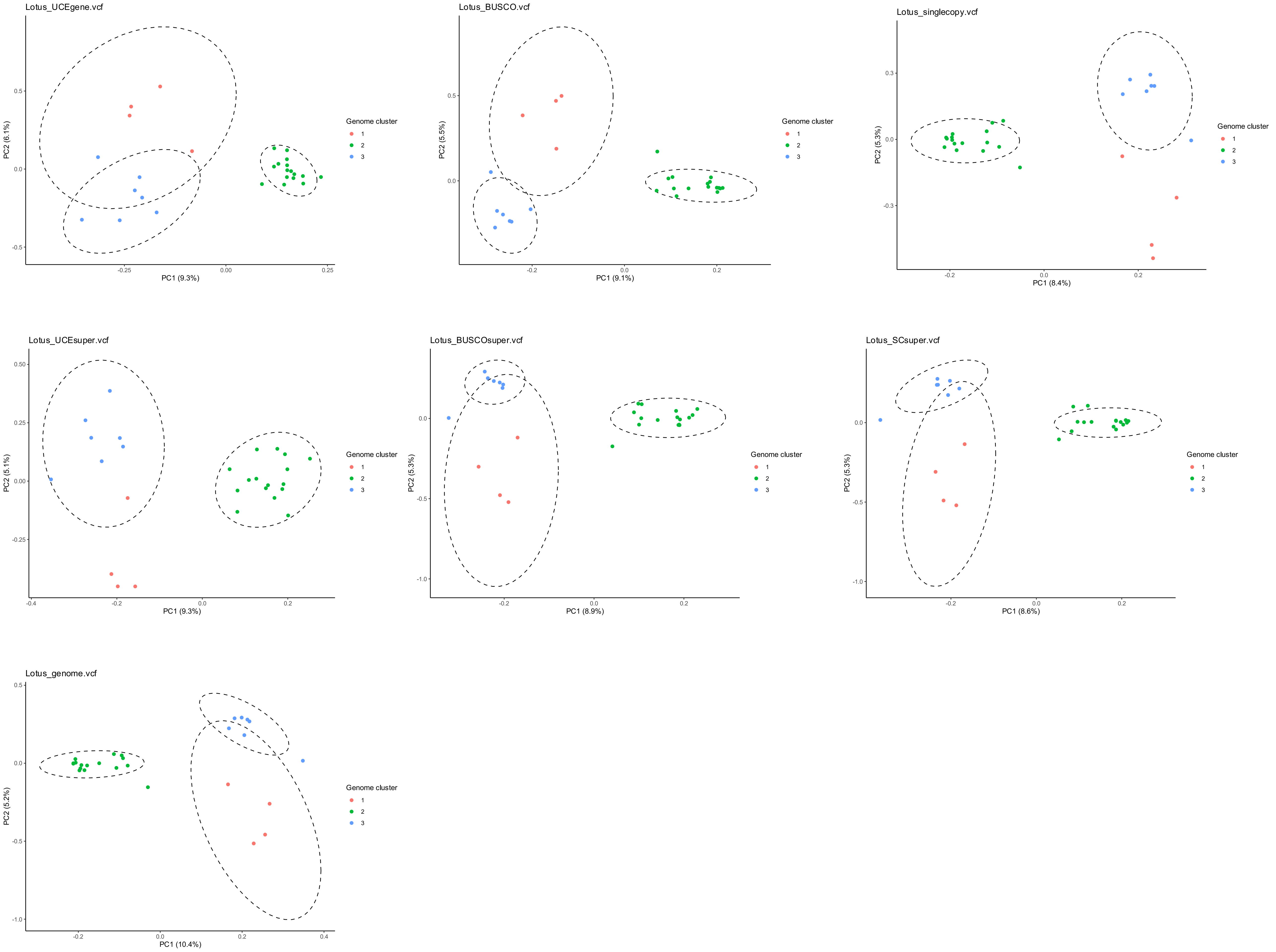

Lotus_UCEgene.vcf
PC1 (9.3%)
PC2 (6.1%)
Lotus_BUSCO.vcf
PC1 (9.1%)
PC2 (5.5%)
Lotus_singlecopy.vcf
PC1 (8.4%)
PC2 (5.3%)
Lotus_UCEsuper.vcf
PC1 (9.3%)
PC2 (5.1%)
Lotus_BUSCOsuper.vcf
PC1 (8.9%)
PC2 (5.3%)
Lotus_SCsuper.vcf
PC1 (8.6%)
PC2 (5.3%)
Lotus_genome.vcf
PC1 (10.4%)
PC2 (5.2%)
Genome cluster
1
2
3

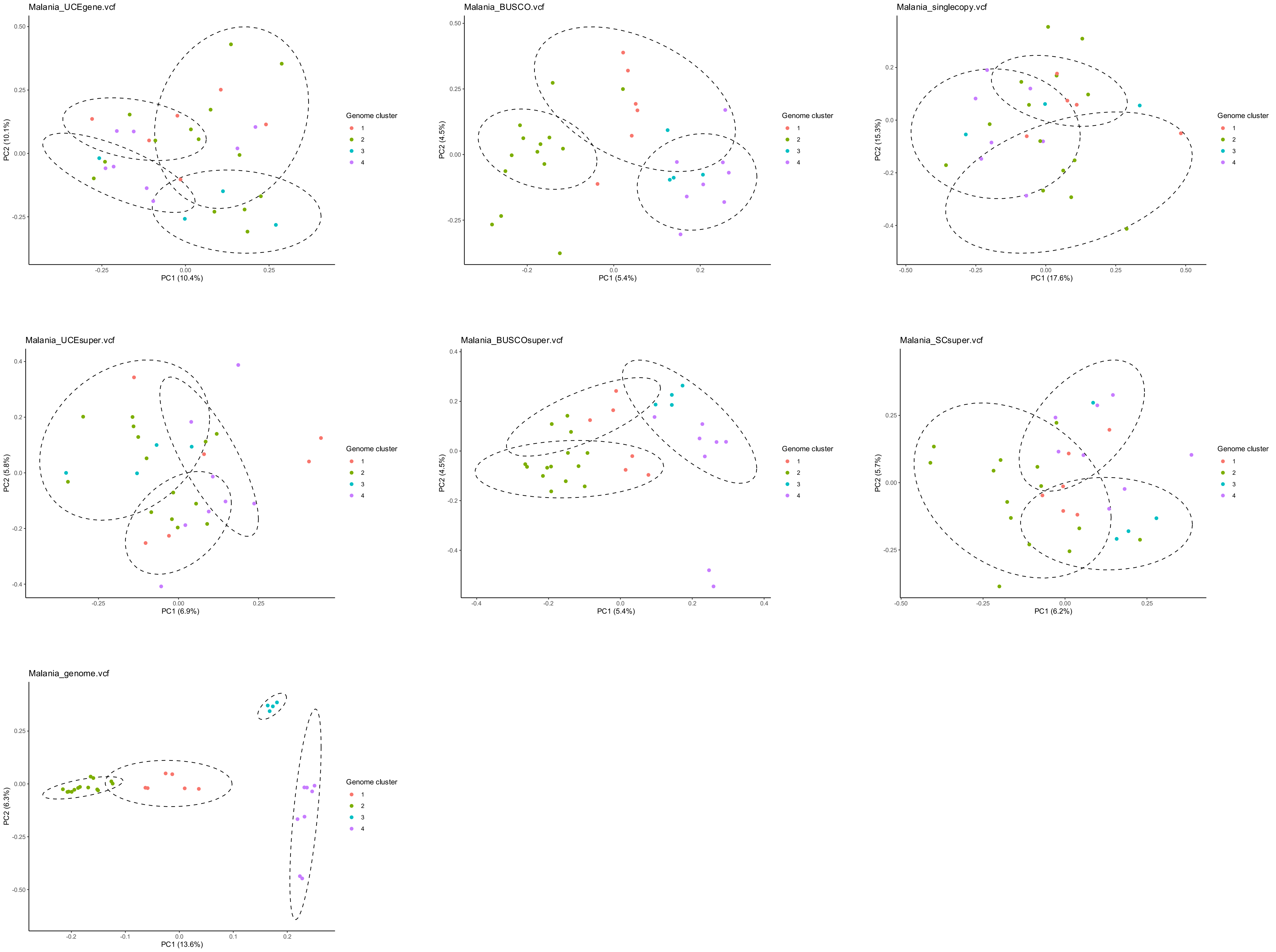

Malania_UCEgene.vcf
PC1 (10.4%)
PC2 (10.1%)
Malania_BUSCO.vcf
PC1 (5.4%)
PC2 (4.5%)
Malania_singlecopy.vcf
PC1 (17.6%)
PC2 (15.3%)
Malania_UCEsuper.vcf
PC1 (6.9%)
PC2 (5.8%)
Malania_BUSCOsuper.vcf
PC1 (5.4%)
PC2 (4.5%)
Malania_SCsuper.vcf
PC1 (6.2%)
PC2 (5.7%)
Malania_genome.vcf
PC1 (13.6%)
PC2 (6.3%)
Genome cluster
1
2
3
4

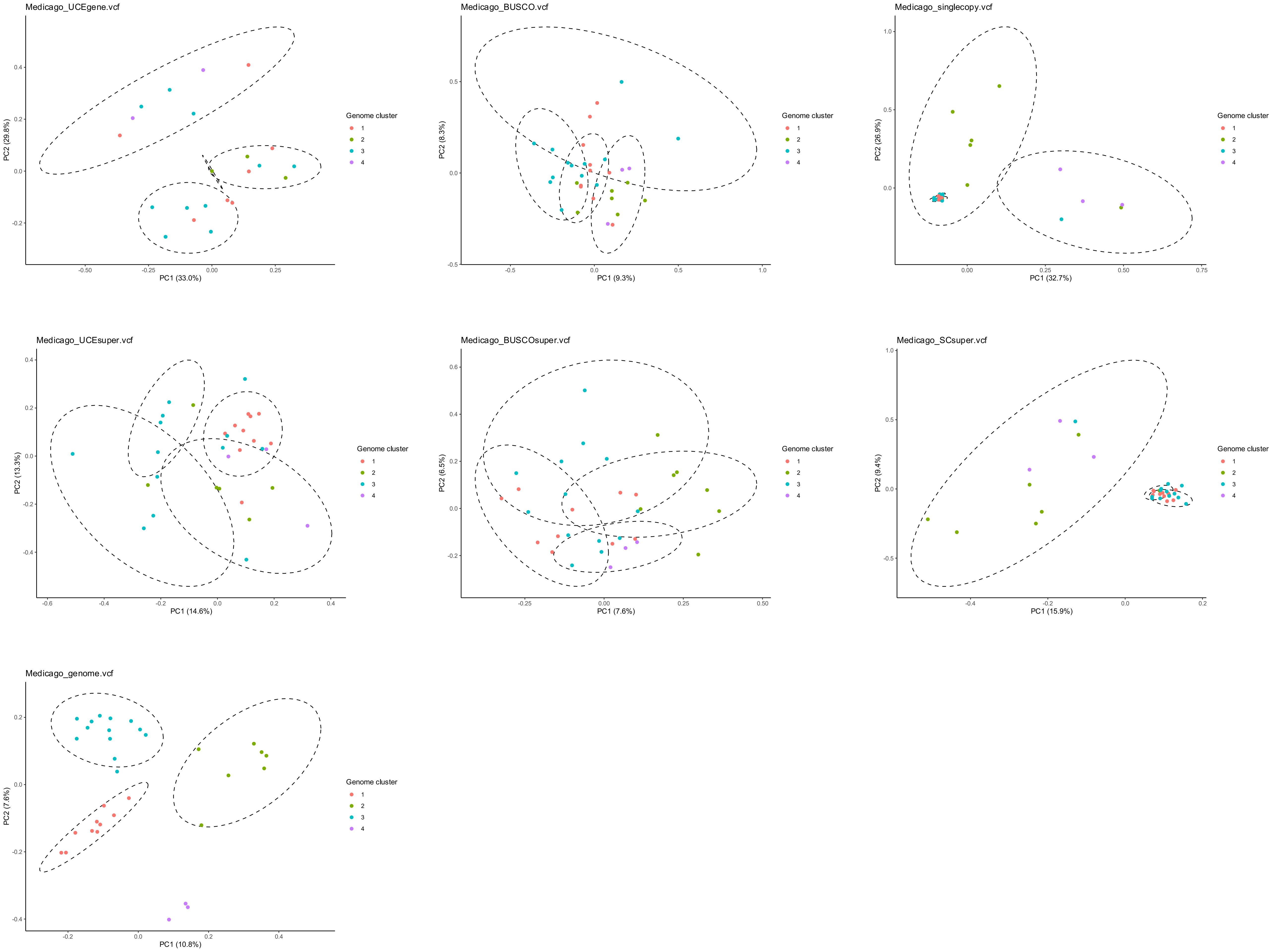

Medicago_UCEgene.vcf
PC1 (33.0%)
PC2 (29.8%)
Medicago_BUSCO.vcf
PC1 (9.3%)
PC2 (8.3%)
Medicago_singlecopy.vcf
PC1 (32.7%)
PC2 (26.9%)
Medicago_UCEsuper.vcf
PC1 (14.6%)
PC2 (13.3%)
Medicago_BUSCOsuper.vcf
PC1 (7.6%)
PC2 (6.5%)
Medicago_SCsuper.vcf
PC1 (15.9%)
PC2 (9.4%)
Medicago_genome.vcf
PC1 (10.8%)
PC2 (7.6%)
Genome cluster
1
2
3
4

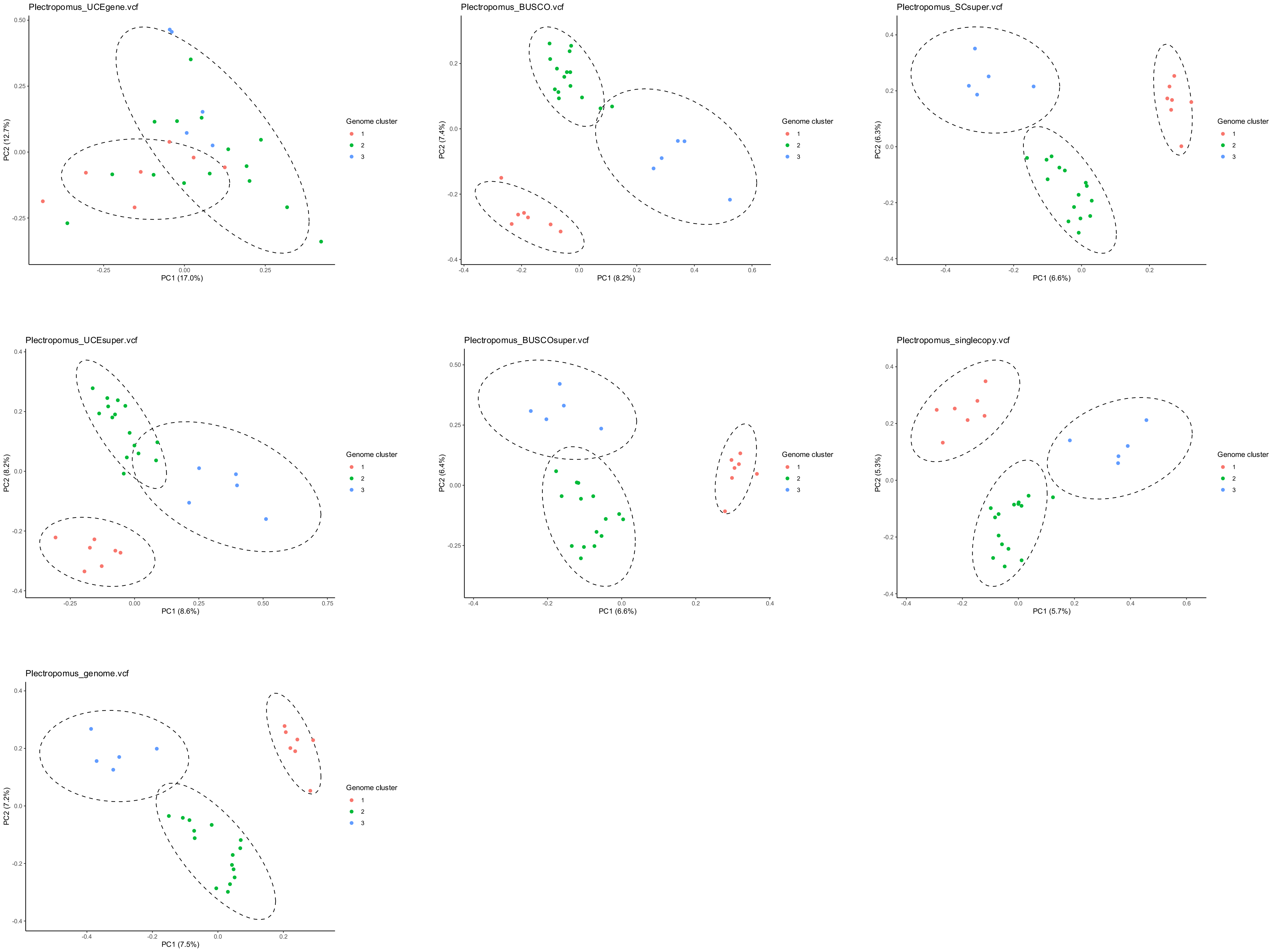
Plectropomus_UCEgene.vcf
PC1 (17.0%)
PC2 (12.7%)
-0.25 0.00 0.25 0.50
-0.25 0.00 0.25
Genome cluster
1
2
3
Plectropomus_BUSCO.vcf
PC1 (8.2%)
PC2 (7.4%)
-0.4 -0.2 0.0 0.2
-0.4 -0.2 0.0 0.2 0.4 0.6
Genome cluster
1
2
3
Plectropomus_SCsuper.vcf
PC1 (6.6%)
PC2 (6.3%)
-0.4 -0.2 0.0 0.2 0.4
-0.4 -0.2 0.0 0.2
Genome cluster
1
2
3
Plectropomus_UCEsuper.vcf
PC1 (8.6%)
PC2 (8.2%)
-0.4 -0.2 0.0 0.2 0.4
-0.25 0.00 0.25 0.50 0.75
Genome cluster
1
2
3
Plectropomus_BUSCOsuper.vcf
PC1 (6.6%)
PC2 (6.4%)
-0.25 0.00 0.25 0.50
-0.4 -0.2 0.0 0.2 0.4
Genome cluster
1
2
3
Plectropomus_singlecopy.vcf
PC1 (5.7%)
PC2 (5.3%)
-0.4 -0.2 0.0 0.2 0.4
-0.4 -0.2 0.0 0.2 0.4 0.6
Genome cluster
1
2
3
Plectropomus_genome.vcf
PC1 (7.5%)
PC2 (7.2%)
-0.4 -0.2 0.0 0.2 0.4
-0.4 -0.2 0.0 0.2
Genome cluster
1
2
3

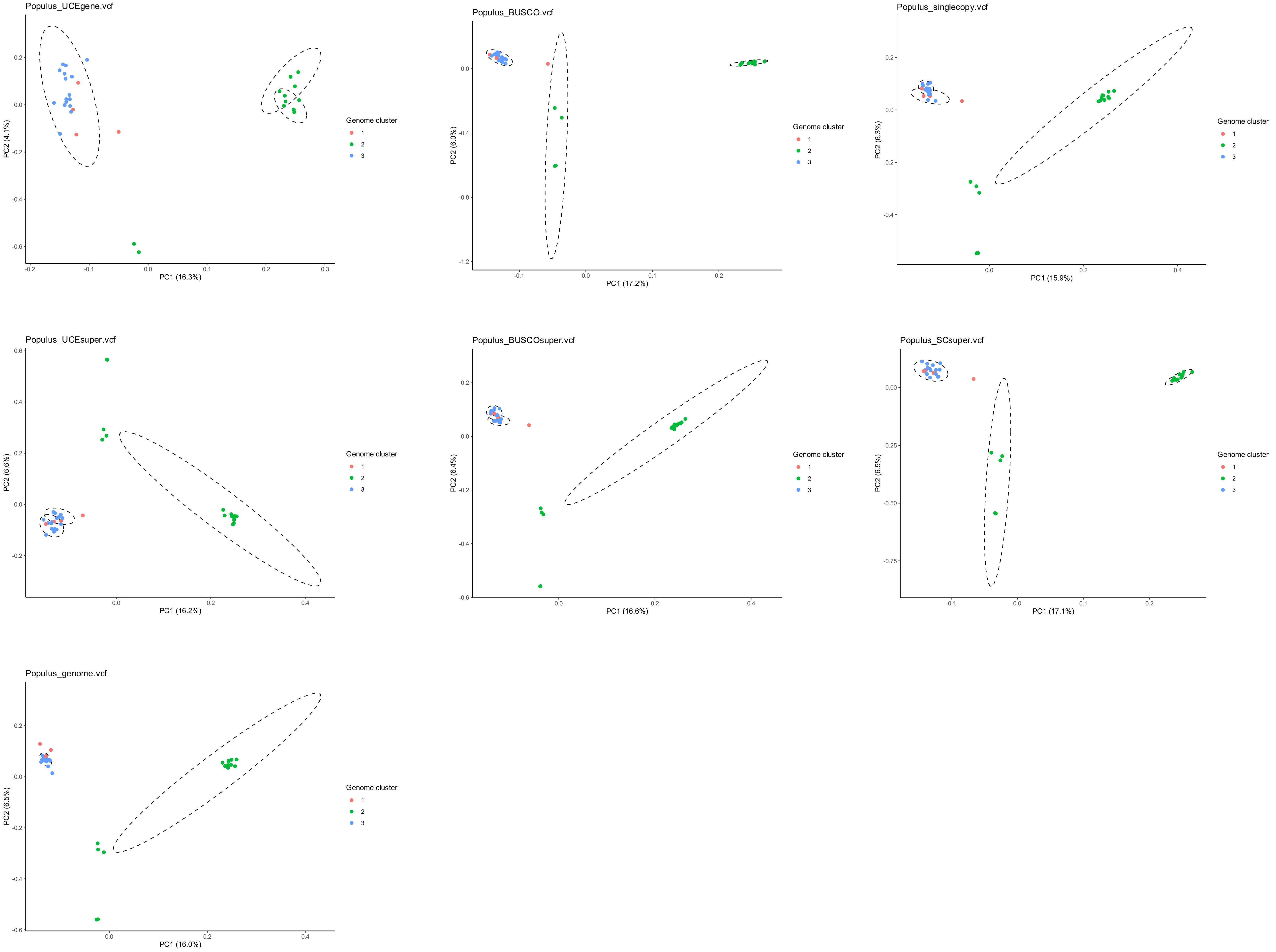
Populus_UCEgene.vcf
PC1 (16.3%)
PC2 (4.1%)
Populus_BUSCO.vcf
PC1 (17.2%)
PC2 (6.0%)
Populus_singlecopy.vcf
PC1 (15.9%)
PC2 (6.3%)
Populus_UCEsuper.vcf
PC1 (16.2%)
PC2 (6.6%)
Populus_BUSCOsuper.vcf
PC1 (16.6%)
PC2 (6.4%)
Populus_SCsuper.vcf
PC1 (17.1%)
PC2 (6.5%)
Populus_genome.vcf
PC1 (16.0%)
PC2 (6.5%)
Genome cluster
1
2
3

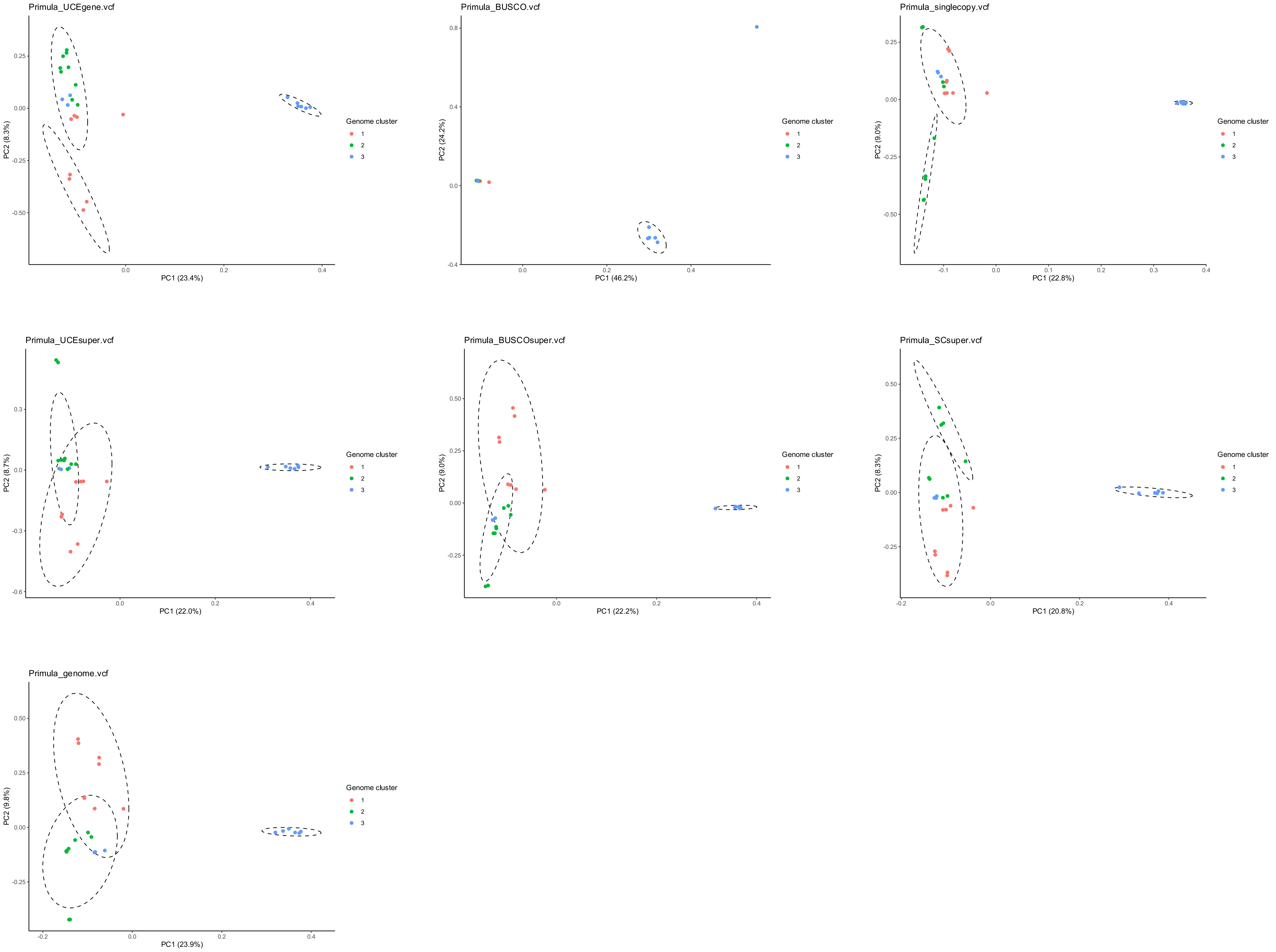
Primula_UCEgene.vcf
PC1 (23.4%)
PC2 (8.3%)
Genome cluster
1
2
3
Primula_BUSCO.vcf
PC1 (46.2%)
PC2 (24.2%)
Genome cluster
1
2
3
Primula_singlecopy.vcf
PC1 (22.8%)
PC2 (9.0%)
Genome cluster
1
2
3
Primula_UCEsuper.vcf
PC1 (22.0%)
PC2 (8.7%)
Genome cluster
1
2
3
Primula_BUSCOsuper.vcf
PC1 (22.2%)
PC2 (9.0%)
Genome cluster
1
2
3
Primula_SCsuper.vcf
PC1 (20.8%)
PC2 (8.3%)
Genome cluster
1
2
3
Primula_genome.vcf
PC1 (23.9%)
PC2 (9.8%)
Genome cluster
1
2
3

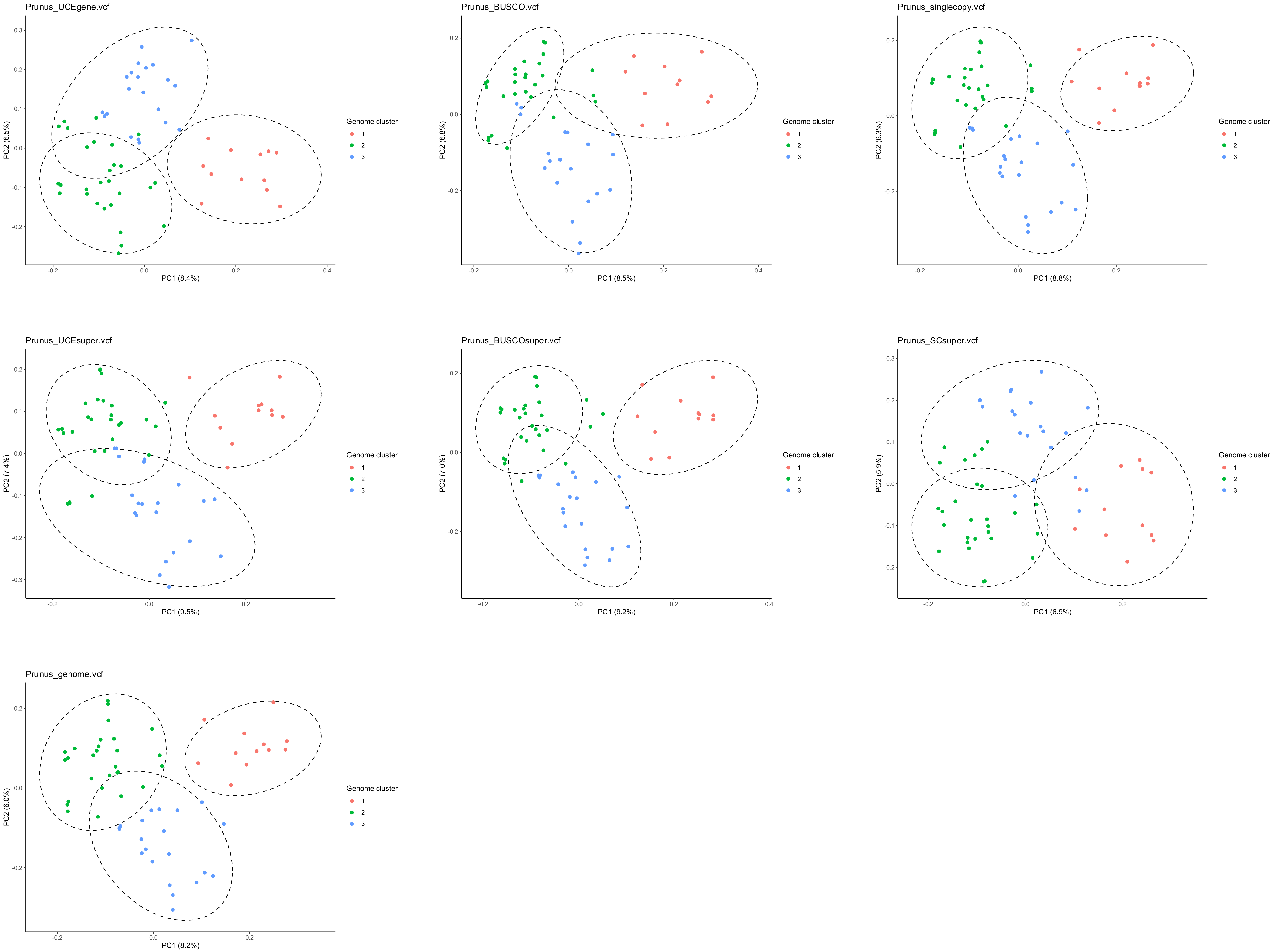

Prunus_UCEgene.vcf
PC1 (8.4%)
PC2 (6.5%)
Prunus_BUSCO.vcf
PC1 (8.5%)
PC2 (6.8%)
Prunus_singlecopy.vcf
PC1 (8.8%)
PC2 (6.3%)
Prunus_UCEsuper.vcf
PC1 (9.5%)
PC2 (7.4%)
Prunus_BUSCOsuper.vcf
PC1 (9.2%)
PC2 (7.0%)
Prunus_SCsuper.vcf
PC1 (6.9%)
PC2 (5.9%)
Prunus_genome.vcf
PC1 (8.2%)
PC2 (6.0%)
Genome cluster
1
2
3

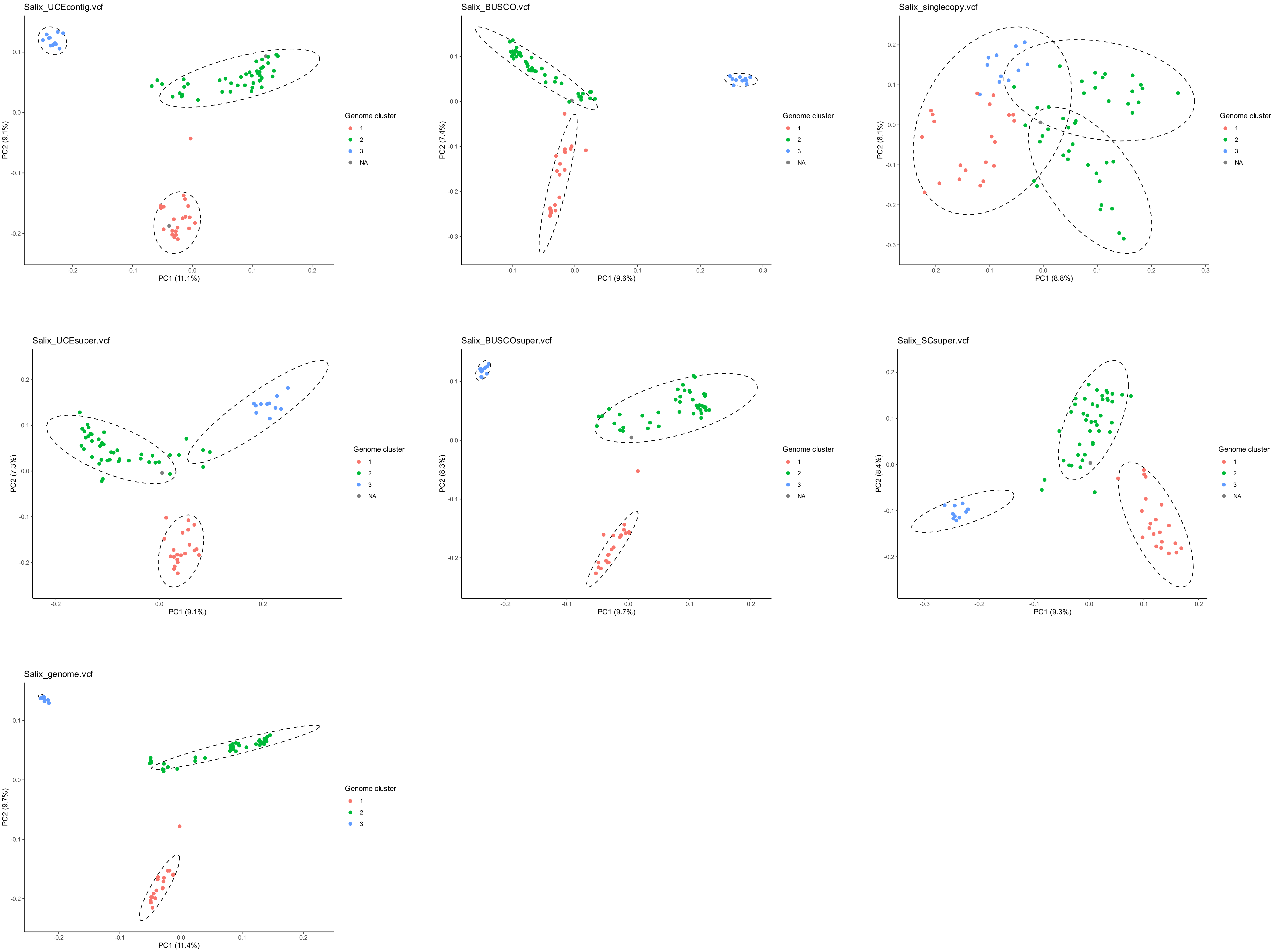

Salix_UCEcontig.vcf
PC1 (11.1%)
PC2 (9.1%)
Genome cluster
1
2
3
NA
Salix_BUSCO.vcf
PC1 (9.6%)
PC2 (7.4%)
Genome cluster
1
2
3
NA
Salix_singlecopy.vcf
PC1 (8.8%)
PC2 (8.1%)
Genome cluster
1
2
3
NA
Salix_UCEsuper.vcf
PC1 (9.1%)
PC2 (7.3%)
Genome cluster
1
2
3
NA
Salix_BUSCOsuper.vcf
PC1 (9.7%)
PC2 (8.3%)
Genome cluster
1
2
3
NA
Salix_SCsuper.vcf
PC1 (9.3%)
PC2 (8.4%)
Genome cluster
1
2
3
NA
Salix_genome.vcf
PC1 (11.4%)
PC2 (9.7%)
Genome cluster
1
2
3

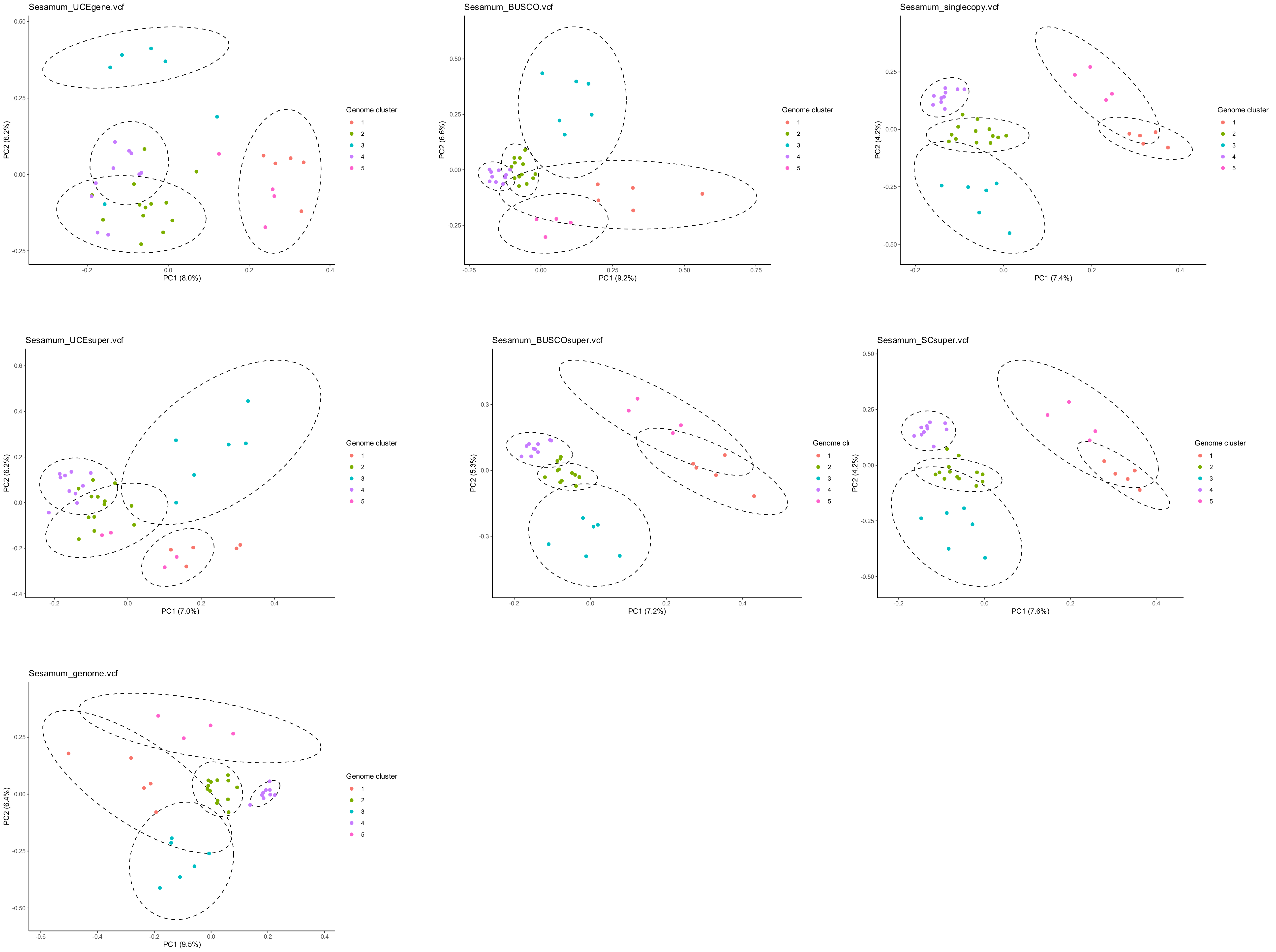
Sesamum_UCEgene.vcf
PC1 (8.0%)
PC2 (6.2%)
Genome cluster
1
2
3
4
5
Sesamum_BUSCO.vcf
PC1 (9.2%)
PC2 (6.6%)
Genome cluster
1
2
3
4
5
Sesamum_singlecopy.vcf
PC1 (7.4%)
PC2 (4.2%)
Genome cluster
1
2
3
4
5
Sesamum_UCEsuper.vcf
PC1 (7.0%)
PC2 (6.2%)
Genome cluster
1
2
3
4
5
Sesamum_BUSCOsuper.vcf
PC1 (7.2%)
PC2 (5.3%)
1
2
3
4
5
Sesamum_SCsuper.vcf
PC1 (7.6%)
PC2 (4.2%)
Genome cluster
1
2
3
4
5
Sesamum_genome.vcf
PC1 (9.5%)
PC2 (6.4%)
Genome cluster
1
2
3
4
5

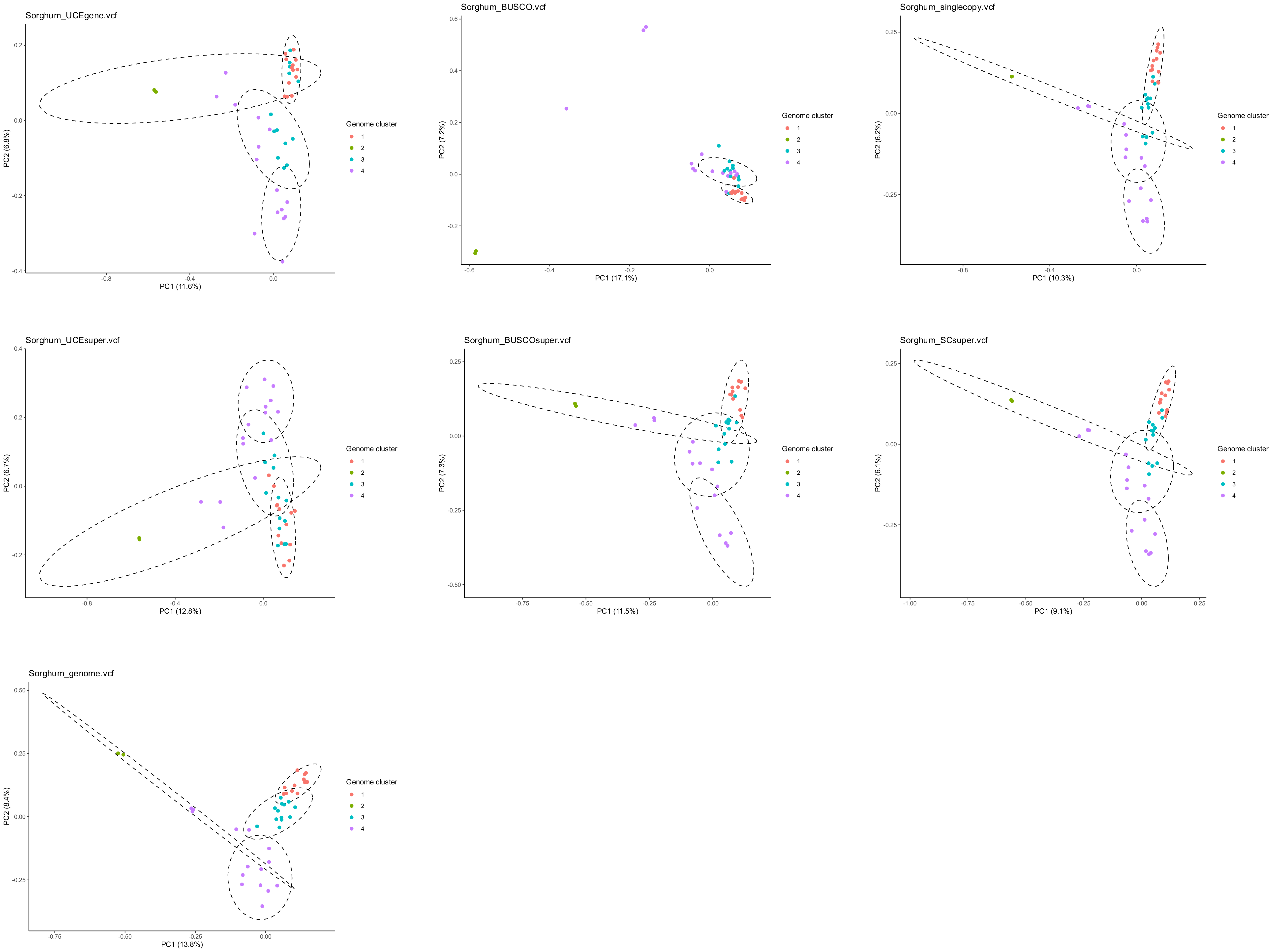
Sorghum_UCEgene.vcf
PC1 (11.6%)
PC2 (6.8%)
Genome cluster
1
2
3
4
Sorghum_BUSCO.vcf
PC1 (17.1%)
PC2 (7.2%)
Genome cluster
1
2
3
4
Sorghum_singlecopy.vcf
PC1 (10.3%)
PC2 (6.2%)
Genome cluster
1
2
3
4
Sorghum_UCEsuper.vcf
PC1 (12.8%)
PC2 (6.7%)
Genome cluster
1
2
3
4
Sorghum_BUSCOsuper.vcf
PC1 (11.5%)
PC2 (7.3%)
Genome cluster
1
2
3
4
Sorghum_SCsuper.vcf
PC1 (9.1%)
PC2 (6.1%)
Genome cluster
1
2
3
4
Sorghum_genome.vcf
PC1 (13.8%)
PC2 (8.4%)
Genome cluster
1
2
3
4

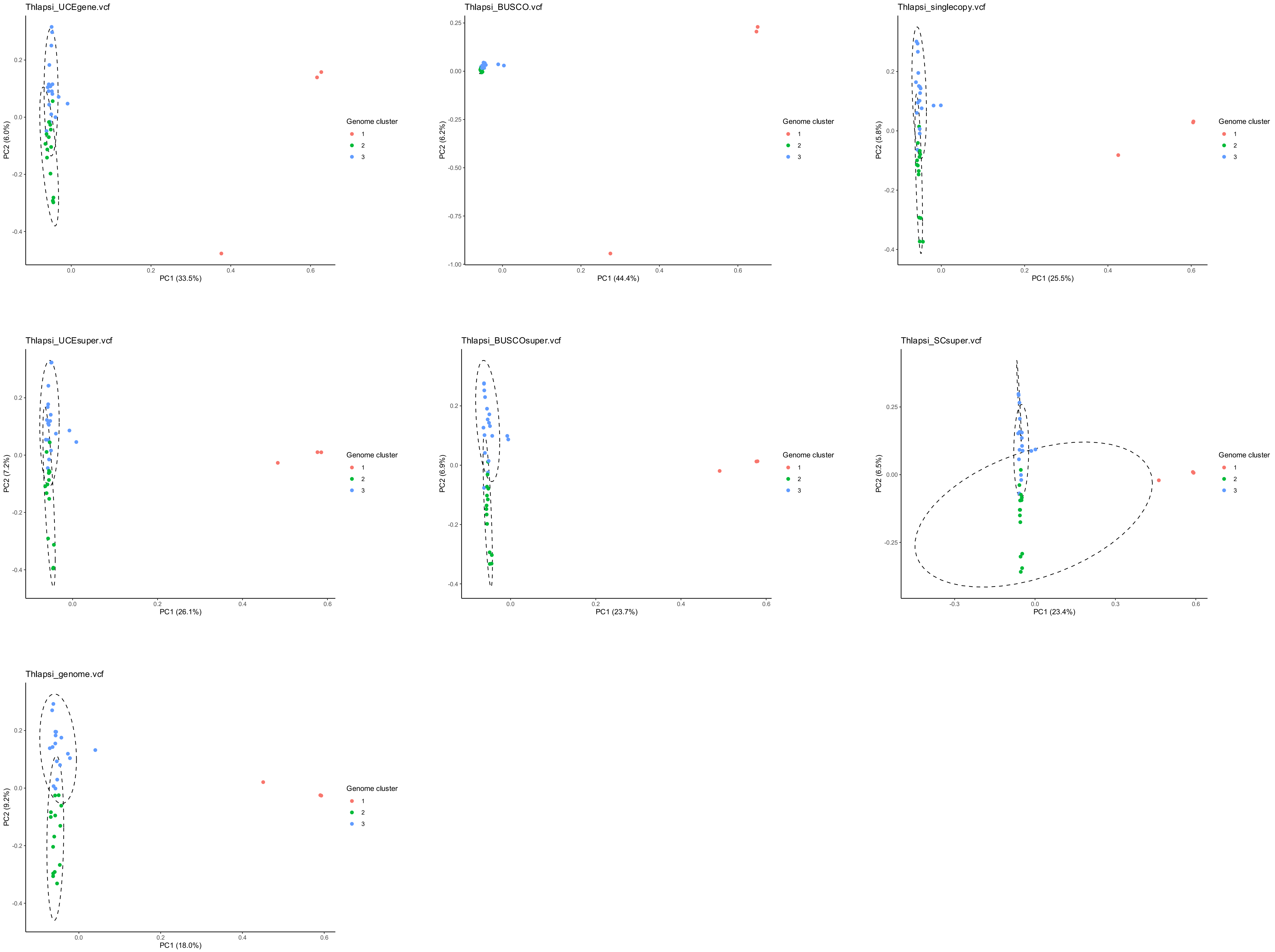

Thlapsi_UCEgene.vcf
PC1 (33.5%)
PC2 (6.0%)
Genome cluster
1
2
3
Thlapsi_BUSCO.vcf
PC1 (44.4%)
PC2 (6.2%)
Thlapsi_singlecopy.vcf
PC1 (25.5%)
PC2 (5.8%)
Thlapsi_UCEsuper.vcf
PC1 (26.1%)
PC2 (7.2%)
Thlapsi_BUSCOsuper.vcf
PC1 (23.7%)
PC2 (6.9%)
Thlapsi_SCsuper.vcf
PC1 (23.4%)
PC2 (6.5%)
Thlapsi_genome.vcf
PC1 (18.0%)
PC2 (9.2%)

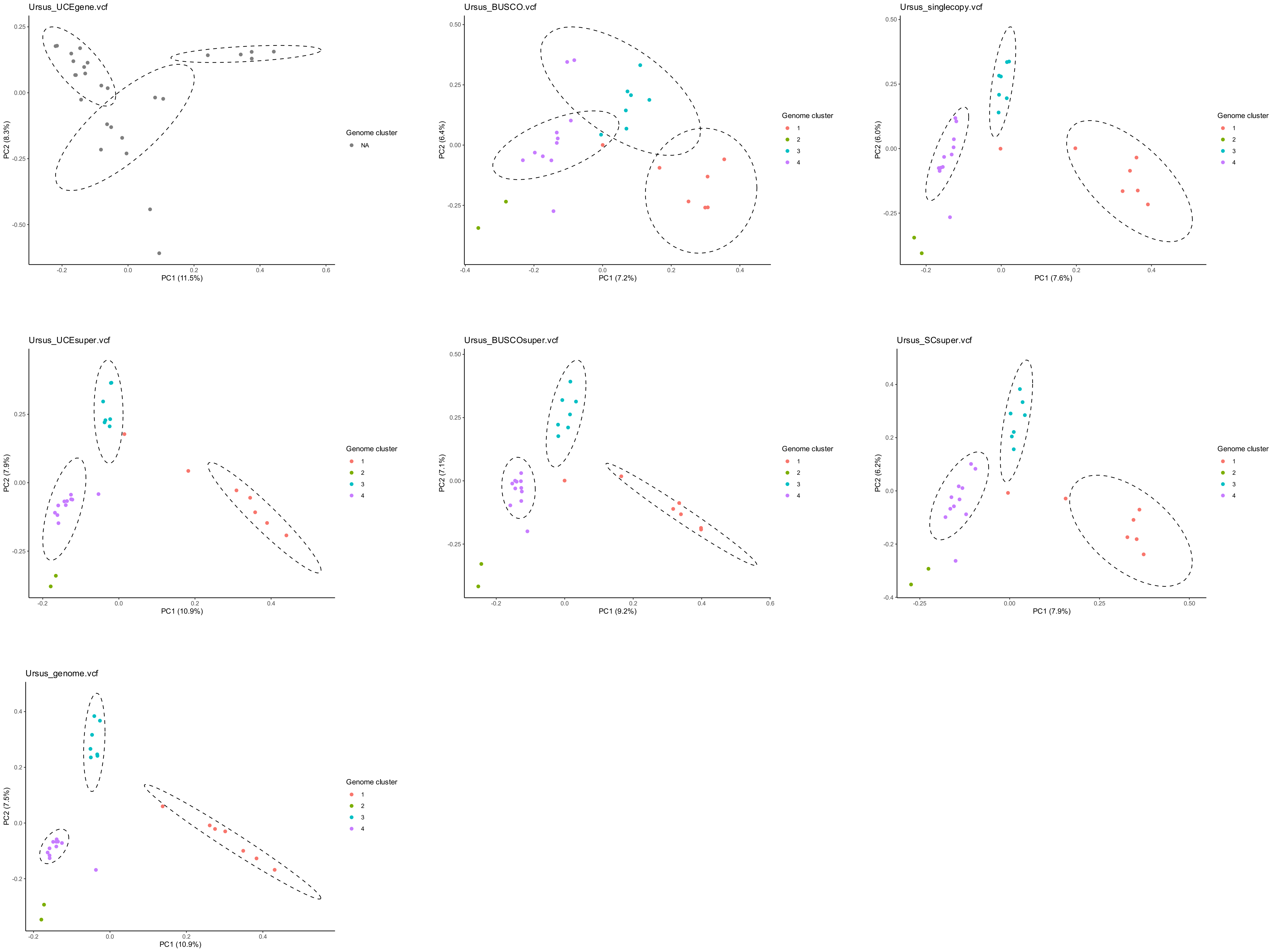

Ursus_UCEgene.vcf
PC1 (11.5%)
PC2 (8.3%)
Genome cluster
NA
Ursus_BUSCO.vcf
PC1 (7.2%)
PC2 (6.4%)
Genome cluster
1
2
3
4
Ursus_singlecopy.vcf
PC1 (7.6%)
PC2 (6.0%)
Genome cluster
1
2
3
4
Ursus_UCEsuper.vcf
PC1 (10.9%)
PC2 (7.9%)
Genome cluster
1
2
3
4
Ursus_BUSCOsuper.vcf
PC1 (9.2%)
PC2 (7.1%)
Genome cluster
1
2
3
4
Ursus_SCsuper.vcf
PC1 (7.9%)
PC2 (6.2%)
Genome cluster
1
2
3
4
Ursus_genome.vcf
PC1 (10.9%)
PC2 (7.5%)
Genome cluster
1
2
3
4

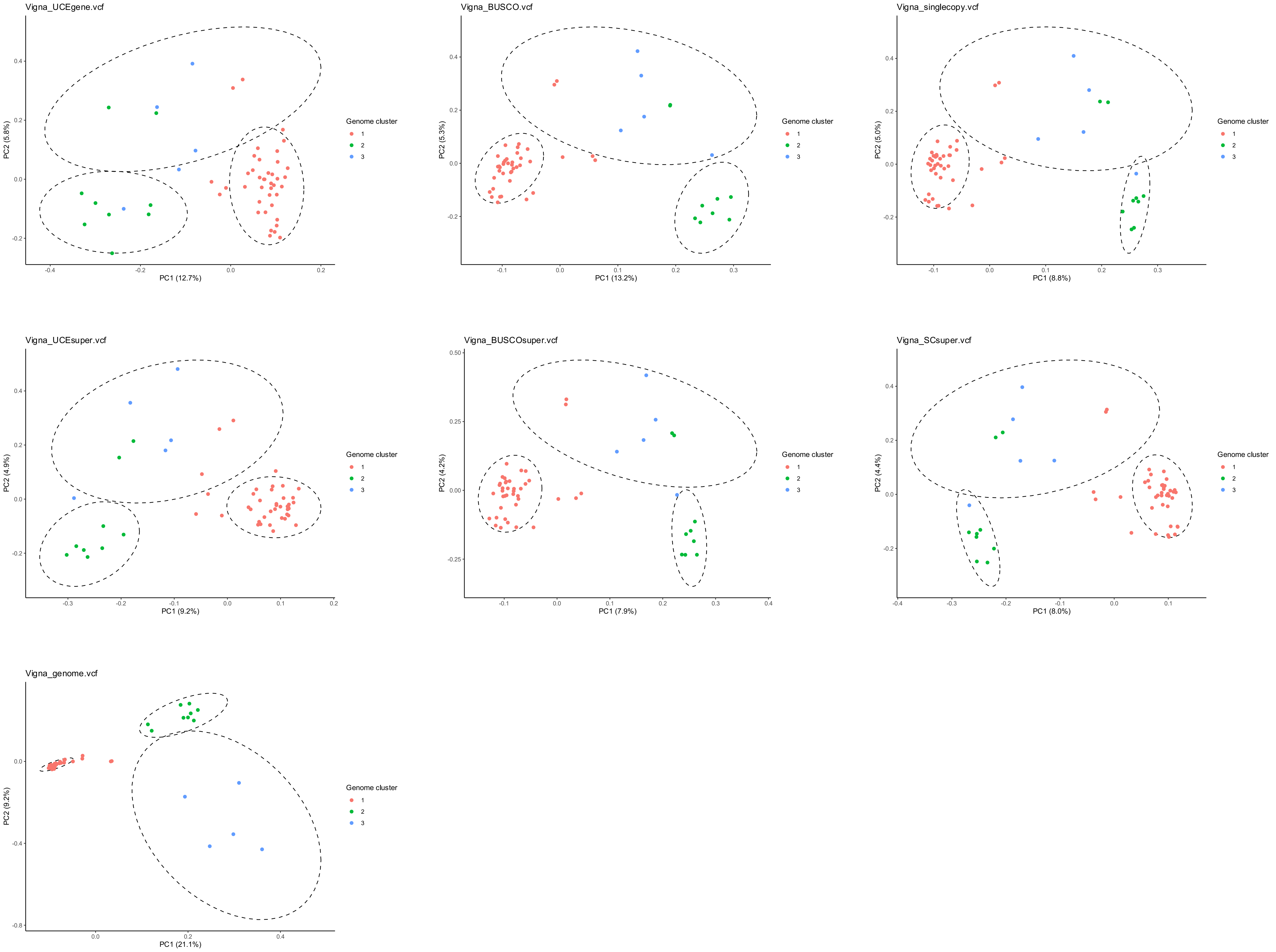
Vigna_UCEgene.vcf
PC1 (12.7%)
PC2 (5.8%)
Genome cluster
1
2
3
Vigna_BUSCO.vcf
PC1 (13.2%)
PC2 (5.3%)
Genome cluster
1
2
3
Vigna_singlecopy.vcf
PC1 (8.8%)
PC2 (5.0%)
Genome cluster
1
2
3
Vigna_UCEsuper.vcf
PC1 (9.2%)
PC2 (4.9%)
Genome cluster
1
2
3
Vigna_BUSCOsuper.vcf
PC1 (7.9%)
PC2 (4.2%)
Genome cluster
1
2
3
Vigna_SCsuper.vcf
PC1 (8.0%)
PC2 (4.4%)
Genome cluster
1
2
3
Vigna_genome.vcf
PC1 (21.1%)
PC2 (9.2%)
Genome cluster
1
2
3

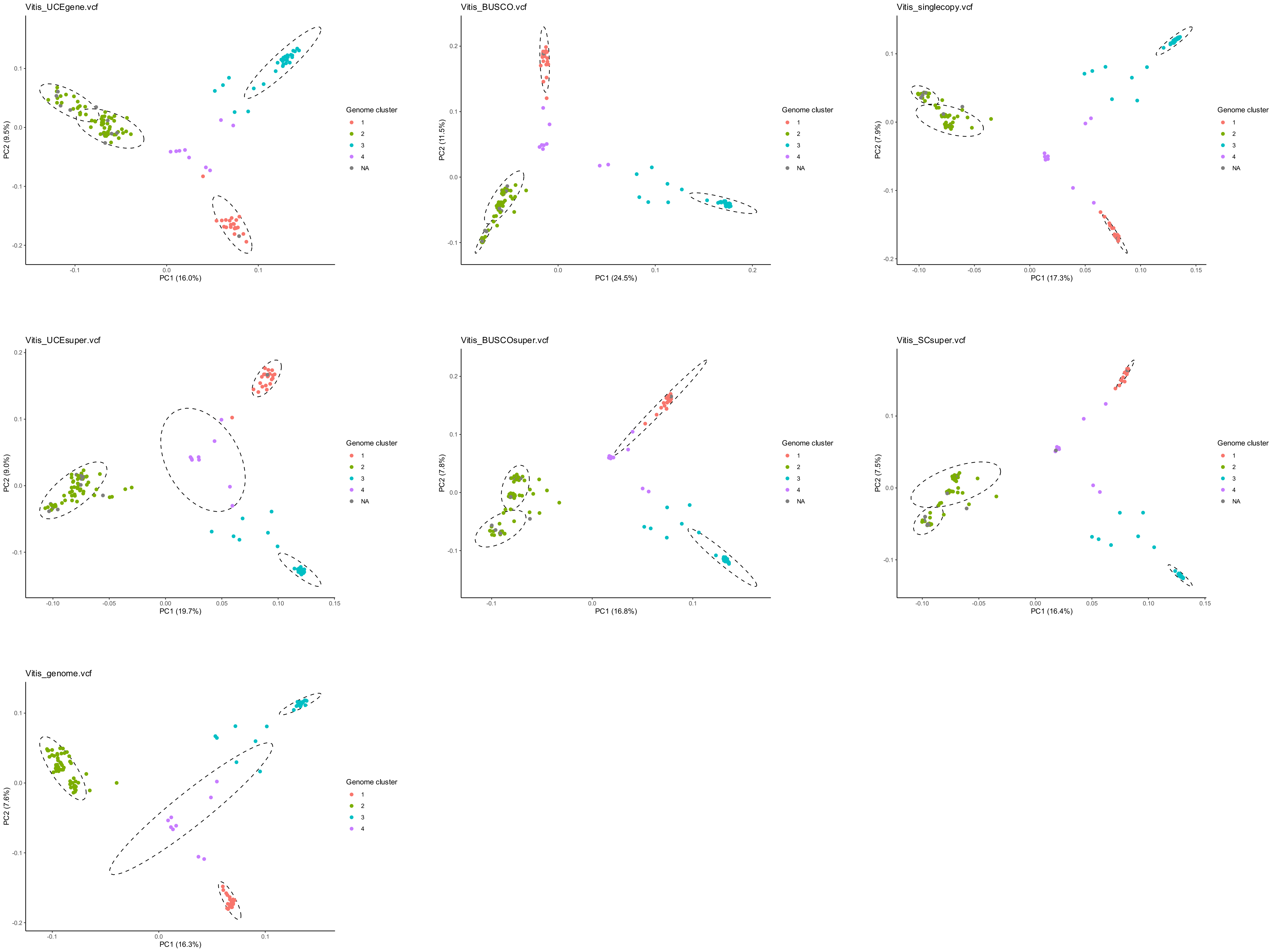
Vitis_UCEgene.vcf
PC1 (16.0%)
PC2 (9.5%)
Genome cluster
1
2
3
4
NA
Vitis_BUSCO.vcf
PC1 (24.5%)
PC2 (11.5%)
Genome cluster
1
2
3
4
NA
Vitis_singlecopy.vcf
PC1 (17.3%)
PC2 (7.9%)
Genome cluster
1
2
3
4
NA
Vitis_UCEsuper.vcf
PC1 (19.7%)
PC2 (9.0%)
Genome cluster
1
2
3
4
NA
Vitis_BUSCOsuper.vcf
PC1 (16.8%)
PC2 (7.8%)
Genome cluster
1
2
3
4
NA
Vitis_SCsuper.vcf
PC1 (16.4%)
PC2 (7.5%)
Genome cluster
1
2
3
4
NA
Vitis_genome.vcf
PC1 (16.3%)
PC2 (7.6%)
Genome cluster
1
2
3
4

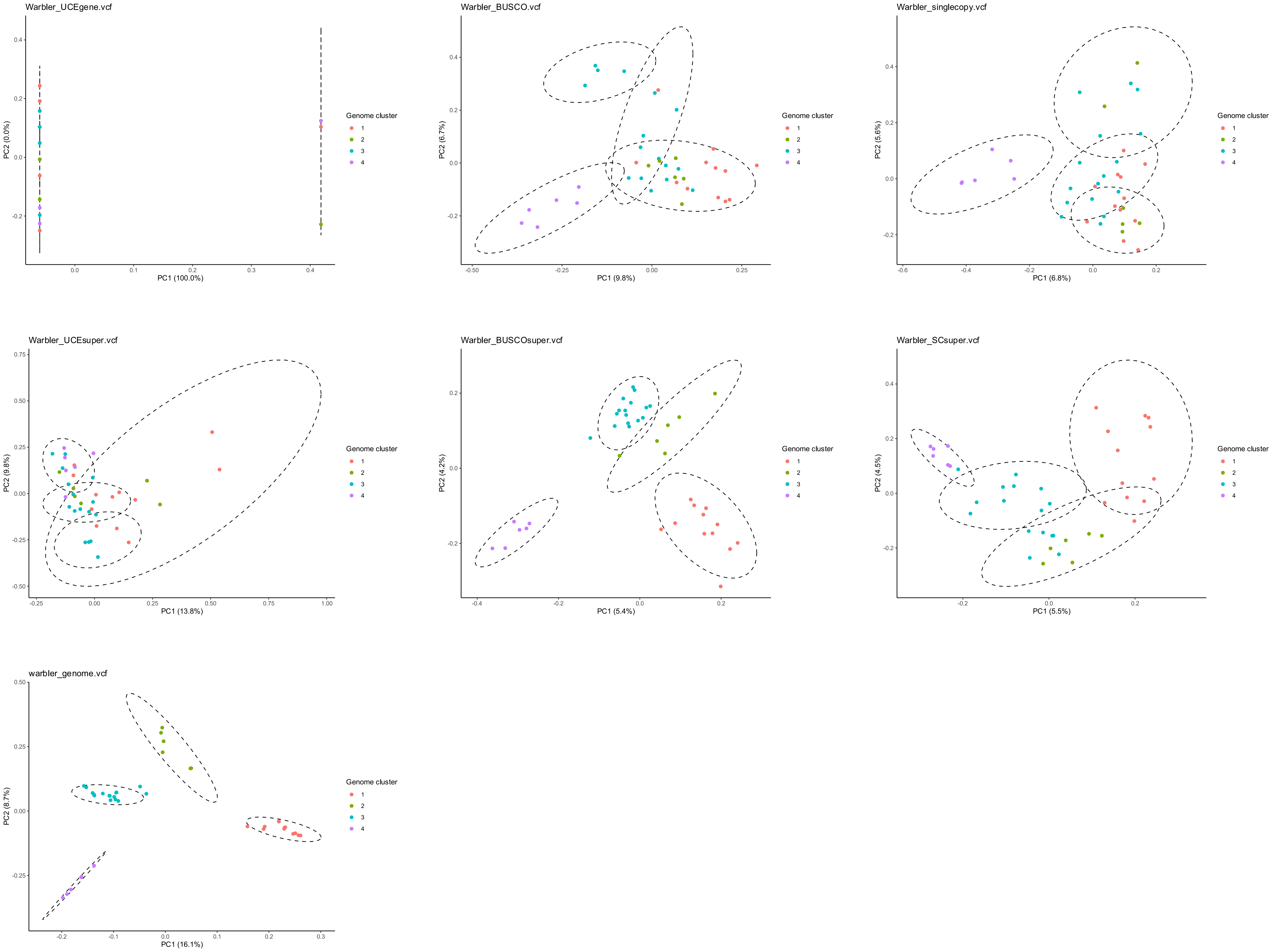
Warbler_UCEgene.vcf
PC1 (100.0%)
PC2 (0.0%)
Warbler_BUSCO.vcf
PC1 (9.8%)
PC2 (6.7%)
Warbler_singlecopy.vcf
PC1 (6.8%)
PC2 (5.6%)
Warbler_UCEsuper.vcf
PC1 (13.8%)
PC2 (9.8%)
Warbler_BUSCOsuper.vcf
PC1 (5.4%)
PC2 (4.2%)
Warbler_SCsuper.vcf
PC1 (5.5%)
PC2 (4.5%)
warbler_genome.vcf
PC1 (16.1%)
PC2 (8.7%)
Genome cluster
1
2
3
4
