## Supplemental Figure 6 for "The power to resolve relationships: identifying incongruence and precision of reduced representation and genome-wide data in phylogenomics and population genomics"

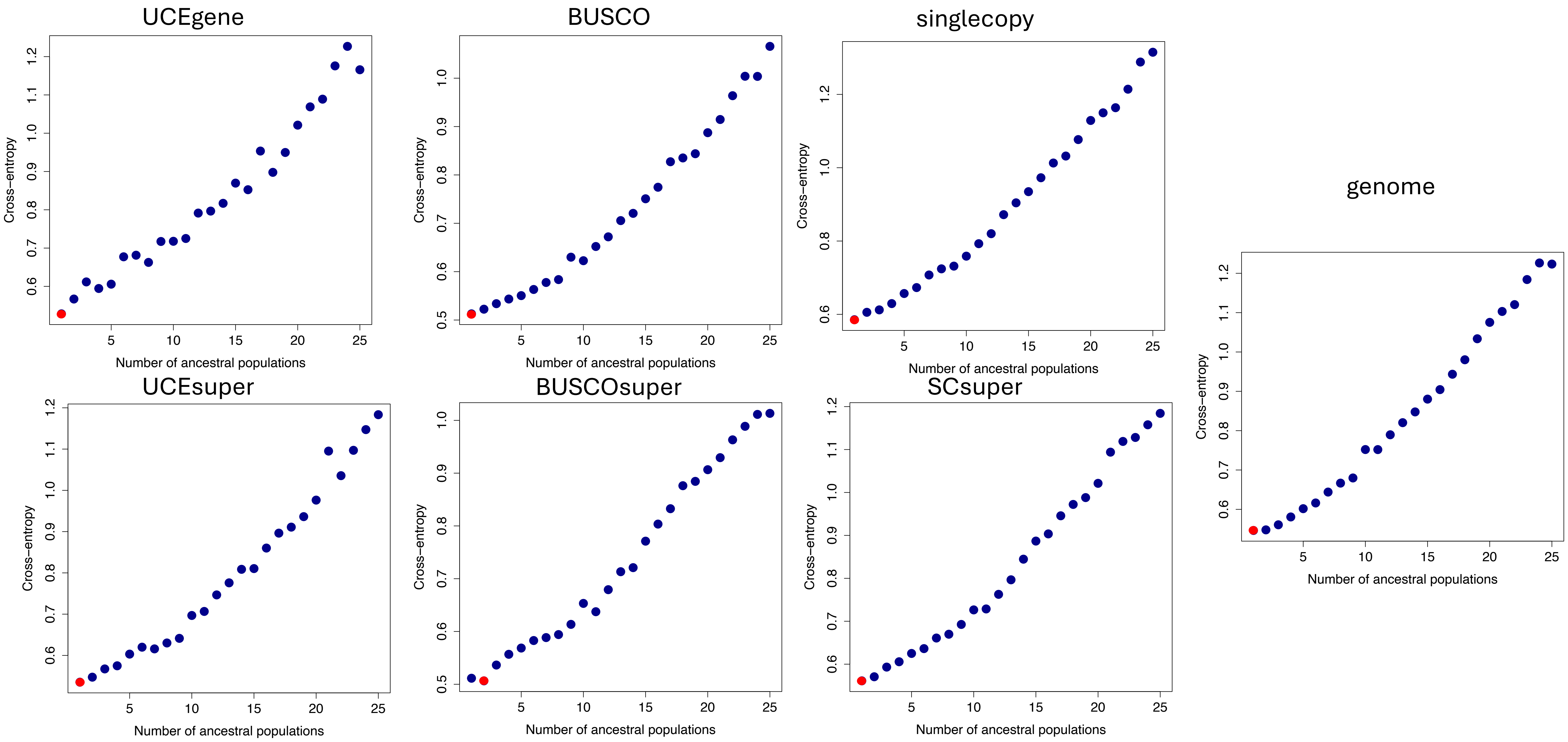
UCEgene
BUSCO
singlecopy
genome
UCEsuper
BUSCOsuper
SCsuper
Cross−entropy
Number of ancestral populations

*Acer*

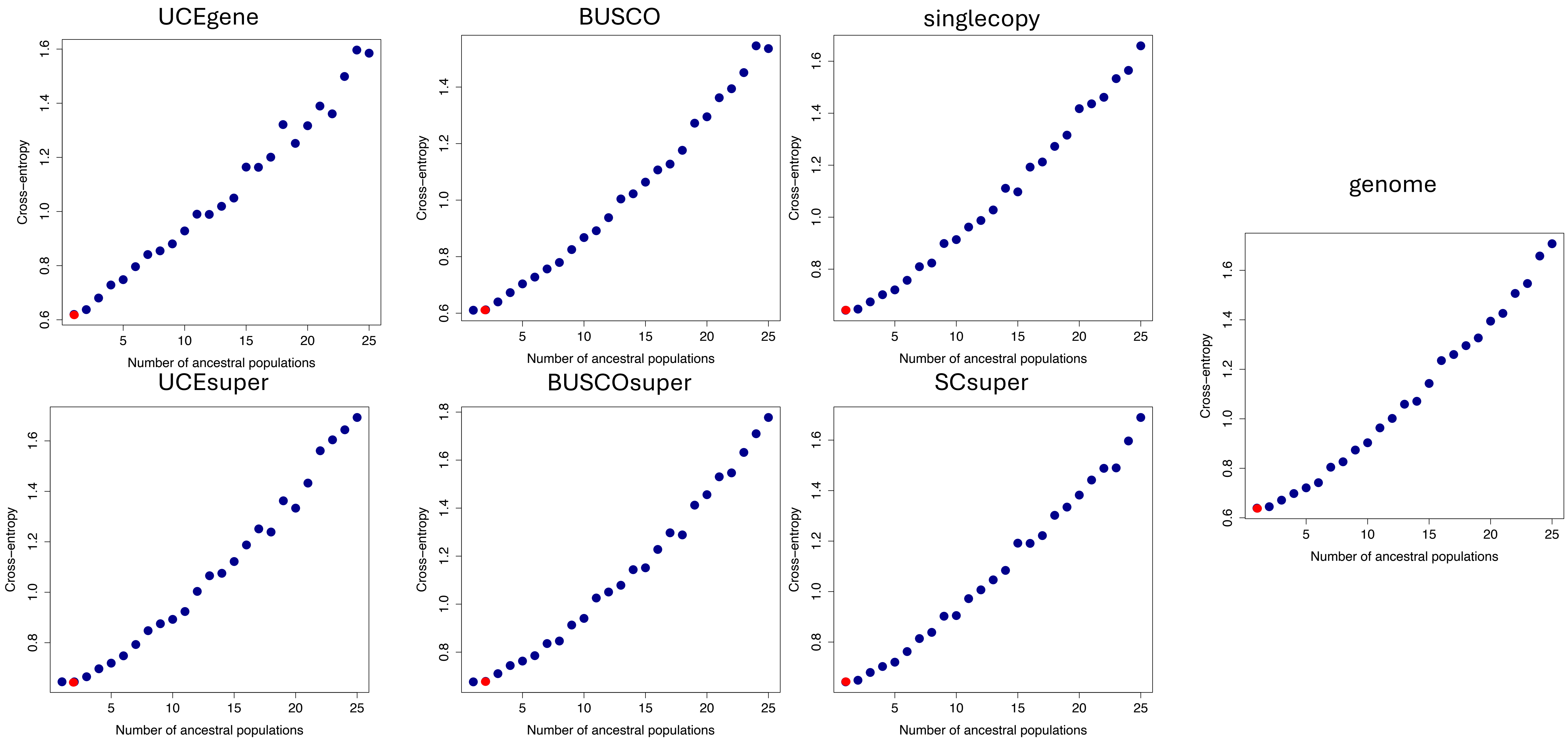
UCEgene
BUSCO
singlecopy
genome
UCEsuper
BUSCOsuper
SCsuper
Cross−entropy
Number of ancestral populations

*Aquilegia*

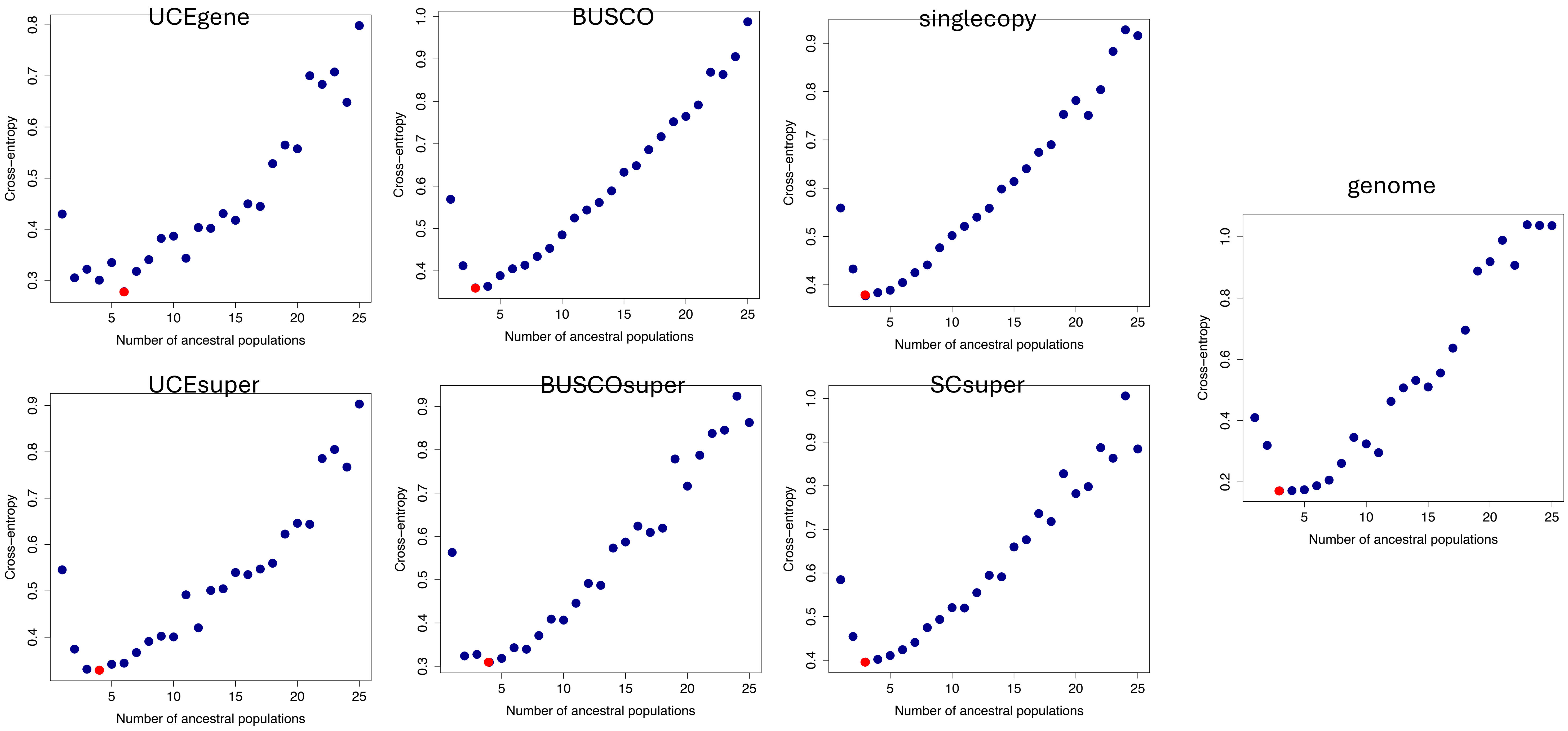
UCEgene
BUSCO
singlecopy
genome
UCEsuper
BUSCOsuper
SCsuper
Cross−entropy
Number of ancestral populations

*Atriplex*

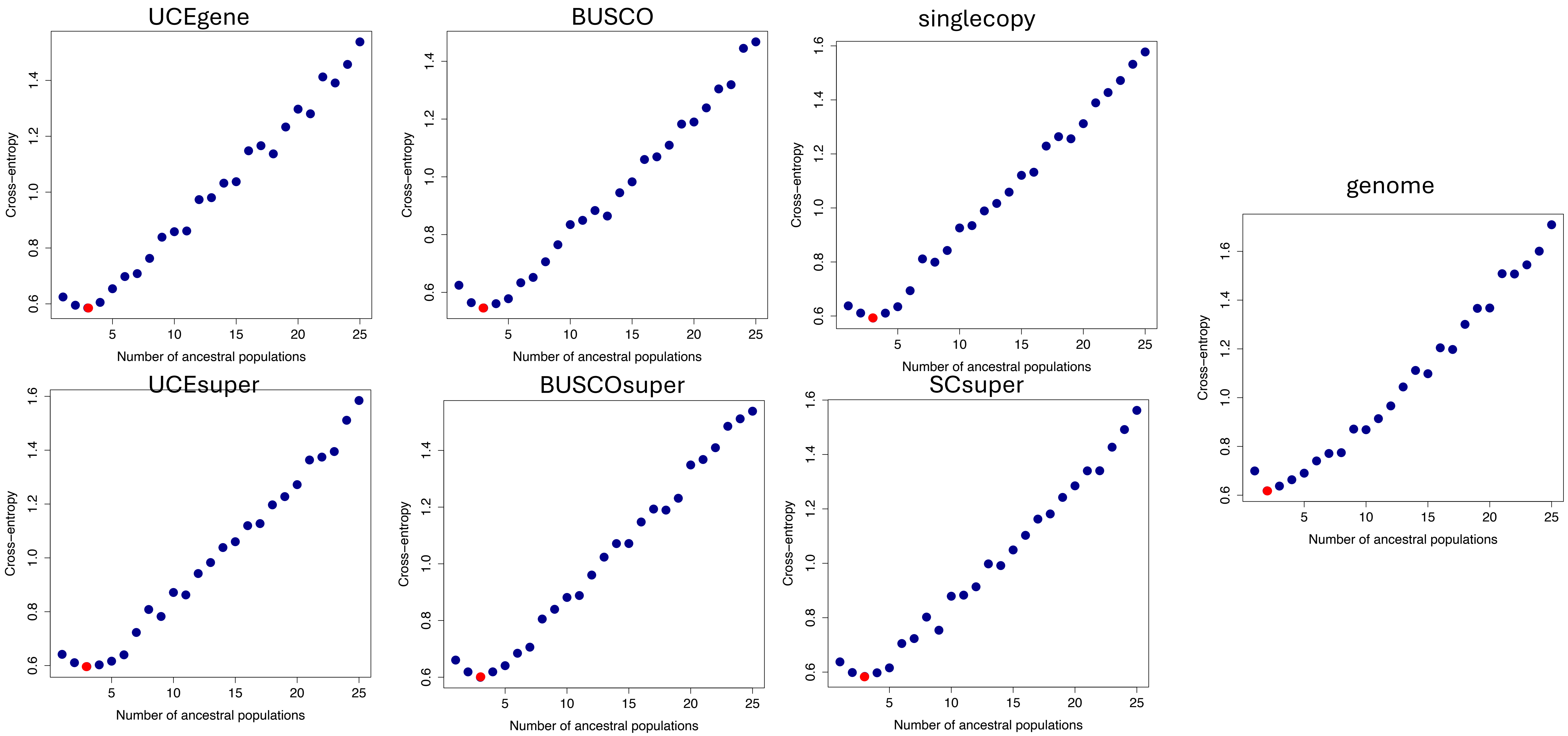
UCEgene
Cross−entropy
Number of ancestral populations
BUSCO
Cross−entropy
Number of ancestral populations
singlecopy
Cross−entropy
Number of ancestral populations
genome
Cross−entropy
Number of ancestral populations
UCEsuper
Cross−entropy
Number of ancestral populations
BUSCOsuper
Cross−entropy
Number of ancestral populations
SCsuper
Cross−entropy
Number of ancestral populations

*Begonia*

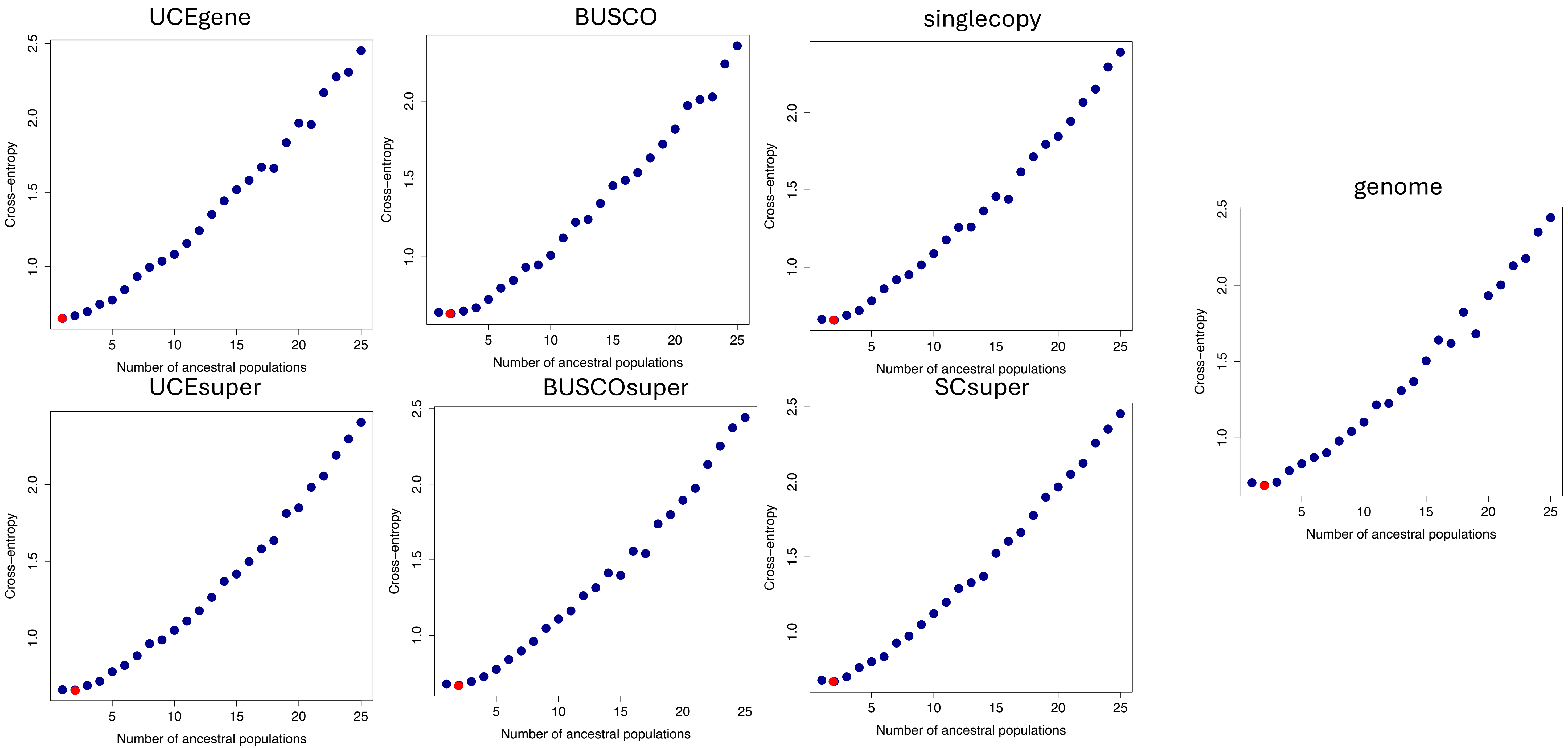
UCEgene
BUSCO
singlecopy
genome
UCEsuper
BUSCOsuper
SCsuper
Cross-entropy
Number of ancestral populations


*Camellia*

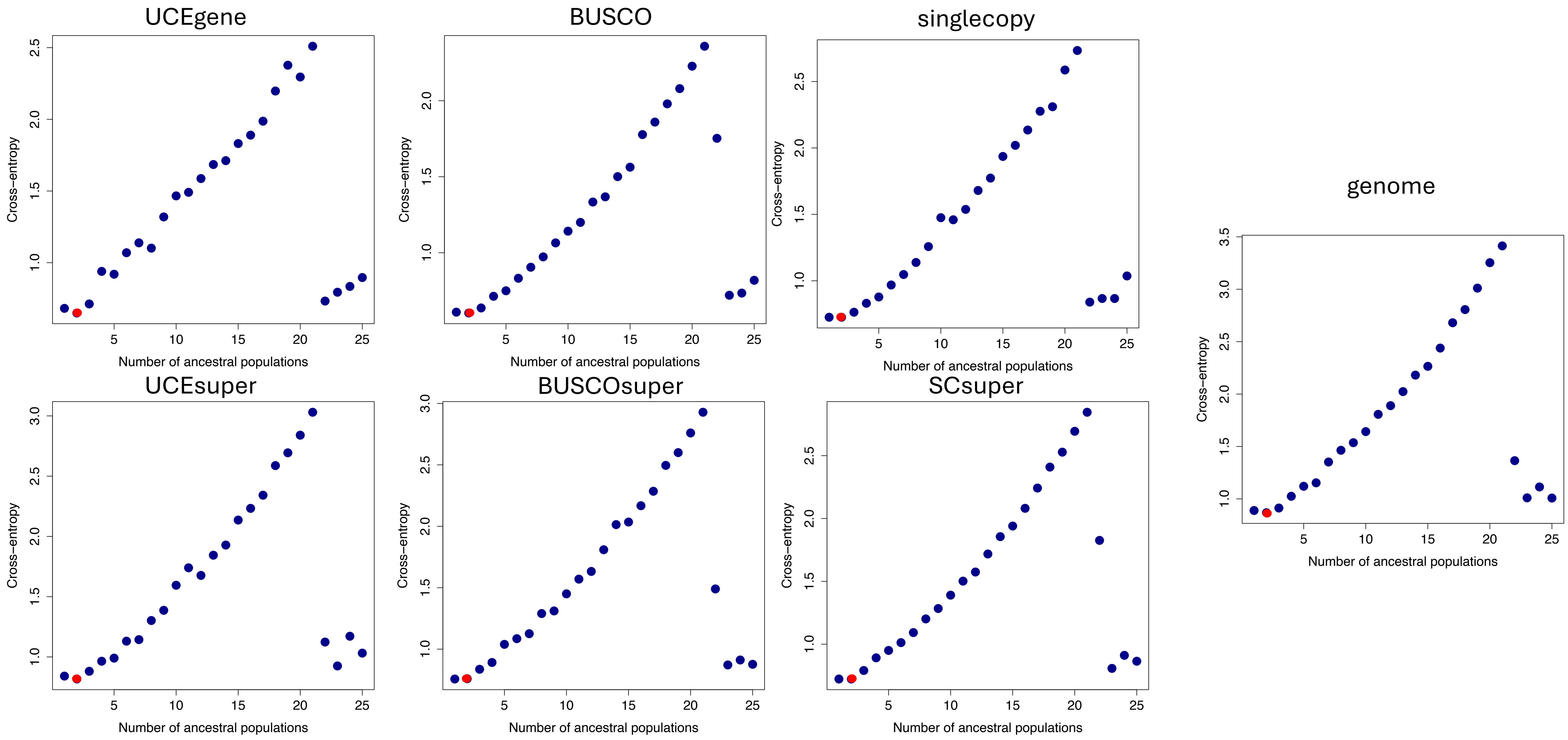

UCEgene
BUSCO
singlecopy
genome
UCEsuper
BUSCOsuper
SCsuper
Cross−entropy
Number of ancestral populations


*Canis*

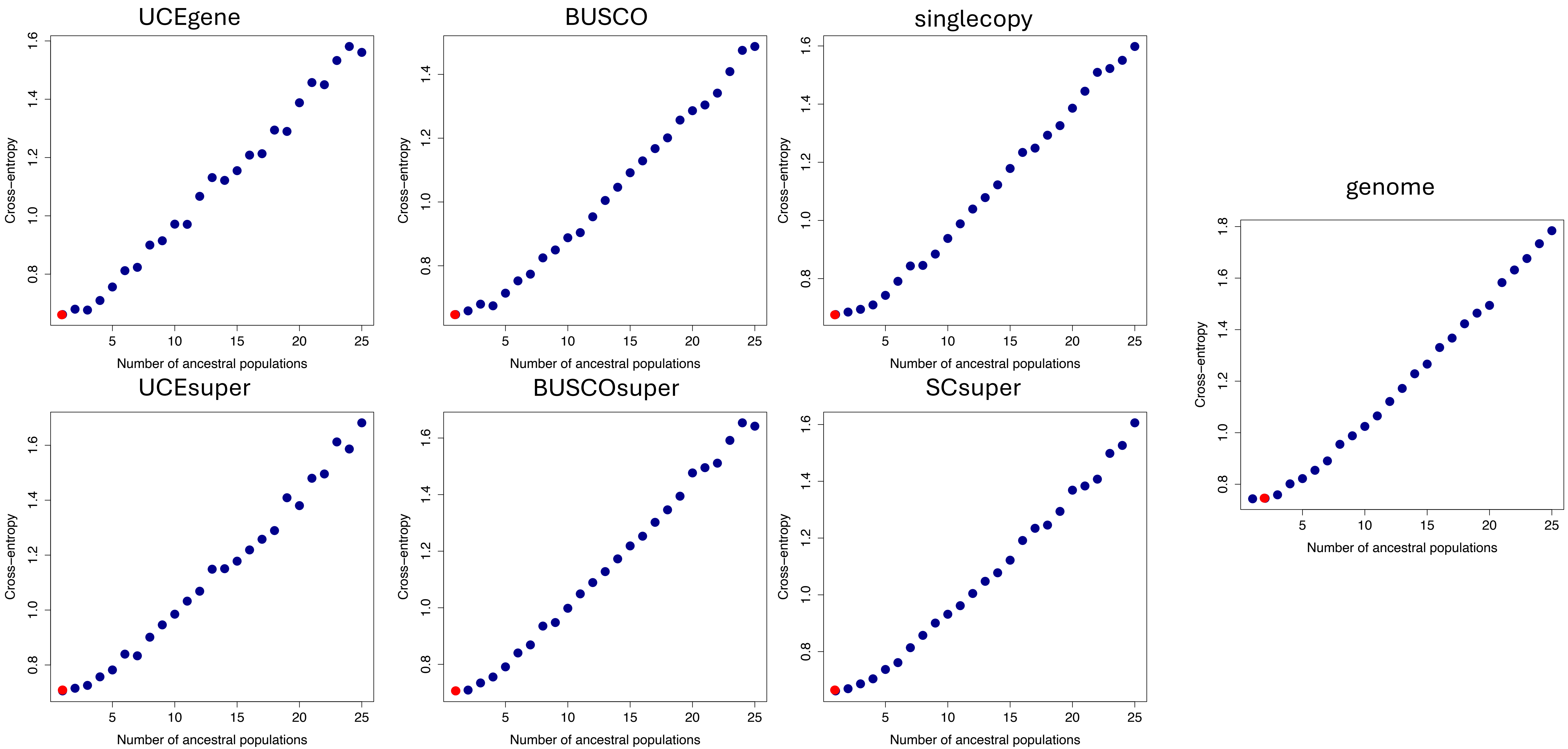
UCEgene
BUSCO
singlecopy
genome
UCEsuper
BUSCOsuper
SCsuper
Cross-entropy
Number of ancestral populations

*Cannabis*

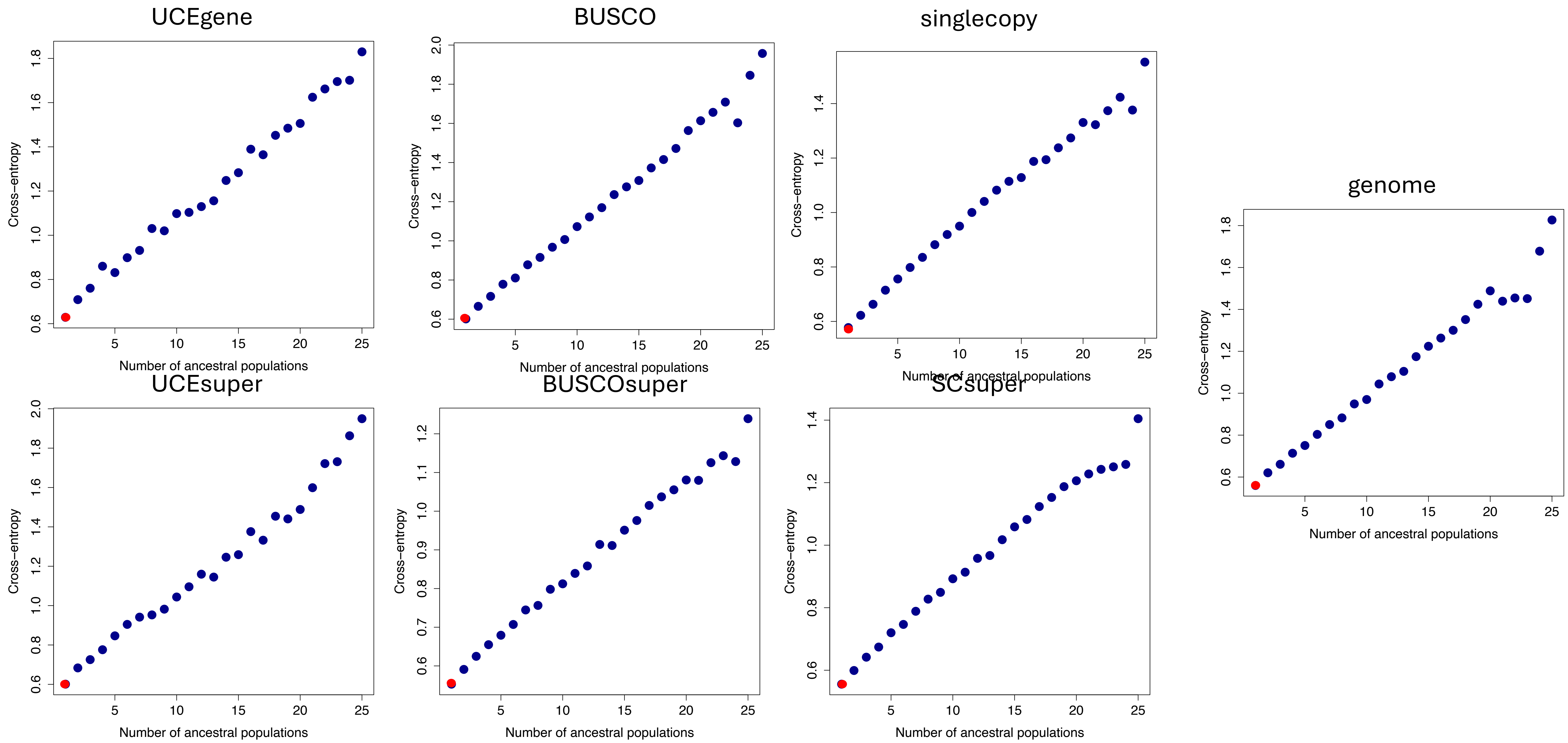
UCEgene
BUSCO
singlecopy
genome
UCEsuper
BUSCOsuper
SCsuper
Cross−entropy
Number of ancestral populations

*Citrus*

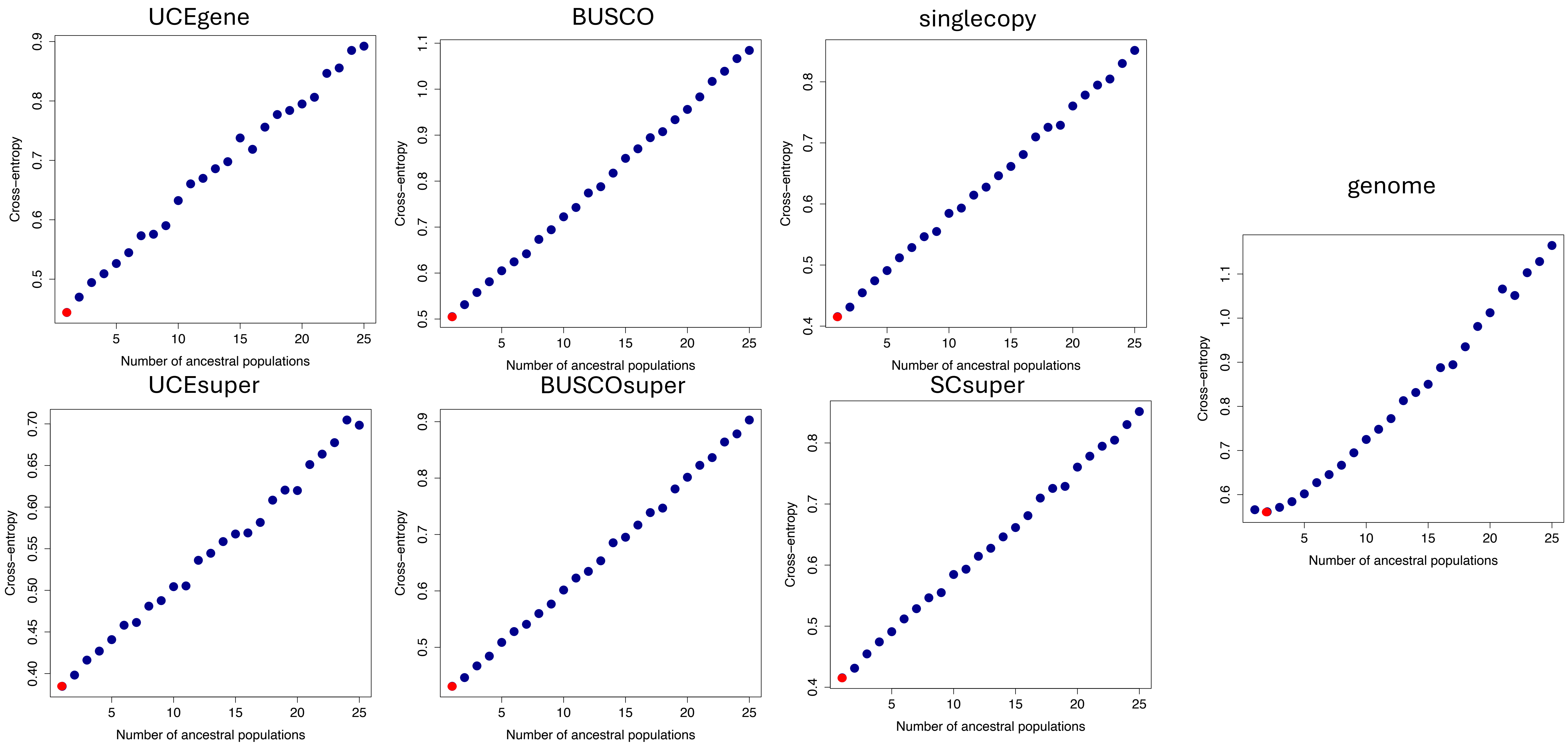
UCEgene
BUSCO
singlecopy
genome
UCEsuper
BUSCOsuper
SCsuper
Cross−entropy
Number of ancestral populations

Coffea

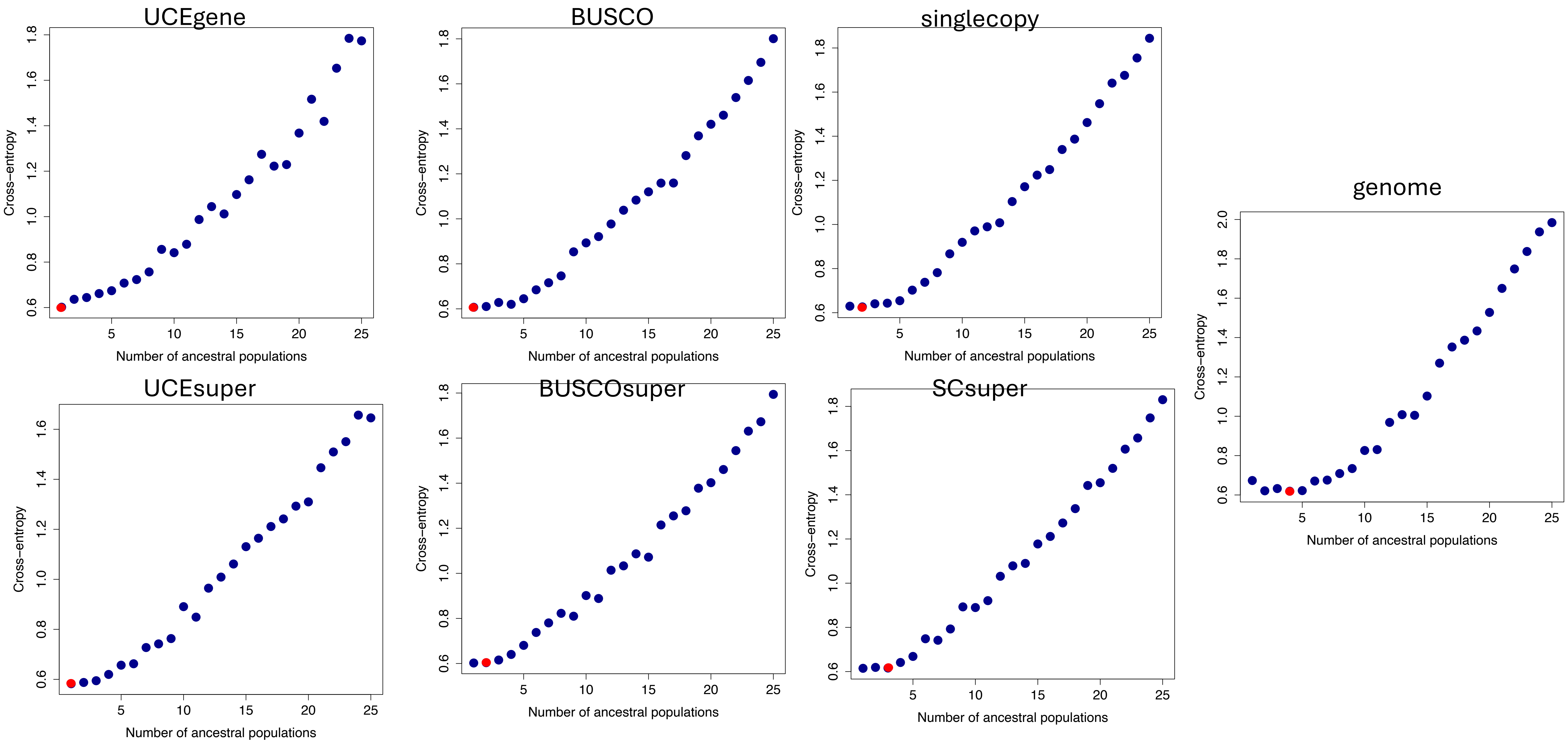
UCEgene
BUSCO
singlecopy
genome
UCEsuper
BUSCOsuper
SCsuper
Cross−entropy
Number of ancestral populations

*Coix*

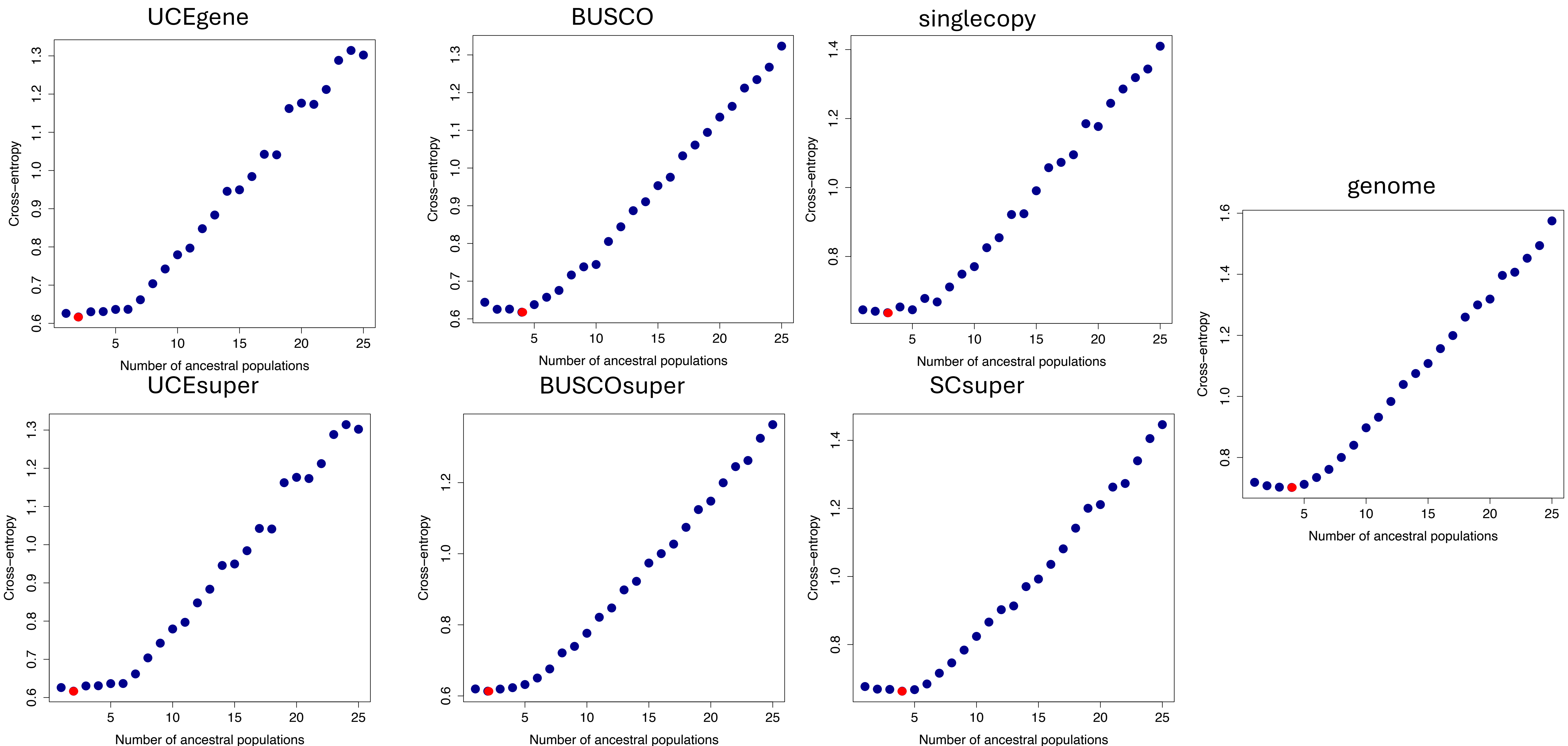
UCEgene
BUSCO
singlecopy
genome
UCEsuper
BUSCOsuper
SCsuper
Cross−entropy
Number of ancestral populations

Corylus

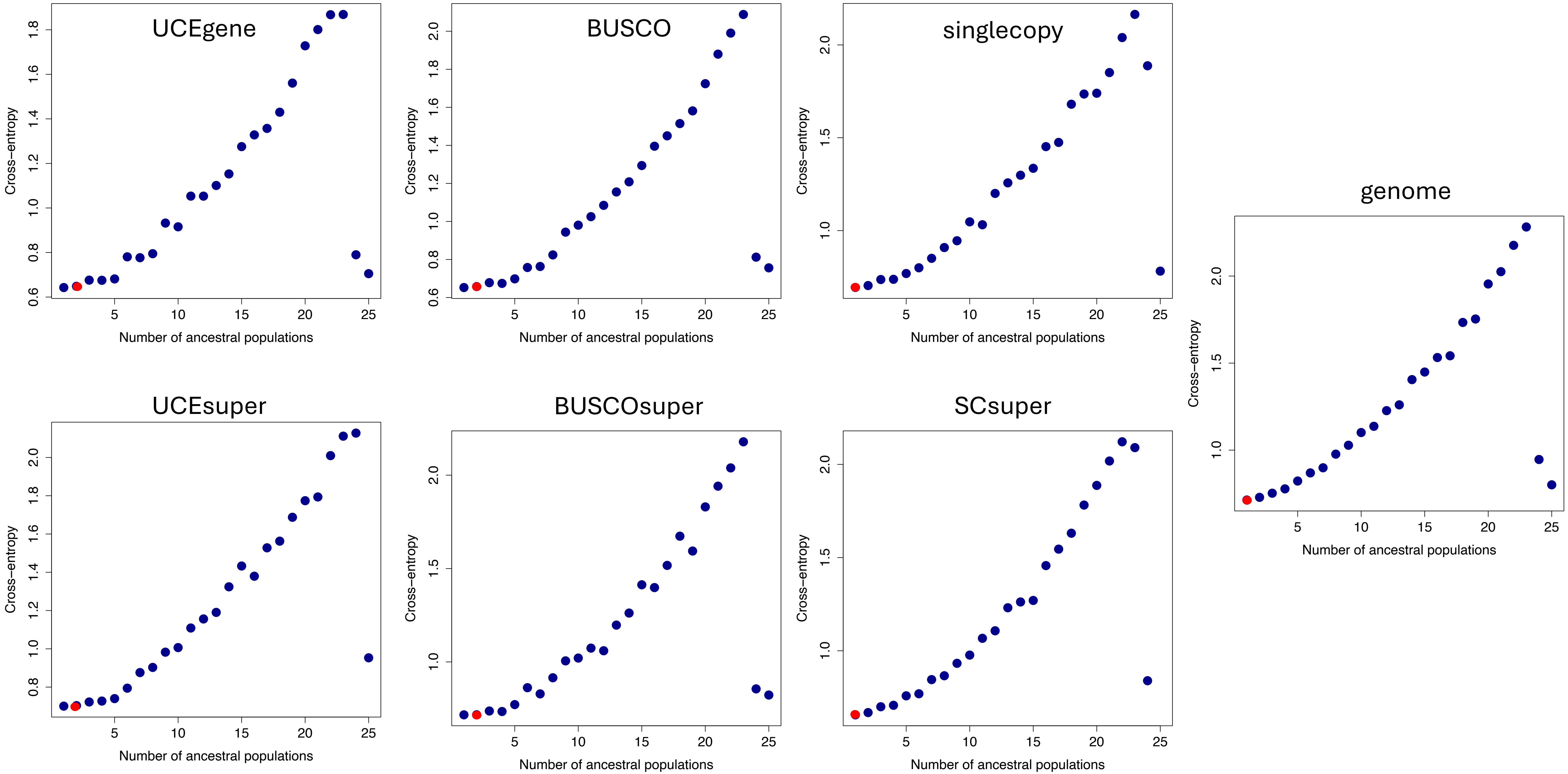
UCEgene
BUSCO
singlecopy
genome
UCEsuper
BUSCOsuper
SCsuper
Cross–entropy
Number of ancestral populations

*Costus*

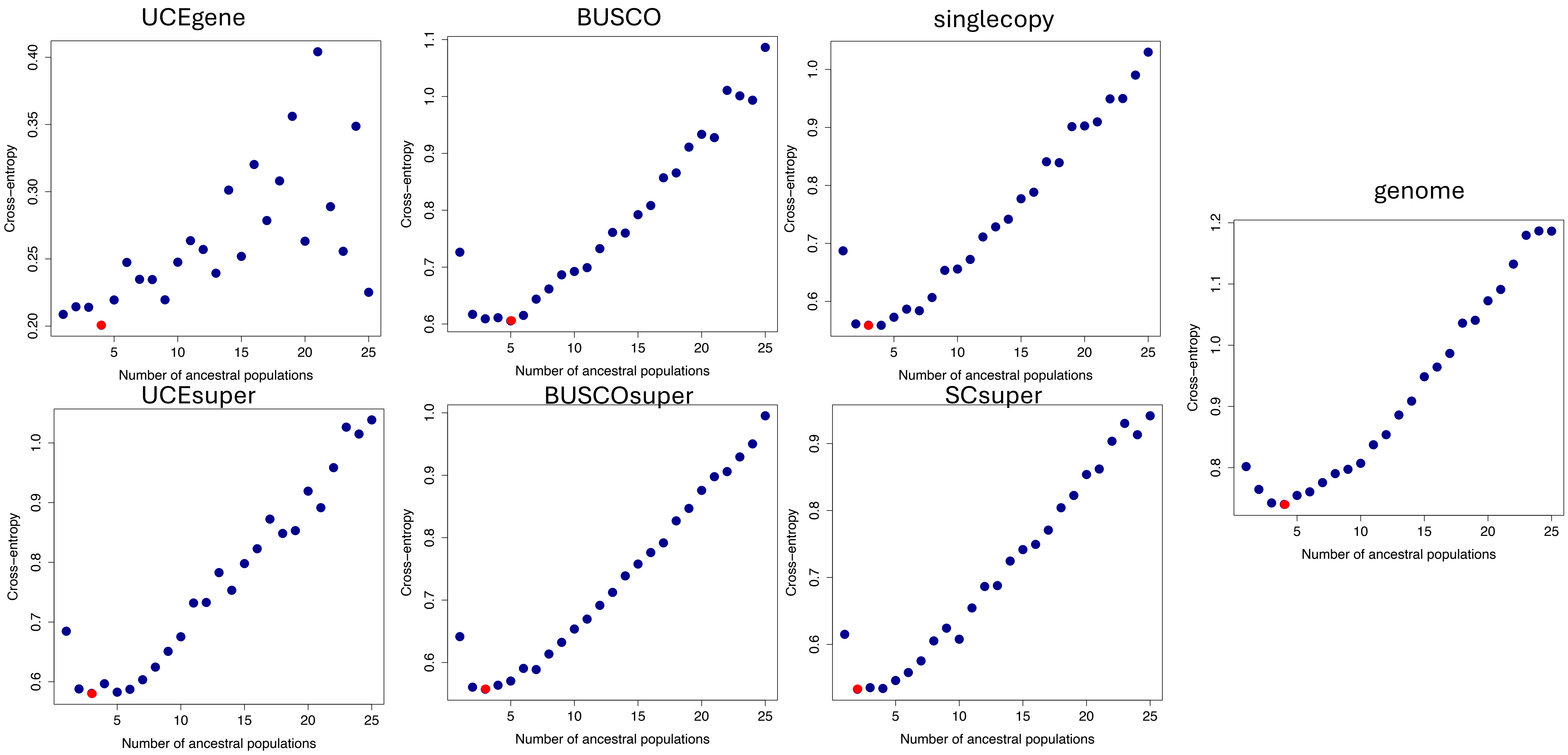

UCEgene
BUSCO
singlecopy
genome
UCEsuper
BUSCOsuper
SCsuper
Cross−entropy
Number of ancestral populations


*Diperion*

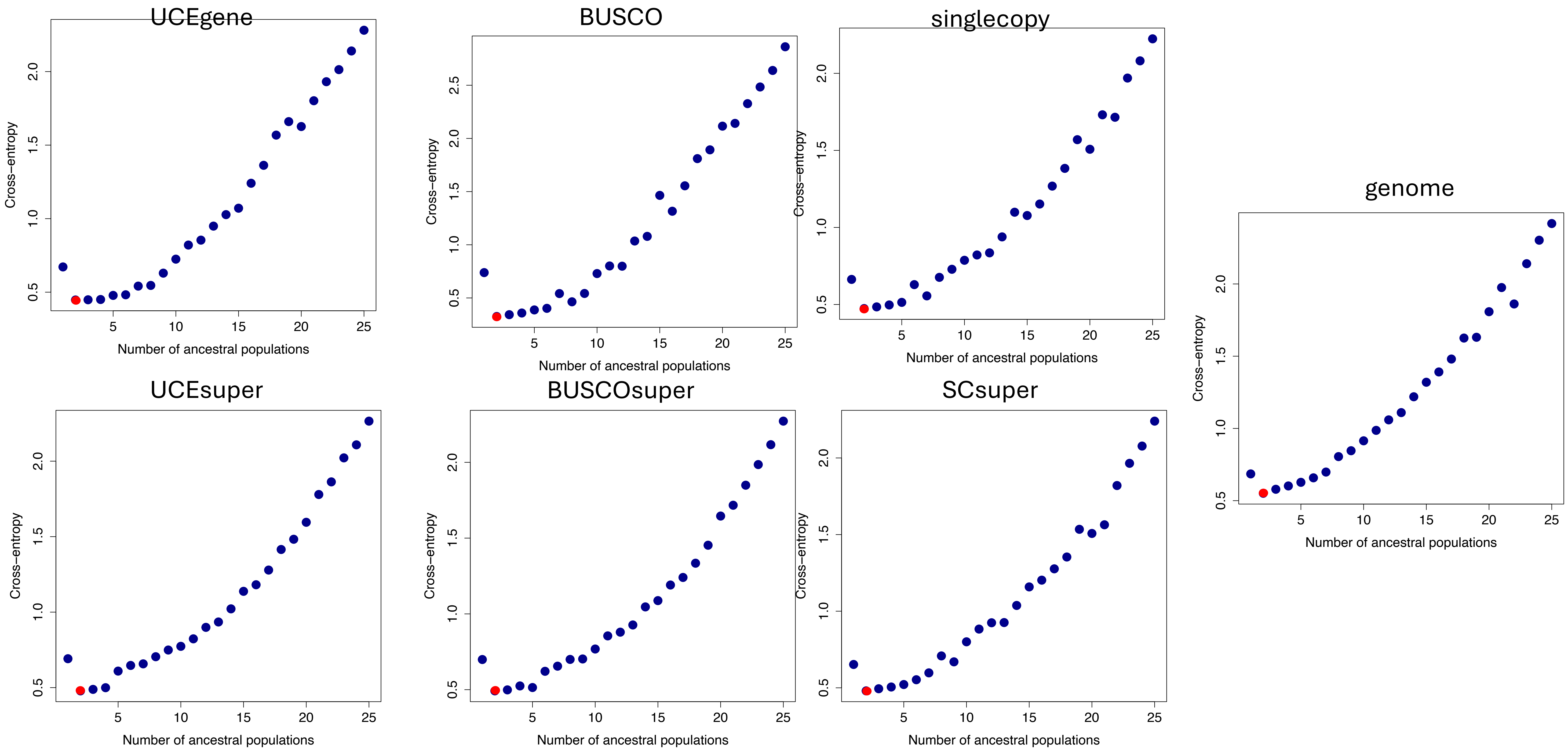
UCEgene
BUSCO
singlecopy
genome
UCEsuper
BUSCOsuper
SCsuper
Cross−entropy
Number of ancestral populations

*Fragaria*

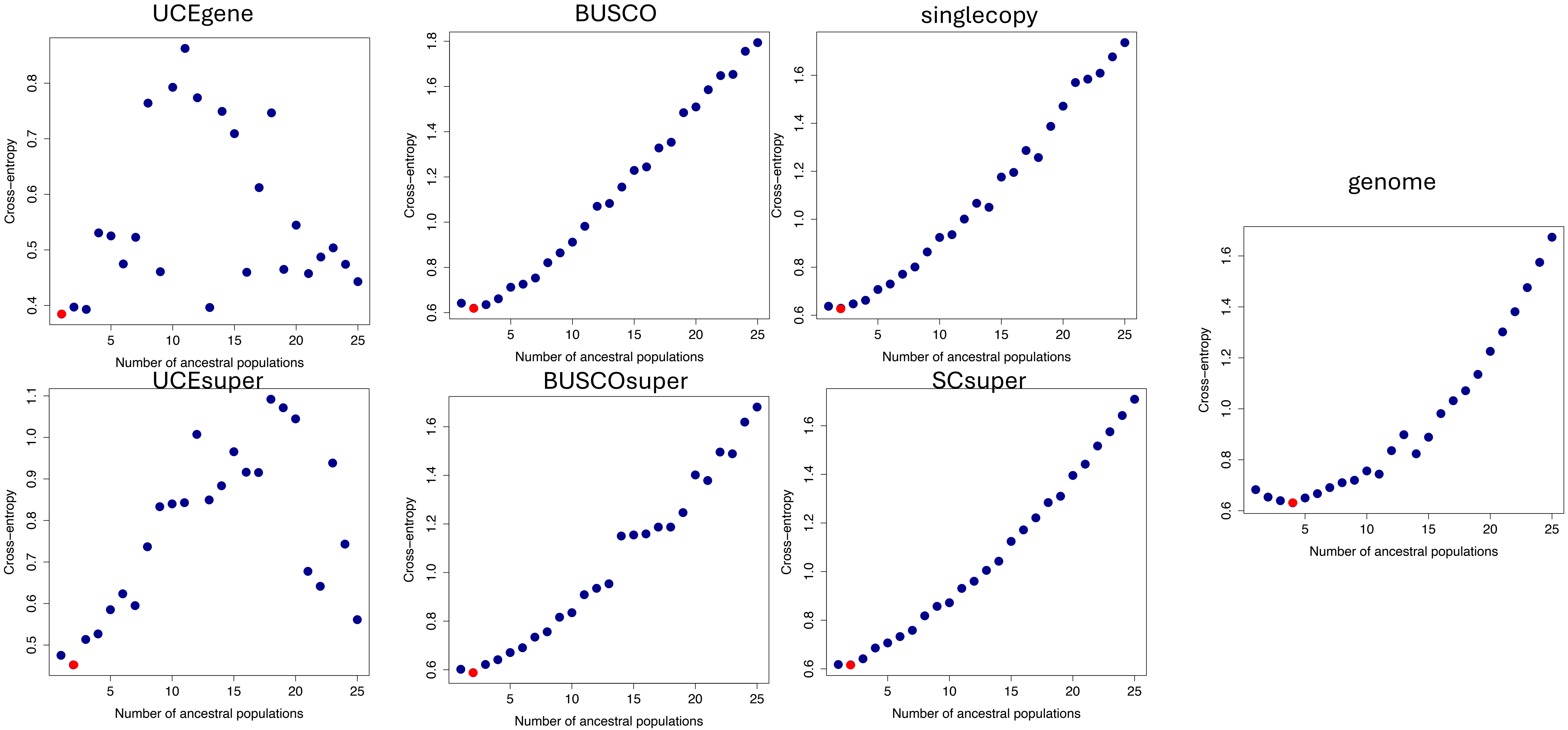
UCEgene
BUSCO
singlecopy
genome
UCEsuper
BUSCOsuper
SCsuper
Cross−entropy
Number of ancestral populations

*Geospiza*

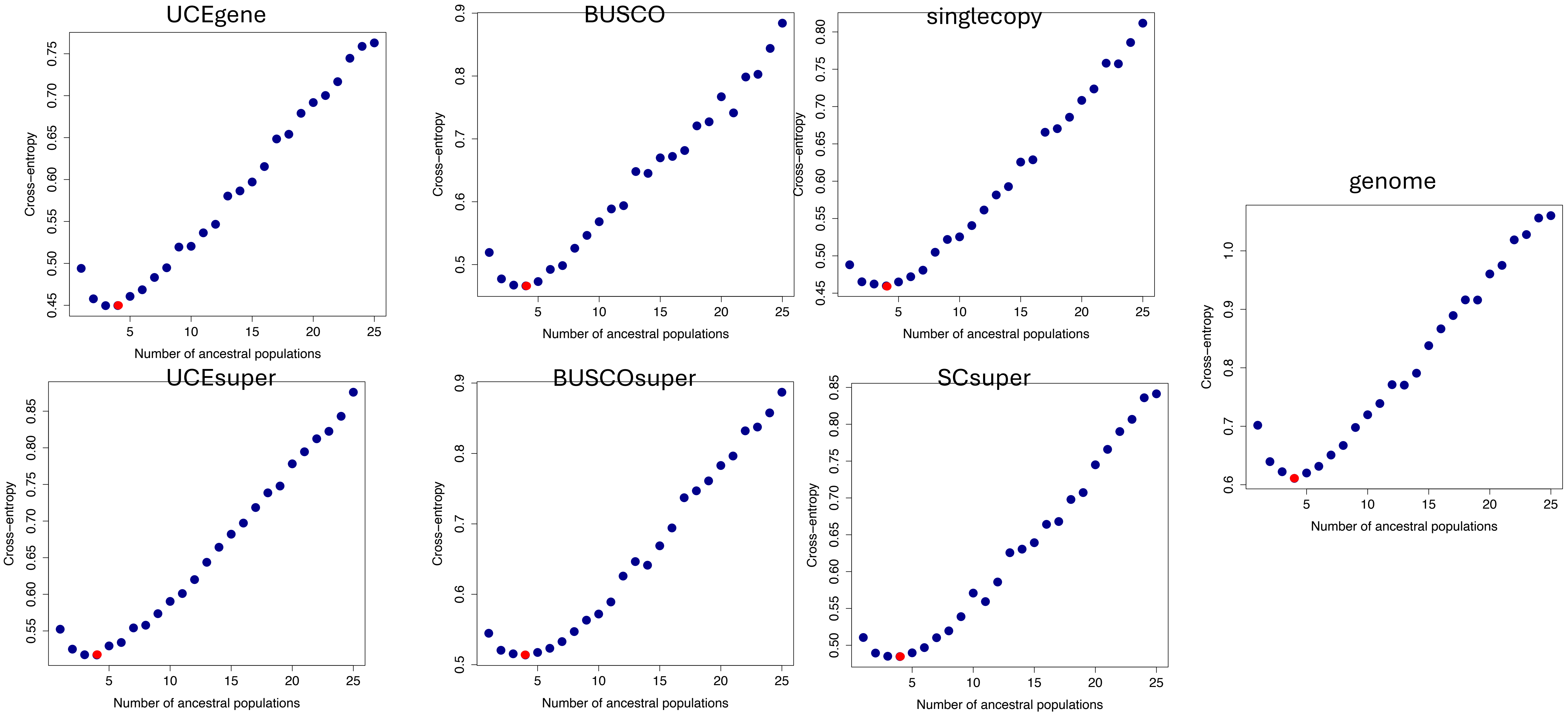
UCEgene
BUSCO
singlecopy
genome
UCEsuper
BUSCOsuper
SCsuper
Cross−entropy
Number of ancestral populations

*Juglans*

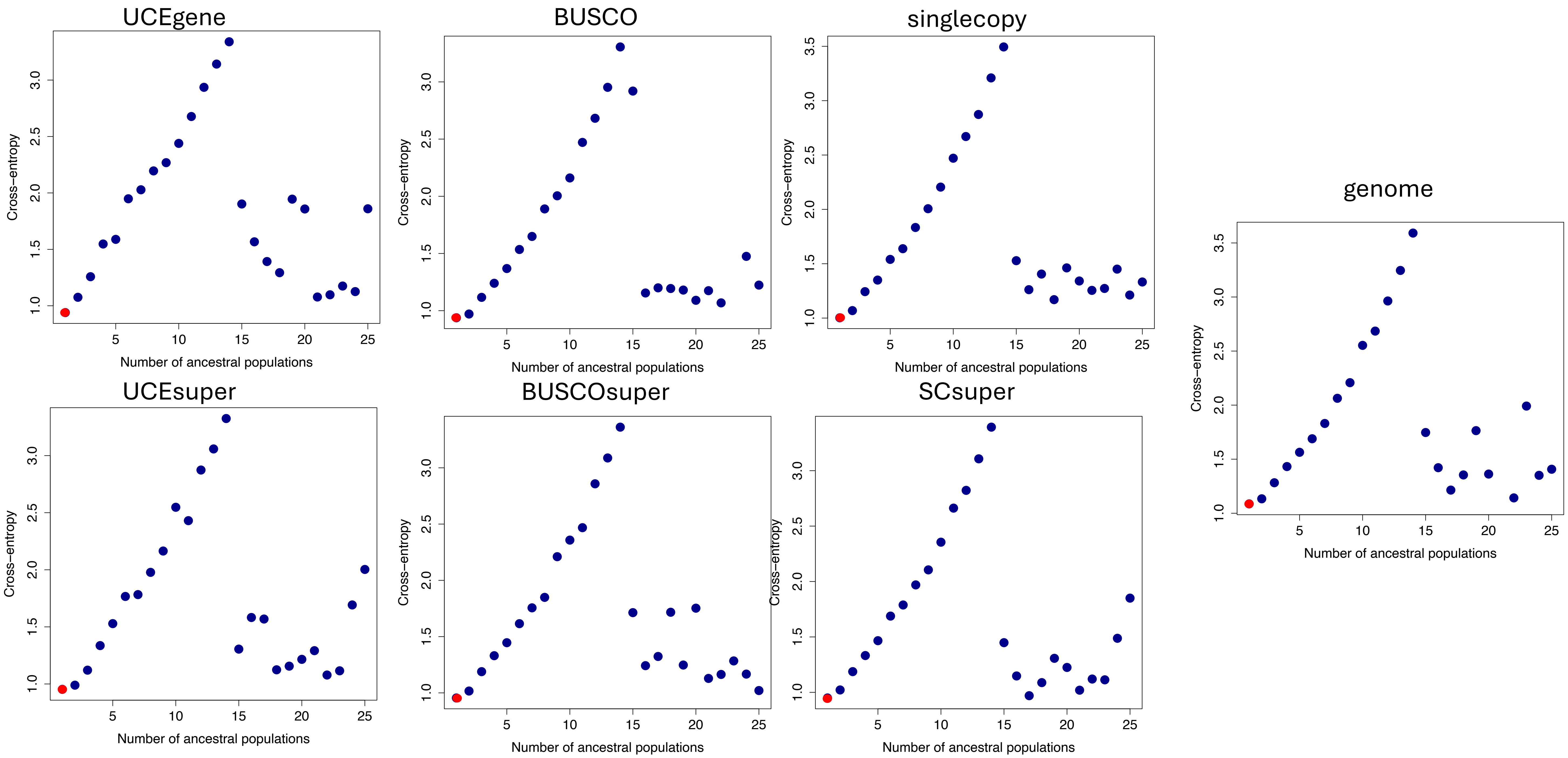


*Liriodendron*

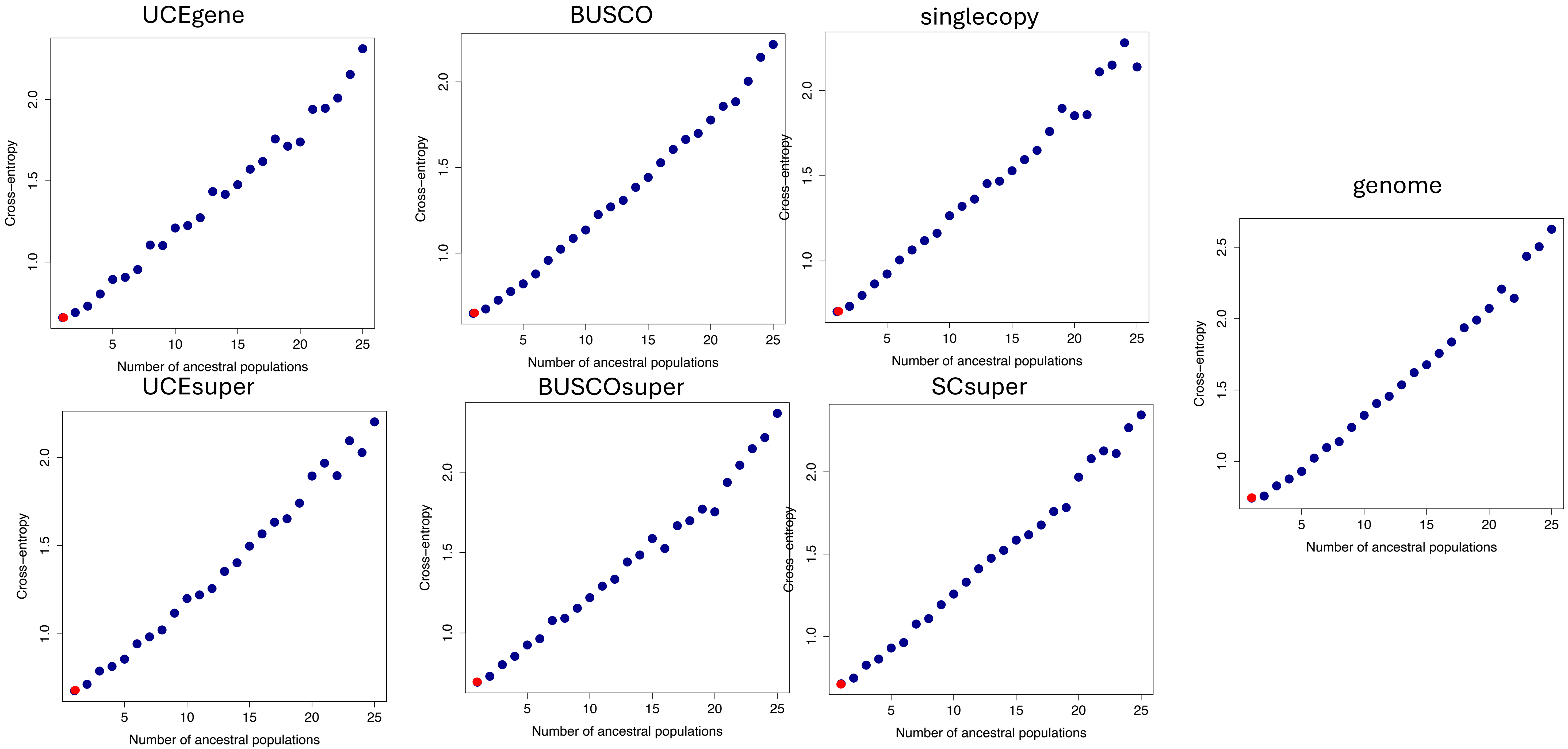

UCEgene
BUSCO
singlecopy
genome
UCEsuper
BUSCOsuper
SCsuper
Cross−entropy
Number of ancestral populations
1.0
1.5
2.0
2.5
5
10
15
20
25


*Lotus*

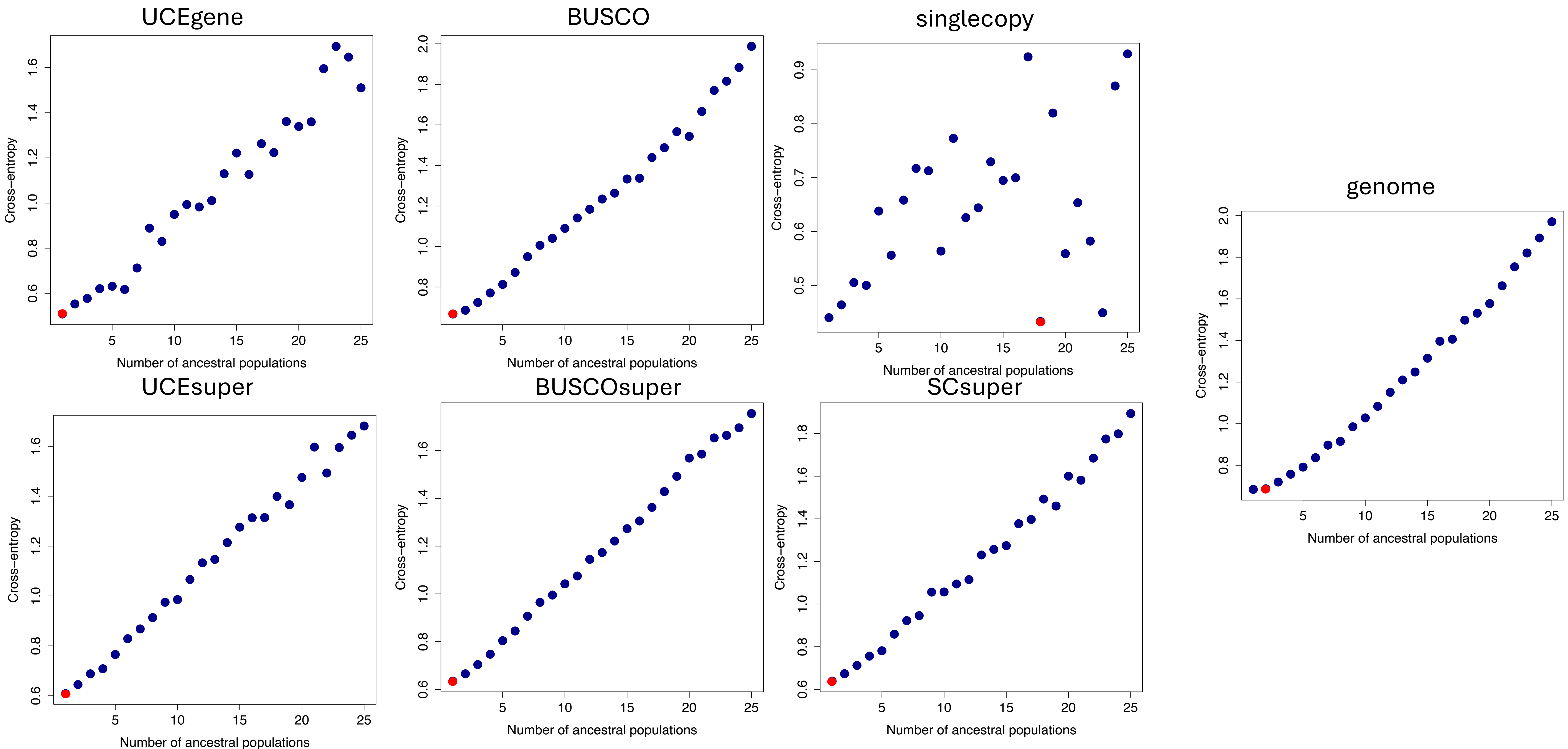
UCEgene
BUSCO
singlecopy
genome
UCEsuper
BUSCOsuper
SCsuper
Cross–entropy
Number of ancestral populations

*Malania*

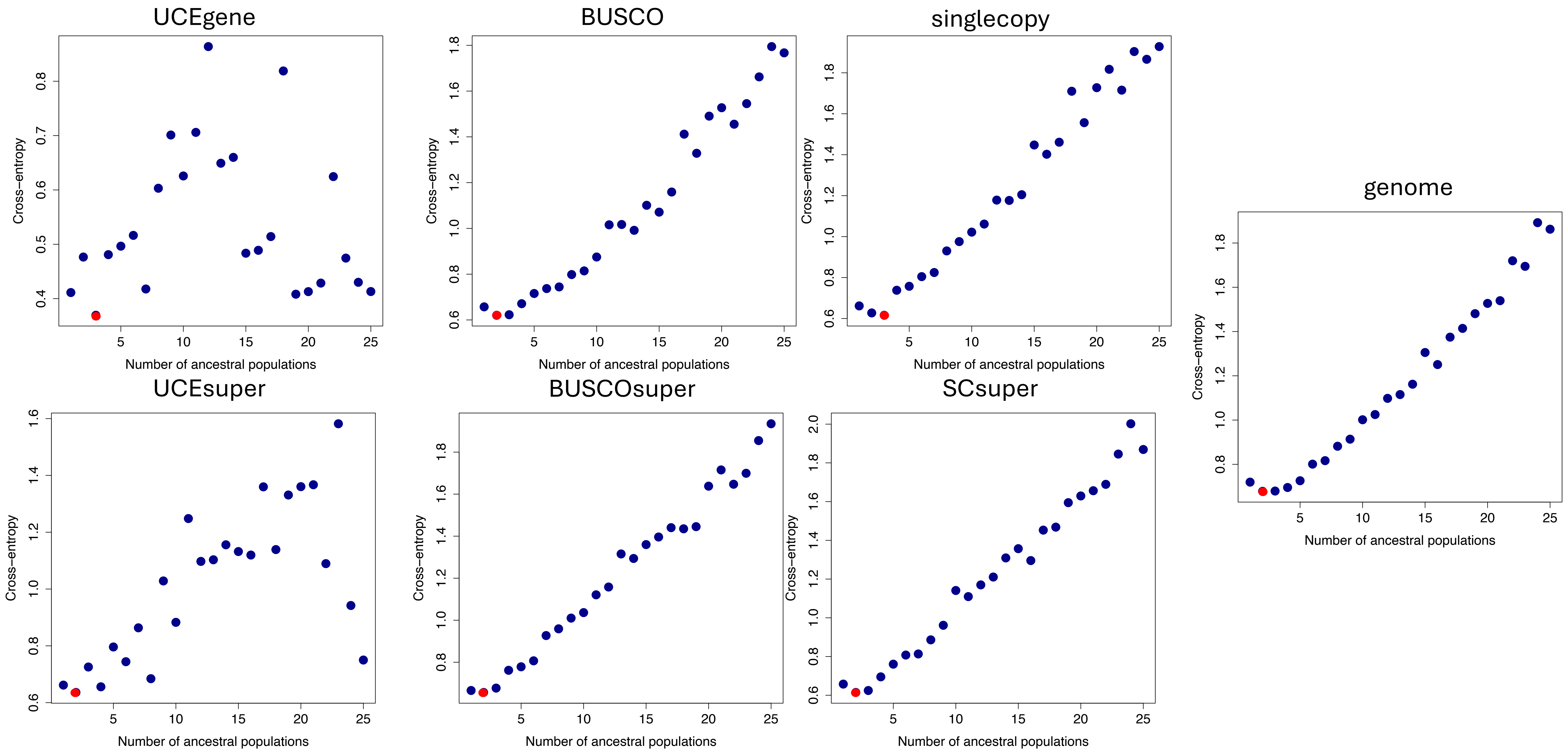
UCEgene
BUSCO
singlecopy
genome
UCEsuper
BUSCOsuper
SCsuper
Cross–entropy
Number of ancestral populations

*Medicago*

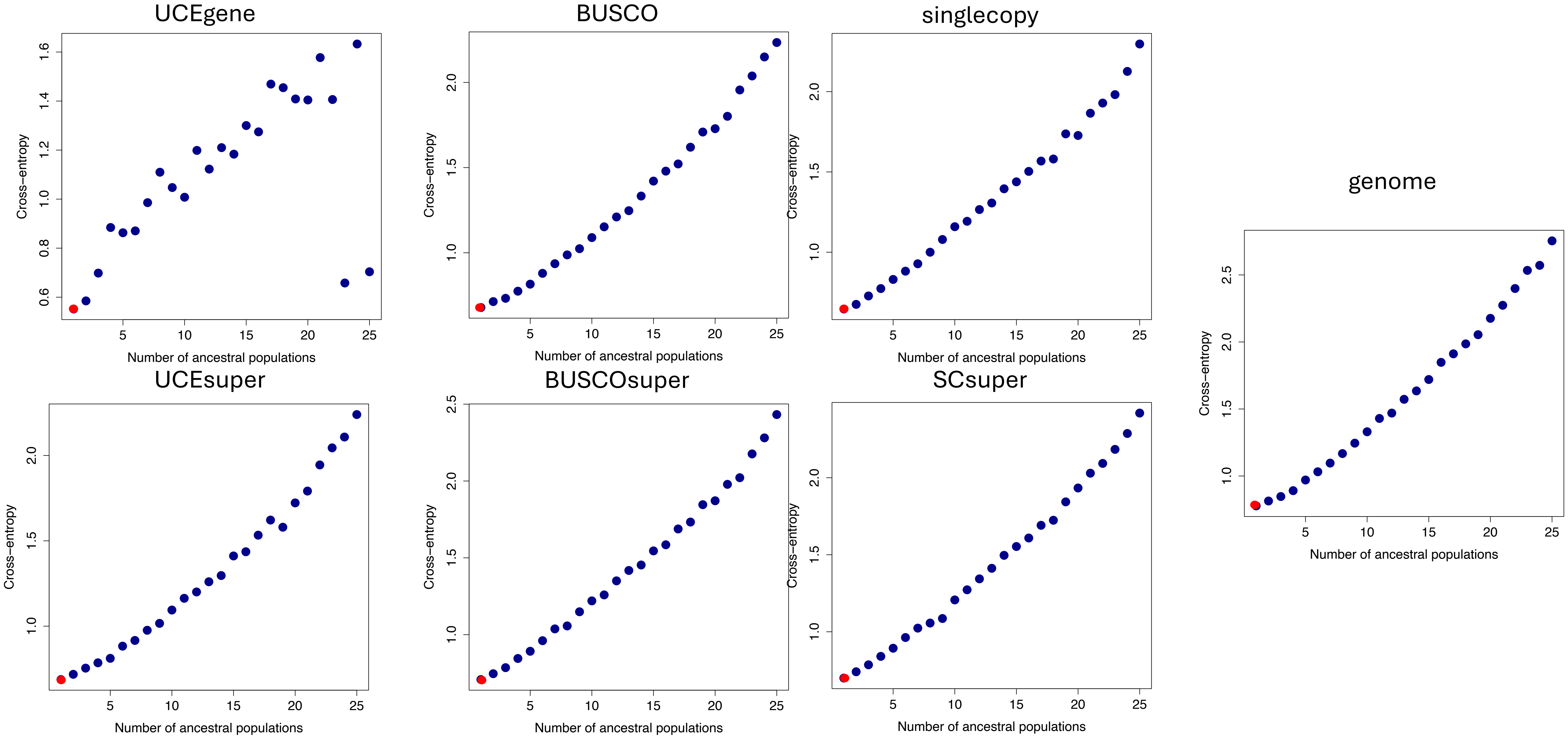


*Plectropomus*

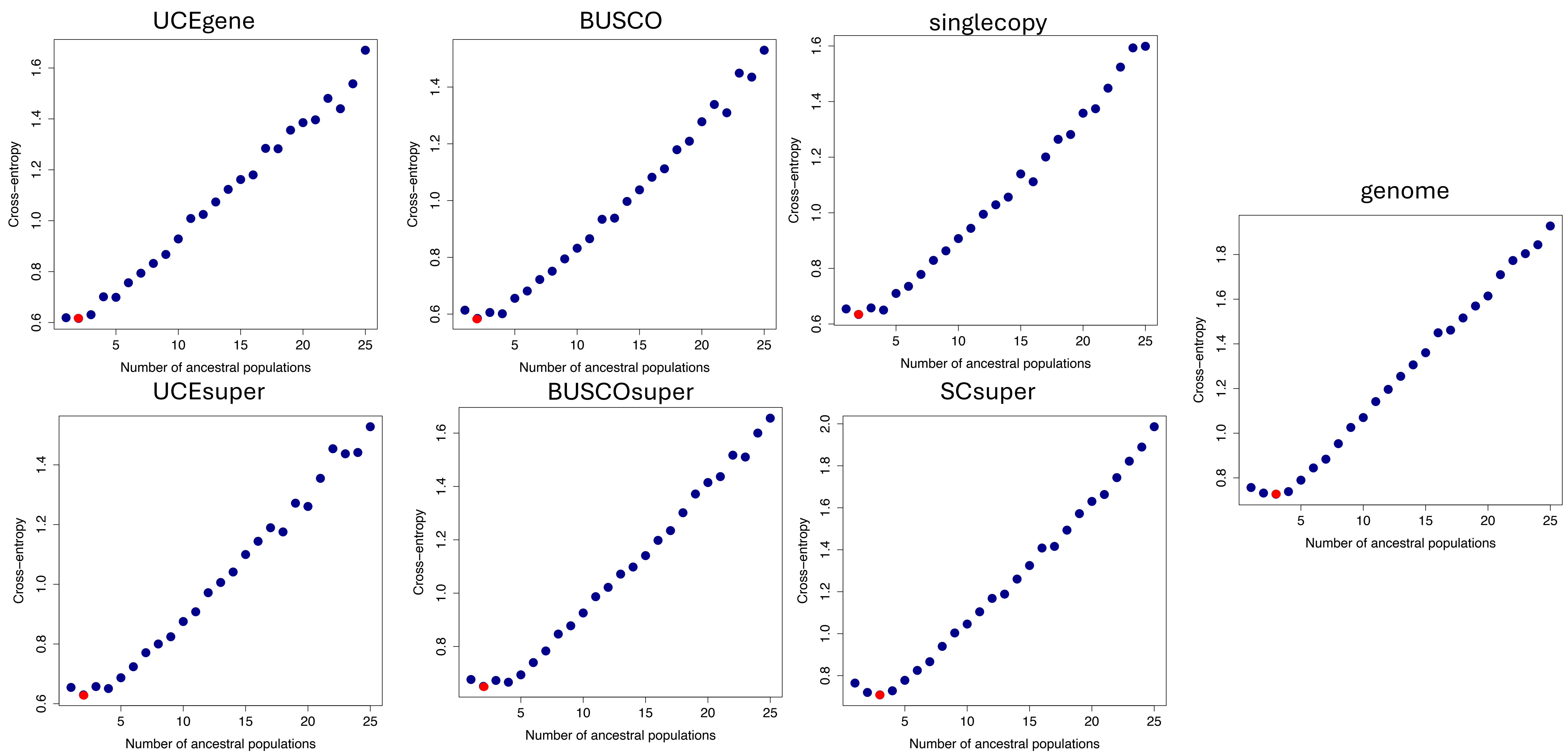
UCEgene
BUSCO
singlecopy
genome
UCEsuper
BUSCOsuper
SCsuper
Cross−entropy
Number of ancestral populations

*Populus*

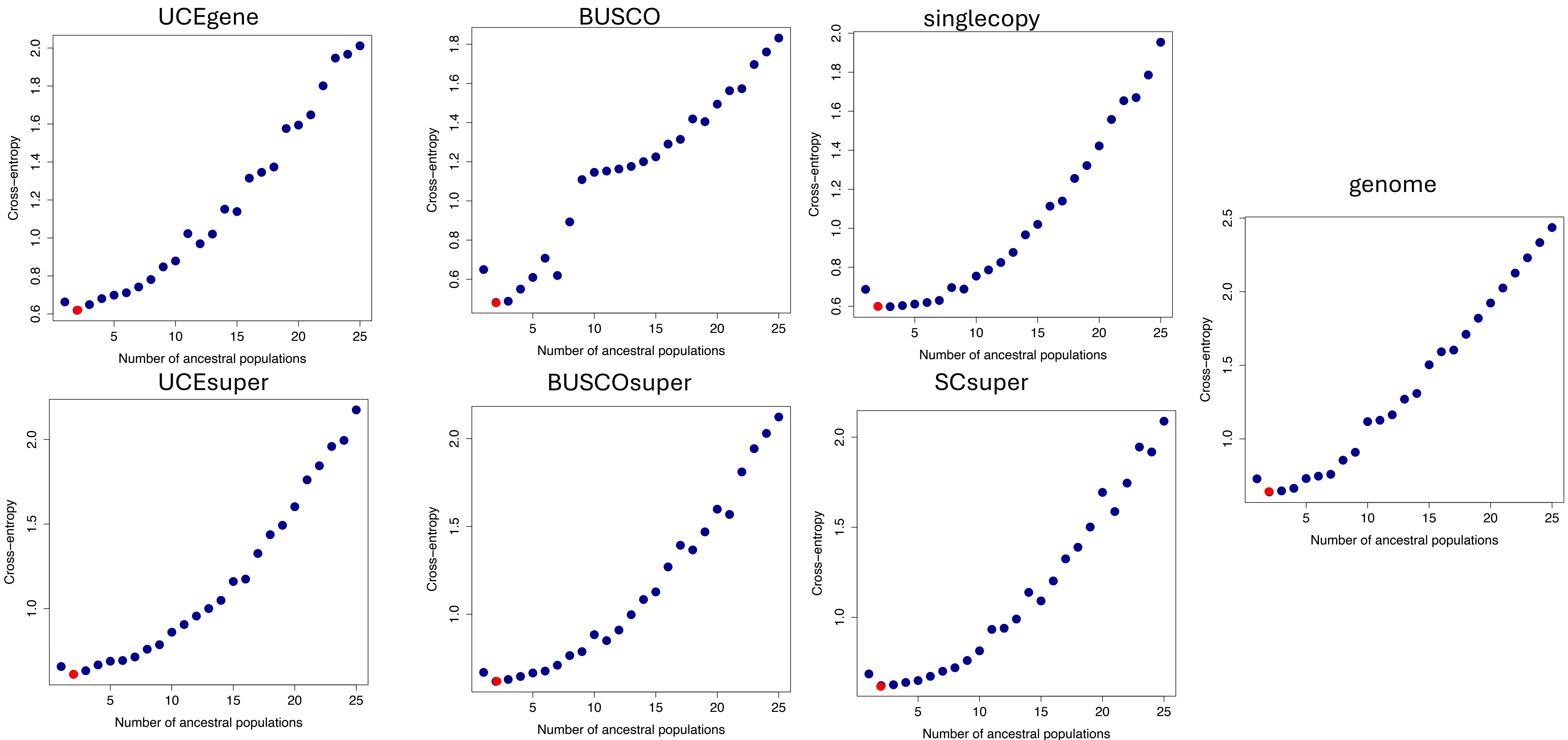
UCEgene
BUSCO
singlecopy
genome
UCEsuper
BUSCOsuper
SCsuper
Cross−entropy
Number of ancestral populations


*Primula*

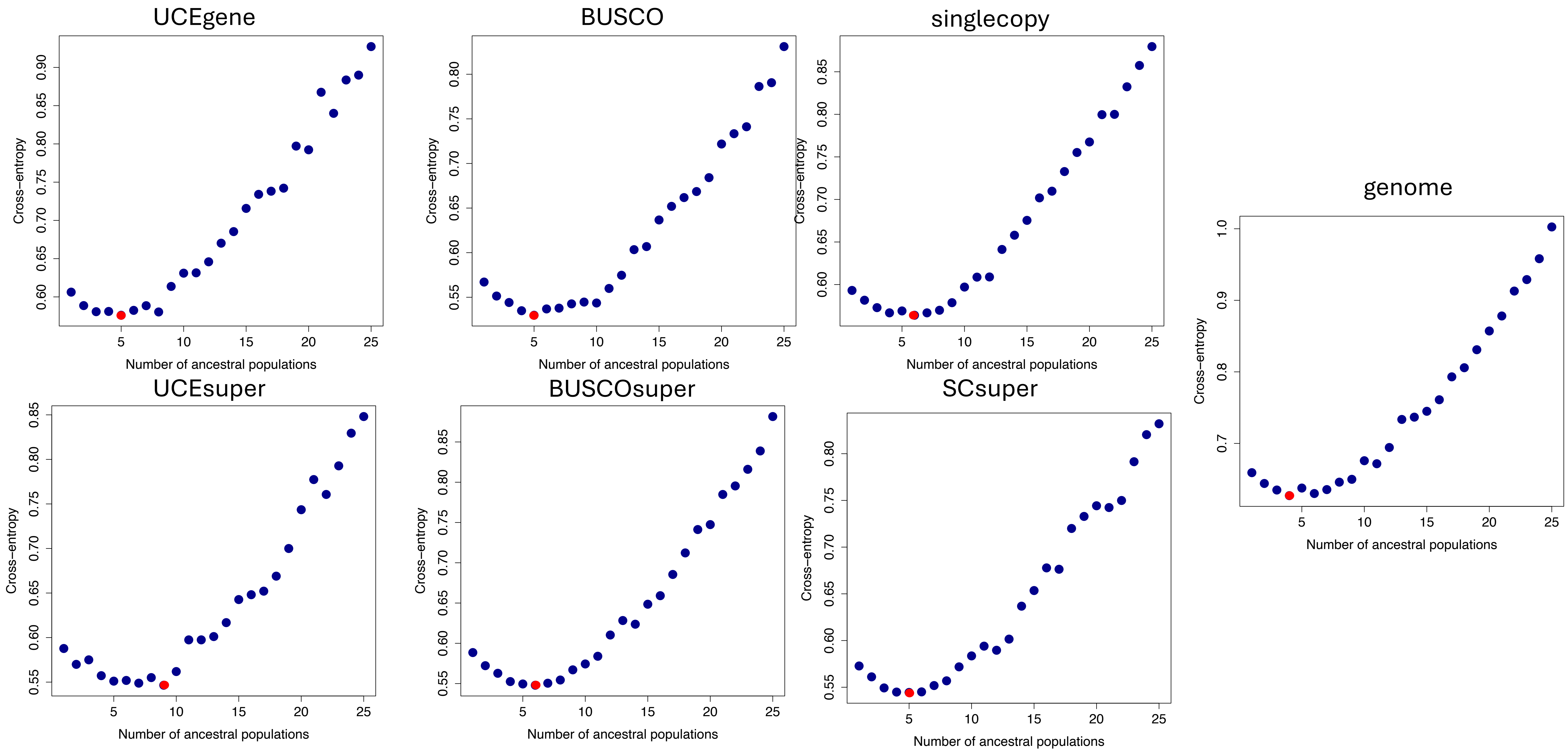
UCEgene
BUSCO
singlecopy
genome
UCEsuper
BUSCOsuper
SCsuper
Cross−entropy
Number of ancestral populations

*Prunus*

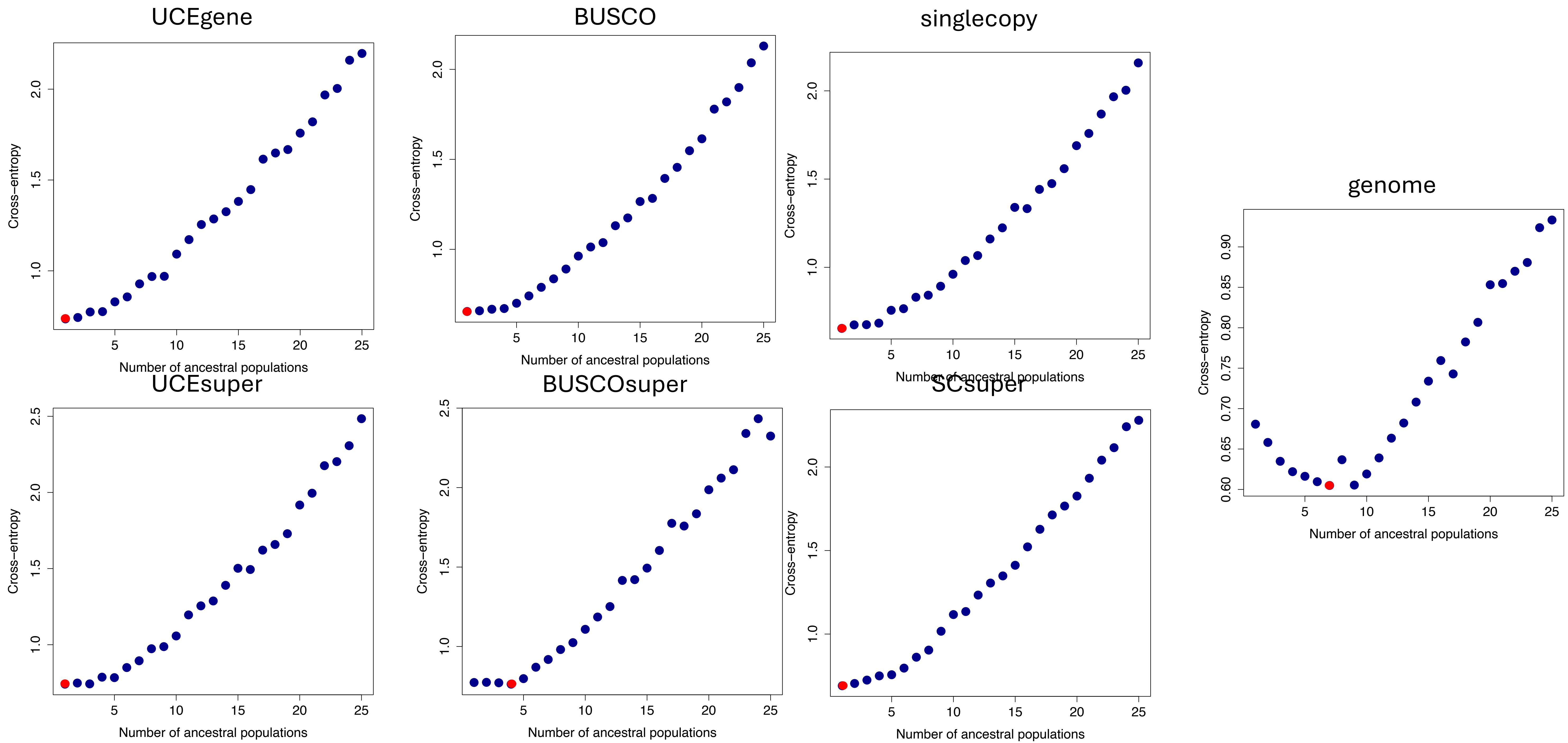
UCEgene
BUSCO
singlecopy
genome
UCEsuper
BUSCOsuper
SCsuper
Cross−entropy
Number of ancestral populations

*Salix*

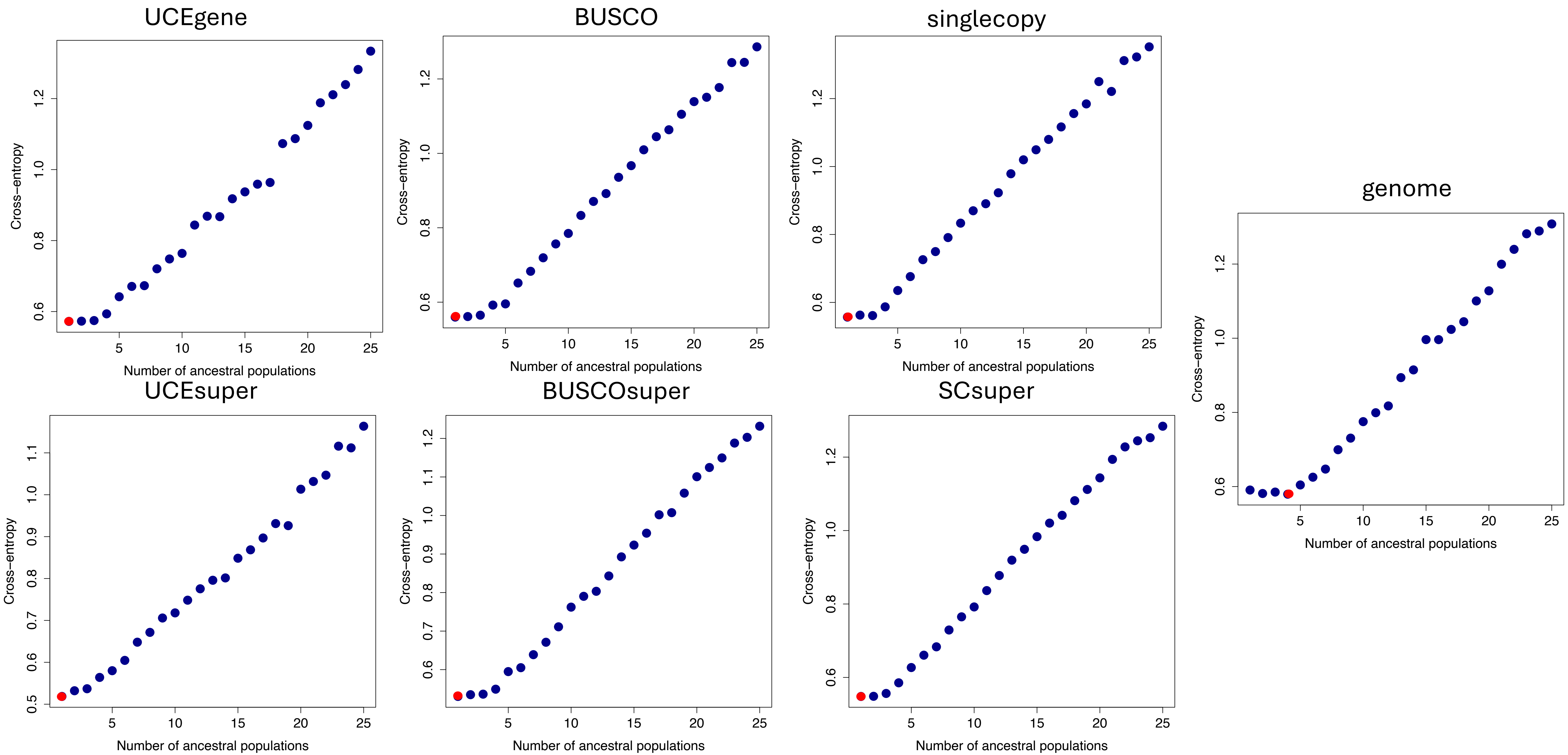
UCEgene
BUSCO
singlecopy
genome
UCEsuper
BUSCOsuper
SCsuper
Cross−entropy
Number of ancestral populations

*Seseamum*

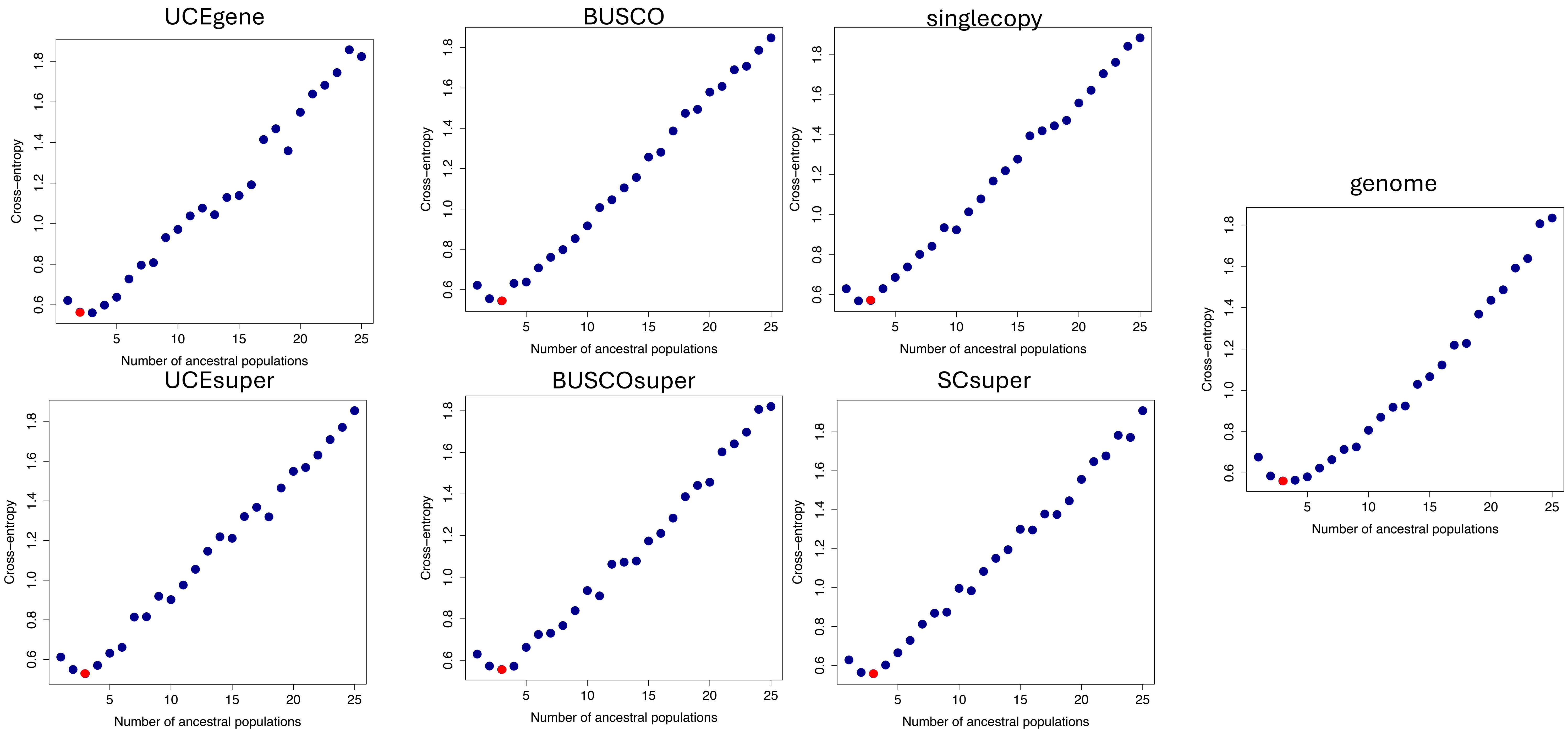
UCEgene
BUSCO
singlecopy
genome
UCEsuper
BUSCOsuper
SCsuper
Cross−entropy
Number of ancestral populations
0.6
0.8
1.0
1.2
1.4
1.6
1.8
5
10
15
20
25

*Solanum*

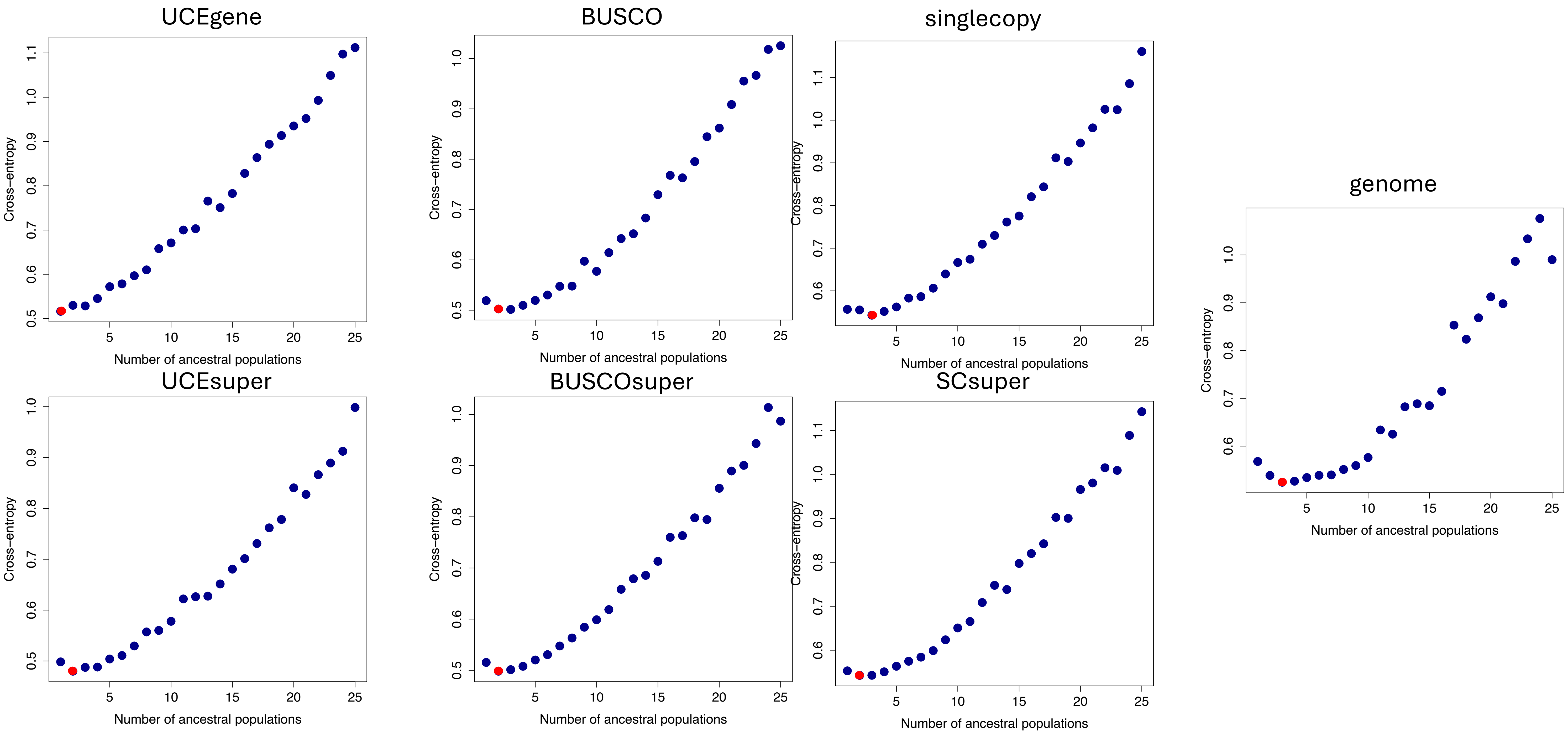
UCEgene
BUSCO
singlecopy
genome
UCEsuper
BUSCOsuper
SCsuper
Cross−entropy
Number of ancestral populations

*Sorghum*

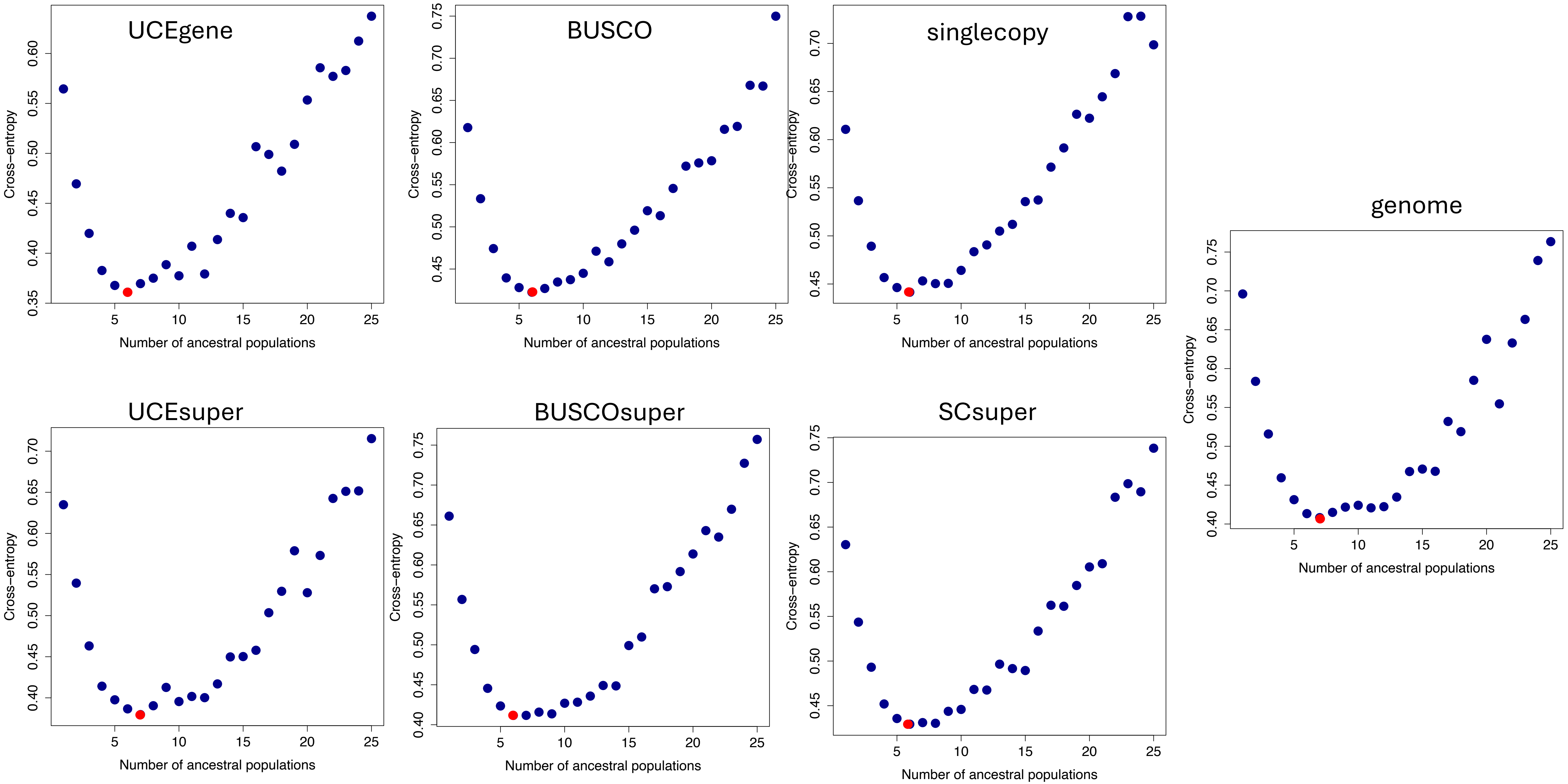


*Spirodela polyrhiza*

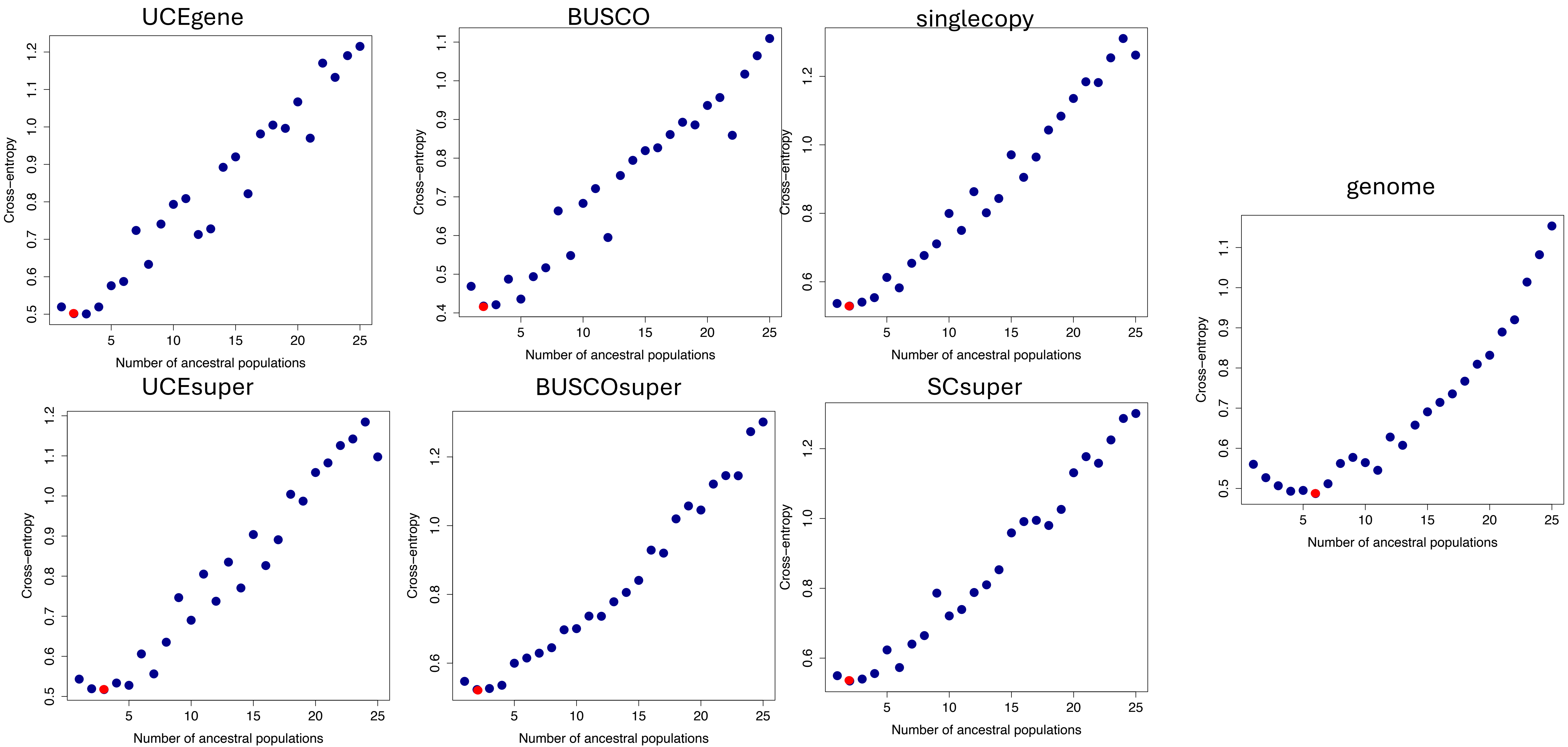
UCEgene
BUSCO
singlecopy
genome
UCEsuper
BUSCOsuper
SCsuper
Cross−entropy
Number of ancestral populations

*Thlaspi*

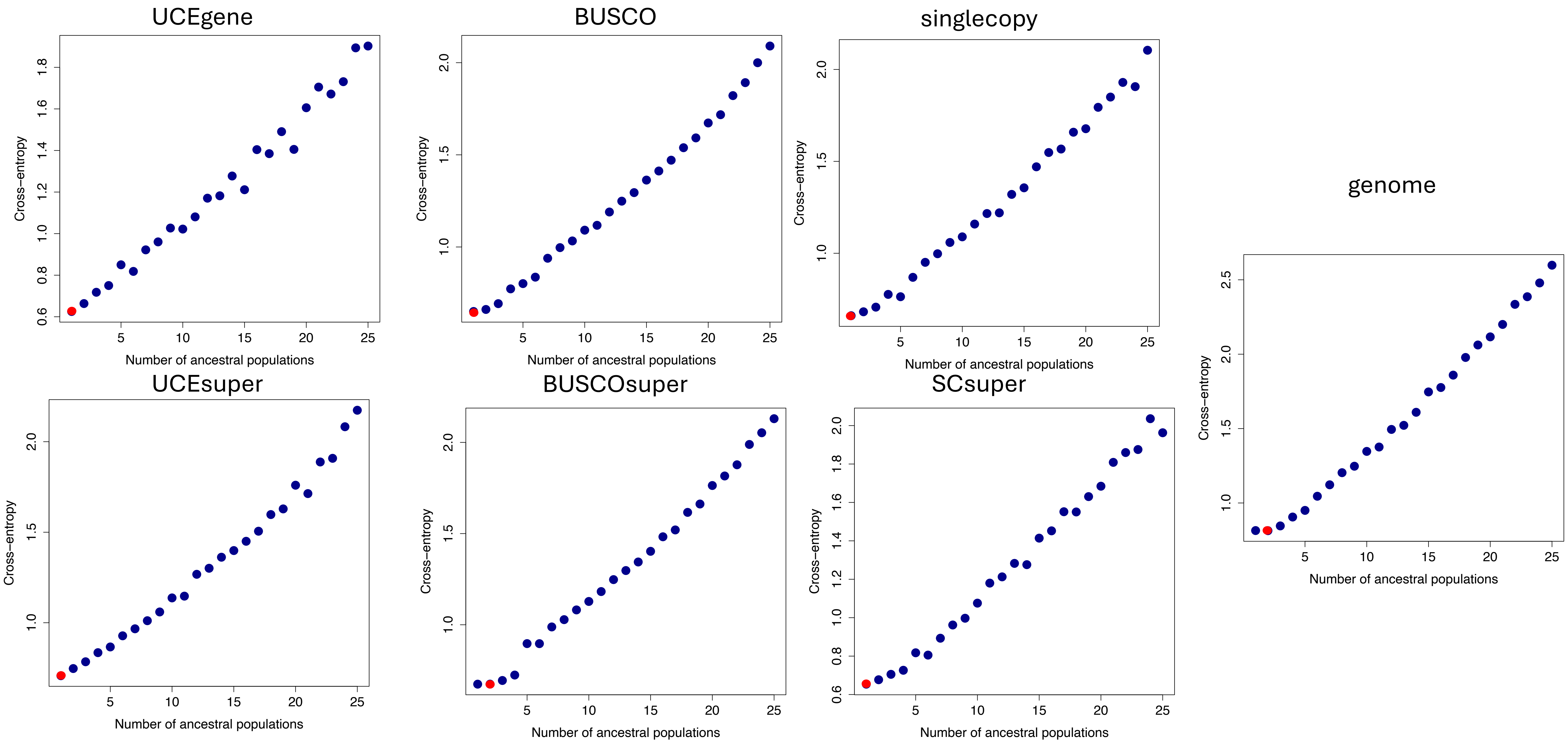
UCEgene
BUSCO
singlecopy
genome
UCEsuper
BUSCOsuper
SCsuper
Cross−entropy
Number of ancestral populations

*Ursus*

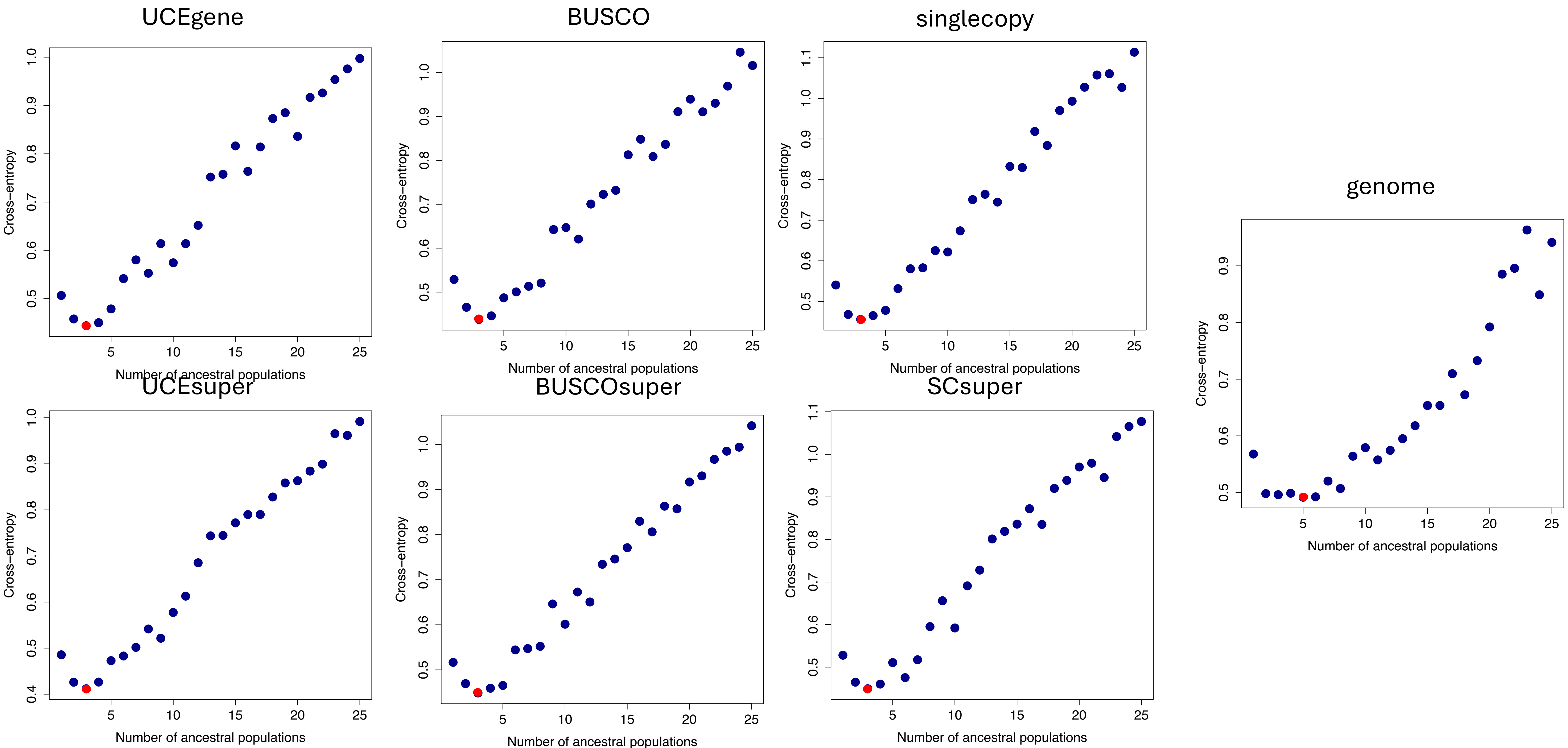

UCEgene
BUSCO
singlecopy
genome
UCEsuper
BUSCOsuper
SCsuper
Cross−entropy
Number of ancestral populations


*Vigna*

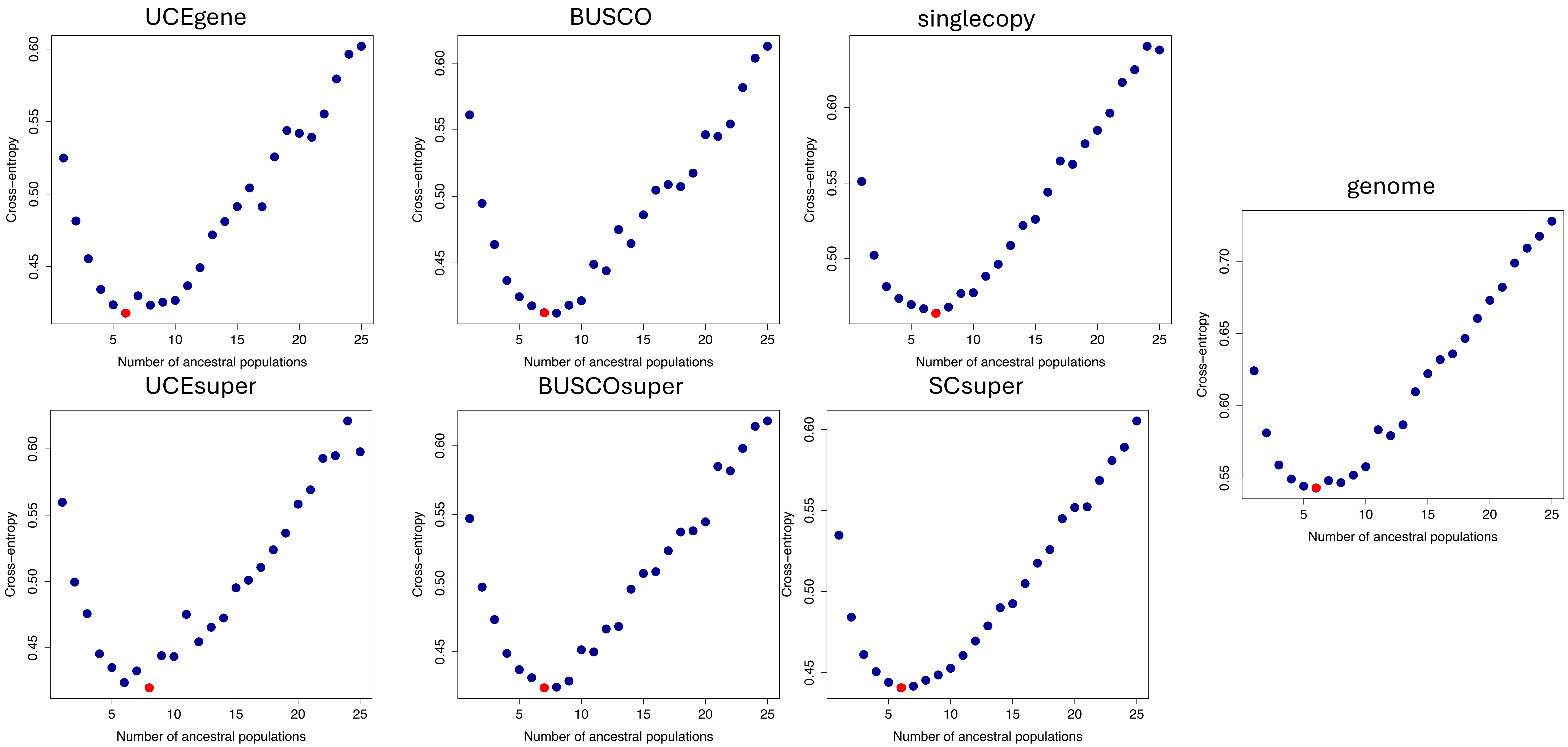

UCEgene
BUSCO
singlecopy
genome
UCEsuper
BUSCOsuper
SCsuper
Cross–entropy
Number of ancestral populations


*Vitis*

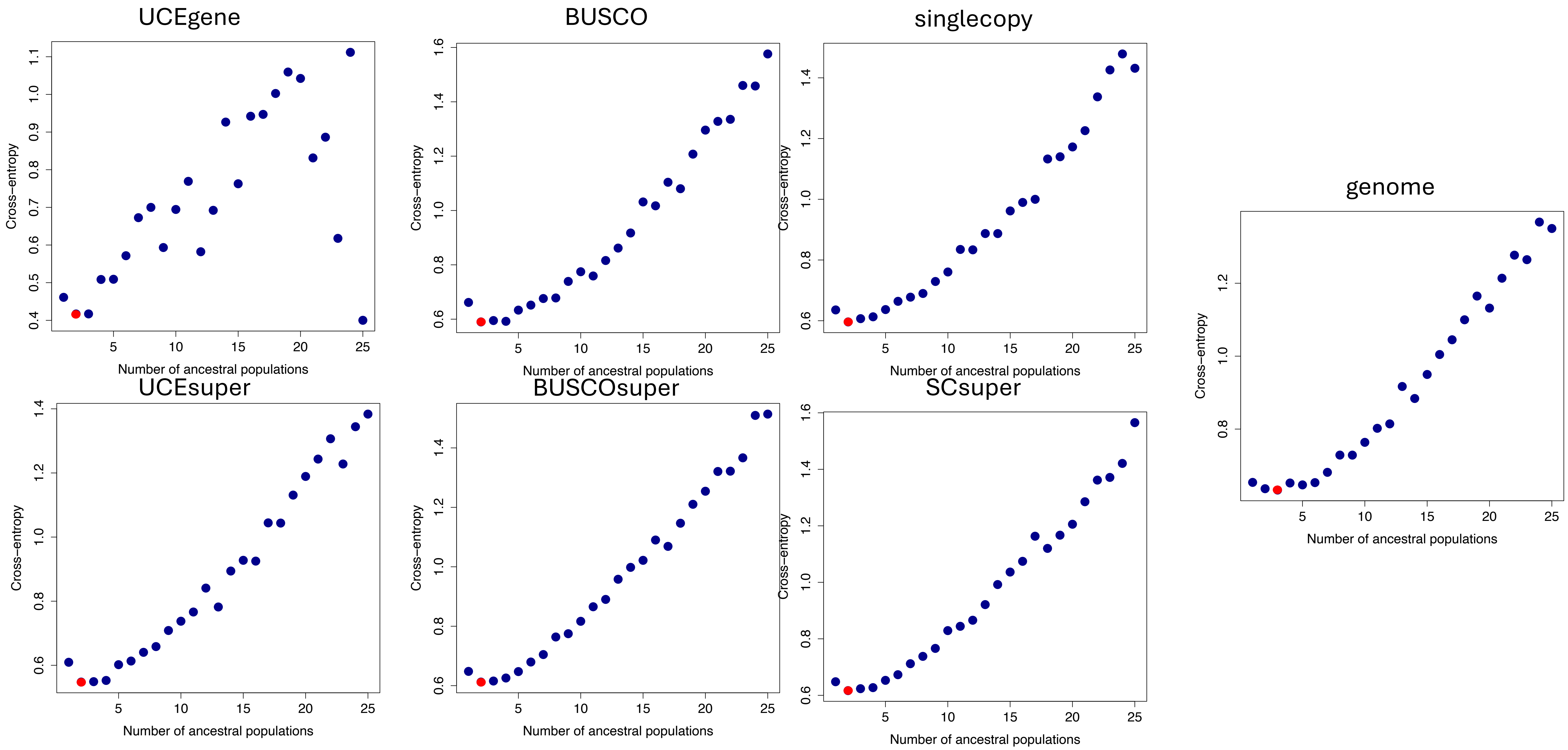
UCEgene
BUSCO
singlecopy
genome
UCEsuper
BUSCOsuper
SCsuper
Cross−entropy
Number of ancestral populations
5
10
15
20
25
0.4
0.5
0.6
0.7
0.8
0.9
1.0
1.1
1.2
1.4
1.6

Warblers
