## Supplemental Figure 7 for "The power to resolve relationships: identifying incongruence and precision of reduced representation and genome-wide data in phylogenomics and population genomics"

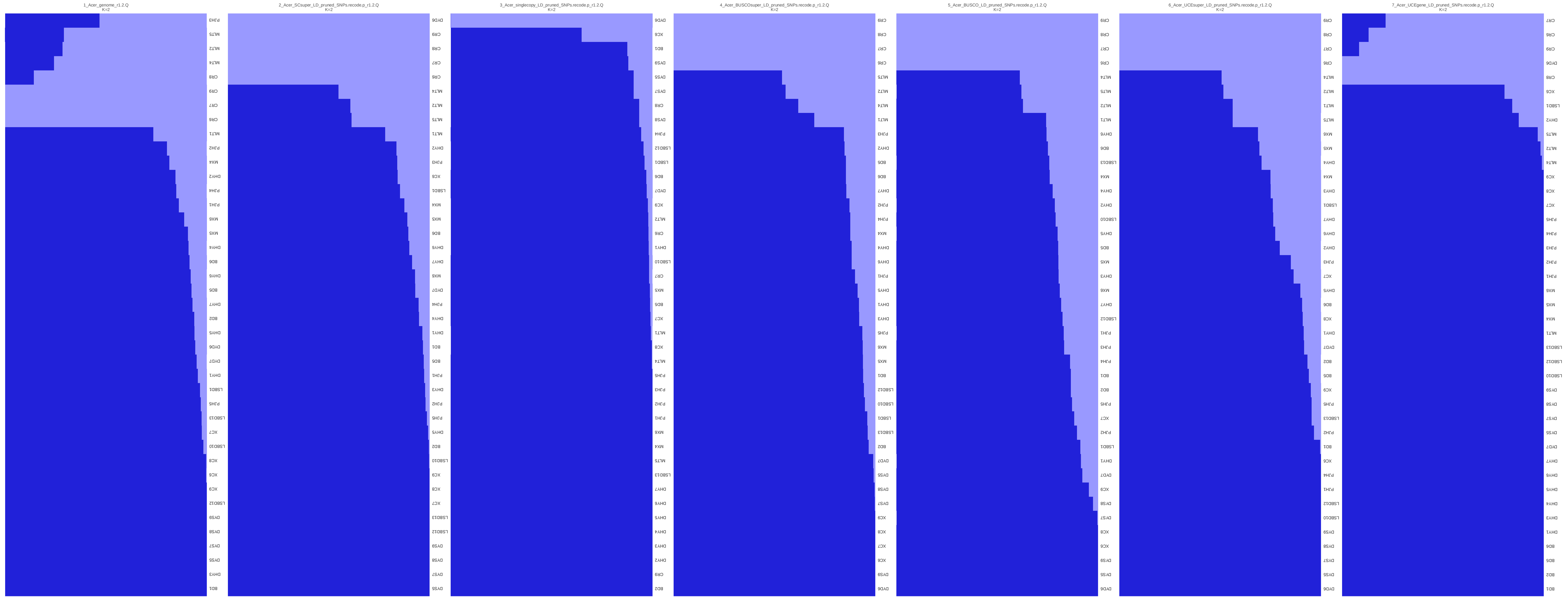
1_Acer_genome_r1.2.Q
K=2
2_Acer_SCsuper_LD_pruned_SNPs.recode.p_r1.2.Q
K=2
3_Acer_singlecopy_LD_pruned_SNPs.recode.p_r1.2.Q
K=2
4_Acer_BUSCOsuper_LD_pruned_SNPs.recode.p_r1.2.Q
K=2
5_Acer_BUSCO_LD_pruned_SNPs.recode.p_r1.2.Q
K=2
6_Acer_UCEsuper_LD_pruned_SNPs.recode.p_r1.2.Q
K=2
7_Acer_UCEgene_LD_pruned_SNPs.recode.p_r1.2.Q
K=2

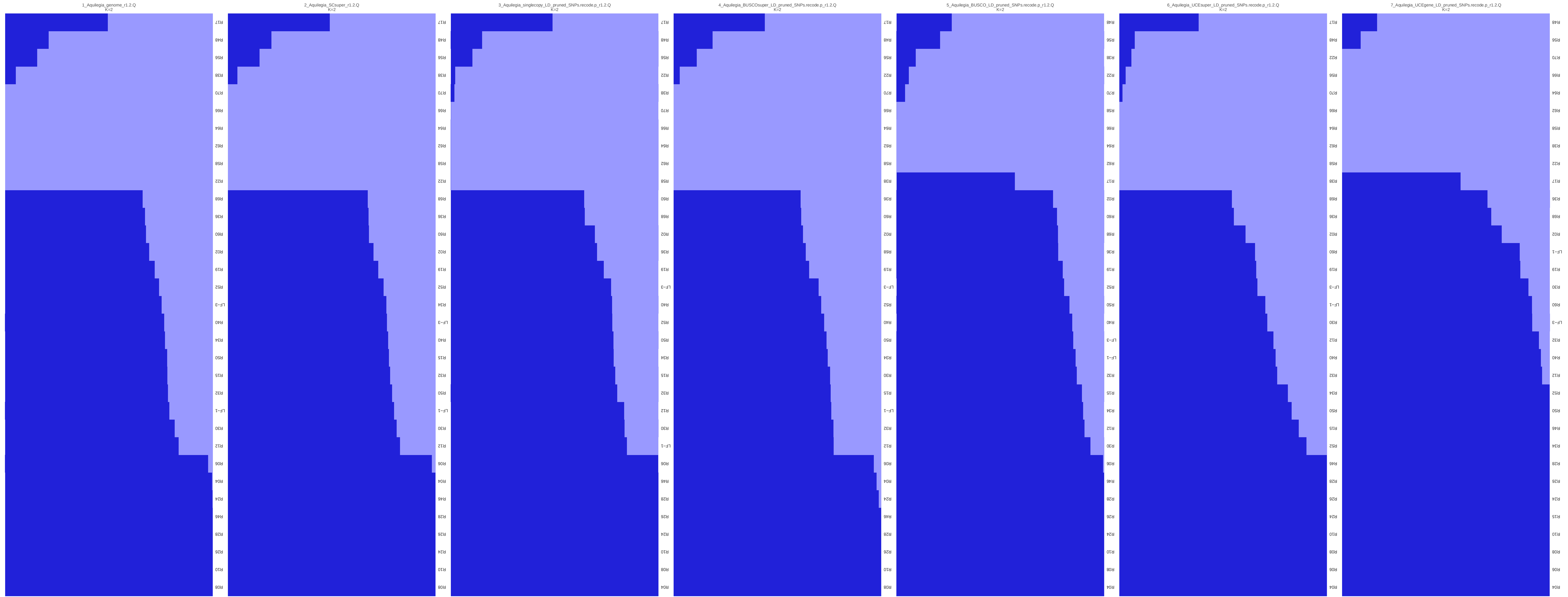
1_Aquilegia_genome_r1.2.Q
K=2
R08 R10 R26 R28 R46 R24 R04 R06 R12 R30 LF−1 R32 R15 R50 R34 R40 LF−3 R52 R19 R02 R60 R36 R68 R22 R58 R62 R64 R66 R70 R38 R56 R48 R17
2_Aquilegia_SCsuper_r1.2.Q
K=2
R08 R10 R24 R26 R28 R46 R04 R06 R12 R30 LF−1 R50 R32 R15 R40 LF−3 R34 R52 R19 R02 R60 R36 R68 R22 R58 R62 R64 R66 R70 R38 R56 R48 R17
3_Aquilegia_singlecopy_LD_pruned_SNPs.recode.p_r1.2.Q
K=2
R04 R08 R10 R24 R26 R28 R46 R06 LF−1 R30 R12 R32 R15 R34 R50 R52 R40 LF−3 R19 R36 R02 R68 R60 R58 R62 R64 R66 R70 R38 R22 R56 R48 R17
4_Aquilegia_BUSCOsuper_LD_pruned_SNPs.recode.p_r1.2.Q
K=2
R08 R10 R26 R28 R46 R24 R04 R06 R12 R32 LF−1 R15 R30 R34 R50 R40 R52 LF−3 R19 R68 R02 R60 R36 R38 R58 R62 R64 R66 R70 R22 R56 R48 R17
5_Aquilegia_BUSCO_LD_pruned_SNPs.recode.p_r1.2.Q
K=2
R04 R08 R10 R24 R26 R28 R46 R06 R30 R12 R34 R15 R32 LF−1 LF−3 R40 R50 R52 R19 R36 R68 R60 R02 R17 R62 R64 R66 R58 R70 R22 R38 R56 R48
6_Aquilegia_UCEsuper_LD_pruned_SNPs.recode.p_r1.2.Q
K=2
R04 R06 R08 R10 R24 R26 R28 R46 R52 R15 R50 R34 R32 R40 R12 R30 LF−1 LF−3 R19 R60 R02 R36 R68 R38 R58 R62 R64 R66 R70 R56 R22 R48 R17
7_Aquilegia_UCEgene_LD_pruned_SNPs.recode.p_r1.2.Q
K=2
R04 R06 R08 R10 R15 R24 R26 R28 R34 R46 R50 R52 R12 R40 R32 LF−3 R60 R30 R19 LF−1 R02 R68 R36 R17 R22 R38 R58 R62 R64 R66 R70 R56 R48

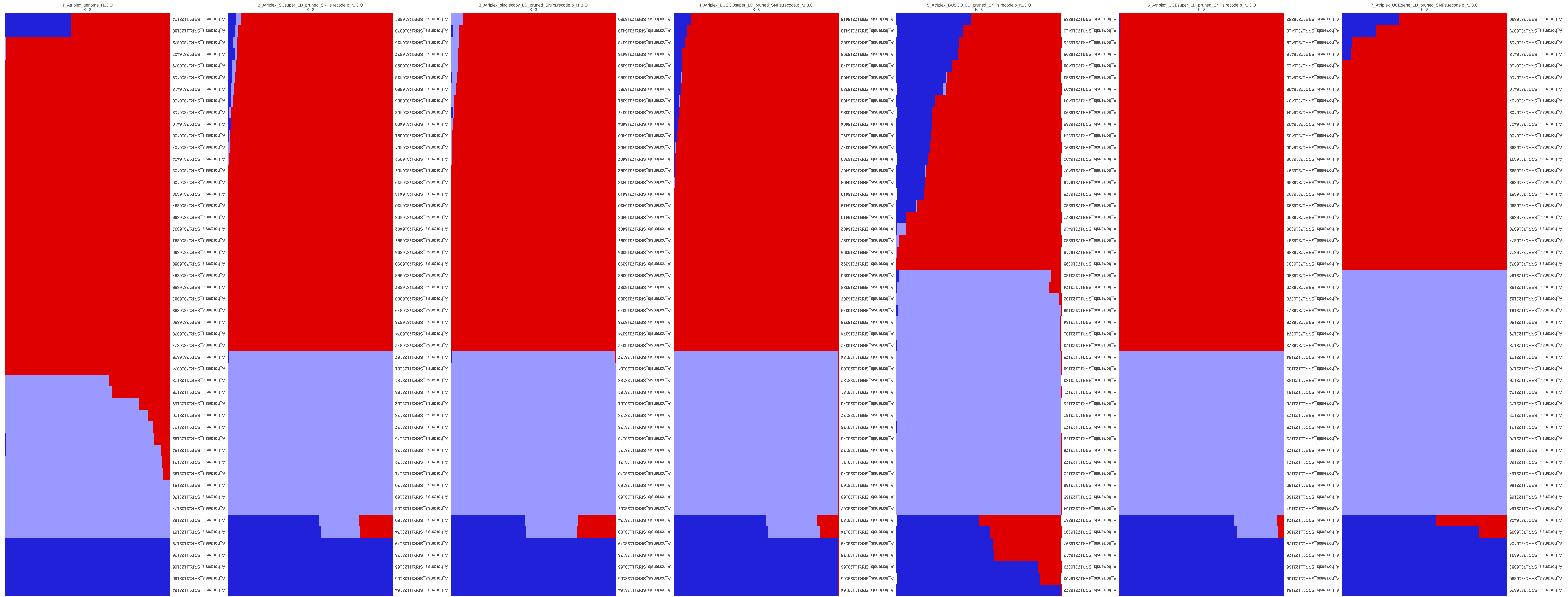
1_Atriplex_genome_r1.3.Q
K=3
2_Atriplex_SCsuper_LD_pruned_SNPs.recode.p_r1.3.Q
K=3
3_Atriplex_singlecopy_LD_pruned_SNPs.recode.p_r1.3.Q
K=3
4_Atriplex_BUSCOsuper_LD_pruned_SNPs.recode.p_r1.3.Q
K=3
5_Atriplex_BUSCO_LD_pruned_SNPs.recode.p_r1.3.Q
K=3
6_Atriplex_UCEsuper_LD_pruned_SNPs.recode.p_r1.3.Q
K=3
7_Atriplex_UCEgene_LD_pruned_SNPs.recode.p_r1.3.Q
K=3

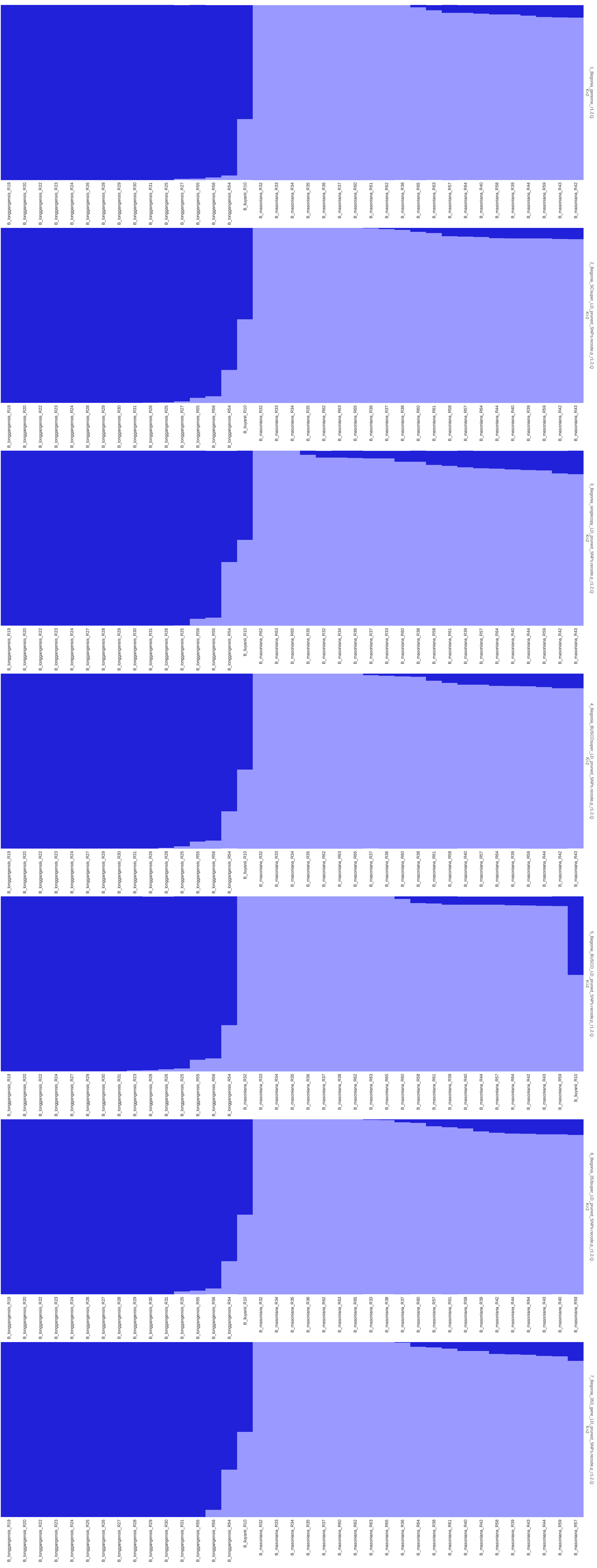

1_Begonia_genome_r1.2.Q
K=2
B_longgangensis_R19
B_longgangensis_R20
B_longgangensis_R22
B_longgangensis_R23
B_longgangensis_R24
B_longgangensis_R26
B_longgangensis_R28
B_longgangensis_R29
B_longgangensis_R30
B_longgangensis_R31
B_longgangensis_R25
B_longgangensis_R27
B_longgangensis_R55
B_longgangensis_R56
B_longgangensis_R54
B_liuyanii_R10
B_masoniana_R32
B_masoniana_R33
B_masoniana_R34
B_masoniana_R35
B_masoniana_R36
B_masoniana_R37
B_masoniana_R60
B_masoniana_R61
B_masoniana_R62
B_masoniana_R38
B_masoniana_R65
B_masoniana_R63
B_masoniana_R57
B_masoniana_R64
B_masoniana_R40
B_masoniana_R58
B_masoniana_R39
B_masoniana_R44
B_masoniana_R59
B_masoniana_R43
B_masoniana_R42
2_Begonia_SCsuper_LD_pruned_SNPs.recode.p_r1.2.Q
K=2
B_longgangensis_R19
B_longgangensis_R20
B_longgangensis_R22
B_longgangensis_R23
B_longgangensis_R24
B_longgangensis_R28
B_longgangensis_R29
B_longgangensis_R30
B_longgangensis_R31
B_longgangensis_R26
B_longgangensis_R25
B_longgangensis_R27
B_longgangensis_R55
B_longgangensis_R56
B_longgangensis_R54
B_liuyanii_R10
B_masoniana_R32
B_masoniana_R33
B_masoniana_R34
B_masoniana_R35
B_masoniana_R62
B_masoniana_R63
B_masoniana_R65
B_masoniana_R36
B_masoniana_R37
B_masoniana_R38
B_masoniana_R60
B_masoniana_R61
B_masoniana_R58
B_masoniana_R57
B_masoniana_R64
B_masoniana_R44
B_masoniana_R40
B_masoniana_R39
B_masoniana_R59
B_masoniana_R42
B_masoniana_R43
3_Begonia_singlecopy_LD_pruned_SNPs.recode.p_r1.2.Q
K=2
B_longgangensis_R19
B_longgangensis_R20
B_longgangensis_R22
B_longgangensis_R23
B_longgangensis_R24
B_longgangensis_R27
B_longgangensis_R28
B_longgangensis_R29
B_longgangensis_R30
B_longgangensis_R31
B_longgangensis_R26
B_longgangensis_R25
B_longgangensis_R56
B_longgangensis_R55
B_longgangensis_R54
B_liuyanii_R10
B_masoniana_R62
B_masoniana_R63
B_masoniana_R65
B_masoniana_R35
B_masoniana_R32
B_masoniana_R34
B_masoniana_R36
B_masoniana_R37
B_masoniana_R33
B_masoniana_R60
B_masoniana_R38
B_masoniana_R58
B_masoniana_R61
B_masoniana_R39
B_masoniana_R57
B_masoniana_R64
B_masoniana_R40
B_masoniana_R44
B_masoniana_R59
B_masoniana_R42
B_masoniana_R43
4_Begonia_BUSCOsuper_LD_pruned_SNPs.recode.p_r1.2.Q
K=2
B_longgangensis_R19
B_longgangensis_R20
B_longgangensis_R22
B_longgangensis_R23
B_longgangensis_R24
B_longgangensis_R27
B_longgangensis_R29
B_longgangensis_R30
B_longgangensis_R31
B_longgangensis_R26
B_longgangensis_R28
B_longgangensis_R25
B_longgangensis_R55
B_longgangensis_R56
B_longgangensis_R54
B_liuyanii_R10
B_masoniana_R32
B_masoniana_R33
B_masoniana_R34
B_masoniana_R35
B_masoniana_R62
B_masoniana_R63
B_masoniana_R65
B_masoniana_R37
B_masoniana_R36
B_masoniana_R60
B_masoniana_R38
B_masoniana_R61
B_masoniana_R58
B_masoniana_R40
B_masoniana_R57
B_masoniana_R64
B_masoniana_R39
B_masoniana_R59
B_masoniana_R44
B_masoniana_R42
B_masoniana_R43
5_Begonia_BUSCO_LD_pruned_SNPs.recode.p_r1.2.Q
K=2
B_longgangensis_R19
B_longgangensis_R20
B_longgangensis_R22
B_longgangensis_R24
B_longgangensis_R27
B_longgangensis_R29
B_longgangensis_R30
B_longgangensis_R31
B_longgangensis_R23
B_longgangensis_R28
B_longgangensis_R26
B_longgangensis_R25
B_longgangensis_R55
B_longgangensis_R56
B_longgangensis_R54
B_masoniana_R32
B_masoniana_R33
B_masoniana_R34
B_masoniana_R35
B_masoniana_R36
B_masoniana_R37
B_masoniana_R38
B_masoniana_R62
B_masoniana_R63
B_masoniana_R65
B_masoniana_R60
B_masoniana_R58
B_masoniana_R61
B_masoniana_R39
B_masoniana_R40
B_masoniana_R44
B_masoniana_R57
B_masoniana_R64
B_masoniana_R42
B_masoniana_R43
B_masoniana_R59
B_liuyanii_R10
6_Begonia_353super_LD_pruned_SNPs.recode.p_r1.2.Q
K=2
B_longgangensis_R19
B_longgangensis_R20
B_longgangensis_R22
B_longgangensis_R23
B_longgangensis_R24
B_longgangensis_R26
B_longgangensis_R27
B_longgangensis_R28
B_longgangensis_R29
B_longgangensis_R30
B_longgangensis_R31
B_longgangensis_R25
B_longgangensis_R55
B_longgangensis_R56
B_longgangensis_R54
B_liuyanii_R10
B_masoniana_R32
B_masoniana_R34
B_masoniana_R35
B_masoniana_R36
B_masoniana_R62
B_masoniana_R63
B_masoniana_R65
B_masoniana_R33
B_masoniana_R38
B_masoniana_R37
B_masoniana_R60
B_masoniana_R57
B_masoniana_R61
B_masoniana_R58
B_masoniana_R39
B_masoniana_R42
B_masoniana_R44
B_masoniana_R64
B_masoniana_R43
B_masoniana_R40
B_masoniana_R59
7_Begonia_353_gene_LD_pruned_SNPs.recode.p_r1.2.Q
K=2
B_longgangensis_R19
B_longgangensis_R20
B_longgangensis_R22
B_longgangensis_R23
B_longgangensis_R24
B_longgangensis_R25
B_longgangensis_R26
B_longgangensis_R27
B_longgangensis_R28
B_longgangensis_R29
B_longgangensis_R30
B_longgangensis_R31
B_longgangensis_R55
B_longgangensis_R56
B_longgangensis_R54
B_liuyanii_R10
B_masoniana_R32
B_masoniana_R33
B_masoniana_R34
B_masoniana_R35
B_masoniana_R37
B_masoniana_R60
B_masoniana_R62
B_masoniana_R63
B_masoniana_R65
B_masoniana_R36
B_masoniana_R64
B_masoniana_R38
B_masoniana_R61
B_masoniana_R40
B_masoniana_R42
B_masoniana_R58
B_masoniana_R39
B_masoniana_R43
B_masoniana_R44
B_masoniana_R59
B_masoniana_R57

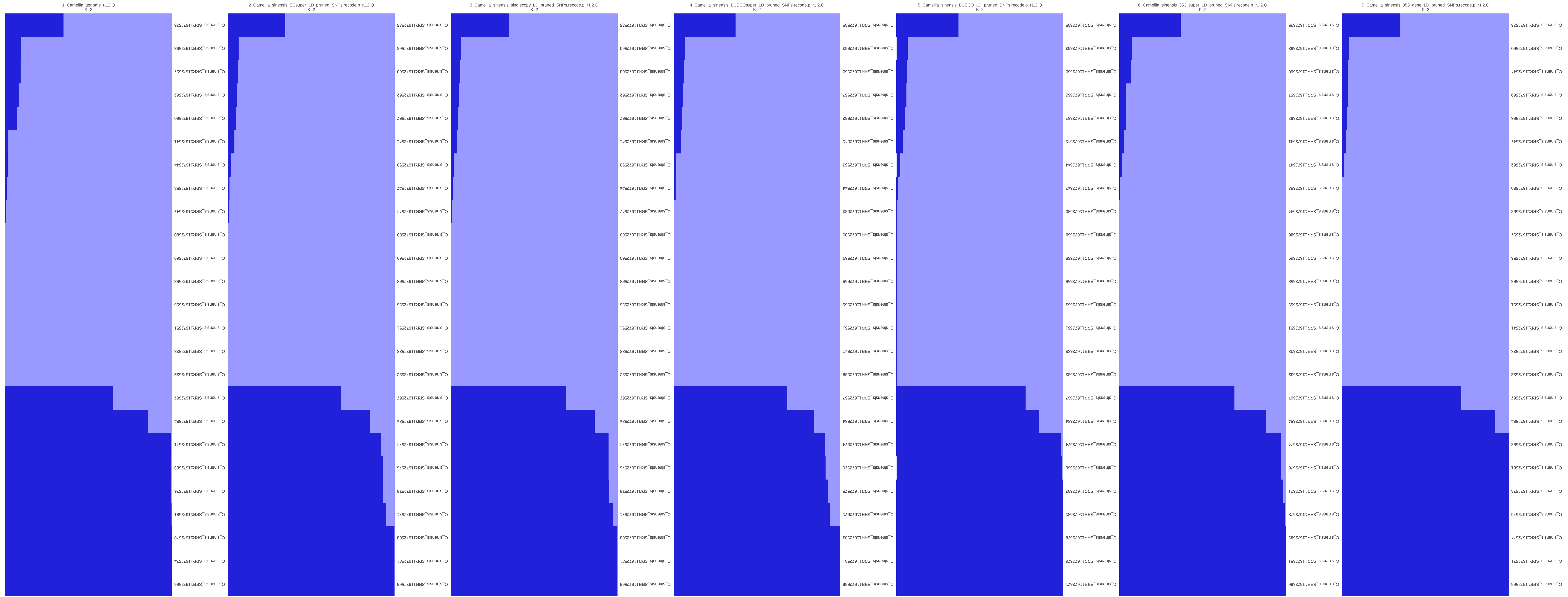
1_Camellia_genome_r1.2.Q
K=2
C_sinensis_SRR11672566
C_sinensis_SRR11672574
C_sinensis_SRR11672578
C_sinensis_SRR11672581
C_sinensis_SRR11672576
C_sinensis_SRR11672583
C_sinensis_SRR11672571
C_sinensis_SRR11672584
C_sinensis_SRR11672567
C_sinensis_SRR11672532
C_sinensis_SRR11672538
C_sinensis_SRR11672551
C_sinensis_SRR11672555
C_sinensis_SRR11672558
C_sinensis_SRR11672569
C_sinensis_SRR11672580
C_sinensis_SRR11672547
C_sinensis_SRR11672553
C_sinensis_SRR11672544
C_sinensis_SRR11672541
C_sinensis_SRR11672560
C_sinensis_SRR11672562
C_sinensis_SRR11672557
C_sinensis_SRR11672563
C_sinensis_SRR11672535
2_Camellia_sinensis_SCsuper_LD_pruned_SNPs.recode.p_r1.2.Q
K=2
C_sinensis_SRR11672566
C_sinensis_SRR11672581
C_sinensis_SRR11672583
C_sinensis_SRR11672571
C_sinensis_SRR11672578
C_sinensis_SRR11672576
C_sinensis_SRR11672574
C_sinensis_SRR11672584
C_sinensis_SRR11672567
C_sinensis_SRR11672532
C_sinensis_SRR11672538
C_sinensis_SRR11672551
C_sinensis_SRR11672555
C_sinensis_SRR11672558
C_sinensis_SRR11672569
C_sinensis_SRR11672580
C_sinensis_SRR11672544
C_sinensis_SRR11672547
C_sinensis_SRR11672553
C_sinensis_SRR11672541
C_sinensis_SRR11672557
C_sinensis_SRR11672562
C_sinensis_SRR11672560
C_sinensis_SRR11672563
C_sinensis_SRR11672535
3_Camellia_sinensis_singlecopy_LD_pruned_SNPs.recode.p_r1.2.Q
K=2
C_sinensis_SRR11672566
C_sinensis_SRR11672581
C_sinensis_SRR11672583
C_sinensis_SRR11672571
C_sinensis_SRR11672578
C_sinensis_SRR11672576
C_sinensis_SRR11672574
C_sinensis_SRR11672584
C_sinensis_SRR11672567
C_sinensis_SRR11672532
C_sinensis_SRR11672538
C_sinensis_SRR11672551
C_sinensis_SRR11672555
C_sinensis_SRR11672558
C_sinensis_SRR11672569
C_sinensis_SRR11672580
C_sinensis_SRR11672547
C_sinensis_SRR11672544
C_sinensis_SRR11672553
C_sinensis_SRR11672541
C_sinensis_SRR11672557
C_sinensis_SRR11672562
C_sinensis_SRR11672563
C_sinensis_SRR11672560
C_sinensis_SRR11672535
4_Camellia_sinensis_BUSCOsuper_LD_pruned_SNPs.recode.p_r1.2.Q
K=2
C_sinensis_SRR11672566
C_sinensis_SRR11672581
C_sinensis_SRR11672583
C_sinensis_SRR11672571
C_sinensis_SRR11672578
C_sinensis_SRR11672576
C_sinensis_SRR11672574
C_sinensis_SRR11672584
C_sinensis_SRR11672567
C_sinensis_SRR11672538
C_sinensis_SRR11672547
C_sinensis_SRR11672551
C_sinensis_SRR11672555
C_sinensis_SRR11672558
C_sinensis_SRR11672569
C_sinensis_SRR11672580
C_sinensis_SRR11672532
C_sinensis_SRR11672544
C_sinensis_SRR11672553
C_sinensis_SRR11672541
C_sinensis_SRR11672562
C_sinensis_SRR11672557
C_sinensis_SRR11672560
C_sinensis_SRR11672563
C_sinensis_SRR11672535
5_Camellia_sinensis_BUSCO_LD_pruned_SNPs.recode.p_r1.2.Q
K=2
C_sinensis_SRR11672571
C_sinensis_SRR11672576
C_sinensis_SRR11672578
C_sinensis_SRR11672581
C_sinensis_SRR11672583
C_sinensis_SRR11672566
C_sinensis_SRR11672574
C_sinensis_SRR11672584
C_sinensis_SRR11672567
C_sinensis_SRR11672532
C_sinensis_SRR11672538
C_sinensis_SRR11672551
C_sinensis_SRR11672553
C_sinensis_SRR11672555
C_sinensis_SRR11672558
C_sinensis_SRR11672569
C_sinensis_SRR11672580
C_sinensis_SRR11672547
C_sinensis_SRR11672544
C_sinensis_SRR11672541
C_sinensis_SRR11672557
C_sinensis_SRR11672562
C_sinensis_SRR11672560
C_sinensis_SRR11672563
C_sinensis_SRR11672535
6_Camellia_sinensis_353_super_LD_pruned_SNPs.recode.p_r1.2.Q
K=2
C_sinensis_SRR11672566
C_sinensis_SRR11672581
C_sinensis_SRR11672583
C_sinensis_SRR11672578
C_sinensis_SRR11672571
C_sinensis_SRR11672576
C_sinensis_SRR11672574
C_sinensis_SRR11672584
C_sinensis_SRR11672567
C_sinensis_SRR11672532
C_sinensis_SRR11672538
C_sinensis_SRR11672551
C_sinensis_SRR11672555
C_sinensis_SRR11672558
C_sinensis_SRR11672569
C_sinensis_SRR11672580
C_sinensis_SRR11672544
C_sinensis_SRR11672553
C_sinensis_SRR11672547
C_sinensis_SRR11672541
C_sinensis_SRR11672562
C_sinensis_SRR11672557
C_sinensis_SRR11672560
C_sinensis_SRR11672563
C_sinensis_SRR11672535
7_Camellia_sinensis_353_gene_LD_pruned_SNPs.recode.p_r1.2.Q
K=2
C_sinensis_SRR11672566
C_sinensis_SRR11672571
C_sinensis_SRR11672574
C_sinensis_SRR11672576
C_sinensis_SRR11672578
C_sinensis_SRR11672581
C_sinensis_SRR11672583
C_sinensis_SRR11672584
C_sinensis_SRR11672567
C_sinensis_SRR11672532
C_sinensis_SRR11672538
C_sinensis_SRR11672541
C_sinensis_SRR11672551
C_sinensis_SRR11672553
C_sinensis_SRR11672555
C_sinensis_SRR11672557
C_sinensis_SRR11672558
C_sinensis_SRR11672580
C_sinensis_SRR11672562
C_sinensis_SRR11672547
C_sinensis_SRR11672563
C_sinensis_SRR11672569
C_sinensis_SRR11672544
C_sinensis_SRR11672560
C_sinensis_SRR11672535

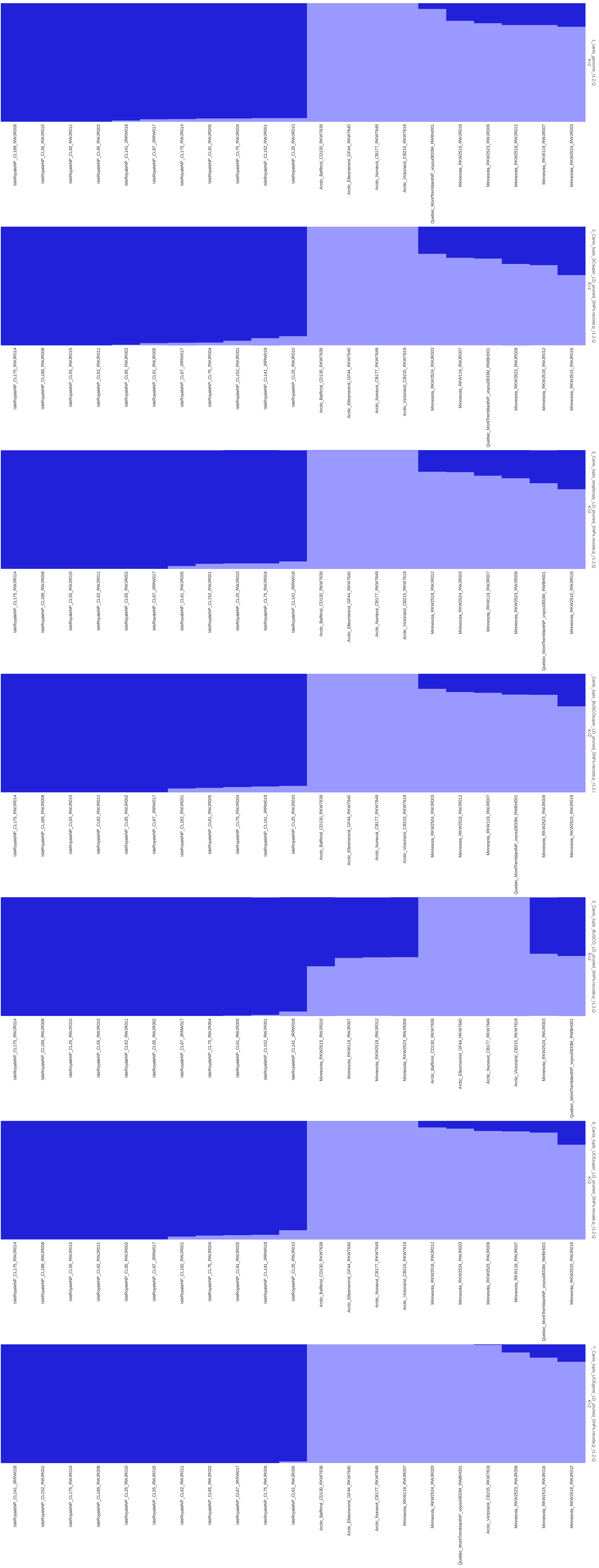
1_canis_genome_r1.2.Q
K=2
IsleRoyaleNP_CL189_RWJR008
IsleRoyaleNP_CL55_RWJR015
IsleRoyaleNP_CL62_RWJR011
IsleRoyaleNP_CL65_RWJR002
IsleRoyaleNP_CL141_JRRW018
IsleRoyaleNP_CL67_JRRW017
IsleRoyaleNP_CL175_RWJR014
IsleRoyaleNP_CL61_RWJR005
IsleRoyaleNP_CL75_RWJR004
IsleRoyaleNP_CL152_RWJR001
IsleRoyaleNP_CL25_RWJR010
Arctic_BaffinIsl_CD130_RKW7639
Arctic_EllesmereIsl_GF44_RKW7640
Arctic_Nunavut_CB177_RKW7649
Arctic_VictoriaIsl_CB215_RKW7619
Quebec_MontTremblantNP_voyou0833M_RWBH001
Minnesota_RKW2515_RWJR016
Minnesota_RKW2523_RWJR009
Minnesota_RKW2518_RWJR012
Minnesota_RKW119_RWJR007
Minnesota_RKW2524_RWJR003
2_Canis_lupis_SCsuper_LD_pruned_SNPs.recode.p_r1.2.Q
K=2
IsleRoyaleNP_CL175_RWJR014
IsleRoyaleNP_CL189_RWJR008
IsleRoyaleNP_CL55_RWJR015
IsleRoyaleNP_CL62_RWJR011
IsleRoyaleNP_CL65_RWJR002
IsleRoyaleNP_CL61_RWJR005
IsleRoyaleNP_CL67_JRRW017
IsleRoyaleNP_CL75_RWJR004
IsleRoyaleNP_CL152_RWJR001
IsleRoyaleNP_CL141_JRRW018
IsleRoyaleNP_CL25_RWJR010
Arctic_BaffinIsl_CD130_RKW7639
Arctic_EllesmereIsl_GF44_RKW7640
Arctic_Nunavut_CB177_RKW7649
Arctic_VictoriaIsl_CB215_RKW7619
Minnesota_RKW2524_RWJR003
Minnesota_RKW119_RWJR007
Quebec_MontTremblantNP_voyou0833M_RWBH001
Minnesota_RKW2523_RWJR009
Minnesota_RKW2518_RWJR012
Minnesota_RKW2515_RWJR016
3_Canis_lupis_singlecopy_LD_pruned_SNPs.recode.p_r1.2.Q
K=2
IsleRoyaleNP_CL175_RWJR014
IsleRoyaleNP_CL189_RWJR008
IsleRoyaleNP_CL55_RWJR015
IsleRoyaleNP_CL62_RWJR011
IsleRoyaleNP_CL65_RWJR002
IsleRoyaleNP_CL67_JRRW017
IsleRoyaleNP_CL61_RWJR005
IsleRoyaleNP_CL152_RWJR001
IsleRoyaleNP_CL25_RWJR010
IsleRoyaleNP_CL75_RWJR004
IsleRoyaleNP_CL141_JRRW018
Arctic_BaffinIsl_CD130_RKW7639
Arctic_EllesmereIsl_GF44_RKW7640
Arctic_Nunavut_CB177_RKW7649
Arctic_VictoriaIsl_CB215_RKW7619
Minnesota_RKW2518_RWJR012
Minnesota_RKW2524_RWJR003
Minnesota_RKW119_RWJR007
Minnesota_RKW2523_RWJR009
Quebec_MontTremblantNP_voyou0833M_RWBH001
Minnesota_RKW2515_RWJR016
_Canis_lupis_BUSCOsuper_LD_pruned_SNPs.recode.p_r1.2.(
K=2
IsleRoyaleNP_CL175_RWJR014
IsleRoyaleNP_CL189_RWJR008
IsleRoyaleNP_CL55_RWJR015
IsleRoyaleNP_CL62_RWJR011
IsleRoyaleNP_CL65_RWJR002
IsleRoyaleNP_CL67_JRRW017
IsleRoyaleNP_CL152_RWJR001
IsleRoyaleNP_CL61_RWJR005
IsleRoyaleNP_CL75_RWJR004
IsleRoyaleNP_CL141_JRRW018
IsleRoyaleNP_CL25_RWJR010
Arctic_BaffinIsl_CD130_RKW7639
Arctic_EllesmereIsl_GF44_RKW7640
Arctic_Nunavut_CB177_RKW7649
Arctic_VictoriaIsl_CB215_RKW7619
Minnesota_RKW2524_RWJR003
Minnesota_RKW2518_RWJR012
Minnesota_RKW119_RWJR007
Quebec_MontTremblantNP_voyou0833M_RWBH001
Minnesota_RKW2523_RWJR009
Minnesota_RKW2515_RWJR016
5_Canis_lupis_BUSCO_LD_pruned_SNPs.recode.p_r1.2.Q
K=2
IsleRoyaleNP_CL175_RWJR014
IsleRoyaleNP_CL189_RWJR008
IsleRoyaleNP_CL25_RWJR010
IsleRoyaleNP_CL55_RWJR015
IsleRoyaleNP_CL62_RWJR011
IsleRoyaleNP_CL65_RWJR002
IsleRoyaleNP_CL67_JRRW017
IsleRoyaleNP_CL75_RWJR004
IsleRoyaleNP_CL61_RWJR005
IsleRoyaleNP_CL152_RWJR001
IsleRoyaleNP_CL141_JRRW018
Minnesota_RKW2515_RWJR016
Minnesota_RKW119_RWJR007
Minnesota_RKW2518_RWJR012
Minnesota_RKW2523_RWJR009
Arctic_BaffinIsl_CD130_RKW7639
Arctic_EllesmereIsl_GF44_RKW7640
Arctic_Nunavut_CB177_RKW7649
Arctic_VictoriaIsl_CB215_RKW7619
Minnesota_RKW2524_RWJR003
Quebec_MontTremblantNP_voyou0833M_RWBH001
6_Canis_lupis_UCEsuper_LD_pruned_SNPs.recode.p_r1.2.Q
K=2
IsleRoyaleNP_CL175_RWJR014
IsleRoyaleNP_CL189_RWJR008
IsleRoyaleNP_CL55_RWJR015
IsleRoyaleNP_CL62_RWJR011
IsleRoyaleNP_CL65_RWJR002
IsleRoyaleNP_CL67_JRRW017
IsleRoyaleNP_CL152_RWJR001
IsleRoyaleNP_CL75_RWJR004
IsleRoyaleNP_CL61_RWJR005
IsleRoyaleNP_CL141_JRRW018
IsleRoyaleNP_CL25_RWJR010
Arctic_BaffinIsl_CD130_RKW7639
Arctic_EllesmereIsl_GF44_RKW7640
Arctic_Nunavut_CB177_RKW7649
Arctic_VictoriaIsl_CB215_RKW7619
Minnesota_RKW2518_RWJR012
Minnesota_RKW2524_RWJR003
Minnesota_RKW2523_RWJR009
Minnesota_RKW119_RWJR007
Quebec_MontTremblantNP_voyou0833M_RWBH001
Minnesota_RKW2515_RWJR016
7_Canis_lupis_UCEgene_LD_pruned_SNPs.recode.p_r1.2.Q
K=2
IsleRoyaleNP_CL141_JRRW018
IsleRoyaleNP_CL152_RWJR001
IsleRoyaleNP_CL175_RWJR014
IsleRoyaleNP_CL189_RWJR008
IsleRoyaleNP_CL25_RWJR010
IsleRoyaleNP_CL55_RWJR015
IsleRoyaleNP_CL62_RWJR011
IsleRoyaleNP_CL65_RWJR002
IsleRoyaleNP_CL67_JRRW017
IsleRoyaleNP_CL75_RWJR004
IsleRoyaleNP_CL61_RWJR005
Arctic_BaffinIsl_CD130_RKW7639
Arctic_EllesmereIsl_GF44_RKW7640
Arctic_Nunavut_CB177_RKW7649
Minnesota_RKW119_RWJR007
Minnesota_RKW2524_RWJR003
Quebec_MontTremblantNP_voyou0833M_RWBH001
Arctic_VictoriaIsl_CB215_RKW7619
Minnesota_RKW2523_RWJR009
Minnesota_RKW2515_RWJR016
Minnesota_RKW2518_RWJR012

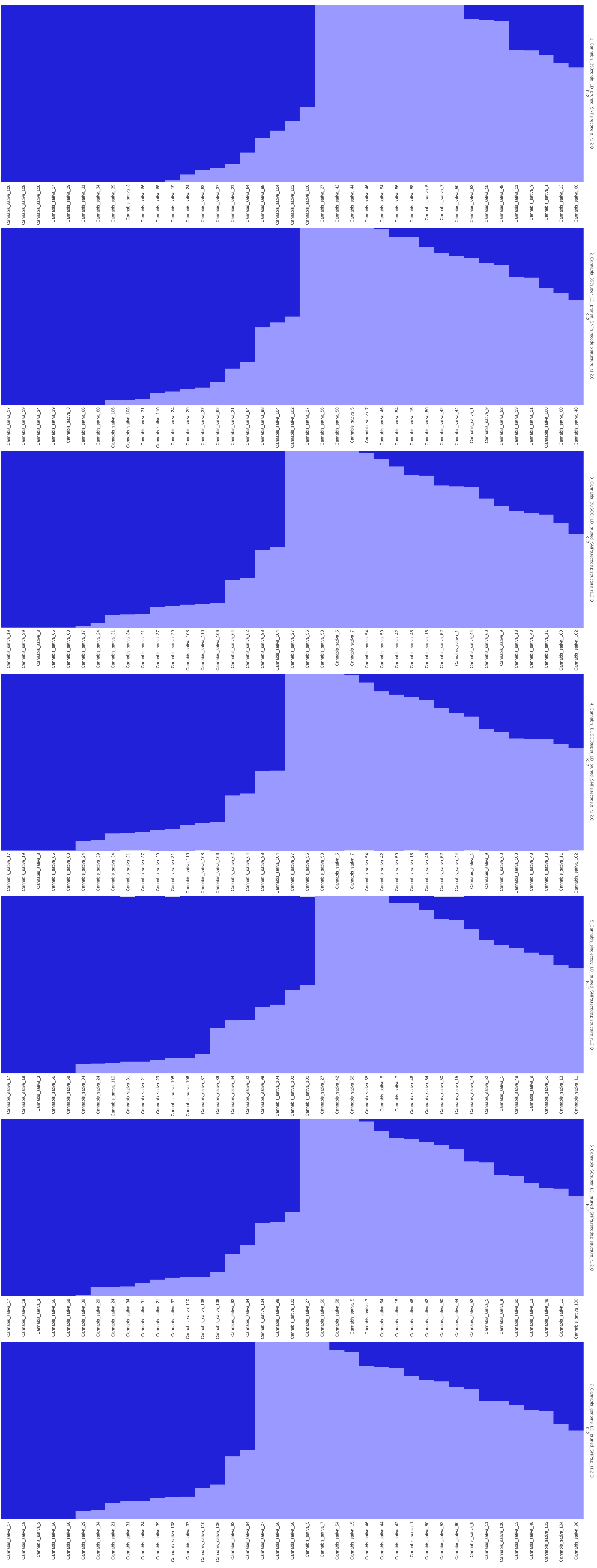
1_Cannabis_353contig_LD_pruned_SNPs.recode.p_r1.2.Q
K=2
Cannabis_sativa_106
Cannabis_sativa_108
Cannabis_sativa_110
Cannabis_sativa_17
Cannabis_sativa_29
Cannabis_sativa_31
Cannabis_sativa_34
Cannabis_sativa_39
Cannabis_sativa_3
Cannabis_sativa_66
Cannabis_sativa_68
Cannabis_sativa_19
Cannabis_sativa_24
Cannabis_sativa_62
Cannabis_sativa_37
Cannabis_sativa_21
Cannabis_sativa_64
Cannabis_sativa_98
Cannabis_sativa_104
Cannabis_sativa_102
Cannabis_sativa_100
Cannabis_sativa_27
Cannabis_sativa_42
Cannabis_sativa_44
Cannabis_sativa_46
Cannabis_sativa_54
Cannabis_sativa_56
Cannabis_sativa_58
Cannabis_sativa_5
Cannabis_sativa_7
Cannabis_sativa_50
Cannabis_sativa_52
Cannabis_sativa_15
Cannabis_sativa_48
Cannabis_sativa_11
Cannabis_sativa_9
Cannabis_sativa_1
Cannabis_sativa_13
Cannabis_sativa_60
2_Cannabis_353super_LD_pruned_SNPs.recode.p.structure_r1.2.Q
K=2
Cannabis_sativa_17
Cannabis_sativa_19
Cannabis_sativa_34
Cannabis_sativa_39
Cannabis_sativa_3
Cannabis_sativa_66
Cannabis_sativa_68
Cannabis_sativa_106
Cannabis_sativa_108
Cannabis_sativa_31
Cannabis_sativa_110
Cannabis_sativa_24
Cannabis_sativa_29
Cannabis_sativa_37
Cannabis_sativa_62
Cannabis_sativa_21
Cannabis_sativa_64
Cannabis_sativa_98
Cannabis_sativa_104
Cannabis_sativa_102
Cannabis_sativa_27
Cannabis_sativa_56
Cannabis_sativa_58
Cannabis_sativa_5
Cannabis_sativa_7
Cannabis_sativa_46
Cannabis_sativa_54
Cannabis_sativa_15
Cannabis_sativa_50
Cannabis_sativa_42
Cannabis_sativa_44
Cannabis_sativa_1
Cannabis_sativa_9
Cannabis_sativa_52
Cannabis_sativa_13
Cannabis_sativa_11
Cannabis_sativa_100
Cannabis_sativa_60
Cannabis_sativa_48
3_Cannabis_BUSCO_LD_pruned_SNPs.recode.p.structure_r1.2.Q
K=2
Cannabis_sativa_19
Cannabis_sativa_39
Cannabis_sativa_3
Cannabis_sativa_66
Cannabis_sativa_68
Cannabis_sativa_17
Cannabis_sativa_24
Cannabis_sativa_31
Cannabis_sativa_34
Cannabis_sativa_21
Cannabis_sativa_37
Cannabis_sativa_29
Cannabis_sativa_108
Cannabis_sativa_110
Cannabis_sativa_106
Cannabis_sativa_64
Cannabis_sativa_62
Cannabis_sativa_98
Cannabis_sativa_104
Cannabis_sativa_27
Cannabis_sativa_56
Cannabis_sativa_58
Cannabis_sativa_5
Cannabis_sativa_7
Cannabis_sativa_54
Cannabis_sativa_50
Cannabis_sativa_42
Cannabis_sativa_46
Cannabis_sativa_15
Cannabis_sativa_52
Cannabis_sativa_1
Cannabis_sativa_44
Cannabis_sativa_60
Cannabis_sativa_9
Cannabis_sativa_13
Cannabis_sativa_48
Cannabis_sativa_11
Cannabis_sativa_100
Cannabis_sativa_102
4_Cannabis_BUSCOsuper_LD_pruned_SNPs.recode.p_r1.2.Q
K=2
Cannabis_sativa_17
Cannabis_sativa_19
Cannabis_sativa_3
Cannabis_sativa_66
Cannabis_sativa_68
Cannabis_sativa_24
Cannabis_sativa_39
Cannabis_sativa_34
Cannabis_sativa_21
Cannabis_sativa_37
Cannabis_sativa_29
Cannabis_sativa_31
Cannabis_sativa_110
Cannabis_sativa_108
Cannabis_sativa_106
Cannabis_sativa_62
Cannabis_sativa_64
Cannabis_sativa_98
Cannabis_sativa_104
Cannabis_sativa_27
Cannabis_sativa_56
Cannabis_sativa_58
Cannabis_sativa_5
Cannabis_sativa_7
Cannabis_sativa_54
Cannabis_sativa_42
Cannabis_sativa_50
Cannabis_sativa_15
Cannabis_sativa_46
Cannabis_sativa_52
Cannabis_sativa_44
Cannabis_sativa_1
Cannabis_sativa_9
Cannabis_sativa_60
Cannabis_sativa_100
Cannabis_sativa_48
Cannabis_sativa_13
Cannabis_sativa_11
Cannabis_sativa_102
5_Cannabis_singlecopy_LD_pruned_SNPs.recode.p.structure_r1.2.Q
K=2
Cannabis_sativa_17
Cannabis_sativa_19
Cannabis_sativa_3
Cannabis_sativa_66
Cannabis_sativa_68
Cannabis_sativa_34
Cannabis_sativa_24
Cannabis_sativa_110
Cannabis_sativa_31
Cannabis_sativa_21
Cannabis_sativa_29
Cannabis_sativa_108
Cannabis_sativa_106
Cannabis_sativa_37
Cannabis_sativa_39
Cannabis_sativa_64
Cannabis_sativa_62
Cannabis_sativa_98
Cannabis_sativa_104
Cannabis_sativa_102
Cannabis_sativa_100
Cannabis_sativa_27
Cannabis_sativa_42
Cannabis_sativa_56
Cannabis_sativa_58
Cannabis_sativa_5
Cannabis_sativa_7
Cannabis_sativa_46
Cannabis_sativa_54
Cannabis_sativa_50
Cannabis_sativa_15
Cannabis_sativa_44
Cannabis_sativa_52
Cannabis_sativa_1
Cannabis_sativa_48
Cannabis_sativa_9
Cannabis_sativa_60
Cannabis_sativa_13
Cannabis_sativa_11
6_Cannabis_SCsuper_LD_pruned_SNPs.recode.p.structure_r1.2.Q
K=2
Cannabis_sativa_17
Cannabis_sativa_19
Cannabis_sativa_3
Cannabis_sativa_66
Cannabis_sativa_68
Cannabis_sativa_39
Cannabis_sativa_29
Cannabis_sativa_24
Cannabis_sativa_34
Cannabis_sativa_31
Cannabis_sativa_21
Cannabis_sativa_37
Cannabis_sativa_110
Cannabis_sativa_108
Cannabis_sativa_106
Cannabis_sativa_62
Cannabis_sativa_64
Cannabis_sativa_104
Cannabis_sativa_98
Cannabis_sativa_102
Cannabis_sativa_27
Cannabis_sativa_56
Cannabis_sativa_58
Cannabis_sativa_5
Cannabis_sativa_7
Cannabis_sativa_54
Cannabis_sativa_15
Cannabis_sativa_46
Cannabis_sativa_42
Cannabis_sativa_50
Cannabis_sativa_44
Cannabis_sativa_52
Cannabis_sativa_1
Cannabis_sativa_9
Cannabis_sativa_60
Cannabis_sativa_13
Cannabis_sativa_48
Cannabis_sativa_11
Cannabis_sativa_100
7_Cannabis_genome_LD_pruned_SNPs.p_r1.2.Q
K=2
Cannabis_sativa_17
Cannabis_sativa_19
Cannabis_sativa_3
Cannabis_sativa_66
Cannabis_sativa_68
Cannabis_sativa_29
Cannabis_sativa_34
Cannabis_sativa_21
Cannabis_sativa_31
Cannabis_sativa_24
Cannabis_sativa_39
Cannabis_sativa_108
Cannabis_sativa_37
Cannabis_sativa_110
Cannabis_sativa_106
Cannabis_sativa_62
Cannabis_sativa_64
Cannabis_sativa_27
Cannabis_sativa_56
Cannabis_sativa_58
Cannabis_sativa_5
Cannabis_sativa_7
Cannabis_sativa_54
Cannabis_sativa_15
Cannabis_sativa_46
Cannabis_sativa_44
Cannabis_sativa_42
Cannabis_sativa_1
Cannabis_sativa_50
Cannabis_sativa_52
Cannabis_sativa_60
Cannabis_sativa_9
Cannabis_sativa_11
Cannabis_sativa_100
Cannabis_sativa_13
Cannabis_sativa_48
Cannabis_sativa_102
Cannabis_sativa_104
Cannabis_sativa_98

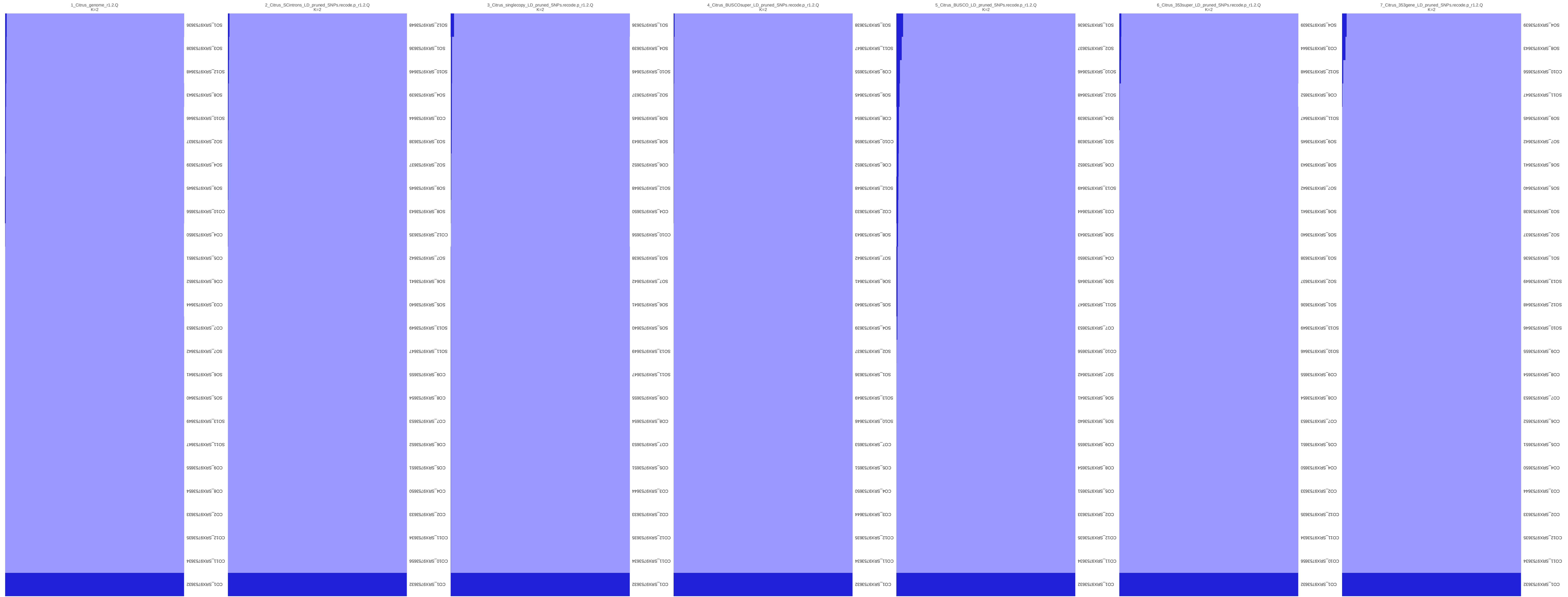
1_Citrus_genome_r1.2.Q
K=2
CO1_SRX9753632
CO11_SRX9753634
CO12_SRX9753635
CO2_SRX9753633
CO8_SRX9753654
CO9_SRX9753655
SO11_SRX9753647
SO13_SRX9753649
SO5_SRX9753640
SO6_SRX9753641
SO7_SRX9753642
CO7_SRX9753653
CO3_SRX9753644
CO6_SRX9753652
CO5_SRX9753651
CO4_SRX9753650
CO10_SRX9753656
SO9_SRX9753645
SO4_SRX9753639
SO2_SRX9753637
SO10_SRX9753646
SO8_SRX9753643
SO12_SRX9753648
SO3_SRX9753638
SO1_SRX9753636
2_Citrus_SCintrons_LD_pruned_SNPs.recode.p_r1.2.Q
K=2
CO1_SRX9753632
CO10_SRX9753656
CO11_SRX9753634
CO2_SRX9753633
CO4_SRX9753650
CO5_SRX9753651
CO6_SRX9753652
CO7_SRX9753653
CO8_SRX9753654
CO9_SRX9753655
SO11_SRX9753647
SO13_SRX9753649
SO5_SRX9753640
SO6_SRX9753641
SO7_SRX9753642
CO12_SRX9753635
SO8_SRX9753643
SO9_SRX9753645
SO2_SRX9753637
SO3_SRX9753638
CO3_SRX9753644
SO4_SRX9753639
SO10_SRX9753646
SO1_SRX9753636
SO12_SRX9753648
3_Citrus_singlecopy_LD_pruned_SNPs.recode.p_r1.2.Q
K=2
CO1_SRX9753632
CO11_SRX9753634
CO12_SRX9753635
CO2_SRX9753633
CO3_SRX9753644
CO5_SRX9753651
CO7_SRX9753653
CO8_SRX9753654
CO9_SRX9753655
SO11_SRX9753647
SO13_SRX9753649
SO5_SRX9753640
SO6_SRX9753641
SO7_SRX9753642
SO3_SRX9753638
CO10_SRX9753656
CO4_SRX9753650
SO12_SRX9753648
CO6_SRX9753652
SO8_SRX9753643
SO9_SRX9753645
SO2_SRX9753637
SO10_SRX9753646
SO4_SRX9753639
SO1_SRX9753636
4_Citrus_BUSCOsuper_LD_pruned_SNPs.recode.p_r1.2.Q
K=2
CO1_SRX9753632
CO11_SRX9753634
CO12_SRX9753635
CO3_SRX9753644
CO4_SRX9753650
CO5_SRX9753651
CO7_SRX9753653
SO10_SRX9753646
SO13_SRX9753649
SO1_SRX9753636
SO2_SRX9753637
SO4_SRX9753639
SO5_SRX9753640
SO6_SRX9753641
SO7_SRX9753642
SO8_SRX9753643
CO2_SRX9753633
SO12_SRX9753648
CO6_SRX9753652
CO10_SRX9753656
CO8_SRX9753654
SO9_SRX9753645
CO9_SRX9753655
SO11_SRX9753647
SO3_SRX9753638
5_Citrus_BUSCO_LD_pruned_SNPs.recode.p_r1.2.Q
K=2
CO1_SRX9753632
CO11_SRX9753634
CO12_SRX9753635
CO2_SRX9753633
CO5_SRX9753651
CO8_SRX9753654
CO9_SRX9753655
SO5_SRX9753640
SO6_SRX9753641
SO7_SRX9753642
CO10_SRX9753656
CO7_SRX9753653
SO11_SRX9753647
SO9_SRX9753645
CO4_SRX9753650
SO8_SRX9753643
CO3_SRX9753644
SO13_SRX9753649
CO6_SRX9753652
SO3_SRX9753638
SO4_SRX9753639
SO12_SRX9753648
SO10_SRX9753646
SO2_SRX9753637
SO1_SRX9753636
6_Citrus_353super_LD_pruned_SNPs.recode.p_r1.2.Q
K=2
CO1_SRX9753632
CO10_SRX9753656
CO11_SRX9753634
CO12_SRX9753635
CO2_SRX9753633
CO4_SRX9753650
CO5_SRX9753651
CO7_SRX9753653
CO8_SRX9753654
CO9_SRX9753655
SO10_SRX9753646
SO13_SRX9753649
SO1_SRX9753636
SO2_SRX9753637
SO3_SRX9753638
SO5_SRX9753640
SO6_SRX9753641
SO7_SRX9753642
SO8_SRX9753643
SO9_SRX9753645
SO11_SRX9753647
CO6_SRX9753652
SO12_SRX9753648
CO3_SRX9753644
SO4_SRX9753639
7_Citrus_353gene_LD_pruned_SNPs.recode.p_r1.2.Q
K=2
CO1_SRX9753632
CO11_SRX9753634
CO12_SRX9753635
CO2_SRX9753633
CO3_SRX9753644
CO4_SRX9753650
CO5_SRX9753651
CO6_SRX9753652
CO7_SRX9753653
CO8_SRX9753654
CO9_SRX9753655
SO10_SRX9753646
SO12_SRX9753648
SO13_SRX9753649
SO1_SRX9753636
SO2_SRX9753637
SO3_SRX9753638
SO5_SRX9753640
SO6_SRX9753641
SO7_SRX9753642
SO9_SRX9753645
SO11_SRX9753647
CO10_SRX9753656
SO8_SRX9753643
SO4_SRX9753639

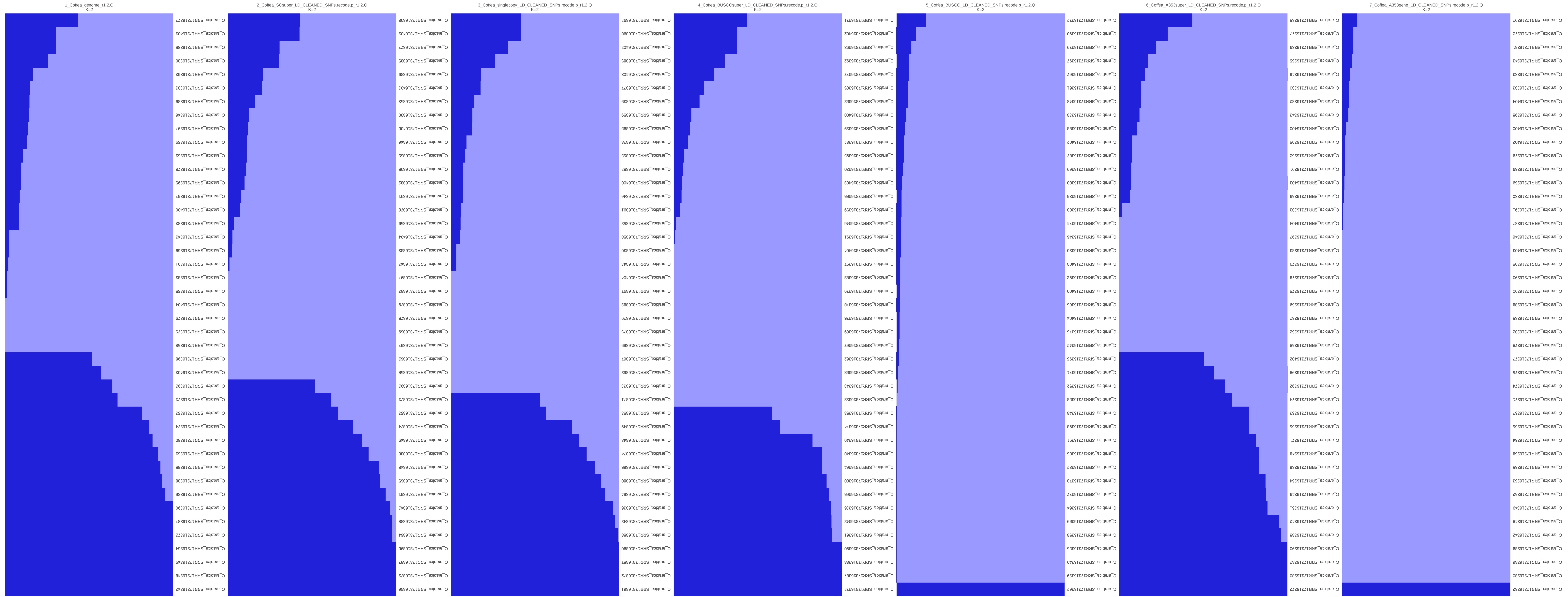
1_Coffea_genome_r1.2.Q
K=2
2_Coffea_SCsuper_LD_CLEANED_SNPs.recode.p_r1.2.Q
K=2
3_Coffea_singlecopy_LD_CLEANED_SNPs.recode.p_r1.2.Q
K=2
4_Coffea_BUSCOsuper_LD_CLEANED_SNPs.recode.p_r1.2.Q
K=2
5_Coffea_BUSCO_LD_CLEANED_SNPs.recode.p_r1.2.Q
K=2
6_Coffea_A353super_LD_CLEANED_SNPs.recode.p_r1.2.Q
K=2
7_Coffea_A353gene_LD_CLEANED_SNPs.recode.p_r1.2.Q
K=2

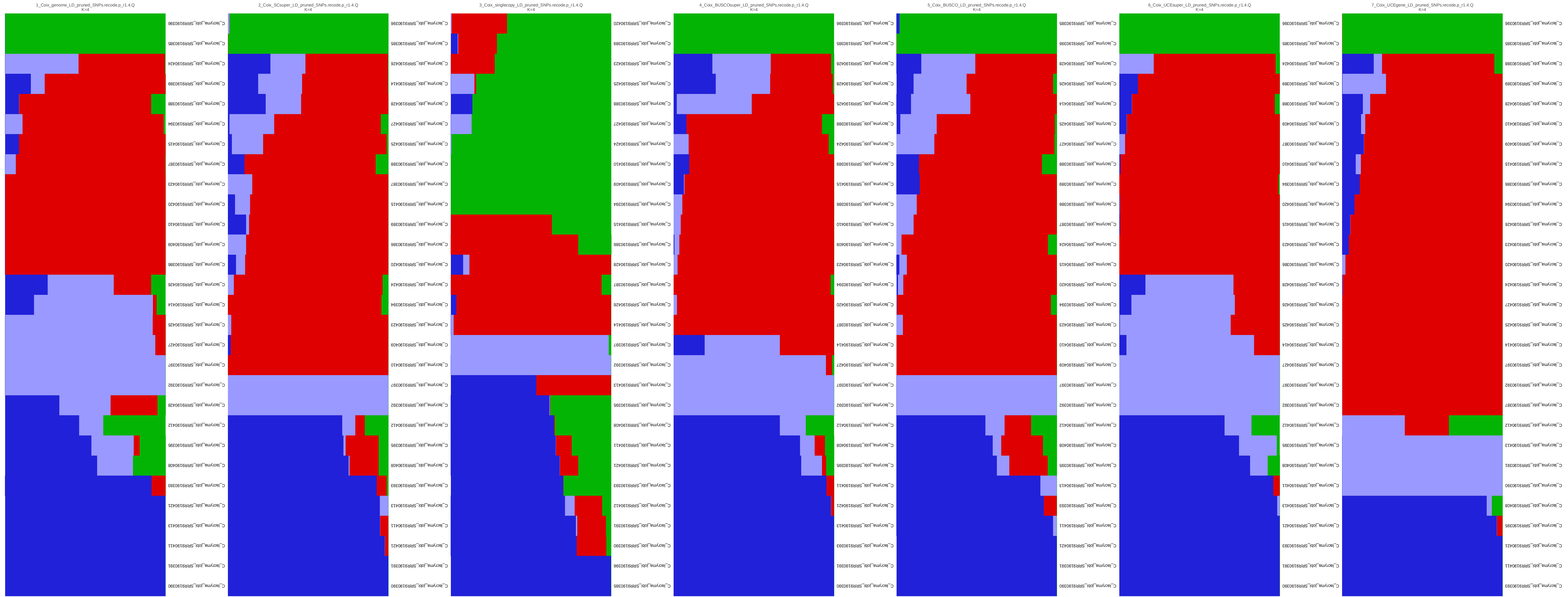
1_Coix_genome_LD_pruned_SNPs.recode.p_r1.4.Q
K=4
2_Coix_SCsuper_LD_pruned_SNPs.recode.p_r1.4.Q
K=4
3_Coix_singlecopy_LD_pruned_SNPs.recode.p_r1.4.Q
K=4
4_Coix_BUSCOsuper_LD_pruned_SNPs.recode.p_r1.4.Q
K=4
5_Coix_BUSCO_LD_pruned_SNPs.recode.p_r1.4.Q
K=4
6_Coix_UCEsuper_LD_pruned_SNPs.recode.p_r1.4.Q
K=4
7_Coix_UCEgene_LD_pruned_SNPs.recode.p_r1.4.Q
K=4

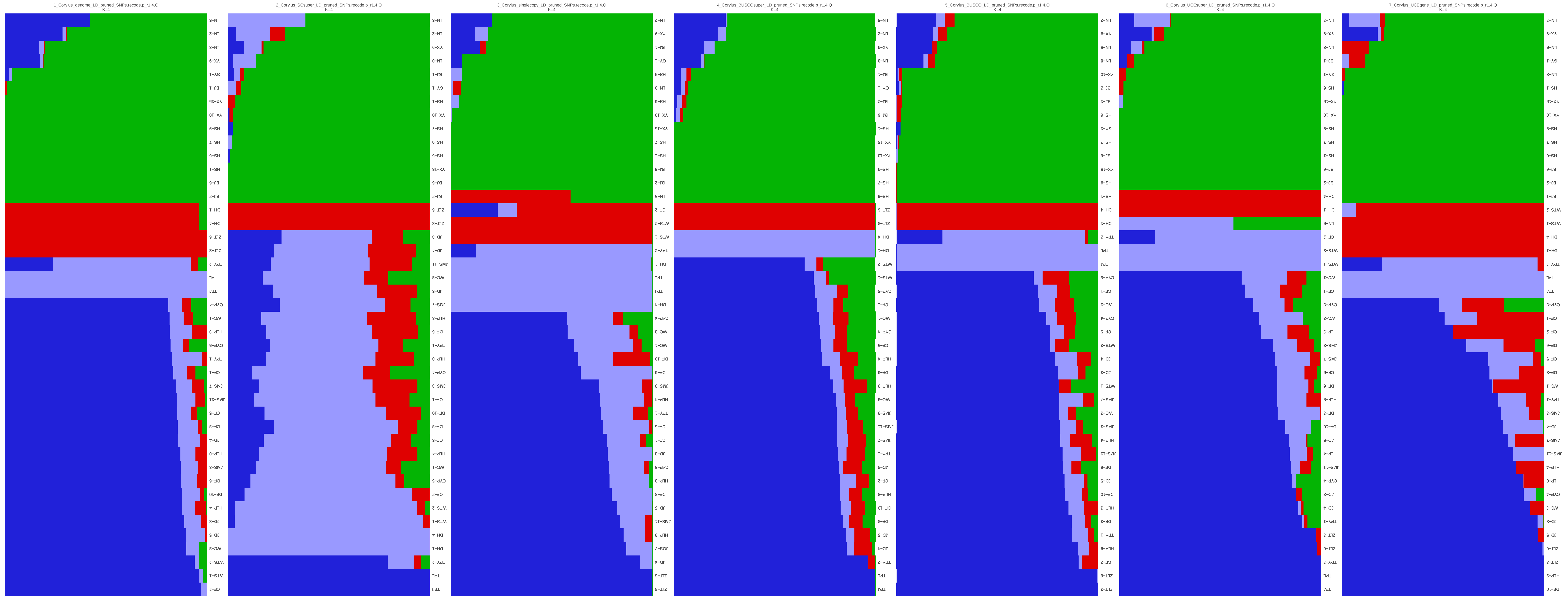
1_Corylus_genome_LD_pruned_SNPs.recode.p_r1.4.Q
K=4
2_Corylus_SCsuper_LD_pruned_SNPs.recode.p_r1.4.Q
K=4
3_Corylus_singlecopy_LD_pruned_SNPs.recode.p_r1.4.Q
K=4
4_Corylus_BUSCOsuper_LD_pruned_SNPs.recode.p_r1.4.Q
K=4
5_Corylus_BUSCO_LD_pruned_SNPs.recode.p_r1.4.Q
K=4
6_Corylus_UCEsuper_LD_pruned_SNPs.recode.p_r1.4.Q
K=4
7_Corylus_UCEgene_LD_pruned_SNPs.recode.p_r1.4.Q
K=4

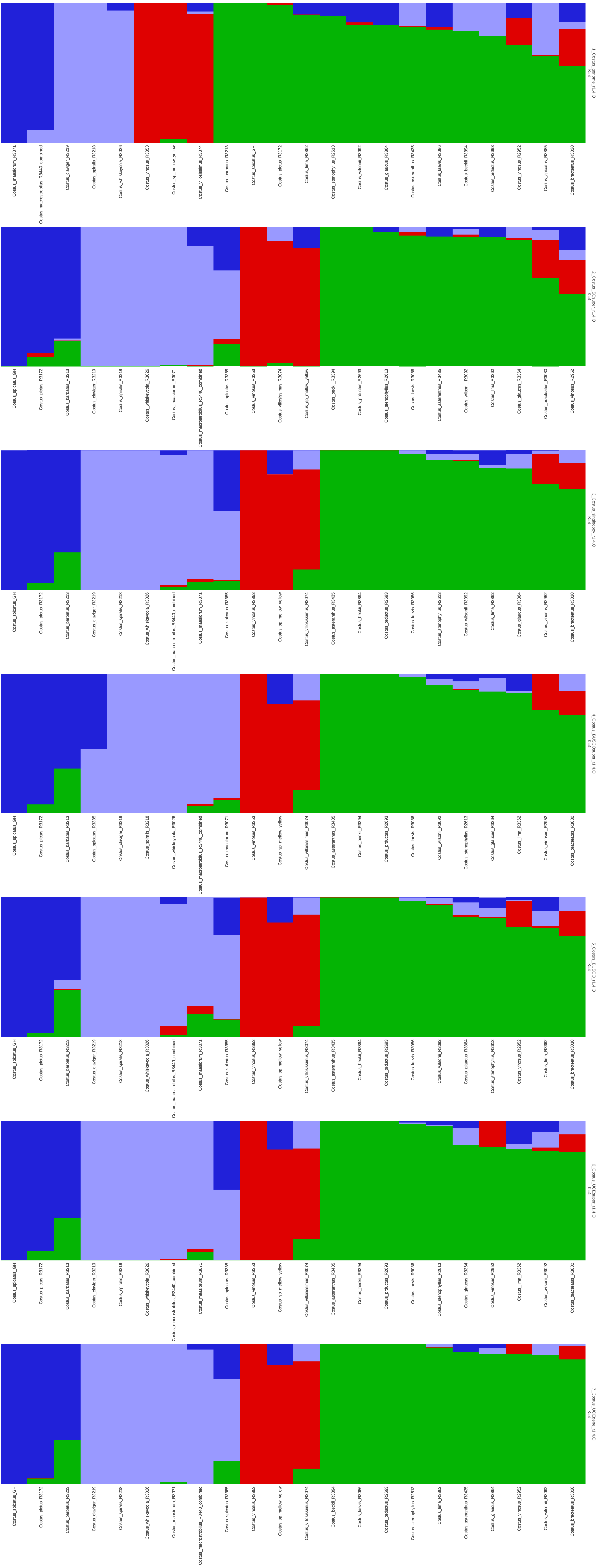

1_Costus_genome_r1.4.Q
K=4
Costus_maasiorum_R3071
Costus_macrostrobilus_R3440_combined
Costus_claviger_R3219
Costus_spiralis_R3218
Costus_whiskeycola_R3026
Costus_vinosus_R3353
Costus_sp_mellow_yellow
Costus_villosissimus_R3074
Costus_barbatus_R3213
Costus_spicatus_GH
Costus_pictus_R3172
Costus_lima_R3362
Costus_stenophyllus_R2613
Costus_wilsonii_R3092
Costus_glaucus_R3364
Costus_asteranthus_R3435
Costus_laevis_R3086
Costus_beckii_R3394
Costus_productus_R2693
Costus_vinosus_R2952
Costus_spicatus_R3385
Costus_bracteatus_R3030
2_Costus_SCsuper_r1.4.Q
K=4
Costus_spicatus_GH
Costus_pictus_R3172
Costus_barbatus_R3213
Costus_claviger_R3219
Costus_spiralis_R3218
Costus_whiskeycola_R3026
Costus_maasiorum_R3071
Costus_macrostrobilus_R3440_combined
Costus_spicatus_R3385
Costus_vinosus_R3353
Costus_villosissimus_R3074
Costus_sp_mellow_yellow
Costus_beckii_R3394
Costus_productus_R2693
Costus_stenophyllus_R2613
Costus_laevis_R3086
Costus_asteranthus_R3435
Costus_wilsonii_R3092
Costus_lima_R3362
Costus_glaucus_R3364
Costus_bracteatus_R3030
Costus_vinosus_R2952
3_Costus_singlecopy_r1.4.Q
K=4
Costus_spicatus_GH
Costus_pictus_R3172
Costus_barbatus_R3213
Costus_claviger_R3219
Costus_spiralis_R3218
Costus_whiskeycola_R3026
Costus_macrostrobilus_R3440_combined
Costus_maasiorum_R3071
Costus_spicatus_R3385
Costus_vinosus_R3353
Costus_sp_mellow_yellow
Costus_villosissimus_R3074
Costus_asteranthus_R3435
Costus_beckii_R3394
Costus_productus_R2693
Costus_laevis_R3086
Costus_stenophyllus_R2613
Costus_wilsonii_R3092
Costus_lima_R3362
Costus_glaucus_R3364
Costus_vinosus_R2952
Costus_bracteatus_R3030
4_Costus_BUSCOsuper_r1.4.Q
K=4
Costus_spicatus_GH
Costus_pictus_R3172
Costus_barbatus_R3213
Costus_spicatus_R3385
Costus_claviger_R3219
Costus_spiralis_R3218
Costus_whiskeycola_R3026
Costus_macrostrobilus_R3440_combined
Costus_maasiorum_R3071
Costus_vinosus_R3353
Costus_sp_mellow_yellow
Costus_villosissimus_R3074
Costus_asteranthus_R3435
Costus_beckii_R3394
Costus_productus_R2693
Costus_laevis_R3086
Costus_wilsonii_R3092
Costus_stenophyllus_R2613
Costus_glaucus_R3364
Costus_lima_R3362
Costus_vinosus_R2952
Costus_bracteatus_R3030
5_Costus_BUSCO_r1.4.Q
K=4
Costus_spicatus_GH
Costus_pictus_R3172
Costus_barbatus_R3213
Costus_claviger_R3219
Costus_spiralis_R3218
Costus_whiskeycola_R3026
Costus_macrostrobilus_R3440_combined
Costus_maasiorum_R3071
Costus_spicatus_R3385
Costus_vinosus_R3353
Costus_sp_mellow_yellow
Costus_villosissimus_R3074
Costus_asteranthus_R3435
Costus_beckii_R3394
Costus_productus_R2693
Costus_laevis_R3086
Costus_wilsonii_R3092
Costus_glaucus_R3364
Costus_stenophyllus_R2613
Costus_vinosus_R2952
Costus_lima_R3362
Costus_bracteatus_R3030
6_Costus_UCEsuper_r1.4.Q
K=4
Costus_spicatus_GH
Costus_pictus_R3172
Costus_barbatus_R3213
Costus_claviger_R3219
Costus_spiralis_R3218
Costus_whiskeycola_R3026
Costus_macrostrobilus_R3440_combined
Costus_maasiorum_R3071
Costus_spicatus_R3385
Costus_vinosus_R3353
Costus_sp_mellow_yellow
Costus_villosissimus_R3074
Costus_asteranthus_R3435
Costus_beckii_R3394
Costus_productus_R2693
Costus_laevis_R3086
Costus_stenophyllus_R2613
Costus_glaucus_R3364
Costus_vinosus_R2952
Costus_lima_R3362
Costus_wilsonii_R3092
Costus_bracteatus_R3030
7_Costus_UCEgene_r1.4.Q
K=4
Costus_spicatus_GH
Costus_pictus_R3172
Costus_barbatus_R3213
Costus_claviger_R3219
Costus_spiralis_R3218
Costus_whiskeycola_R3026
Costus_maasiorum_R3071
Costus_macrostrobilus_R3440_combined
Costus_spicatus_R3385
Costus_vinosus_R3353
Costus_sp_mellow_yellow
Costus_villosissimus_R3074
Costus_beckii_R3394
Costus_laevis_R3086
Costus_productus_R2693
Costus_stenophyllus_R2613
Costus_lima_R3362
Costus_asteranthus_R3435
Costus_glaucus_R3364
Costus_vinosus_R2952
Costus_wilsonii_R3092
Costus_bracteatus_R3030

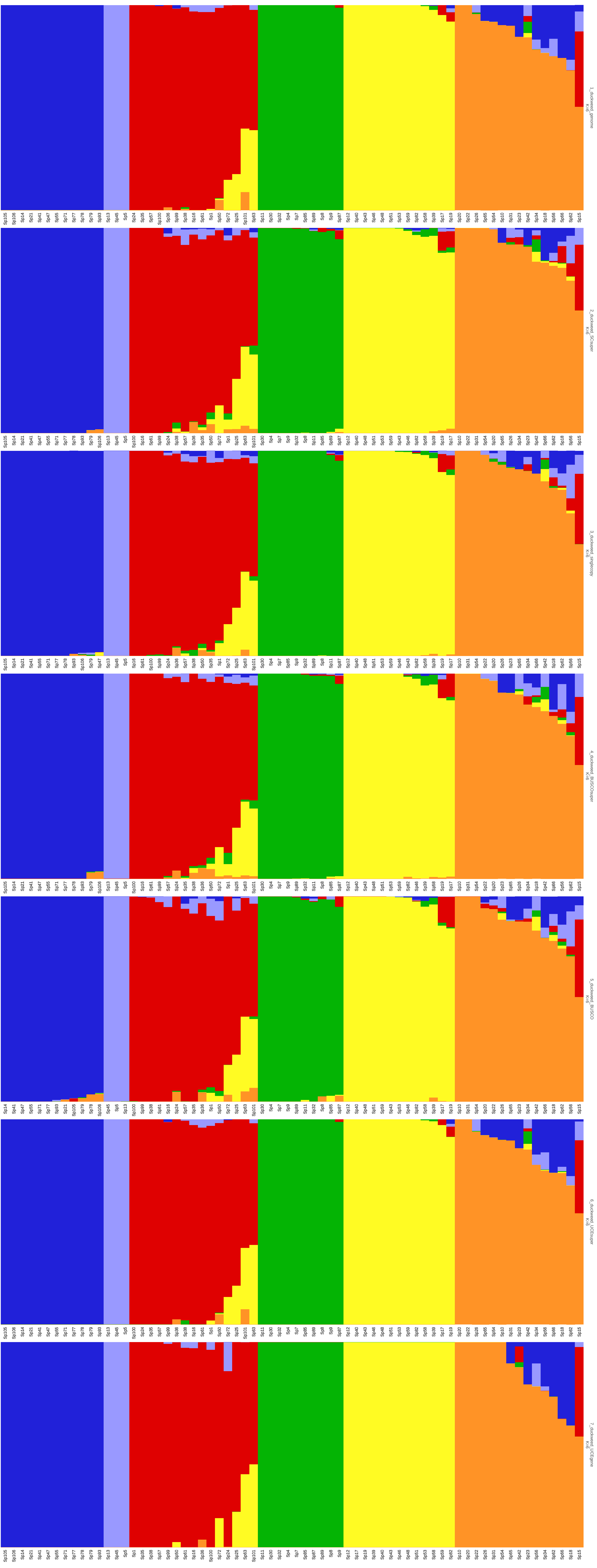
1_duckweed_genome
K=6
2_duckweed_SCsuper
K=6
3_duckweed_singlecopy
K=6
4_duckweed_BUSCOsuper
K=6
5_duckweed_BUSCO
K=6
6_duckweed_UCEsuper
K=6
7_duckweed_UCEgene
K=6

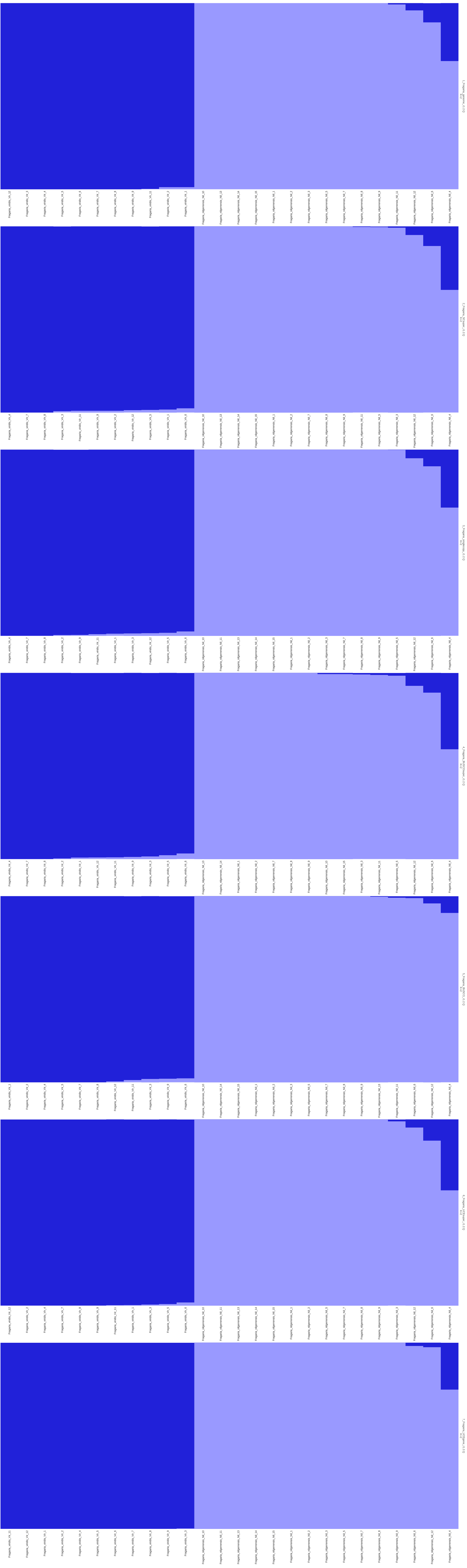

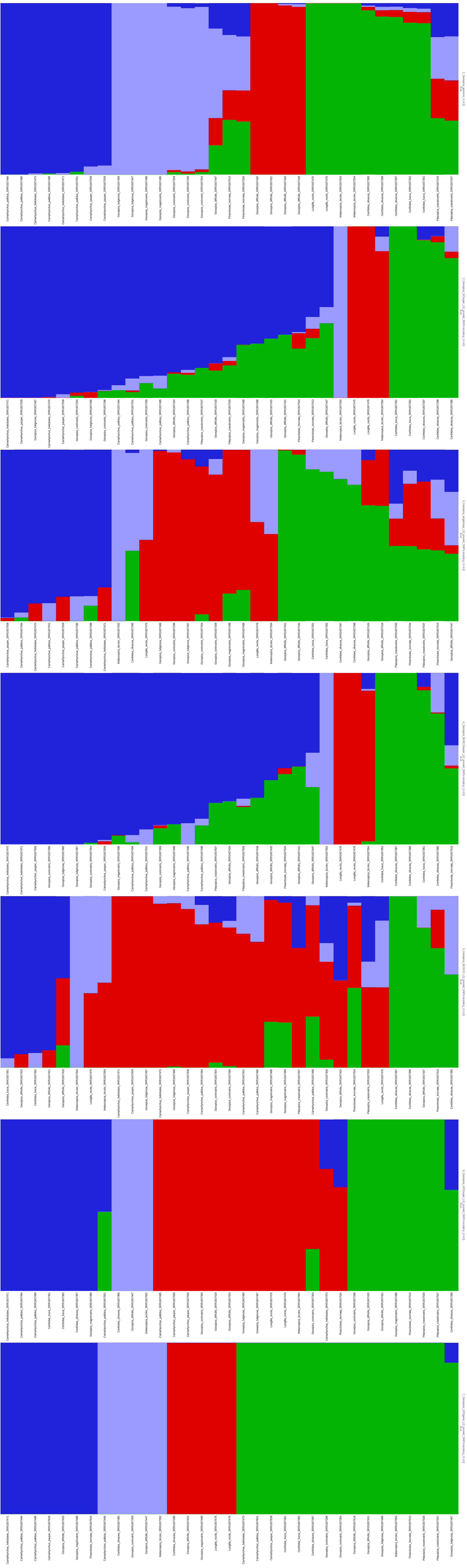

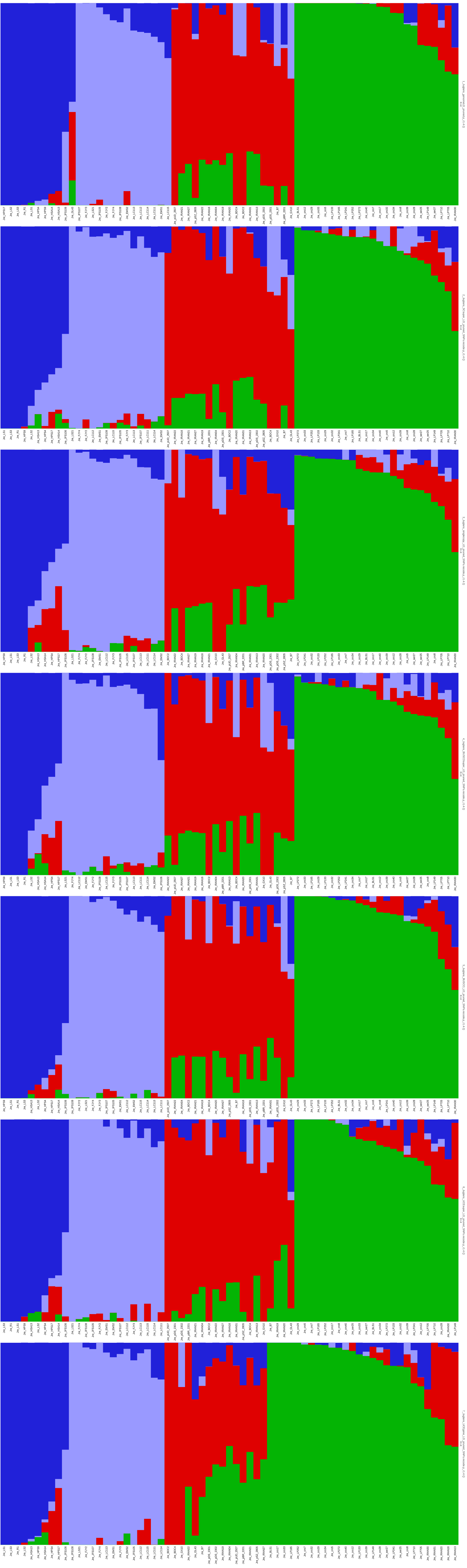

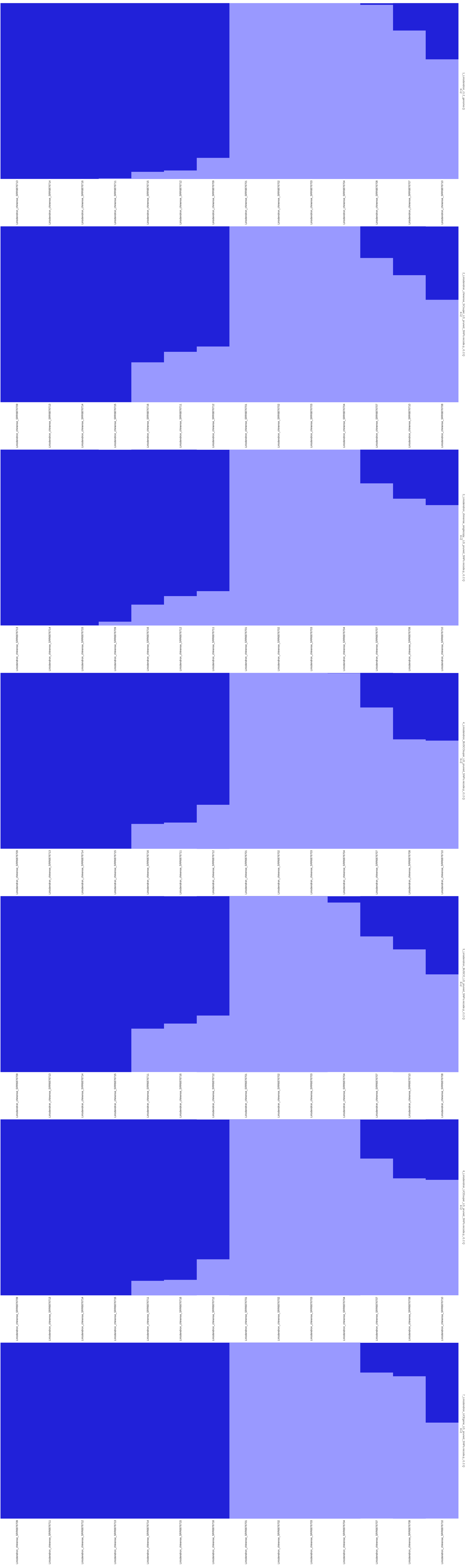

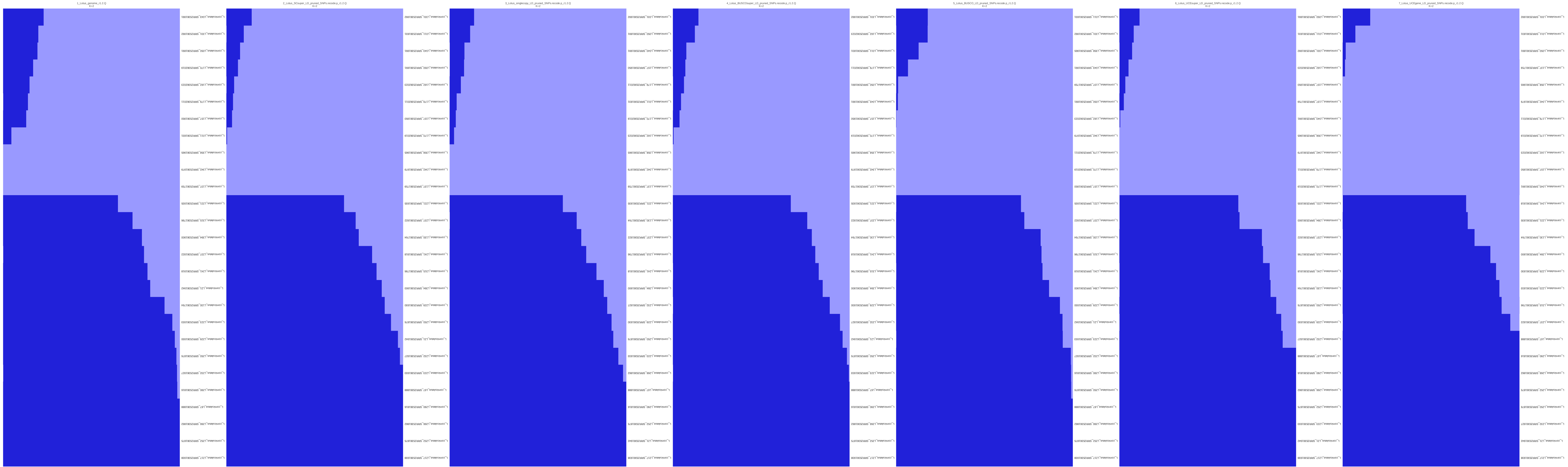
1_Lotus_genome_r1.2.Q
K=2
2_Lotus_SCsuper_LD_pruned_SNPs.recode.p_r1.2.Q
K=2
3_Lotus_singlecopy_LD_pruned_SNPs.recode.p_r1.2.Q
K=2
4_Lotus_BUSCOsuper_LD_pruned_SNPs.recode.p_r1.2.Q
K=2
5_Lotus_BUSCO_LD_pruned_SNPs.recode.p_r1.2.Q
K=2
6_Lotus_UCEsuper_LD_pruned_SNPs.recode.p_r1.2.Q
K=2
7_Lotus_UCEgene_LD_pruned_SNPs.recode.p_r1.2.Q
K=2

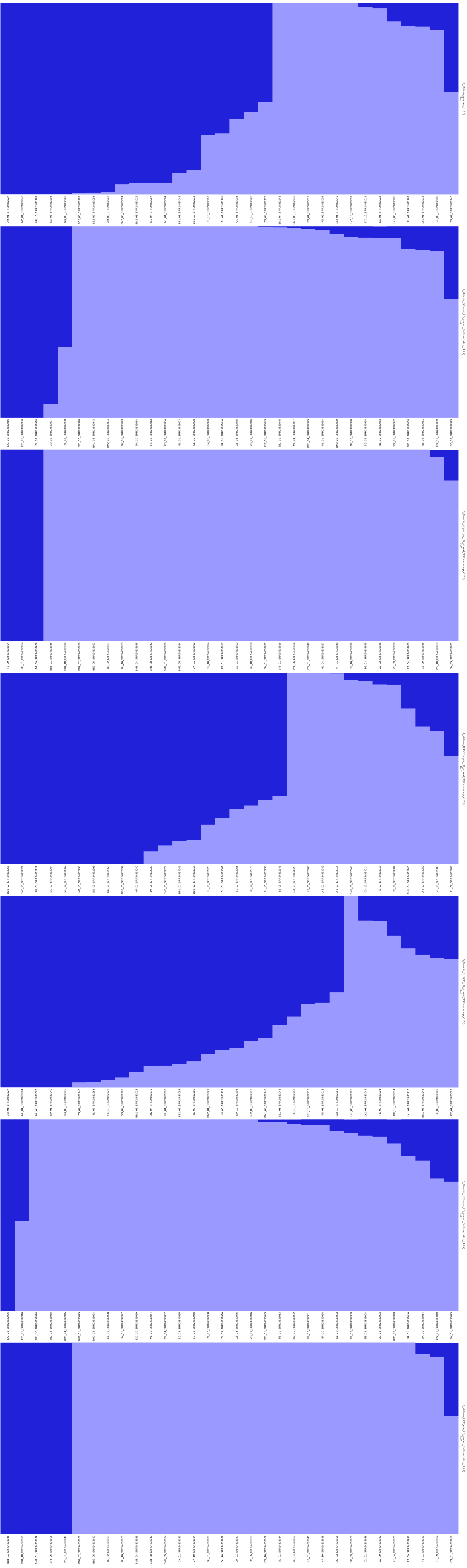

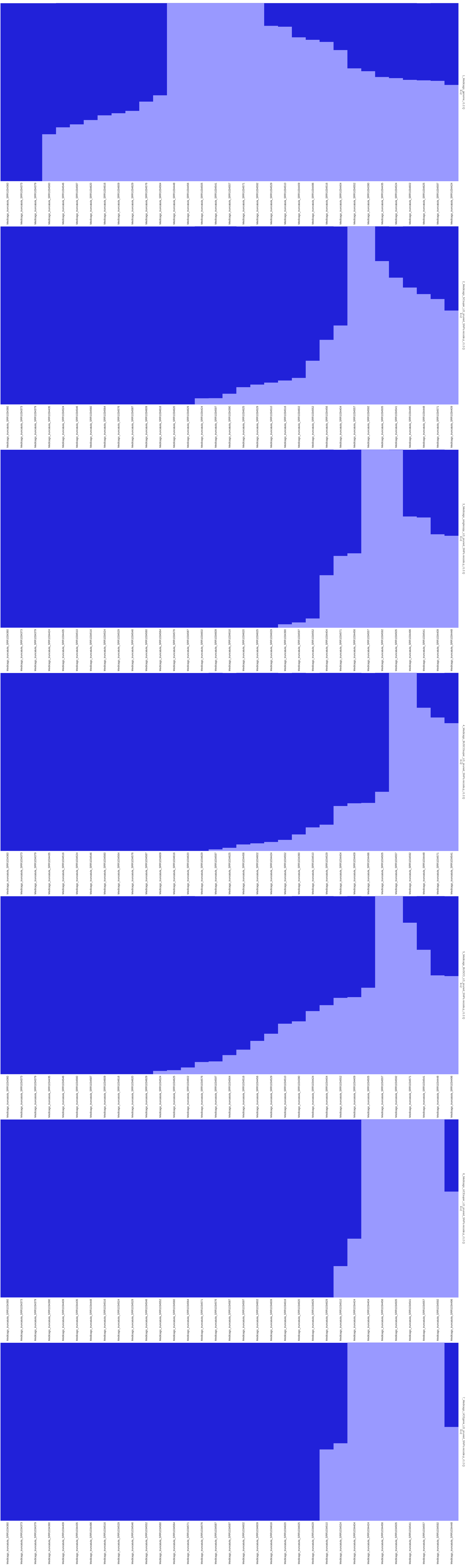

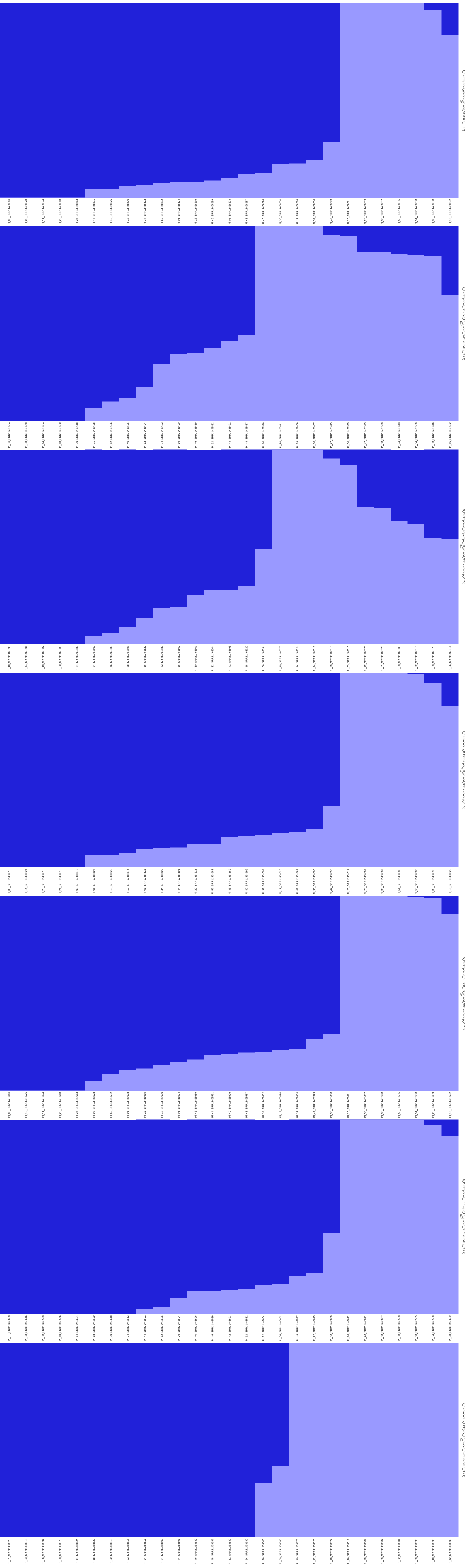

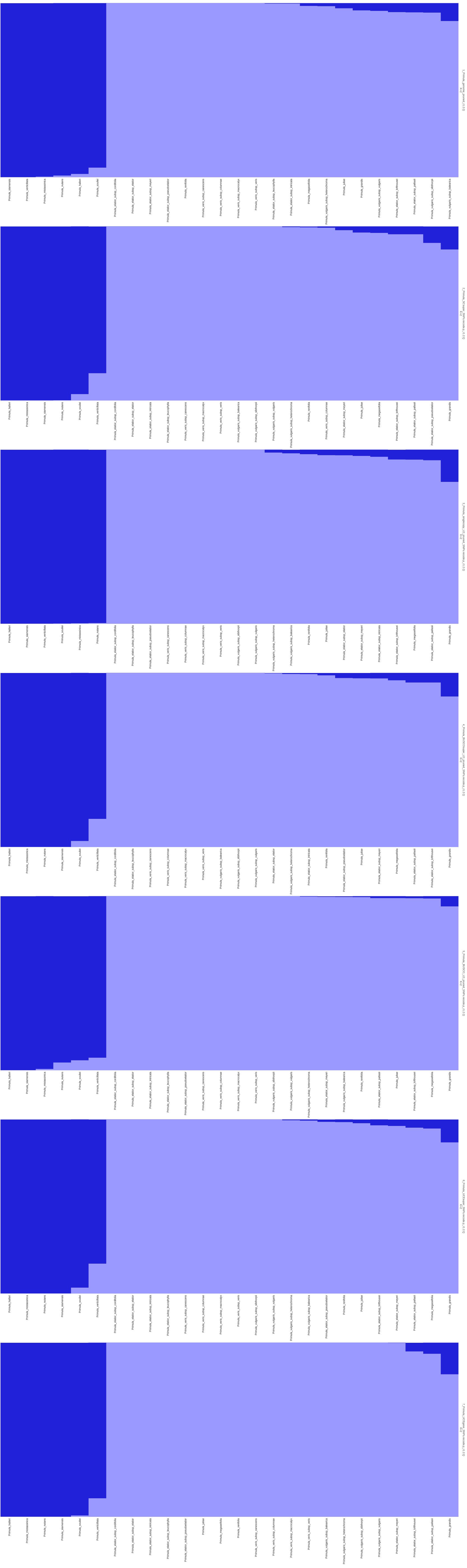
1_Primula_genome_pruned_r1.2.Q
K=2
Primula_siamensis
Primula_verticillata
Primula_mistassinica
Primula_nutans
Primula_halleri
Primula_souliei
Primula_elatior_subsp_cordifolia
Primula_elatior_subsp_elatior
Primula_elatior_subsp_meyeri
Primula_elatior_subsp_pseudoelatior
Primula_renifolia
Primula_veris_subsp_canescens
Primula_veris_subsp_columnae
Primula_veris_subsp_macrocalyx
Primula_veris_subsp_veris
Primula_elatior_subsp_leucophylla
Primula_elatior_subsp_intricata
Primula_megaseifolia
Primula_vulgaris_subsp_heterochroma
Primula_juliae
Primula_grandis
Primula_vulgaris_subsp_vulgaris
Primula_elatior_subsp_lofthousei
Primula_elatior_subsp_pallasii
Primula_vulgaris_subsp_sibthorpii
Primula_vulgaris_subsp_balearica
2_Primula_SCsuper_SNPs.recode.p_r1.2.Q
K=2
Primula_halleri
Primula_mistassinica
Primula_siamensis
Primula_nutans
Primula_souliei
Primula_verticillata
Primula_elatior_subsp_cordifolia
Primula_elatior_subsp_elatior
Primula_elatior_subsp_intricata
Primula_elatior_subsp_leucophylla
Primula_veris_subsp_canescens
Primula_veris_subsp_macrocalyx
Primula_veris_subsp_veris
Primula_vulgaris_subsp_balearica
Primula_vulgaris_subsp_sibthorpii
Primula_vulgaris_subsp_vulgaris
Primula_vulgaris_subsp_heterochroma
Primula_renifolia
Primula_veris_subsp_columnae
Primula_elatior_subsp_meyeri
Primula_juliae
Primula_megaseifolia
Primula_elatior_subsp_lofthousei
Primula_elatior_subsp_pallasii
Primula_elatior_subsp_pseudoelatior
Primula_grandis
3_Primula_singlecopy_LD_pruned_SNPs.recode.p_r1.2.Q
K=2
Primula_halleri
Primula_siamensis
Primula_verticillata
Primula_souliei
Primula_mistassinica
Primula_nutans
Primula_elatior_subsp_cordifolia
Primula_elatior_subsp_leucophylla
Primula_elatior_subsp_pseudoelatior
Primula_veris_subsp_canescens
Primula_veris_subsp_columnae
Primula_veris_subsp_macrocalyx
Primula_veris_subsp_veris
Primula_vulgaris_subsp_sibthorpii
Primula_vulgaris_subsp_vulgaris
Primula_vulgaris_subsp_heterochroma
Primula_vulgaris_subsp_balearica
Primula_renifolia
Primula_juliae
Primula_elatior_subsp_elatior
Primula_elatior_subsp_meyeri
Primula_elatior_subsp_intricata
Primula_elatior_subsp_lofthousei
Primula_megaseifolia
Primula_elatior_subsp_pallasii
Primula_grandis
4_Primula_BUSCOsuper_LD_pruned_SNPs.recode.p_r1.2.Q
K=2
Primula_halleri
Primula_mistassinica
Primula_nutans
Primula_siamensis
Primula_souliei
Primula_verticillata
Primula_elatior_subsp_cordifolia
Primula_elatior_subsp_leucophylla
Primula_veris_subsp_canescens
Primula_veris_subsp_columnae
Primula_veris_subsp_macrocalyx
Primula_veris_subsp_veris
Primula_vulgaris_subsp_balearica
Primula_vulgaris_subsp_sibthorpii
Primula_vulgaris_subsp_vulgaris
Primula_elatior_subsp_elatior
Primula_vulgaris_subsp_heterochroma
Primula_elatior_subsp_intricata
Primula_renifolia
Primula_elatior_subsp_pseudoelatior
Primula_juliae
Primula_elatior_subsp_meyeri
Primula_megaseifolia
Primula_elatior_subsp_pallasii
Primula_elatior_subsp_lofthousei
Primula_grandis
5_Primula_BUSCO_LD_pruned_SNPs.recode.p_r1.2.Q
K=2
Primula_halleri
Primula_siamensis
Primula_mistassinica
Primula_nutans
Primula_souliei
Primula_verticillata
Primula_elatior_subsp_cordifolia
Primula_elatior_subsp_elatior
Primula_elatior_subsp_intricata
Primula_elatior_subsp_leucophylla
Primula_elatior_subsp_pseudoelatior
Primula_veris_subsp_canescens
Primula_veris_subsp_columnae
Primula_veris_subsp_macrocalyx
Primula_veris_subsp_veris
Primula_vulgaris_subsp_sibthorpii
Primula_vulgaris_subsp_vulgaris
Primula_vulgaris_subsp_heterochroma
Primula_elatior_subsp_meyeri
Primula_vulgaris_subsp_balearica
Primula_renifolia
Primula_elatior_subsp_pallasii
Primula_juliae
Primula_elatior_subsp_lofthousei
Primula_megaseifolia
Primula_grandis
K=2
Primula_halleri
Primula_mistassinica
Primula_nutans
Primula_siamensis
Primula_souliei
Primula_verticillata
Primula_elatior_subsp_cordifolia
Primula_elatior_subsp_elatior
Primula_elatior_subsp_intricata
Primula_elatior_subsp_leucophylla
Primula_veris_subsp_canescens
Primula_veris_subsp_columnae
Primula_veris_subsp_macrocalyx
Primula_veris_subsp_veris
Primula_vulgaris_subsp_sibthorpii
Primula_vulgaris_subsp_vulgaris
Primula_vulgaris_subsp_heterochroma
Primula_vulgaris_subsp_balearica
Primula_elatior_subsp_pseudoelatior
Primula_renifolia
Primula_juliae
Primula_elatior_subsp_lofthousei
Primula_elatior_subsp_meyeri
Primula_elatior_subsp_pallasii
Primula_megaseifolia
Primula_grandis
K=2
Primula_halleri
Primula_mistassinica
Primula_nutans
Primula_siamensis
Primula_souliei
Primula_verticillata
Primula_elatior_subsp_cordifolia
Primula_elatior_subsp_elatior
Primula_elatior_subsp_intricata
Primula_elatior_subsp_leucophylla
Primula_elatior_subsp_pseudoelatior
Primula_juliae
Primula_megaseifolia
Primula_renifolia
Primula_veris_subsp_canescens
Primula_veris_subsp_columnae
Primula_veris_subsp_macrocalyx
Primula_veris_subsp_veris
Primula_vulgaris_subsp_balearica
Primula_vulgaris_subsp_heterochroma
Primula_vulgaris_subsp_sibthorpii
Primula_vulgaris_subsp_vulgaris
Primula_elatior_subsp_meyeri
Primula_elatior_subsp_lofthousei
Primula_elatior_subsp_pallasii
Primula_grandis

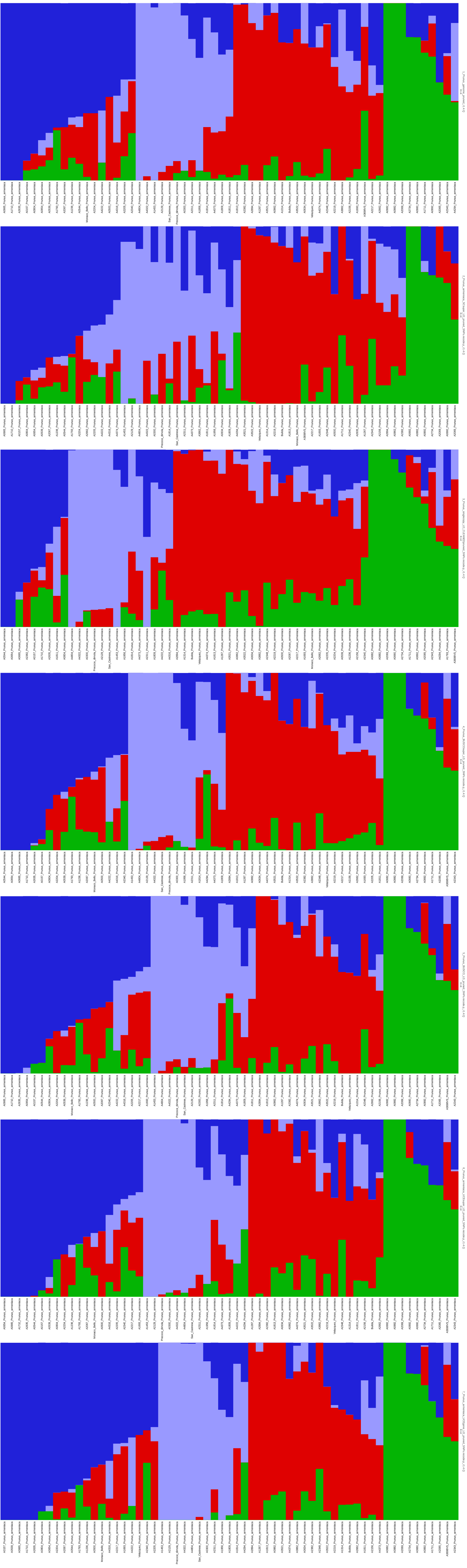

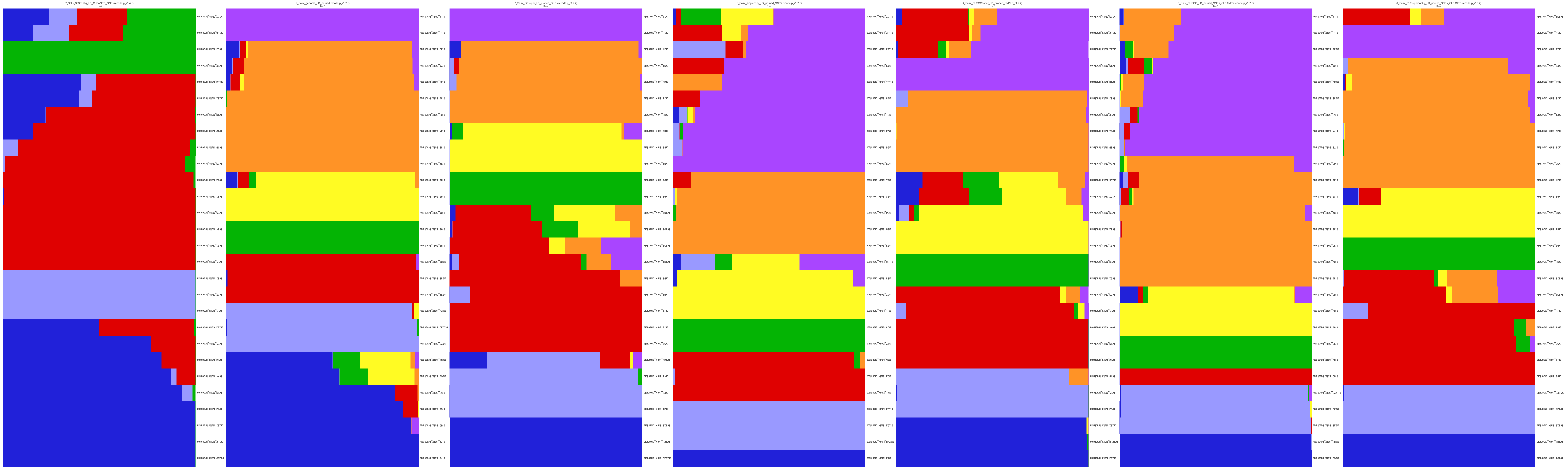
7_Salix_353contig_LD_CLEANED_SNPs.recode.p_r1.4.Q
K=4
1_Salix_genome_LD_pruned.recode.p_r1.7.Q
K=7
2_Salix_SCsuper_LD_pruned_SNPs.recode.p_r1.7.Q
K=7
3_Salix_singlecopy_LD_pruned_SNPs.recode.p_r1.7.Q
K=7
4_Salix_BUSCOsuper_LD_pruned_SNPs.p_r1.7.Q
K=7
5_Salix_BUSCO_LD_pruned_SNPs_CLEANED.recode.p_r1.7.Q
K=7
6_Salix_353Supercontig_LD_pruned_SNPs_CLEANED.recode.p_r1.7.Q
K=7

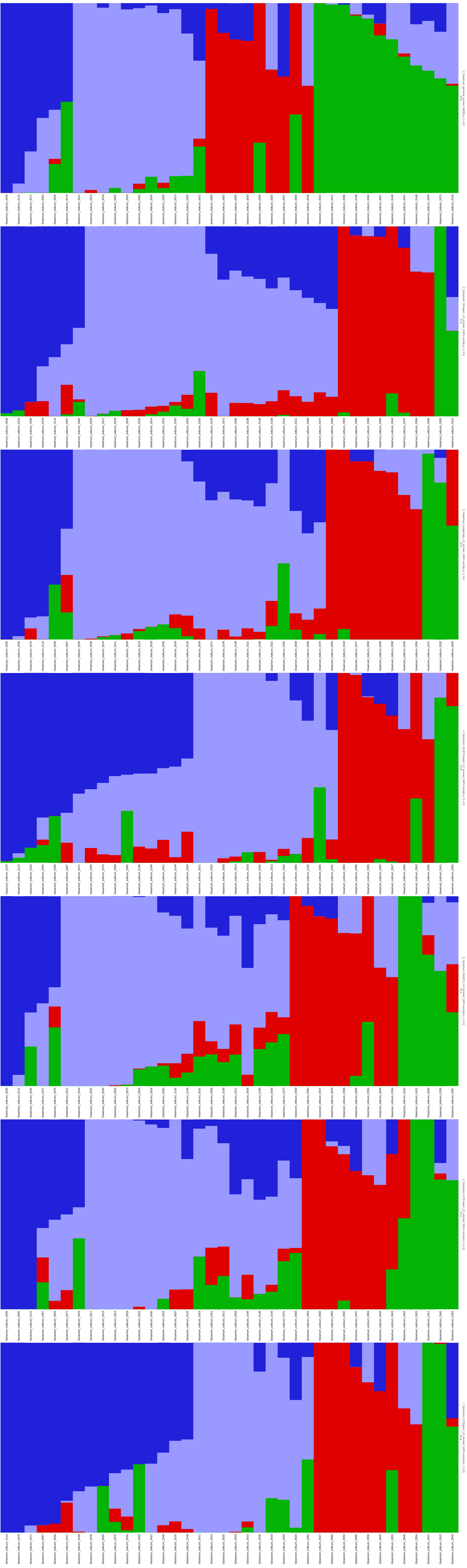

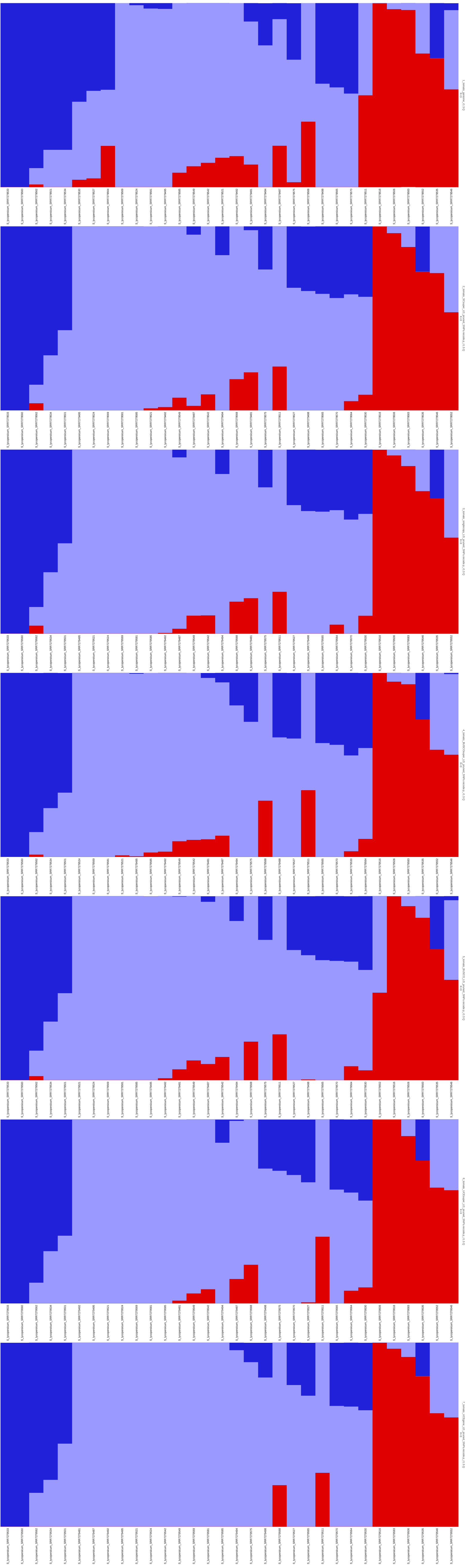

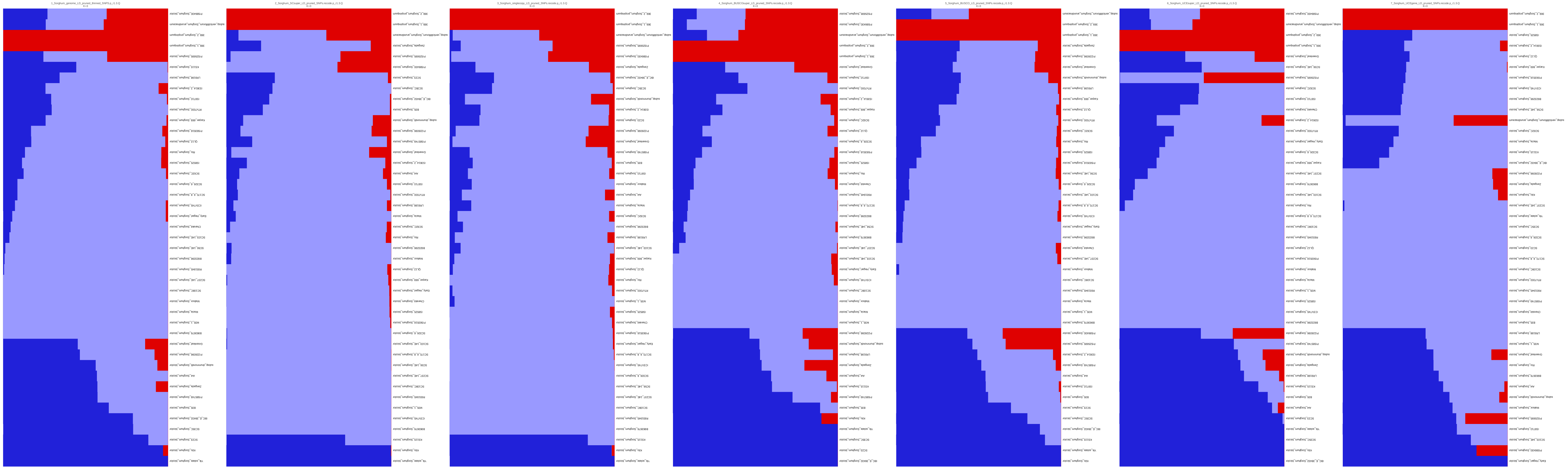

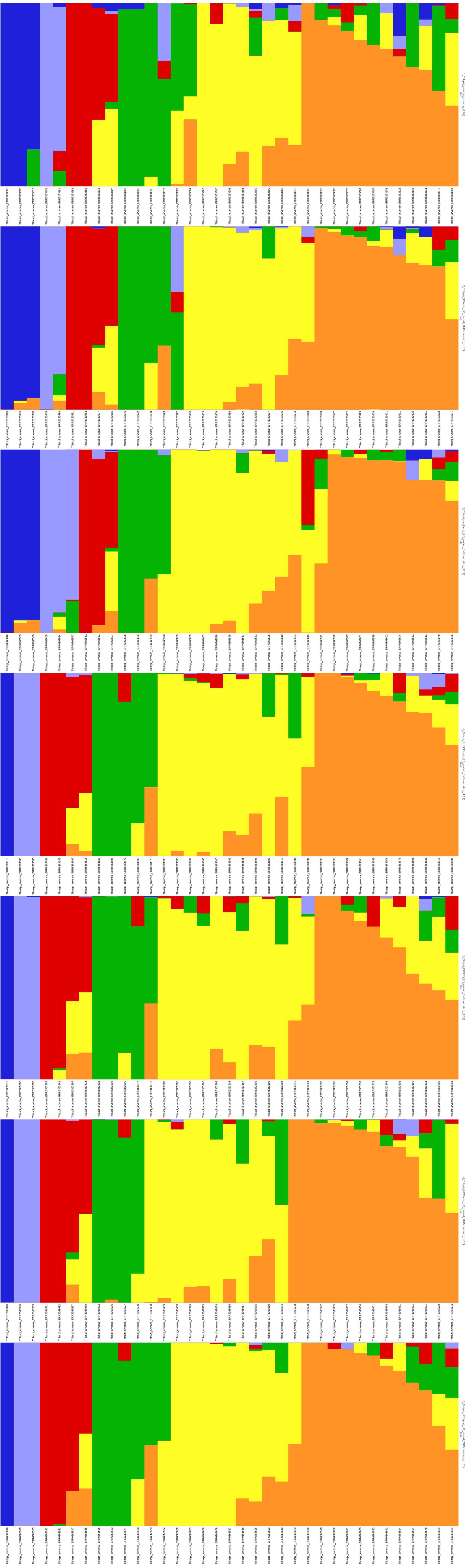

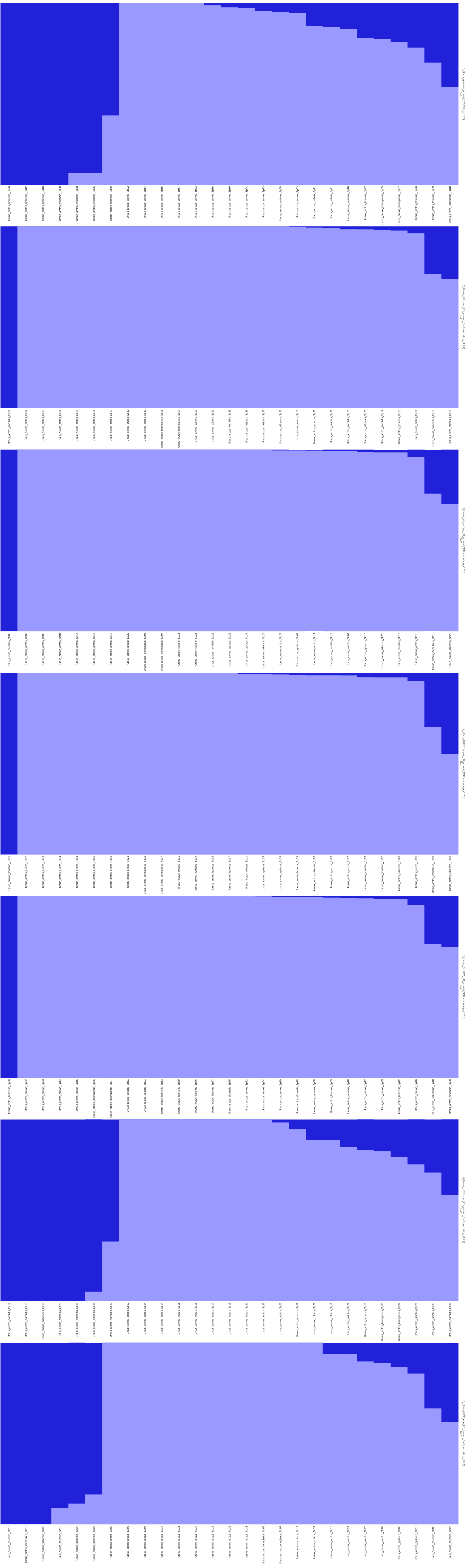

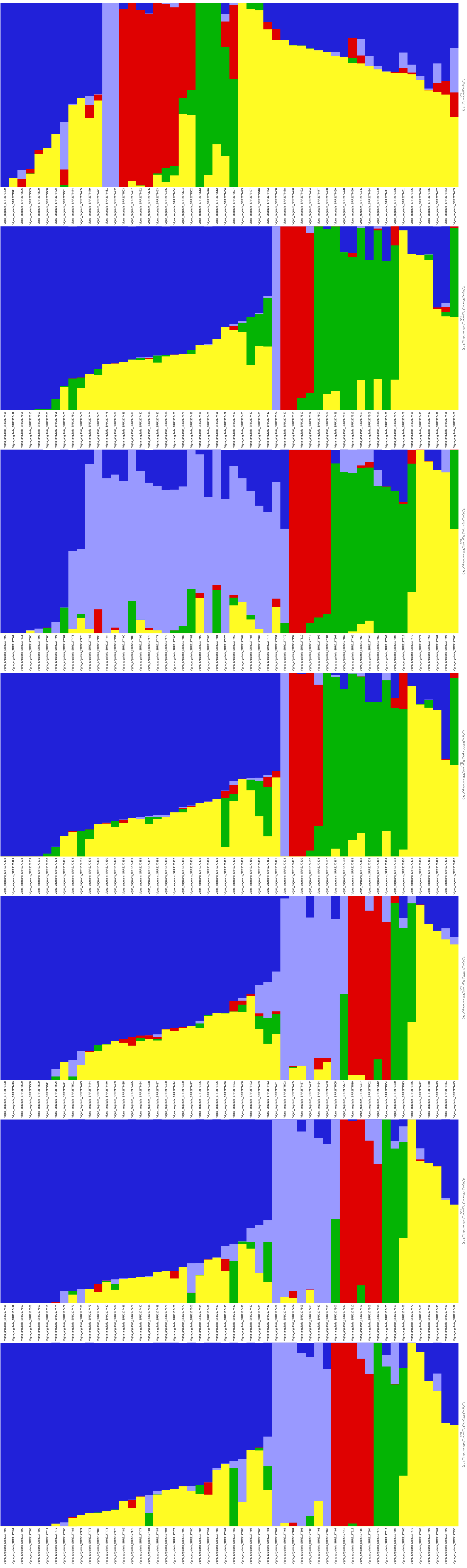

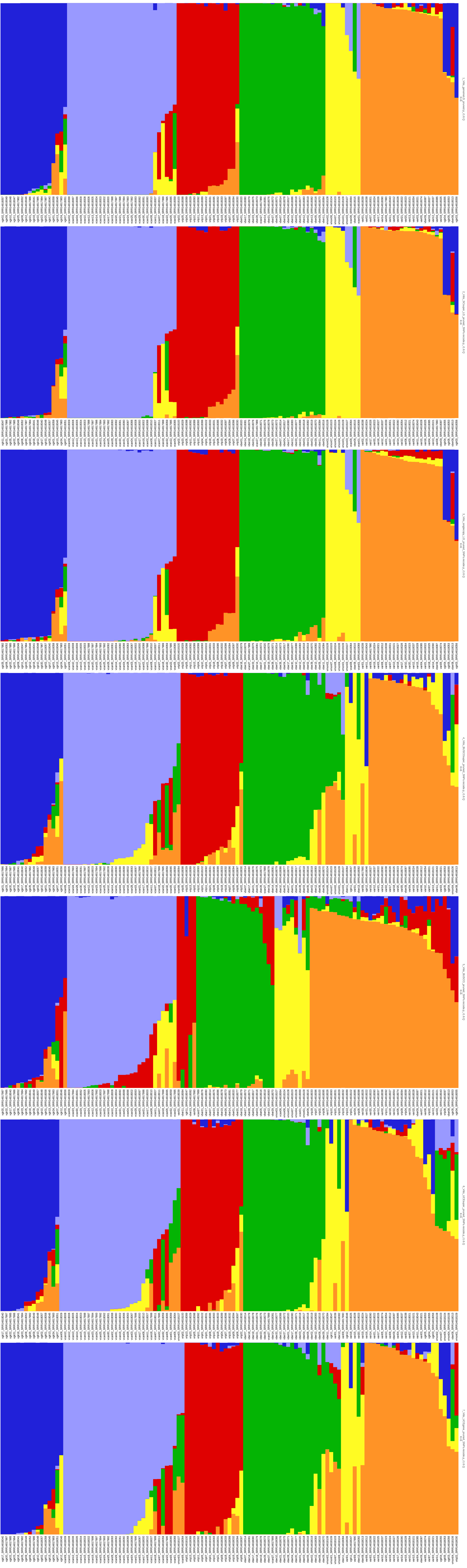

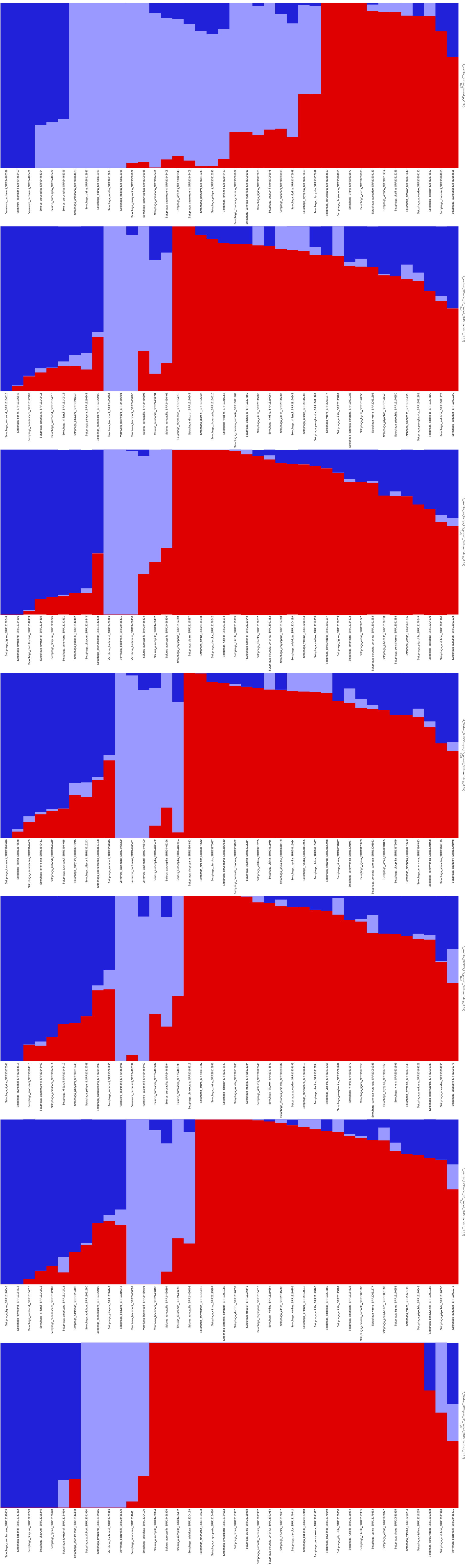
